# Proteomic Analysis Links Truncated Tau to Lysosome Motility, Autophagy and Endo-lysosomal Dysfunction

**DOI:** 10.1101/2025.03.18.643906

**Authors:** Despoina Goniotaki, Maximilian Wulf, Steven Lynham, Ayushin Ale, George Chennell, Stefania Markotti, Katrin Marcus, Wendy Noble, Diane P. Hanger, Graham Fraser, Deepak P. Srivastava

## Abstract

Tauopathies are characterized by the progressive accumulation of abnormal tau species, which disrupt the autophagy-lysosomal pathway (ALP), a critical system for degrading intracellular macromolecules and aggregated proteins, causing toxicity and cell death. This study investigates the impact of the N-terminally truncated Tau35 protein overexpression on proteolytic pathways, including effects on autophagy and endo-lysosomal processes. Using a Tau35 mouse model and SH-SY5Y cell lines stably expressing either the Tau35 fragment or full-length tau, we employed western blotting, proteomic analysis of lysosome-enriched brain fractions, proteolysis/endocytosis assays, and live-cell imaging with the lysotracker reporter to assess protein degradation and lysosomal function. Our findings identify early pathological changes in endo-lysosomal processes, including increased endocytosis, proteolytic dysfunction and lysosomal motility abnormalities, associated with Tau35-induced toxicity. This work extends previous research by providing new insights into the mechanisms of Tau35-induced neurotoxicity, offering a foundation for developing targeted therapeutic strategies to address tauopathies.

## Main

Tauopathies are progressive neurodegenerative disorders characterized by the intracellular deposition of abnormal tau protein aggregates and age-related neuronal loss ^1^. The accumulation of tau within cells reflects disruptions in protein homeostasis and impaired clearance mechanisms mediated by the endo-lysosomal and autophagic pathways^2^. Growing genetic and molecular evidence suggests that dysfunction in these systems extends beyond the degradation of aggregation-prone proteins, implicating their broader roles in the pathophysiology of tauopathies and other neurodegenerative diseases^2,3^. While aberrant post-translational modifications of tau, especially hyperphosphorylation, have traditionally been considered key drivers of tau aggregation and neurotoxicity^4^, recent findings highlight the critical role of tau truncation in neurodegeneration^5^. The process of tau proteolysis has attracted growing interest for its potential role in driving disease progression through mechanisms specific to individual fragments, such as aggregation and cell-to-cell propagation^5,6^. Among these, the Tau35 fragment, initially identified in human brain tissue and associated with primary human tauopathies^7,8^, has been shown to disrupt kinase activity, as well as lysosomal and synaptic functions in the Tau35 mouse model. Notably, these mice express the Tau35 fragment at relatively low levels compared to most other overexpressing animal models^9^.

The effect of Tau35 overexpression in CHO cells, primary cortical neurons, and its sub-endogenous expression in Tau35 mice was recently reported, demonstrating a progressive accumulation of abnormally phosphorylated tau species across all models^9–11^, accompanied by structural and functional synaptic alterations ^10,12^, behavioural abnormalities ^9^, and disruptions involving critical elements of the three primary cellular protein degradation pathways: the proteasome, lysosomes, and autophagy ^9,13^. In 14-month-old Tau35 mice, representing late stages of the disease, notable elevations in p62 and LC3I/II markers have been observed, accompanied by a decrease in active cathepsin D, a critical lysosomal enzyme responsible for protein degradation (summarized in Table 1)^9^.

**Table 1.**
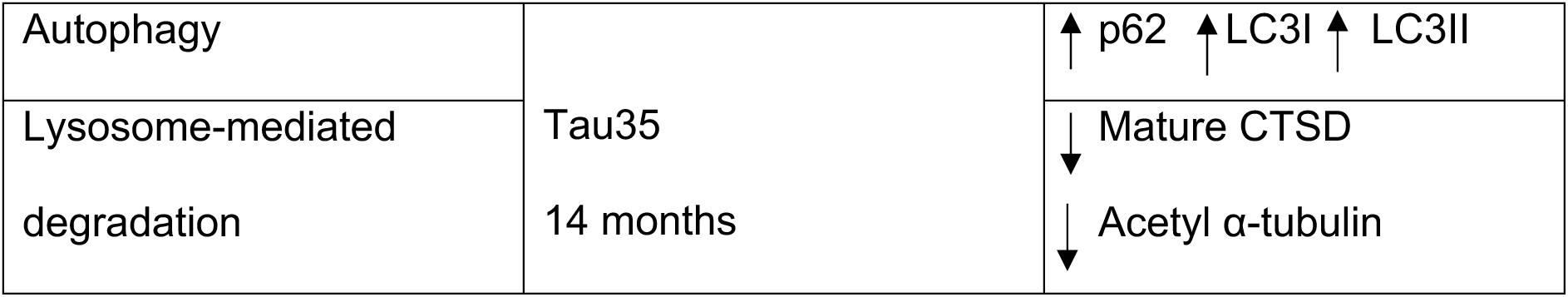
Summary of autophagy-lysosomal changes reported in the Tau35 mouse model at late disease stages (14M)^9^.

This study investigates the impact of Tau35 overexpression on physiological proteolytic pathways, with a focus on alterations in autophagy and endo-lysosomal processes. To achieve this, the Tau35 mouse model was utilized alongside the generation of human neuroblastoma (SH-SY5Y) cell lines stably expressing either the N-terminally truncated 35kDa human tau (187-441) with a C-terminal HA tag (Tau35) fragment or full-length (2N4R) human tau. Western blotting and proteomic analysis of lysosome-enriched brain fractions from Tau35 mice in early (4 month) and advanced (10 month) disease stages, as well as proteolysis and endocytosis assays and lysotracker-based live-cell imaging in differentiated SH-SY5Y cells, were employed to assess the effects of Tau35 fragment overexpression on protein degradation pathways. This work extends prior research by uncovering early pathological events and highlighting the role of altered endocytosis, proteolytic dysfunction and lysosomal motility disruptions in the presence of disease-associated human tau fragments. Insights into the downstream neurotoxic mechanisms of tau alterations may inform the development of targeted therapeutic strategies.

## Results

### Reduced LAMP2 expression and altered balance of active Cathepsin B (decreased) and Cathepsin L (increased) in the brains of Tau35 mice

Tau35 mice, illustrated in Figure 1a with a schematic comparing the expressed protein to full-length human tau, serve as a model for primary tauopathies^9^. To investigate the impact of Tau35 expression on alterations in the autophagy-lysosomal pathway (ALP) during disease progression, we euthanized mice at early (4 months; WT, n=8; Tau35, n=8) and advanced (10 months; WT, n=8–12; Tau35, n=8–12) pathological stages. The brains of these mice, which exhibit progressive tauopathy characterized by elevated tau phosphorylation, accumulation of abnormal tau species, cognitive and motor impairments, and reduced lifespan^9^, were analysed for the expression of selected lysosomal markers.

**Fig. 1:**
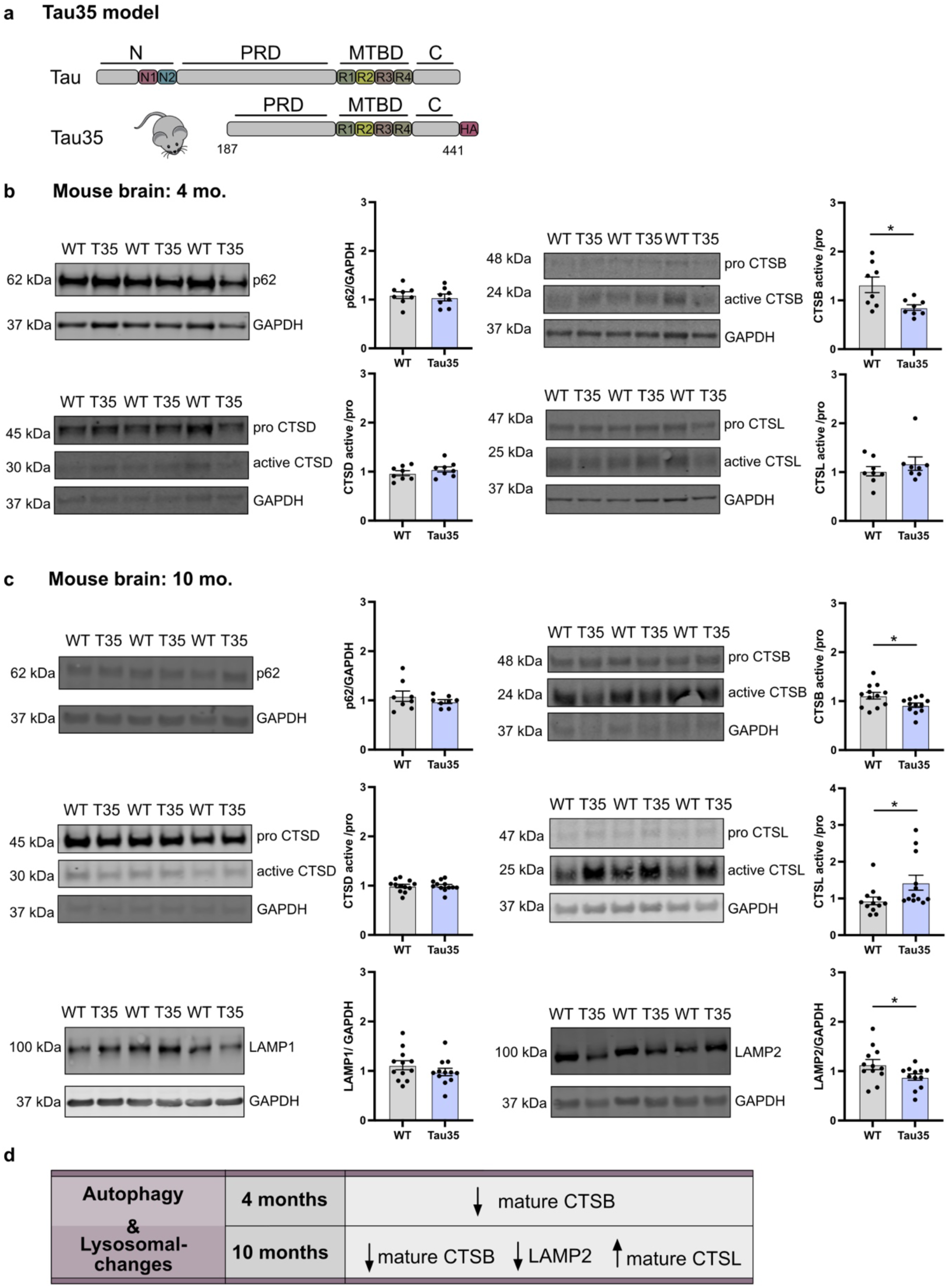
Reduced LAMP2 expression and imbalanced active cathepsins B↓/L↑ in Tau35 mouse brains a Tau35 model. **a,** Schematic representation of the Tau35-HA construct that is expressed in Tau35 mice, in comparison to full length human tau (441 amino acids). N: represents the amino-terminal domain, N1, N2: represent the two amino-terminal inserts, R1-R4: represent the four microtubule-binding domain repeats, PRD: represents the proline-rich domain, C: represents the carboxy-terminal domain. **b,c,** Western blots of total brain homogenates from WT and Tau35 mice aged 4 (**b**) and 10 (**c**) months respectively, were probed with antibodies to p62, CTSB, CTSD, CTSL, LAMP1, LAMP2 and GAPDH. Quantification of the blots is shown in the graphs as mean ± SEM, *n* = 8-12 brains per group. Student *t* test, \**P* < 0.05. p62/SQSTM1, Sequestosome-1; CTSB, Cathepsin B; CTSD, Cathepsin D; CTSL, Cathepsin L; LAMP1, lysosomal-associated membrane protein 1; LAMP2, lysosomal-associated membrane protein 2; GAPDH, glyceraldehyde 3-phosphate dehydrogenase; SEM, standard error of the mean; WT, wild type. **d,** Table summarizing changes in key lysosomal and autophagy markers (p62, LAMP1, LAMP2, CTSB, CTSD, CTSL) at early (4M) and advanced (10M) disease stages in Tau35 mice.

Western blot analysis of whole-brain tissue revealed a significant reduction in the expression of mature Cathepsin B (CTSB) at early disease stages (4 months) (Fig. 1b), while no significant changes were observed in other tested cathepsins, including their mature and immature forms, or in other key autophagy and lysosomal markers (Fig. 1b, Extended Data Fig. 1a). At advanced disease stages (10 months), the reduction in mature CTSB expression persisted, accompanied by a decrease in the lysosomal marker LAMP2 and a significant increase in the expression of mature Cathepsin L (CTSL). No significant changes were observed in the expression of Cathepsin D (CTSD), LAMP1, p62, or the premature forms of any tested cathepsins (Fig. 1c, Extended Data Fig. 1b). These findings indicate that in transgenic mice, early-stage pathology is associated with a selective reduction in mature CTSB expression, while other markers remain unchanged. At advanced stages, reductions in both LAMP2 and mature CTSB are observed, alongside an increase in mature CTSL (Fig. 1d), potentially reflecting a compensatory mechanism to mitigate the effects of CTSB deficiency and maintain tissue integrity^14^.

### Early Tau35 pathology alters endo-lysosomal pathways involved in mitochondria and energy/metabolism dynamics

The endo-lysosomal system is essential for executing metabolic tasks, including the uptake, intracellular trafficking, processing, appropriation, degradation, and disposal of molecules. In the context of tauopathies and other neurodegenerative diseases, it orchestrates the internalization, trafficking, and clearance of aggregated proteins^2,15^. Given its critical role in maintaining cellular homeostasis, the endo-lysosomal system is proposed to be a key regulator of neurodegeneration progression.

To assess the effect of Tau35 overexpression on the lysosomal proteome in the brain during disease progression, lysosome-enriched fractions were isolated from the brains of WT and Tau35 mice at early (4 months) and advanced (10 months) stages of tau pathology. This was achieved through differential and subsequent discontinuous iodixanol gradient centrifugation. The process is outlined in a flowchart, accompanied by representative images of the iodixanol gradient, showing enriched subcellular fractions (lysosomal and mitochondrial) from both WT and Tau35 mice (Fig. 2a and Extended Data Fig. 2a). Western blotting analysis of brain lysates and different fractions obtained showed highest enrichment of lysosomes in the LF fraction characterised by the lysosomal membrane protein LAMP2 (Fig. 2b) and highest enrichment of mitochondria in the MF fraction characterised by the mitochondrial transmembrane protein MCU (Extended Data Fig. 2b), indicating sufficient separation between the fractions. The enrichment and yield of fractions were comparable between WT and Tau35 preparations (Fig. 2b and Extended Data Fig. 2b). Specifically, we observed 10-fold and 10 to 20-fold enrichment in the WT/Tau35 LF fractions in the early and advanced disease stages respectively (Fig. 2b).

**Fig. 2:**
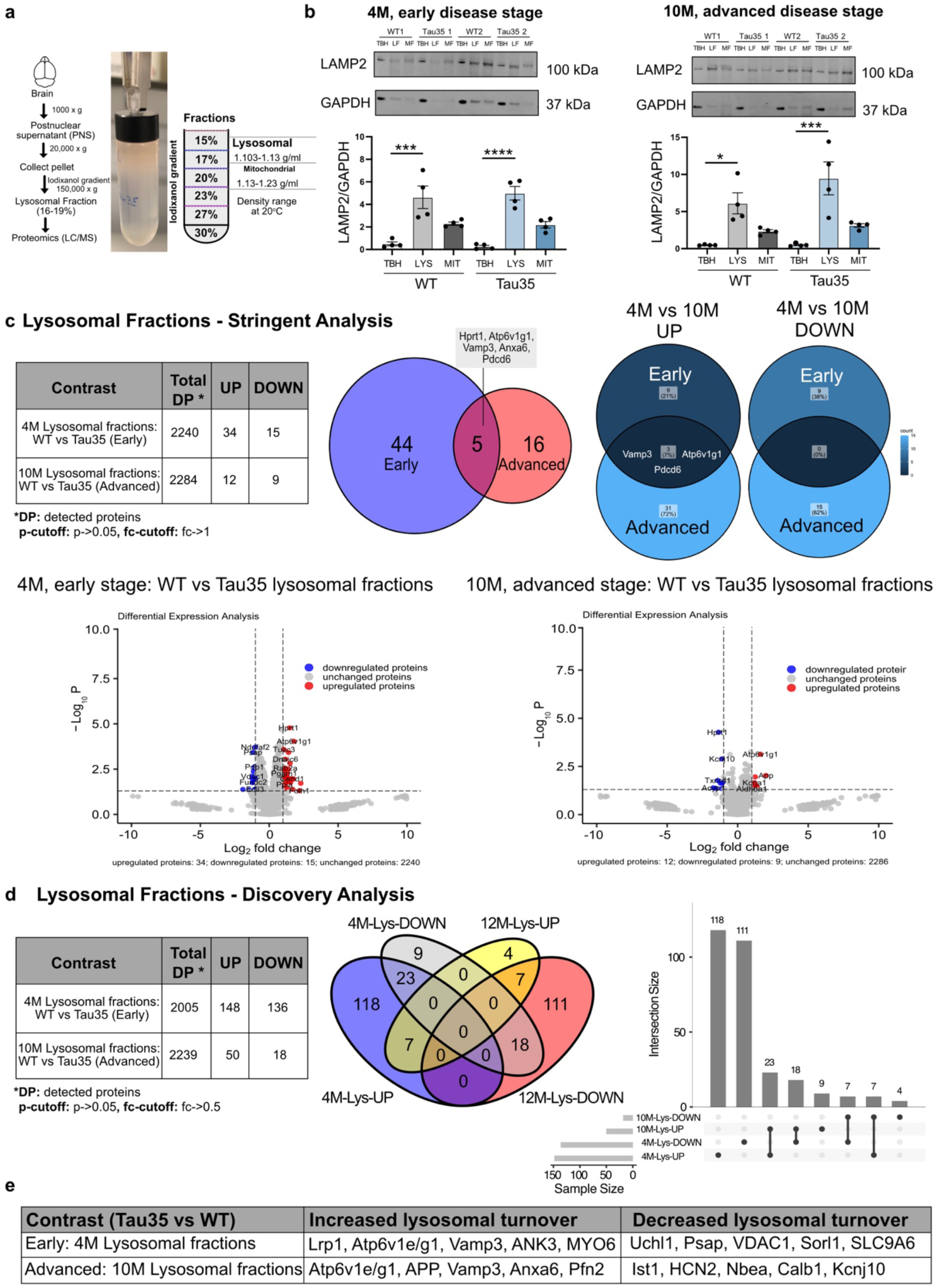
Lysosomal proteome alterations in Tau35 mouse brains during disease progression. **a,** Schematic diagram of the applied workflow for subcellular fractionation of lysosomes from mouse brain. Representative images of the discontinuous iodixanol gradient showing enriched fractions from WT and Tau35 mouse brain samples. The positions of the lysosomal and mitochondrial fractions within the gradient are marked, with the lysosome-enriched fraction selected for subsequent experiments. **b,** Western blots of total brain homogenates (TBH), lysosomal fractions (LF) and mitochondrial fractions (MF) from WT and Tau35 mice aged 4 and 10 months respectively, were probed with antibodies to LAMP2 and GAPDH. Western blotting analysis of mouse brain extracts and the different subcellular fractions from WT and Tau35 brain reveal the enrichment of the lysosomal marker protein LAMP2 in LF. Quantification of the blots is shown in the graphs as mean ± SEM, n = 4 brains per group. Ordinary one-way ANOVA, *P < 0.05, ***P < 0.001, ****P < 0.0001. LAMP2, lysosomal-associated membrane protein 2; GAPDH, glyceraldehyde 3-phosphate dehydrogenase; SEM, standard error of the mean; WT, wild type. **c,** The limma package in R was used to analyze differentially expressed proteins and produce Volcano plots (stringent analysis: p-cutoff: p->0.05, fc-cutoff: fc->1) from early (4M) and advanced (10M) sample cohorts. Table summarizing the differentially expressed proteins between early and advanced cohorts of wild-type (WT) and Tau35 mice. Venn diagrams from the two cohorts illustrate distinct protein alterations specific to each disease stage. Five proteins were significantly altered in both cohorts (Vamp3, Hprt1, Atp6v1g1, Anxa6, Pdcd6), three of which (Vamp3, Atp6v1g1, Pdcd6) exhibit upregulated lysosomal processing in Tau35 samples. Volcano plots show log-fold change (Log2FC) on the x-axis, and -log10 transformed *p*-values, adjusted for multiple testing using the Benjamini-Hochberg procedure on the y-axis. **d,** The limma package in R was used to analyze differentially expressed proteins (discovery analysis: p-cutoff: p->0.05, fc-cutoff: fc->0.5) from early and advanced sample cohorts. Table summarizing differentially expressed proteins between two cohorts, early and advanced, of wild-type (WT) and transgenic Tau35 mice. The upset plot is an interactive Venn diagram that shows the overlap of the four datasets (4M-Lys-UP, 4M-Lys-DOWN, 10M-Lys-UP and 10M-Lys-DOWN) by arranging them as bar charts of their frequencies. The heights of the vertical bars correspond to intersection size—the number of differentially expressed proteins that are shared among the corresponding datasets. For example, the first vertical grey bar shows that the “4M-Lys-UP” dataset contains 118 unique differentially expressed proteins, while the fifth vertical grey bar shows that the “10M-Lys-UP” dataset only contains 9 unique differentially expressed proteins. The third bar shows that “4M-Lys-UP” and “10M-Lys-UP” share 23 differentially expressed proteins, while the lack of an all-datasets bar shows that there are zero differentially expressed genes that are shared across all four datasets. **e,** Table summarizing key proteins that are up- or down-regulated in early and advanced cohorts. Vamp3, vesicle associated membrane protein 3; Atp6v1g1, Vacuolar ATPase H+ transporting V1 subunit G1; HPRT1, Hypoxanthine Phosphoribosyltransferase 1; ANXA6, Annexin A6; PDCD6, programmed cell death 6; Lrp1, low-density lipoprotein receptor-related protein 1; ANK3, ankyrin-G; Myo6, myosin VI; Uchl1, ubiquitin C-terminal hydrolase L1; Psap, prosaponin; Vdac1, voltage-dependent anion channel 1; Sorl1, sortilin-related receptor 1; Ndufaf2, NADH:ubiquinone oxidoreductase complex assembly factor 2; APP, amyloid precursor protein; Anxa6, annexin A6; Pfn2, profilin 2; Ist1, IST1 factor associated with ESCRT-III; Hcn2, hyperpolarization-activated cyclic nucleotide-gated channel 2; Nbea, neurobeactin; Calb1, calbindin 1.

We initially conducted a rigorous (p-cutoff: p->0.05, fc-cutoff: fc->1) analysis of the proteomic dataset obtained through liquid chromatography-tandem mass spectrometry (LC-MS/MS) using the limma package in R. This analysis identified five proteins that were differentially expressed in both the early and advanced disease stage cohorts (Early cohort: 34 up, 15 down; Advanced cohort: 12 up, 9 down) (Supplementary files 1 and 3), with three of these proteins (Vamp3, Atp6v1g1, Pdcd6) showing upregulated lysosomal processing in Tau35 samples across both stages (Fig. 2c).

We then relaxed the analysis parameters and conducted a discovery (p-cutoff: p->0.05, fc-cutoff: fc->0.05) analysis to identify intracellular mechanisms and pathways influenced by Tau35 overexpression. This approach revealed fifty-five proteins that were differentially expressed in both the early and advanced disease stage cohorts (Early cohort: 148 up, 136 down; Advanced cohort: 50 up, 18 down) (Fig. 2c, Extended Data Fig. 2c and Supplementary files 1 and 3).

The differentially expressed proteins identified through this discovery analysis using the limma package were subsequently cross-referenced with a curated list of autophagy and endo-lysosomal pathway-associated proteins^16^ to identify specific ALP components exhibiting differential expression in Tau35 brains. Notably, several proteins involved in autophagy and endo-lysosomal processes showed altered lysosomal processing in Tau35 samples across both stages (Early cohort: 31 altered, 13 up, 18 down; Advanced cohort: 7 altered, 6 up, 1 down). Many of them showed upregulated lysosomal processing in Tau35 samples across both stages (Extended Data Fig. 2d), alongside key modulators of tau (e.g. Lrp1) and intracellular organelle acidification (e.g. components of the V-ATPase) (Fig. 2e). Gene Ontology (GO) analysis of overrepresented terms in both cohorts revealed associations with mitochondria, energy/metabolism dynamics, and neuronal cellular homeostasis (Extended Data Fig. 2e-f), underscoring the interconnectedness of intracellular processes and the critical role of the endo-lysosomal system in maintaining cellular homeostasis during disease progression.

### Excessive endocytic activity disrupts lysosomal proteolytic function in human neuroblastoma SH-SY5Y tau models

To explore the impact of disease-associated tau cleavage on proteolysis and endocytosis, we assessed the proteolytic capacity of differentiated SH-SY5Y cells overexpressing N-terminally truncated Tau35, compared to WT control cells and cells overexpressing full-length tau (FL-Tau).

For this purpose, we generated six novel tauopathy cell lines, each stably expressing either Tau35 or full length (FL)-Tau fused to epitope tags or fluorescent reporters (construct maps are provided in Supplementary file 2), along with a control SH-SY5Y line stably expressing EGFP (Extended Data Fig. 3a). In this study, we focused on two of these newly generated SH-SY5Y tauopathy lines: one expressing Tau35-HA (referred to as Tau35) and one expressing Avi-FL tau (referred to as FL-tau) (Extended Data Fig. 3a). Western blotting was employed to analyze protein expression and confirm that the fusion proteins were full-length and not degraded upon overexpression (Extended Data Fig. 3b).

We then differentiated the cells, as differentiated SH-SY5Y cells exhibit a polarised morphology similar to neurons (protocol schematic, Extended Data Fig. 3c). Overexpression of Tau35 and FL-tau led to a significant increase in p62 levels (Fig. 3a), a marker of autophagic flux, while EGFP overexpression alone did not induce any changes in p62 levels (Extended Data Fig. 3d). This suggests that the observed p62 increase is specific to tau overexpression, making the Tau35- and FL-tau-overexpressing lines suitable models for studying neuronal-like alterations in autophagy and endo-lysosomal processes.

**Fig. 3:**
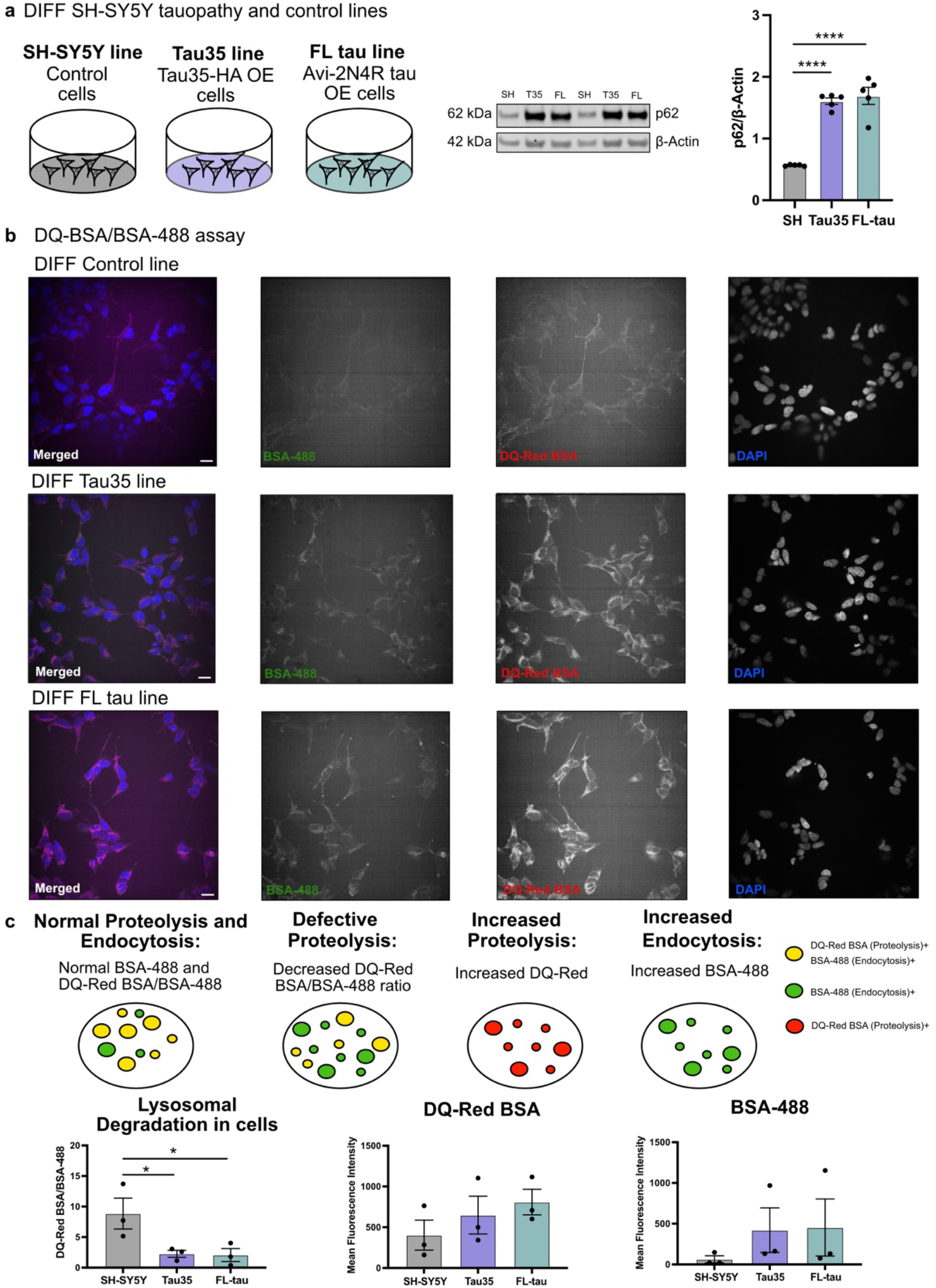
Endocytic dysregulation and defective proteolysis in SH-SY5Y tau models. **a,** Schematic illustrating the three differentiated SH-SY5Y cell lines; control SH-SY5Y cells, cells overexpressing (OE) Tau35-HA (referred to as Tau35) and cells overexpressing (OE) Avi-FL tau (referred to as FL-tau). Western blot of total cell lysates from control, Tau35, and FL-tau differentiated cells, collected at 14 days in vitro (DIV), were probed with antibodies to p62 and GAPDH. Quantification of the blots is shown in the graphs as mean ± SEM, *n* = 5 independent experiments. Ordinary one-way ANOVA, \*\*\*\**P* < 0.0001. p62/SQSTM1, Sequestosome-1; GAPDH, glyceraldehyde 3-phosphate dehydrogenase; SEM, standard error of the mean. **b,** Representative images of control, Tau35, and FL-tau differentiated cells at DIV14 after 4h incubation with both DQ-Red BSA and Alexa-488 BSA probes and counterstained with NucBlue reagent. Scale bar: 10 μm. **c,** Representative schematic illustrating normal, defective, and altered proteolysis and endocytosis. Quantifications showing the mean DQ-Red BSA/BSA-488 ratio, the mean DQ-Red BSA fluorescent intensity and the mean BSA-488 fluorescent intensity in control, Tau35, and FL-tau differentiated cells. Quantification of the blots is shown in the graphs as mean ± SEM, *n* = 3 independent experiments; SEM, standard error of the mean.

We treated the cells with both DQ-Red BSA and Alexa-488 BSA for 4 hours to investigate endo-lysosomal changes and counterstained with NucBlue reagent to facilitate the detection and segmentation of nuclei (representative images for each cell line are shown in Fig. 3b). Measurements of lysosomal proteolytic activity by DQ-Red BSA fluorescence normalised to BSA-488 in differentiated SH-SY5Y cells, revealed that Tau35- and FL-tau-overexpressing cells have significantly reduced activity compared to controls (Fig. 3b and c). Although tau overexpressing cells showed decreased lysosomal proteolytic activity, as indicated by lower DQ-Red BSA fluorescence normalized to Alexa-488-BSA fluorescence, the substantial reduction was primarily driven by a marked increase in endocytic activity as indicated by Alexa-488-BSA fluorescence (Fig. 3c). Tau overexpressing lines exhibited a 7-fold increase in Alexa-488-BSA fluorescence compared to controls, while DQ-Red BSA fluorescence increased by 1.6-fold (Tau35) and 2-fold (FL-tau). These findings suggest that excessive endocytic activity can negatively impact lysosomal proteolytic function in differentiated SH-SY5Y cells with neuronal-like morphology. Neuronal endosomal dysfunction, a characteristic feature of early Alzheimer’s disease (AD), involves increased activity within the endocytic pathway, driven by elevated expression of Rab GTPases^17^ or enhanced trafficking through early endosomes^18^. This dysfunction is a critical factor in AD progression and is consistent with our findings of markedly elevated endocytic activity, highlighting its potential contribution to the underlying disease pathology.

### Altered endo-lysosomal trafficking dynamics in human neuroblastoma SH-SY5Y tau models

To further investigate the impact of Tau35 overexpression on endo-lysosomal motility and trafficking, we performed super-resolution live-cell imaging of lysosomes using Lysotracker on differentiated control, Tau35 or FL-tau SH-SY5Y cells. Given the excellent cell permeability, high specificity, and low cytotoxicity of Lysotracker Deep Red (647), we utilized this probe for live-cell iSIM imaging. By applying the probe in iSIM imaging, we were able to record the dynamic processes of lysosome motility and interactions in live cells with a spatial resolution of ∼200 nm (representative images for each cell line are shown in Fig. 4a).

**Fig. 4:**
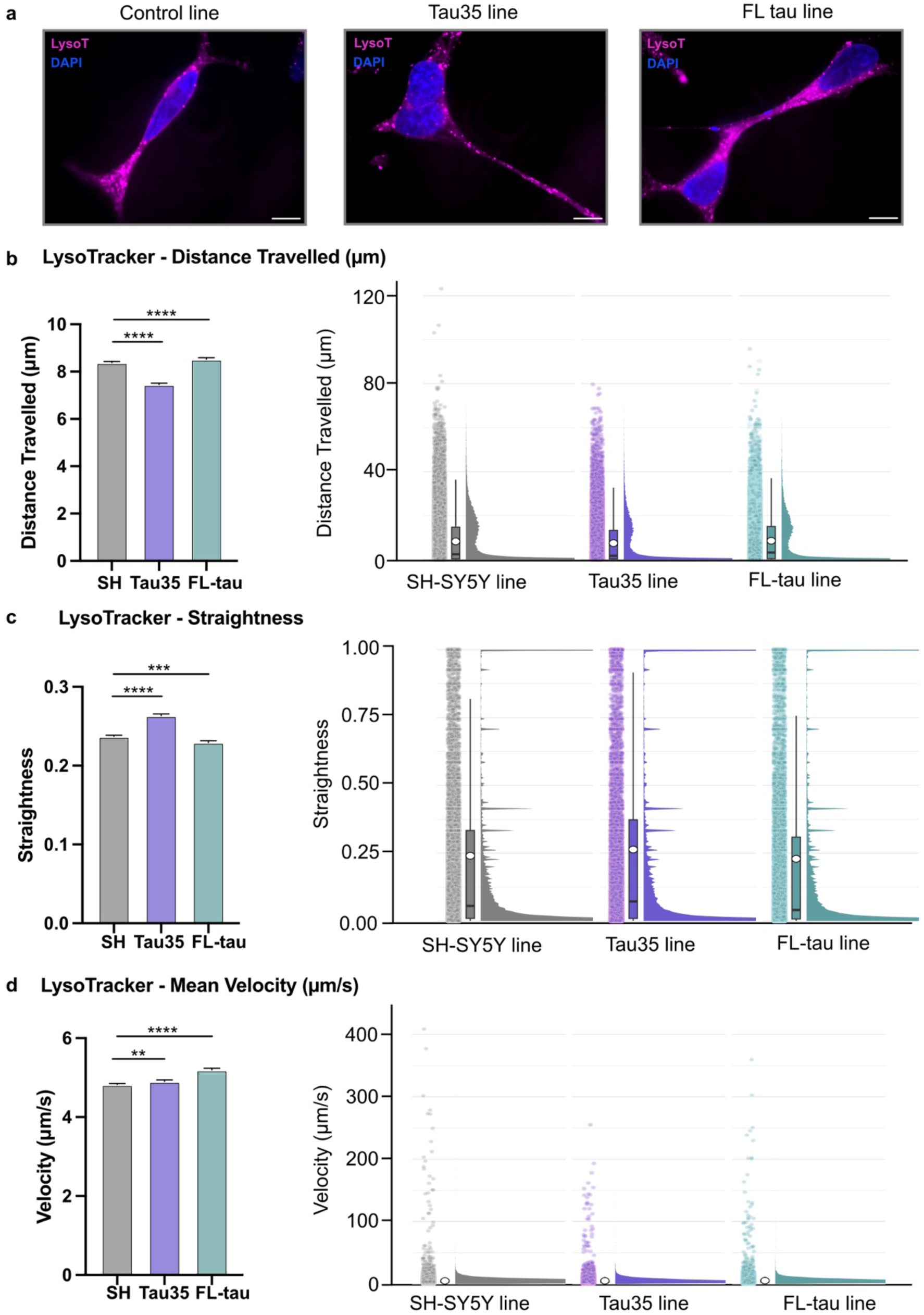
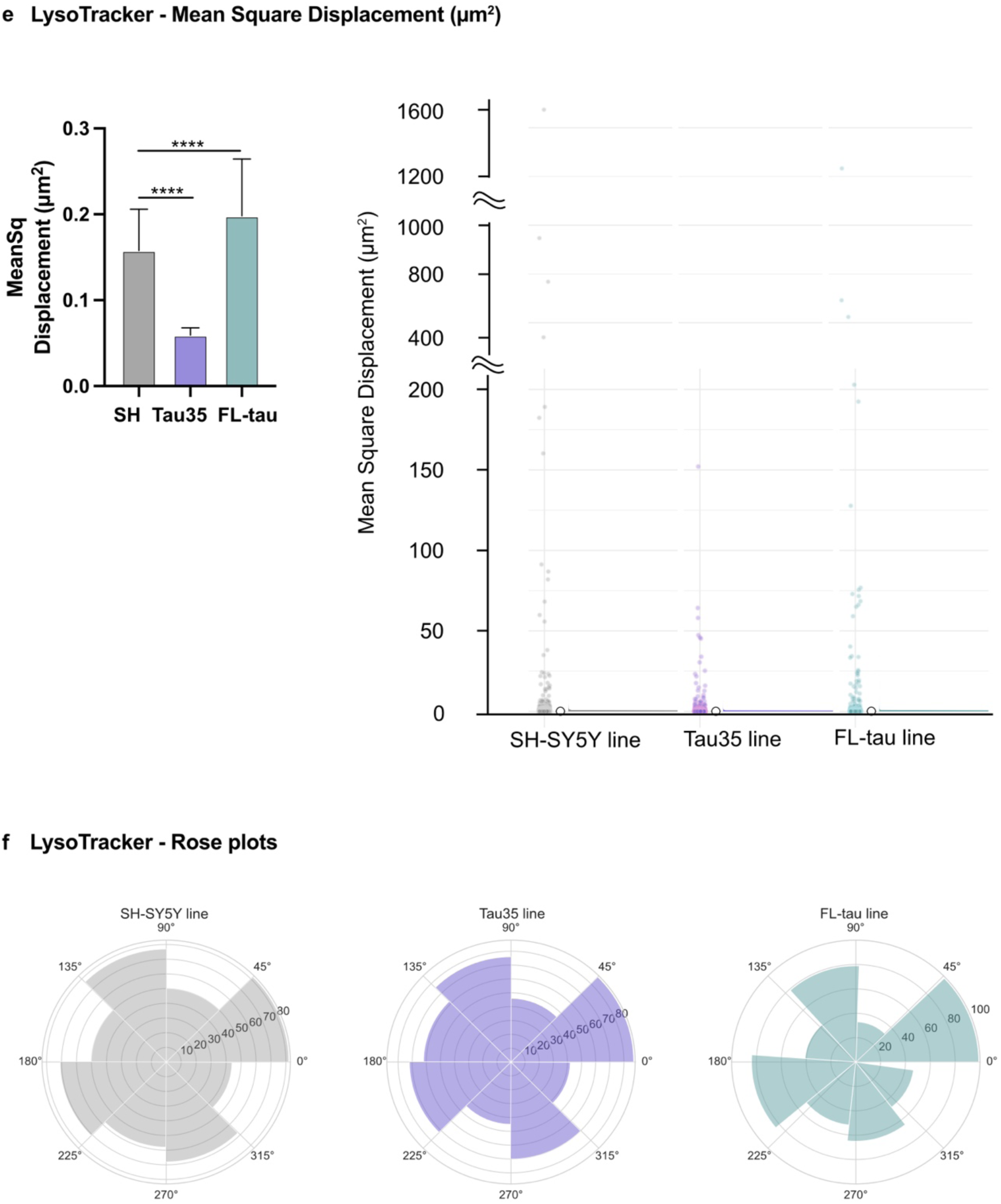
Lysosomal motility dynamics in SH-SYSY tau models. **a,** Super-resolution microscopy images of lysotracker staining in control, Tau35, and FL-tau differentiated cells at DIV14. Cells were stained with Lysotracker Deep Red (magenta) and counterstained for nuclei with NucBlue reagent (blue). Scale bar: 10 μm. **b,c,d,e** Quantification of lysosome motility parameters obtained by analyzing their trajectories. Simple box plots (left) and Raincloud plots (right) combining a split-half violin, raw jittered data points and a standard visualisation of central tendency with boxplot of Distance travelled (μm) (**b**), Straightness of the measured track (**c**), Mean Velocity (μm/s) (**d**) and Mean Square Displacement (μm^2^) (**e**) parameters in all three cell lines. **e**, A segmented Raincloud plot for Mean Square Displacement (μm²) is shown to better illustrate the distribution of values within specific ranges (0–200, 400– 1000, 1200–1600) across the different cell lines. **f,** Rose plots illustrating the angles of displacement for Lysotracker-detected lysosome motility in all three cell lines are presented. The bootstrapping technique was used to create random subsets of data (1,000 bootstrap replicates for each parameter), derived from the original dataset containing 24,000–43,000 data points (spots) per cell line, in order to minimize the impact of large datasets potentially obscuring differences. For **b-f**: Quantification of the blots is shown in the graphs as mean ± SEM, *n* = 6-7 independent experiments; Kruskal-Wallis’s test; \*\**P* < 0.01, \*\*\**P* < 0.001, \*\*\*\**P* < 0.0001.

Lysosomes, dynamic organelles with both stationary and mobile pools, play key biological roles, and their dysfunction is linked to diseases such as cancer, autoimmune and neurodegenerative disorders^19^. Therefore, understanding changes in lysosome motility can provide insights into how disruptions in lysosomal dynamics contribute to disease progression. Using Lysotracker imaging, we initially examined basic lysosomal parameters. No differences were observed in total cell area, the number of Lysotracker spots per cell or per area, or the mean spot intensity across all cell lines. (Extended Data Fig. 4a). We then investigated the trafficking of lysosomes, by acquiring images of stained live organelles at 100 frames per second (sec) over 10msec. Their motility was calculated by the assumption that immobile lysosomes move less than 0.2 µm in their track displacement. Lysosomes in the Tau35 cell line exhibited a shorter travel distance, moving 12.5% less than those in the control line. In contrast, lysosomes in the control and FL-tau lines travelled similar distances, with lysosomes in the FL-tau line covering slightly greater distances, moving on average 1.65% farther than those in the control line. It is important to note, however, that the highest absolute distance values, exceeding 100 μm, were observed in the control line, as compared to the Tau35 and FL-tau lines (Fig. 4b). We then investigated whether the lysosomes in the tested cell lines exhibited directed motion or a more diffusive pattern. Lysosomes in the control and FL-tau lines moved in a more organised manner, with the FL-tau line showing more linear movement compared to the others. In contrast, lysosomes in the Tau35 line displayed a more diffusive motion than those in the other lines (Fig. 4c). Additionally, the mean speed of lysosomes was higher in both tau overexpressing cell lines compared to control SH-SY5Y cells. Specifically, lysosomes in the Tau35 and FL-tau cell lines showed a 1.66% and 6.12% increase, respectively, in mean velocity. It is noteworthy, however, that the highest observed absolute speeds, exceeding 400 μm/s, were recorded in the control line, in contrast to the Tau35 and FL-tau lines. These findings suggest that lysosomes in the mobile pools of tau-overexpressing cells, particularly in the Tau35 line (with a maximum detected speed of 250 μm/s), exhibit slower movement compared to the control (Fig. 4d). This observation is further corroborated by the mean square displacement measurements, which show a 166% decrease in mean square displacement for lysosomes in the Tau35 line compared to the control line. Notably, the maximum mean square displacement value for lysosomes in the Tau35 line was only 150 μm^2^, which is substantially lower than the maximum values observed in the control (1593 μm^2^) and FL-tau (over 1291 μm^2^) lines, highlighting a significant reduction in lysosomal mobility in the Tau35 line (Fig. 4e and Extended Data Fig. 4f). Figure 4e shows a segmented Raincloud plot for Mean Square Displacement (μm²), highlighting value distributions within specific ranges (0–200, 400–1000, 1200–1600)across cell lines, while Extended Data Figure 4f presents the original Raincloud plot, illustrating the full range of values in proportion.

Next, to examine the directedness of the lysosomes, we constructed cellular histograms (rose plots). Rose plots depicting the displacement angles of Lysotracker-detected lysosome motility across the three cell lines are presented, illustrating the movement of individual lysosomes (spots) from their initial positions at the start of video acquisition. Lysosomes in the control line appeared to travel farther from their starting positions compared to those in the tau-overexpressing lines. Among the tau-overexpressing lines, lysosomes in the FL-tau line exhibited the least deviation, with most of their movement restricted to range of angles between 0°-45°, 90°-135° and 175°-220° (Fig. 4f). These findings may reflect the stochastic shared movement orientation among lysosomes, which reduces independent angle sampling and produces a multimodal distribution of movement angles. The bootstrapping technique, renowned for improving statistical analysis by facilitating robust comparisons across scenarios and distributions while supporting careful hypothesis evaluation^20^, was employed to generate 1,000 random subsets per parameter from datasets containing 24,000–43,000 data points per cell line. This approach minimized the risk of large datasets masking subtle differences and validated our findings on lysosome motility (Extended Data Fig. 4b-e).

## Discussion

Tau normally regulates microtubule assembly and axonal transport by modulating dynein and kinesin motor proteins. However, under pathological conditions, tau through mutation, truncation or posttranslational modifications (PTMs), detaches from microtubules and forms aggregates^21^. Evidence from Alzheimer’s disease (AD)^22,23^, frontotemporal dementia^24–26^ and Niemann-Pick type C (NPC), a lysosomal storage disorder^27^, suggests that endo-lysosomal dysfunction contributes to tau aggregate accumulation and neurofibrillary tangle (NFT) formation.

Tau truncation has garnered attention for its potential role in disease progression, particularly through mechanisms unique to specific tau fragments^5,6^. Overexpression of the Tau35 fragment in various models results in progressive tau phosphorylation, synaptic alterations, behavioural abnormalities, and disruptions in protein degradation pathways, including the proteasome, lysosomes, and autophagy^9–13^. The C-terminal Tau35 fragment, associated with human tauopathies^7,8^, disrupts kinase activity, synaptic processes and lysosomal protein degradation; with late-stage Tau35 mice showing increased p62 and LC3I/II levels, alongside reduced active cathepsin D^9^.

Our findings reveal that early-stage pathology (4M) in Tau35 mouse brains is marked by a selective reduction in mature CTSB protein expression, which persists into advanced stages (10M), alongside an increase in mature CTSL protein expression. This imbalance in cathepsin expression may promote tau pathology and drive disease progression^14^. Notably, unlike earlier studies in the Tau35 model^9,13^, we did not observe changes in mature CTSD expression at the examined timepoints, potentially underscoring the delicate balance among cathepsins during disease progression. In advanced stages (10M), the CTSB/CTSL disequilibrium is further associated with a significant reduction in LAMP2 levels, aligning with prior observations in the Tau35-overexpressing CHO cell model^13^. The decline in LAMP2 likely reflects compromised lysosomal function, as previously described in mice^28^.

Autophagic-endolysosomal networks (AELN) are essential for tau clearance, including its degradation, release, and uptake by neurons and glial cells, ensuring proper tau homeostasis^29^. Therefore, increasing evidence indicates that alterations or dysfunctions in the endo-lysosomal and autophagic systems may coincide with or even precede tau development^30^.

To examine the effect of Tau35 overexpression on the lysosomal proteome, lysosome-enriched brain fractions from WT and Tau35 mice at early (4 months) and advanced (10 months) tau pathology stages were analysed using discontinuous iodixanol gradient centrifugation, followed by rigorous and discovery proteomic analyses with stringent and lenient selection criteria.

Our stringent proteomic analysis of Tau35 lysosome-enriched fractions identified several differentially expressed proteins, which can be grouped into four key categories with roles in AELN processes and tau processing.

**(1) pH Maintenance and Ionic Homeostasis:** Proteins such as ATP6V1E1, ATP6V1G1, and SLC9A6 were dysregulated. ATP6V1E1 and ATP6V1G1, subunits of v-ATPase, play essential roles in lysosomal acidification and autophagic flux maintenance, with their dysfunction linked to impaired autophagy and tau aggregation. Notably, a brain proteomic analysis found that the expression of v-ATPase H+ Transporting V1 Subunit E1 (ATP6V1E1), which regulates the acidification of intracellular organelles, is altered during the progression of tau tangle pathologies^31,32^. SLC9A6 regulates endosomal pH and volume, and its loss-of-function mutations are associated with tau inclusions. Interestingly a SLC9A6 gene mutation has been reported to induce abnormalities in vesicular targeting and tau aggregation, with potential implications for tauopathies^33^.
**(2) Tau Regulation:** Proteins like LRP1, SORL1, FKBP1A, and UCHL1 were altered. SORL1, a receptor involved in protein trafficking, plays a role in the trafficking and seeding of pathogenic tau^34^. LRP1, often downregulated in tauopathies, regulates tau clearance and endolysosomal trafficking^35^. FKBP1A modulates tau phosphorylation and calcium homeostasis^36^, while UCHL1, vital for proteostasis, contributes to tau aggregation and oxidative stress when downregulated^37^.
**(3) Intracellular Trafficking:** Proteins such as Myo6, LRP1, and VAMP3 were implicated. Myo6, which plays a role in vesicle transport and lysosomal degradation^38,39^, has been shown to co-localize with fibrillary tau protein^40^. VAMP3 dysregulation has been reported to impair vesicle transport, autophagy, and consequently tau clearance. In late-onset Alzheimer’s disease (LOAD), PICALM dysregulation disrupts the interaction and endocytosis of SNARE proteins, including VAMP3, further affecting tau removal^41^. Additionally, LRP1 dysfunction is recognized as a contributor to increased tau accumulation^35^, as described above.
**(4) Cytoskeletal Dynamics:** ANK3 and PFN2, critical regulators of neuronal structure, were also impacted. ANK3, a gene essential for microtubule stability and neuronal integrity, plays a key role in maintaining neuronal function. Notably, ankyrins have been shown to interact with tau in Drosophila, leading to reduced lifespan and memory deficits^42^. Similarly, PFN2, through its role in actin cytoskeleton dynamics and intracellular trafficking, has been reported to interact with and modulate tau^43^.

To explore the pathways impacted by Tau35 overexpression in greater detail, we relaxed the analysis parameters and performed a discovery analysis. Differentially expressed proteins were compared against a curated list of autophagy and endo-lysosomal pathway-associated proteins^16^, uncovering disruptions in ALP components at both early and advanced stages of disease. Consistent with our stringent analysis, key categories such as endo-lysosomal ionic homeostasis, endo-lysosomal trafficking, and cellular homeostasis were highlighted. The discovery analysis further revealed additional pathways, such as PI3K-related processes, heat shock protein regulation, and Rab-dependent autophagosome formation. Notably, recent research demonstrated that Rab5 overactivation, independent of APP-βCTF, can replicate key aspects of Alzheimer’s disease, including synaptic plasticity impairments and tau hyperphosphorylation^44^. Gene Ontology (GO) analysis further identified overrepresented terms associated with mitochondria, energy/metabolism, and neuronal cellular homeostasis, underscoring the interconnected nature of these processes and the essential role of the endo-lysosomal system in maintaining cellular balance during disease progression.

We next explored the impact of tau cleavage on proteolysis and endocytosis by assessing the proteolytic capacity of differentiated SH-SY5Y cells overexpressing either N-terminally truncated Tau35 or full-length tau. Our results showed that overexpression of tau, both Tau35 and full-length tau, led to a significant increase in p62 levels, a marker of autophagic flux, indicating altered autophagy and endo-lysosomal processes. Additionally, tau-overexpressing cells exhibited a notable reduction in lysosomal proteolytic activity, which was primarily attributed to a substantial increase in endocytic activity.

Successful autophagic degradation of aggregated proteins depends on proper early and late endosomal sorting and maturation^45^. Additionally, autophagosomal biogenesis shares regulatory mechanisms with endosomal compartments and may rely on recycling endosomes for membrane sources. Key regulators involved in endosomal recycling also play a role at the intersection of these pathways^45–47^. Together, these findings highlight the interconnected nature of autophagic and endo-lysosomal trafficking, suggesting they cannot be considered separate, independent processes^48^. These findings further suggest that excessive endocytic activity, characteristic of early Alzheimer’s disease^17,18^, could impair lysosomal function and contribute to disease pathology by disrupting cellular homeostasis^30,41^.

Lysosomes are dynamic organelles with distinct pools: a stationary pool near the microtubule-organising centre and a mobile pool at the cell periphery. Their movement is essential for various cellular functions, including autophagy, protein degradation, and organelle turnover^19^. Disruptions in lysosome motility, such as impaired trafficking or distribution, can lead to the accumulation of toxic proteins, cellular stress, and organelle dysfunction, which contribute to diseases like neurodegenerative disorders, cancer, and autoimmune conditions^49^. Understanding lysosome motility changes can provide insights into disease progression and help identify therapeutic strategies to restore cellular function^50^.

To explore the specific impact of truncated tau overexpression on endo-lysosomal dynamics, we performed live-cell imaging to monitor Lysotracker trafficking in differentiated SH-SY5Y cells overexpressing truncated (Tau35) or full-length tau, alongside control SH-SY5Y cells. Super-resolution Lysotracker imaging revealed no significant differences in basic lysosomal parameters, such as spot number or mean fluorescence intensity, across the cell lines. However, analysis of lysosomal movement showed that lysosomes in the Tau35 line exhibited significantly slower movement compared to the control, with a marked reduction in travel distance and mean square displacement. Conversely, lysosomes in both the control and full-length tau lines displayed more organised, directed movement. Taken together these findings suggest that truncated tau overexpression impairs lysosomal motility, potentially disrupting lysosomal function, which may contribute to cellular dysfunction in tauopathies.

In conclusion, this correlative study investigated the effects of Tau35 overexpression on proteolytic pathways, autophagy, and endo-lysosomal processes. The findings highlight that truncated tau induced early pathological changes, such as increased endocytosis, impaired proteolysis, and lysosomal motility abnormalities. This research provides new insights into the mechanisms underlying Tau35-induced neurodegeneration, emphasising the critical role of endo-lysosomal processes in tauopathy progression. Key processes identified include endo-lysosomal trafficking, cellular homeostasis, and autophagosome formation, with disrupted lysosomal function linked to excessive endocytic activity and impaired lysosomal motility. These results suggest that altered endo-lysosomal dynamics may contribute to tauopathy pathology by disrupting cellular homeostasis, providing a foundation for future therapeutic strategies targeting these pathways.

## Methods

### Mice

#### Ethics statement

All experimental procedures adhered to the Animals (Scientific Procedures) Act, 1986, and received approval from the local ethical review committee. The study was conducted in compliance with ARRIVE guidelines 2.0^51^. Mice were generated via targeted knock-in of the Tau35 cDNA construct at the Hprt locus on the X chromosome, under the regulation of the human tau promoter, as previously described^9^. The construct encodes an N-terminally truncated fragment of wild-type human tau protein (amino acids 187-441) with a haemagglutinin (HA) tag appended to the C-terminus. Male hemizygous transgenic and wild-type mice were used exclusively in this study to circumvent potential complications arising from incomplete X chromosome inactivation in female mice.

### Preparation of mouse brain homogenates for western blots

Mice were euthanized by cervical dislocation, and brains were promptly extracted, snap-frozen on dry ice, and stored at -80°C. Brain tissue was lysed through ultrasonication (parameters: 40% amplitude, 4-second pulses, 30-second duration) using a Vibra-Cell ultrasonic liquid processor (Model no. VCX 130, Sonics and Materials, Newton, CT, USA) in ice-cold RIPA buffer (150 mM NaCl, 1 mM ethylenediaminetetraacetic acid [EDTA], 50mM Tris-HCl, 1% [v/v] NP-40, 0.5% [w/v] sodium deoxycholate, 0.1% [w/v] sodium dodecyl sulfate [SDS]) supplemented with protease (cOmplete^™^, EDTA-free, Merck Millipore, Cat#11873580001) and phosphatase (PhosSTOP, Sigma-Aldrich, Cat#4906845001) inhibitors. This process was performed in a refrigerated chamber and repeated three times, with incubation on ice for 3 min between cycles. The lysates were centrifuged at 10,000g for 15 min at 4°C, and the resulting supernatants were stored at -80°C for further analysis.

### Isolation of lysosome enriched fractions from mouse brain tissue

Mice were sacrificed by cervical dislocation, and their brains were promptly extracted, snap-frozen on dry ice, and stored at -80°C. Lysosome enriched fractions were isolated from Tau35 and wild-type (WT) control mouse brain tissue using the Lysosome Enrichment Kit for Tissue and Cultured Cells (Thermo Fisher Scientific, Cat# 89839) following the manufacturer’s instructions. Briefly, 500mg of brain tissue per sample was washed with 1x Phospate Buffered Saline (PBS), minced and homogenized in 1mL of Lysosome Enrichment Reagent A using a glass Dounce homogenizer. Next, 1mL of Lysosome Enrichment Reagent B was added and the mixture was inverted 5-6 times for thorough mixing. All steps were conducted in a refrigerated chamber and performed on ice. Samples were then centrifuged at 500 x g for 10 min at 4°C, and the supernatants were collected and kept on ice. In a 6.3 mL quick-seal polypropylene ultracentrifuge tube (Beckman Coulter Inc., Cat# 345830), OptiPrep (Iodixanol) gradients were prepared in descending concentrations: 30%, 27%, 23%, 20% and 17%. The sample was diluted in 15% OptiPrep media and overlaid on top of the density gradients. After ultracentrifugation at 145,000 x g for 2h at 4°C, the lysosome pellet was washed in 2-3 volumes of 1xPBS to reduce the OptiPrep media concentration and collected by adding 1mL of Gradient Dilution Buffer. Lysosome purity was evaluated by western blotting using antibodies against the lysosomal marker LAMP2. Samples were stored at -80°C in lysosomal buffer (2% CHAPS (3-[(3-**ch**olamidopropyl)dimethyl**a**mmonio]-1-**p**ropane**s**ulfonate) in 1x Tris-Buffered Saline (TBS)) for subsequent analysis.

### Preparation of lysosome enriched fractions for proteomic analysis

Sample lysis, reduction, alkylation and enzymatic digestion were performed before peptide purification. The protein concentration of lysosome-enriched fractions from transgenic and control mouse brain samples was measured using the Bradford Protein Assay (Pierce, Thermo Fisher Scientific, Cat# 23200) and 12μg of protein were used for each replicate. To improve protein separation 8M urea and 5mM Dithiothreitol (DTT) were added, followed by incubation at 37°C, in a thermomixer (750rpm) for 30min. To alkylate free cysteines iodoacetamide (IAA) was added to a final concentration of 20mM, followed by vortexing and sample incubation at room temperature in the dark for 20min. Proteins were then precipitated by addition of methanol/chloroform/18MΩ water (4:1:3), vigorous vortexing and centrifugation at 14,000 rpm for 1min. The liquid phases were discarded and the protein pellets were washed once with methanol and centrifuged at 14,000 rpm for 1min, followed by removal of methanol. Protein pellets were then air-dried and solubilised in 0.2M EPPS (4-(2-Hydroxyethyl)-1-piperazinepropanesulfonic acid, 4-(2-Hydroxyethyl) piperazine-1-propanesulfonic acid, *N*-(2-Hydroxyethyl)piperazine-*N*ʹ-(3-propanesulfonic acid)) buffer. Proteins were digested by addition of trypsin (0.5μg, Thermo Fisher Scientific, Cat# 90057), rigorous vortexing and overnight incubation in a thermomixer (750rpm, ThermoMixer®C, Eppendorf SE). The following day, samples were dried to completion in a Speedvac (Thermo Fisher Scientific, IL, USA) and peptides were cleaned up using C18 spin columns (Cat# 89852; Thermo Fisher Scientific) according to manufacturer’s instructions. Briefly, samples were resuspended in 300ul of 0.1% trifluoroacetic acid (TFA) and eluted in 50% acetonitrile/0.1% TFA. Following elution, samples were dried to completion by Speedvac and stored at -80°C.

### Proteomic Analysis

#### Liquid Chromatography with Tandem Mass Spectrometry (LC-MS/MS)

The extracted peptide samples were individually resuspended in MS sample buffer (2% acetonitrile (ACN) in 0.05% formic acid (FA)) to a concentration of 1mg/ml, 6μl of which was injected to be analysed by LC-MS/MS. Chromatographic separation was performed using a U3000 UHPLC NanoLC system (ThermoFisherScientific, UK). Peptides were resolved by reversed phase chromatography on a 75μm C18 Pepmap column (50cm length) using a three-step linear gradient of 80% acetonitrile in 0.1% formic acid. The gradient was delivered to elute the peptides at a flow rate of 250nl/min over 60 min starting at 5% B (0-5 minutes) and increasing solvent to 40% B (5-40 minutes) prior to a wash step at 99% B (40-45 minutes) followed by an equilibration step at 5% B (45-60 minutes).

The eluate was ionized by electrospray ionization using an Orbitrap Fusion Lumos (ThermoFisherScientific, UK) operating under Xcalibur v4.3. The instrument was first programmed to acquire using an Orbitrap-Ion Trap method by defining a 3s cycle time between a full MS scan and MS/MS fragmentation by collision induced dissociation. Orbitrap spectra (FTMS1) were collected at a resolution of 120,000 over a scan range of m/z 375-1800 with an automatic gain control (AGC) setting of 4.0e5 (100%) with a maximum injection time of 35 ms. Monoisotopic precursor ions were filtered using charge state (+2 to +7) with an intensity threshold set between 5.0e3 to 1.0e20 and a dynamic exclusion window of 35s and ±10 ppm. MS2 precursor ions were isolated in the quadrupole set to a mass width filter of 1.6 m/z. Ion trap fragmentation spectra (ITMS2) were collected with an AGC target setting of 1.0e4 (100%) with a maximum injection time of 35 ms with CID collision energy set at 35%.

### Raw Proteomic Data Processing and Analysis

Raw mass spectrometry data were processed into peak list files using Proteome Discoverer (ThermoScientific; v2.5). The raw data file was searched using the Sequest^52^ search algorithm against the Uniprot Mouse Taxonomy database (37,716 entries). Database searching was performed at a stringency of 1% false discovery rate (FDR) including a decoy search. Posttranslational modifications for carbamidomethylation (C, static), oxidation (M, variable) and phosphorylation (S, T & Y; variable) were included in the database search. Protein/peptide identification, along with peak intensities, were exported as Excel files for subsequent analysis using R. The raw proteome data (intensity values) were log-2 transformed and converted into an expression set object (iBAQ values). Differential expression analysis at the peptide level was conducted using R (R Core Team (2023). _R: A Language and Environment for Statistical Computing_. R Foundation for Statistical Computing, Vienna, Austria. https://www.R-project.org/), RStudio (Version: 2024.12.0+467) (Posit team (2024). RStudio: Integrated Development Environment for R. Posit Software, PBC, Boston, MA. URL http://www.posit.co/), and the limma package^53,54^, which employs empirical Bayes methods to generate accurate variance estimates, even with a small sample size. Lists of differentially expressed proteins were generated following stringent (p-value: p_cutoff <- 0.05, fold change: fc_cutoff <- 1) and lenient (p-value: p_cutoff <- 0.05, fold change: fc_cutoff <- 0.5) criteria (discovery analysis) and used the Cluster Profiler package^55^ to perform gene ontology (GO) analysis in R, aiming to identify the biological processes and molecular functions that are impacted in lysosome-enriched fractions. The complete set of detected proteins, along with the differentially expressed proteins identified via discovery analysis using the limma package, was also cross-referenced with a curated list of autophagy and endo-lysosomal pathway-associated proteins^16^, aiming to pinpoint specific ALP components exhibiting differential expression in Tau35 brains.

### Western Blots

Protein concentrations were measured using the bicinchoninic acid (BCA) protein assay following the manufacturer’s protocol (Pierce™ BCA Protein Assay Kit, Thermo Fisher Scientific, Cat# 23225). Samples prepared in NuPAGE LDS Sample Buffer (4X) (Thermo Fisher Scientific, Cat# NP0007) were heated at 95°C for 10 minutes and resolved using Bolt™ Bis-Tris Plus Mini Protein Gels (4–12%, 1.0 mm, WedgeWell™ format; Thermo Fisher Scientific, Cat# NW04125BOX or NW04127BOX). Proteins were transferred onto nitrocellulose membranes (Amersham Protran 0.45 NC, Cytiva, Cat# 10600007) and blocked with Intercept® (TBS) Blocking Buffer (LI-COR Biosciences, Cat# 927-60001). Membranes were incubated with primary antibodies overnight at 4°C, washed with TBS containing 0.02% (v/v) Tween 20, and treated with fluorophore-conjugated secondary antibodies for antigen detection. Imaging was performed using the Odyssey® imaging system (LI-COR Biosciences), and ImageStudio™ Lite software (LI-COR Biosciences) was used for western blot quantification.

## Human neuroblastoma (SH-SY5Y) cell lines

### Cell Culture

Human neuroblastoma SH-SY5Y cells were sourced from the American Type Culture Collection (ATCC, passage 16) and cultured in DMEM/F12 (Dulbecco’s Modified Eagle Medium/Nutrient Mixture F-12) GlutaMAX^TM^ (Thermo Fisher Scientific, Cat# 31331-028) supplemented with 10% heat-inactivated serum (Gibco, Cat# 16000044), and 1% penicillin-streptomycin (Gibco, Cat# 15140922). Cells were cultivated in T75 flasks, maintained at 37°C in a humidified incubator with 5% CO₂ and kept below ATCC passage + 3 to avoid cell senescence.

### Cloning and Expression of Fusion Proteins in SH-SY5Y cells

The codon-optimised synthetic gene sequences (Tau35-HA, Tau35-mKO_2_, mKO_2_-Tau35, Avi-FL tau, FL tau-eGFP and eGFP-FL tau) were generated by GenScript and subsequently cloned into the pLVX-TetOne-Puro vector (detailed map sequences for all plasmids are included in Supplementary data 2). All constructs were confirmed through restriction enzyme digestion and sequencing. Low passage, undifferentiated SH-SY5Y cells were transiently transfected with each of the newly generated plasmids using Lipofectamine 3000 reagent (Thermo Fisher Scientific, Cat# L3000015) following the manufacturer’s protocol.

### Cell Transfection, Clonal Isolation and Stable SH-SY5Y Cell Line Production

Human neuroblastoma SH-SY5Y cells were cultured in 6-well plates (1×10^4^ cells/well) and maintained until they reached 70–80% confluence. At this point, the cells were transfected with the plasmids of interest [pLVX-TetOne-Puro - (Tau35-HA, Tau35-mKO_2_, mKO_2_-Tau35, Avi-FL tau, FL tau-eGFP and eGFP-FL tau) generated by GenScript or pLVX-TetOne-Puro-GFP (Addgene plasmid # 171123; http://n2t.net/addgene:171123;RRID:Addgene_171123)^56^] using Lipofectamine 3000 reagent (Thermo Fisher Scientific, Cat# L3000015) according to manufacturer’s instructions. Approximately 48 hours post-transfection, puromycin dihydrochloride (2 µg/ml, Thermo Fisher Scientific, Cat# A1113803) was added to the culture medium to initiate selection. The medium was then replaced every 2–3 days with fresh selection medium containing 2 µg/ml puromycin for up to 2 weeks to ensure selection. After selection with puromycin, stable cell pools were cultured in the presence of 1 µg/ml doxycycline hyclate (Sigma-Aldrich, Cat# D5207) for five days to induce construct expression. Colony isolation was performed as follows: colonies of interest were marked on the underside of the culture plate. The tissue culture medium was carefully aspirated, and the plate was washed with Dulbecco’s phosphate-buffered saline (DPBS - no calcium, no magnesium, Thermo Fisher Scientific, Cat# D5207). Sterile cloning discs (SP Bel-Art, Cat# F37847-0001) were dipped in trypsin solution (Trypsin 0.25% EDTA, Thermo Fisher Scientific, Cat# 25200072) and placed onto the marked colonies. The plate was incubated for 3 to 10 minutes to ensure complete trypsinization of the colonies. Meanwhile, a 24-well plate containing 1 mL of growth medium supplemented with puromycin (1.5µg/ml) and doxycycline (1µg/ml) was prepared for colony transfer. After trypsinisation, the cloning discs were removed and transferred into the prepared 24-well plate. The cells adhered to the discs and were maintained in individual wells. Once the cells reached near confluence, they were trypsinised and passaged into larger culture vessels for further expansion and characterization. The cloning discs were left in the 24-well plate for subsequent monitoring. To assess colony quality, one well of a 6-well plate was used for routine passaging, while a second well of a 24-well plate, containing a 13-mm coverslip, was used for colony screening. The coverslips were fixed after 1–2 days to evaluate colony morphology and assess whether the colonies should be retained or subjected to further sub-cloning to achieve >95% homogeneity.

For those colonies of interest, cells were passaged from the 6-well plate into at least three separate dishes: one for cryopreservation as passage 0, one for continued culture (if the cell line was already clonal), and one for sub-cloning (if necessary). Sub-cloning was performed by seeding the cells at clonal density. Once a desired clone or sub-clone was identified, a reduced concentration of puromycin (0.75-1µg/ml) was used for routine maintenance. Following selection, clones were cultured in the presence of 1 μg/ml doxycycline and 1 μg/ml puromycin for five days and expression was analysed by western blotting to evaluate expression levels and confirm that fusion proteins are full-length and not degraded when overexpressed (supplementary figure 3B). In brief, cells were lysed in RIPA Lysis and Extraction buffer (89900, ThermoFischer Scientific) supplemented with protease (cOmplete^™^, EDTA-free, Merck Millipore, Cat# 11873580001) and phosphatase (PhosSTOP, Sigma-Aldrich, Cat# 4906845001) inhibitors. Protein levels were assessed by the bicinchoninic acid protein assay, according to the manufacturer’s instructions (Pierce™ BCA Protein Assay Kit, ThermoFisher Scientific, Cat# 23227). Lysates were boiled in NuPAGE LDS Sample Buffer (4X) and processed as described above.

### SH-SY5Y Cell Line Differentiation

SH-SY5Y cell lines were re-cultured, expanded for 2-3 passages, and then differentiated according to the Shipley *et al.*^57^ protocol, with minor modifications, including the replacement of retinoic acid with the synthetic retinoid EC23 (Sigma-Aldrich, Cat# SML2404). The detailed media formulations are listed in the tables below, and a schematic representation of the SH-SY5Y differentiation protocol is provided in Extended Data Figure 3 (Extended Data Fig 3C). Stock solutions of the synthetic retinoid EC23 are prepared by dissolving EC23 in DMSO to a final concentration of 5–10 mM. Aliquots of the stock solutions are stored at −20°C, protected from light.

The differentiation process spanned 15-20 days and involved gradual serum starvation followed by the introduction of neurotrophic factors. On day 0, 25,000-100,000 cells were plated onto uncoated 35mm dishes, or 50,000-100,000 cells per well in 6-well plates. For 96-well plates, 4,000-5,000 cells were seeded per well. These ranges are indicative, with the final cell number determined by the experimental design. Differentiation began on day 1 with the replacement of basic growth media by differentiation media #1, which was refreshed on days 1, 3, and 5. On day 7, cells were split 1:1 and replated onto uncoated dishes or plates or Ibidi chambers in differentiation media #1. On day 8, differentiation media #2 was introduced and replaced again on day 10. On day 11, cells were split 1:1 and plated onto ECM (MAXGEL (TM) ECM Mixture, Merck, Cat# E0282)-coated dishes, plates, or Ibidi chambers in differentiation media #2. From days 12, 15, and 18, the media was replaced with differentiation media #3, supplemented with 1 μg/ml doxycycline and 1 μg/ml puromycin to induce transgene expression. Fully differentiated SH-SY5Y cells were utilized on days 14, 15 and 20 for western blot, qPCR, immunocytochemistry (ICC), and live cell imaging experiments.

### Media Recipes

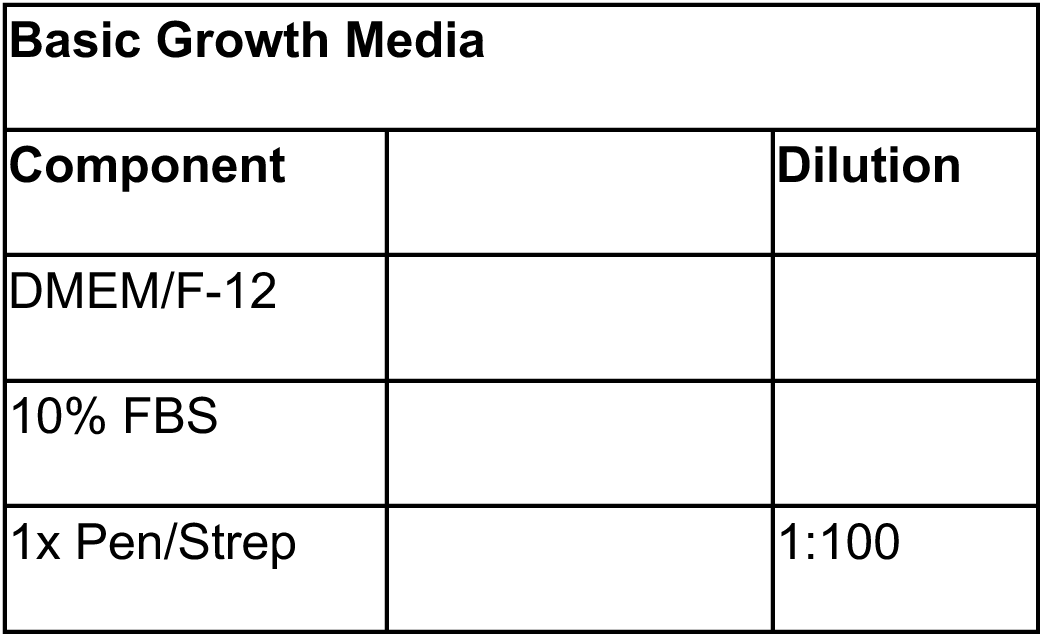

| Basic Growth Media |  |  |
| --- | --- | --- |
| Component |  | Dilution |
| DMEM/F-12 |  |  |
| 10% FBS |  |  |
| 1x Pen/Strep |  | 1:100 |

| Differentiation Media #1 |  |  |
| --- | --- | --- |
| Component | For 25ml | Dilution |
| DMEM/F-12 | Up to 25ml |  |
| 2.5% FBS | 625µl |  |
| 1x Pen/Strep | 250µl | 1:100 |
| 10µM EC23 | 10µl/5µl<br>(5mM stock)/(10mM stock)10µl/5µl | 1:500/<br>1:1000 |
| *Add EC23 immediately prior to use |  |  |
| **Once EC23 is added, keep media for max 3 days |  |  |

| Differentiation Media #2 |  |  |
| --- | --- | --- |
| Component |  | Dilution |
| DMEM/F-12 | Up to 25ml |  |
| 1% FBS | 250µl |  |
| 1x Pen/Strep | 250µl | 1:100 |
| 10µM EC23 | 10µl/5µl<br>(5mM stock)/ (10mM stock) | 1:500/<br>1:1000 |
| *Add EC23 immediately prior to use |  |  |
| **Once EC23 is added, keep media for max 3 days |  |  |

| Differentiation Media #3 |  |  |
| --- | --- | --- |
| Component |  | Dilution |
| Neurobasal | Up to 25ml |  |
| 1x B-27 | 500µl (50x stock) | 1:50 |
| 20mM KCL | 250µl (1M stock) | 1:50 |
| 1x Pen/Strep | 250µl | 1:100 |
| 2mM Glutamax | 250µl (100x stock) | 1:100 |
| 50ng/ml BDNF | 12.5µl (10µg/ml stock) | 1:200 |
| 2 mM dibutyryl cyclic AMP (db-cAMP) | 250µl (0.2M stock) | 1:500 |
| 10µM EC23 | 10µl/5µl | 1:500/<br>1:1000 |
|  | (5mM stock)/ (10mM stock) |  |
| *Add EC23 immediately prior to use |  |  |
| **Once EC23 is added, keep media for max 3 days |  |  |

### Antibodies and Reagents

The following primary antibodies were used in this study: rabbit anti-Tau (Agilent, Cat# A0024, western blot, 1/5000, RRID: AB_10013724), mouse anti-GAPDH (6C5) (Santa Cruz Biotechnology, Cat# sc-32233, western blot, 1/5000, RRID: AB_627679), rabbit anti-MCU (D2Z3B) (Cell Signaling Technology, Cat# 14997, western blot, 1/1000, RRID: AB_2721812, goat anti-Cathepsin B (R and D Systems, Cat# AF953, western blot, 1/1000, RRID: AB_355738), goat anti-Cathepsin D (R and D Systems, Cat# AF1029, western blot, 1/1000, RRID: AB_2087094), goat anti-Cathepsin L (Novus, Cat# AF1515, western blot, 1/1000, RRID: AB_2665930), mouse anti-p62/SQSTM1 (2C11) (Novus, Cat# H00008878-M01, western blot, 1/2000, RRID: AB_548364), rabbit anti-LAMP1 (ThermoFisher Scientific, Cat# PA1-654A, western blot, 1/1000, RRID: AB_2134611) and rabbit anti-LAMP2 (ThermoFisher Scientific, Cat# PA1-655, western blot, 1/1000, RRID: AB_2134625). For western blots secondary antibodies were purchased from LI-COR Biosciences and used at 1/10,000 dilution.

The following additional reagents were used in this study: Dulbecco’s Phosphate-Buffered Saline (DPBS)-no calcium, no magnesium (ThermoFisher Scientific, Cat#14190094), KCL 2M RNase free (Invitrogen, Cat# 10606365), Neurobasal (w/o phenol red (ThermoFisher Scientific, Cat# 12348-017), B27 supplement (Gibco, Cat# 17504-044), dibutyryl-cAMP (db-cAMP) (Insight Biotechnology Limited, Cat#sc-201567A), human BDNF (Cambridge Bioscience, Cat# GFH1-10), CHAPS (Scientific Laboratory Supplies, Cat# C3023-1G, Dithiothreitol (DTT) (Merck, Cat# 10197777001), EPPS (Thermo Fisher Scientific, Cat# A13714.14), Acetonitrile (ACN) (Merck, Cat# 360457), Formic acid (Thermo Fisher Scientific, Cat# 10559570), Trifluoroacetic acid (TFA) (Merck, Cat# 8082600026), Iodoacetamide (IAA) (Merck, Cat# I1149).

### BSA (Bovine Serum Albumin) and DQ (dye-quenched)-BSA experiments

Control, Tau35-HA and Avi-FL tau SH-SY5Y lines were plated on 96-well plates (Falcon, Corning, Cat# 353219) and, on day 14, differentiated cells were treated with 10μg/ml red DQ-Red BSA (Thermo Fisher Scientific, Cat# D12051) and 50 μg/ml Alexa-488 BSA (Thermo Fisher Scientific, Cat# A13100) for 4h in a humidified incubator. Following the incubation, cells were washed three times with warm culture media and counterstained with NucBlue Live ReadyProbes Reagent (Invitrogen, Cat# R37605) to visualise, count and segment nuclei. The cells were then washed with DPBS and fixed in 2% (w/v) parafolmaldehyde (PFA) for 10 min. Imaging was performed using a spinning disk confocal microscope (Opera Phenix HCS System, PerkinElmer). Images were acquired using a 40× water-immersion objective (NA 1.1) with a binning factor of two. Images were captured sequentially at each wavelength, utilising a 405 nm laser for NucBlue, a 488 nm laser for Alexa-488 BSA, and a 568 nm laser for DQ-Red BSA. To encompass the entire cell depth, 10 μm z-stacks were acquired with 0.4 μm intervals. Data analysis was conducted using PerkinElmer’s Harmony HCA Software.

### Lysotracker staining and imaging

Control, Tau35-HA and Avi-FL tau SH-SY5Y lines were plated on 8-well chamber slides (Ibidi, Cat# 80841) and, on day 14, differentiated cells were treated with 100nM Lysotracker Deep Red (Thermo Fisher Scientific, Cat# L12492) for 1h in a humidified incubator, according to manufacturer’s instructions. Following the incubation, cells were washed three times with warm culture media and counterstained with NucBlue Live Ready Probes Reagent (Invitrogen, Cat# R37605) to visualise, count and segment nuclei. Warm imaging media (Live Cell Imaging Solution, Invitrogen, Cat# 12363603) was added and cells were returned to the incubator for 10min before imaging. Live cell imaging microscopy was performed in imaging media at 37°C with humidified CO_2_ (Okolab incubator with CO_2_ control) using the Vt-iSIM super-resolution microscope with Hamamatsu Flash 4.0 sCMOS camera. Images were acquired using a 100× silicone-immersion objective (NA 1.35). Prior to image acquisition the lens was calibrated as previously described^58^, to minimize uneven background and nonspecific labelling, ensuring an optimal signal-to-background ratio (SBR). Staining with Lysotracker-647 allowed clear visualisation of lysosome morphology in live-cell SIM images, with minimal fluorescence background. To assist with nuclear detection and segmentation, cells were counterstained with NucBlue reagent. For lysotracker spot detection, intensity and motility parameter analysis, 100 frames were acquired per imaging session (10sec), at a fast acquisition mode, at a rate of 100 frames per second (fps), utilising a 640nm laser for Lysotracker Deep Red. Following video acquisition, an overview image was captured sequentially at each wavelength, utilising a 640nm laser for Lysotracker Deep Red, a 405 nm laser for NucBlue and brightfield. Acquisition was controlled and data stored using NIS-Elements v5.2 (Nikon). Data were analysed using NIS-Elements Advanced Research software (Nikon, RRID:SCR_014329), Image J^59^ and the Python programming language (Python Software Foundation, https://www.python.org/, Van Rossum, G., & Drake Jr, F. L. (1995). Python reference manual. Centrum voor Wiskunde en Informatica Amsterdam). The analysis protocol began by analysing video files acquired at the 647 nm wavelength with NIS Elements. The DAPI and brightfield channels from corresponding overview images were transferred to these video files to provide additional context. Subsequently, the far-red channel (647 nm) was selected for processing, followed by preprocessing using median (radius: 2) and rolling ball (radius: 5) filters to reduce noise. The bright spot detection (diameter: 0.6μm) algorithm (NIS-Elements AR 6.02.01) was used to detect bright spots, which were used to generate binary objects for further analysis. Regions of interest (ROIs) corresponding to individual cells were delineated based on the brightfield channel. Quantitative measurements included total object (spot) count, number of spots per cell, total area, mean intensity of objects (spots), total cell area, and motility parameters such as distance travelled, straightness, mean velocity, and mean square displacement. The exported .csv files were subsequently uploaded to Python for advanced statistical analysis and visualisation. This included the generation of raincloud plots to illustrate the distribution of quantitative data and rose plots to represent directional motility patterns of lysosomes. This comprehensive approach enabled detailed analysis of lysosomal behaviour across all cell lines.

### Quantification and Statistical Analysis

For all datasets the statistical significance was assessed as follows: for two group comparisons, two-tailed unpaired Student’s t-tests were used to estimate statistical significance between means; for three group comparisons, the ordinary one-way ANOVA test was used to estimate statistical significance between means of parametric datasets and Kruskal-Wallis test was used to estimate statistical significance between means of not normally distributed datasets. Dataset normality was tested with either the Shapiro-Wilk test (for small n numbers) or the Kolmogorov-Smirnov test (for large n numbers). Statistical analyses were performed using Prism (GraphPad, version 10) and R packages. Sample number, number of experiments and statistical information are stated in the corresponding figure legends. In figures, asterisks denote statistical significance as follows: \**P* < 0.05, \*\**P* < 0.01, \*\*\**P* < 0.001, \*\*\*\**P* < 0.0001. Error bars represent the standard error of the mean (SEM).

## Acknowledgements

We thank Dr Ian Chessell (Head of Neuroscience, AstraZeneca) and Dr Alison Darke (Associate Director, Global Programme for Postdoctoral Researchers, AstraZeneca) for their support and guidance. We also thank the Wohl Cellular Imaging Centre at King’s College London for assistance with super resolution microscopy. The authors acknowledge use of King’s Computational Research, Engineering and Technology Environment (CREATE). For the purposes of open access, the authors have applied a Creative Commons Attribution (CC BY) licence to any Accepted Author Manuscript version arising from this submission.

## Contributions

D.G., D.P.S. and D.P.H. led the project. D.G., D.P.S., D.P.H. and G.F. designed and supervised the research. D.G., M.W., S.L., A. A., and G.C performed the experiments and contributed to data acquisition. D.G., M.W., S.L., A. A., S.M. and D.P.S. contributed to data analysis and interpretation. D.G. wrote the paper. M.W., S.L., G.C., S.M., K.M., W.N., G.F. and D.P.S. reviewed the paper. All authors read the manuscript.

## Extended Figures

**Extended Data Fig. 1:**
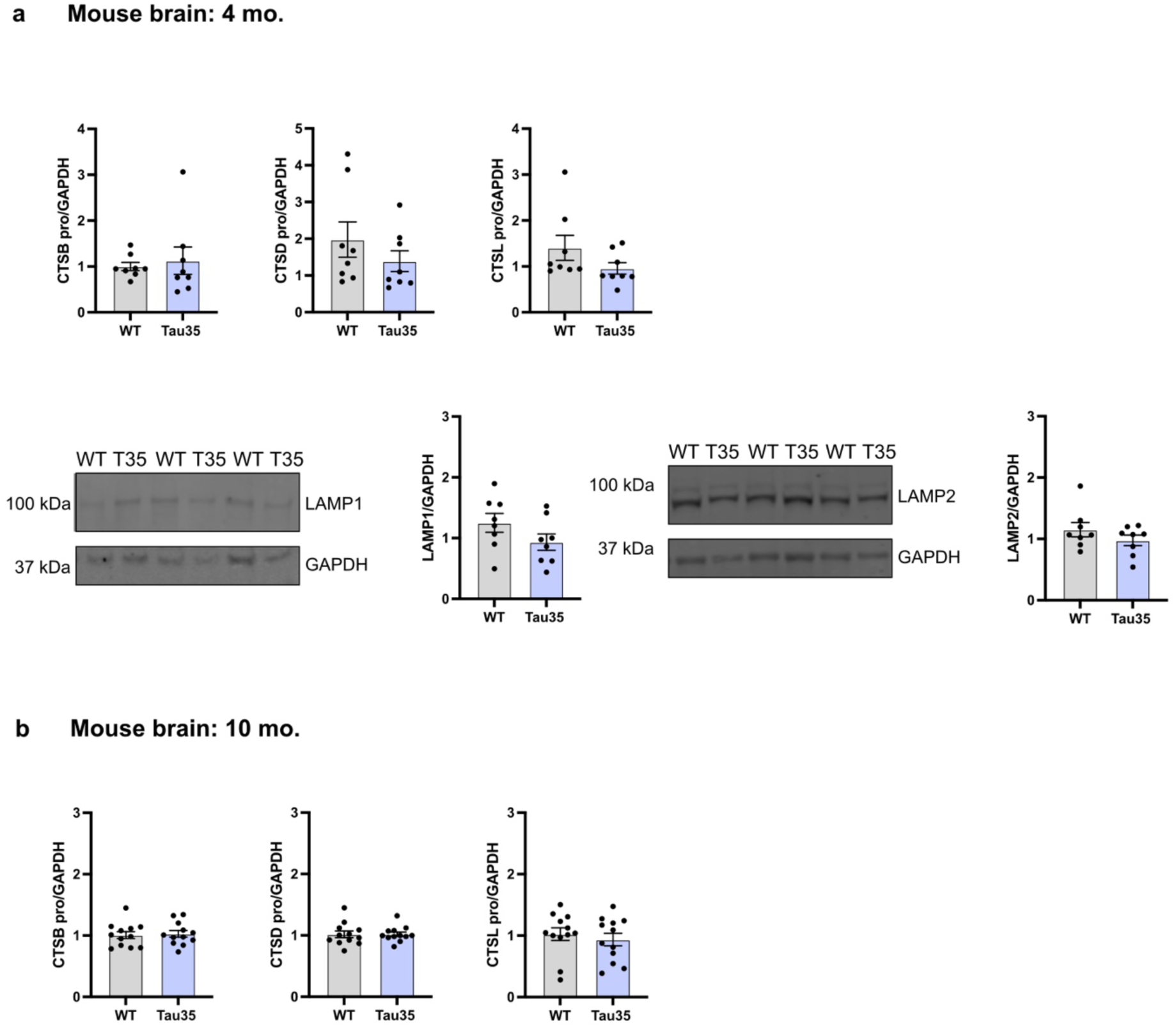
Pro-cathepsin and lysosomal marker expression across disease stages. **a,b,** Western blots of total brain homogenates from WT and Tau35 mice aged 4 (**a**) and 10 (**b**) months respectively, were probed with antibodies to CTSB, CTSD, CTSL, LAMP1, LAMP2 and GAPDH. Quantification of the blots is shown in the graphs as mean ± SEM, *n* = 8-12 brains per group. Student *t* test. CTSB, Cathepsin B; CTSD, Cathepsin D; CTSL, Cathepsin L; LAMP1, lysosomal-associated membrane protein 1; LAMP2, lysosomal-associated membrane protein 2; GAPDH, glyceraldehyde 3-phosphate dehydrogenase; SEM, standard error of the mean; WT, wild type.

**Extended Data Fig. 2:**
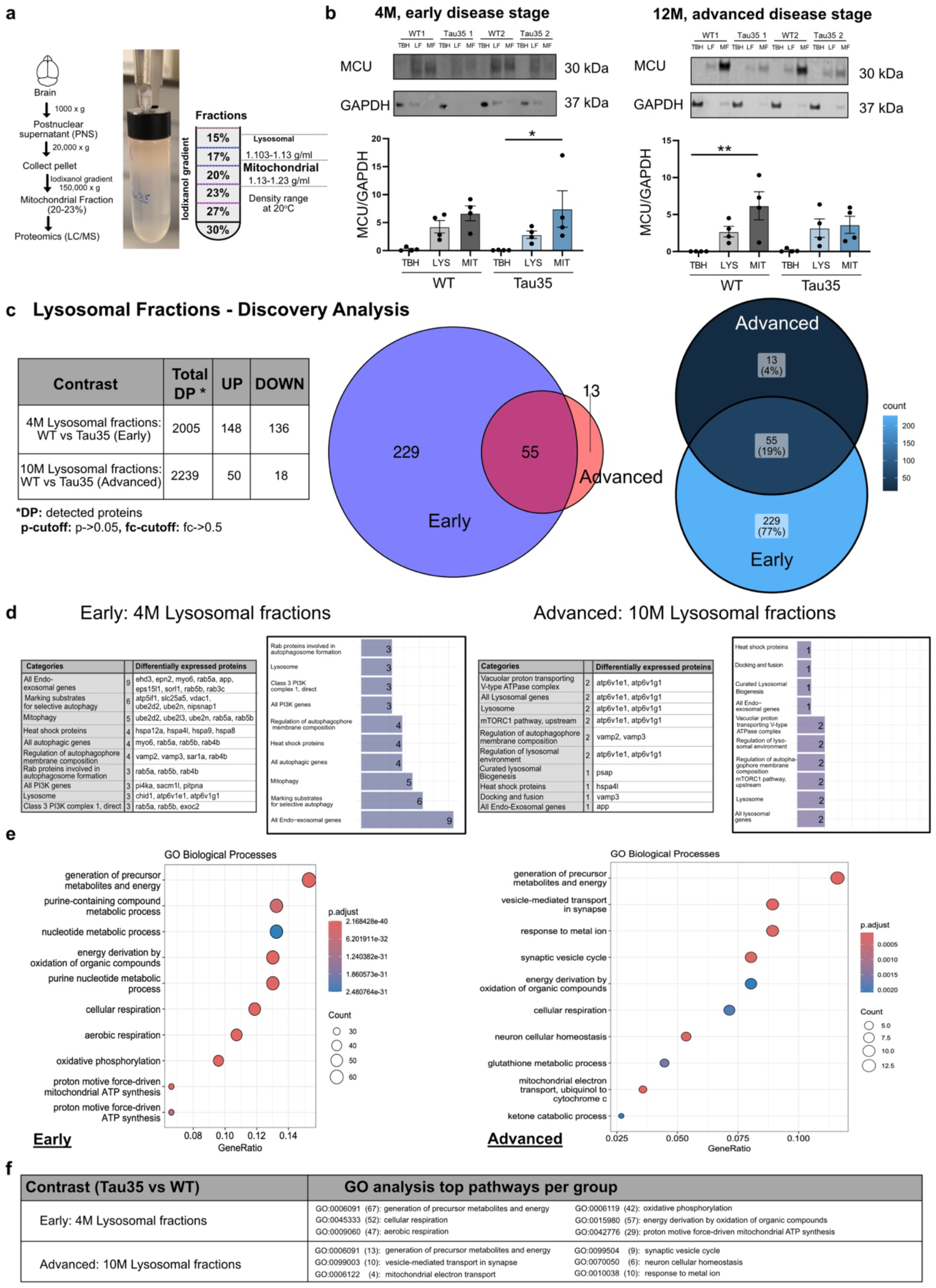
Broad discovery analysis of lysosome protein dynamics in Tau35 mouse brains. **a**, Schematic diagram of the applied workflow for subcellular fractionation of lysosomes from mouse brain. Representative images of the discontinuous iodixanol gradient showing enriched fractions from WT and Tau35 mouse brain samples. The positions of lysosomal and mitochondrial fractions on the gradient are indicated. **b,** Western blots of total brain homogenates (TBH), lysosomal fractions (LF) and mitochondrial fractions (MF) from WT and Tau35 mice aged 4 and 10 months respectively, were probed with antibodies to MCU and GAPDH. Western blotting analysis of mouse brain extracts and the different subcellular fractions from WT and Tau35 brain reveal the enrichment of the mitochondrial marker protein MCU in MF. Quantification of the blots is shown in the graphs as mean ± SEM, n = 4 brains per group. Ordinary one-way ANOVA, *P < 0.05, **P < 0.01. MCU, mitochondrial calcium uniporter; GAPDH, glyceraldehyde 3-phosphate dehydrogenase; SEM, standard error of the mean; WT, wild type. **c,** The limma package in R was used to analyze differentially expressed proteins (discovery analysis: p-cutoff: p->0.05, fc-cutoff: fc->0.5) from early and advanced sample cohorts. Table summarizing differentially expressed proteins between two cohorts, early and advanced, of wild-type (WT) and transgenic Tau35 mice. Venn diagrams illustrating protein alterations in lysosomal fractions from transgenic mice at 4 and 10 months. A total of 55 proteins are significantly altered across all datasets. **d,** Tables and graphs highlight the key autophagy-lysosomal pathway (ALP) categories and proteins identified by comparing the limma-identified differentially expressed list of proteins (discovery analysis) at early and advanced disease stages with a curated list of autophagy and endo-lysosomal pathway-associated proteins. **e,** Dot plots of the top 10 gene ontology (GO) terms in the GO enrichment analysis for the early and advanced cohorts. **f,** Table summarizing top pathways that are altered in early and advanced cohorts, including number of differentially expressed proteins that were detected per pathway.

**Extended Data Fig. 3:**
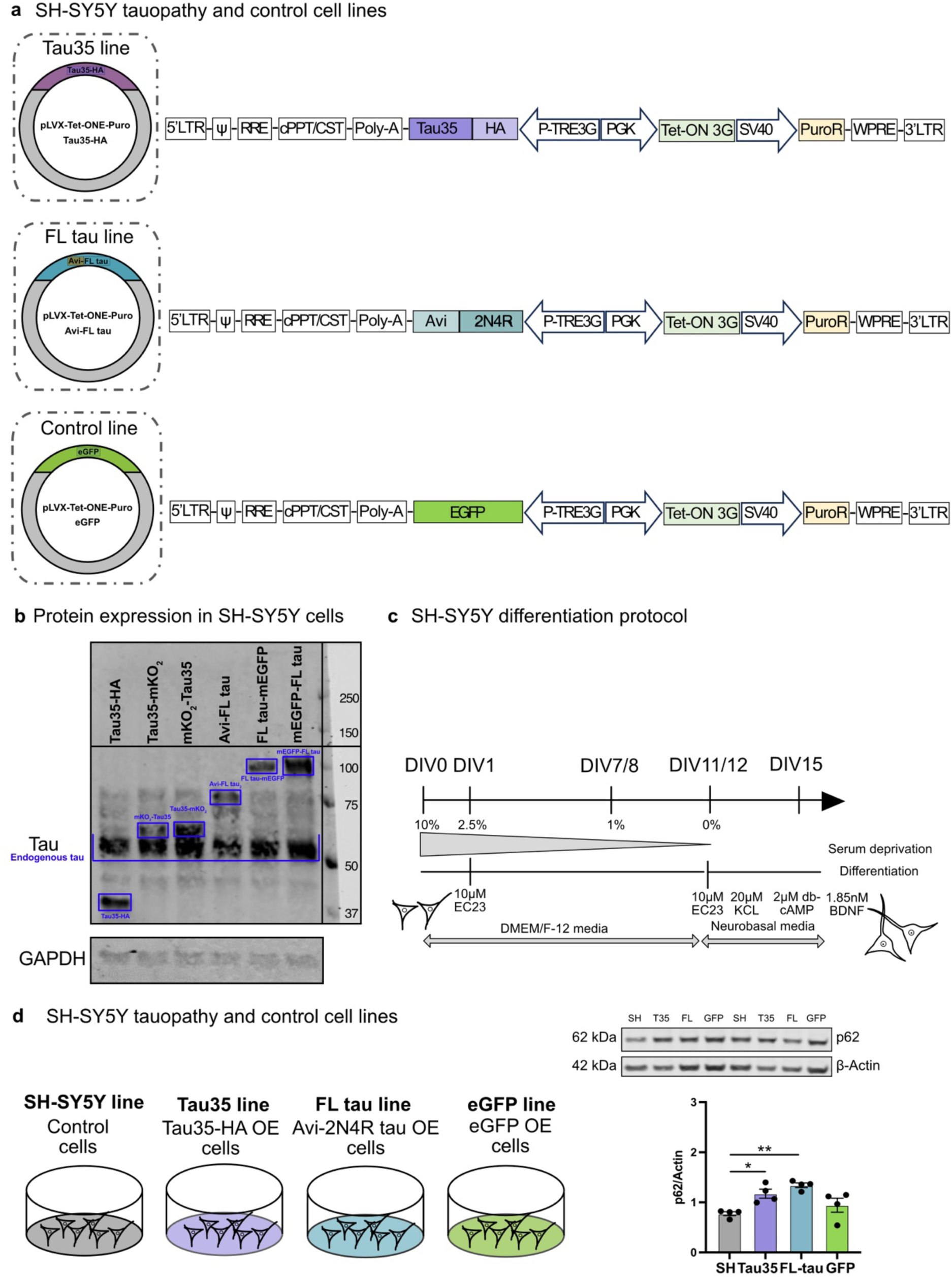
Novel SH-SYSY tau models generation, characterisation and differentiation. **a,** Generation of stable cell lines harboring an inducible pLVX-Tet-ONE vector expressing Tau35-HA (Tau35 line), Avi-FL tau (FL-tau line) and EGFP (GFP line). **b,** Western blot analysis cell lysates from all stable cell lines overexpressing tau (line maps provided in Figure S5) was conducted using antibodies specific to tau and GAPDH. The blot revealed both endogenous tau and the overexpressed fusion proteins, which are highlighted. **c,** Diagrammatic representation of the SH-SY5Y differentiation protocol, detailing the timeline of media changes and the various supplements administered. **d,** Schematic illustrating the four differentiated SH-SY5Y cell lines; control SH-SY5Y cells, cells overexpressing (OE) Tau35-HA (referred to as Tau35) and cells overexpressing (OE) Avi-FL tau (referred to as FL-tau) and EGFP (referred to as GFP). Western blot of total cell lysates from control, Tau35, and FL-tau, GFP differentiated cells, collected at 14 days in vitro (DIV), were probed with antibodies to p62 and GAPDH. Quantification of the blots is shown in the graphs as mean ± SEM, *n* = 4 independent experiments. Ordinary one-way ANOVA, \**P* < 0.05, \*\**P* < 0.01. p62/SQSTM1, Sequestosome-1; GAPDH, glyceraldehyde 3-phosphate dehydrogenase; SEM, standard error of the mean.

**Extended Data Fig. 4:**
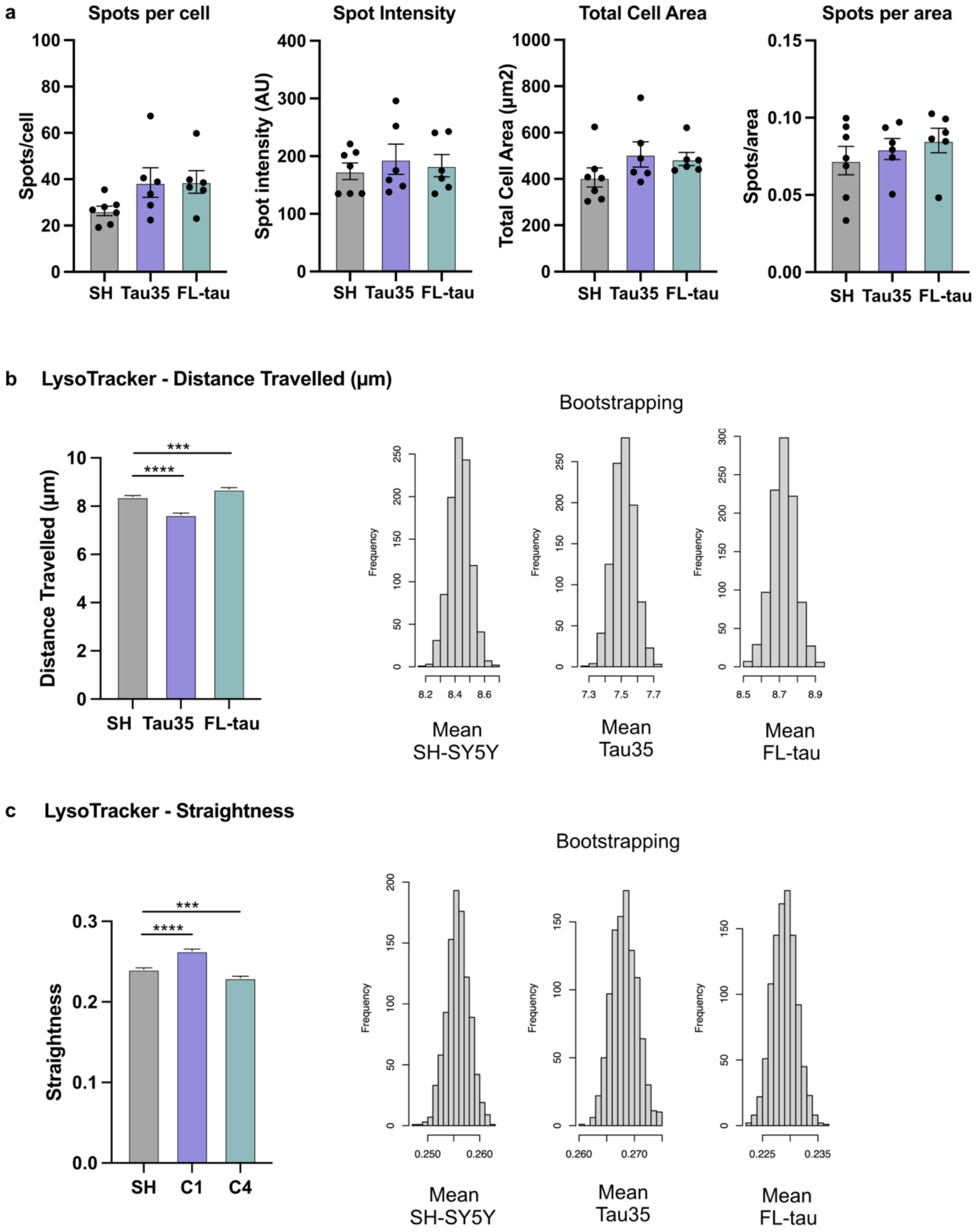
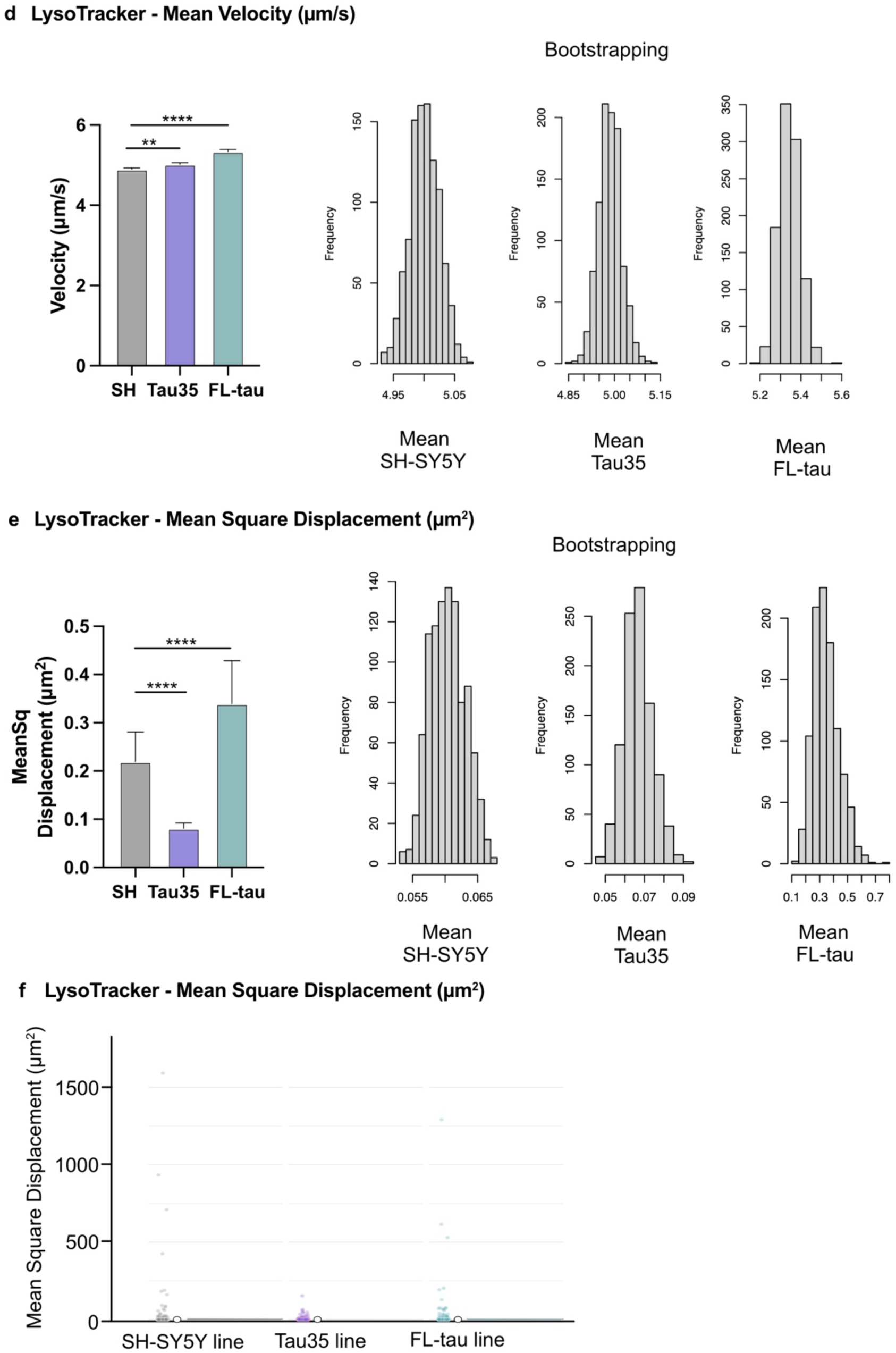
Bootstrapping analysis of lysosomal motility data for sampling distributions and robust comparisons. **a,** Quantification of basic lysosome motility parameters, following lysotracker staining. Simple box plots of the number of spots per cell, spot intensity, total cell area and number of spots per area; AU: arbitrary units; Ordinary one-way ANOVA, SEM, standard error of the mean. **b,c,d,e,** Simple box plots (left) and histograms (right) are shown to display the frequency distributions of lysosome motility parameters, Distance travelled (μm) (**b**), Straightness (**c**), Mean Velocity (μm/s) (**d**) and Mean Square Displacement (μm^2^) (**e**), for comparison between the three cell lines. The bootstrapping analysis was performed using random subsets of the data (1,000 bootstrap replicates for each parameter), which were derived from the original dataset containing 24,000–43,000 data points (spots) per line. For **b-f**: Quantification of the blots is shown in the graphs as mean ± SEM, *n* = 6-7 independent experiments; Kruskal-Wallis’s test; \*\**P* < 0.01, \*\*\**P* < 0.001, \*\*\*\**P* < 0.0001. **f,** The original Raincloud plot for Mean Square Displacement (μm²) is presented to accurately depict the full range of values in proportion across the different cell lines.

