## Supplemental Data for "Proteomic Analysis Links Truncated Tau to Lysosome Motility, Autophagy and Endo-lysosomal Dysfunction"

### Supplementary Information

#### Supplementary data 1 – Differential Expression Analysis of Lysosome-Enriched Brain Fractions – Differentially Expressed Proteins

##### Stringent Analysis (Early stage (4M) samples)

| Proteins | logFC | AveExpr | t | P.Value | adj.P.Val | B | Category |
|---|---|---|---|---|---|---|---|
| **Hprt1** | 1.482654748 | 23.9755517 | 10.72384963 | 1.65E-05 | 0.02445837 | 3.409095466 | upregulated proteins |
| **Atp6v1g1** | 1.801287028 | 24.167866 | 8.184564052 | 9.22E-05 | 0.05202668 | 1.988688014 | upregulated proteins |
| **Tusc3** | 1.065376825 | 24.5783636 | 6.915070538 | 0.00025991 | 0.05408401 | 1.05103871 | upregulated proteins |
| **Ank3** | 1.395740215 | 24.0841377 | 6.491685002 | 0.00037985 | 0.06210586 | 0.695640945 | upregulated proteins |
| **Dnajc6** | 1.175166328 | 23.1239242 | 5.611381578 | 0.00089067 | 0.07550888 | -0.1219185 | upregulated proteins |
| **Pdcd6** | 1.506357912 | 24.0803382 | 5.101027818 | 0.00152477 | 0.08725505 | -0.64901814 | upregulated proteins |
| **Rab2a** | 1.17285903 | 24.7912644 | 4.569387824 | 0.00277215 | 0.10057711 | -1.24277796 | upregulated proteins |
| **Vamp3** | 1.376682813 | 23.6340417 | 4.465101891 | 0.0031319 | 0.10057711 | -1.36475055 | upregulated proteins |
| **Anxa5** | 1.086172925 | 22.2627323 | 4.341223795 | 0.00362794 | 0.10265279 | -1.5120012 | upregulated proteins |
| **Slc38a3** | 1.09626222 | 23.8244998 | 4.293951184 | 0.00383961 | 0.10325773 | -1.56886897 | upregulated proteins |
| **Napa** | 1.048673575 | 20.730417 | 4.115476387 | 0.00477065 | 0.10416692 | -1.78692344 | upregulated proteins |
| **Cyc1** | 1.017870803 | 24.1474367 | 4.115197663 | 0.00477229 | 0.10416692 | -1.78726811 | upregulated proteins |
| **Gpi** | 1.315873443 | 22.6933896 | 4.069501199 | 0.00504899 | 0.10416692 | -1.8439487 | upregulated proteins |
| **Pgam1** | 1.081659018 | 22.7614465 | 4.007430901 | 0.00545346 | 0.10528471 | -1.92148821 | upregulated proteins |
| **Atp5pf** | 1.148758425 | 22.3920259 | 3.710910354 | 0.00794413 | 0.11085336 | -2.30047725 | upregulated proteins |
| **Prps1** | 1.229831708 | 23.53826 | 3.665947814 | 0.00842019 | 0.11158678 | -2.35915098 | upregulated proteins |
| **Pura** | 1.1258595 | 19.6750244 | 3.584817541 | 0.00935969 | 0.11501283 | -2.46579682 | upregulated proteins |
| **Itgav** | 1.269248428 | 22.9278385 | 3.510063476 | 0.0103268 | 0.11915364 | -2.56492443 | upregulated proteins |
| **Ndufb9** | 1.48077869 | 21.5462391 | 3.489333488 | 0.01061381 | 0.11915364 | -2.59255682 | upregulated proteins |
| **Cend1** | 1.658282058 | 23.8296867 | 3.478330228 | 0.01076965 | 0.11915364 | -2.60724871 | upregulated proteins |
| **Them4** | 1.76207438 | 22.7022354 | 3.458119284 | 0.01106241 | 0.11981767 | -2.63427967 | upregulated proteins |
| **Lrp1** | 1.117698145 | 20.9665289 | 3.304917886 | 0.01358236 | 0.13517402 | -2.84100162 | upregulated proteins |
| **Itgb1** | 1.479496723 | 21.6000895 | 3.244465434 | 0.0147415 | 0.14062518 | -2.92342082 | upregulated proteins |
| **Myo6** | 1.54710877 | 23.2549639 | 3.134726815 | 0.0171263 | 0.14519301 | -3.07417214 | upregulated proteins |

| | | | | | | | |
|---|---|---|---|---|---|---|---|
| **Camkv** | 1.129415442 | 23.8445987 | 3.082459811 | 0.01840463 | 0.14647242 | -3.14645629 | upregulated proteins |
| **Dcps** | 1.18926942 | 20.9786022 | 3.076031475 | 0.01856876 | 0.14707228 | -3.15536691 | upregulated proteins |
| **Hycc2** | 2.286445488 | 21.7644682 | 3.051686175 | 0.01920466 | 0.15012594 | -3.18915227 | upregulated proteins |
| **Ppia** | 1.10397588 | 22.2358438 | 2.942244827 | 0.02236399 | 0.16199736 | -3.34175318 | upregulated proteins |
| **Atp5f1c** | 1.008060025 | 20.0695853 | 2.637065353 | 0.0344478 | 0.2021821 | -3.77216378 | upregulated proteins |
| **Myl12b** | 1.285262173 | 23.275955 | 2.62894994 | 0.03485036 | 0.20298338 | -3.78367687 | upregulated proteins |
| **Fkbp1a** | 1.08347151 | 22.9234956 | 2.567109974 | 0.03808373 | 0.2112128 | -3.87146876 | upregulated proteins |
| **Grm1** | 1.698594888 | 20.3198975 | 2.510722323 | 0.04130434 | 0.21408511 | -3.95158337 | upregulated proteins |
| **Hnrnpm** | 14.25046414 | 17.6634911 | 2.487762563 | 0.04269549 | 0.21653082 | -3.98421146 | upregulated proteins |
| **Folh1** | 2.18272715 | 24.2569966 | 2.393134724 | 0.04896005 | 0.23233046 | -4.11865065 | upregulated proteins |
| **Ndufaf2** | -1.00697562 | 24.2984689 | -7.27789635 | 0.00019043 | 0.05202668 | 1.337745875 | downregulated proteins |
| **Psap** | -1.18205985 | 24.9800101 | -6.50778668 | 0.00037428 | 0.06210586 | 0.709585216 | downregulated proteins |
| **Pdp1** | -1.07250894 | 24.2240072 | -4.70291004 | 0.0023767 | 0.09933821 | -1.08926076 | downregulated proteins |
| **Uchl1** | -1.04154539 | 24.4911962 | -4.6106582 | 0.00264263 | 0.09933821 | -1.19501 | downregulated proteins |
| **Anxa6** | -1.14089919 | 21.4818466 | -4.20277652 | 0.00428742 | 0.10416692 | -1.67960207 | downregulated proteins |
| **Vdac1** | -1.20541044 | 21.5370307 | -3.7136009 | 0.00791659 | 0.11085336 | -2.29697607 | downregulated proteins |
| **Gdi2** | -1.00356953 | 23.4059239 | -3.54314677 | 0.00988603 | 0.11702908 | -2.52095374 | downregulated proteins |
| **Nipsnap1** | -1.22135801 | 24.1899699 | -3.47558858 | 0.01080887 | 0.11915364 | -2.61091213 | downregulated proteins |
| **Fundc2** | -1.16041125 | 26.2147083 | -3.16241455 | 0.01648793 | 0.14424435 | -3.03600372 | downregulated proteins |
| **Mtfp1** | -1.03752944 | 21.7447425 | -3.16110109 | 0.01651763 | 0.14424435 | -3.03781238 | downregulated proteins |
| **Slc9a6** | -1.20193702 | 25.8919655 | -3.10863101 | 0.01775224 | 0.14623364 | -3.11022478 | downregulated proteins |
| **Itm2b** | -13.552999 | 15.7889577 | -2.76927239 | 0.02853335 | 0.18294912 | -3.5849988 | downregulated proteins |
| **Edil3** | -1.03550717 | 22.500499 | -2.53007678 | 0.04016817 | 0.21262684 | -3.92408083 | downregulated proteins |
| **Sorl1** | -1.91116318 | 24.8287551 | -2.50546268 | 0.04161883 | 0.21408511 | -3.95905771 | downregulated proteins |
| **Rps14** | -1.45735263 | 21.9830646 | -2.46747618 | 0.04396499 | 0.22119273 | -4.01304034 | downregulated proteins |

#### Stringent Analysis (Advanced stage (10M) samples)

| Proteins | logFC | AveExpr | t | P.Value | adj.P.Val | B | Category |
|---|---|---|---|---|---|---|---|
| **Atp6v1g1** | 1.609594225 | 22.36168014 | 5.794993101 | 0.0007389 | 0.43340542 | -1.57910923 | upregulated proteins |
| **App** | 2.0042263 | 22.00218631 | 3.585990398 | 0.00933867 | 0.84328435 | -2.70522762 | upregulated proteins |
| **Got1** | 1.23645398 | 19.44205638 | 3.456363736 | 0.01108087 | 0.84328435 | -2.79695289 | upregulated proteins |
| **Pdcd6** | 15.11918016 | 12.78877465 | 3.07661944 | 0.01854386 | 0.84328435 | -3.08389703 | upregulated proteins |
| **Kcna1** | 1.18960055 | 22.59141177 | 2.950011374 | 0.02211175 | 0.84328435 | -3.18544942 | upregulated proteins |
| **Krt73** | 1.071967377 | 22.44166053 | 2.742167319 | 0.02964024 | 0.84328435 | -3.35811789 | upregulated proteins |
| **Vamp3** | 1.045562138 | 24.5937094 | 2.7088304 | 0.03108017 | 0.84328435 | -3.38646164 | upregulated proteins |
| **Pde1b** | 1.323900823 | 21.37399286 | 2.688854983 | 0.031978 | 0.84328435 | -3.40352645 | upregulated proteins |
| **Atp6v1e1** | 1.257094013 | 25.44076545 | 2.634038421 | 0.03458415 | 0.84328435 | -3.45066008 | upregulated proteins |
| **Anxa6** | 13.08346744 | 15.85681677 | 2.589679508 | 0.03685527 | 0.84328435 | -3.48911743 | upregulated proteins |
| **Pfn2** | 1.383959255 | 22.21154203 | 2.567623666 | 0.03804186 | 0.84328435 | -3.50834008 | upregulated proteins |
| **Aldh6a1** | 1.003151 | 23.39061424 | 2.435549304 | 0.04602768 | 0.84328435 | -3.62475432 | upregulated proteins |
| **Hprt1** | -1.35913237 | 24.17226624 | -8.9238509 | 5.34E-05 | 0.12313318 | -0.85610949 | downregulated proteins |
| **Kcnj10** | -1.11930644 | 21.53325515 | -5.25872301 | 0.00128496 | 0.5928822 | -1.78658915 | downregulated proteins |
| **Ist1** | -16.9752603 | 13.99056497 | -3.28373277 | 0.01396858 | 0.84328435 | -2.9240517 | downregulated proteins |
| **Txnrd1** | -1.41301874 | 21.73151016 | -3.14622628 | 0.01684868 | 0.84328435 | -3.02929323 | downregulated proteins |
| **Hcn2** | -1.08254029 | 24.37010389 | -2.93652001 | 0.02253293 | 0.84328435 | -3.19643718 | downregulated proteins |
| **Atp5if1** | -1.21104421 | 20.64253287 | -2.90217285 | 0.0236441 | 0.84328435 | -3.22455201 | downregulated proteins |

| | | | | | | | |
|---|---|---|---|---|---|---|---|
| **Acyp1** | -1.72285251 | 23.2514449 | -2.51585334 | 0.04098574 | 0.84328435 | -3.55371334 | downregulated proteins |
| **Nbea** | -1.545281 | 23.19386344 | -2.47320307 | 0.04358818 | 0.84328435 | -3.59134814 | downregulated proteins |
| **Calb1** | -1.12256586 | 21.03986195 | -2.45340048 | 0.04485404 | 0.84328435 | -3.60889642 | downregulated proteins |

#### Discovery Analysis (Early stage (4M) samples)

| Proteins | logFC | AveExpr | t | P.Value | adj.P.Val | B | Category |
|---|---|---|---|---|---|---|---|
| **Hprt1** | 1.48265475 | 23.9755517 | 10.7238496 | 1.65097822452125E-05 | 0.02445837 | 3.40909547 | upregulated proteins |
| **Glo1** | 0.88921776 | 24.5264764 | 10.3034024 | 2.13703548473205E-05 | 0.02445837 | 3.20826276 | upregulated proteins |
| **Atp6v1g1** | 1.80128703 | 24.167866 | 8.18456405 | 9.21583025353879E-05 | 0.05202668 | 1.98868801 | upregulated proteins |
| **S100a1** | 0.81660577 | 26.8149709 | 8.01810459 | 0.000104754 | 0.05202668 | 1.87575835 | upregulated proteins |
| **Kcnj10** | 0.57032684 | 32.395901 | 7.34756403 | 0.000179646 | 0.05202668 | 1.3910004 | upregulated proteins |
| **Ndufaf2** | -1.0069756 | 24.2984689 | -7.2778964 | 0.000190431 | 0.05202668 | 1.33774588 | downregulated proteins |
| **Rida** | 0.76643256 | 25.68442 | 7.23929894 | 0.000196721 | 0.05202668 | 1.30799701 | upregulated proteins |
| **Ppp1ca** | 0.58788606 | 33.5578061 | 7.20083179 | 0.000203224 | 0.05202668 | 1.2781735 | upregulated proteins |
| **Fabp3** | 0.75358984 | 28.0613895 | 7.19309444 | 0.000204561 | 0.05202668 | 1.27215354 | upregulated proteins |
| **Lrch2** | 0.61208152 | 24.6448789 | 6.93293718 | 0.000255879 | 0.05408401 | 1.06553337 | upregulated proteins |
| **Tusc3** | 1.06537683 | 24.5783636 | 6.91507054 | 0.000259906 | 0.05408401 | 1.05103871 | upregulated proteins |
| **Gsta4** | -0.6867246 | 19.9832786 | -6.5846584 | 0.000348927 | 0.06210586 | 0.77568209 | downregulated proteins |
| **Psap** | -1.1820599 | 24.9800101 | -6.5077867 | 0.000374282 | 0.06210586 | 0.70958522 | downregulated proteins |
| **Ank3** | 1.39574022 | 24.0841377 | 6.491685 | 0.000379852 | 0.06210586 | 0.69564094 | upregulated proteins |
| **Arf6** | 0.67016129 | 26.3132387 | 6.37488627 | 0.000423171 | 0.06342825 | 0.59344817 | upregulated proteins |
| **Sacm1l** | 0.74457391 | 21.3244015 | 6.27735452 | 0.000463635 | 0.06342825 | 0.50669043 | upregulated proteins |
| **Aco1** | 0.6340852 | 21.851129 | 6.26047697 | 0.00047107 | 0.06342825 | 0.49154427 | upregulated proteins |
| **App** | 0.61887613 | 25.1380994 | 6.13420733 | 0.000531154 | 0.06754514 | 0.37696742 | upregulated proteins |
| **Dclk1** | -0.8340945 | 25.1093881 | -6.0055858 | 0.000601368 | 0.07244906 | 0.25793787 | downregulated proteins |
| **Chp1** | 0.54454911 | 28.6907657 | 5.77302955 | 0.000756447 | 0.07550888 | 0.0366386 | upregulated proteins |
| **Gsr** | -0.5753545 | 31.0372655 | -5.7598019 | 0.000766533 | 0.07550888 | 0.0238114 | downregulated proteins |
| **Got1** | -0.8774215 | 27.573672 | -5.7436705 | 0.000779037 | 0.07550888 | 0.00813298 | downregulated proteins |
| **Cds1** | 0.53500724 | 22.7326039 | 5.70218735 | 0.000812258 | 0.07550888 | -0.0323644 | upregulated proteins |
| **Ppp2r5c** | 0.71957055 | 26.1589525 | 5.63901063 | 0.000865947 | 0.07550888 | -0.094538 | upregulated proteins |
| **Rab5a** | -0.6320557 | 23.4053561 | -5.6179465 | 0.000884724 | 0.07550888 | -0.1154022 | downregulated proteins |
| **Dnajc6** | 1.17516633 | 23.1239242 | 5.61138158 | 0.000890668 | 0.07550888 | -0.1219185 | upregulated proteins |
| **Tpt1** | -0.5410861 | 28.1665027 | -5.5046935 | 0.00099383 | 0.07769799 | -0.2287422 | downregulated proteins |

| | | | | | | | |
|---|---|---|---|---|---|---|---|
| **Timm10** | 0.94194164 | 22.7471944 | 5.48561804 | 0.001013651 | 0.07769799 | -0.2480265 | upregulated proteins |
| **Cbarp** | 0.87567699 | 26.2610788 | 5.48118338 | 0.001018322 | 0.07769799 | -0.2525177 | upregulated proteins |
| **Ist1** | -0.6975562 | 29.6554428 | -5.382166 | 0.001129157 | 0.08337551 | -0.3535956 | downregulated proteins |
| **Uqcrh** | 0.5037877 | 25.412371 | 5.32711556 | 0.001196574 | 0.08444618 | -0.4104553 | upregulated proteins |
| **Cct2** | 0.71745459 | 26.1063589 | 5.31077598 | 0.001217442 | 0.08444618 | -0.4274238 | upregulated proteins |
| **Rab18** | -0.6651339 | 26.0305222 | -5.1683887 | 0.001417524 | 0.08725505 | -0.5770853 | downregulated proteins |
| **Calb2** | -0.7080722 | 27.191278 | -5.1390527 | 0.001463161 | 0.08725505 | -0.6083227 | downregulated proteins |
| **Vamp2** | -0.6873281 | 27.5600146 | -5.1311363 | 0.001475755 | 0.08725505 | -0.6167758 | downregulated proteins |
| **Pdcd6** | 1.50635791 | 24.0803382 | 5.10102782 | 0.001524772 | 0.08725505 | -0.6490181 | upregulated proteins |
| **Ncald** | 0.77014992 | 25.0371729 | 4.99797915 | 0.001706791 | 0.09085686 | -0.7604805 | upregulated proteins |
| **Sez6l2** | 0.5880655 | 22.1403674 | 4.97292817 | 0.001754617 | 0.09127995 | -0.7878378 | upregulated proteins |
| **Lgi3** | -0.8686743 | 29.0507037 | -4.9292856 | 0.001841542 | 0.09264566 | -0.8357432 | downregulated proteins |
| **Uso1** | 0.63090223 | 25.393019 | 4.8582819 | 0.001993396 | 0.09264566 | -0.9143499 | upregulated proteins |
| **Akr1b1** | -0.8370796 | 28.9094793 | -4.8480056 | 0.002016506 | 0.09264566 | -0.9257953 | downregulated proteins |
| **Plec** | 0.91209025 | 24.4801612 | 4.84772142 | 0.002017149 | 0.09264566 | -0.926112 | upregulated proteins |
| **Gstm1** | -0.5869978 | 24.2442124 | -4.8370675 | 0.002041432 | 0.09264566 | -0.9379969 | downregulated proteins |
| **Kcna1** | -0.6697445 | 27.2864501 | -4.82721 | 0.002064189 | 0.09264566 | -0.94901 | downregulated proteins |
| **Hcn2** | -0.8912277 | 27.5315521 | -4.7218579 | 0.00232584 | 0.09933821 | -1.0677159 | downregulated proteins |
| **Pdp1** | -1.0725089 | 24.2240072 | -4.70291 | 0.002376702 | 0.09933821 | -1.0892608 | downregulated proteins |
| **Lgi2** | -0.5710203 | 22.216167 | -4.69799 | 0.00239011 | 0.09933821 | -1.0948649 | downregulated proteins |
| **Atp5if1** | -0.6182418 | 22.3684778 | -4.6683257 | 0.002472751 | 0.09933821 | -1.1287391 | downregulated proteins |
| **Arpc4** | 0.82177925 | 24.1548782 | 4.65840636 | 0.002501089 | 0.09933821 | -1.1400989 | upregulated proteins |
| **Ndrg4** | -0.7023022 | 28.7830597 | -4.6301474 | 0.002583814 | 0.09933821 | -1.1725512 | downregulated proteins |
| **Smpd3** | -0.6360064 | 31.0260403 | -4.6284417 | 0.002588904 | 0.09933821 | -1.1745143 | downregulated proteins |
| **Uchl1** | -1.0415454 | 24.4911962 | -4.6106582 | 0.002642633 | 0.09933821 | -1.19501 | downregulated proteins |
| **Ube2l3** | -0.9670504 | 21.9990242 | -4.6091382 | 0.002647283 | 0.09933821 | -1.1967643 | downregulated proteins |
| **Rab2a** | 1.17285903 | 24.7912644 | 4.56938782 | 0.002772152 | 0.10057711 | -1.242778 | upregulated proteins |
| **Krt73** | 0.76924148 | 24.2245667 | 4.54815929 | 0.002841502 | 0.10057711 | -1.2674595 | upregulated proteins |
| **Tkt** | -0.5966042 | 29.2760678 | -4.5351016 | 0.002885112 | 0.10057711 | -1.2826785 | downregulated proteins |
| **Psma6** | -0.6724336 | 27.9566507 | -4.4862483 | 0.003054971 | 0.10057711 | -1.3398708 | downregulated proteins |
| **Vamp3** | 1.37668281 | 23.6340417 | 4.46510189 | 0.003131903 | 0.10057711 | -1.3647505 | upregulated proteins |
| **DSCC1** | 0.58750931 | 25.421489 | 4.4532578 | 0.003175928 | 0.10057711 | -1.3787184 | upregulated proteins |
| **Pfn1** | 0.55850415 | 19.6636186 | 4.43432297 | 0.003247737 | 0.10057711 | -1.4010971 | upregulated proteins |
| **Vat1** | 0.79243311 | 24.7966748 | 4.41318603 | 0.003330026 | 0.10057711 | -1.4261492 | upregulated proteins |
| **Ldhb** | -0.6517663 | 24.1816152 | -4.4025669 | 0.003372236 | 0.10057711 | -1.4387635 | downregulated proteins |
| **Pde1b** | 0.67824973 | 24.6630713 | 4.39352121 | 0.003408658 | 0.10057711 | -1.4495237 | upregulated proteins |
| **Nrxn2** | -0.7602648 | 24.5542851 | -4.3930344 | 0.00341063 | 0.10057711 | -1.4501032 | downregulated proteins |
| **Ca8** | 0.72145164 | 21.5420139 | 4.39072499 | 0.003420004 | 0.10057711 | -1.4528526 | upregulated proteins |
| **Arpc2** | 0.78633532 | 22.3066835 | 4.38894051 | 0.003427267 | 0.10057711 | -1.4549778 | upregulated proteins |

| | | | | | | | |
|---|---|---|---|---|---|---|---|
| **Ppp2r1a** | -0.7669361 | 28.8571869 | -4.3592343 | 0.003550711 | 0.10265279 | -1.4904332 | downregulated proteins |
| **Anxa5** | 1.08617293 | 22.2627323 | 4.34122379 | 0.00362794 | 0.10265279 | -1.5120012 | upregulated proteins |
| **Atp5me** | 0.74422764 | 20.9637672 | 4.3401653 | 0.003632536 | 0.10265279 | -1.5132705 | upregulated proteins |
| **Slc39a12** | 0.6039702 | 29.4809369 | 4.31288058 | 0.003753259 | 0.10325773 | -1.5460526 | upregulated proteins |
| **Slc38a3** | 1.09626222 | 23.8244998 | 4.29395118 | 0.003839613 | 0.10325773 | -1.568869 | upregulated proteins |
| **Kcnab2** | -0.6836487 | 23.4307202 | -4.2879963 | 0.003867231 | 0.10325773 | -1.576059 | downregulated proteins |
| **Marcksl1** | -0.9715984 | 22.6324844 | -4.2853668 | 0.003879495 | 0.10325773 | -1.5792358 | downregulated proteins |
| **Atp6v1e1** | 0.6790977 | 26.0085288 | 4.26967857 | 0.003953566 | 0.10401969 | -1.5982132 | upregulated proteins |
| **Ddt** | -0.9920749 | 20.0750888 | -4.2264689 | 0.004165766 | 0.10416692 | -1.6506943 | downregulated proteins |
| **Hpcal4** | -0.6108881 | 25.4525551 | -4.2178842 | 0.004209402 | 0.10416692 | -1.6611579 | downregulated proteins |
| **Pex5l** | -0.9850309 | 22.0339253 | -4.2053528 | 0.004274007 | 0.10416692 | -1.6764541 | downregulated proteins |
| **Hspa4l** | -0.6134817 | 26.4501494 | -4.2041687 | 0.004280168 | 0.10416692 | -1.6779009 | downregulated proteins |
| **Anxa6** | -1.1408992 | 21.4818466 | -4.2027765 | 0.004287423 | 0.10416692 | -1.6796021 | downregulated proteins |
| **Pfn2** | 0.50220858 | 26.9547177 | 4.16636098 | 0.004482114 | 0.10416692 | -1.7242156 | upregulated proteins |
| **Mpp3** | -0.8555979 | 25.5038185 | -4.1521823 | 0.004560533 | 0.10416692 | -1.7416457 | downregulated proteins |
| **Nectin1** | -0.6644981 | 29.2179201 | -4.1379985 | 0.004640495 | 0.10416692 | -1.7591154 | downregulated proteins |
| **Acyp1** | -0.5927846 | 26.166695 | -4.1361893 | 0.004650805 | 0.10416692 | -1.7613461 | downregulated proteins |
| **Praf2** | 0.53764647 | 28.251948 | 4.1332966 | 0.004667342 | 0.10416692 | -1.7649139 | upregulated proteins |
| **Napa** | 1.04867358 | 20.730417 | 4.11547639 | 0.004770652 | 0.10416692 | -1.7869234 | upregulated proteins |
| **Cyc1** | 1.0178708 | 24.1474367 | 4.11519766 | 0.004772287 | 0.10416692 | -1.7872681 | upregulated proteins |
| **Zfyve28** | 0.74000975 | 25.8301674 | 4.09037906 | 0.00492044 | 0.10416692 | -1.8180097 | upregulated proteins |
| **Nbea** | -0.5915865 | 29.3197876 | -4.089531 | 0.004925591 | 0.10416692 | -1.819062 | downregulated proteins |
| **Gpi** | 1.31587344 | 22.6933896 | 4.0695012 | 0.005048988 | 0.10416692 | -1.8439487 | upregulated proteins |
| **Gnai2** | -0.8903206 | 27.7041308 | -4.0669587 | 0.005064893 | 0.10416692 | -1.8471124 | downregulated proteins |
| **Ndufab1** | 0.55996893 | 26.0546015 | 4.05923495 | 0.005113548 | 0.10416692 | -1.8567299 | upregulated proteins |
| **Calb1** | -0.8471172 | 21.2520204 | -4.0477105 | 0.005187104 | 0.10416692 | -1.8710982 | downregulated proteins |
| **Trpv2** | -0.6214032 | 27.8360097 | -4.039793 | 0.005238311 | 0.10416692 | -1.880982 | downregulated proteins |
| **Pgam1** | 1.08165902 | 22.7614465 | 4.0074309 | 0.005453463 | 0.10528471 | -1.9214882 | upregulated proteins |
| **Lmbrd2** | -0.6127329 | 27.4905644 | -3.9934858 | 0.005549147 | 0.10528471 | -1.9389953 | downregulated proteins |
| **Slc12a2** | 0.89747707 | 21.6402576 | 3.98979226 | 0.005574799 | 0.10528471 | -1.9436377 | upregulated proteins |
| **Oxct1** | -0.7798032 | 27.7823803 | -3.9771965 | 0.005663257 | 0.10528471 | -1.9594856 | downregulated proteins |
| **Apmap** | -0.6167554 | 29.3098344 | -3.9737739 | 0.005687558 | 0.10528471 | -1.9637964 | downregulated proteins |
| **Ugp2** | -0.6948793 | 27.2145094 | -3.9628211 | 0.005766099 | 0.10528471 | -1.9776043 | downregulated proteins |
| **Capza2** | 0.66116126 | 24.7420976 | 3.95079636 | 0.005853697 | 0.10528471 | -1.9927859 | upregulated proteins |
| **Ephb3** | 0.84795328 | 21.367996 | 3.91493612 | 0.006123712 | 0.10650789 | -2.0381994 | upregulated proteins |
| **Ndufs4** | 0.8322696 | 23.5237935 | 3.91229232 | 0.006144153 | 0.10650789 | -2.0415557 | upregulated proteins |
| **Uqcrfs1** | -0.8270405 | 24.2077068 | -3.9058951 | 0.006193922 | 0.10650789 | -2.0496816 | downregulated proteins |
| **Idh1** | -0.7314133 | 29.9971087 | -3.8953807 | 0.006276685 | 0.10650789 | -2.0630518 | downregulated proteins |
| **Lactb** | 0.51628396 | 27.9357334 | 3.86971296 | 0.006483854 | 0.10895989 | -2.0957652 | upregulated proteins |

| | | | | | | | |
|---|---|---|---|---|---|---|---|
| **Rab5b** | -0.6670558 | 28.2768898 | -3.8492285 | 0.006654559 | 0.10895989 | -2.1219481 | downregulated proteins |
| **Nrcam** | -0.6324692 | 28.6558368 | -3.8491203 | 0.006655473 | 0.10895989 | -2.1220865 | downregulated proteins |
| **Hmox2** | 0.59480531 | 25.4293949 | 3.84808717 | 0.006664213 | 0.10895989 | -2.1234089 | upregulated proteins |
| **Adsl** | -0.542735 | 29.7295738 | -3.8394761 | 0.006737551 | 0.10904645 | -2.1344368 | downregulated proteins |
| **Ckb** | 0.65780871 | 24.4612015 | 3.82998718 | 0.006819389 | 0.10904645 | -2.1466028 | upregulated proteins |
| **Osbpl1a** | 0.67245746 | 25.6545822 | 3.81595151 | 0.00694244 | 0.10904645 | -2.1646244 | upregulated proteins |
| **Lrpap1** | -0.8139047 | 25.5137265 | -3.8156604 | 0.006945018 | 0.10904645 | -2.1649985 | downregulated proteins |
| **Mdh1** | -0.5423317 | 29.8590478 | -3.8091452 | 0.007002983 | 0.10904645 | -2.1733747 | downregulated proteins |
| **Rad23b** | 0.61479195 | 24.7224829 | 3.79502926 | 0.007130394 | 0.11028021 | -2.191546 | upregulated proteins |
| **Svip** | 0.52522811 | 30.2649903 | 3.77144744 | 0.007348928 | 0.11085336 | -2.2219723 | upregulated proteins |
| **Ehd3** | -0.5550309 | 29.4685505 | -3.7597014 | 0.007460499 | 0.11085336 | -2.2371601 | downregulated proteins |
| **Gdi1** | -0.5326956 | 24.4144877 | -3.7540465 | 0.007514871 | 0.11085336 | -2.2444797 | downregulated proteins |
| **Nnt** | -0.6867434 | 27.1578973 | -3.7508042 | 0.00754624 | 0.11085336 | -2.2486786 | downregulated proteins |
| **Mcam** | 0.51788917 | 26.5066716 | 3.72781865 | 0.007772766 | 0.11085336 | -2.2784932 | upregulated proteins |
| **Vdac1** | -1.2054104 | 21.5370307 | -3.7136009 | 0.007916591 | 0.11085336 | -2.2969761 | downregulated proteins |
| **Tppp3** | -0.5749736 | 27.9498256 | -3.712333 | 0.007929557 | 0.11085336 | -2.2986258 | downregulated proteins |
| **Atp5pf** | 1.14875843 | 22.3920259 | 3.71091035 | 0.007944133 | 0.11085336 | -2.3004772 | upregulated proteins |
| **Itfg1** | 0.54199775 | 28.7035595 | 3.70731415 | 0.007981109 | 0.11085336 | -2.3051587 | upregulated proteins |
| **Rev1** | -0.9232934 | 21.5288733 | -3.6998518 | 0.008058437 | 0.11085336 | -2.3148793 | downregulated proteins |
| **Endod1** | -0.5442246 | 28.6002902 | -3.6988988 | 0.008068372 | 0.11085336 | -2.3161214 | downregulated proteins |
| **Aqp4** | -0.5370287 | 25.166431 | -3.694211 | 0.008117431 | 0.11085336 | -2.3222328 | downregulated proteins |
| **Adam11** | -0.6310452 | 27.2345267 | -3.6863859 | 0.008200051 | 0.11085336 | -2.332442 | downregulated proteins |
| **Mag** | -0.565855 | 26.5231436 | -3.6824876 | 0.008241552 | 0.11085336 | -2.3375314 | downregulated proteins |
| **Prps1** | 1.22983171 | 23.53826 | 3.66594781 | 0.008420194 | 0.11158678 | -2.359151 | upregulated proteins |
| **Lamp5** | 0.75548814 | 24.6255888 | 3.65559387 | 0.008534168 | 0.11158678 | -2.372706 | upregulated proteins |
| **Sar1a** | -0.5372479 | 29.3352754 | -3.6536683 | 0.008555549 | 0.11158678 | -2.3752287 | downregulated proteins |
| **Rap1gds1** | -0.6081903 | 29.4923423 | -3.6513587 | 0.00858127 | 0.11158678 | -2.3782553 | downregulated proteins |
| **Psmd6** | 0.52350875 | 28.7148571 | 3.64633826 | 0.008637474 | 0.11158678 | -2.384837 | upregulated proteins |
| **Cadps** | 0.65589016 | 26.2093298 | 3.64344587 | 0.008670035 | 0.11158678 | -2.3886306 | upregulated proteins |
| **Cab39** | 0.98373341 | 23.2495091 | 3.64279804 | 0.008677347 | 0.11158678 | -2.3894804 | upregulated proteins |
| **Kcnc3** | -0.6219098 | 30.5292328 | -3.629337 | 0.008830796 | 0.11229829 | -2.4071535 | downregulated proteins |
| **Etfdh** | -0.7561109 | 26.2368074 | -3.6205116 | 0.008933001 | 0.11253002 | -2.4187552 | downregulated proteins |
| **Npepps** | -0.6412135 | 27.6906142 | -3.6007706 | 0.009166307 | 0.11403086 | -2.4447486 | downregulated proteins |
| **Ndufb8** | 0.50002273 | 25.9400215 | 3.5885873 | 0.009313594 | 0.11501283 | -2.4608197 | upregulated proteins |
| **Pura** | 1.1258595 | 19.6750244 | 3.58481754 | 0.009359687 | 0.11501283 | -2.4657968 | upregulated proteins |
| **Agpat3** | -0.8381432 | 23.8775397 | -3.5818634 | 0.00939598 | 0.11501283 | -2.4696986 | downregulated proteins |
| **Rufy3** | -0.5752557 | 26.0731899 | -3.5674555 | 0.009575195 | 0.11658309 | -2.4887468 | downregulated proteins |
| **Psat1** | 0.67843887 | 24.9429422 | 3.55538918 | 0.009728141 | 0.11702908 | -2.5047228 | upregulated proteins |

| | | | | | | | |
|---|---|---|---|---|---|---|---|
| **Gdi2** | -1.0035695 | 23.4059239 | -3.5431468 | 0.009886034 | 0.11702908 | -2.5209537 | downregulated proteins |
| **Tnc** | 0.57535477 | 24.2182701 | 3.54203501 | 0.00990051 | 0.11702908 | -2.5224288 | upregulated proteins |
| **Cntnap2** | -0.6569151 | 27.8897705 | -3.5374534 | 0.009960409 | 0.11702908 | -2.5285095 | downregulated proteins |
| **Emc1** | 0.66214944 | 22.9813817 | 3.52968085 | 0.010062925 | 0.11752059 | -2.538832 | upregulated proteins |
| **Itgav** | 1.26924843 | 22.9278385 | 3.51006348 | 0.010326796 | 0.11915364 | -2.5649244 | upregulated proteins |
| **Ppp1r7** | -0.5019489 | 23.8874674 | -3.498412 | 0.010487063 | 0.11915364 | -2.5804479 | downregulated proteins |
| **Ndufb9** | 1.48077869 | 21.5462391 | 3.48933349 | 0.010613807 | 0.11915364 | -2.5925568 | upregulated proteins |
| **Psma4** | 0.56260426 | 28.1689796 | 3.48748302 | 0.010639844 | 0.11915364 | -2.5950264 | upregulated proteins |
| **Cend1** | 1.65828206 | 23.8296867 | 3.47833023 | 0.010769651 | 0.11915364 | -2.6072487 | upregulated proteins |
| **Nipsnap1** | -1.221358 | 24.1899699 | -3.4755886 | 0.010808867 | 0.11915364 | -2.6109121 | downregulated proteins |
| **Them4** | 1.76207438 | 22.7022354 | 3.45811928 | 0.011062405 | 0.11981767 | -2.6342797 | upregulated proteins |
| **Prrt1** | 0.69989154 | 23.9577341 | 3.45576003 | 0.011097137 | 0.11981767 | -2.6374388 | upregulated proteins |
| **Arhgdia** | 0.57572522 | 27.4169954 | 3.41108582 | 0.011777569 | 0.12310034 | -2.6974054 | upregulated proteins |
| **Eps15l1** | -0.6286692 | 26.1301163 | -3.394264 | 0.012045361 | 0.12475942 | -2.7200568 | downregulated proteins |
| **Ube2n** | 0.78144195 | 21.347346 | 3.37787142 | 0.012312659 | 0.1263842 | -2.7421672 | upregulated proteins |
| **Eif5a** | -0.624961 | 28.3427823 | -3.3478554 | 0.01281885 | 0.12983339 | -2.7827466 | downregulated proteins |
| **Lrp1** | 1.11769815 | 20.9665289 | 3.30491789 | 0.013582361 | 0.13517402 | -2.8410016 | upregulated proteins |
| **Me1** | -0.6134638 | 28.1877574 | -3.2926845 | 0.013808762 | 0.13683228 | -2.857643 | downregulated proteins |
| **Atp1b3** | -0.5675099 | 28.6316218 | -3.2655033 | 0.014326473 | 0.14014229 | -2.8946862 | downregulated proteins |
| **Tpi1** | -0.5835155 | 29.3117358 | -3.2503304 | 0.014624517 | 0.14062518 | -2.9154047 | downregulated proteins |
| **Psmb1** | -0.5447699 | 29.599437 | -3.249125 | 0.014648479 | 0.14062518 | -2.9170518 | downregulated proteins |
| **Kpnb1** | 0.57249032 | 26.1796638 | 3.24668492 | 0.014697114 | 0.14062518 | -2.9203867 | upregulated proteins |
| **Creld1** | -0.5340874 | 27.1868876 | -3.24617 | 0.0147074 | 0.14062518 | -2.9210906 | downregulated proteins |
| **Itgb1** | 1.47949672 | 21.6000895 | 3.24446543 | 0.014741504 | 0.14062518 | -2.9234208 | upregulated proteins |
| **Etfb** | 0.82379363 | 23.7273361 | 3.24431843 | 0.014744449 | 0.14062518 | -2.9236218 | upregulated proteins |
| **Sh3bgrl3** | 0.93620302 | 24.6087334 | 3.22916611 | 0.015051437 | 0.14195782 | -2.9443518 | upregulated proteins |
| **Alb** | -0.6140064 | 26.0484596 | -3.2267613 | 0.015100786 | 0.14195782 | -2.9476444 | downregulated proteins |
| **Aars1** | -0.6226111 | 22.5020521 | -3.2209403 | 0.015220964 | 0.14195782 | -2.9556173 | downregulated proteins |
| **Timm9** | -0.634639 | 29.9644668 | -3.2194776 | 0.015251323 | 0.14195782 | -2.9576214 | downregulated proteins |
| **Trpc3** | -0.6117487 | 28.7523035 | -3.2011395 | 0.015637506 | 0.14310023 | -2.9827687 | downregulated proteins |
| **Pea15** | -0.626578 | 27.5902955 | -3.194766 | 0.015774174 | 0.14310023 | -2.9915182 | downregulated proteins |
| **Ywhaq** | 0.52120322 | 24.9068393 | 3.19136847 | 0.015847551 | 0.14310023 | -2.9961843 | upregulated proteins |
| **Basp1** | 0.65609816 | 23.8599123 | 3.18218301 | 0.01604777 | 0.14310023 | -3.0088061 | upregulated proteins |
| **Slc8a1** | 0.50059375 | 24.7711454 | 3.18024837 | 0.016090285 | 0.14310023 | -3.0114658 | upregulated proteins |
| **Fundc2** | -1.1604113 | 26.2147083 | -3.1624145 | 0.016487931 | 0.14424435 | -3.0360037 | downregulated proteins |
| **Csnk2a1** | -0.5342361 | 27.7472747 | -3.1612311 | 0.01651469 | 0.14424435 | -3.0376334 | downregulated proteins |
| **Mtfp1** | -1.0375294 | 21.7447425 | -3.1611011 | 0.016517631 | 0.14424435 | -3.0378124 | downregulated proteins |
| **Zmpste24** | -0.815762 | 24.3286043 | -3.1395436 | 0.017013374 | 0.14500449 | -3.0675258 | downregulated proteins |
| **Crym** | 0.51546647 | 29.9293483 | 3.13924902 | 0.017020257 | 0.14500449 | -3.0679322 | upregulated proteins |

| | | | | | | | |
|---|---|---|---|---|---|---|---|
| **Rexo2** | -0.6209731 | 22.3833367 | -3.1383741 | 0.017040719 | 0.14500449 | -3.0691393 | downregulated proteins |
| **Myo6** | 1.54710877 | 23.2549639 | 3.13472681 | 0.017126305 | 0.14519301 | -3.0741721 | upregulated proteins |
| **Bcap31** | 0.69422854 | 22.8798697 | 3.12967407 | 0.017245632 | 0.14566513 | -3.0811469 | upregulated proteins |
| **Nsdhl** | -0.733907 | 25.5454747 | -3.1265598 | 0.017319621 | 0.14568756 | -3.0854471 | downregulated proteins |
| **Gnas** | -0.5043987 | 29.2696707 | -3.1193506 | 0.017492209 | 0.14568756 | -3.095406 | downregulated proteins |
| **Prrt2** | -0.5878173 | 21.40244 | -3.1156685 | 0.017581067 | 0.14580819 | -3.1004948 | downregulated proteins |
| **Slc9a6** | -1.201937 | 25.8919655 | -3.108631 | 0.017752244 | 0.14623364 | -3.1102248 | downregulated proteins |
| **Uggt1** | 0.62803202 | 23.9495455 | 3.10274365 | 0.017896815 | 0.14623364 | -3.1183688 | upregulated proteins |
| **Mtatp8** | -0.9011461 | 24.4371739 | -3.1010227 | 0.017939314 | 0.14623364 | -3.1207502 | downregulated proteins |
| **Uqcrc2** | -0.5980995 | 30.1955702 | -3.0940423 | 0.018112799 | 0.14623364 | -3.1304124 | downregulated proteins |
| **Mtcl3** | 0.59080738 | 22.815713 | 3.09249404 | 0.018151519 | 0.14623364 | -3.1325562 | upregulated proteins |
| **Camkv** | 1.12941544 | 23.8445987 | 3.08245981 | 0.018404627 | 0.14647242 | -3.1464563 | upregulated proteins |
| **Lonp1** | -0.5494978 | 25.6090922 | -3.0818093 | 0.018421166 | 0.14647242 | -3.1473578 | downregulated proteins |
| **Hspa9** | -0.8152296 | 26.5408346 | -3.0815001 | 0.018429033 | 0.14647242 | -3.1477863 | downregulated proteins |
| **Dcps** | 1.18926942 | 20.9786022 | 3.07603147 | 0.018568758 | 0.14707228 | -3.1553669 | upregulated proteins |
| **Eno2** | -0.8562782 | 20.9843183 | -3.0628791 | 0.018909463 | 0.14925435 | -3.1736116 | downregulated proteins |
| **Hycc2** | 2.28644549 | 21.7644682 | 3.05168618 | 0.019204665 | 0.15012594 | -3.1891523 | upregulated proteins |
| **Dlg2** | -0.5825207 | 27.1814717 | -3.0463519 | 0.019347077 | 0.15063082 | -3.1965631 | downregulated proteins |
| **Pgrmc2** | 0.59799296 | 25.1581211 | 3.03928208 | 0.019537565 | 0.15098281 | -3.2063896 | upregulated proteins |
| **Ndufs2** | -0.6205498 | 28.1900791 | -3.0386904 | 0.019553598 | 0.15098281 | -3.2072122 | downregulated proteins |
| **Ptpa** | -0.6481831 | 24.9552537 | -3.0373427 | 0.019590168 | 0.15098281 | -3.2090861 | downregulated proteins |
| **Exoc2** | 0.64171435 | 23.6882598 | 3.0278588 | 0.019849589 | 0.15195889 | -3.2222777 | upregulated proteins |
| **Cd9** | 0.97201851 | 22.6936838 | 3.02211667 | 0.020008436 | 0.15215717 | -3.230269 | upregulated proteins |
| **Gstk1** | 0.50555804 | 27.1508788 | 3.00084537 | 0.020608778 | 0.1556881 | -3.2599006 | upregulated proteins |
| **Taldo1** | -0.5580091 | 27.7321038 | -2.9955168 | 0.020762153 | 0.15633081 | -3.2673303 | downregulated proteins |
| **Syt2** | -0.528018 | 29.1861083 | -2.9715718 | 0.021466519 | 0.15991055 | -3.3007502 | downregulated proteins |
| **Timm50** | -0.602443 | 27.2765111 | -2.9526178 | 0.022042076 | 0.16163023 | -3.3272415 | downregulated proteins |
| **Atcay** | -0.5317845 | 28.6953242 | -2.9468675 | 0.022219917 | 0.16197895 | -3.335285 | downregulated proteins |
| **Atp2a3** | -0.8222782 | 23.6624202 | -2.9430513 | 0.022338779 | 0.16199736 | -3.3406245 | downregulated proteins |
| **Ppia** | 1.10397588 | 22.2358438 | 2.94224483 | 0.022363986 | 0.16199736 | -3.3417532 | upregulated proteins |
| **Psmc2** | -0.807544 | 23.670047 | -2.9357005 | 0.022569646 | 0.16297136 | -3.3509135 | downregulated proteins |
| **Psmd8** | 0.60577887 | 24.1507808 | 2.91348081 | 0.023283021 | 0.16654636 | -3.3820424 | upregulated proteins |
| **Prdx2** | -0.5166998 | 27.502797 | -2.8955057 | 0.023877614 | 0.1692349 | -3.4072552 | downregulated proteins |
| **Ktn1** | 0.58335807 | 26.7302483 | 2.89321458 | 0.023954549 | 0.1692349 | -3.4104707 | upregulated proteins |
| **Ube2d2** | 0.52620264 | 26.836314 | 2.86115819 | 0.025058975 | 0.17280242 | -3.4555044 | upregulated proteins |
| **Pi4ka** | -0.5857924 | 28.1871088 | -2.8561408 | 0.025236658 | 0.17347361 | -3.4625602 | downregulated proteins |
| **Hspa12a** | 0.60911152 | 26.2284845 | 2.85400017 | 0.025312871 | 0.17347653 | -3.465571 | upregulated proteins |
| **Hspd1** | -0.7226156 | 25.3930766 | -2.8130937 | 0.026816957 | 0.17771215 | -3.5231704 | downregulated proteins |
| **Lingo1** | 0.90402743 | 20.9195797 | 2.81221582 | 0.026850251 | 0.17771215 | -3.5244077 | upregulated proteins |

| | | | | | | | |
|---|---|---|---|---|---|---|---|
| **Epn2** | 0.74457709 | 23.9717406 | 2.80920124 | 0.026964915 | 0.17787519 | -3.5286572 | upregulated proteins |
| **Grm5** | -0.6603754 | 23.9283484 | -2.8055206 | 0.027105615 | 0.17828952 | -3.5338465 | downregulated proteins |
| **Pitpna** | 0.67302889 | 19.5153978 | 2.79992106 | 0.027321156 | 0.17919234 | -3.5417429 | upregulated proteins |
| **Cnot11** | -0.5331357 | 23.3749328 | -2.7777707 | 0.028191658 | 0.18211057 | -3.572999 | downregulated proteins |
| **Rab4b** | -0.6509078 | 24.7650478 | -2.7761152 | 0.028257879 | 0.18211057 | -3.5753362 | downregulated proteins |
| **Uba1** | 0.66662987 | 23.3312121 | 2.77449107 | 0.028323007 | 0.18211057 | -3.5776293 | upregulated proteins |
| **Itm2b** | -13.552999 | 15.7889577 | -2.7692724 | 0.02853335 | 0.18294912 | -3.5849988 | downregulated proteins |
| **Nefh** | 0.53598474 | 26.4302971 | 2.76504161 | 0.028705081 | 0.18353612 | -3.5909744 | upregulated proteins |
| **Slc44a1** | -0.6809051 | 27.6130318 | -2.7291553 | 0.030206215 | 0.18962708 | -3.6417018 | downregulated proteins |
| **Chid1** | 0.92563411 | 22.1292308 | 2.72902414 | 0.030211853 | 0.18962708 | -3.6418874 | upregulated proteins |
| **Acot2** | 0.82313066 | 21.8567387 | 2.72866101 | 0.03022746 | 0.18962708 | -3.6424011 | upregulated proteins |
| **Blmh** | 0.70346955 | 25.1899807 | 2.72672107 | 0.030310986 | 0.18962708 | -3.6451454 | upregulated proteins |
| **Gprc5b** | 0.70281783 | 23.6379111 | 2.72127565 | 0.03054674 | 0.18962708 | -3.6528496 | upregulated proteins |
| **Plppr4** | -0.6452166 | 25.6475495 | -2.7133468 | 0.030893462 | 0.19112199 | -3.6640701 | downregulated proteins |
| **Arpc5l** | -0.6261542 | 23.0863684 | -2.7099714 | 0.031042316 | 0.19152523 | -3.6688478 | downregulated proteins |
| **Cnnm1** | 0.78847614 | 20.8800024 | 2.69439147 | 0.031739205 | 0.19364722 | -3.6909071 | upregulated proteins |
| **Hspa8** | 0.59842246 | 25.2489646 | 2.69369774 | 0.031770615 | 0.19364722 | -3.6918896 | upregulated proteins |
| **F3** | -0.7117845 | 29.9368144 | -2.6827091 | 0.032272508 | 0.19489255 | -3.7074552 | downregulated proteins |
| **Glod4** | 0.50944474 | 25.4203672 | 2.67757604 | 0.032509797 | 0.19504426 | -3.7147282 | upregulated proteins |
| **Dld** | 0.71000353 | 23.2188931 | 2.66407499 | 0.033142679 | 0.19623884 | -3.7338628 | upregulated proteins |
| **Rps6** | 0.83669647 | 24.2679858 | 2.66163458 | 0.033258447 | 0.19623884 | -3.7373223 | upregulated proteins |
| **Atp5f1c** | 1.00806003 | 20.0695853 | 2.63706535 | 0.034447803 | 0.2021821 | -3.7721638 | upregulated proteins |
| **Ndrg1** | 0.51263235 | 23.9558526 | 2.62897646 | 0.034849034 | 0.20298338 | -3.7836393 | upregulated proteins |
| **Myl12b** | 1.28526217 | 23.275955 | 2.62894994 | 0.034850357 | 0.20298338 | -3.7836769 | upregulated proteins |
| **Got2** | 0.81622217 | 24.1760475 | 2.620747 | 0.035262249 | 0.20382648 | -3.7953163 | upregulated proteins |
| **Lmnb1** | 0.56246561 | 25.9946873 | 2.60807569 | 0.035908526 | 0.20651913 | -3.8133 | upregulated proteins |
| **Ndufb3** | 0.61628748 | 26.5857354 | 2.60207438 | 0.036218912 | 0.20706455 | -3.821819 | upregulated proteins |
| **Plpbp** | 0.66922495 | 25.7634331 | 2.60100062 | 0.036274741 | 0.20706455 | -3.8233433 | upregulated proteins |
| **Slc25a22** | 0.64928692 | 23.2603492 | 2.5825512 | 0.037248141 | 0.2108132 | -3.8495387 | upregulated proteins |
| **Cplx1** | 0.55990797 | 23.9439139 | 2.57888859 | 0.037444598 | 0.2108132 | -3.85474 | upregulated proteins |
| **Rab3c** | -0.5171515 | 27.8910175 | -2.5709311 | 0.037875155 | 0.2112128 | -3.8660415 | downregulated proteins |
| **Atp5f1b** | 0.73451503 | 21.6842667 | 2.56897182 | 0.037981952 | 0.2112128 | -3.8688243 | upregulated proteins |
| **Fkbp1a** | 1.08347151 | 22.9234956 | 2.56710997 | 0.038083731 | 0.2112128 | -3.8714688 | upregulated proteins |
| **Col7a1** | 0.58554618 | 22.7707183 | 2.56657126 | 0.038113233 | 0.2112128 | -3.8722339 | upregulated proteins |
| **Mapk3** | 0.51712622 | 24.0307712 | 2.55635117 | 0.038677447 | 0.21182959 | -3.8867514 | upregulated proteins |
| **Bcat1** | 0.58707692 | 21.9062371 | 2.55436463 | 0.038788122 | 0.21189979 | -3.8895734 | upregulated proteins |
| **Gap43** | -0.6431048 | 28.2996077 | -2.5490999 | 0.039083028 | 0.2124313 | -3.8970528 | downregulated proteins |
| **Hsd17b11** | -0.7361154 | 23.0796289 | -2.5414198 | 0.03951741 | 0.21262684 | -3.907964 | downregulated proteins |
| **Ndufv1** | -0.5638663 | 25.7723351 | -2.5373065 | 0.039752121 | 0.21262684 | -3.9138083 | downregulated proteins |
| **Dnajc3** | 0.53283185 | 23.3387235 | 2.53471473 | 0.039900752 | 0.21262684 | -3.9174908 | upregulated proteins |

| | | | | | | | |
|---|---|---|---|---|---|---|---|
| **Ndufb5** | 0.59925782 | 23.9148325 | 2.53205973 | 0.040053607 | 0.21262684 | -3.9212633 | upregulated proteins |
| **Edil3** | -1.0355072 | 22.500499 | -2.5300768 | 0.040168168 | 0.21262684 | -3.9240808 | downregulated proteins |
| **Asrgl1** | -0.5613587 | 26.0653894 | -2.5223523 | 0.040617683 | 0.21373305 | -3.9350569 | downregulated proteins |
| **Grm1** | 1.69859489 | 20.3198975 | 2.51072232 | 0.041304344 | 0.21408511 | -3.9515834 | upregulated proteins |
| **Cnp** | 0.94456488 | 21.9666849 | 2.50821545 | 0.041453925 | 0.21408511 | -3.9551458 | upregulated proteins |
| **Gcsh** | -0.6052461 | 26.6434681 | -2.5058757 | 0.041594042 | 0.21408511 | -3.9584708 | downregulated proteins |
| **Sorl1** | -1.9111632 | 24.8287551 | -2.5054627 | 0.041618826 | 0.21408511 | -3.9590577 | downregulated proteins |
| **Ndufa3** | 0.53454154 | 25.3157738 | 2.50450528 | 0.041676337 | 0.21408511 | -3.9604182 | upregulated proteins |
| **Pcmt1** | 0.53410462 | 19.995122 | 2.50283713 | 0.041776739 | 0.21408511 | -3.9627889 | upregulated proteins |
| **Tomm22** | -0.7765633 | 24.7483491 | -2.5023365 | 0.041806922 | 0.21408511 | -3.9635003 | downregulated proteins |
| **Alg2** | -0.7697522 | 21.9878793 | -2.4934944 | 0.042343709 | 0.21634989 | -3.9760659 | downregulated proteins |
| **Hnrnpm** | 14.2504641 | 17.6634911 | 2.48776256 | 0.042695486 | 0.21653082 | -3.9842115 | upregulated proteins |
| **Rps14** | -1.4573526 | 21.9830646 | -2.4674762 | 0.043964992 | 0.22119273 | -4.0130403 | downregulated proteins |
| **Aldh18a1** | -0.611222 | 23.5984593 | -2.4640936 | 0.044180438 | 0.22177417 | -4.0178471 | downregulated proteins |
| **Txndc5** | 0.70655331 | 22.678672 | 2.45572806 | 0.04471796 | 0.22344028 | -4.0297346 | upregulated proteins |
| **Slc25a5** | 0.52802604 | 23.7499019 | 2.45381662 | 0.044841724 | 0.22344028 | -4.0324507 | upregulated proteins |
| **Atp1a3** | 0.60921456 | 26.5622563 | 2.44604385 | 0.045348656 | 0.2241967 | -4.0434952 | upregulated proteins |
| **Uqcr10** | 0.75066558 | 23.3251669 | 2.41799336 | 0.047227768 | 0.22758813 | -4.0833462 | upregulated proteins |
| **Nfu1** | -0.60874 | 19.5880912 | -2.4049098 | 0.048131494 | 0.23028741 | -4.1019293 | downregulated proteins |
| **Sparcl1** | -0.6461925 | 24.302886 | -2.3996171 | 0.048502122 | 0.23081363 | -4.1094457 | downregulated proteins |
| **Folh1** | 2.18272715 | 24.2569966 | 2.39313472 | 0.048960051 | 0.23233046 | -4.1186506 | upregulated proteins |
| **Pfkm** | 0.62786828 | 26.6219769 | 2.39223644 | 0.049023858 | 0.23233046 | -4.1199261 | upregulated proteins |
| **Rpl27** | -0.7549196 | 23.1490517 | -2.3868273 | 0.049409888 | 0.2336761 | -4.1276061 | downregulated proteins |

#### Discovery Analysis (Advanced stage (10M) samples)

| Proteins | logFC | AveExpr | t | P.Value | adj.P.Val | B | Category |
|---|---|---|---|---|---|---|---|
| **Hprt1** | -1.3591324 | 24.1722662 | -8.9238509 | 5.33737222260029E-05 | 0.12313318 | -0.8561095 | downregulated proteins |
| **Glo1** | 0.91313895 | 21.8790261 | 6.09250387 | 0.000551956 | 0.43340542 | -1.47807 | upregulated proteins |
| **Atp6v1g1** | 1.60959423 | 22.3616801 | 5.7949931 | 0.000738898 | 0.43340542 | -1.5791092 | upregulated proteins |
| **S100a1** | -0.5542213 | 23.4467489 | -5.7781205 | 0.000751461 | 0.43340542 | -1.585123 | downregulated proteins |
| **Kcnj10** | -1.1193064 | 21.5332552 | -5.258723 | 0.001284964 | 0.5928822 | -1.7865892 | downregulated proteins |
| **Ndufaf2** | 0.53613059 | 25.7252552 | 4.73051395 | 0.002300385 | 0.79570878 | -2.0278838 | upregulated proteins |
| **Rida** | 0.5633735 | 22.3010775 | 4.68818246 | 0.002414374 | 0.79570878 | -2.0489936 | upregulated proteins |

| | | | | | | | |
|---|---|---|---|---|---|---|---|
| **Ppp1ca** | 0.78565381 | 23.9099429 | 4.53657282 | 0.00287709 | 0.82968093 | -2.126905 | upregulated proteins |
| **Fabp3** | 0.88162628 | 21.5240438 | 4.37895571 | 0.003464728 | 0.84328435 | -2.2118457 | upregulated proteins |
| **Tusc3** | 0.81559011 | 25.4148102 | 4.12456277 | 0.004713361 | 0.84328435 | -2.3578067 | upregulated proteins |
| **Gsta4** | 0.51273865 | 23.4088259 | 4.11585821 | 0.004764078 | 0.84328435 | -2.3630008 | upregulated proteins |
| **Psap** | 0.58561107 | 26.7122038 | 4.06469839 | 0.00507456 | 0.84328435 | -2.3938011 | upregulated proteins |
| **Arf6** | 0.61333384 | 23.4197484 | 3.82702019 | 0.006839718 | 0.84328435 | -2.5431096 | upregulated proteins |
| **Sacm1l** | -0.6786373 | 27.3460579 | -3.7656408 | 0.007398092 | 0.84328435 | -2.5833593 | downregulated proteins |
| **App** | 2.0042263 | 22.0021863 | 3.5859904 | 0.009338667 | 0.84328435 | -2.7052276 | upregulated proteins |
| **Dclk1** | 0.72723989 | 18.7310656 | 3.57724352 | 0.009446346 | 0.84328435 | -2.7113168 | upregulated proteins |
| **Samm50** | 0.6878859 | 24.4888419 | 3.5244965 | 0.010124956 | 0.84328435 | -2.748344 | upregulated proteins |
| **Got1** | 1.23645398 | 19.4420564 | 3.45636374 | 0.01108087 | 0.84328435 | -2.7969529 | upregulated proteins |
| **Cds1** | -0.9986552 | 26.0466543 | -3.4469656 | 0.011220224 | 0.84328435 | -2.8037271 | downregulated proteins |
| **Ppp2r5c** | 0.90904093 | 21.6671231 | 3.43272106 | 0.011435066 | 0.84328435 | -2.8140264 | upregulated proteins |
| **Tpt1** | 0.80749832 | 19.0451233 | 3.32889843 | 0.013141945 | 0.84328435 | -2.8902551 | upregulated proteins |
| **Ist1** | -16.97526 | 13.990565 | -3.2837328 | 0.013968584 | 0.84328435 | -2.9240517 | downregulated proteins |
| **Uqcrh** | 0.52354015 | 21.8145164 | 3.26249844 | 0.01437642 | 0.84328435 | -2.9400734 | upregulated proteins |
| **Sncb** | 0.55302926 | 21.1520685 | 3.22433412 | 0.015141968 | 0.84328435 | -2.9690812 | upregulated proteins |
| **Txnrd1** | -1.4130187 | 21.7315102 | -3.1462263 | 0.01684868 | 0.84328435 | -3.0292932 | downregulated proteins |
| **Calb2** | 0.61475863 | 23.4622562 | 3.13922606 | 0.017011412 | 0.84328435 | -3.0347446 | upregulated proteins |
| **Vamp2** | -0.8009784 | 29.6407305 | -3.1157294 | 0.017570048 | 0.84328435 | -3.053108 | downregulated proteins |
| **Uqcrq** | -0.6282777 | 25.636655 | -3.0833216 | 0.018372967 | 0.84328435 | -3.0786011 | downregulated proteins |
| **Pdcd6** | 15.1191802 | 12.7887747 | 3.07661944 | 0.01854386 | 0.84328435 | -3.083897 | upregulated proteins |
| **Lgi3** | 0.86566723 | 24.9640087 | 3.02756051 | 0.019847632 | 0.84328435 | -3.122909 | upregulated proteins |
| **Pvalb** | -0.5485745 | 21.8407185 | -3.0164737 | 0.020155604 | 0.84328435 | -3.131785 | downregulated proteins |
| **Kcna1** | 1.18960055 | 22.5914118 | 2.95001137 | 0.022111748 | 0.84328435 | -3.1854494 | upregulated proteins |
| **Hcn2** | -1.0825403 | 24.3701039 | -2.93652 | 0.022532931 | 0.84328435 | -3.1964372 | downregulated proteins |
| **Pdp1** | 0.76053644 | 23.4722155 | 2.92369716 | 0.022941144 | 0.84328435 | -3.2069096 | upregulated proteins |
| **Atp5if1** | -1.2110442 | 20.6425329 | -2.9021729 | 0.023644104 | 0.84328435 | -3.224552 | downregulated proteins |
| **Arpc4** | -0.6046128 | 23.9102639 | -2.8901904 | 0.024045298 | 0.84328435 | -3.2344076 | downregulated proteins |
| **Ube2l3** | 0.89605884 | 24.7569467 | 2.79416116 | 0.027532805 | 0.84328435 | -3.3142587 | upregulated proteins |
| **Cndp2** | 0.52157892 | 24.2390238 | 2.77210898 | 0.02840669 | 0.84328435 | -3.3328084 | upregulated proteins |
| **Rab2a** | 0.58777799 | 25.6048852 | 2.75400788 | 0.029145819 | 0.84328435 | -3.3480923 | upregulated proteins |

| | | | | | | | |
|---|---|---|---|---|---|---|---|
| **Krt73** | 1.07196738 | 22.4416605 | 2.74216732 | 0.029640235 | 0.84328435 | -3.3581179 | upregulated proteins |
| **Acyp2** | 0.7931481 | 21.8162555 | 2.72515736 | 0.03036599 | 0.84328435 | -3.3725587 | upregulated proteins |
| **Vamp3** | 1.04556214 | 24.5937094 | 2.7088304 | 0.031080174 | 0.84328435 | -3.3864616 | upregulated proteins |
| **Rpl28** | 0.90315087 | 22.0418269 | 2.70792166 | 0.031120439 | 0.84328435 | -3.3872367 | upregulated proteins |
| **Vat1** | 0.61808603 | 20.2796694 | 2.69887867 | 0.031524117 | 0.84328435 | -3.3949558 | upregulated proteins |
| **Ldhb** | 0.64804367 | 23.1090864 | 2.69229948 | 0.03182126 | 0.84328435 | -3.4005796 | upregulated proteins |
| **Pde1b** | 1.32390082 | 21.3739929 | 2.68885498 | 0.031977997 | 0.84328435 | -3.4035265 | upregulated proteins |
| **Nrxn2** | 0.69220047 | 22.9354773 | 2.68278262 | 0.032256285 | 0.84328435 | -3.4087259 | upregulated proteins |
| **Slc38a3** | 0.67236117 | 20.7282983 | 2.64931163 | 0.03383641 | 0.84328435 | -3.4374834 | upregulated proteins |
| **Kcnab2** | 0.63982274 | 22.1007952 | 2.64205921 | 0.034189341 | 0.84328435 | -3.4437361 | upregulated proteins |
| **Marcksl1** | 0.87866852 | 24.8236682 | 2.6391932 | 0.034329871 | 0.84328435 | -3.4462091 | upregulated proteins |
| **Atp6v1e1** | 1.25709401 | 25.4407655 | 2.63403842 | 0.034584148 | 0.84328435 | -3.4506601 | upregulated proteins |
| **Pex5l** | 0.68569177 | 19.7381547 | 2.60789377 | 0.035904453 | 0.84328435 | -3.4732931 | upregulated proteins |
| **Hspa4l** | 0.52718493 | 24.0352174 | 2.5932763 | 0.036665455 | 0.84328435 | -3.4859889 | upregulated proteins |
| **Anxa6** | 13.0834674 | 15.8568168 | 2.58967951 | 0.036855273 | 0.84328435 | -3.4891174 | upregulated proteins |
| **Pfn2** | 1.38395926 | 22.211542 | 2.56762367 | 0.038041856 | 0.84328435 | -3.5083401 | upregulated proteins |
| **Acyp1** | -1.7228525 | 23.2514449 | -2.5158533 | 0.040985745 | 0.84328435 | -3.5537133 | downregulated proteins |
| **Praf2** | 0.64988461 | 26.3906922 | 2.51278405 | 0.041167538 | 0.84328435 | -3.5564142 | upregulated proteins |
| **Nme1** | 0.87877659 | 24.0273759 | 2.50343339 | 0.041726543 | 0.84328435 | -3.5646497 | upregulated proteins |
| **Napa** | 0.56675199 | 20.6400703 | 2.49167497 | 0.042440671 | 0.84328435 | -3.5750211 | upregulated proteins |
| **Cyc1** | -0.9108774 | 21.2011109 | -2.4824622 | 0.043009029 | 0.84328435 | -3.583159 | downregulated proteins |
| **Zfyve28** | 0.67724851 | 22.9971235 | 2.47990186 | 0.043168375 | 0.84328435 | -3.5854224 | upregulated proteins |
| **Nbea** | -1.545281 | 23.1938634 | -2.4732031 | 0.043588181 | 0.84328435 | -3.5913481 | downregulated proteins |
| **Ndufab1** | 0.50558271 | 22.26684 | 2.45546667 | 0.044720203 | 0.84328435 | -3.6070633 | upregulated proteins |
| **Calb1** | -1.1225659 | 21.0398619 | -2.4534005 | 0.044854038 | 0.84328435 | -3.6088964 | downregulated proteins |
| **Aldh6a1** | 1.003151 | 23.3906142 | 2.4355493 | 0.04602768 | 0.84328435 | -3.6247543 | upregulated proteins |
| **Uqcrb** | 0.68695573 | 23.0320414 | 2.42284057 | 0.046882489 | 0.84328435 | -3.6360659 | upregulated proteins |
| **Enoph1** | -0.9657279 | 28.8423718 | -2.4111622 | 0.04768241 | 0.84328435 | -3.646476 | downregulated proteins |
| **Cntnap1** | 0.93791659 | 23.2589386 | 2.40852314 | 0.047865116 | 0.84328435 | -3.6488305 | upregulated proteins |

#### Supplementary data 2 - SH-SY5Y tauopathy cell line maps

Tau35-HA

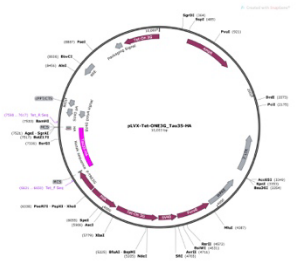

(Tau35 overexpressing)

Avi-FL tau

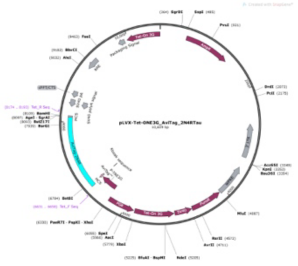

(FL tau overexpressing)

$mKO_2$-Tau35

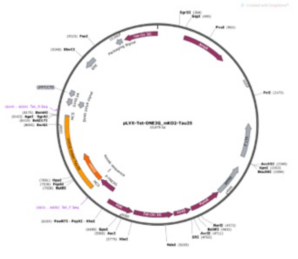

(Tau35 overexpressing)

Tau35-$mKO_2$

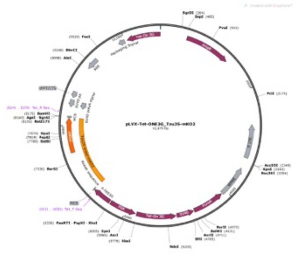

(Tau35 overexpressing)

mEGFP-FL tau

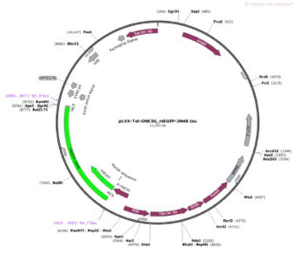

(FL tau overexpressing)

FL tau-mEGFP

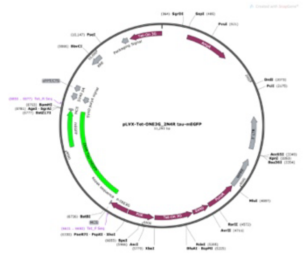

(FL tau overexpressing)

Schematic of expression plasmids for all six novel SH-SY5Y tauopathy cell lines. Generalised maps of plasmids used for the expression of truncated (Tau35) or full length (2N4R) tau.

#### Supplementary data 3 – Differential Expression Analysis of Lysosome-Enriched Brain Fractions (full list of detected proteins)

#### Early stage (4M) samples

| Proteins | logFC | AveExpr | t | P.Value | adj.P.Val | B |
|---|---|---|---|---|---|---|
| **Hprt1** | 1.48265474750001 | 23.97555172125 | 10.7238496297493 | 1.65097822452125E-05 | 0.0244583711227583 | 3.40909546573355 |
| **Glo1** | 0.889217764999994 | 24.526476365 | 10.30340242567 | 2.13703548473205E-05 | 0.0244583711227583 | 3.20826276189975 |
| **Atp6v1g1** | 1.8012870275 | 24.16786601875 | 8.18456405197047 | 9.21583025353879E-05 | 0.0520266819285532 | 1.98868801380223 |
| **S100a1** | 0.816605772499997 | 26.81497089375 | 8.01810459038053 | 0.00010475415333572 | 0.0520266819285532 | 1.87575834792631 |
| **Kcnj10** | 0.570326844999997 | 32.395900995 | 7.34756402716859 | 0.000179645523242229 | 0.0520266819285532 | 1.39100039739627 |
| **Ndufaf2** | -1.006975615 | 24.2984688625 | -7.27789635242004 | 0.000190431070901172 | 0.0520266819285532 | 1.3377458751486 |
| **Rida** | 0.766432560000002 | 25.684420015 | 7.23929893736295 | 0.000196720699718147 | 0.0520266819285532 | 1.30799700987382 |
| **Ppp1ca** | 0.587886057500008 | 33.55780611625 | 7.20083178583311 | 0.000203223796626246 | 0.0520266819285532 | 1.27817350197535 |
| **Fabp3** | 0.753589842499998 | 28.06138945375 | 7.19309443741693 | 0.00020456100365093 | 0.0520266819285532 | 1.27215353957641 |
| **Lrch2** | 0.612081519999997 | 24.64487892 | 6.93293718478017 | 0.000255879192652784 | 0.054084014974879 | 1.06553337193749 |
| **Tusc3** | 1.065376825 | 24.5783635625 | 6.91507053762176 | 0.000259905707611913 | 0.054084014974879 | 1.05103870964271 |
| **Gsta4** | -0.686724620000003 | 19.98327855 | -6.58465841088366 | 0.000348926966061932 | 0.0621058597641011 | 0.77568208895301 |
| **Psap** | -1.18205985 | 24.9800101075 | -6.50778668043794 | 0.000374282149672305 | 0.0621058597641011 | 0.709585215810784 |
| **Ank3** | 1.395740215 | 24.0841377275 | 6.49168500221645 | 0.000379852353297255 | 0.0621058597641011 | 0.69564094463875 |
| **Arf6** | 0.670161287499997 | 26.31323874375 | 6.37488627277888 | 0.000423171209924599 | 0.0634282520926951 | 0.593448165450214 |
| **Sacm1l** | 0.744573912499998 | 21.32440148875 | 6.27735451817071 | 0.000463635146187405 | 0.0634282520926951 | 0.506690425230598 |
| **Aco1** | 0.634085200000008 | 21.85112901 | 6.26047696781887 | 0.000471070461151515 | 0.0634282520926951 | 0.491544273848067 |
| **App** | 0.618876132500006 | 25.13809943375 | 6.13420732861739 | 0.00053115444391367 | 0.0675451401176883 | 0.376967423315459 |
| **Dclk1** | -0.834094519999997 | 25.1093881225 | -6.00558578132736 | 0.000601368380385799 | 0.0724490643527945 | 0.257937865059906 |
| **Samm50** | 0.48943154749999 | 27.52194465375 | 5.84288928141523 | 0.000705588594104078 | 0.0755088831917057 | 0.103951965285762 |
| **Chp1** | 0.544549112500004 | 28.69076565625 | 5.77302955456013 | 0.000756446629177741 | 0.0755088831917057 | 0.0366385953261767 |
| **Gsr** | -0.575354497499994 | 31.03726549125 | -5.75980191835246 | 0.000766532546747565 | 0.0755088831917057 | 0.0238113953194006 |

| | | | | | | |
|---|---|---|---|---|---|---|
| **Got1** | -0.877421544999997 | 27.573672035 | -5.74367053327992 | 0.000779037267058084 | 0.0755088831917057 | 0.00813298015163788 |
| **Cds1** | 0.535007235000005 | 22.7326039275 | 5.70218735303273 | 0.000812258151686023 | 0.0755088831917057 | -0.0323644293185676 |
| **Ppp2r5c** | 0.719570549999993 | 26.158952465 | 5.63901062779707 | 0.000865947405538872 | 0.0755088831917057 | -0.0945380192647098 |
| **Rab5a** | -0.632055702499994 | 23.40535611125 | -5.61794645000374 | 0.0008847236265795 | 0.0755088831917057 | -0.115402173602872 |
| **Dnajc6** | 1.17516632750001 | 23.12392417125 | 5.6113815784299 | 0.000890668346953279 | 0.0755088831917057 | -0.12191850290915 |
| **Tpt1** | -0.541086050000011 | 28.166502735 | -5.5046935013512 | 0.000993830422676245 | 0.0776979885195725 | -0.228742164462814 |
| **Timm10** | 0.941941642499994 | 22.74719440125 | 5.48561803718178 | 0.00101365119412772 | 0.0776979885195725 | -0.248026457314737 |
| **Cbarp** | 0.875676990000002 | 26.2610788025 | 5.4811833800126 | 0.0010183222610691 | 0.0776979885195725 | -0.252517728796449 |
| **Ist1** | -0.69755623 | 29.65544284 | -5.3821660410743 | 0.00112915719538902 | 0.0833755103304988 | -0.353595604781727 |
| **Uqcrh** | 0.503787702499999 | 25.41237104375 | 5.32711555603469 | 0.00119657430845305 | 0.0844461824880777 | -0.410455305087639 |
| **Cct2** | 0.717454589999999 | 26.1063588875 | 5.31077597988642 | 0.00121744168724621 | 0.0844461824880777 | -0.427423750439385 |
| **Sncb** | 0.393215335000001 | 25.0462270225 | 5.24193515466333 | 0.00130993817558438 | 0.0872550537501411 | -0.499378365408614 |
| **Txnrd1** | -0.417494129999998 | 29.808185265 | -5.18552938196024 | 0.00139159573546986 | 0.0872550537501411 | -0.558897687604356 |
| **Rab18** | -0.665133942500002 | 26.03052220875 | -5.16838870231199 | 0.00141752409052035 | 0.0872550537501411 | -0.577085343899318 |
| **Calb2** | -0.708072177499993 | 27.19127800625 | -5.13905269925527 | 0.00146316050105096 | 0.0872550537501411 | -0.608322659874132 |
| **Vamp2** | -0.687328122499999 | 27.56001458625 | -5.13113631155816 | 0.00147575481280546 | 0.0872550537501411 | -0.61677583267093 |
| **Uqcrq** | -0.410014515 | 24.1190844475 | -5.11102231350977 | 0.00150830198146673 | 0.0872550537501411 | -0.63829911357409 |
| **Pdcd6** | 1.50635791249999 | 24.08033818125 | 5.10102781773545 | 0.00152477158147909 | 0.0872550537501411 | -0.649018141225592 |
| **Mdp1** | 0.431668332500003 | 22.59505031125 | 5.0697640934563 | 0.00157759888656171 | 0.0880761914960915 | -0.682652502291971 |
| **Pa2g4** | 0.43915419499999 | 29.2249935525 | 5.014153824816 | 0.00167668380902789 | 0.0908568620783198 | -0.742871080352536 |

| | | | | | | |
|---|---|---|---|---|---|---|
| **Ncald** | 0.770149924999995 | 25.03717294 | 4.99797915486734 | 0.0017067912054905 | 0.0908568620783198 | -0.760480502937173 |
| **Sez6l2** | 0.588065502499994 | 22.14036744875 | 4.97292817287463 | 0.00175461675510548 | 0.09127994891901 | -0.787837787101505 |
| **Lgi3** | -0.868674297500007 | 29.05070374875 | -4.92928562418458 | 0.00184154243049063 | 0.0926456596493285 | -0.83574321423395 |
| **Pvalb** | 0.495677945000004 | 26.148496065 | 4.90990513509714 | 0.00188167237363335 | 0.0926456596493285 | -0.857116763774388 |
| **Uso1** | 0.630902234999997 | 25.39301901 | 4.85828190079791 | 0.00199339606359557 | 0.0926456596493285 | -0.914349901079986 |
| **Akr1b1** | -0.837079562500005 | 28.90947929125 | -4.84800555097048 | 0.00201650595858693 | 0.0926456596493285 | -0.925795327518736 |
| **Plec** | 0.912090252499993 | 24.48016117875 | 4.84772141808474 | 0.00201714914773139 | 0.0926456596493285 | -0.926112031708529 |
| **Gstm1** | -0.586997847499998 | 24.24421241625 | -4.83706747641718 | 0.00204143182032467 | 0.0926456596493285 | -0.937996885502494 |
| **Kcna1** | -0.669744527500008 | 27.28645012875 | -4.82720997120448 | 0.00206418900922488 | 0.0926456596493285 | -0.949009963139556 |
| **Hcn2** | -0.891227685000011 | 27.531552075 | -4.72185793305411 | 0.00232583966628182 | 0.0993382092601092 | -1.06771588266425 |
| **C1qbp** | 0.379807502500004 | 25.48041010375 | 4.70621375854751 | 0.00236774540067593 | 0.0993382092601092 | -1.08549994770805 |
| **Pdp1** | -1.07250894250001 | 24.22400715875 | -4.70291004345572 | 0.00237670178316634 | 0.0993382092601092 | -1.08926076226254 |
| **Lgi2** | -0.571020335 | 22.2161669875 | -4.69798999744105 | 0.00239010982911312 | 0.0993382092601092 | -1.09486490459551 |
| **Atp5if1** | -0.618241842500005 | 22.36847781875 | -4.66832568463132 | 0.00247275133853889 | 0.0993382092601092 | -1.12873913062344 |
| **Arpc4** | 0.821779247500011 | 24.15487819875 | 4.65840635923303 | 0.00250108896378256 | 0.0993382092601092 | -1.14009886138115 |
| **Ndrg4** | -0.702302167499997 | 28.78305971375 | -4.63014740776234 | 0.00258381394247633 | 0.0993382092601092 | -1.17255120258503 |
| **Smpd3** | -0.636006362499998 | 31.02604026375 | -4.62844165334197 | 0.002588903602705 | 0.0993382092601092 | -1.17451433536082 |
| **Uchl1** | -1.041545385 | 24.49119615 | -4.61065819901914 | 0.00264263344589524 | 0.0993382092601092 | -1.19500999892029 |
| **Ube2l3** | -0.96705043 | 21.9990241525 | -4.60913817715125 | 0.00264728299033056 | 0.0993382092601092 | -1.19676429073225 |
| **Cndp2** | 0.444231140000003 | 27.3544722375 | 4.58710766193123 | 0.00271569962890971 | 0.10026187823507 | -1.22223349209661 |
| **Rab2a** | 1.17285902999999 | 24.791264405 | 4.56938782350122 | 0.00277215233893911 | 0.100577114249108 | -1.2427779583149 |

| | | | | | | |
|---|---|---|---|---|---|---|
| **Krt73** | 0.769241480000005 | 24.2245667275 | 4.54815928736063 | 0.0028415018009382 | 0.100577114249108 | -1.26745946475438 |
| **Tkt** | -0.59660418499999 | 29.2760677625 | -4.53510161210906 | 0.00288511207905809 | 0.100577114249108 | -1.28267845979615 |
| **Acyp2** | 0.46815933500001 | 24.8814847025 | 4.51941003538062 | 0.00293850251616449 | 0.100577114249108 | -1.30100501779114 |
| **Hagh** | -0.490599192500003 | 28.03604049625 | -4.501272892974 | 0.00300158117470814 | 0.100577114249108 | -1.32223909027053 |
| **Psma6** | -0.672433617499994 | 27.95665071875 | -4.48624831137814 | 0.00305497072134559 | 0.100577114249108 | -1.33987078216909 |
| **Vamp3** | 1.3766828125 | 23.63404169875 | 4.46510189104753 | 0.00313190309784015 | 0.100577114249108 | -1.36475054737243 |
| **DSCC1** | 0.587509310000009 | 25.4214889725 | 4.45325779637518 | 0.00317592764861987 | 0.100577114249108 | -1.37871836054003 |
| **Pfn1** | 0.558504152500007 | 19.66361864375 | 4.43432296965569 | 0.00324773723460061 | 0.100577114249108 | -1.40109705437456 |
| **Rpl28** | -0.439632724999996 | 30.25488037 | -4.42109443502153 | 0.00329897089920667 | 0.100577114249108 | -1.41676717630447 |
| **Vat1** | 0.792433114999994 | 24.7966748375 | 4.41318602929957 | 0.00333002627394432 | 0.100577114249108 | -1.42614921284374 |
| **Ldhb** | -0.651766302500004 | 24.18161521375 | -4.40256692080761 | 0.00337223570728322 | 0.100577114249108 | -1.43876350859139 |
| **Pde1b** | 0.678249727500003 | 24.66307125125 | 4.3935212080051 | 0.00340865777615044 | 0.100577114249108 | -1.44952366187326 |
| **Nrxn2** | -0.760264847500004 | 24.55428514875 | -4.39303436519864 | 0.00341063031363524 | 0.100577114249108 | -1.45010316435365 |
| **Ca8** | 0.721451639999994 | 21.542013855 | 4.39072499286877 | 0.00342000437997981 | 0.100577114249108 | -1.45285261408399 |
| **Arpc2** | 0.786335315000002 | 22.3066835275 | 4.38894051471081 | 0.00342726732696828 | 0.100577114249108 | -1.45497775577449 |
| **Ppp2r1a** | -0.76693613749999 | 28.85718693375 | -4.35923425390348 | 0.00355071131005239 | 0.102652789663778 | -1.49043324738405 |
| **Anxa5** | 1.086172925 | 22.26273234 | 4.34122379467673 | 0.00362794025630156 | 0.102652789663778 | -1.51200119540041 |
| **Atp5me** | 0.744227635000001 | 20.9637671825 | 4.34016529802455 | 0.00363253646254521 | 0.102652789663778 | -1.51327045620627 |
| **Slc39a12** | 0.603970200000006 | 29.48093691 | 4.31288058296702 | 0.0037532592622055 | 0.103257731372039 | -1.54605258870414 |
| **Dpm3** | 0.373966932500004 | 27.53102378125 | 4.2941074556842 | 0.00383889166388028 | 0.103257731372039 | -1.56868036385229 |

| | | | | | | |
|---|---|---|---|---|---|---|
| **Slc38a3** | 1.09626222 | 23.824499845 | 4.29395118420294 | 0.00383961347275248 | 0.103257731372039 | -1.56886896904117 |
| **Kcnab2** | -0.683648715000004 | 23.4307202475 | -4.287996285188 | 0.00386723068111242 | 0.103257731372039 | -1.57605901539799 |
| **Marcksl1** | -0.971598354999998 | 22.632484415 | -4.28536676518762 | 0.00387949536828106 | 0.103257731372039 | -1.57923582515442 |
| **Atp6v1e1** | 0.6790977 | 26.008528775 | 4.26967857267428 | 0.00395356624991399 | 0.104019691333944 | -1.59821320704053 |
| **Ddt** | -0.992074944999999 | 20.075088785 | -4.22646887777477 | 0.00416576595302011 | 0.104166921985806 | -1.65069425787346 |
| **Hpcal4** | -0.61088806250001 | 25.45255508625 | -4.21788418458875 | 0.00420940231736178 | 0.104166921985806 | -1.66115794019477 |
| **Pex5l** | -0.985030869999996 | 22.03392526 | -4.20535283118127 | 0.00427400656558109 | 0.104166921985806 | -1.67645410994978 |
| **Hspa4l** | -0.613481742499999 | 26.45014936875 | -4.20416866217948 | 0.00428016770152229 | 0.104166921985806 | -1.67790089387971 |
| **Anxa6** | -1.14089919 | 21.4818465975 | -4.20277652441286 | 0.00428742339677737 | 0.104166921985806 | -1.67960206603034 |
| **Ddah2** | 0.485959570000002 | 23.8901462425 | 4.18479343990351 | 0.00438237748929245 | 0.104166921985806 | -1.70160606368563 |
| **Pfn2** | 0.502208579999991 | 26.95471774 | 4.16636097543639 | 0.00448211351654922 | 0.104166921985806 | -1.72421560638267 |
| **Actr1b** | 0.415623777499995 | 25.63355239875 | 4.16407778016177 | 0.00449464058800834 | 0.104166921985806 | -1.72702012915291 |
| **Mpp3** | -0.855597937500001 | 25.50381851375 | -4.15218227720072 | 0.00456053347661319 | 0.104166921985806 | -1.74164573085909 |
| **Gda** | 0.448858677499999 | 25.77779856125 | 4.14752161120779 | 0.00458663962056413 | 0.104166921985806 | -1.74738243956212 |
| **Nectin1** | -0.664498099999999 | 29.2179200525 | -4.13799853322767 | 0.00464049475323075 | 0.104166921985806 | -1.75911535671288 |
| **Acyp1** | -0.5927846075 | 26.16669495625 | -4.13618930677531 | 0.00465080477419308 | 0.104166921985806 | -1.76134611143015 |
| **Praf2** | 0.537646465000002 | 28.251948015 | 4.13329659800508 | 0.0046673415023582 | 0.104166921985806 | -1.76491391137558 |
| **Nme1** | -0.370213132500002 | 29.47153226875 | -4.13293363437368 | 0.00466942101749524 | 0.104166921985806 | -1.76536167998082 |
| **Cyrib** | -0.461952225000001 | 31.0231673575 | -4.12283049281123 | 0.00472771560672074 | 0.104166921985806 | -1.77783411648119 |
| **Napa** | 1.048673575 | 20.730416975 | 4.11547638689495 | 0.00477065175466584 | 0.104166921985806 | -1.78692344168214 |
| **Cyc1** | 1.01787080250001 | 24.14743674125 | 4.11519766279553 | 0.00477228746811587 | 0.104166921985806 | -1.7872681070256 |

| | | | | | | |
|---|---|---|---|---|---|---|
| **Zfyve28** | 0.7400097475 | 25.83016737875 | 4.0903790636018 | 0.00492043970657293 | 0.104166921985806 | -1.81800971902121 |
| **Ppa1** | 0.44237444749999 | 27.62125127625 | 4.09021539213734 | 0.00492143335753928 | 0.104166921985806 | -1.81821278797365 |
| **Nbea** | -0.591586487500003 | 29.31978755625 | -4.08953095637674 | 0.00492559096374724 | 0.104166921985806 | -1.81906202260353 |
| **Gpi** | 1.3158734425 | 22.69338960875 | 4.06950119894965 | 0.00504898781836285 | 0.104166921985806 | -1.8439486961433 |
| **Otub1** | 0.336808545000004 | 25.052823935 | 4.06890041974102 | 0.00505274112540502 | 0.104166921985806 | -1.84469617442232 |
| **Gnai2** | -0.890320577500002 | 27.70413082375 | -4.06695871404202 | 0.00506489271531167 | 0.104166921985806 | -1.84711241402447 |
| **Ndufab1** | 0.559968927500002 | 26.05460147375 | 4.05923495455869 | 0.00511354844236453 | 0.104166921985806 | -1.85672991632473 |
| **Calb1** | -0.8471171775 | 21.25202036875 | -4.04771045226911 | 0.00518710423460336 | 0.104166921985806 | -1.87109824850024 |
| **Aldh6a1** | 0.4333836375 | 27.99998636125 | 4.0471095122671 | 0.00519097152097952 | 0.104166921985806 | -1.87184807633158 |
| **Uqcrb** | 0.291228747499989 | 27.69882043375 | 4.04296852545596 | 0.00521770665629623 | 0.104166921985806 | -1.8770166361475 |
| **Trpv2** | -0.621403244999989 | 27.8360097375 | -4.03979302924782 | 0.00523831081303015 | 0.104166921985806 | -1.88098202560252 |
| **Enoph1** | -0.335808525000004 | 29.19638333 | -4.03357999473728 | 0.00527888289661578 | 0.104166921985806 | -1.88874530726586 |
| **Cntnap1** | 0.498257872500005 | 27.04447436875 | 4.02316556825622 | 0.00534766681132203 | 0.104622301975352 | -1.90177246353635 |
| **Pgam1** | 1.0816590175 | 22.76144653625 | 4.00743090091814 | 0.00545346264427972 | 0.105284712867177 | -1.92148820946651 |
| **Lmbrd2** | -0.612732852499999 | 27.49056443125 | -3.99348584895813 | 0.00554914746986021 | 0.105284712867177 | -1.93899531547513 |
| **Slc12a2** | 0.897477067499995 | 21.64025758875 | 3.98979226137713 | 0.00557479875177389 | 0.105284712867177 | -1.94363768304404 |
| **Oxct1** | -0.779803189999999 | 27.7823803375 | -3.97719651253924 | 0.00566325677665632 | 0.105284712867177 | -1.95948562564599 |
| **Apmap** | -0.616755397499993 | 29.30983435875 | -3.97377392169266 | 0.00568755829297861 | 0.105284712867177 | -1.96379637898458 |
| **Prkaca** | -0.461883332500005 | 28.45814157625 | -3.9651062981891 | 0.00574961431506199 | 0.105284712867177 | -1.97472176144401 |
| **Nme2** | 0.447631262500003 | 23.19465888375 | 3.96442765277861 | 0.00575450433909189 | 0.105284712867177 | -1.97557769648291 |
| **Ugp2** | -0.694879337499998 | 27.21450942875 | -3.96282107416662 | 0.00576609883181629 | 0.105284712867177 | -1.97760427622498 |

| | | | | | | |
|---|---|---|---|---|---|---|
| **Capza2** | 0.66116126 | 24.742097635 | 3.95079635788368 | 0.0058536974502533 | 0.105284712867177 | -1.99278585110791 |
| **Slc16a1** | 0.460525537500008 | 27.27746917625 | 3.95010008395928 | 0.00585881421643081 | 0.105284712867177 | -1.99366563589743 |
| **Pak1** | 0.351267907500013 | 24.82719837875 | 3.9462115276512 | 0.00588748066710295 | 0.105284712867177 | -1.99858050822774 |
| **Sgta** | 0.320733882500001 | 24.68939024125 | 3.93365249138091 | 0.00598112096576616 | 0.106130123183246 | -2.01447098636327 |
| **Aldoc** | 0.486269787499985 | 28.64565442875 | 3.92244759898251 | 0.00606604339439918 | 0.106507887515503 | -2.02866962020767 |
| **Ephb3** | 0.847953282500004 | 21.36799596375 | 3.91493612431288 | 0.00612371241836885 | 0.106507887515503 | -2.03819935766634 |
| **Ndufs4** | 0.832269597500005 | 23.52379353625 | 3.9122923186326 | 0.00614415278085394 | 0.106507887515503 | -2.04155569128951 |
| **Uqcrfs1** | -0.82704051499999 | 24.2077067775 | -3.90589512746334 | 0.0061939219592623 | 0.106507887515503 | -2.04968163293788 |
| **Idh1** | -0.731413345000004 | 29.997108715 | -3.89538065769131 | 0.00627668473977961 | 0.106507887515503 | -2.06305178080375 |
| **Ca2** | 0.450737255000004 | 26.91501617 | 3.89476195642785 | 0.00628159231742808 | 0.106507887515503 | -2.06383907132932 |
| **Lactb** | 0.516283964999992 | 27.9357334325 | 3.86971296074513 | 0.00648385373059703 | 0.108959889954564 | -2.0957651562071 |
| **Ociad2** | 0.471916327500001 | 29.65020519625 | 3.84931587837116 | 0.00665382024180781 | 0.108959889954564 | -2.12183623003933 |
| **Rab5b** | -0.667055802499991 | 28.27688975625 | -3.84922850064803 | 0.00665455875233998 | 0.108959889954564 | -2.12194805680801 |
| **Nrcam** | -0.632469222500003 | 28.65583676875 | -3.84912030343619 | 0.00665547335199969 | 0.108959889954564 | -2.12208653022282 |
| **Hmox2** | 0.594805310000002 | 25.4293949425 | 3.8480871682811 | 0.00666421345287851 | 0.108959889954564 | -2.12340885545628 |
| **Adsl** | -0.54273502249999 | 29.72957377625 | -3.83947614458935 | 0.00673755108480145 | 0.109046452162365 | -2.13443683733936 |
| **Ckb** | 0.657808707499996 | 24.46120149875 | 3.82998717573604 | 0.00681938944384911 | 0.109046452162365 | -2.14660282639357 |
| **Ndufb1** | 0.350231052499996 | 27.88845400875 | 3.82574139064267 | 0.00685635893762074 | 0.109046452162365 | -2.15205105277757 |
| **Osbpl1a** | 0.672457459999986 | 25.654582185 | 3.81595151243153 | 0.00694244042188731 | 0.109046452162365 | -2.16462438270632 |
| **Lrpap1** | -0.813904702499993 | 25.51372647125 | -3.81566042997324 | 0.00694501789942396 | 0.109046452162365 | -2.16499845760153 |
| **Ganab** | 0.438939547499999 | 20.80617356125 | 3.81326661418299 | 0.00696625437076245 | 0.109046452162365 | -2.16807529781995 |

| | | | | | | |
|---|---|---|---|---|---|---|
| **Mdh1** | -0.542331730000004 | 29.8590478425 | -3.80914523629086 | 0.00700298316639043 | 0.109046452162365 | -2.17337474389773 |
| **Rad23b** | 0.614791952499999 | 24.72248287375 | 3.79502926323176 | 0.00713039382524073 | 0.110280212607946 | -2.19154597448761 |
| **Neo1** | 0.454400907500002 | 27.57368483375 | 3.78396839051558 | 0.00723199880465859 | 0.110853359028242 | -2.2058063444986 |
| **Svip** | 0.525228107499998 | 30.26499031125 | 3.77144744089808 | 0.00734892783011849 | 0.110853359028242 | -2.22197230587803 |
| **Adgrg1** | 0.496177305 | 28.1796205225 | 3.76517269749242 | 0.0074083006427985 | 0.110853359028242 | -2.23008294187643 |
| **Clta** | 0.399551322499988 | 24.43536557375 | 3.76215828013883 | 0.00743700965369275 | 0.110853359028242 | -2.23398151934887 |
| **Ehd3** | -0.555030907500008 | 29.46855049875 | -3.75970136706565 | 0.00746049887606508 | 0.110853359028242 | -2.23716012028194 |
| **Gdi1** | -0.532695565000012 | 24.414487655 | -3.75404645291557 | 0.00751487069411486 | 0.110853359028242 | -2.24447967395987 |
| **Nnt** | -0.686743402500003 | 27.15789734125 | -3.75080422049449 | 0.00754623958621193 | 0.110853359028242 | -2.24867857193077 |
| **Cox6c** | -0.298941017499995 | 24.98146457625 | -3.74893351273628 | 0.00756440383499871 | 0.110853359028242 | -2.2511020026879 |
| **Mcam** | 0.517889170000004 | 26.506671575 | 3.72781865149225 | 0.00777276628412587 | 0.110853359028242 | -2.27849321127625 |
| **Vdac1** | -1.2054104425 | 21.53703067625 | -3.7136008989184 | 0.00791659051594752 | 0.110853359028242 | -2.29697606681179 |
| **Tppp3** | -0.574973607500009 | 27.94982562125 | -3.71233299078483 | 0.00792955666684864 | 0.110853359028242 | -2.29862584329949 |
| **Atp5pf** | 1.148758425 | 22.39202588 | 3.7109103537024 | 0.00794413270661284 | 0.110853359028242 | -2.30047724502428 |
| **Ptprn2** | 0.420000625000004 | 25.235245415 | 3.70857190385775 | 0.00796815547273889 | 0.110853359028242 | -2.30352115142684 |
| **Itfg1** | 0.54199775499999 | 28.7035595475 | 3.7073141511797 | 0.00798110903610103 | 0.110853359028242 | -2.30515868751315 |
| **Rev1** | -0.923293367499994 | 21.52887330625 | -3.6998518050154 | 0.00805843717402896 | 0.110853359028242 | -2.31487932504318 |
| **Endod1** | -0.544224629999988 | 28.6002902475 | -3.69889876309814 | 0.00806837167375683 | 0.110853359028242 | -2.31612139761051 |
| **Aqp4** | -0.537028702500006 | 25.16643095625 | -3.69421101181302 | 0.00811743141027326 | 0.110853359028242 | -2.3222328413094 |
| **Gde1** | 0.426710052500006 | 31.76706124625 | 3.69234779586992 | 0.00813702112029198 | 0.110853359028242 | -2.32466286123571 |
| **Pcyox1l** | -0.334278150000003 | 30.2671799325 | -3.68762254886548 | 0.00818693335347989 | 0.110853359028242 | -2.33082794897802 |

| | | | | | | |
|---|---|---|---|---|---|---|
| **Adam11** | -0.63104516500001 | 27.2345267225 | -3.68638589657192 | 0.00820005097064001 | 0.110853359028242 | -2.33244198900656 |
| **Vamp1** | -0.294305207499995 | 28.85871236625 | -3.68542145071973 | 0.00821029708777308 | 0.110853359028242 | -2.33370091613139 |
| **Mag** | -0.565855042500001 | 26.52314362625 | -3.68248761025979 | 0.00824155157465412 | 0.110853359028242 | -2.33753144228374 |
| **Dpysl3** | 0.483260447500001 | 27.28163764625 | 3.67877273973364 | 0.0082813125355305 | 0.110853359028242 | -2.3423835968676 |
| **Prps1** | 1.2298317075 | 23.53825998125 | 3.66594781375325 | 0.00842019395358724 | 0.111586780148226 | -2.35915097530441 |
| **Lamp5** | 0.755488144999998 | 24.625588815 | 3.65559387127827 | 0.0085341676623694 | 0.111586780148226 | -2.37270603640187 |
| **Sar1a** | -0.537247872499997 | 29.33527535375 | -3.65366828861496 | 0.0085555486502 | 0.111586780148226 | -2.37522874414161 |
| **Rap1gds1** | -0.608190292499998 | 29.49234231375 | -3.65135871175754 | 0.00858127031294595 | 0.111586780148226 | -2.37825526427964 |
| **Psmd6** | 0.523508749999998 | 28.7148570625 | 3.64633826336706 | 0.00863747365131617 | 0.111586780148226 | -2.38483695588522 |
| **Cadps** | 0.655890157499993 | 26.20932980375 | 3.64344586633287 | 0.00867003549474529 | 0.111586780148226 | -2.38863055227583 |
| **Cab39** | 0.983733409999992 | 23.249509145 | 3.64279804267322 | 0.00867734681799224 | 0.111586780148226 | -2.38948039504623 |
| **Tcp1** | 0.419605047500006 | 26.31320681125 | 3.63314712461037 | 0.0087870650025379 | 0.112298285635653 | -2.40214838121383 |
| **Kcnc3** | -0.621909797499999 | 30.52923275125 | -3.62933701222493 | 0.00883079572495304 | 0.112298285635653 | -2.40715347408207 |
| **Etfdh** | -0.756110872499995 | 26.23680744875 | -3.62051163920755 | 0.00893300128293567 | 0.112530020197376 | -2.41875516417589 |
| **Tmem163** | -0.435044592500002 | 22.39341961625 | -3.61928220123484 | 0.00894734105544886 | 0.112530020197376 | -2.4203722907269 |
| **Cox5b** | 0.474020102500003 | 25.27401257375 | 3.60399860062009 | 0.00912770914736752 | 0.114030855667173 | -2.44049427714668 |
| **Npepps** | -0.641213529999995 | 27.69061422 | -3.60077061515414 | 0.00916630731444292 | 0.114030855667173 | -2.44474861918779 |
| **Ndufb8** | 0.500022727500003 | 25.94002147625 | 3.58858730131085 | 0.00931359378730746 | 0.115012828347377 | -2.46081965632292 |
| **Pura** | 1.12585949999999 | 19.675024415 | 3.58481754082738 | 0.00935968663113295 | 0.115012828347377 | -2.46579681994249 |
| **Agpat3** | -0.838143200000001 | 23.87753972 | -3.58186338473757 | 0.00939598029749216 | 0.115012828347377 | -2.46969862322561 |
| **Rufy3** | -0.57525571 | 26.0731898975 | -3.56745547919685 | 0.00957519519484453 | 0.116583094686166 | -2.48874683859919 |

| | | | | | | |
|---|---|---|---|---|---|---|
| **Psat1** | 0.678438872500003 | 24.94294215375 | 3.55538918053074 | 0.00972814116644819 | 0.117029076941751 | -2.50472280434342 |
| **Lasp1** | -0.457241607499991 | 31.16719080125 | -3.55090376068739 | 0.00978567048233033 | 0.117029076941751 | -2.51066701236191 |
| **Gdi2** | -1.0035695325 | 23.40592389875 | -3.54314677443107 | 0.00988603384059266 | 0.117029076941751 | -2.52095373510493 |
| **Tnc** | 0.575354767500009 | 24.21827011125 | 3.542035007155 | 0.00990050968162517 | 0.117029076941751 | -2.52242879530644 |
| **Atp5pd** | -0.463089069999992 | 28.625317685 | -3.53898489095418 | 0.00994034217608946 | 0.117029076941751 | -2.52647652465882 |
| **Cntnap2** | -0.656915095000006 | 27.8897705475 | -3.53745335727589 | 0.00996040854271794 | 0.117029076941751 | -2.5285094936458 |
| **Hars1** | 0.441647322500003 | 26.51434273625 | 3.53674444659755 | 0.00996971166607312 | 0.117029076941751 | -2.52945062238011 |
| **Emc1** | 0.66214944 | 22.9813816575 | 3.52968085165433 | 0.0100629247888635 | 0.117520585927084 | -2.5388320292856 |
| **Rab10** | -0.494091702499993 | 21.06477483875 | -3.5120875396523 | 0.0102992260877229 | 0.119153639477846 | -2.56222972516271 |
| **Itgav** | 1.2692484275 | 22.92783851625 | 3.51006347634574 | 0.0103267955484692 | 0.119153639477846 | -2.56492443120219 |
| **Cacybp** | 0.307950230000003 | 26.4703691875 | 3.50383217445972 | 0.0104121746958317 | 0.119153639477846 | -2.57322407750514 |
| **Gpr37** | 0.462358705 | 24.88784006 | 3.49856403685293 | 0.0104849539424821 | 0.119153639477846 | -2.5802452034367 |
| **Ppp1r7** | -0.501948887499999 | 23.88746739875 | -3.49841196153715 | 0.0104870630322996 | 0.119153639477846 | -2.58044794128567 |
| **Ndufb9** | 1.48077869 | 21.5462390525 | 3.48933348798359 | 0.0106138066806207 | 0.119153639477846 | -2.59255681507005 |
| **Psma4** | 0.562604257500006 | 28.16897963125 | 3.48748301940091 | 0.0106398439794518 | 0.119153639477846 | -2.59502641395992 |
| **St13** | 0.466933049999998 | 23.4505128675 | 3.48196373447652 | 0.0107179158056301 | 0.119153639477846 | -2.60239523746862 |
| **Pllp** | -0.362950184999995 | 28.1102627225 | -3.47850747031586 | 0.0107671214841106 | 0.119153639477846 | -2.60701191622068 |
| **Cend1** | 1.65828205750001 | 23.82968673125 | 3.47833022783283 | 0.0107696514136929 | 0.119153639477846 | -2.60724871222182 |
| **Nipsnap1** | -1.2213580125 | 24.18996985375 | -3.4755885755278 | 0.0108088674226558 | 0.119153639477846 | -2.61091212761688 |
| **Gosr2** | -0.465841007499996 | 30.08944368375 | -3.47134605501997 | 0.0108698566995043 | 0.119153639477846 | -2.61658311178877 |
| **Mcee** | -0.4346587625 | 23.44460618375 | -3.47067957401134 | 0.010879471669231 | 0.119153639477846 | -2.61747422943615 |

| | | | | | | |
|---|---|---|---|---|---|---|
| **Mmut** | 0.371283199999993 | 29.0564284325 | 3.46268276909478 | 0.0109955583214239 | 0.119817668298873 | -2.62817123770465 |
| **Them4** | 1.76207438 | 22.7022353825 | 3.45811928391915 | 0.0110624052534264 | 0.119817668298873 | -2.63427966773527 |
| **Prrt1** | 0.699891542499994 | 23.95773409625 | 3.45576002505795 | 0.0110971366008567 | 0.119817668298873 | -2.6374387894325 |
| **Btbd17** | 0.269233772499991 | 26.34721179375 | 3.44679695936036 | 0.011230164440829 | 0.120456295471087 | -2.64944772883662 |
| **Uqcrc1** | 0.438888120000009 | 25.515432575 | 3.44470022119428 | 0.0112615322109273 | 0.120456295471087 | -2.65225861288222 |
| **Scarb2** | -0.488455585000011 | 26.6532283125 | -3.43110489867342 | 0.0114672362792605 | 0.122086064387103 | -2.67049934254566 |
| **Ndufb11** | 0.388722295000001 | 28.43562757 | 3.42562497956328 | 0.0115512964213066 | 0.122411655131347 | -2.67785895960495 |
| **Tollip** | 0.329859492499992 | 25.54759480875 | 3.41967202258447 | 0.011643368608013 | 0.12281875918775 | -2.68585857818573 |
| **Arhgdia** | 0.575725217499997 | 27.41699541625 | 3.41108582114846 | 0.0117775686565838 | 0.123100336055741 | -2.69740536517967 |
| **Ndufa8** | 0.386903875000005 | 20.781180735 | 3.41108251710068 | 0.0117776206187013 | 0.123100336055741 | -2.69740981043745 |
| **Psma3** | -0.427072347499994 | 29.11069403875 | -3.40442158004249 | 0.0118828814316135 | 0.123635979986196 | -2.7063744606251 |
| **Eps15l1** | -0.628669164999994 | 26.130116265 | -3.39426399973628 | 0.0120453609531349 | 0.124759417292877 | -2.72005675426069 |
| **Pebp1** | -0.330537695000004 | 25.0099952675 | -3.38323439234858 | 0.0122245117366299 | 0.126044627770927 | -2.7349295563211 |
| **Ube2n** | 0.781441954999998 | 21.3473460475 | 3.37787142041461 | 0.0123126590165311 | 0.126384199501524 | -2.74216716762355 |
| **Gstm5** | 0.3630856825 | 27.92782083875 | 3.36636048439456 | 0.0125041820796032 | 0.127777110625945 | -2.75771484830195 |
| **Adam10** | 0.493673870000002 | 25.20601587 | 3.35254780955793 | 0.012738257163611 | 0.129590536211136 | -2.77639485816448 |
| **Eif5a** | -0.62496097 | 28.3427822575 | -3.34785536017004 | 0.0128188497489737 | 0.129833394138941 | -2.78274661583098 |
| **Sars1** | -0.395718782500012 | 33.83259710125 | -3.33543283398775 | 0.0130348734856573 | 0.131439759509557 | -2.79957595260918 |
| **Mal2** | 0.374988345000006 | 19.4164407275 | 3.32670542869275 | 0.0131889867006371 | 0.132410484902449 | -2.81141148283497 |
| **Grb2** | -0.396519775000009 | 27.0508430975 | -3.3147048567401 | 0.0134041150479296 | 0.133982617225812 | -2.82770210471811 |
| **Lrp1** | 1.117698145 | 20.96652885 | 3.3049178858413 | 0.0135823610235957 | 0.135174019056567 | -2.84100162170544 |

| | | | | | | |
|---|---|---|---|---|---|---|
| **Me1** | -0.613463767500001 | 28.18775739125 | -3.29268445326368 | 0.0138087621043127 | 0.136832279033644 | -2.8576429777943 |
| **Vmp1** | -0.496890445000005 | 28.13694133 | -3.28876784185416 | 0.0138821020678735 | 0.136966084626563 | -2.86297486233339 |
| **Cyb5a** | 0.414427679999992 | 26.14599261 | 3.28036975443133 | 0.0140407767835989 | 0.137937073208832 | -2.87441419261849 |
| **Atp1b3** | -0.567509887500005 | 28.63162177375 | -3.26550326057099 | 0.0143264730884806 | 0.140142294442445 | -2.89468623926934 |
| **Tpi1** | -0.583515527499998 | 29.31173578125 | -3.25033036861039 | 0.0146245169472683 | 0.140625181861578 | -2.91540465225105 |
| **Psmb1** | -0.544769944999999 | 29.5994370175 | -3.24912496303416 | 0.0146484792128124 | 0.140625181861578 | -2.91705184762955 |
| **Kpnb1** | 0.572490320000014 | 26.1796637725 | 3.24668491684203 | 0.0146971142153298 | 0.140625181861578 | -2.92038673880746 |
| **Creld1** | -0.534087357499995 | 27.18688762625 | -3.24616999732287 | 0.0147073997575491 | 0.140625181861578 | -2.92109059027994 |
| **Itgb1** | 1.4794967225 | 21.60008954375 | 3.24446543382653 | 0.0147415037401948 | 0.140625181861578 | -2.92342081817502 |
| **Etfb** | 0.823793632500006 | 23.72733609875 | 3.24431842712405 | 0.0147444489500999 | 0.140625181861578 | -2.92362180086315 |
| **Fkbp2** | 0.290396117500006 | 23.98404385375 | 3.22931919613945 | 0.0150483015312726 | 0.141957821327128 | -2.94414221252467 |
| **Sh3bgrl3** | 0.936203017499988 | 24.60873337125 | 3.22916610860788 | 0.0150514372328745 | 0.141957821327128 | -2.94435179318649 |
| **Alb** | -0.614006404999991 | 26.0484596475 | -3.22676134190909 | 0.0151007864545189 | 0.141957821327128 | -2.94764435374462 |
| **Aars1** | -0.622611082500001 | 22.50205211125 | -3.22094029327217 | 0.015220963686049 | 0.141957821327128 | -2.95561733308554 |
| **Timm9** | -0.634639019999995 | 29.9644668175 | -3.21947760004243 | 0.0152513226391727 | 0.141957821327128 | -2.95762140212028 |
| **Psmb6** | -0.261535589999998 | 28.2956530975 | -3.21791161873531 | 0.0152838974887238 | 0.141957821327128 | -2.95976727538624 |
| **Ppid** | -0.262415300000001 | 27.873851085 | -3.21626180558531 | 0.0153182970152034 | 0.141957821327128 | -2.96202834444441 |
| **Wdr1** | 0.481102212499998 | 25.39419772375 | 3.2094277473046 | 0.0154616789692592 | 0.142708803067073 | -2.9713979151807 |
| **Trpc3** | -0.611748712500003 | 28.75230353625 | -3.20113951907521 | 0.0156375062940583 | 0.143100225698563 | -2.98276868624888 |
| **Chl1** | 0.444806354999997 | 24.6238518075 | 3.20018827221582 | 0.0156578230273739 | 0.143100225698563 | -2.98407424388728 |
| **Pea15** | -0.626577997500007 | 27.59029547875 | -3.19476602909153 | 0.0157741735791374 | 0.143100225698563 | -2.99151816404108 |

| | | | | | | |
|---|---|---|---|---|---|---|
| **Mtnd5** | 0.477926989999993 | 23.64234131 | 3.19448644382862 | 0.0157801980217526 | 0.143100225698563 | -2.99190208695611 |
| **Ywhaq** | 0.521203222499999 | 24.90683925875 | 3.1913684721463 | 0.015847550939079 | 0.143100225698563 | -2.99618427302542 |
| **Basp1** | 0.656098160000003 | 23.8599122875 | 3.18218300885189 | 0.0160477697200176 | 0.143100225698563 | -3.0088061478575 |
| **Snca** | -0.3645659775 | 28.43146218625 | -3.1818397731863 | 0.0160553037436467 | 0.143100225698563 | -3.00927798504918 |
| **Slc8a1** | 0.500593754999997 | 24.77114536 | 3.18024837124525 | 0.0160902846219116 | 0.143100225698563 | -3.01146582449184 |
| **Psma7** | -0.432001202499993 | 31.01296884375 | -3.17736611521189 | 0.0161538482332881 | 0.143100225698563 | -3.01542906823655 |
| **Inpp5a** | 0.350605287499995 | 23.31843069875 | 3.1757939785722 | 0.0161886326771695 | 0.143100225698563 | -3.01759124232398 |
| **Lamp1** | 0.2367984475 | 26.61761686625 | 3.17565242115919 | 0.0161917686570239 | 0.143100225698563 | -3.01778594169159 |
| **Fundc2** | -1.16041125 | 26.214708315 | -3.16241454838299 | 0.0164879307984059 | 0.14424435015871 | -3.03600371551846 |
| **Csnk2a1** | -0.534236067499993 | 27.74727467125 | -3.1612310565595 | 0.016514689626491 | 0.14424435015871 | -3.03763340671743 |
| **Mtfp1** | -1.0375294375 | 21.74474249875 | -3.16110108989052 | 0.0165176310163508 | 0.14424435015871 | -3.03781238316409 |
| **Ndufa12** | -0.292151562499999 | 28.76500259625 | -3.1586463732429 | 0.0165732914337007 | 0.14424435015871 | -3.04119312576689 |
| **Krt222** | 0.443527367499993 | 25.88916964625 | 3.14541087256288 | 0.0168768882735813 | 0.145004485824718 | -3.05943350985258 |
| **Idi1** | 0.285279875000004 | 26.2606819525 | 3.14518042075187 | 0.0168822268706611 | 0.145004485824718 | -3.05975128149292 |
| **Nsfl1c** | -0.307889039999999 | 27.27414258 | -3.14217421104909 | 0.0169520340029143 | 0.145004485824718 | -3.06389711619991 |
| **Zmpste24** | -0.815761977499999 | 24.32860433125 | -3.13954357161216 | 0.0170133736507864 | 0.145004485824718 | -3.06752584338225 |
| **Crym** | 0.515466470000007 | 29.9293482775 | 3.13924902341628 | 0.0170202565276157 | 0.145004485824718 | -3.06793219434151 |
| **Rexo2** | -0.620973077500004 | 22.38333670125 | -3.13837408127254 | 0.0170407193913714 | 0.145004485824718 | -3.06913929918586 |
| **Myo6** | 1.54710877 | 23.2549639225 | 3.13472681480916 | 0.0171263046843032 | 0.145193005268037 | -3.07417213997701 |
| **Bcap31** | 0.694228542499989 | 22.87986968125 | 3.1296740682496 | 0.0172456315792472 | 0.145665131678586 | -3.08114685837149 |
| **Nsdhl** | -0.73390700500001 | 25.5454747 | -3.12655982791281 | 0.0173196212984291 | 0.14568756497249 | -3.08544711861179 |

| | | | | | | |
|---|---|---|---|---|---|---|
| **Ctsb** | 0.334767175000014 | 25.5851764125 | 3.11967064421324 | 0.0174845078148181 | 0.14568756497249 | -3.09496379144105 |
| **Gnas** | -0.504398730000005 | 29.269670745 | -3.11935059881821 | 0.0174922085342635 | 0.14568756497249 | -3.09540602777443 |
| **Mog** | 0.360550484999994 | 29.5280897975 | 3.1189075537983 | 0.0175028747782589 | 0.14568756497249 | -3.09601824273857 |
| **Prrt2** | -0.5878173125 | 21.40244000625 | -3.11566848489369 | 0.0175810666077399 | 0.145808193714191 | -3.1004947581021 |
| **Slc9a6** | -1.20193702249999 | 25.89196551875 | -3.10863101383134 | 0.0177522435622233 | 0.146233637800113 | -3.11022478188218 |
| **Pafah1b2** | 0.4136696375 | 25.39965832625 | 3.103358233152 | 0.017881664610427 | 0.146233637800113 | -3.11751848762425 |
| **Uggt1** | 0.628032017500001 | 23.94954548125 | 3.10274365091565 | 0.0178968151337431 | 0.146233637800113 | -3.11836882069706 |
| **Mtatp8** | -0.901146050000001 | 24.4371739225 | -3.10102265201523 | 0.0179393138328222 | 0.146233637800113 | -3.1207502046913 |
| **Cds2** | 0.354283107500002 | 25.08088842625 | 3.09942794533953 | 0.0179787901687009 | 0.146233637800113 | -3.12295712181273 |
| **Ndufa2** | 0.387563992499999 | 21.29066818125 | 3.09755885651616 | 0.0180251769769225 | 0.146233637800113 | -3.12554410691291 |
| **Uqcrc2** | -0.598099487500008 | 30.19557016375 | -3.09404225390079 | 0.0181127987328928 | 0.146233637800113 | -3.13041241913612 |
| **Mtcl3** | 0.590807377499999 | 22.81571299375 | 3.09249403857011 | 0.0181515192272692 | 0.146233637800113 | -3.13255615844649 |
| **Gclm** | 0.240216070000002 | 23.06371822 | 3.09026853226168 | 0.0182073336710494 | 0.146233637800113 | -3.13563816067118 |
| **Camkv** | 1.12941544249999 | 23.84459873875 | 3.08245981132847 | 0.018404627257598 | 0.14647241584617 | -3.14645628904736 |
| **Lonp1** | -0.549497825 | 25.609092155 | -3.08180929478911 | 0.0184211658408511 | 0.14647241584617 | -3.14735780140694 |
| **Hspa9** | -0.815229595000002 | 26.54083463 | -3.08150008594032 | 0.0184290326621656 | 0.14647241584617 | -3.14778633138706 |
| **Dcps** | 1.18926942 | 20.97860217 | 3.07603147497741 | 0.0185687584500841 | 0.147072277135787 | -3.15536690610911 |
| **Eno2** | -0.856278220000004 | 20.984318345 | -3.06287906900395 | 0.0189094629333651 | 0.149254347084389 | -3.17361160607331 |
| **Hsph1** | -0.342083729999992 | 34.010497785 | -3.05854948053841 | 0.0190230732775481 | 0.149635102172878 | -3.179621448383 |
| **Hycc2** | 2.2864454875 | 21.76446824875 | 3.05168617509688 | 0.0192046648295536 | 0.150125941946504 | -3.1891522708567 |
| **Ndufs6** | 0.322317140000003 | 20.8878378175 | 3.05123580909531 | 0.0192166452557123 | 0.150125941946504 | -3.18977784683862 |

| | | | | | | |
|---|---|---|---|---|---|---|
| **Dlg2** | -0.582520749999993 | 27.1814717025 | -3.04635192351097 | 0.0193470774060838 | 0.150630816947367 | -3.19656308576908 |
| **Pgrmc2** | 0.597992955000002 | 25.1581211025 | 3.03928208466016 | 0.0195375647149174 | 0.150982807269481 | -3.20638959055812 |
| **Ndufs2** | -0.620549822500003 | 28.19007911375 | -3.03869038051566 | 0.0195535978528024 | 0.150982807269481 | -3.20721224074721 |
| **Ptpa** | -0.648183102500003 | 24.95525369875 | -3.03734270298655 | 0.0195901676535762 | 0.150982807269481 | -3.20908605740477 |
| **Vapa** | -0.483016682500001 | 23.90390841875 | -3.03154485744315 | 0.0197483284529319 | 0.151691019559601 | -3.21714948714467 |
| **Exoc2** | 0.641714352499999 | 23.68825983625 | 3.02785880008154 | 0.019849588755701 | 0.151958891845483 | -3.22227766708183 |
| **Ran** | 0.253839092500005 | 27.12176840875 | 3.0239119251745 | 0.0199586278902965 | 0.152157174918399 | -3.22777020323465 |
| **Cd9** | 0.9720185075 | 22.69368383375 | 3.02211667316951 | 0.0200084358455387 | 0.152157174918399 | -3.23026901571508 |
| **Akr1a1** | 0.358720989999998 | 25.918906205 | 3.01386496501902 | 0.0202390830496106 | 0.153401526823042 | -3.24175865208853 |
| **Gstk1** | 0.505558042499992 | 27.15087882875 | 3.00084536797975 | 0.0206087781519408 | 0.155688096335949 | -3.25990055678398 |
| **Taldo1** | -0.558009082499996 | 27.73210377375 | -2.99551678288048 | 0.0207621526179179 | 0.156330813626362 | -3.26733027332911 |
| **Ermp1** | 0.323635727499997 | 30.02798154625 | 2.9764532750718 | 0.0213208878594672 | 0.15991055187751 | -3.29393275341658 |
| **Syt2** | -0.528017974999997 | 29.186108295 | -2.97157175467472 | 0.0214665186352137 | 0.15991055187751 | -3.30075019744776 |
| **Tnr** | -0.421896919999998 | 25.065273565 | -2.96839070024294 | 0.0215619882403854 | 0.15991055187751 | -3.3051939804844 |
| **Ptprs** | 0.296015927500001 | 26.34280271375 | 2.96765262527072 | 0.0215842037573338 | 0.15991055187751 | -3.3062251688377 |
| **Rps24** | 0.438738187499993 | 24.44285413625 | 2.96756387889346 | 0.021586876596833 | 0.15991055187751 | -3.30634916261658 |
| **Adam22** | 0.280108305000006 | 28.435233555 | 2.95901195827572 | 0.0218460989220622 | 0.160793682679701 | -3.31830100964063 |
| **Ociad1** | -0.429528662500005 | 25.63371028875 | -2.95899599153276 | 0.0218465859822573 | 0.160793682679701 | -3.31832333036069 |
| **Timm50** | -0.602443037500002 | 27.27651114375 | -2.95261783073496 | 0.0220420757099692 | 0.1616302258451 | -3.32724151667022 |
| **Apoe** | 0.294715762500005 | 27.23475484875 | 2.9506919556308 | 0.0221014681911387 | 0.1616302258451 | -3.32993506343021 |
| **Atcay** | -0.531784469999991 | 28.69532423 | -2.94686745681252 | 0.022219917339684 | 0.161978951562219 | -3.33528502002898 |

| | | | | | | |
|---|---|---|---|---|---|---|
| **Atp2a3** | -0.82227816499999 | 23.66242021 | -2.9430513428237 | 0.0223387790615311 | 0.161997355740374 | -3.34062453621125 |
| **Ppia** | 1.10397588 | 22.23584384 | 2.942244826855 | 0.0223639862009429 | 0.161997355740374 | -3.34175317940284 |
| **Psmc2** | -0.807543995000007 | 23.6700469775 | -2.93570047193857 | 0.0225696460972783 | 0.162971356204006 | -3.35091349196648 |
| **Pgm1** | 0.390326197499988 | 23.35414648875 | 2.92256017335802 | 0.0229886738550639 | 0.16547507690013 | -3.36931751058117 |
| **Mecp2** | -0.354767072500007 | 26.08792769375 | -2.91920565162808 | 0.0230969627715967 | 0.165733378633808 | -3.37401814248411 |
| **Psmd8** | 0.605778874999999 | 24.1507808 | 2.91348081396508 | 0.0232830205629845 | 0.166546356464598 | -3.38204244567798 |
| **Cd81** | -0.394344090000004 | 26.051264305 | -2.9085213496219 | 0.0234454886914767 | 0.167186054874736 | -3.38899616567017 |
| **Prdx2** | -0.516699764999998 | 27.5027969675 | -2.89550570224613 | 0.0238776136163213 | 0.169234900314238 | -3.40725518377994 |
| **Ktn1** | 0.583358069999996 | 26.73024826 | 2.89321458421902 | 0.0239545493972786 | 0.169234900314238 | -3.41047070792209 |
| **Stxbp5** | -0.364274544999997 | 24.4156742925 | -2.89245907034371 | 0.0239799771151187 | 0.169234900314238 | -3.41153114452269 |
| **Cisd1** | 0.387560195000002 | 27.9184661825 | 2.89101832658924 | 0.0240285463530482 | 0.169234900314238 | -3.41355349437828 |
| **Hspa4** | 0.376637385000002 | 23.33763739 | 2.88666395736716 | 0.0241759719322681 | 0.169750919487613 | -3.41966666422773 |
| **Slc6a11** | -0.379570409999996 | 30.8844729625 | -2.88100389133801 | 0.0243690378703869 | 0.170583265092709 | -3.42761517218872 |
| **Hdhd2** | -0.49051549499999 | 25.363445585 | -2.87255724645144 | 0.0246601962966704 | 0.17209508939963 | -3.43948158166734 |
| **Pdcd5** | 0.407757424999996 | 24.94179923 | 2.86757468391112 | 0.0248336714877783 | 0.172778948436244 | -3.44648400696817 |
| **Picalm** | 0.369474697499999 | 23.73817956875 | 2.86474946036549 | 0.024932609069672 | 0.172802419820992 | -3.45045538175243 |
| **Ube2d2** | 0.526202644999991 | 26.83631402 | 2.86115819466799 | 0.0250589753340201 | 0.172802419820992 | -3.45550444555187 |
| **Dnajc5** | 0.421325572499995 | 22.01992068875 | 2.86102931915716 | 0.0250635226651678 | 0.172802419820992 | -3.45568565346178 |
| **Pi4ka** | -0.585792394999995 | 28.1871087875 | -2.85614080704465 | 0.0252366583558092 | 0.173473606535877 | -3.46256016204612 |
| **Hspa12a** | 0.609111524999996 | 26.228484545 | 2.85400016645955 | 0.0253128709054717 | 0.173476531444984 | -3.46557101704368 |

| | | | | | | |
|---|---|---|---|---|---|---|
| **Rap1b** | 0.434652945000003 | 25.88520777 | 2.84026514895497 | 0.0258077003090749 | 0.175730510022681 | -3.48489762865005 |
| **Ndufv3** | 0.254479195000009 | 33.28775996 | 2.8384727074578 | 0.0258730266344355 | 0.175730510022681 | -3.48742079321812 |
| **Atic** | 0.338222307500004 | 21.27677456125 | 2.83842594054856 | 0.0258747334123248 | 0.175730510022681 | -3.48748662864115 |
| **Nipsnap2** | 0.353181609999996 | 20.7864525775 | 2.83639829486923 | 0.0259488477010337 | 0.175730510022681 | -3.49034116736239 |
| **Ndufs1** | 0.285002802499999 | 28.20159356375 | 2.83125322072056 | 0.0261379182989949 | 0.176488775771089 | -3.49758576512786 |
| **Cox7a2** | -0.442763142499992 | 23.07781612375 | -2.8289234860824 | 0.0262240092594897 | 0.176549285867564 | -3.5008667978402 |
| **D11Wsu47e** | 0.254496169999999 | 26.33175706 | 2.82077291571778 | 0.0265275622500069 | 0.1777121516724 | -3.51234846344701 |
| **Hspd1** | -0.722615617500004 | 25.39307655375 | -2.81309365980359 | 0.0268169572882292 | 0.1777121516724 | -3.52317037195478 |
| **Gstp1** | -0.331656917500009 | 24.03531498875 | -2.8123626212918 | 0.0268446798605805 | 0.1777121516724 | -3.52420078868539 |
| **Lingo1** | 0.904027432500001 | 20.91957970125 | 2.81221581737819 | 0.0268502506301394 | 0.1777121516724 | -3.52440771674701 |
| **Thop1** | -0.3649892225 | 27.86421047125 | -2.81216892592739 | 0.0268520302775338 | 0.1777121516724 | -3.52447381309051 |
| **Nceh1** | 0.372026022499995 | 27.87091305625 | 2.81189149481654 | 0.0268625620264965 | 0.1777121516724 | -3.52486487209264 |
| **Epn2** | 0.744577085000007 | 23.9717406175 | 2.80920124240273 | 0.0269649148563173 | 0.177875187625678 | -3.52865724330805 |
| **Grm5** | -0.660375417499999 | 23.92834835625 | -2.80552059731373 | 0.0271056150198876 | 0.178289519484261 | -3.53384652501608 |
| **Pitpna** | 0.673028890000001 | 19.515397835 | 2.7999210567331 | 0.0273211560771131 | 0.179192338855335 | -3.54174294260024 |
| **Spata20** | -0.444641914999988 | 28.2288804 | -2.79045859924625 | 0.0276895087934918 | 0.181089387509437 | -3.55509143830377 |
| **Txndc12** | 0.227714307499994 | 27.11062838875 | 2.78489634957632 | 0.0279084748407579 | 0.181919471191861 | -3.56294065545474 |
| **Dlg4** | -0.250329627499998 | 21.05020947875 | -2.7832058715576 | 0.0279753839491198 | 0.181919471191861 | -3.56532657325112 |
| **Asah1** | 0.263176812500006 | 26.12605716375 | 2.77905467953278 | 0.0281404062055453 | 0.182110570902655 | -3.57118626281955 |
| **Cnot11** | -0.533135687499996 | 23.37493281875 | -2.77777065142392 | 0.0281916575617644 | 0.182110570902655 | -3.57299897106418 |

| | | | | | | |
|---|---|---|---|---|---|---|
| **Rab4b** | -0.650907774999993 | 24.765047825 | -2.77611519962809 | 0.028257879252266 | 0.182110570902655 | -3.57533618048746 |
| **Uba1** | 0.666629870000001 | 23.33121211 | 2.77449107315281 | 0.028323007095389 | 0.182110570902655 | -3.57762932652276 |
| **Itm2b** | -13.5529989675 | 15.78895771875 | -2.76927238828532 | 0.02853334970879 | 0.182949124603418 | -3.58499880266221 |
| **Nefh** | 0.535984735 | 26.43029708 | 2.76504160519954 | 0.0287050813098047 | 0.18353612044174 | -3.59097442663052 |
| **Kcnj3** | -0.499786992500002 | 26.52736192875 | -2.75535947868355 | 0.0291021947940958 | 0.18511824335822 | -3.60465356650033 |
| **Psmb5** | 0.273641355000002 | 21.3177929175 | 2.75506721416764 | 0.029114271563547 | 0.18511824335822 | -3.60506656909985 |
| **Gng3** | 0.394693689999997 | 29.53714809 | 2.74004626477178 | 0.0297421099936636 | 0.188586398270072 | -3.62629933449408 |
| **Slc44a1** | -0.680905087500001 | 27.61303182125 | -2.72915534335528 | 0.0302062152603732 | 0.18962707618988 | -3.641701842641 |
| **Chid1** | 0.925634112499999 | 22.12923081125 | 2.72902413712046 | 0.0302118526445041 | 0.18962707618988 | -3.64188743970308 |
| **Acot2** | 0.823130662499999 | 21.85673869375 | 2.72866101261904 | 0.0302274603516377 | 0.18962707618988 | -3.64240110018 |
| **Blmh** | 0.70346955250001 | 25.18998065625 | 2.72672106500726 | 0.030310986216412 | 0.18962707618988 | -3.64514538418102 |
| **Prdx6** | -0.357262229999993 | 26.861821945 | -2.72176350644916 | 0.0305255407994521 | 0.18962707618988 | -3.65215931832452 |
| **Cnpy2** | 0.291779549999994 | 24.65849924 | 2.72128138462642 | 0.0305464911311018 | 0.18962707618988 | -3.65284148959814 |
| **Gprc5b** | 0.702817827500006 | 23.63791105375 | 2.72127564973164 | 0.0305467404285705 | 0.18962707618988 | -3.65284960417549 |
| **Adam23** | 0.301644422500008 | 30.45755153625 | 2.72076428454814 | 0.0305689782062323 | 0.18962707618988 | -3.65357316599544 |
| **Plppr4** | -0.64521657249999 | 25.64754950125 | -2.7133467793258 | 0.0308934622744678 | 0.191121986881775 | -3.66407012787468 |
| **Arpc5l** | -0.626154229999997 | 23.086368365 | -2.7099713863827 | 0.0310423163741358 | 0.191525234987593 | -3.66884775543 |
| **Mecr** | 0.395841637500002 | 24.01457123375 | 2.70551135826846 | 0.0312401579134785 | 0.19222774587084 | -3.67516145673088 |
| **Cnnm1** | 0.788476135 | 20.880002355 | 2.69439146859822 | 0.0317392051378129 | 0.193647222520428 | -3.6909070939903 |
| **Diras2** | -0.265530325 | 31.52587728 | -2.69415619198496 | 0.0317498539447303 | 0.193647222520428 | -3.69124030497371 |
| **Hspa8** | 0.598422464999992 | 25.2489646425 | 2.69369773920075 | 0.0317706146459039 | 0.193647222520428 | -3.69188959691932 |

| | | | | | | |
|---|---|---|---|---|---|---|
| **Rps5** | -0.240468602499995 | 27.10430412375 | -2.69284557035251 | 0.0318092423187771 | 0.193647222520428 | -3.69309652169445 |
| **Hpca** | 0.411926057499993 | 28.36459107125 | 2.68505217090189 | 0.0321647990988396 | 0.194885899403603 | -3.70413580870057 |
| **Ndufs5** | 0.246232455000005 | 25.0136992825 | 2.684655822623 | 0.0321829925620629 | 0.194885899403603 | -3.70469730475297 |
| **F3** | -0.711784492499998 | 29.93681437625 | -2.68270913425288 | 0.0322725075346562 | 0.194892554593468 | -3.70745522587664 |
| **Slc6a9** | 0.38698192 | 26.4346079575 | 2.68093364352851 | 0.0323543777831009 | 0.194892554593468 | -3.70997075042931 |
| **Glod4** | 0.509444744999996 | 25.4203672225 | 2.67757604232261 | 0.0325097966504999 | 0.195044256450295 | -3.71472818711398 |
| **Nptx1** | -0.372863339999991 | 27.008360995 | -2.67578732237919 | 0.0325929132147297 | 0.195044256450295 | -3.71726284719691 |
| **Tpm4** | -0.192341392499998 | 26.67455085875 | -2.67487932914392 | 0.0326351901356325 | 0.195044256450295 | -3.71854954737061 |
| **Rab5c** | 0.371570015 | 19.1251770875 | 2.67217965876902 | 0.0327612286093986 | 0.195287636163837 | -3.72237540252674 |
| **Igsf8** | -0.435755074999996 | 28.8390436375 | -2.66646360563731 | 0.0330297768519523 | 0.196238835942769 | -3.73047693099937 |
| **Dld** | 0.710003525000001 | 23.2188931475 | 2.66407498989284 | 0.0331426791999469 | 0.196238835942769 | -3.73386277581167 |
| **Rps6** | 0.836696474999997 | 24.267985775 | 2.66163457655434 | 0.0332584471098861 | 0.196238835942769 | -3.73732227663375 |
| **Sptb** | 0.224355322500003 | 24.87772935625 | 2.66152351271288 | 0.033263725795454 | 0.196238835942769 | -3.73747972497229 |
| **Sez6l** | 0.305992462499997 | 25.39423646125 | 2.65577115081736 | 0.033538328437045 | 0.197350215404617 | -3.74563514512472 |
| **Atp5f1c** | 1.00806002500001 | 20.06958531 | 2.63706535347906 | 0.0344478027633045 | 0.202182103910779 | -3.77216378200114 |
| **Ndufb4** | 0.271653210000007 | 22.937517305 | 2.62957600972103 | 0.0348191272271479 | 0.202983375726195 | -3.78278861039622 |
| **Ndrg1** | 0.512632354999994 | 23.95585263 | 2.62897645603265 | 0.0348490335333698 | 0.202983375726195 | -3.78363925266595 |
| **Myl12b** | 1.2852621725 | 23.27595496125 | 2.62894994043134 | 0.034850356776057 | 0.202983375726195 | -3.78367687307016 |
| **Tmem33** | -0.47844274500001 | 27.0035686425 | -2.62573744731513 | 0.0350110629796552 | 0.203111645957337 | -3.78823493338285 |
| **Slc24a2** | 0.301912072499995 | 24.13312241375 | 2.62496446394868 | 0.0350498471617073 | 0.203111645957337 | -3.78933173322907 |

| | | | | | | |
|---|---|---|---|---|---|---|
| **Got2** | 0.816222167500001 | 24.17604746625 | 2.62074699688176 | 0.0352622487435842 | 0.203826483267839 | -3.79531630030896 |
| **Stx12** | 0.437531382500008 | 22.99297029625 | 2.61783361991736 | 0.0354097577074957 | 0.204163565220296 | -3.79945068724537 |
| **Lmnb1** | 0.562465607499998 | 25.99468728875 | 2.60807568798857 | 0.0359085257656369 | 0.206519134365686 | -3.81330003144094 |
| **Grid2** | 0.272705719999998 | 25.4176287075 | 2.60287055019127 | 0.0361775740981474 | 0.207064546082843 | -3.82068873050166 |
| **Ndufb3** | 0.616287477499998 | 26.58573535375 | 2.60207438178716 | 0.0362189122594676 | 0.207064546082843 | -3.82181895685167 |
| **Plpbp** | 0.66922495 | 25.7634331375 | 2.60100062217368 | 0.0362747413626999 | 0.207064546082843 | -3.82334327322938 |
| **Tmco1** | -0.488895867500005 | 27.59697659125 | -2.59620687214998 | 0.0365250844501471 | 0.207974921160166 | -3.83014887793397 |
| **Rhog** | 0.291019755000001 | 26.713997755 | 2.58486469502292 | 0.037124601822285 | 0.210813202999123 | -3.84625343036989 |
| **Slc25a22** | 0.649286915000001 | 23.260349245 | 2.58255120116614 | 0.0372481407747868 | 0.210813202999123 | -3.84953868667741 |
| **Rab22a** | -0.423456032499988 | 26.51193482625 | -2.58126827537004 | 0.0373168325925911 | 0.210813202999123 | -3.85136054520461 |
| **Cplx1** | 0.559907967500003 | 23.94391389875 | 2.57888858696602 | 0.0374445979975923 | 0.210813202999123 | -3.85473999021796 |
| **Nfasc** | -0.32109358500001 | 26.952123755 | -2.57644877810461 | 0.037576064244584 | 0.210813202999123 | -3.85820493869167 |
| **B3gat3** | 0.351597437500001 | 24.28219639875 | 2.57644735563687 | 0.0375761410326091 | 0.210813202999123 | -3.85820695887699 |
| **Sh3gl2** | 0.289529417500006 | 27.85167183375 | 2.57271880790681 | 0.0377779789399699 | 0.211212804723759 | -3.86350237581408 |
| **Rab3c** | -0.517151475000006 | 27.8910174525 | -2.57093106312824 | 0.037875155078338 | 0.211212804723759 | -3.86604149374947 |
| **Atp5f1b** | 0.734515025 | 21.684266745 | 2.56897181863914 | 0.0379819522300987 | 0.211212804723759 | -3.86882426267059 |
| **Fkbp1a** | 1.08347151 | 22.923495605 | 2.56710997430515 | 0.0380837305628781 | 0.211212804723759 | -3.87146875922068 |
| **Col7a1** | 0.5855461775 | 22.77071832875 | 2.56657125653913 | 0.0381132326498233 | 0.211212804723759 | -3.87223394660958 |
| **Sema4b** | -0.339482462500005 | 28.17206063375 | -2.56497106081612 | 0.0382010053104571 | 0.211212804723759 | -3.87450687435897 |
| **Gnb5** | 0.282491174999997 | 27.03529431 | 2.56215144089228 | 0.0383561769304745 | 0.211559732515316 | -3.87851199202043 |

| | | | | | | |
|---|---|---|---|---|---|---|
| **Romo1** | 0.381158619999994 | 18.9821768325 | 2.55887498320714 | 0.0385373134770723 | 0.211829590807156 | -3.88316619720417 |
| **Mapk3** | 0.517126215000005 | 24.030771245 | 2.55635117090688 | 0.0386774469800314 | 0.211829590807156 | -3.88675139310953 |
| **Eno1** | 0.349684052499992 | 23.81236231625 | 2.55625620959145 | 0.0386827299944916 | 0.211829590807156 | -3.88688629222313 |
| **Bcat1** | 0.587076915000001 | 21.90623706 | 2.5543646253569 | 0.0387881215955607 | 0.211899785995796 | -3.88957344841849 |
| **Gap43** | -0.643104769999994 | 28.299607725 | -2.54909986030848 | 0.0390830279539217 | 0.212431296455517 | -3.89705278508824 |
| **Ndufb10** | 0.448163667500012 | 26.25108306125 | 2.54846777269215 | 0.0391185907691747 | 0.212431296455517 | -3.89795078174465 |
| **Plp1** | 0.421812950000003 | 27.7375390475 | 2.54766455868495 | 0.0391638301023277 | 0.212431296455517 | -3.8990919031348 |
| **Opa3** | 0.3748759275 | 22.58057373625 | 2.54211196041623 | 0.0394780590263252 | 0.212626840317974 | -3.90698068919315 |
| **Hsd17b11** | -0.736115427500007 | 23.07962892125 | -2.54141983571043 | 0.0395174104703915 | 0.212626840317974 | -3.90796404541317 |
| **Ndufa7** | 0.450526592500012 | 24.98167452125 | 2.54081666883038 | 0.0395517373535846 | 0.212626840317974 | -3.9088210171702 |
| **Slc30a3** | 0.280658057500005 | 30.48232823375 | 2.53871602950786 | 0.0396715289711504 | 0.212626840317974 | -3.91180561372775 |
| **Ndufv1** | -0.563866297500006 | 25.77233509625 | -2.53730648093563 | 0.0397521214621958 | 0.212626840317974 | -3.91380833569733 |
| **Dnajc3** | 0.532831849999994 | 23.3387234575 | 2.53471473231657 | 0.0399007515450762 | 0.212626840317974 | -3.91749081721967 |
| **Cand1** | 0.367542965000005 | 27.3005678275 | 2.53364191861237 | 0.0399624433816315 | 0.212626840317974 | -3.91901514469227 |
| **Ndufb5** | 0.599257817500003 | 23.91483252875 | 2.53205972968131 | 0.0400536074633671 | 0.212626840317974 | -3.92126325012838 |
| **Ndufa4** | -0.41567255 | 23.1379541775 | -2.53149880197627 | 0.0400859793493712 | 0.212626840317974 | -3.92206026911717 |
| **Edil3** | -1.03550716749999 | 22.50049896375 | -2.5300767767401 | 0.0401681680769315 | 0.212626840317974 | -3.92408083111995 |
| **Rpl10** | 0.397760187500005 | 24.61782328875 | 2.52915256610308 | 0.0402216784000362 | 0.212626840317974 | -3.92539405697986 |
| **Hint1** | 0.344053317499991 | 28.45755555125 | 2.52303671271588 | 0.0405776461312026 | 0.21373304733262 | -3.93408437285833 |
| **Asrgl1** | -0.561358737499997 | 26.06538943375 | -2.52235233366075 | 0.0406176826516775 | 0.21373304733262 | -3.93505686087123 |
| **Fahd1** | -0.368789960000008 | 24.78456676 | -2.5201768150995 | 0.040745223983755 | 0.213912425914714 | -3.93814825111338 |

| | | | | | | |
|---|---|---|---|---|---|---|
| **Gabarapl2** | -0.366587592500004 | 28.22918352125 | -2.51825021154373 | 0.0408585193257588 | 0.214016363241789 | -3.94088596566029 |
| **Faim2** | 0.429214447499998 | 19.03823661375 | 2.5121801439427 | 0.0412176150452042 | 0.214085110018982 | -3.94951173072526 |
| **Grm1** | 1.6985948875 | 20.31989754875 | 2.51072232309567 | 0.0413043437463055 | 0.214085110018982 | -3.95158337497985 |
| **Cnp** | 0.944564877500003 | 21.96668492875 | 2.50821545017145 | 0.0414539252735489 | 0.214085110018982 | -3.95514580627681 |
| **Tspan2** | 0.212874930000012 | 27.66386344 | 2.50700625691298 | 0.0415262767216929 | 0.214085110018982 | -3.95686416017504 |
| **Gcsh** | -0.605246057499997 | 26.64346814875 | -2.50587569465866 | 0.0415940417059867 | 0.214085110018982 | -3.9584707793527 |
| **Sorl1** | -1.9111631775 | 24.82875512125 | -2.5054626791021 | 0.0416188260928408 | 0.214085110018982 | -3.9590577087434 |
| **Ndufa3** | 0.534541542500005 | 25.31577384625 | 2.50450528369119 | 0.0416763366915759 | 0.214085110018982 | -3.96041824969526 |
| **Eef1b** | -0.431182262499991 | 25.94643079125 | -2.50445886334312 | 0.0416791272471469 | 0.214085110018982 | -3.96048421707887 |
| **Pcmt1** | 0.534104620000008 | 19.99512201 | 2.50283712777329 | 0.0417767393787373 | 0.214085110018982 | -3.96278885038806 |
| **Tomm22** | -0.776563299999992 | 24.7483491275 | -2.5023364679203 | 0.0418069218778876 | 0.214085110018982 | -3.96350033527515 |
| **Alg2** | -0.769752207499995 | 21.98787931125 | -2.4934943895503 | 0.0423437085309369 | 0.216349885775256 | -3.97606586826259 |
| **Hadh** | -0.459838277499998 | 22.28753080375 | -2.48796360514307 | 0.0426830960165573 | 0.216530817705007 | -3.98392575325962 |
| **Hnrnpm** | 14.2504641375 | 17.66349107625 | 2.48776256342333 | 0.0426954855152871 | 0.216530817705007 | -3.98421145688651 |
| **Cdh10** | -0.215174770000004 | 28.10160271 | -2.48773914751215 | 0.0426969287980592 | 0.216530817705007 | -3.98424473361417 |
| **Fasn** | 0.426211532499998 | 28.28522262625 | 2.48675708872833 | 0.0427575052873146 | 0.216530817705007 | -3.98564035290438 |
| **Coa3** | 0.332035122500002 | 27.84678572875 | 2.48205904242963 | 0.0430485284233044 | 0.21752335885418 | -3.99231680834928 |
| **Rps14** | -1.45735263000001 | 21.983064615 | -2.4674761769399 | 0.0439649917831533 | 0.221192725195174 | -4.01304033893356 |
| **Cdh2** | 0.304225635000002 | 26.2595966325 | 2.46742925563043 | 0.0439679728981233 | 0.221192725195174 | -4.01310701676122 |
| **Aldh18a1** | -0.611221990000004 | 23.5984593175 | -2.46409359268583 | 0.04418043826959 | 0.221774173682218 | -4.01784714923878 |
| **Txndc5** | 0.706553312500002 | 22.67867195375 | 2.45572806204 | 0.0447179601272429 | 0.223440278593206 | -4.02973462486732 |

| | | | | | | |
|---|---|---|---|---|---|---|
| **Gpx1** | -0.392191674999992 | 26.3193724525 | -2.45572326154082 | 0.0447182705118308 | 0.223440278593206 | -4.02974144625212 |
| **Slc25a5** | 0.528026042500009 | 23.74990194875 | 2.4538166161327 | 0.0448417240468585 | 0.223440278593206 | -4.03245072406679 |
| **Ndufv2** | 0.281254830000005 | 25.6359906575 | 2.45287522022203 | 0.0449028082799802 | 0.223440278593206 | -4.0337884034156 |
| **Sucla2** | 0.455919144999999 | 27.5902897625 | 2.44850178786366 | 0.0451877151516347 | 0.224064553910889 | -4.04000273401939 |
| **Ndufs8** | 0.438596567499999 | 21.57105456125 | 2.4479462582816 | 0.0452240384040327 | 0.224064553910889 | -4.04079208751384 |
| **Atp1a3** | 0.609214557499996 | 26.56225632875 | 2.44604384791788 | 0.0453486557798086 | 0.224196702116592 | -4.04349520261314 |
| **Ak1** | -0.341884189999988 | 29.366716675 | -2.44089331877163 | 0.0456878224652122 | 0.225331902384009 | -4.05081332992221 |
| **Stx7** | 0.470155215000013 | 26.5181639075 | 2.43957345490304 | 0.0457751571029114 | 0.225331902384009 | -4.05268860642585 |
| **Cct4** | 0.260720212500008 | 29.36809452125 | 2.43601787639468 | 0.0460112854212568 | 0.225501254026041 | -4.05774029759037 |
| **Prkacb** | 0.350492997499991 | 21.75737215125 | 2.4353126219776 | 0.0460582709220349 | 0.225501254026041 | -4.05874228803225 |
| **Tmem35a** | 0.282429572500003 | 25.94970088125 | 2.43345410508623 | 0.0461823262512313 | 0.225501254026041 | -4.06138274319053 |
| **Cul3** | -0.331918977499996 | 23.52843165625 | -2.43181619808342 | 0.046291941515782 | 0.225501254026041 | -4.06370972962434 |
| **Tm9sf4** | -0.2132279625 | 21.41551742375 | -2.43166407146125 | 0.0463021360385494 | 0.225501254026041 | -4.06392585502811 |
| **Dgke** | -0.313047442500011 | 30.28680759125 | -2.43007715102275 | 0.0464086190639845 | 0.225539976724969 | -4.06618036295947 |
| **Letm1** | 0.354698009999996 | 24.300389675 | 2.42521263829357 | 0.0467366052479856 | 0.226652731806439 | -4.07309104018293 |
| **Psma5** | 0.283761834999993 | 24.1453951225 | 2.42063062944986 | 0.0470477282767281 | 0.227588130270061 | -4.07960002550279 |
| **Micos10** | -0.270262090000003 | 26.5606640025 | -2.4186891582743 | 0.0471801988705955 | 0.227588130270061 | -4.08235787649976 |
| **Uqcr10** | 0.750665584999993 | 23.3251669275 | 2.41799335522486 | 0.0472277684046654 | 0.227588130270061 | -4.08334624486264 |
| **Dynll2** | -0.490920212500008 | 23.11884294875 | -2.41566228013777 | 0.0473874957718455 | 0.227878104667551 | -4.08665740462263 |
| **Vps35** | 0.4680876125 | 21.89613546375 | 2.40775281921407 | 0.0479336162030594 | 0.230021063917826 | -4.09789154888134 |
| **Nfu1** | -0.608739997499992 | 19.58809117125 | -2.40490981825406 | 0.0481314943380572 | 0.230287405628073 | -4.10192926580777 |

| | | | | | | |
|---|---|---|---|---|---|---|
| **Lrrc59** | 0.167448892499991 | 24.20844123625 | 2.40406680765179 | 0.0481903308413486 | 0.230287405628073 | -4.10312650124455 |
| **Slc7a14** | -0.411591885000007 | 27.34301481 | -2.39981696886052 | 0.0484880697464661 | 0.23081363096462 | -4.10916183614126 |
| **Sparcl1** | -0.646192527500006 | 24.30288600875 | -2.39961705960307 | 0.048502121666222 | 0.23081363096462 | -4.10944572353525 |
| **Folh1** | 2.18272715 | 24.25699661 | 2.39313472385706 | 0.0489600512285721 | 0.232330457151875 | -4.11865064519409 |
| **Pfkm** | 0.62786827750001 | 26.62197686375 | 2.39223644275148 | 0.0490238579311295 | 0.232330457151875 | -4.11992612283703 |
| **Rpl27** | -0.754919557499992 | 23.14905170375 | -2.38682733852069 | 0.0494098882620925 | 0.233676103784979 | -4.12760611672843 |
| **Ndufb6** | 0.229929635000008 | 26.13505868 | 2.38437342700321 | 0.0495860445019908 | 0.234025682195994 | -4.13108998881826 |
| **Slc44a2** | 0.499477670000001 | 21.9224647675 | 2.37145578023556 | 0.0505240405011797 | 0.236945862276781 | -4.14942660309271 |
| **Prdx1** | 0.266796387500008 | 25.95330017625 | 2.36931821716062 | 0.0506810039645915 | 0.236945862276781 | -4.15246038753265 |
| **Etfa** | 0.521099290000002 | 18.7524892475 | 2.36925256882261 | 0.0506858325011198 | 0.236945862276781 | -4.15255355811205 |
| **Nol3** | 0.253134332499997 | 24.31945164375 | 2.36730336603828 | 0.0508294146202586 | 0.236945862276781 | -4.15531987646609 |
| **Atp5pb** | 0.552240900000001 | 21.4921868525 | 2.36607727141122 | 0.0509199448778826 | 0.236945862276781 | -4.15705989283067 |
| **Mcts1** | -0.139062565 | 27.3812641225 | -2.36580265034355 | 0.0509402445346961 | 0.236945862276781 | -4.15744961548863 |
| **Dpysl5** | -0.609040459999996 | 25.2012572025 | -2.36438976552118 | 0.051044814551523 | 0.236945862276781 | -4.15945464151992 |
| **Rps12** | 0.6514749125 | 20.45478791375 | 2.36223597597505 | 0.0512046439543815 | 0.236945862276781 | -4.16251095749365 |
| **Crmp1** | 0.829719387499992 | 21.68258863375 | 2.36032667247992 | 0.0513467595011372 | 0.236945862276781 | -4.16522020583881 |
| **Hras** | 0.442861417499998 | 26.14175731125 | 2.35933941717431 | 0.0514204026136438 | 0.236945862276781 | -4.16662104425233 |
| **Glg1** | 0.365119362500007 | 24.89061812125 | 2.35919143193034 | 0.0514314507218317 | 0.236945862276781 | -4.16683102087118 |
| **Ddah1** | -0.298260822499994 | 30.06174610375 | -2.35898373362029 | 0.0514469609224815 | 0.236945862276781 | -4.16712572321417 |
| **Twf1** | 0.898650507500001 | 25.76805922875 | 2.35185940388295 | 0.0519818943636231 | 0.238928827707497 | -4.17723347740922 |

| | | | | | | |
|---|---|---|---|---|---|---|
| **Pde10a** | 0.273740562499999 | 25.63333818375 | 2.34984245004819 | 0.0521343710044354 | 0.239149449357019 | -4.18009472557508 |
| **Atp5f1d** | 0.220696617500003 | 24.19127574125 | 2.34097301377839 | 0.0528103337401653 | 0.241765707862477 | -4.19267504980033 |
| **Gjc3** | -0.431314180000008 | 22.6899995275 | -2.33229699367656 | 0.0534802380890078 | 0.243945053173481 | -4.20497797780326 |
| **Adcy5** | -1.3957822725 | 23.99085384375 | -2.32932473331463 | 0.0537117304255002 | 0.243945053173481 | -4.20919202661871 |
| **Tmx2** | 0.633632869999992 | 25.60217232 | 2.32869503554953 | 0.0537609053955296 | 0.243945053173481 | -4.21008475827441 |
| **Cs** | 0.271932585000005 | 26.3509433475 | 2.32654359692904 | 0.0539292652731308 | 0.243945053173481 | -4.21313475254786 |
| **Epb41l1** | -0.464485777499998 | 23.78738278875 | -2.32405794952497 | 0.054124449879783 | 0.243945053173481 | -4.21665828278511 |
| **Pld3** | 0.445836745000008 | 24.7621663425 | 2.32320190571539 | 0.054191837488521 | 0.243945053173481 | -4.21787170357122 |
| **Atp6v0d1** | 0.388919099999995 | 25.1477907525 | 2.32288686548953 | 0.054216658980292 | 0.243945053173481 | -4.21831825687759 |
| **Rab14** | 0.620166195000003 | 25.14654019 | 2.32259596482544 | 0.0542395888759023 | 0.243945053173481 | -4.21873058958583 |
| **Gsn** | 0.505465689999998 | 24.5819330775 | 2.32252053613001 | 0.0542455360704682 | 0.243945053173481 | -4.21883750420961 |
| **Cox5a** | -0.419313957500009 | 24.82036684125 | -2.31627543518484 | 0.0547402515900911 | 0.24533943988402 | -4.22768856243934 |
| **Dnm1l** | 0.263364287500004 | 26.27953125625 | 2.31590216092645 | 0.0547699667019371 | 0.24533943988402 | -4.22821753872611 |
| **Dpp10** | 0.542375982499998 | 25.42511951625 | 2.31412398723104 | 0.0549117475263913 | 0.245494121265449 | -4.23073734179918 |
| **Cox6b1** | 0.262703127500004 | 26.36190767625 | 2.31149979761783 | 0.0551216697291826 | 0.245952245633721 | -4.23445573313255 |
| **Gnaq** | -0.487801042500003 | 28.33627906125 | -2.30714411201968 | 0.0554719137551861 | 0.246932459681352 | -4.24062685141787 |
| **Ank1** | -0.392029242499998 | 29.03683957375 | -2.30506985876123 | 0.0556395038429272 | 0.246932459681352 | -4.24356531171243 |
| **Crocc** | 0.508707240000003 | 26.7965920825 | 2.30475504644481 | 0.0556649843580504 | 0.246932459681352 | -4.24401126687071 |
| **Decr1** | -0.143218187500004 | 21.02518455375 | -2.30223438021611 | 0.0558694338379196 | 0.246939746538087 | -4.2475817958614 |
| **Try5** | 0.44428774 | 26.106845215 | 2.30107270138294 | 0.0559639145916957 | 0.246939746538087 | -4.24922720597846 |

| | | | | | | |
|---|---|---|---|---|---|---|
| **Ppp1r1b** | -0.296146207500001 | 27.45086405375 | -2.30074900572322 | 0.0559902701849136 | 0.246939746538087 | -4.24968567829606 |
| **Ddx6** | 0.62298048000001 | 24.5855780975 | 2.29638324703473 | 0.0563469738124724 | 0.248035044339902 | -4.2558686625836 |
| **Dnm1** | -0.757778357500001 | 21.95831391875 | -2.29194327903157 | 0.0567121182908978 | 0.249163222203196 | -4.26215569877612 |
| **Hint2** | 0.241683172500004 | 29.37103115125 | 2.28669194408377 | 0.0571471055214714 | 0.250593342028061 | -4.26959023907334 |
| **L1cam** | -0.278254789999984 | 25.89145968 | -2.28417712585992 | 0.0573566197157982 | 0.250759434566794 | -4.27315002601286 |
| **Sema4d** | 0.805942590000004 | 24.8160962825 | 2.2827939538216 | 0.0574721878006271 | 0.250759434566794 | -4.27510778584383 |
| **Hcn1** | -1.09065373000001 | 22.4315131375 | -2.27937065118107 | 0.0577592359331296 | 0.250759434566794 | -4.27995269660147 |
| **Fbxo2** | 0.168292200000007 | 29.1153478925 | 2.2789321594089 | 0.0577961093601725 | 0.250759434566794 | -4.28057323302977 |
| **Gaa** | 1.35091207 | 22.07901527 | 2.27649421568736 | 0.0580015572795508 | 0.250759434566794 | -4.2840231084025 |
| **Sv2c** | 0.406956590000004 | 25.6433544 | 2.27645972965834 | 0.0580044687791655 | 0.250759434566794 | -4.28407190621649 |
| **Cyfip2** | 0.359513625000016 | 28.6122107925 | 2.27552630678366 | 0.0580833300020137 | 0.250759434566794 | -4.28539267481694 |
| **Ola1** | -0.932509957499995 | 22.03496969625 | -2.27539456073881 | 0.058094469494296 | 0.250759434566794 | -4.28557908777102 |
| **Oxr1** | -0.311228722499997 | 28.14019952625 | -2.27449095579371 | 0.0581709304303048 | 0.250759434566794 | -4.28685760783998 |
| **Vat1l** | -0.802124707499985 | 28.07628774375 | -2.27067872974998 | 0.058494641517495 | 0.25168089179238 | -4.2922510213334 |
| **Cct3** | 0.499823655000004 | 24.42583788 | 2.2690141521512 | 0.0586365618138093 | 0.251818180097204 | -4.29460573223643 |
| **Ndufa13** | -0.3275249 | 27.76294304 | -2.26576334393657 | 0.058914733691896 | 0.252128312988148 | -4.29920382743419 |
| **Clec2l** | -1.2302331025 | 22.47782864125 | -2.26559619683733 | 0.0589290727167581 | 0.252128312988148 | -4.29944023018799 |
| **Phb1** | -0.4537368075 | 22.86587833125 | -2.2612385812113 | 0.059304154104044 | 0.252833474789184 | -4.3056027517189 |
| **Emc4** | 0.759853580000001 | 22.9499676875 | 2.2598538831793 | 0.0594238494795383 | 0.252833474789184 | -4.30756073062535 |
| **Rad18** | 0.4238181375 | 20.86024010875 | 2.25576482480282 | 0.059778749365777 | 0.252833474789184 | -4.31334197613828 |
| **Atp5f1a** | -0.272413639999989 | 25.45104185 | -2.25558931945391 | 0.0597940300578443 | 0.252833474789184 | -4.31359008675194 |

| | | | | | | |
|---|---|---|---|---|---|---|
| **Pdhb** | 0.367847772499999 | 24.34638128125 | 2.25457624121612 | 0.0598823131837153 | 0.252833474789184 | -4.31502222799478 |
| **Canx** | 11.152732635 | 14.6871368 | 2.25435588908216 | 0.0599015329609082 | 0.252833474789184 | -4.3153337204413 |
| **Ppt1** | 0.477365364999997 | 23.237866225 | 2.25410048471041 | 0.0599238179442416 | 0.252833474789184 | -4.31569475910383 |
| **Lrpprc** | 0.704240910000003 | 23.73810865 | 2.25190416025327 | 0.0601158035028812 | 0.252833474789184 | -4.31879929478721 |
| **Atp1b2** | 0.462998704999997 | 23.6749804 | 2.2500915894808 | 0.060274714941063 | 0.252833474789184 | -4.32136114420431 |
| **Pfdn5** | -0.406026435000001 | 25.11907928 | -2.24908179760678 | 0.0603634302933051 | 0.252833474789184 | -4.32278826594684 |
| **Itgb8** | -0.649226015 | 28.3119128375 | -2.24690714772715 | 0.0605549349942592 | 0.252833474789184 | -4.32586142456185 |
| **Echdc1** | 0.322855959999998 | 24.851507335 | 2.24628048334003 | 0.0606102349540647 | 0.252833474789184 | -4.32674694987502 |
| **Pfkl** | 0.212596042499996 | 26.08069943125 | 2.24622447332548 | 0.0606151800503906 | 0.252833474789184 | -4.32682609504006 |
| **Tufm** | 0.334085940000001 | 27.974693115 | 2.24594036846428 | 0.060640269838035 | 0.252833474789184 | -4.32722754715015 |
| **Ndufa5** | 0.409745544999993 | 23.8529676475 | 2.24182962502031 | 0.0610044790099037 | 0.253889549915763 | -4.33303556986703 |
| **Bsg** | 0.618536277500006 | 22.90034362125 | 2.23326649866058 | 0.0617703126737841 | 0.25657583595077 | -4.34513045008506 |
| **Nlgn3** | 0.2758674325 | 28.90460078375 | 2.23211407642906 | 0.0618741203341307 | 0.25657583595077 | -4.3467577673953 |
| **Mtch2** | -0.599460412500001 | 22.88579396375 | -2.22836609313583 | 0.0622129554271382 | 0.25739446085731 | -4.35204955459184 |
| **Svop** | 0.3016727975 | 26.26268352375 | 2.22744591231826 | 0.0622964313302533 | 0.25739446085731 | -4.35334859958937 |
| **Ywhae** | 0.268284657499994 | 30.18836216875 | 2.22175401684411 | 0.0628153114339347 | 0.258403337683644 | -4.36138257875328 |
| **Maob** | 0.207444477500001 | 27.74105763375 | 2.22062356605415 | 0.0629188848945133 | 0.258403337683644 | -4.36297788787978 |
| **Aco2** | 0.189620345000002 | 25.4242099075 | 2.21924616876563 | 0.0630453178216702 | 0.258403337683644 | -4.36492155743926 |
| **Dctn2** | 0.433194805000007 | 25.3289412725 | 2.21860994500937 | 0.063103804424009 | 0.258403337683644 | -4.36581929399739 |
| **Kcna4** | -0.456593245000001 | 23.7681215875 | -2.21859637199548 | 0.0631050527589151 | 0.258403337683644 | -4.36583844570478 |
| **Chchd3** | 0.588285599999995 | 21.2523260875 | 2.21618005820151 | 0.0633276846379036 | 0.258851910957431 | -4.36924766664012 |

| | | | | | | |
|---|---|---|---|---|---|---|
| **Cox7c** | 0.198355517499998 | 28.05765927375 | 2.21261972481473 | 0.0636571734373414 | 0.25973488413204 | -4.37427016231771 |
| **Slc25a27** | -0.229592132500002 | 25.55872900125 | -2.20787897470824 | 0.0640985990168709 | 0.261070628380102 | -4.38095626688365 |
| **Rplp2** | 0.507542355000005 | 24.4438395875 | 2.20580665763551 | 0.0642925306205308 | 0.261395386483828 | -4.38387837782944 |
| **Fscn1** | -0.598793604999997 | 26.144978295 | -2.20262012614969 | 0.0645918915774633 | 0.262146879114918 | -4.38837091540526 |
| **Acot13** | -0.22171078500001 | 24.20930029 | -2.19916599410256 | 0.0649179852614064 | 0.263004014625415 | -4.39323977031154 |
| **Eif4b** | -0.748002332499993 | 22.71006497375 | -2.19534233094139 | 0.0652809070294261 | 0.26400706040699 | -4.39862832586614 |
| **Trp53i11** | -0.945143649999999 | 21.116142745 | -2.1835253018155 | 0.0664155289522735 | 0.268121950214734 | -4.41527361212086 |
| **Ndufb2** | -0.565919217499996 | 26.69219259125 | -2.18018800247625 | 0.0667395546820846 | 0.268840715197346 | -4.4199722258057 |
| **Pcp4** | 0.359250059999994 | 25.295861455 | 2.17927508380368 | 0.0668284696143686 | 0.268840715197346 | -4.42125735563103 |
| **Rab6b** | -0.304269825000006 | 23.6954928 | -2.17656510420568 | 0.0670931165158309 | 0.269431831060942 | -4.42507178540712 |
| **Opa1** | -0.603788242500006 | 24.50090005125 | -2.16973159576241 | 0.0677651569699683 | 0.270828093749665 | -4.43468725228959 |
| **Immt** | 0.710354419999991 | 22.892567505 | 2.1694961642663 | 0.0677884310478816 | 0.270828093749665 | -4.43501845089324 |
| **Aldh2** | -0.331936992500001 | 25.50710444125 | -2.16942203071779 | 0.0677957613449359 | 0.270828093749665 | -4.43512273885647 |
| **Cltb** | -0.570760987500002 | 26.75205070625 | -2.16700245226601 | 0.0680354475737766 | 0.271234408626055 | -4.43852621278118 |
| **Tom1l2** | -0.875151785000003 | 22.503675565 | -2.16600542490174 | 0.0681344626299614 | 0.271234408626055 | -4.43992850759516 |
| **Acsl3** | 12.023578745 | 16.1980900875 | 2.16362627284673 | 0.0683713241069392 | 0.271704793195805 | -4.44327434028609 |
| **Aldh5a1** | -0.448417607499998 | 23.28975836875 | -2.15752413924078 | 0.068982635113236 | 0.2730992306141 | -4.45185332871795 |
| **Rab24** | 0.182430989999993 | 25.7051077 | 2.15677759271205 | 0.0690578010917872 | 0.2730992306141 | -4.45290264643318 |
| **S100b** | 0.405148847500001 | 25.76080845125 | 2.1554352269472 | 0.069193164646635 | 0.2730992306141 | -4.45478928575559 |
| **Suclg1** | 0.1840894475 | 33.97319104875 | 2.15537290598123 | 0.0691994555509732 | 0.2730992306141 | -4.45487687091854 |

| | | | | | | |
|---|---|---|---|---|---|---|
| **Slc6a1** | 0.503568072499995 | 19.49814621125 | 2.15150627900018 | 0.0695908952759365 | 0.274049025685389 | -4.46031022127599 |
| **Gabra6** | 0.229865372500004 | 26.16037233375 | 2.15063340735634 | 0.0696795687850136 | 0.274049025685389 | -4.46153656408978 |
| **Paics** | -0.672468265000006 | 21.2679416475 | -2.14841309789712 | 0.0699056390306282 | 0.274466565593667 | -4.4646556430386 |
| **Gad2** | -0.657439527500003 | 25.28038219125 | -2.14653678813107 | 0.0700972588151975 | 0.274673086418639 | -4.46729107973961 |
| **Mapk10** | -0.246477142500005 | 20.50196665125 | -2.14555013952249 | 0.0701982330951961 | 0.274673086418639 | -4.46867676626934 |
| **Cox4i1** | -1.3471914225 | 22.51777659375 | -2.13520319224377 | 0.071265998128534 | 0.27796857178672 | -4.48320228196301 |
| **Ywhab** | -0.444112302499999 | 24.68908771375 | -2.13252804025889 | 0.0715447099024168 | 0.27796857178672 | -4.4869559328427 |
| **Ckap4** | 0.211115720000002 | 27.956226875 | 2.13117297594625 | 0.0716863054293644 | 0.27796857178672 | -4.48885700249544 |
| **Gsk3b** | 0.268576452499996 | 30.27016129375 | 2.13076828341835 | 0.0717286477708939 | 0.27796857178672 | -4.48942472194817 |
| **Sod2** | 0.369562607500004 | 26.41089037625 | 2.12976957525031 | 0.0718332485640701 | 0.27796857178672 | -4.49082567495147 |
| **Necap1** | -0.471079587499997 | 23.66773219625 | -2.12943508988092 | 0.0718683154950422 | 0.27796857178672 | -4.49129485507331 |
| **Hsdl1** | 0.298596440000004 | 23.6195663875 | 2.12922340139206 | 0.0718905174738918 | 0.27796857178672 | -4.49159178262119 |
| **Rab6a** | 0.265155744999994 | 27.1738748075 | 2.12523201476204 | 0.0723104274926983 | 0.279120688922068 | -4.49718943168951 |
| **Ndufa9** | -0.190626252500003 | 20.50902446625 | -2.12039165070731 | 0.0728229584879351 | 0.280245322596284 | -4.50397534614255 |
| **Acat2** | -0.524386867499995 | 22.55904129875 | -2.12016879587724 | 0.0728466434883306 | 0.280245322596284 | -4.50428771279147 |
| **Nherf1** | 0.191080602499998 | 31.47066711625 | 2.11447078005824 | 0.0734548603944372 | 0.28211103262226 | -4.51227248312642 |
| **Bpnt1** | 0.370352454999999 | 24.863868425 | 2.11229520666861 | 0.0736884265133847 | 0.282534017234736 | -4.51532019007158 |
| **Dlst** | 0.311123285000004 | 24.75394601 | 2.10809425804284 | 0.0741415428619058 | 0.283795972593482 | -4.52120364627671 |
| **Rps27a** | 0.300727060000003 | 29.47310465 | 2.10243472349641 | 0.0747564010491394 | 0.285287080474693 | -4.52912659394095 |
| **Dbnl** | 0.552482709999996 | 23.2124009975 | 2.1016936659207 | 0.0748372877000671 | 0.285287080474693 | -4.53016374125133 |

| | | | | | | |
|---|---|---|---|---|---|---|
| **Rpl30** | -0.232907282500005 | 30.88201722375 | -2.09826601864787 | 0.0752125581622156 | 0.285287080474693 | -4.5349600511838 |
| **Mrps26** | -0.253432315000005 | 28.32450637 | -2.09795108013748 | 0.0752471330749262 | 0.285287080474693 | -4.53540067410097 |
| **Slc25a12** | 1.029503885 | 21.5386692225 | 2.09795017048209 | 0.0752472329627205 | 0.285287080474693 | -4.53540194676069 |
| **Eif5** | 0.559833614999995 | 24.0105462275 | 2.0958284371633 | 0.0754805786958646 | 0.285287080474693 | -4.53837010019942 |
| **Actb** | -0.6339325 | 21.560114355 | -2.09555585478682 | 0.0755106094445866 | 0.285287080474693 | -4.53875138392091 |
| **Sod1** | -11.0000342025 | 15.16399024125 | -2.09539655290505 | 0.0755281654729857 | 0.285287080474693 | -4.53897420857783 |
| **Ykt6** | 0.331179567500001 | 24.95854172875 | 2.0929143768613 | 0.075802244572984 | 0.285850638925141 | -4.54244577055454 |
| **Ppp1cb** | 0.199362502500001 | 28.06808217625 | 2.08969826659285 | 0.076158844504745 | 0.286723018209476 | -4.54694268633668 |
| **Capzb** | 0.425348940000003 | 23.7491757475 | 2.08849578525202 | 0.0762926053638015 | 0.286754964988082 | -4.54862372387937 |
| **Vps26b** | -0.831585419999989 | 26.6779436175 | -2.08357632759081 | 0.0768422840133963 | 0.288073899817428 | -4.55549912004716 |
| **Gria3** | 0.181440374999994 | 25.7940800725 | 2.0831044477767 | 0.0768952174698333 | 0.288073899817428 | -4.55615845601248 |
| **Ermn** | 0.557199319999999 | 24.8696169675 | 2.08165804495807 | 0.0770576959348949 | 0.288210892148651 | -4.5581792728736 |
| **H1-3** | -0.488424889999997 | 24.847087485 | -2.07721168556444 | 0.0775593194900969 | 0.289613837378192 | -4.56438976180667 |
| **Idh3a** | 0.911016859999997 | 24.4660603625 | 2.06716756268924 | 0.0787045020820518 | 0.293411409227714 | -4.57840957361401 |
| **Mtx2** | 11.6337612575 | 16.21291764375 | 2.06267687098164 | 0.0792219539558157 | 0.294153050801939 | -4.58467348808966 |
| **Hepacam** | 11.4122574425 | 15.87724163875 | 2.06222747051261 | 0.0792739237599977 | 0.294153050801939 | -4.58530019349775 |
| **Ass1** | -0.244121379999999 | 22.21262242 | -2.06144668105839 | 0.0793642970938583 | 0.294153050801939 | -4.58638896852824 |
| **Pcyox1** | -0.620737504999994 | 21.6125525 | -2.06098771559712 | 0.079417468499606 | 0.294153050801939 | -4.58702893654109 |
| **Ndufa10** | -0.212170259999997 | 23.2069435225 | -2.05029704421858 | 0.0806660860088094 | 0.298295106420298 | -4.6019275938409 |
| **Ctnnd2** | -0.516877607499996 | 24.03349847875 | -2.04285335550937 | 0.0815470022138174 | 0.300278679331146 | -4.61229187732623 |
| **Cd99l2** | 0.307549702500005 | 21.81633110625 | 2.04157689053091 | 0.0816990232732656 | 0.300278679331146 | -4.6140683903353 |

| | | | | | | |
|---|---|---|---|---|---|---|
| **Cpt1c** | 0.269699052499998 | 24.54701277375 | 2.04138586761959 | 0.0817217974645422 | 0.300278679331146 | -4.61433422550198 |
| **Slc30a9** | 0.293233342500002 | 27.99364173375 | 2.04134035387557 | 0.0817272246497614 | 0.300278679331146 | -4.61439756349334 |
| **Hpcal1** | 0.328739634999998 | 21.0664103325 | 2.03941062490395 | 0.0819576611465661 | 0.300642766609759 | -4.6170827479376 |
| **Gstz1** | 0.254062022500005 | 31.14600788875 | 2.0376159414013 | 0.0821725514406756 | 0.30094875239633 | -4.61957954111062 |
| **Map2k1** | -10.890897485 | 15.321808155 | -2.03238882546675 | 0.0828016295800268 | 0.302768258959555 | -4.62684894531568 |
| **Micu1** | 0.358033207499997 | 25.76956684125 | 2.02753691321155 | 0.0833898359389878 | 0.304360902484443 | -4.63359298728259 |
| **Napb** | 0.494161437500001 | 23.00168058375 | 2.0266063604953 | 0.0835031222194102 | 0.304360902484443 | -4.63488603740286 |
| **Tpbg** | -1.1980343975 | 21.09260168625 | -2.02452830501764 | 0.0837566587054339 | 0.304647440118986 | -4.6377731374895 |
| **Vdac2** | 0.636365532499994 | 23.92015598625 | 2.02378183681771 | 0.0838479192988035 | 0.304647440118986 | -4.63881006971511 |
| **Vdac3** | 0.384281402500008 | 23.60261341875 | 2.02088287065024 | 0.0842032741590233 | 0.30545371561015 | -4.64283628545099 |
| **Acaa1a** | 0.302346042500002 | 25.80284199375 | 2.01904370373966 | 0.084429493809522 | 0.305789733117082 | -4.64538994962225 |
| **Ndufc2** | 0.431735742499995 | 25.45563977375 | 2.01757154437943 | 0.084611005745264 | 0.305963020775528 | -4.64743365960433 |
| **Slc6a5** | 11.15912913 | 15.7537875 | 2.01493764444293 | 0.084936722894435 | 0.306086564357833 | -4.65108932318156 |
| **Hk1** | -0.609850039999998 | 23.3675277375 | -2.01472058225938 | 0.084963620978463 | 0.306086564357833 | -4.65139054273234 |
| **Atp1b1** | 0.299157950000001 | 23.8278979475 | 2.0140535442121 | 0.0850463324296993 | 0.306086564357833 | -4.65231615361771 |
| **Ppm1h** | 0.435339962500002 | 22.03353127125 | 2.01238764857961 | 0.0852532493431346 | 0.306349588298956 | -4.65462752492894 |
| **Lancl1** | 0.2061198225 | 30.25096865625 | 2.0072460989339 | 0.0858950225269751 | 0.307954485664248 | -4.66175854550876 |
| **Flot2** | -0.724312224999995 | 20.63606258 | -2.00665631453733 | 0.0859689455392985 | 0.307954485664248 | -4.66257627882735 |
| **Ccny** | -10.171936095 | 14.44922314 | -2.00325193672279 | 0.0863968810915651 | 0.309003845029051 | -4.66729536965652 |
| **Tagln3** | 0.240467550000005 | 25.3955930825 | 2.00011696342722 | 0.0867928185115275 | 0.309935665480322 | -4.67163940380198 |
| **Kcnc1** | 0.516550999999996 | 23.39640416 | 1.9972899347751 | 0.0871514042895384 | 0.310233136025398 | -4.675555392185 |

| | | | | | | |
|---|---|---|---|---|---|---|
| **Pfkp** | -0.529718412499999 | 21.82061118875 | -1.99565308334045 | 0.0873596958745789 | 0.310233136025398 | -4.67782216886479 |
| **Pptc7** | 0.724589202499999 | 20.91742069125 | 1.9955541583597 | 0.087372299994967 | 0.310233136025398 | -4.67795915033784 |
| **Cox6a1** | 0.9700086625 | 21.66356614125 | 1.99519363890382 | 0.0874182493387425 | 0.310233136025398 | -4.6784583485244 |
| **Ndufa6** | 0.194829642500004 | 25.99078099125 | 1.98967015315848 | 0.0881252325701944 | 0.31179307156994 | -4.68610389678744 |
| **Rpl15** | 10.49102636 | 14.963684135 | 1.9895588878408 | 0.0881395320446966 | 0.31179307156994 | -4.68625785818789 |
| **Ndufb7** | 0.327004649999999 | 23.0014048975 | 1.98857208982448 | 0.0882664527642293 | 0.31179307156994 | -4.68762323402007 |
| **Abcb1a** | 0.369324720000009 | 28.2073160125 | 1.98499519705064 | 0.088728025185753 | 0.312886266817817 | -4.69257103946242 |
| **Bri3bp** | 0.237592337500004 | 20.10128832125 | 1.98405835979249 | 0.088849311241407 | 0.312886266817817 | -4.69386658987339 |
| **Rmdn3** | 0.374515955 | 25.0328348975 | 1.97866033040113 | 0.089551357245987 | 0.314874127090728 | -4.70132867944868 |
| **Ccdc136** | 0.55781209500001 | 24.1781130875 | 1.97421545314069 | 0.0901335525023212 | 0.316070090786029 | -4.70746949822475 |
| **Epha4** | 0.210677494999995 | 23.2201643925 | 1.9739559479589 | 0.0901676580529824 | 0.316070090786029 | -4.70782791457469 |
| **Rps10** | 9.840505805 | 14.1160666375 | 1.97136754028444 | 0.0905085385037667 | 0.316279291210942 | -4.71140227613868 |
| **Gabbr2** | 0.561162834999998 | 23.7711544325 | 1.97121401159148 | 0.0905287973922317 | 0.316279291210942 | -4.71161424985562 |
| **Gng2** | -0.611780740000007 | 21.990361245 | -1.9703578155709 | 0.0906418589053638 | 0.316279291210942 | -4.71279630728722 |
| **Atp5po** | -0.443176829999992 | 26.229167315 | -1.96611823091805 | 0.0912037567920443 | 0.316513641216278 | -4.71864759376336 |
| **Guk1** | 0.221673632499996 | 27.79210555875 | 1.96577033204991 | 0.0912500182313656 | 0.316513641216278 | -4.71912761102932 |
| **Hsp90ab1** | -0.194582440000005 | 28.633970015 | -1.96507854094933 | 0.0913420771209959 | 0.316513641216278 | -4.7200820552209 |
| **Tprg1l** | 0.464046522499999 | 23.83588545625 | 1.96450638729766 | 0.0914182846785426 | 0.316513641216278 | -4.72087137652046 |
| **Rap1a** | 0.503353550000003 | 22.08401922 | 1.96366508248605 | 0.0915304555267985 | 0.316513641216278 | -4.72203190542206 |
| **Mras** | -1.11004202000001 | 23.1176399625 | -1.96136534440042 | 0.091837771807916 | 0.316513641216278 | -4.72520362663672 |
| **Nckap1** | -0.233592900000001 | 20.64867982 | -1.96093522390421 | 0.0918953620686856 | 0.316513641216278 | -4.72579673197676 |

| | | | | | | |
|---|---|---|---|---|---|---|
| **Mcu** | 0.342532700000003 | 24.6044538175 | 1.95886125174391 | 0.0921735531850863 | 0.316513641216278 | -4.72865613654445 |
| **Abhd6** | -0.494329162500001 | 22.44309457375 | -1.95862826399761 | 0.09220485668758 | 0.316513641216278 | -4.72897731176949 |
| **Usp5** | 0.242149402500001 | 25.08822744125 | 1.95797149150249 | 0.0922931549896553 | 0.316513641216278 | -4.7298826258479 |
| **Mettl26** | -0.259575945000002 | 24.0092500025 | -1.95677948175388 | 0.0924536249869395 | 0.316513641216278 | -4.731525532313 |
| **Actn1** | 0.199198349999996 | 23.60734603 | 1.95641156946973 | 0.0925032093503609 | 0.316513641216278 | -4.73203256267474 |
| **Slc25a4** | -0.681046435000006 | 22.503163585 | -1.95638635801128 | 0.0925066081143249 | 0.316513641216278 | -4.73206730642601 |
| **Slc2a1** | 10.12743584 | 14.61185383 | 1.95524069808054 | 0.092661184764375 | 0.31656933123232 | -4.73364601452115 |
| **Psmb2** | -0.668914462499998 | 21.10204289125 | -1.95246052305691 | 0.093037354979945 | 0.316726602714654 | -4.73747610027747 |
| **Sirt2** | 0.273628857499993 | 23.24235489375 | 1.9500558991805 | 0.093363924162207 | 0.316726602714654 | -4.7407876997101 |
| **Acat1** | 10.847365905 | 15.6748568775 | 1.94865717470718 | 0.0935544022472726 | 0.316726602714654 | -4.74271351814953 |
| **Adss1** | 0.2151486725 | 27.75467866625 | 1.94679979096803 | 0.0938079318681097 | 0.316726602714654 | -4.74527029227249 |
| **Ndufs3** | 0.246276282499998 | 21.43181284125 | 1.94557745922363 | 0.0939751468192133 | 0.316726602714654 | -4.746952548144 |
| **Bin1** | 0.992357532499994 | 22.37756896125 | 1.94550512969734 | 0.0939850506833791 | 0.316726602714654 | -4.74705208446701 |
| **Ppm1e** | 0.209515497500004 | 27.36322577375 | 1.94527826320758 | 0.0940161214919856 | 0.316726602714654 | -4.74736428077668 |
| **Cd82** | 0.717732467499992 | 21.80624300125 | 1.94518462681324 | 0.0940289485383458 | 0.316726602714654 | -4.747493133299 |
| **Slc25a51** | 0.594640017500002 | 25.10339951875 | 1.94478845403153 | 0.0940832384463724 | 0.316726602714654 | -4.74803828685953 |
| **Eif4a2** | 0.396953107500003 | 26.60949917375 | 1.94473250013349 | 0.094090908626459 | 0.316726602714654 | -4.74811527992543 |
| **Mpc1** | -0.369277954999994 | 22.2351621025 | -1.94147840433229 | 0.0945380410698697 | 0.317310878150554 | -4.75259196452845 |
| **Ntrk2** | -0.244566040000002 | 30.2418295325 | -1.94145162382025 | 0.0945417295319692 | 0.317310878150554 | -4.75262879866635 |
| **Pc** | -0.242178185000004 | 28.286508525 | -1.9386945282935 | 0.0949222207318366 | 0.318121468894838 | -4.75642022693567 |
| **Emd** | 0.525841049999997 | 25.1223330975 | 1.93632495851428 | 0.0952504344626116 | 0.318754743399003 | -4.75967762974044 |

| | | | | | | |
|---|---|---|---|---|---|---|
| **Slc25a3** | -10.84910825 | 15.7775219225 | -1.92980402573087 | 0.0961594302875654 | 0.320911739675 | -4.76863644805637 |
| **Hsd17b4** | 0.296120460000004 | 21.314542645 | 1.92843892326854 | 0.0963507968123072 | 0.320911739675 | -4.77051089358502 |
| **Rgs17** | 0.33252435 | 22.7097845875 | 1.92551566067919 | 0.0967618522463009 | 0.320911739675 | -4.77452369217993 |
| **Syt7** | -10.6923106175 | 15.58647580875 | -1.92511994440275 | 0.0968176281368639 | 0.320911739675 | -4.77506677238385 |
| **Gpd1** | -0.2884954425 | 25.83010755875 | -1.92359978322951 | 0.0970321872344286 | 0.320911739675 | -4.77715276156297 |
| **Hsp90aa1** | -0.256342395000004 | 22.6588494225 | -1.92298337687641 | 0.0971193211759775 | 0.320911739675 | -4.77799847865182 |
| **Gprin1** | 0.224494114999999 | 24.7826118 | 1.92254930449209 | 0.0971807268282643 | 0.320911739675 | -4.77859398777745 |
| **Rpl3** | 0.32345613 | 20.925403925 | 1.92231558436337 | 0.0972138056091173 | 0.320911739675 | -4.77891461632349 |
| **Anxa7** | 0.348768784999997 | 27.3209089925 | 1.92153703086267 | 0.097324075236564 | 0.320911739675 | -4.77998259819877 |
| **Ncam1** | 9.7609589625 | 14.24125036375 | 1.92070271530519 | 0.0974423788796343 | 0.320911739675 | -4.78112694288625 |
| **Rps4x** | 0.562994077500004 | 24.50600017125 | 1.91985771923429 | 0.0975623407808345 | 0.320911739675 | -4.78228580078695 |
| **Maoa** | -0.109722065 | 26.2674883775 | -1.9186198359869 | 0.0977383414491835 | 0.320911739675 | -4.78398323032203 |
| **Ncam2** | 0.410171237499998 | 24.43655638125 | 1.91717744606624 | 0.0979438114374562 | 0.320911739675 | -4.78596071396144 |
| **Lmna** | 1.083019175 | 20.3667537425 | 1.91688147245754 | 0.0979860255397725 | 0.320911739675 | -4.78636643730509 |
| **Retreg2** | 1.1022790575 | 22.41827446125 | 1.91545373152618 | 0.0981899119834251 | 0.320911739675 | -4.78832335934029 |
| **Baiap2** | 10.70314266 | 15.652669295 | 1.91535006154525 | 0.0982047326087665 | 0.320911739675 | -4.78846543839806 |
| **Col9a1** | 11.295291645 | 16.4929001625 | 1.91429250278748 | 0.0983560461930407 | 0.320911739675 | -4.78991469669825 |
| **Lpin3** | -9.9004868575 | 14.48254386125 | -1.91349767775776 | 0.0984699187100953 | 0.320911739675 | -4.7910037664422 |
| **Ap3b2** | 9.9566543175 | 14.56750143875 | 1.91195978109176 | 0.0986906161562788 | 0.320911739675 | -4.7931106434003 |
| **Cap2** | 9.8954379075 | 14.46910337875 | 1.91095283323821 | 0.0988353817534028 | 0.320911739675 | -4.79448988418064 |
| **Cpd** | 0.647994497499997 | 22.61067639875 | 1.91092675671951 | 0.0988391334516709 | 0.320911739675 | -4.79452559917812 |

| | | | | | | |
|---|---|---|---|---|---|---|
| **Sntb1** | 0.471145744999998 | 22.53146401 | 1.90687724145301 | 0.0994234441926844 | 0.322070579206751 | -4.80007028472423 |
| **Sfxn5** | -0.839267432500002 | 25.11154468125 | -1.90629801278089 | 0.099507298069429 | 0.322070579206751 | -4.8008631127665 |
| **Kcnab1** | -0.234286645000005 | 22.169357225 | -1.9045891098814 | 0.099755096552322 | 0.322070579206751 | -4.80320181182817 |
| **Plcb1** | 10.908265915 | 16.00879297 | 1.90456315747012 | 0.0997588644200903 | 0.322070579206751 | -4.8032373242247 |
| **Cntn1** | 0.349113480000007 | 25.2367266925 | 1.90326377271247 | 0.0999476921532928 | 0.32222572864632 | -4.80501518667463 |
| **Fkbp8** | 0.355831807500003 | 26.76739655125 | 1.9013655815676 | 0.100224167919244 | 0.322645889865208 | -4.80761175073753 |
| **Jam3** | 0.618789515000007 | 21.1308463 | 1.89994849460971 | 0.100431057570493 | 0.322645889865208 | -4.80954973479778 |
| **Ddx3x** | 9.5774230425 | 14.08491118125 | 1.89929846904036 | 0.100526098731959 | 0.322645889865208 | -4.81043856380331 |
| **Ap1b1** | 0.330222909999996 | 19.8792868475 | 1.89850769743245 | 0.100641837205661 | 0.322645889865208 | -4.81151973084103 |
| **Iars2** | 10.3735088025 | 15.26425147125 | 1.89720443853076 | 0.100832868551559 | 0.322658085811168 | -4.81330131096244 |
| **Tmem65** | 0.226016009999995 | 25.358663035 | 1.89655931336857 | 0.100927562009959 | 0.322658085811168 | -4.81418308254868 |
| **Kcna2** | 11.157845425 | 16.4586149575 | 1.8891232844857 | 0.102025341510922 | 0.325712701141562 | -4.82434070174362 |
| **Stmn1** | -0.34948511999999 | 26.8572126475 | -1.88618603552754 | 0.102462174257484 | 0.326189611223717 | -4.82834986130855 |
| **Map7d2** | 0.238209982499995 | 27.69701895875 | 1.88350502307164 | 0.102862493293172 | 0.326189611223717 | -4.83200771528695 |
| **Ahcy** | -1.26595482750001 | 19.50874184125 | -1.8833778489545 | 0.102881520359402 | 0.326189611223717 | -4.83218118887426 |
| **Scg2** | 0.212510145000003 | 26.978016215 | 1.88307552492539 | 0.102926766150702 | 0.326189611223717 | -4.8325935645615 |
| **Mapre2** | 0.878310724999999 | 21.51307948 | 1.88269562048417 | 0.102983650163321 | 0.326189611223717 | -4.83311173438635 |
| **Mccc2** | -0.177595677499998 | 27.08763454125 | -1.88200767986179 | 0.10308673529047 | 0.326189611223717 | -4.8340499728499 |
| **Septin6** | 0.596250967500001 | 23.28752629875 | 1.87969813049654 | 0.103433549083393 | 0.326189611223717 | -4.83719909645069 |
| **Arf5** | -0.736497102500007 | 25.11287945625 | -1.87922062292472 | 0.10350539598446 | 0.326189611223717 | -4.83785004948602 |
| **Asph** | 0.738493447499991 | 20.17855068875 | 1.87915654297376 | 0.103515041308181 | 0.326189611223717 | -4.83793740160434 |

| | | | | | | |
|---|---|---|---|---|---|---|
| **Eef1g** | -0.317221539999998 | 20.2097037125 | -1.87835409687507 | 0.10363589989018 | 0.326189611223717 | -4.83903120209678 |
| **Apoo** | 1.08099788749999 | 25.01124364875 | 1.87599544809851 | 0.103991939886837 | 0.326189611223717 | -4.84224545003667 |
| **Slc9a1** | 11.861258095 | 17.5958167375 | 1.87481398320848 | 0.104170731182756 | 0.326189611223717 | -4.84385504887281 |
| **Ahsa1** | -9.506949135 | 14.10157139 | -1.87475432708886 | 0.104179766900009 | 0.326189611223717 | -4.84393631506324 |
| **Shisa4** | 10.540413075 | 15.610190585 | 1.87411833746262 | 0.104276143552652 | 0.326189611223717 | -4.84480264116315 |
| **Ncan** | 0.444445924999997 | 20.4245554875 | 1.87388005964417 | 0.104312274100376 | 0.326189611223717 | -4.84512719398102 |
| **Clptm1** | -10.540834645 | 15.63183648 | -1.87162235709763 | 0.104655220357272 | 0.326469508554685 | -4.84820175979365 |
| **Tmed2** | 0.749989662500003 | 21.80507516125 | 1.87141328990371 | 0.10468703332422 | 0.326469508554685 | -4.84848641496543 |
| **Septin4** | 0.335429160000004 | 23.4603324625 | 1.86869299644859 | 0.105101830243605 | 0.327109336772292 | -4.85218937434319 |
| **Syt3** | 9.8538231925 | 14.64323069875 | 1.86819451685313 | 0.105178013046923 | 0.327109336772292 | -4.85286775024076 |
| **Acad8** | 0.394952989999993 | 24.5744226325 | 1.86718805652676 | 0.105331994722677 | 0.327143739376131 | -4.85423726908508 |
| **Gng12** | 11.1541434375 | 16.60166490125 | 1.86540745610134 | 0.105604952724927 | 0.327547068817557 | -4.8566596469934 |
| **Aldoa** | 0.187797355000001 | 25.3062509275 | 1.86213581863459 | 0.106108276681245 | 0.328356117192705 | -4.86110868293115 |
| **Git1** | 0.497530585000003 | 23.85395039 | 1.86041362959276 | 0.106374163747611 | 0.328356117192705 | -4.86344971850869 |
| **Mrrf** | -10.7707071125 | 16.06155774875 | -1.86007879924633 | 0.106425933117332 | 0.328356117192705 | -4.86390479063265 |
| **Hdhd3** | 10.557299935 | 15.7443886325 | 1.85999044734945 | 0.1064395976221 | 0.328356117192705 | -4.86402486671506 |
| **Chmp5** | 11.3379089625 | 16.91450239875 | 1.85871319569331 | 0.106637328219785 | 0.328523343600387 | -4.86576054537642 |
| **Pdxp** | -0.415019277499997 | 22.51733528125 | -1.85474090363176 | 0.107254563198268 | 0.329980773065639 | -4.87115626796465 |
| **Rpl6** | -1.105754135 | 21.821825205 | -1.84952033564684 | 0.108071047237302 | 0.332046479364007 | -4.87824226127331 |
| **Timm8a1** | 0.364813872500005 | 28.45199077875 | 1.8466354347945 | 0.108524826826406 | 0.332993738077271 | -4.8821553917158 |
| **Ttyh1** | -0.358999369999999 | 26.56079411 | -1.84566705007929 | 0.108677563154624 | 0.333015986694692 | -4.88346850679148 |

| | | | | | | |
|---|---|---|---|---|---|---|
| **Sdha** | 0.260356402500001 | 25.19790345875 | 1.84284360052105 | 0.109124078134427 | 0.333189566000857 | -4.88729585406439 |
| **Thy1** | 0.500144527499991 | 25.87029689375 | 1.84268146696303 | 0.109149772790306 | 0.333189566000857 | -4.88751558061591 |
| **Rpl8** | 11.70234666 | 17.575668055 | 1.83972934960888 | 0.109618647767883 | 0.333189566000857 | -4.89151530336074 |
| **Mtco2** | 1.16726754250001 | 23.80559498625 | 1.83942289217019 | 0.109667433238536 | 0.333189566000857 | -4.89193039837936 |
| **Rab39b** | 0.466171317500002 | 23.33606051125 | 1.83923735188401 | 0.109696979945392 | 0.333189566000857 | -4.8921817013238 |
| **Hspe1** | 0.19022017 | 20.358908435 | 1.83916617135986 | 0.109708317270108 | 0.333189566000857 | -4.89227810889483 |
| **Prxl2a** | 0.244577420000002 | 24.7924782625 | 1.83888483803858 | 0.109753137948731 | 0.333189566000857 | -4.89265913806597 |
| **Lmnb2** | 0.167861857499997 | 28.39200129125 | 1.83362254923707 | 0.110594781805715 | 0.335299941130175 | -4.89978287616154 |
| **Nudt3** | 0.466096869999987 | 24.4889903275 | 1.83169573550348 | 0.110904516763742 | 0.335794231312441 | -4.90238967338972 |
| **Cttn** | -10.0347062025 | 15.13118404625 | -1.82722783143946 | 0.111625970259426 | 0.337532161061858 | -4.90843100934862 |
| **Acadl** | 0.335837377499999 | 25.41828937875 | 1.82537780858061 | 0.111926031562368 | 0.337992989770792 | -4.91093117803388 |
| **Park7** | 0.281688772500004 | 26.68338358375 | 1.82345633942026 | 0.112238507976681 | 0.338490045795288 | -4.91352705055971 |
| **Krt42** | 0.205060237499996 | 23.57413507875 | 1.82203204891345 | 0.112470676325477 | 0.338527977701052 | -4.91545068186653 |
| **Emc8** | 0.2923868725 | 26.32916045875 | 1.82156522647299 | 0.112546872446702 | 0.338527977701052 | -4.9160810629584 |
| **Psmc3** | -0.696718260000001 | 22.4816684075 | -1.819173389333 | 0.112938059586514 | 0.339201197577988 | -4.91931010864066 |
| **Rpl32** | 0.221378935000004 | 24.078811 | 1.81816390007419 | 0.113103557047958 | 0.339201197577988 | -4.92067253922421 |
| **Wfs1** | 11.21367864 | 16.98266805 | 1.817451493344 | 0.113220491472979 | 0.339201197577988 | -4.92163387419393 |
| **Lta4h** | 0.371086847500003 | 25.05136604375 | 1.81658156036109 | 0.113363440868135 | 0.339201197577988 | -4.92280761391737 |
| **Sdhc** | 1.3690939 | 22.44962677 | 1.81561313579368 | 0.113522779980603 | 0.339234521378069 | -4.9241140288313 |
| **Srsf2** | -0.621753674999994 | 20.755167385 | -1.81455032684931 | 0.113697897869612 | 0.339314847749075 | -4.92554751095313 |
| **Coro2b** | -0.177308909999997 | 31.630490785 | -1.81303029139092 | 0.113948806514796 | 0.339620856917147 | -4.92759721446935 |

| | | | | | | |
|---|---|---|---|---|---|---|
| **Tmed10** | -10.662231235 | 16.1978720125 | -1.80903706958592 | 0.114610510981455 | 0.340723616788743 | -4.93297925344922 |
| **Ache** | -0.258237449999999 | 27.846579935 | -1.80726876850276 | 0.114904715529508 | 0.340723616788743 | -4.93536132546995 |
| **Slc30a1** | -0.245088917500006 | 22.61394910375 | -1.80700064512818 | 0.114949388732926 | 0.340723616788743 | -4.93572244716592 |
| **Ca14** | 10.45477963 | 15.89764381 | 1.80656360983707 | 0.115022241049187 | 0.340723616788743 | -4.93631103020452 |
| **C2orf72** | -0.186796064999999 | 30.1886888425 | -1.80631882707868 | 0.11506306499681 | 0.340723616788743 | -4.93664067428902 |
| **Dixdc1** | -0.387440559999998 | 22.8675223475 | -1.80374941943883 | 0.115492426353599 | 0.341001584316362 | -4.94009996229583 |
| **Gtpbp1** | -10.12556125 | 15.4169671075 | -1.80291466362388 | 0.115632251034274 | 0.341001584316362 | -4.94122347720183 |
| **Vti1b** | 0.677754880000005 | 22.00254814 | 1.80236272618301 | 0.115724792331398 | 0.341001584316362 | -4.94196624723213 |
| **Cask** | 0.562679652500002 | 22.80637989625 | 1.80219557903248 | 0.115752831373444 | 0.341001584316362 | -4.94219117082378 |
| **Atp5mf** | -0.405668357500002 | 23.91156084625 | -1.80091890513127 | 0.115967210672345 | 0.341194017003853 | -4.94390891601565 |
| **Clu** | 9.3800934625 | 14.30592478125 | 1.79800881677271 | 0.116457306275424 | 0.341982788534486 | -4.94782289378472 |
| **Dnajc11** | -10.0049277375 | 15.26377846125 | -1.79755385015359 | 0.116534108805985 | 0.341982788534486 | -4.94843462040301 |
| **Cnrip1** | -0.661528079999993 | 24.899543915 | -1.79508414211343 | 0.116951871286478 | 0.342769312899807 | -4.95175437694669 |
| **Dync1h1** | 0.222944232500002 | 27.65104457625 | 1.79221398861986 | 0.117439185406475 | 0.343757410991587 | -4.95561049959361 |
| **Cyb5r3** | 9.7794446775 | 14.96045157875 | 1.78651406872054 | 0.118412765112384 | 0.346114566800642 | -4.96326235763919 |
| **Pcbp2** | 0.172362747500003 | 26.17402893375 | 1.78532745580626 | 0.118616421031086 | 0.346114566800642 | -4.96485429508966 |
| **Rpn2** | 0.189104357500007 | 32.81466087625 | 1.78398515290781 | 0.118847204250521 | 0.346114566800642 | -4.96665467268796 |
| **Trap1** | -0.170154639999996 | 23.568510335 | -1.78293117536178 | 0.119028718372026 | 0.346114566800642 | -4.96806800984295 |
| **Psma2** | 10.2540855075 | 15.73698874875 | 1.78192318051758 | 0.1192025630531 | 0.346114566800642 | -4.96941942142464 |
| **Odf2l** | 1.0021385375 | 22.34070724375 | 1.77985963429412 | 0.119559216750844 | 0.346114566800642 | -4.972185193666 |
| **Gpd2** | 0.919631749999994 | 21.9575854125 | 1.77939916648173 | 0.119638941863493 | 0.346114566800642 | -4.97280221016196 |

| | | | | | | |
|---|---|---|---|---|---|---|
| **Fam131b** | 0.243037602500003 | 24.79272928875 | 1.77922078382594 | 0.119669840668969 | 0.346114566800642 | -4.97304122428846 |
| **Pdha1** | 0.248659630000002 | 29.8673361775 | 1.77882944303028 | 0.119737654219217 | 0.346114566800642 | -4.97356555146882 |
| **Csnk2b** | 1.0267079225 | 21.99701903375 | 1.77790074982565 | 0.119898730947023 | 0.346114566800642 | -4.97480967800912 |
| **Atp8a1** | -10.3775277075 | 15.94873618875 | -1.77775820228843 | 0.119923473477042 | 0.346114566800642 | -4.97500062259882 |
| **Psmb7** | 0.272575545000009 | 22.9698451175 | 1.77647790083705 | 0.120145920606784 | 0.346114566800642 | -4.97671537128859 |
| **Echs1** | 10.8308589875 | 16.64288794875 | 1.77610853200679 | 0.120210170645046 | 0.346114566800642 | -4.97721000024016 |
| **Vim** | 0.202244504999996 | 28.2266394375 | 1.77511925855265 | 0.120382413066151 | 0.346175054658818 | -4.97853458194957 |
| **Coq9** | 0.746979929999998 | 23.5160102025 | 1.77169403638759 | 0.120980613240912 | 0.347170154714228 | -4.98311880448355 |
| **Fmnl2** | 0.715494887499997 | 23.28142040625 | 1.77083127984979 | 0.121131739738523 | 0.347170154714228 | -4.98427301333058 |
| **Slc4a10** | 0.196472597499998 | 24.97659236875 | 1.77053622075891 | 0.121183465974953 | 0.347170154714228 | -4.984667703371 |
| **Atp5f1e** | -0.680041417500004 | 23.35493887875 | -1.7676223282427 | 0.121695435619574 | 0.34797989745966 | -4.98856429231721 |
| **Srsf1** | 0.157396610000006 | 27.7268346225 | 1.76719801284682 | 0.121770160709999 | 0.34797989745966 | -4.98913152064057 |
| **Ftl1** | -0.224275664999993 | 26.52248667 | -1.76342153643531 | 0.122437169930205 | 0.349449728142443 | -4.99417785955562 |
| **Tmx4** | 0.460561729999998 | 24.2434839675 | 1.76030117646557 | 0.122990938798574 | 0.350593099514242 | -4.99834461654366 |
| **Acot7** | 1.2499370275 | 20.36364843875 | 1.75666147283056 | 0.123639911122714 | 0.352004672338176 | -5.00320160559733 |
| **Cdh13** | 0.736551502499999 | 23.74621733125 | 1.75397990029611 | 0.124120144389224 | 0.352905154589032 | -5.0067777536736 |
| **Mpp2** | -0.343255719999998 | 26.1989729675 | -1.75298092714492 | 0.124299503558889 | 0.352905154589032 | -5.00810949155413 |
| **Acot9** | 0.257583092500006 | 26.59417769125 | 1.75231767009872 | 0.124418724226015 | 0.352905154589032 | -5.00899353555712 |
| **Cspg5** | 1.142988125 | 21.2014899425 | 1.74607585084966 | 0.125546071608518 | 0.355509966021146 | -5.01730731811518 |
| **Gatd3** | 9.2059595475 | 14.32276832375 | 1.74551585623356 | 0.125647690044171 | 0.355509966021146 | -5.01805268445071 |
| **Rnaseh2a** | 0.191229844999985 | 26.2217839425 | 1.74445069839758 | 0.125841194336238 | 0.355617893624259 | -5.01947019880565 |

| | | | | | | |
|---|---|---|---|---|---|---|
| **Dmtn** | 10.70786139 | 16.656715995 | 1.74245940265813 | 0.12620371265743 | 0.355769170608547 | -5.02211938647966 |
| **Gng7** | 1.565250525 | 21.396467515 | 1.74244917419277 | 0.126205577341258 | 0.355769170608547 | -5.02213299145752 |
| **Chchd6** | 0.155675077500003 | 19.39076555625 | 1.74095071405623 | 0.126479036608542 | 0.356101494215195 | -5.02412579672377 |
| **Actr2** | 0.377188802499997 | 23.79069867875 | 1.73543650774687 | 0.127490230281921 | 0.358507539453708 | -5.03145381533984 |
| **Ly6h** | 10.0320916075 | 15.65374478125 | 1.73109540095348 | 0.128291731648383 | 0.360297591067264 | -5.03721691791278 |
| **Phb2** | 10.3485858975 | 16.18390820125 | 1.73028652855556 | 0.128441605203533 | 0.360297591067264 | -5.03829016640034 |
| **Tmem9b** | 0.717913617499999 | 23.24011207375 | 1.72824701260935 | 0.128820243487711 | 0.360917426368873 | -5.04099547273567 |
| **Calm2** | 0.146940320000006 | 25.8681816225 | 1.72536169879486 | 0.12935772509289 | 0.361665960687573 | -5.04482068172981 |
| **Npm1** | 10.4171897275 | 16.34261065875 | 1.72511694220102 | 0.129403417126746 | 0.361665960687573 | -5.04514505987715 |
| **Slc4a3** | -0.286667914999995 | 24.52143715 | -1.72375603676076 | 0.129657756733653 | 0.361934884345526 | -5.04694837065821 |
| **Rpl19** | 0.215926279999998 | 23.71453528 | 1.71933262574566 | 0.130487745002012 | 0.363808097819251 | -5.05280611301137 |
| **Ptges3** | -0.568881582499998 | 22.10896405875 | -1.71839641212167 | 0.130664060014235 | 0.363856488287814 | -5.05404518611421 |
| **Ncdn** | 0.154260624999992 | 29.79651519 | 1.71602452958316 | 0.131111768006317 | 0.364659583191323 | -5.05718323317746 |
| **Rps8** | 0.611868752500001 | 21.32245907625 | 1.71474947375588 | 0.131353046513258 | 0.364887285763163 | -5.05886948838559 |
| **Fxyd1** | -0.371631515000004 | 20.67669028 | -1.70595795886127 | 0.133028204478662 | 0.369092800062615 | -5.07048340658871 |
| **Map1b** | 0.308067317500004 | 22.71564555625 | 1.69482700694119 | 0.135178278467993 | 0.373128946957797 | -5.08515529305624 |
| **Asphd2** | 0.217864742499991 | 26.02366866125 | 1.69452684179079 | 0.135236713438097 | 0.373128946957797 | -5.0855504365773 |
| **Tomm40** | -0.350207812500003 | 24.35966435875 | -1.69377422408947 | 0.135383335537041 | 0.373128946957797 | -5.08654107950087 |
| **Ndrg3** | 0.327170937500014 | 21.72309785375 | 1.69366583790132 | 0.135404463344024 | 0.373128946957797 | -5.08668373029401 |
| **Rtn1** | -0.788842095000007 | 23.636674935 | -1.69261053166701 | 0.135610338659682 | 0.373128946957797 | -5.0880724713931 |
| **Tm9sf2** | 0.412909167500004 | 22.60816582625 | 1.69217646725574 | 0.135695104671767 | 0.373128946957797 | -5.08864358609618 |

| | | | | | | |
|---|---|---|---|---|---|---|
| **Fis1** | 0.174419537499993 | 27.82413669625 | 1.69194559091936 | 0.135740211700502 | 0.373128946957797 | -5.08894733567029 |
| **Glul** | -0.20994915999999 | 23.561755895 | -1.6912532078214 | 0.135875570063019 | 0.373128946957797 | -5.08985816495239 |
| **Rogdi** | 0.535505507500009 | 23.37834283625 | 1.69087278208662 | 0.135949996401399 | 0.373128946957797 | -5.09035855335245 |
| **Scn1b** | -0.662061157499998 | 21.93885060125 | -1.68963932314389 | 0.136191575868559 | 0.373344331931894 | -5.09198066940622 |
| **Cpt1a** | 0.206339635000006 | 27.2184708875 | 1.68729261660507 | 0.13665231532207 | 0.374090551179871 | -5.09506554616473 |
| **Prkar1b** | -0.691423735000004 | 23.80199451 | -1.68658950862704 | 0.136790647154894 | 0.374090551179871 | -5.09598949695443 |
| **Spock2** | -0.275381544999995 | 25.349872115 | -1.68562691145511 | 0.1369802470473 | 0.374162035192447 | -5.09725419849506 |
| **Stim1** | 0.202590332500005 | 26.71110504375 | 1.68136038844253 | 0.137823613473189 | 0.376016985983468 | -5.10285635089877 |
| **Pdhx** | 0.259702239999999 | 25.2936030325 | 1.67822741000487 | 0.138446040633533 | 0.377265460726378 | -5.10696656603186 |
| **Ssr1** | 0.225261627500011 | 24.08846684375 | 1.67457163322484 | 0.139175695695247 | 0.378380688729797 | -5.11175883914838 |
| **Vps45** | 9.9680648725 | 15.94397635125 | 1.67415485976048 | 0.139259110019065 | 0.378380688729797 | -5.11230491612022 |
| **Lsamp** | 0.58341579 | 22.22881124 | 1.67369496130093 | 0.139351210397212 | 0.378380688729797 | -5.1129074352457 |
| **Clpp** | 0.284453580000001 | 27.2200594825 | 1.66600846390936 | 0.140899078132237 | 0.381553938048259 | -5.1229678925866 |
| **Ptprd** | -0.191965787500003 | 26.79516449125 | -1.66566792071221 | 0.14096802946146 | 0.381553938048259 | -5.12341318531108 |
| **Sfxn1** | -0.151375074999997 | 23.9366969125 | -1.66478665922537 | 0.141146610468276 | 0.381553938048259 | -5.1245653499336 |
| **Sult4a1** | 0.143587927499993 | 26.73002405375 | 1.66458933800939 | 0.141186625394004 | 0.381553938048259 | -5.1248232950392 |
| **Myh9** | 0.172235012499996 | 25.74754749625 | 1.66159680278226 | 0.141794798620553 | 0.382745629767036 | -5.12873373881325 |
| **Rps3** | -0.919910067499995 | 24.27110794125 | -1.66051664144528 | 0.14201492671732 | 0.382888300654823 | -5.13014452736325 |
| **Ddx39b** | 0.148628135000003 | 26.369995135 | 1.65846805480234 | 0.142433298105437 | 0.383564493368641 | -5.13281915305051 |
| **Ogdh** | 0.192158065000005 | 27.88048969 | 1.65497576366469 | 0.143149190792572 | 0.38503936277814 | -5.13737560041547 |
| **Scn2a** | 0.190643572500001 | 25.32149717625 | 1.65184828032523 | 0.143793178904051 | 0.386317589802082 | -5.14145277426338 |

| | | | | | | |
|---|---|---|---|---|---|---|
| **Hspa1b** | 0.52839861750001 | 25.80107087625 | 1.64788933849741 | 0.144612290072947 | 0.387737631226022 | -5.14660938514389 |
| **Tubb2a** | -0.370973849999999 | 27.82933922 | -1.64765687744064 | 0.14466052296506 | 0.387737631226022 | -5.1469120139315 |
| **Kcna10** | 0.271822960000002 | 26.3260550825 | 1.64594648658082 | 0.145015874632277 | 0.38823548191027 | -5.14913814448177 |
| **Adcy2** | 0.2285982225 | 22.54524142375 | 1.64328123737544 | 0.145571246303911 | 0.389267035969221 | -5.1526051652773 |
| **Bcan** | 0.848327994999991 | 23.8927262075 | 1.64145153388641 | 0.145953668886542 | 0.389628913294195 | -5.15498395084799 |
| **Rps17** | 0.308334609999996 | 24.7257860975 | 1.64029982513915 | 0.14619486957585 | 0.389628913294195 | -5.15648071993002 |
| **Cbln3** | 0.265375532499998 | 23.03427020875 | 1.64019315306505 | 0.146217228711102 | 0.389628913294195 | -5.15661932980848 |
| **Auh** | 1.5319766425 | 21.54875456875 | 1.63917636365639 | 0.146430515605719 | 0.389743546769175 | -5.15794036091067 |
| **Pycr2** | 0.862757665 | 22.365946885 | 1.62946261130266 | 0.148482912189645 | 0.394496650296107 | -5.17054349602496 |
| **Gapdh** | -0.296040352499993 | 24.62412951875 | -1.6290113785938 | 0.148578905603191 | 0.394496650296107 | -5.17112819008614 |
| **Ndufa11** | -0.287147542499994 | 20.83649021375 | -1.62814100393702 | 0.148764230105856 | 0.394496650296107 | -5.17225580445487 |
| **Spryd7** | 0.561447657500004 | 23.60103557375 | 1.6274773614132 | 0.148905681894206 | 0.394496650296107 | -5.173115417241 |
| **Hadhb** | 0.239549132500002 | 25.27349703875 | 1.62572860675503 | 0.14927902398706 | 0.395028538620094 | -5.17537986599342 |
| **Prepl** | 0.234147997499996 | 19.49333535375 | 1.62399925823368 | 0.149649086302243 | 0.395550529498654 | -5.17761817956364 |
| **Stxbp1** | 0.197334157499998 | 30.17352548375 | 1.62066966965923 | 0.150364006421623 | 0.396981788580272 | -5.18192487192872 |
| **Hibadh** | 0.742310890000006 | 23.2361379375 | 1.61532280055979 | 0.151518770572211 | 0.399403087156704 | -5.18883300475901 |
| **Fth1** | 0.177758727499992 | 22.67252292125 | 1.61480936639014 | 0.151630092939788 | 0.399403087156704 | -5.18949584909665 |
| **Septin7** | 0.268155345 | 25.8467072025 | 1.61273420085519 | 0.15208080902234 | 0.399794464219018 | -5.19217397593531 |
| **Tmem109** | -0.668786865000001 | 21.7751080475 | -1.61251728678738 | 0.152127994030041 | 0.399794464219018 | -5.19245383184944 |
| **Itpr1** | 0.533370877499991 | 25.77631023875 | 1.61079764220126 | 0.152502550851153 | 0.400306712495459 | -5.19467189543674 |
| **Aldh7a1** | 0.15918285250001 | 26.24882900625 | 1.61001786808885 | 0.152672678029068 | 0.400306712495459 | -5.19567734429698 |

| | | | | | | |
|---|---|---|---|---|---|---|
| **Sytl4** | -0.582922085 | 23.3978176525 | -1.60779587694693 | 0.153158434750412 | 0.401120889180427 | -5.19854125792126 |
| **Tf** | 0.205765535000001 | 29.5734828575 | 1.60453395033957 | 0.153874152889368 | 0.402018389679128 | -5.20274245875193 |
| **Omp** | 0.877068419999997 | 24.8593153825 | 1.60416469768853 | 0.15395536936386 | 0.402018389679128 | -5.20321780636802 |
| **Me2** | 0.224744995000002 | 24.152147185 | 1.60383456390327 | 0.15402801561756 | 0.402018389679128 | -5.20364275538228 |
| **Kif2a** | -0.203255980000005 | 27.682916675 | -1.59579885906098 | 0.155806182161505 | 0.405674461715673 | -5.21397464478848 |
| **Ppp2ca** | 0.183235864999993 | 25.5032012375 | 1.59490008226551 | 0.156006253018854 | 0.405674461715673 | -5.21512884199501 |
| **Map2** | -0.397778880000008 | 21.8257827925 | -1.59466331416424 | 0.156058998284253 | 0.405674461715673 | -5.2154328492426 |
| **Cpox** | 0.1719086875 | 27.65776182875 | 1.59357580233183 | 0.15630147896372 | 0.405674461715673 | -5.21682894756758 |
| **Cplx2** | -0.309473392499999 | 24.86805807375 | -1.59346924648683 | 0.156325256427197 | 0.405674461715673 | -5.2169657167461 |
| **Ppib** | 1.1851511675 | 23.79500024125 | 1.59272172206572 | 0.15649215801439 | 0.405674461715673 | -5.21792508526988 |
| **Grm2** | -0.232570167500008 | 25.65815600125 | -1.59106989327292 | 0.156861555043732 | 0.406172058252377 | -5.22004433412311 |
| **Epn1** | 0.200278672499998 | 26.06844285125 | 1.58957943808251 | 0.157195561450686 | 0.406576994531773 | -5.22195571800759 |
| **Surf1** | -0.550465177499998 | 25.55166715125 | -1.5863319970026 | 0.157925598553532 | 0.408004170529385 | -5.2261175623602 |
| **Hnrnpdl** | 0.261451654999995 | 22.8824410525 | 1.58297196205147 | 0.158684268913714 | 0.409259723284701 | -5.23041975351496 |
| **Prkar2b** | 0.122018802499994 | 31.25071518875 | 1.58259687071266 | 0.158769171811627 | 0.409259723284701 | -5.23089977052367 |
| **Faf2** | 0.699353079999987 | 22.8170075525 | 1.5778058932063 | 0.159857345016786 | 0.411482506530023 | -5.237026506165 |
| **Mff** | 0.555804905000006 | 21.3535511175 | 1.57659581902916 | 0.160133283501492 | 0.411482506530023 | -5.23857265074716 |
| **Nlgn2** | 0.350657730000005 | 23.90829765 | 1.576021414137 | 0.160264422423192 | 0.411482506530023 | -5.2393063983804 |
| **Sdhb** | 0.214006552499995 | 28.49942975875 | 1.57564442722677 | 0.160350544265959 | 0.411482506530023 | -5.23978789848415 |
| **Kif5c** | 0.579983962500002 | 21.43690943875 | 1.57387478845932 | 0.160755389335297 | 0.411538872597143 | -5.24204745185164 |
| **Exog** | -0.246371687500002 | 27.06577691125 | -1.57343201972544 | 0.160856831129899 | 0.411538872597143 | -5.2426126210962 |

| | | | | | | |
|---|---|---|---|---|---|---|
| **Ak3** | 0.472414424999997 | 23.2399659325 | 1.57273961102549 | 0.161015586584861 | 0.411538872597143 | -5.24349629897345 |
| **Cacna2d1** | 0.265398839999996 | 25.672419115 | 1.57188345789973 | 0.161212086529937 | 0.411538872597143 | -5.24458871272874 |
| **Rpl10a** | 0.223499829999994 | 23.8113667975 | 1.57162496355845 | 0.161271458593114 | 0.411538872597143 | -5.24491848774854 |
| **Atp2b4** | -0.306689165000002 | 24.651789385 | -1.5700001316602 | 0.161645121834327 | 0.412033055544291 | -5.24699081521769 |
| **Serpinb1a** | 0.719446167499996 | 24.75119905625 | 1.56752868901976 | 0.162215021283387 | 0.413025788340014 | -5.25014107119472 |
| **Eci1** | 11.18190288 | 10.8569297325 | 1.56590745413715 | 0.162589879886365 | 0.413124199045601 | -5.25220638453792 |
| **Dlat** | 0.243018655 | 26.13998415 | 1.56580050500323 | 0.162614636671073 | 0.413124199045601 | -5.25234259460539 |
| **Slc12a5** | 0.232966800000007 | 24.8486371575 | 1.56104016421579 | 0.163720115362848 | 0.415471556613702 | -5.25840108547457 |
| **Ggt7** | -0.196683787499992 | 26.67706543625 | -1.55328310870967 | 0.165536436007171 | 0.419313204466568 | -5.26825551782068 |
| **Hoxa2** | 0.232143242500005 | 24.08380749625 | 1.55301170440919 | 0.165600321903791 | 0.419313204466568 | -5.26859989918635 |
| **Isoc2a** | -0.504237912500002 | 24.10604043125 | -1.5513002966604 | 0.166003695974619 | 0.419870121641881 | -5.27077084662221 |
| **Tacc1** | 0.357814507499988 | 24.88190650375 | 1.54882762172267 | 0.166588103350801 | 0.420883188267089 | -5.27390553167286 |
| **Coq7** | -0.351129732499999 | 23.52839587375 | -1.54730212442277 | 0.166949596723395 | 0.421331451929273 | -5.2758383029697 |
| **Cyp2d26** | 0.255010582499992 | 23.92921051125 | 1.54275142163461 | 0.168032275085391 | 0.423118660804573 | -5.28159871649455 |
| **Yars1** | 0.381988122500001 | 23.21030913875 | 1.54192924317794 | 0.168228573347118 | 0.423118660804573 | -5.2826386164285 |
| **Map4** | 0.176732617499997 | 22.32502956375 | 1.54154891859828 | 0.168319448878308 | 0.423118660804573 | -5.28311956796698 |
| **Fgf14** | 0.211253307499998 | 22.86233767375 | 1.54122383749343 | 0.16839716032895 | 0.423118660804573 | -5.28353061596895 |
| **Spr** | 0.101056527499999 | 21.96003984875 | 1.53940850733272 | 0.168831728486091 | 0.423552946764328 | -5.2858252652862 |
| **Stip1** | 0.294962989999995 | 25.7675636925 | 1.53676770484067 | 0.169465751945892 | 0.423552946764328 | -5.28916109664666 |
| **Apool** | 0.934914007499998 | 21.59769003625 | 1.5361992237196 | 0.169602523966964 | 0.423552946764328 | -5.28987884613342 |
| **Sympk** | 0.371265887500009 | 26.57935752375 | 1.53604973714532 | 0.169638506137614 | 0.423552946764328 | -5.29006756341227 |

| | | | | | | |
|---|---|---|---|---|---|---|
| **Fxyd6** | 0.197081817499996 | 22.91899886625 | 1.53599999455648 | 0.169650480989369 | 0.423552946764328 | -5.29013035836106 |
| **Gk** | 0.323952162499996 | 23.59314581875 | 1.53525466755001 | 0.169830001659093 | 0.423552946764328 | -5.29107114399773 |
| **Dctn1** | 0.332665975000001 | 24.3982822 | 1.53426258494442 | 0.17006922790941 | 0.423552946764328 | -5.2923230646265 |
| **Ccdc51** | 0.194184089999993 | 26.14905894 | 1.53387526953852 | 0.170162707622855 | 0.423552946764328 | -5.29281171971392 |
| **Impa1** | 0.320216312500005 | 25.79949910375 | 1.5335744342917 | 0.170235347760237 | 0.423552946764328 | -5.29319122757397 |
| **Cacna2d2** | 0.167428774999998 | 26.752250805 | 1.53166701938876 | 0.170696580244269 | 0.424239383473541 | -5.2955966463439 |
| **Rack1** | -0.340095835000003 | 24.0482948475 | -1.52399964506377 | 0.172562261326397 | 0.427613875494839 | -5.30525166690976 |
| **Acsbg1** | -0.229500697499997 | 22.31452145375 | -1.52378493377462 | 0.172614775429109 | 0.427613875494839 | -5.30552170944975 |
| **Cpne5** | -0.229500697499997 | 22.31452145375 | -1.52378493377462 | 0.172614775429109 | 0.427613875494839 | -5.30552170944975 |
| **Rhoa** | -0.308742020000004 | 23.258151005 | -1.52108495704533 | 0.173276391112708 | 0.428684302299558 | -5.30891593311138 |
| **Homer1** | 0.239918442499999 | 25.40746953125 | 1.52011853239865 | 0.173513774463038 | 0.428684302299558 | -5.31013016098892 |
| **Cep164** | 0.854155814999999 | 22.2317440625 | 1.51973235543788 | 0.173608714823805 | 0.428684302299558 | -5.31061525595789 |
| **Rps2** | 7.1778993375 | 18.52736199875 | 1.5181854229617 | 0.173989501832872 | 0.428834431761032 | -5.31255784244363 |
| **Lman1** | 0.603095700000001 | 21.690008035 | 1.51796344566388 | 0.174044205813018 | 0.428834431761032 | -5.31283651690112 |
| **Tmx3** | -0.121175517499992 | 28.55953741375 | -1.51539671239714 | 0.174677898466631 | 0.429740303977155 | -5.31605743042518 |
| **Sema4a** | -0.166842077500004 | 22.43556695625 | -1.5149542889422 | 0.17478734076135 | 0.429740303977155 | -5.31661235088399 |
| **Rgs6** | 0.128766399999996 | 27.531963015 | 1.513449872276 | 0.175159959298462 | 0.429808882161341 | -5.31849872264873 |
| **Rab11b** | -0.690726340000001 | 23.064482485 | -1.51332558249571 | 0.175190776346235 | 0.429808882161341 | -5.31865452815178 |
| **Ttyh3** | -0.432441644999997 | 24.66903927 | -1.51248695717826 | 0.175398839467065 | 0.429858611927315 | -5.31970564056549 |
| **Dnajb11** | -0.224154014999996 | 19.4262157525 | -1.5111656646558 | 0.175727112353348 | 0.430202524253278 | -5.32136114951665 |
| **Scn1a** | 0.247416072499991 | 26.72234250375 | 1.50586607481619 | 0.177049454829826 | 0.432976711651145 | -5.32799426015465 |

| | | | | | | |
|---|---|---|---|---|---|---|
| **Rps9** | 0.858621769999996 | 24.2450084025 | 1.50250790354433 | 0.177892090620664 | 0.43411513968319 | -5.33219161709009 |
| **Abr** | 0.271044762500001 | 25.78893417375 | 1.50176296671116 | 0.178079507801312 | 0.43411513968319 | -5.3331220941647 |
| **Sdcbp** | 0.125152767500005 | 32.14867352375 | 1.5017453975533 | 0.178083930171479 | 0.43411513968319 | -5.33314403654538 |
| **Astn1** | 0.265177549999997 | 20.4443542525 | 1.49944433696391 | 0.178664004322438 | 0.434611179491491 | -5.33601678839765 |
| **Lpgat1** | 0.253067512500003 | 25.54118524125 | 1.49934422066484 | 0.178689281829347 | 0.434611179491491 | -5.33614172970728 |
| **Acot11** | 0.542059510000005 | 24.279453605 | 1.49804940054744 | 0.179016494730651 | 0.434611179491491 | -5.3377572501064 |
| **Dnaja2** | -1.2037992275 | 24.62761133625 | -1.49792921368582 | 0.179046894827643 | 0.434611179491491 | -5.33790717033505 |
| **Cyfip1** | -0.406347612499996 | 21.47868634125 | -1.49508768061155 | 0.17976700936965 | 0.435896911490601 | -5.34144997039672 |
| **Naxe** | 0.204450247500002 | 25.16780476125 | 1.49229368266596 | 0.180477657250437 | 0.437156992006614 | -5.34493029997928 |
| **Lrrc8a** | 0.389862954999995 | 21.63022176 | 1.49062127364382 | 0.180904257932174 | 0.437676712001095 | -5.34701200270897 |
| **Cpe** | 0.1702861375 | 27.05863421375 | 1.48971645618995 | 0.181135443789813 | 0.437676712001095 | -5.3481377815209 |
| **Isyna1** | 0.376860892499998 | 23.14126491625 | 1.4892065491349 | 0.181265846647898 | 0.437676712001095 | -5.34877206277965 |
| **Dpysl2** | -0.207169272500003 | 27.86479694875 | -1.48733540433628 | 0.18174510479643 | 0.438110643419507 | -5.3510986939583 |
| **Ctsd** | -0.940710877499999 | 21.55790750125 | -1.48701082224193 | 0.181828357906742 | 0.438110643419507 | -5.35150214145229 |
| **Copa** | -0.393198174999988 | 23.56265734 | -1.48460219326612 | 0.182447242629676 | 0.43913957768594 | -5.35449465129101 |
| **Pitpnm1** | -0.178590942499998 | 24.36409865875 | -1.48326504873653 | 0.18279164357009 | 0.439506378289849 | -5.35615490205991 |
| **Ctbp1** | -0.292289882500004 | 23.56144631125 | -1.48095871317048 | 0.183387066480835 | 0.440475335964985 | -5.35901679887956 |
| **Vma21** | 0.864625055000001 | 22.8700844875 | 1.47986814156063 | 0.18366923251415 | 0.440690642793384 | -5.36036930416839 |
| **Sv2b** | -0.297364580000007 | 24.3429896 | -1.47811821951694 | 0.184122820178474 | 0.441316372134584 | -5.36253848921699 |
| **Prkar1a** | -0.181704660000005 | 27.2814068425 | -1.47589874815303 | 0.184699584194217 | 0.442235719895985 | -5.36528788648703 |
| **Ap2a1** | 0.268233094999996 | 24.613587 | 1.47359999362533 | 0.185298684099619 | 0.443206570432631 | -5.36813332572267 |

| | | | | | | |
|---|---|---|---|---|---|---|
| **Cep112** | 0.183180257499991 | 27.91782903125 | 1.47056112675633 | 0.18609338428923 | 0.444642752231782 | -5.37189148685205 |
| **Snx3** | 0.219922127499999 | 26.12305250125 | 1.46916754768993 | 0.186458857148309 | 0.445020281592675 | -5.37361362387113 |
| **Gnb4** | 0.136075772499996 | 20.90268202625 | 1.46834954532531 | 0.186673685904846 | 0.445020281592675 | -5.37462410146194 |
| **Hmgcl** | -0.264225490000001 | 22.29594406 | -1.46773724207383 | 0.18683463984734 | 0.445020281592675 | -5.37538029416747 |
| **Lgi1** | -0.213867957499996 | 25.69062252625 | -1.46609651418859 | 0.187266553704865 | 0.445585386102325 | -5.37740580647648 |
| **Pcbp1** | 0.364091619999989 | 23.470656405 | 1.4623569786197 | 0.188254358480891 | 0.447103179699553 | -5.38201807332449 |
| **Alcam** | 0.621946592500002 | 23.35678917125 | 1.46220315682479 | 0.188295091843761 | 0.447103179699553 | -5.38220766666284 |
| **Tomm40l** | -0.307338674999997 | 24.198749295 | -1.45170182426626 | 0.19109490396093 | 0.452897227091907 | -5.39512715833249 |
| **Septin3** | -0.389199997499993 | 24.80983675625 | -1.45156757415288 | 0.191130939873649 | 0.452897227091907 | -5.39529201538533 |
| **Timm29** | 0.190476947500002 | 25.14595311125 | 1.44974047049781 | 0.191621989566111 | 0.453591245208715 | -5.39753489527378 |
| **Ank2** | -0.493202797500004 | 22.61344182125 | -1.44871084470347 | 0.191899212859927 | 0.45377820065741 | -5.3987981860394 |
| **Prkcsh** | -0.187641667499989 | 28.45781959875 | -1.4470340597028 | 0.192351458064599 | 0.454378212084485 | -5.40085451862985 |
| **Grik2** | 6.87042048 | 18.4798087175 | 1.44206077958752 | 0.193698471859348 | 0.45700909769186 | -5.40694632081529 |
| **Pccb** | -0.212539557499994 | 23.52769204125 | -1.44144996582161 | 0.193864497098644 | 0.45700909769186 | -5.40769376531865 |
| **Rpl23** | 0.6797150875 | 23.80485786125 | 1.43920683366442 | 0.194475305524089 | 0.457977339860741 | -5.41043725221383 |
| **Abcd3** | -0.329079414999995 | 23.9387185775 | -1.436394841619 | 0.195243468408545 | 0.459313771004274 | -5.41387336511943 |
| **Hnrnpa0** | 0.144657134999992 | 28.9311072925 | 1.43459870760635 | 0.195735556584383 | 0.459998654026337 | -5.41606632592574 |
| **Ugt8** | 0.505257954999998 | 22.9473041475 | 1.43146785973826 | 0.196595990819129 | 0.461546895369216 | -5.41988547168453 |
| **Clip2** | 0.143136500000008 | 26.6359394025 | 1.42854327651942 | 0.197402814940556 | 0.462966232990711 | -5.42344907547383 |
| **Vsnl1** | 0.247875385000011 | 23.5652705325 | 1.42533351459089 | 0.198291743057481 | 0.463392658256778 | -5.42735578061434 |

| | | | | | | |
|---|---|---|---|---|---|---|
| **Gltp** | -0.206034719999998 | 23.89429969 | -1.4252931109049 4 | 0.198302955580442 | 0.463392658256778 | -5.42740492789772 |
| **Epb41l2** | 0.766096807500006 | 22.86597214625 | 1.42513913581909 | 0.1983456908009 | 0.463392658256778 | -5.42759221740642 |
| **Clic4** | 0.227493369999998 | 24.2026947875 | 1.42496363589634 | 0.198394410262841 | 0.463392658256778 | -5.42780567597259 |
| **Pclo** | 0.142056365000002 | 28.376364055 | 1.42287469761697 | 0.198975134650672 | 0.464232549505762 | -5.43034536777868 |
| **Wdr33** | 0.126479822500002 | 30.39247095375 | 1.42221223828795 | 0.199159617131786 | 0.464232549505762 | -5.431150364327 |
| **Ywhah** | 0.222831777499994 | 23.35717696375 | 1.41997653916949 | 0.199783352758438 | 0.465212710543301 | -5.43386564758905 |
| **H1-4** | 0.230739057500003 | 20.82681802125 | 1.41604577766159 | 0.200884248210178 | 0.467300857879165 | -5.43863413668104 |
| **Afg3l2** | 0.168819877499999 | 30.75657432625 | 1.41413130080715 | 0.201422408685714 | 0.468077049219899 | -5.44095409538556 |
| **Lamp2** | -0.370279924999995 | 25.02913181 | -1.41320325944111 | 0.201683746799065 | 0.468209022741441 | -5.44207809486683 |
| **Mtnd1** | 0.178379652500006 | 25.06029110375 | 1.41116594813 | 0.202258524164457 | 0.46839440182044 | -5.44454421466953 |
| **Gad1** | -0.952887847500001 | 22.03417431375 | -1.41089310223634 | 0.202335612346485 | 0.46839440182044 | -5.44487434490739 |
| **Amph** | -0.338431382500001 | 24.10929472375 | -1.41074493686488 | 0.202377485102846 | 0.46839440182044 | -5.44505360355968 |
| **Psmd12** | -0.26048673 | 24.1982447925 | -1.40770659630847 | 0.203237859424586 | 0.46990367554402 | -5.44872734417075 |
| **Mlec** | 0.630943630000001 | 22.43412902 | 1.40699383717212 | 0.203440167087865 | 0.46990367554402 | -5.4495885489318 |
| **Cmtr1** | -0.198143680000001 | 22.8080914275 | -1.40194777293354 | 0.204877592528314 | 0.472746783565838 | -5.45567887695657 |
| **Cct5** | 0.813271935000003 | 22.73261039 | 1.39904118095688 | 0.205709684187863 | 0.474188788626402 | -5.45918164762403 |
| **Ksr2** | 0.472657987499993 | 25.11781558875 | 1.39771957167173 | 0.206089029187366 | 0.474585299607525 | -5.46077304210329 |
| **Ldha** | 0.342656637500003 | 23.06560924375 | 1.39301494657691 | 0.207444486879622 | 0.477226563283874 | -5.46643145229508 |
| **Rps3a** | 6.21015773 | 17.1909683525 | 1.38978666384147 | 0.208379190831305 | 0.478895550012909 | -5.47030824132109 |
| **Rab21** | -0.596441132500004 | 22.62528706625 | -1.38462404545371 | 0.209881760411105 | 0.480992180049807 | -5.47649777754502 |
| **Tmem43** | 0.150035465000002 | 25.2102191425 | 1.38447430554752 | 0.209925485589044 | 0.480992180049807 | -5.47667711553035 |

| | | | | | | |
|---|---|---|---|---|---|---|
| **Ech1** | -1.0186378925 | 20.85946580875 | -1.38430414109355 | 0.209975184736068 | 0.480992180049807 | -5.47688090239848 |
| **Me3** | -0.452870882500001 | 23.42200487125 | -1.3830533679876 | 0.210340813918666 | 0.480992180049807 | -5.47837839322349 |
| **Txn** | 0.19240709 | 20.582550355 | 1.38271421786411 | 0.210440052783974 | 0.480992180049807 | -5.47878431400231 |
| **Cct8** | 6.5158195975 | 18.28313240375 | 1.38098838589623 | 0.210945695118564 | 0.480992180049807 | -5.48084907806296 |
| **Rtn4ip1** | -0.193094752500006 | 26.67756189375 | -1.38076569171834 | 0.211011019794776 | 0.480992180049807 | -5.48111540382599 |
| **Actr3** | 0.174680032500003 | 26.45890026375 | 1.38066378775936 | 0.211040918105639 | 0.480992180049807 | -5.48123726557927 |
| **Gh1** | -0.67721912999999 | 22.050699785 | -1.38018080831535 | 0.211182674071671 | 0.480992180049807 | -5.4818147690865 |
| **Dnaja3** | -0.239982234999992 | 25.5865687025 | -1.37416252769382 | 0.212956172294514 | 0.48441805750093 | -5.4890015826419 |
| **Cct7** | 0.260941159999998 | 24.6837265825 | 1.37364233602826 | 0.213110084710981 | 0.48441805750093 | -5.48962196532409 |
| **Rab1b** | 0.5602193175 | 23.40600436875 | 1.37159272259572 | 0.213717478682675 | 0.485316774508574 | -5.49206508146492 |
| **Tubb2b** | 0.304189632499998 | 22.03093236375 | 1.37008181050622 | 0.214166214324791 | 0.485853780564368 | -5.49386478139224 |
| **Actc1** | 0.1672950325 | 20.62216685875 | 1.36848158863086 | 0.214642386069412 | 0.486451902686024 | -5.4957696646552 |
| **Tomm70** | 0.268846212500012 | 24.72520980375 | 1.36647845382414 | 0.215239774263049 | 0.487323287129692 | -5.49815243154355 |
| **Emilin3** | 0.25132846999999 | 19.4880813525 | 1.36358966314461 | 0.216103883891696 | 0.48853007257858 | -5.50158528958863 |
| **Kcnk1** | -6.80336642000001 | 19.29795104 | -1.36327019079278 | 0.21619963456623 | 0.48853007257858 | -5.50196468243523 |
| **Capns1** | 0.171752232500008 | 25.69630627125 | 1.36218557723605 | 0.216524989974649 | 0.488782743640997 | -5.50325235770035 |
| **Ndufs7** | 0.855839782500006 | 20.50923066125 | 1.3605829374895 | 0.217006533140146 | 0.489387147150535 | -5.50515399852398 |
| **Fah** | 0.2469298575 | 25.92326975875 | 1.35934010417402 | 0.217380618060753 | 0.489748262540418 | -5.50662784454372 |
| **Kgd4** | 0.150060777500009 | 24.38489094375 | 1.35314109433811 | 0.219255010909348 | 0.493326047848354 | -5.51396783919375 |
| **Gabrb2** | 0.111404197500008 | 29.27044477125 | 1.35232803853582 | 0.219501910903728 | 0.493326047848354 | -5.51492914889365 |
| **Mapt** | 0.171124102499995 | 21.82479460125 | 1.35162518734606 | 0.219715543008354 | 0.493326047848354 | -5.51575989727875 |

| | | | | | | |
|---|---|---|---|---|---|---|
| **Idh2** | 0.611584097500007 | 23.33483499125 | 1.35124643170362 | 0.219830742160472 | 0.493326047848354 | -5.5162074739965 |
| **Sfxn3** | -0.384848077499999 | 21.67442226625 | -1.34825965249283 | 0.220741045602896 | 0.494483825881444 | -5.51973448404881 |
| **Pdpr** | -0.649996380000005 | 23.314297915 | -1.34813630279165 | 0.220778711249819 | 0.494483825881444 | -5.51988004987049 |
| **Mtco1** | 6.025015105 | 17.20821265 | 1.34728622108134 | 0.221038443533614 | 0.494581619988702 | -5.52088303261185 |
| **Osbpl8** | 0.281199357499993 | 25.57697183125 | 1.34577366137813 | 0.221501254452123 | 0.495133175235262 | -5.52266676686906 |
| **Glrx5** | 6.8125948775 | 19.52691466125 | 1.34500732868853 | 0.221736061803639 | 0.495174483383932 | -5.52357005660851 |
| **Aldh3b1** | -0.207731447500002 | 22.75762158125 | -1.34270817922393 | 0.222441847238229 | 0.495933683081809 | -5.52627835545729 |
| **Plpp3** | 0.100979880000001 | 26.2987764775 | 1.34248863445365 | 0.222509345795115 | 0.495933683081809 | -5.52653683232968 |
| **Cbr1** | -0.226567889999998 | 21.0359214675 | -1.34161421717575 | 0.222778362524103 | 0.496050264414078 | -5.52756607292506 |
| **Rhot1** | -0.461906750000001 | 20.759220715 | -1.33912439401515 | 0.223545932453503 | 0.49620415060819 | -5.53049465413067 |
| **Htra2** | -6.35521775 | 17.8319798725 | -1.33883432133414 | 0.223635508131668 | 0.49620415060819 | -5.53083564235565 |
| **Pls3** | 1.137365465 | 23.519087865 | 1.33792022367234 | 0.223917992009751 | 0.49620415060819 | -5.53190991360221 |
| **Ap2m1** | 0.178023222499998 | 19.59907288375 | 1.33725677423785 | 0.224123214559001 | 0.49620415060819 | -5.53268935478528 |
| **Set** | 0.355178470000002 | 22.6613794775 | 1.33723187840002 | 0.224130918723421 | 0.49620415060819 | -5.53271859890662 |
| **Ndrg2** | 0.266697367500008 | 22.11600547375 | 1.33717623197756 | 0.224148139680589 | 0.49620415060819 | -5.53278396335858 |
| **Atm** | 0.518018570000002 | 22.34399987 | 1.33588783813918 | 0.224547185823382 | 0.496607254444175 | -5.53429692710084 |
| **Cetn2** | 0.267340085000004 | 23.4804105475 | 1.33460011180595 | 0.224946649287949 | 0.496947994038011 | -5.53580827512942 |
| **Stx1a** | -0.396394372499998 | 26.89829762125 | -1.33399215092805 | 0.225135460820191 | 0.496947994038011 | -5.53652152282088 |
| **Slc25a1** | 0.123280349999995 | 27.3838099525 | 1.33143880814253 | 0.225929961899968 | 0.497831882024147 | -5.53951502372457 |
| **Calu** | 0.141401297500003 | 22.47015489375 | 1.33062836724299 | 0.226182654008997 | 0.497831882024147 | -5.54046448567395 |
| **Rpl12** | 0.282575932500002 | 24.76061588625 | 1.32990145093581 | 0.226409514580834 | 0.497831882024147 | -5.54131581340178 |

| | | | | | | |
|---|---|---|---|---|---|---|
| **Hspa2** | -0.126648712499996 | 25.53426173375 | -1.32900730963859 | 0.226688837720085 | 0.497831882024147 | -5.54236262045532 |
| **Arrb1** | 1.084566315 | 23.112527195 | 1.32854077646542 | 0.226834699281482 | 0.497831882024147 | -5.54290864904544 |
| **Cst3** | 0.115788817500004 | 31.49377500875 | 1.3283876982554 | 0.226882577115515 | 0.497831882024147 | -5.54308778718507 |
| **Ndel1** | -0.133232687500001 | 28.03697726625 | -1.32782605770573 | 0.227058315785587 | 0.497831882024147 | -5.54374493943819 |
| **Mtarc2** | -6.9535736025 | 18.18288778 | -1.2597456579952 | 0.25606053121927 | 0.530908112283432 | -5.54743731740947 |
| **Slc23a2** | 0.361774609999998 | 22.9373997625 | 1.32314126194331 | 0.228528857068728 | 0.500502594821475 | -5.54922018970992 |
| **Dpp6** | -0.674255432500011 | 20.37871507125 | -1.32255419097661 | 0.22871372397696 | 0.500502594821475 | -5.54990552887284 |
| **Ptprz1** | 0.445132505 | 20.9533602275 | 1.32016159635843 | 0.229468500507336 | 0.501309548350378 | -5.55269679513732 |
| **Gnaz** | -0.178826562499992 | 25.21271533625 | -1.31999703973039 | 0.229520492211095 | 0.501309548350378 | -5.55288866394218 |
| **Adgrl1** | 0.391175207500005 | 23.29040586125 | 1.31810883234433 | 0.230117810187442 | 0.502135050065829 | -5.55508927374072 |
| **Cryl1** | 0.416491940000004 | 21.8802278025 | 1.31708963584454 | 0.2304407890918 | 0.502280252014409 | -5.55627633652193 |
| **Frrs1l** | 0.470427717499998 | 22.85143155125 | 1.31651451447963 | 0.230623217504213 | 0.502280252014409 | -5.55694594780096 |
| **Abat** | 0.254817452499999 | 23.40565664375 | 1.31110171146291 | 0.232346353599242 | 0.505552094475918 | -5.56323971410806 |
| **Mt-Cyb** | 0.171709275000005 | 26.871221725 | 1.3097236487555 | 0.232786844501696 | 0.506029522378331 | -5.56483964996378 |
| **Slc1a4** | 6.48602676 | 18.891070095 | 1.30884547147434 | 0.233067929690976 | 0.506159858693212 | -5.56585870431911 |
| **Tspan9** | 0.489625712500011 | 23.69303205375 | 1.30435441217779 | 0.234510053345656 | 0.508302007052347 | -5.57106396250482 |
| **Anp32a** | 0.255807900000004 | 20.2049949625 | 1.30388725879903 | 0.23466050674461 | 0.508302007052347 | -5.57160480346476 |
| **Psmc1** | 0.933973797500002 | 20.63781055375 | 1.3037010584332 | 0.234720498669433 | 0.508302007052347 | -5.57182034290026 |
| **Ncln** | 6.33774275 | 18.62803709 | 1.30178787926731 | 0.235337680700994 | 0.508712896176663 | -5.57403392867102 |
| **Negr1** | 0.363921602499996 | 22.08601771875 | 1.30173511780476 | 0.235354721297985 | 0.508712896176663 | -5.5740949476375 |
| **Rasal1** | 0.342974679999998 | 19.1895836825 | 1.29564240814332 | 0.237329748449026 | 0.511898782474971 | -5.58113139898316 |

| | | | | | | |
|---|---|---|---|---|---|---|
| **Stx1b** | 0.286433022499999 | 22.40987543375 | 1.29550361042933 | 0.237374908906801 | 0.511898782474971 | -5.58129146894921 |
| **Emc2** | 0.251826952500004 | 20.20697357875 | 1.29512060356635 | 0.237499566181048 | 0.511898782474971 | -5.58173312315239 |
| **Sepsecs** | -0.39006048249999 | 23.30872204875 | -1.29168740736006 | 0.238619512048768 | 0.513828845794571 | -5.58568856817013 |
| **Gna11** | -0.266782077500004 | 20.13273553125 | -1.29080020084977 | 0.23890967333843 | 0.513915588460923 | -5.58670972180359 |
| **Tomm20** | -0.229622199999991 | 22.3937223925 | -1.29019180790314 | 0.239108825561766 | 0.513915588460923 | -5.58740972685118 |
| **Fyn** | 0.1466944525 | 27.78139157375 | 1.28787912523856 | 0.239867178011838 | 0.515061885993524 | -5.59006886398488 |
| **Pdia3** | 6.559950575 | 19.450667725 | 1.28659873431522 | 0.240287928187507 | 0.51548178783618 | -5.59153984428911 |
| **Glud1** | 0.413067925 | 22.027148495 | 1.28527431296934 | 0.2407238205258 | 0.515933356913442 | -5.59306049149011 |
| **Gls** | -0.253986757500002 | 23.90501181375 | -1.28376081681078 | 0.241222780082545 | 0.516036657843136 | -5.5947970825458 |
| **Atl2** | 0.356096525000002 | 21.0632132425 | 1.28362772052693 | 0.241266701242135 | 0.516036657843136 | -5.59494973926127 |
| **Pdxk** | -0.158891819999997 | 20.7927484225 | -1.28261934561212 | 0.241599685472616 | 0.516036657843136 | -5.59610600221001 |
| **Ppm1a** | 0.848736972499999 | 21.45183184375 | 1.28239510965037 | 0.241673786460394 | 0.516036657843136 | -5.59636305073518 |
| **Tom1** | 0.328190802499996 | 25.44478457875 | 1.27996366914317 | 0.242478544607759 | 0.517272496372005 | -5.59914855788906 |
| **Alyref** | 0.196874327500005 | 22.46521200625 | 1.27902741193354 | 0.242789045048494 | 0.51745262953073 | -5.60022030836499 |
| **Ywhag** | -1.05586376 | 20.713174 | -1.27776648800701 | 0.243207761640289 | 0.517862852460113 | -5.60166296720338 |
| **Srsf3** | 0.51700194 | 20.5816197575 | 1.27698729397305 | 0.243466821733389 | 0.517932671884505 | -5.60255403812388 |
| **Cfl1** | 0.325062349999996 | 17.89343835 | 1.2761373344976 | 0.243749681438572 | 0.518052944115962 | -5.60352566256888 |
| **Stk39** | 0.122013389999996 | 25.901817945 | 1.27452291422142 | 0.244287729943774 | 0.518714855140351 | -5.60537010290159 |
| **Timm44** | 0.219365357499996 | 18.73166501375 | 1.27303648213034 | 0.244784030202726 | 0.519286974174271 | -5.60706707634078 |
| **Acad9** | 0.114206720000006 | 27.17799375 | 1.27151405512014 | 0.245293251287866 | 0.519885418701783 | -5.6088039054103 |
| **Slc12a7** | -0.191960517500004 | 21.84666570625 | -1.27050601883145 | 0.245630922142712 | 0.520119501188408 | -5.60995321229537 |

| | | | | | | |
|---|---|---|---|---|---|---|
| **Kcnj16** | 0.152053942500004 | 21.30977523625 | 1.26897393481559 | 0.24614490616164 | 0.520402672794247 | -5.61169895447143 |
| **Pcp4l1** | 0.235055624999998 | 22.95803144 | 1.26875225822939 | 0.246219351086138 | 0.520402672794247 | -5.61195143968619 |
| **Slc3a2** | -5.5565254575 | 16.77385755875 | -1.26654208090607 | 0.246962650404343 | 0.521492164922087 | -5.61446732711031 |
| **Cacna1a** | 0.625543310000005 | 21.75474812 | 1.26522179316811 | 0.247407595523217 | 0.521950217652206 | -5.61596896765389 |
| **Fh** | 0.248309439999996 | 20.22655924 | 1.26364429849645 | 0.247940126602382 | 0.522351174285042 | -5.61776189833709 |
| **Pacsin1** | -0.505314414999997 | 23.5690434575 | -1.26330719924222 | 0.248054052620289 | 0.522351174285042 | -5.6181448585255 |
| **Arpc1a** | 0.112016704999995 | 28.86413715 | 1.2623135157601 | 0.248390140105601 | 0.522578153218492 | -5.61927336706417 |
| **Nln** | 0.374288735000007 | 23.7306486325 | 1.26135162117366 | 0.24871584921252 | 0.522782900686372 | -5.62036525899181 |
| **Snap91** | 0.166288569999999 | 27.874739295 | 1.26013209017981 | 0.249129325560194 | 0.523171583676408 | -5.62174887759699 |
| **Psma1** | 0.169517959999993 | 25.10360558 | 1.25798654070791 | 0.249858199772 | 0.524104899308012 | -5.6241811297124 |
| **Rassf4** | 0.245462282499997 | 22.17332170875 | 1.25747662472703 | 0.250031695082721 | 0.524104899308012 | -5.62475881208349 |
| **Cnih2** | 0.219542247500005 | 23.55898722875 | 1.25115086465281 | 0.252192612973153 | 0.528150861020628 | -5.63191332814826 |
| **Fdps** | 0.195299802500003 | 24.32950983625 | 1.24834735576032 | 0.253155423288043 | 0.529682599548749 | -5.63507705187954 |
| **Atp1a2** | 0.297385947500004 | 20.49248953875 | 1.2465001738218 | 0.253791522763925 | 0.530528580462671 | -5.63715918636542 |
| **Uqcr11** | 7.3904947425 | 22.64402920375 | 1.24565645659832 | 0.254082522378724 | 0.53065227529644 | -5.63810958827787 |
| **Atp6v1b2** | 0.154825249999998 | 22.97737679 | 1.24469170950593 | 0.254415615637232 | 0.530720471803597 | -5.63919583743677 |
| **Crip2** | 0.114898597499998 | 26.41180894375 | 1.24421921001128 | 0.254578889489012 | 0.530720471803597 | -5.63972765493555 |
| **Rps7** | 0.130616204999999 | 25.7460223625 | 1.24271045310331 | 0.255100846436458 | 0.530908112283432 | -5.64142498758319 |
| **Mapk1** | -5.781604085 | 17.7297546225 | -1.24074681901503 | 0.255781540343575 | 0.530908112283432 | -5.64363214404613 |
| **Sbf1** | -0.174043687500003 | 28.51418299625 | -1.24074589948378 | 0.255781859462773 | 0.530908112283432 | -5.64363317710815 |
| **Cyp46a1** | 0.188855677500008 | 26.49292074125 | 1.24070230245651 | 0.255796990006525 | 0.530908112283432 | -5.64368215633822 |

| | | | | | | |
|---|---|---|---|---|---|---|
| **Mccc1** | -0.533759677500001 | 23.07724998875 | -1.24017350492949 | 0.255980572519146 | 0.530908112283432 | -5.64427615087684 |
| **Snx6** | 0.206177075000003 | 26.7388108025 | 1.23887742100976 | 0.256431010215217 | 0.531195097178853 | -5.6457313692439 |
| **Slc25a20** | 0.634441032500003 | 24.28128624625 | 1.23679389594855 | 0.257156533139592 | 0.532130611277309 | -5.6480687331164 |
| **Idh3g** | -0.325883222499993 | 21.78931279875 | -1.23559838496903 | 0.257573624015418 | 0.532130611277309 | -5.64940879158891 |
| **Slc1a3** | 6.0516802525 | 18.59294073125 | 1.23497886165968 | 0.257789990994125 | 0.532130611277309 | -5.65010290362659 |
| **Slc17a7** | 6.583632615 | 20.2964545625 | 1.23491439287819 | 0.257812515468124 | 0.532130611277309 | -5.65017512181652 |
| **Ak4** | 0.461077752500003 | 23.62896243125 | 1.23267058731096 | 0.258597516593989 | 0.533270013949226 | -5.65268717946546 |
| **Fbxl8** | 0.195851397499997 | 27.82661662125 | 1.22974155983372 | 0.25962531187344 | 0.534907595750049 | -5.6559620924149 |
| **Kras** | 0.166556370000009 | 26.5465005825 | 1.22736957765978 | 0.260460189819961 | 0.536145120951341 | -5.65861061200115 |
| **Trim56** | 0.947776477499996 | 21.98882215125 | 1.22477448177742 | 0.261376216426186 | 0.537547313027438 | -5.66150458939311 |
| **Agfg1** | 0.109993585000005 | 26.03467334 | 1.22173189978466 | 0.262453688401339 | 0.539278718806701 | -5.66489269561055 |
| **Dhx57** | -6.73564719 | 20.9368705925 | -1.21992234522358 | 0.263096296644444 | 0.539395740572026 | -5.66690523581608 |
| **Serpina3n** | -0.822733784999993 | 22.7333128525 | -1.2196492502796 | 0.263193394031581 | 0.539395740572026 | -5.66720880195356 |
| **Phyhipl** | 6.2864824775 | 19.58100403375 | 1.21953472877517 | 0.263234120574771 | 0.539395740572026 | -5.66733608872029 |
| **Snx1** | -0.124748625000002 | 20.1807665625 | -1.21866628961787 | 0.263543132252718 | 0.539395740572026 | -5.66830108433605 |
| **Mblac2** | -0.213317660000008 | 26.9458945025 | -1.21709131881988 | 0.264104331026136 | 0.539395740572026 | -5.67005006082491 |
| **Kctd12** | 0.6127149 | 20.8954677125 | 1.21628774906113 | 0.264391052302847 | 0.539395740572026 | -5.67094186063329 |
| **Anpep** | 0.150523075000002 | 25.622272295 | 1.21627150459541 | 0.264396851204899 | 0.539395740572026 | -5.67095988486838 |
| **Atp6v0a1** | 0.525724817500006 | 21.27313774875 | 1.21579887827056 | 0.264565615234811 | 0.539395740572026 | -5.6714842263252 |
| **Sco1** | -0.292547657500009 | 23.71485073875 | -1.21501847255143 | 0.264844480306396 | 0.539395740572026 | -5.6723497428191 |
| **Abcb7** | 0.167729737499997 | 19.99593178375 | 1.21495517671023 | 0.264867108957168 | 0.539395740572026 | -5.67241992628508 |

| | | | | | | |
|---|---|---|---|---|---|---|
| **Surf4** | 6.1762275325 | 19.32451567875 | 1.21267500051478 | 0.265683380191139 | 0.540577117562238 | -5.67494668057073 |
| **Hnrnpu** | -0.176521504999997 | 25.0460642725 | -1.21099652722551 | 0.266285612009225 | 0.541258854180146 | -5.67680474304941 |
| **Ap3d1** | -5.8339000925 | 18.30586916625 | -1.2104238196981 | 0.266491362455668 | 0.541258854180146 | -5.67743835442163 |
| **Ctnna2** | 0.29390047 | 23.020529815 | 1.20890621269594 | 0.267037228002099 | 0.541886715334047 | -5.67911642956955 |
| **Lnpk** | 5.9029487775 | 18.48123649625 | 1.20533833408173 | 0.268324281685454 | 0.544016192008862 | -5.68305629527966 |
| **Brk1** | 0.123088349999996 | 23.112502455 | 1.20408404655158 | 0.268777990090417 | 0.544453822404393 | -5.68443959344266 |
| **Galnt2** | -5.4435892175 | 17.13666163125 | -1.20326700579483 | 0.269073883651848 | 0.544571281767532 | -5.68534017764981 |
| **Uba6** | -5.3654183775 | 16.93404457875 | -1.20197622615107 | 0.269541903991623 | 0.54493648442184 | -5.68676214622904 |
| **Gjb6** | -5.7350521825 | 17.95471061625 | -1.20095297601804 | 0.269913410003288 | 0.54493648442184 | -5.68788870223172 |
| **Hnrnpa3** | 0.52911379 | 20.702465585 | 1.20080124888532 | 0.269968533566783 | 0.54493648442184 | -5.68805569532659 |
| **Slc7a11** | -0.180411277499999 | 25.91063413625 | -1.20005036839498 | 0.270241473815221 | 0.545006813711931 | -5.68888192678798 |
| **Slc32a1** | 0.0953781749999933 | 26.8042561775 | 1.19834707215762 | 0.270861474654984 | 0.545776334053924 | -5.69075492326741 |
| **Prkcg** | -5.508834325 | 17.44732858 | -1.19716094543036 | 0.271293934011257 | 0.5461669436691 | -5.69205821807244 |
| **Slc25a13** | -0.134765782499997 | 25.44823301125 | -1.19561299897895 | 0.271859187797754 | 0.546296651378103 | -5.69375782925541 |
| **Arfgap1** | -0.196810307499995 | 24.02674488375 | -1.19548520358662 | 0.27190589836671 | 0.546296651378103 | -5.69389808274239 |
| **Vwa8** | 6.031816535 | 19.01255872 | 1.19502449606375 | 0.27207434799958 | 0.546296651378103 | -5.69440362231733 |
| **Arl3** | -5.79291009416667 | 16.5808300428571 | -1.12305242524753 | 0.305725042923282 | 0.566375225794997 | -5.69499803602008 |
| **Cadm2** | 6.04020156000001 | 19.073342715 | 1.19354644899112 | 0.272615363763169 | 0.54690321442059 | -5.6960246552055 |
| **Adh5** | -0.622657280000006 | 22.2483112825 | -1.19204600446296 | 0.273165504379397 | 0.547527004837514 | -5.69766893299767 |
| **Rtn3** | 0.229978265 | 25.128212 | 1.18984207462901 | 0.273975273588196 | 0.54866964238266 | -5.70008171795008 |
| **Ptpra** | 5.6601783025 | 18.05297530125 | 1.18748009095934 | 0.274845356000092 | 0.549930961437246 | -5.70266433822739 |

| | | | | | | |
|---|---|---|---|---|---|---|
| **M6pr** | 0.332210152499997 | 21.84061732875 | 1.18653926547367 | 0.275192573656181 | 0.549945234146312 | -5.70369212489539 |
| **Stum** | 0.473643152500003 | 20.57540220375 | 1.18583817763655 | 0.275451554235018 | 0.549945234146312 | -5.70445767244883 |
| **Aak1** | 0.124795027500003 | 24.88108888875 | 1.18550890865347 | 0.275573256254181 | 0.549945234146312 | -5.70481711425389 |
| **Gpm6b** | -6.3879631675 | 20.43438806125 | -1.18137000770853 | 0.277106901933022 | 0.552047302233938 | -5.70932977280966 |
| **Acadsb** | 5.541519205 | 17.7358323325 | 1.18136453606749 | 0.277108934148883 | 0.552047302233938 | -5.70933573178772 |
| **Pcca** | -5.55687457 | 17.73810706 | -1.1804214291473 | 0.277459399223777 | 0.552264838976717 | -5.71036256980558 |
| **Syn1** | 0.205386624999996 | 25.4938142275 | 1.17952507598324 | 0.277792834726038 | 0.552448130919115 | -5.71133800957966 |
| **Dbt** | -0.840471364999999 | 20.51318141 | -1.17694644252996 | 0.278753937229236 | 0.553477386064524 | -5.71414147313008 |
| **Nt5dc3** | -0.109418907500004 | 28.28423562625 | -1.17659745799791 | 0.278884223734126 | 0.553477386064524 | -5.71452057874815 |
| **Ap3m2** | 1.3166522125 | 21.37534886625 | 1.17619166856414 | 0.279035781353631 | 0.553477386064524 | -5.71496130007326 |
| **Adgrl3** | -0.243310862499996 | 20.95525391625 | -1.17377511142976 | 0.279939766008721 | 0.554768136671082 | -5.71758383119846 |
| **Atad3** | 5.9108999325 | 19.05624196125 | 1.17216039477548 | 0.280545163251681 | 0.554768136671082 | -5.71933421551229 |
| **Slc25a10** | 5.5564219175 | 17.92512443875 | 1.17066389354918 | 0.281107215749051 | 0.554768136671082 | -5.72095504600402 |
| **Tardbp** | 6.08623111499999 | 19.5768422275 | 1.17003161254993 | 0.281344969073855 | 0.554768136671082 | -5.72163944989078 |
| **Eef2** | 0.181616420000005 | 19.682736375 | 1.16929376322572 | 0.281622631131072 | 0.554768136671082 | -5.72243781876769 |
| **Dhrs4** | -0.0960150175000081 | 21.01714858625 | -1.16895731499298 | 0.281749316840004 | 0.554768136671082 | -5.72280175354178 |
| **Cdc42** | 5.44891768 | 17.5733139225 | 1.16894703308192 | 0.281753189124739 | 0.554768136671082 | -5.72281287436576 |
| **Vcp** | -0.148873232499987 | 25.38884872375 | -1.16884438518716 | 0.281791849927481 | 0.554768136671082 | -5.72292389390735 |
| **Abcb8** | 0.143952877500002 | 19.82383276125 | 1.16864282461966 | 0.281867777609641 | 0.554768136671082 | -5.72314187455291 |
| **Ptges2** | 0.105117089999997 | 29.3962300725 | 1.16678153915136 | 0.282569729777836 | 0.554968432389802 | -5.72515362466578 |

| | | | | | | |
|---|---|---|---|---|---|---|
| **Ak2** | 5.122348375 | 16.583099775 | 1.16661518972278 | 0.282632536613647 | 0.554968432389802 | -5.72533331925567 |
| **Dhrs1** | 5.558792195 | 18.0068454125 | 1.16559228983856 | 0.283018998485238 | 0.554968432389802 | -5.72643791029657 |
| **Pcsk1n** | 0.239665415000001 | 23.37721975 | 1.16520798764042 | 0.283164305617614 | 0.554968432389802 | -5.72685273935173 |
| **Arcn1** | 0.311328165000006 | 24.784195635 | 1.16516174323057 | 0.283181795120703 | 0.554968432389802 | -5.72690265110754 |
| **Gabrg2** | 5.0280271125 | 16.29986759125 | 1.16388606317868 | 0.283664609014531 | 0.55533728185067 | -5.72827898387991 |
| **Ptn** | 0.339252582500002 | 22.97128778875 | 1.16304191384446 | 0.283984476252382 | 0.55533728185067 | -5.72918919260294 |
| **Map1a** | 0.112900967500003 | 27.01870621875 | 1.16229084182204 | 0.284269327101474 | 0.55533728185067 | -5.72999867490735 |
| **Fam234b** | 5.50257317 | 17.87301153 | 1.16210340356285 | 0.284340451869369 | 0.55533728185067 | -5.7302006364268 |
| **Slc25a11** | 0.206600014999999 | 25.3586692825 | 1.15994287649201 | 0.285161349972939 | 0.555896607028107 | -5.73252701815123 |
| **Mdh2** | 5.0770682025 | 16.51764864875 | 1.15936629125446 | 0.285380758423415 | 0.555896607028107 | -5.73314738296502 |
| **Lancl2** | -5.70328392 | 18.46576912 | -1.15905099791336 | 0.285500796681292 | 0.555896607028107 | -5.73348653017539 |
| **Fkbp3** | 0.261792210000003 | 24.124071625 | 1.15879508957016 | 0.28559825682178 | 0.555896607028107 | -5.73376175479691 |
| **Rab7a** | 0.112675069999995 | 24.1136410125 | 1.1527862334625 | 0.287894635812361 | 0.559442863463716 | -5.74021261545908 |
| **Cacna1e** | -0.405499320000008 | 25.2871572275 | -1.15274878167515 | 0.287908996575036 | 0.559442863463716 | -5.7402527525191 |
| **Qdpr** | 0.156608477500001 | 21.17077041375 | 1.15209075674203 | 0.288161411100973 | 0.559458413918683 | -5.74095781641738 |
| **Fech** | -0.372498592500001 | 24.74351193125 | -1.15102115566347 | 0.28857209588561 | 0.559739579708613 | -5.74210330808323 |
| **Prdx5** | -0.124811869999991 | 22.99503789 | -1.15044036312554 | 0.288795300845728 | 0.559739579708613 | -5.74272501327704 |
| **Selenof** | -0.250974035000002 | 23.9595583225 | -1.14925552656564 | 0.289251090804146 | 0.559807633872428 | -5.74399266668287 |
| **Gdpd1** | 5.9146379225 | 19.38439426125 | 1.14886754513456 | 0.289400471433909 | 0.559807633872428 | -5.74440757827601 |
| **D2hgdh** | 0.0864134225000086 | 23.59919800125 | 1.1484427331396 | 0.289564105943624 | 0.559807633872428 | -5.74486177004182 |
| **Lpcat4** | 5.36001721 | 17.4914783575 | 1.14773551733303 | 0.289836690533069 | 0.559861759181599 | -5.74561764867353 |

| | | | | | | |
|---|---|---|---|---|---|---|
| **Syn3** | 5.5334476875 | 18.16155005625 | 1.1465966210138 6 | 0.29027610629575 | 0.560237780194748 | -5.74683425993 |
| **Diras1** | 4.98336347499999 | 16.37331957 | 1.1457145453440 7 | 0.29061681323000 6 | 0.56042281843596 | -5.7477759730381 |
| **Pgrmc1** | -0.46448474499998 6 | 24.0181713875 | -1.1433790743224 9 | 0.29152050253236 4 | 0.561692281394428 | -5.7502670122777 2 |
| **Kars1** | 0.28809025750000 2 | 21.18056899875 | 1.1418258073605 | 0.29212281208762 8 | 0.561941073211583 | -5.7519218650225 8 |
| **Opalin** | 6.018326215 | 19.787599265 | 1.1417799260588 6 | 0.29214061910082 3 | 0.561941073211583 | -5.7519707241692 8 |
| **Skp1** | -5.8464505275 | 19.27320473625 | -1.1410243039133 5 | 0.29243401314199 4 | 0.562033128532346 | -5.7527751998344 7 |
| **Abhd12** | 0.34602411750000 1 | 20.16063267875 | 1.1394095612928 | 0.29306180444124 2 | 0.562767173125842 | -5.7544931468062 1 |
| **Manf** | 0.119985555 | 24.70347554 | 1.1379673910924 3 | 0.29362344311691 1 | 0.562928076300294 | -5.7560261143117 7 |
| **Txndc15** | 0.37606507250000 5 | 21.93328392625 | 1.1379314534379 5 | 0.29363745002295 8 | 0.562928076300294 | -5.7560642979452 5 |
| **Sugt1** | 5.640070715 | 18.6690560575 | 1.1359205770165 8 | 0.29442208038554 3 | 0.563959951466533 | -5.7581995584561 1 |
| **Dag1** | 0.10562013000000 9 | 29.4862626125 | 1.1327265994093 8 | 0.29567190541301 7 | 0.565161096255617 | -5.7615858871870 3 |
| **Syngr1** | 5.5439504425 | 18.39259984375 | 1.1323247232430 1 | 0.29582947172218 2 | 0.565161096255617 | -5.7620115114283 5 |
| **Tspan7** | -0.94064326999999 2 | 20.4416776275 | -1.1321147167595 4 | 0.29591183791964 3 | 0.565161096255617 | -5.7622338873935 7 |
| **Map2k4** | 6.05671889 | 20.090984705 | 1.1317962868366 2 | 0.29603676470532 3 | 0.565161096255617 | -5.7625710200242 8 |
| **Bcl2l13** | 5.93756141 | 19.69940807 | 1.1310334013948 | 0.29633623760146 2 | 0.565261373224789 | -5.7633784531143 6 |
| **Atp6v1h** | 5.608315755 | 18.557840845 | 1.1294448286186 3 | 0.29696063750135 8 | 0.565589001774887 | -5.7650586110351 5 |
| **Tppp** | 0.18895662750000 5 | 25.01158058375 | 1.1293392465028 8 | 0.29700217568082 8 | 0.565589001774887 | -5.7651702237949 9 |
| **Rpl7a** | 0.31760705250000 1 | 23.15408346125 | 1.1281009041957 3 | 0.29748972225123 5 | 0.565725098900735 | -5.7664787722871 3 |
| **Akr7a2** | 0.15374014999999 4 | 25.80794237 | 1.1279023859199 9 | 0.29756794192943 9 | 0.565725098900735 | -5.7666884551784 1 |
| **Clybl** | 0.20748350499999 9 | 21.028728 | 1.1271383058947 8 | 0.29786916080017 8 | 0.56582780835818 | -5.7674952743244 2 |
| **Ttc7b** | 0.33820576749999 6 | 21.50409712125 | 1.1256479697542 5 | 0.29845740928016 4 | 0.566162878766941 | -5.7690679098768 6 |

| | | | | | | |
|---|---|---|---|---|---|---|
| **Erp29** | 0.309292882499996 | 23.73506080125 | 1.12464676395629 | 0.298853130028672 | 0.566162878766941 | -5.77012361410818 |
| **Nefm** | 0.138818312499993 | 25.33141579375 | 1.12449164129338 | 0.298914479929396 | 0.566162878766941 | -5.77028712366107 |
| **Gabra1** | -5.3773262375 | 17.94996155625 | -1.12418719841882 | 0.29903491499748 | 0.566162878766941 | -5.77060798221811 |
| **Efhd2** | 6.2081171075 | 20.74848975125 | 1.12348658878048 | 0.299312221829203 | 0.566219566749625 | -5.77134614556123 |
| **Vgf** | -0.179207704999989 | 27.4176196625 | -1.12272482566845 | 0.299613973414257 | 0.566322365933306 | -5.77214838658217 |
| **Phyhip** | 5.648150985 | 18.916378095 | 1.1209482500527 | 0.300318686105086 | 0.566375225794997 | -5.77401793090524 |
| **Kcnip4** | 0.12972445250001 | 22.75486505875 | 1.12008035679231 | 0.300663446983138 | 0.566375225794997 | -5.77493051089576 |
| **Rps26** | 4.96459844 | 16.6373705925 | 1.11988066920936 | 0.30074281658401 | 0.566375225794997 | -5.77514041223533 |
| **Map6** | 0.1129393075 | 30.55508824375 | 1.11974736159243 | 0.300795811782677 | 0.566375225794997 | -5.77528052421559 |
| **Scrn3** | 5.9839992825 | 19.76727846875 | 1.11882781561224 | 0.30116157743715 | 0.566375225794997 | -5.77624669754725 |
| **Lrp6** | 0.476534497499998 | 22.58295294375 | 1.11827249930602 | 0.301382640931165 | 0.566375225794997 | -5.77682991100081 |
| **Sirpa** | 0.286142914999999 | 26.7475255975 | 1.11637130040316 | 0.302140488785381 | 0.566375225794997 | -5.77882512813551 |
| **Stam** | -5.977313625 | 20.08605625 | -1.11636051436185 | 0.302144792722939 | 0.566375225794997 | -5.77883644097818 |
| **Map1lc3a** | 0.401944399999998 | 20.742677015 | 1.11381019214051 | 0.303163854737866 | 0.566375225794997 | -5.78150923302898 |
| **Stoml2** | -0.4389964175 | 22.77416189875 | -1.1136446980481 | 0.303230080266449 | 0.566375225794997 | -5.78168253038237 |
| **Rps11** | 0.540053260000004 | 23.672244355 | 1.1133397978492 | 0.303352122739407 | 0.566375225794997 | -5.78200176102861 |
| **Ddost** | -0.370141485000005 | 23.640354725 | -1.11248889248663 | 0.30369292748847 | 0.566375225794997 | -5.78289234381204 |
| **Rpl14** | -5.4052126 | 18.2150743675 | -1.11227779107559 | 0.303777526346823 | 0.566375225794997 | -5.78311321685169 |
| **Hnrnpl** | 6.06553793 | 20.42530151 | 1.11222906843737 | 0.3037970546751 | 0.566375225794997 | -5.78316419073716 |
| **Srgap2** | 0.459641815000001 | 24.8480146375 | 1.11162545048152 | 0.304039073594188 | 0.566375225794997 | -5.78379557261451 |

| | | | | | | |
|---|---|---|---|---|---|---|
| **Psmb4** | 0.429238954999999 | 23.7034375575 | 1.11127364184729 | 0.304180203005823 | 0.566375225794997 | -5.78416345499559 |
| **Pde2a** | -5.6567053275 | 19.06919496875 | -1.11022463847872 | 0.304601332871657 | 0.566375225794997 | -5.7852599137468 |
| **Aifm1** | 5.11904097 | 17.27819229 | 1.1099165711163 | 0.304725099218332 | 0.566375225794997 | -5.78558178314511 |
| **Prkar2a** | -0.2033429725 | 20.64143819375 | -1.1094815978251 | 0.304899920131351 | 0.566375225794997 | -5.78603614008446 |
| **Pdzd8** | 0.1723214025 | 24.02360422625 | 1.10843695403134 | 0.305320109653797 | 0.566375225794997 | -5.78712683838573 |
| **Wbp2** | -0.219568814999999 | 20.655561715 | -1.10752582460597 | 0.305686981522067 | 0.566375225794997 | -5.78807756181386 |
| **Stx6** | 0.25409851500001 | 23.4335998225 | 1.10704806854759 | 0.305879496778554 | 0.566375225794997 | -5.78857586517848 |
| **Selenom** | 5.55664434 | 18.8214104125 | 1.10582194394635 | 0.306374025604163 | 0.566375225794997 | -5.78985404839139 |
| **Dguok** | 6.1339657 | 20.8125166625 | 1.10380021473921 | 0.307190867784078 | 0.566375225794997 | -5.79195949333993 |
| **Lman2** | 5.076700245 | 17.2127565025 | 1.10371149759968 | 0.307226752944315 | 0.566375225794997 | -5.79205182348881 |
| **Ahcyl1** | -5.59542922249999 | 18.98582707125 | -1.10368401380549 | 0.307237870546738 | 0.566375225794997 | -5.79208042552751 |
| **Cpne6** | 0.2615259775 | 23.63157063125 | 1.10318263189453 | 0.307440744553959 | 0.566375225794997 | -5.79260212148627 |
| **Cmtm4** | 0.2228645675 | 25.08290414375 | 1.10281098724896 | 0.307591193484945 | 0.566375225794997 | -5.79298871860832 |
| **Nudc** | 0.659958064999998 | 22.8416916125 | 1.10118562018177 | 0.308249878746792 | 0.566375225794997 | -5.79467842699045 |
| **Eif2s2** | 4.79283136 | 16.242585355 | 1.10110168091245 | 0.308283926592253 | 0.566375225794997 | -5.7947656424998 |
| **Cyb5r1** | -0.216067410000004 | 19.88751276 | -1.10107384275526 | 0.308295219114409 | 0.566375225794997 | -5.79479456619997 |
| **Mpst** | 5.669205935 | 19.205031875 | 1.10066718531551 | 0.308460217737114 | 0.566375225794997 | -5.79521702378427 |
| **Gucy1b1** | -5.7393219225 | 19.49469686875 | -1.1006209109747 | 0.308478997808533 | 0.566375225794997 | -5.7952650892473 |
| **Trim9** | -0.231420122500005 | 24.34103558875 | -1.10035191602011 | 0.308588185703138 | 0.566375225794997 | -5.79554446853183 |
| **Ryr2** | 0.149285352499998 | 20.4469446342857 | 1.0236323248596 | 0.346778100543972 | 0.579611774337155 | -5.79556570104784 |
| **Icam5** | 0.363009292499996 | 18.81715608125 | 1.09989416248531 | 0.308774065051269 | 0.566375225794997 | -5.79601978517127 |

| | | | | | | |
|---|---|---|---|---|---|---|
| **Mthfd1l** | 0.579536155 | 21.1131293625 | 1.09882984069184 | 0.309206605150769 | 0.566375225794997 | -5.79712441621072 |
| **Pafah1b1** | 0.2174702125 | 20.28172849625 | 1.09862400667274 | 0.30929031293119 | 0.566375225794997 | -5.79733796076897 |
| **Csrp1** | -0.0968388724999976 | 29.38056231125 | -1.09801078762785 | 0.309539803793181 | 0.566375225794997 | -5.79797398750032 |
| **Aldh3a2** | 5.2465631525 | 17.88814370875 | 1.09759466673122 | 0.309709197652949 | 0.566375225794997 | -5.79840544585657 |
| **Erc2** | 5.2456300775 | 17.77780328875 | 1.09740431340319 | 0.309786711531383 | 0.566375225794997 | -5.79860277765383 |
| **Ap2s1** | -0.164100274999988 | 19.5612059925 | -1.0942531329843 | 0.311072199280145 | 0.567497903779104 | -5.80186604776556 |
| **Hadha** | 0.876909157500002 | 20.06888923625 | 1.09332875488871 | 0.311450109897691 | 0.567497903779104 | -5.80282207612091 |
| **Tmem126a** | -0.262462845000005 | 21.3290124275 | -1.09277326894714 | 0.311677386795346 | 0.567497903779104 | -5.80339631272622 |
| **Dtd1** | 5.51332168 | 18.8602238875 | 1.09267039565793 | 0.311719492145724 | 0.567497903779104 | -5.80350263638051 |
| **Mtpn** | 5.5472963575 | 18.98991353125 | 1.09251741786151 | 0.31178211347666 | 0.567497903779104 | -5.80366073223141 |
| **Ephx1** | -0.206886524999998 | 23.1077422725 | -1.09225802434678 | 0.311888319333383 | 0.567497903779104 | -5.80392876904127 |
| **Slc1a6** | 5.52117018 | 18.93842612 | 1.08963494904646 | 0.31296395965338 | 0.568955976234268 | -5.80663677015171 |
| **Pdia4** | 0.130179314999996 | 19.3092936225 | 1.08909250003102 | 0.313186775908771 | 0.568955976234268 | -5.80719621830377 |
| **Ppme1** | -5.1851295675 | 17.80789289625 | -1.08822094122423 | 0.313545046565748 | 0.569155124178427 | -5.80809468490868 |
| **Adcy1** | 0.159296500000004 | 27.3041261725 | 1.08350045894567 | 0.3154912633952 | 0.571310777072368 | -5.81295221347168 |
| **Brinp1** | 5.32593718 | 18.356978415 | 1.08347793630092 | 0.315500572683627 | 0.571310777072368 | -5.81297535476855 |
| **Mpc2** | 4.845856365 | 16.7116457275 | 1.08266769986437 | 0.315835615649105 | 0.571310777072368 | -5.81380762385517 |
| **Arl8a** | 5.69212843 | 19.661155785 | 1.08099722619573 | 0.316527285624802 | 0.571310777072368 | -5.81552215223566 |
| **Tpd52l2** | 5.3686701275 | 18.45892063625 | 1.08086075767473 | 0.316583845317493 | 0.571310777072368 | -5.81566213817737 |
| **Rps15a** | 5.6670652075 | 19.57696057375 | 1.08016382435609 | 0.316872818303042 | 0.571310777072368 | -5.81637684229469 |

| | | | | | | |
|---|---|---|---|---|---|---|
| **Slc1a2** | -4.83157566 | 16.7016686425 | -1.07946210065174 | 0.317163992818198 | 0.571310777072368 | -5.81709613371298 |
| **Hdgf** | 5.9312264625 | 20.49713059125 | 1.07933827896752 | 0.317215394038704 | 0.571310777072368 | -5.81722302139416 |
| **C2cd2l** | -5.55334475 | 19.1903080425 | -1.07866660847575 | 0.31749433710039 | 0.571310777072368 | -5.81791114615321 |
| **Mfn2** | -0.219876215000003 | 24.73061321 | -1.07816950533575 | 0.31770091028664 | 0.571310777072368 | -5.81842023429655 |
| **Hsd17b12** | 5.7510768575 | 19.91092429875 | 1.07810535586901 | 0.317727575755289 | 0.571310777072368 | -5.81848591843508 |
| **Hyou1** | 0.498399900000006 | 22.207625225 | 1.07728926623339 | 0.318066963287372 | 0.571310777072368 | -5.81932129311613 |
| **Smap2** | -5.70054581 | 19.7205092125 | -1.07699776620906 | 0.318188260412832 | 0.571310777072368 | -5.8196195743221 |
| **Fundc1** | 0.214169142499998 | 25.18785187375 | 1.076468237691 | 0.318408699912576 | 0.571310777072368 | -5.82016127677386 |
| **Slc25a23** | 0.217851247500011 | 24.35120411125 | 1.07630559589213 | 0.318476431430468 | 0.571310777072368 | -5.82032762029689 |
| **Vars1** | 5.3949484425 | 18.72289784625 | 1.07491013728042 | 0.319058042211456 | 0.571388445921262 | -5.82175411605754 |
| **Matr3** | 0.164241987500002 | 26.58153973875 | 1.07465835426898 | 0.319163073561389 | 0.571388445921262 | -5.82201136102842 |
| **Atp6v0c** | 5.22088773000001 | 18.127448515 | 1.07440545488922 | 0.319268598660242 | 0.571388445921262 | -5.82226970406592 |
| **Chmp4b** | 5.9024750275 | 20.50945232125 | 1.07370651101418 | 0.319560387082078 | 0.571463848461622 | -5.82298347098924 |
| **Khsrp** | 5.0889464375 | 17.68008634875 | 1.0728925871888 | 0.319900447010634 | 0.571625388920641 | -5.82381424573441 |
| **Rpsa** | -0.250299854999998 | 20.63876174 | -1.07197958234047 | 0.32028225012359 | 0.57186120946404 | -5.82474562664948 |
| **Prnp** | -5.5667475375 | 19.37132295375 | -1.07011055047868 | 0.321064991791585 | 0.572720956825528 | -5.82665053936161 |
| **S1pr1** | -0.383571787499996 | 23.91414736125 | -1.06830540953499 | 0.321822436704891 | 0.572720956825528 | -5.82848811551878 |
| **Cct6a** | 5.599576395 | 19.41846976 | 1.06738360365588 | 0.322209784268852 | 0.572720956825528 | -5.82942564235264 |
| **Actbl2** | 5.603160415 | 19.5722720225 | 1.06699115182525 | 0.322374808231071 | 0.572720956825528 | -5.82982461426339 |
| **Ilvbl** | -5.0925483275 | 17.79177695875 | -1.06657746540144 | 0.322548834752642 | 0.572720956825528 | -5.83024506163722 |
| **Tmod2** | 6.05468334 | 21.1458920475 | 1.06642098394568 | 0.322614681877737 | 0.572720956825528 | -5.8304040705462 |

| | | | | | | |
|---|---|---|---|---|---|---|
| **Tmem11** | -5.24354751 | 18.2832701975 | -1.06572004737485 | 0.322909767307215 | 0.572720956825528 | -5.83111612665516 |
| **Ergic1** | 0.291850437499996 | 21.88979667875 | 1.06534372411233 | 0.323068284055897 | 0.572720956825528 | -5.83149828364818 |
| **Sdc4** | 0.183646862500005 | 26.45570328125 | 1.06498246369855 | 0.323220514738541 | 0.572720956825528 | -5.83186505472877 |
| **Rcn2** | 4.96616706 | 17.350000965 | 1.06478972710539 | 0.323301755119419 | 0.572720956825528 | -5.83206069542688 |
| **Tmed9** | -5.0522308775 | 17.68599700625 | -1.06428155811917 | 0.323516031968287 | 0.572720956825528 | -5.8325764015794 |
| **Rala** | -0.350063349999996 | 25.31421072 | -1.06185321694428 | 0.324541551458403 | 0.573253562611121 | -5.83503835925938 |
| **Tmem30a** | 5.369411915 | 18.82801992 | 1.06117362585216 | 0.324829017903658 | 0.573253562611121 | -5.83572664632327 |
| **Snta1** | 5.440311035 | 19.08739727 | 1.06107619491944 | 0.32487024782987 | 0.573253562611121 | -5.83582529842909 |
| **Syp** | 0.172742369999995 | 23.244191115 | 1.06073513218005 | 0.3250146086694 | 0.573253562611121 | -5.8361705854817 |
| **Acaa2** | 5.5082438175 | 19.34111917625 | 1.06031465984185 | 0.325192651787043 | 0.573253562611121 | -5.83659615757905 |
| **Atp2b1** | 5.375655105 | 18.883782255 | 1.06001515990726 | 0.325319518493598 | 0.573253562611121 | -5.83689921722036 |
| **Ywhaz** | 5.2080864525 | 18.30586722625 | 1.0591692986546 | 0.325678035343233 | 0.573435015211562 | -5.83775480461549 |
| **Atp2a2** | 5.57770076 | 19.6308643825 | 1.05755978588222 | 0.326361098484273 | 0.573435015211562 | -5.83938148794214 |
| **Prdx3** | -0.531378135000001 | 22.9402591175 | -1.05737326862339 | 0.326440328877679 | 0.573435015211562 | -5.83956988161974 |
| **Wasf3** | 0.137048092499999 | 28.35353724125 | 1.05625688166467 | 0.326914879120811 | 0.573435015211562 | -5.84069700701307 |
| **Acadvl** | 4.887927595 | 17.236372135 | 1.05503142036218 | 0.32743642957228 | 0.573435015211562 | -5.84193328295226 |
| **Sec61b** | 4.91731397 | 17.3406967475 | 1.05482119306127 | 0.327525968130583 | 0.573435015211562 | -5.8421452630892 |
| **Vcan** | -0.510231320000003 | 21.6973264925 | -1.05391875392173 | 0.327910551156206 | 0.573435015211562 | -5.84305488564561 |
| **Tm9sf3** | 0.148821992499997 | 25.77913201375 | 1.05310671341115 | 0.328256918362194 | 0.573435015211562 | -5.84387291697575 |
| **Krt1** | -0.342488762499993 | 22.88012253375 | -1.05307659292988 | 0.328269771552237 | 0.573435015211562 | -5.84390325104734 |
| **Prxl2b** | -1.51097267999999 | 21.1765586925 | -1.05228424668118 | 0.328608030655601 | 0.573435015211562 | -5.84470099433107 |

| | | | | | | |
|---|---|---|---|---|---|---|
| **Mtfr1l** | -0.175918545000002 | 27.5356120275 | -1.0519958340076 | 0.328731225476029 | 0.573435015211562 | -5.84499126546192 |
| **Ezr** | 5.0081669375 | 17.71943629125 | 1.05120080932096 | 0.329071009428976 | 0.573435015211562 | -5.84579111984289 |
| **Pck2** | -5.0742869225 | 17.88040280125 | -1.05097836789836 | 0.329166128390501 | 0.573435015211562 | -5.84601483549253 |
| **Emc10** | -0.145856279999997 | 26.733607945 | -1.05060412476715 | 0.329326209337328 | 0.573435015211562 | -5.84639114634122 |
| **Psmb3** | -0.250898934999995 | 21.60796595 | -1.05052801931013 | 0.329358770732193 | 0.573435015211562 | -5.84646766060527 |
| **Tecr** | 0.141178467499998 | 26.19192045125 | 1.05035974464743 | 0.329430775449194 | 0.573435015211562 | -5.84663682515248 |
| **Asap2** | 5.0966217425 | 18.05503471125 | 1.04887118002011 | 0.330068279287764 | 0.573677121642526 | -5.8481324206002 |
| **Nptn** | 0.0853464050000063 | 23.8240347725 | 1.04814347389386 | 0.330380290050541 | 0.573677121642526 | -5.84886301339106 |
| **Ahcyl2** | -5.0087446225 | 17.77137727125 | -1.04767381880243 | 0.330581783860508 | 0.573677121642526 | -5.84933433941819 |
| **Txn2** | -5.0366614825 | 17.86350920875 | -1.04749425050094 | 0.330658849045929 | 0.573677121642526 | -5.8495145067469 |
| **Arhgap1** | 5.3391642125 | 18.94734735375 | 1.04711192351278 | 0.330822979715218 | 0.573677121642526 | -5.84989803571282 |
| **Atp6v1d** | 0.275094930000005 | 22.91968396 | 1.04619639752767 | 0.331216273367495 | 0.573743284978061 | -5.85081603359631 |
| **Nrxn3** | -0.175955230000003 | 26.3081176625 | -1.04478019309736 | 0.331825383116923 | 0.573743284978061 | -5.85223493142058 |
| **Bdh1** | 0.185993672499997 | 25.58454440875 | 1.04468968430997 | 0.331864341262665 | 0.573743284978061 | -5.85232556562713 |
| **Dynlrb1** | 5.4927321675 | 19.55467440875 | 1.04234989649354 | 0.332872731803659 | 0.573743284978061 | -5.85466664550656 |
| **Pygm** | 0.273012184999999 | 21.15021619 | 1.04222409477007 | 0.332927018199341 | 0.573743284978061 | -5.85479241023668 |
| **Inpp4a** | 0.149267814999998 | 26.2624201125 | 1.0408872158951 | 0.333504347805555 | 0.573743284978061 | -5.85612822402632 |
| **Rdx** | 5.84784586 | 20.87016673 | 1.04074240944969 | 0.333566930041328 | 0.573743284978061 | -5.85627284133792 |
| **Tpm1** | 0.879417074999999 | 21.0742092125 | 1.04068744053478 | 0.333590688869581 | 0.573743284978061 | -5.85632773467979 |
| **Cept1** | 0.282785945000001 | 26.2141357275 | 1.04028197877974 | 0.333765980305356 | 0.573743284978061 | -5.8567325747653 |
| **Rpl18a** | 0.159792637500001 | 21.65399523875 | 1.03939558243133 | 0.334149446886729 | 0.573743284978061 | -5.85761721793411 |

| | | | | | | |
|---|---|---|---|---|---|---|
| **Slc17a6** | 5.5238152375 | 18.07982679 | 0.958747664858528 | 0.375913771340296 | 0.599016214397003 | -5.85775568300895 |
| **Psip1** | -0.0930172250000041 | 21.4726657875 | -1.03858161373165 | 0.334501888464054 | 0.573743284978061 | -5.85842910003275 |
| **Vamp4** | 0.356442627500002 | 22.53792707625 | 1.03838552647813 | 0.334586836694227 | 0.573743284978061 | -5.85862461636554 |
| **Rpl35** | 5.377098105 | 19.2107708225 | 1.03834286728828 | 0.33460531962661 | 0.573743284978061 | -5.85866714783951 |
| **Dmxl2** | 5.2723570825 | 18.82315778125 | 1.03740073426968 | 0.335013723929274 | 0.573743284978061 | -5.85960614028771 |
| **Dlg3** | -5.386954995 | 19.2791138325 | -1.03714834305871 | 0.335123199905104 | 0.573743284978061 | -5.85985758615389 |
| **Gars1** | 4.84064931 | 17.34184238 | 1.03574953216747 | 0.335730455942803 | 0.573743284978061 | -5.86125036028213 |
| **Arxes2** | 0.132820580000004 | 23.701116335 | 1.03561681203428 | 0.335788118112902 | 0.573743284978061 | -5.86138243742624 |
| **Lap3** | -5.25212526 | 18.824394245 | -1.0350255982612 | 0.336045074848517 | 0.573743284978061 | -5.86197063899956 |
| **Rftn2** | 5.5940734825 | 20.05603710875 | 1.03416859389379 | 0.336417827748046 | 0.573743284978061 | -5.86282284774809 |
| **Rplp0** | 5.29713428 | 19.0018999475 | 1.03382361252085 | 0.336567969405093 | 0.573743284978061 | -5.86316575519082 |
| **Enpp6** | -0.214931182499996 | 24.62555547625 | -1.03372150821998 | 0.336612417084125 | 0.573743284978061 | -5.86326722986716 |
| **Ensa** | 0.248134884999992 | 23.162817595 | 1.03363226050358 | 0.336651271889965 | 0.573743284978061 | -5.8633559213262 |
| **Gabbr1** | 0.212121754999995 | 21.5004533825 | 1.03341530383932 | 0.336745740767372 | 0.573743284978061 | -5.86357150275328 |
| **Atp5mg** | 5.2160465575 | 18.72333757125 | 1.03284784501826 | 0.336992927187341 | 0.573743284978061 | -5.86413521041223 |
| **Camk2d** | -0.0741266049999965 | 26.28830158 | -1.03253917813616 | 0.337127443554168 | 0.573743284978061 | -5.86444174316641 |
| **Abce1** | -0.436944149999999 | 22.543074135 | -1.02841029074311 | 0.338930893775492 | 0.576222177919824 | -5.86853572534548 |
| **Grm3** | -5.082829315 | 18.28935913 | -1.02753445802306 | 0.339314426366843 | 0.576222177919824 | -5.86940263032173 |
| **Eef1a1** | 0.556090794999999 | 20.275627145 | 1.02708303584411 | 0.339512240839244 | 0.576222177919824 | -5.86984924235774 |
| **Esyt1** | 0.3178239125 | 24.17003067375 | 1.02690344217888 | 0.339590964619416 | 0.576222177919824 | -5.87002688288332 |
| **Grhpr** | -0.142191149999999 | 19.758266945 | -1.02626339656897 | 0.339871641950767 | 0.576271250685412 | -5.8706597849869 |

| | | | | | | |
|---|---|---|---|---|---|---|
| **Rpl7** | 5.7157869275 | 20.61880645375 | 1.02540034417324 | 0.340250403830475 | 0.576335681242949 | -5.8715127526462 |
| **Ddx1** | 0.113034885000005 | 25.199074585 | 1.02487156323585 | 0.340482630870329 | 0.576335681242949 | -5.87203509805216 |
| **Slc24a3** | 0.144944389999996 | 21.9380744275 | 1.02420441275265 | 0.34077580472721 | 0.576335681242949 | -5.87269385046589 |
| **Ptrh2** | -4.7920975675 | 17.34738820125 | -1.02333664420672 | 0.34115743653955 | 0.576335681242949 | -5.87355023033132 |
| **Atp6v1f** | -4.9369310675 | 17.87937704875 | -1.0230020079139 | 0.341304694680191 | 0.576335681242949 | -5.87388033420134 |
| **Dlg1** | 0.255816492500003 | 22.93279015375 | 1.02273926457507 | 0.341420351142612 | 0.576335681242949 | -5.87413946397967 |
| **Vapb** | -0.195039959999999 | 21.11750816 | -1.02084935677545 | 0.342253176533905 | 0.577030437199295 | -5.87600195597043 |
| **Dnajb2** | -4.63043768499999 | 16.801419455 | -1.02066137847716 | 0.342336100356768 | 0.577030437199295 | -5.87618707060461 |
| **Hsd17b10** | 5.0576059875 | 18.37048440625 | 1.01939333075036 | 0.342895894221633 | 0.577548713666901 | -5.87743515371702 |
| **Numbl** | -5.2218886775 | 18.88618394875 | -1.01857166576991 | 0.343259012080948 | 0.577735204892125 | -5.87824327983548 |
| **Ethe1** | -0.476543719999988 | 22.0858276175 | -1.01656799860642 | 0.344145760078705 | 0.578561291154771 | -5.88021194376908 |
| **Pgs1** | 5.0500142075 | 18.39115521875 | 1.01632067213604 | 0.344255342312276 | 0.578561291154771 | -5.88045475384829 |
| **Agk** | -0.566908662499998 | 20.81295668375 | -1.01467923338582 | 0.344983305172135 | 0.578991539768629 | -5.8820651274417 |
| **Nr3c2** | 0.149674459999993 | 25.788447285 | 1.01460278433333 | 0.345017239075758 | 0.578991539768629 | -5.88214008340562 |
| **Cyp4a10** | 0.551455042499999 | 23.04927306375 | 1.0126197077531 | 0.345898395700434 | 0.579611774337155 | -5.8840829896782 |
| **Oat** | 0.114258347499998 | 27.11273383375 | 1.01193901318756 | 0.346201261237904 | 0.579611774337155 | -5.88474925596483 |
| **Atp5mk** | 5.0271110675 | 18.37335059625 | 1.0113698422076 | 0.346454665625979 | 0.579611774337155 | -5.88530611132475 |
| **Camk2a** | -0.149832800000002 | 25.8366297425 | -1.0108192368273 | 0.346699942744816 | 0.579611774337155 | -5.8858445849376 |
| **Tpm3** | 5.25835535749999 | 19.20688402375 | 1.01035661855947 | 0.346906129681914 | 0.579611774337155 | -5.8862968443908 |
| **Gria1** | 5.73316173 | 21.0219714725 | 1.00848741504378 | 0.347740204625065 | 0.580343123276405 | -5.88812265099563 |
| **Nap1l1** | 4.7301896275 | 17.33405280375 | 1.0082395515407 | 0.34785092404335 | 0.580343123276405 | -5.88836457408622 |

| | | | | | | |
|---|---|---|---|---|---|---|
| **Sort1** | -0.159439152499999 | 26.71430637375 | -1.00702718404417 | 0.348392880591661 | 0.580601581065585 | -5.88954725789715 |
| **Olfm1** | 5.1139452775 | 18.78311531625 | 1.00636894648522 | 0.348687404932452 | 0.580601581065585 | -5.8901889423953 |
| **Sdhd** | 0.166631382500004 | 24.10921546875 | 1.00560090805131 | 0.349031305312526 | 0.580601581065585 | -5.89093727770018 |
| **Ncs1** | -5.179291255 | 19.0398488075 | -1.00536369672798 | 0.349137573759016 | 0.580601581065585 | -5.8911683189572 |
| **Atp1a1** | -5.30159205 | 17.9683263485714 | -0.920953838775194 | 0.393755201442685 | 0.612694167025697 | -5.89263397361934 |
| **Timm13** | 5.23952131 | 19.193779305 | 1.0030462213379 | 0.350177112811388 | 0.580601581065585 | -5.89342341015772 |
| **Plp2** | 0.190081160000002 | 22.2751136133333 | 0.840519717083479 | 0.440446578639096 | 0.647878837942065 | -5.89352958284103 |
| **Gna13** | 0.639607677500003 | 23.22598958125 | 1.00283224517804 | 0.350273216973677 | 0.580601581065585 | -5.89363143356456 |
| **Hp1bp3** | 4.44961499 | 16.3957853875 | 1.00259708345054 | 0.350378860085191 | 0.580601581065585 | -5.89386001555299 |
| **Grpel1** | -0.193322102500005 | 24.23805425375 | -1.00258027293057 | 0.350386412929139 | 0.580601581065585 | -5.89387635421224 |
| **Wasf1** | -5.09447482749999 | 18.77653643875 | -1.00232268272504 | 0.35050216223829 | 0.580601581065585 | -5.89412668884632 |
| **Fam162a** | -5.4820488475 | 20.20230154375 | -1.0022275182939 | 0.35054493235662 | 0.580601581065585 | -5.89421916081202 |
| **Pygb** | 0.1971141225 | 26.38913438875 | 1.00166912597029 | 0.350795974929534 | 0.580601581065585 | -5.89476162463156 |
| **Impact** | 0.185429287499996 | 22.97988954625 | 1.00059729862222 | 0.351278241522524 | 0.58094774763839 | -5.89580225447396 |
| **Slc8a2** | -0.226009434999995 | 23.5730268125 | -0.999607298795081 | 0.351724149650416 | 0.58094774763839 | -5.8967627102582 |
| **Sdr39u1** | -5.46093677 | 20.150721105 | -0.999204515923345 | 0.351905694379246 | 0.58094774763839 | -5.89715327279241 |
| **Mtx1** | 0.322682749999998 | 23.33297363 | 0.998753734343912 | 0.352108960055196 | 0.58094774763839 | -5.89759024030734 |
| **Slc27a1** | -4.9650266025 | 18.30245044625 | -0.998357610553444 | 0.35228765511578 | 0.58094774763839 | -5.89797410511187 |
| **Pigk** | -0.301934162500004 | 21.62539573875 | -0.99782523604607 | 0.352527925500097 | 0.58094774763839 | -5.89848982730599 |
| **Reep1** | 0.395295674999993 | 22.5484758025 | 0.996016710191907 | 0.353345100559633 | 0.581875492935971 | -5.90024026886391 |
| **Add3** | -0.548307050000012 | 23.0740281725 | -0.9950494238984 | 0.353782770249077 | 0.582177398346611 | -5.9011755272166 |

| | | | | | | |
|---|---|---|---|---|---|---|
| **Cpt2** | 0.129903580000001 | 27.1628963975 | 0.994439214338536 | 0.354059089886043 | 0.582213546515196 | -5.90176518693244 |
| **Nptxr** | -0.133243870000001 | 25.4168503175 | -0.991836838622415 | 0.35523940296817 | 0.583573577681868 | -5.90427692212436 |
| **Ndufaf3** | -0.204275497500003 | 25.02457358875 | -0.991491957581127 | 0.355396053861304 | 0.583573577681868 | -5.90460942573086 |
| **Add1** | 0.167488152499999 | 25.12745418625 | 0.990327979156565 | 0.355925149283119 | 0.584023416995742 | -5.90573099826034 |
| **Cep135** | -0.10045390749999 | 24.20968762125 | -0.988178460753875 | 0.356903837684933 | 0.585209802622359 | -5.90779964044075 |
| **Farp1** | 0.588460150000003 | 21.1771272225 | 0.983263926407386 | 0.359149297652134 | 0.588470109037749 | -5.91251672998504 |
| **Ncl** | -0.259747230000002 | 22.8501036575 | -0.982095515605345 | 0.359684753519452 | 0.588904948523518 | -5.91363562682371 |
| **Camk2b** | 5.2791922275 | 19.78802549625 | 0.981562479705955 | 0.359929236777808 | 0.588904948523518 | -5.9141457452947 |
| **Pgd** | -0.227205847500006 | 28.70357365375 | -0.980033224252571 | 0.360631361491115 | 0.589195198617027 | -5.91560810702123 |
| **Api5** | 0.387299872499998 | 27.29097346625 | 0.979780815777941 | 0.360747351165197 | 0.589195198617027 | -5.91584931113948 |
| **Pakap** | 0.135942612499996 | 26.13053401125 | 0.97949475147457 | 0.360878841616895 | 0.589195198617027 | -5.91612262113125 |
| **Bcas1** | -0.255418949999992 | 23.6314115975 | -0.977072093199989 | 0.361993910346729 | 0.590360289094183 | -5.91843487711597 |
| **Pnp** | 0.448090030000003 | 20.814535825 | 0.976306154188291 | 0.362346999468917 | 0.590360289094183 | -5.9191650224996 |
| **Aldh1l1** | 0.207941940000008 | 21.68407708 | 0.976264544202281 | 0.362366188806172 | 0.590360289094183 | -5.91920467573441 |
| **Macf1** | 0.177224885000012 | 26.477164345 | 0.975092871644197 | 0.362906852710387 | 0.590820615827934 | -5.92032073084022 |
| **Ppp3r1** | -5.139678895 | 19.4102944125 | -0.9742437993369 | 0.363299042608983 | 0.591038740960883 | -5.9211288729922 |
| **Rpl17** | -0.40074499 | 22.020927015 | -0.973447055836967 | 0.363667358725635 | 0.591217744405525 | -5.92188673004294 |
| **Prdm11** | -0.411261305000004 | 22.54625494 | -0.971114643626064 | 0.364747233102964 | 0.592280313701551 | -5.92410263658504 |
| **Septin5** | 4.4815836 | 16.9712978725 | 0.970917840112868 | 0.36483846322376 | 0.592280313701551 | -5.92428942747112 |
| **S100a6** | -0.439768245000003 | 23.13694871 | -0.969322912829776 | 0.365578454748102 | 0.592824421770326 | -5.92580216382935 |
| **Rhob** | 4.56435771 | 17.3259867025 | 0.968659700640669 | 0.365886502080928 | 0.592824421770326 | -5.92643064981412 |

| | | | | | | |
|---|---|---|---|---|---|---|
| **Ap2b1** | 0.124875469999996 | 26.8639000075 | 0.968521769158481 | 0.365950593255339 | 0.592824421770326 | -5.92656131857547 |
| **Tubb4b** | 0.102195729999998 | 26.5555397975 | 0.966989792609535 | 0.366663021072373 | 0.592934810580657 | -5.92801169035314 |
| **Gnb1** | 0.224069954999997 | 22.4850644425 | 0.96673734002106 | 0.36678052340912 | 0.592934810580657 | -5.9282505299087 |
| **Sgip1** | 0.261183800000001 | 22.56851306 | 0.966214607589575 | 0.367023917605879 | 0.592934810580657 | -5.92874492602353 |
| **Dbn1** | 4.7758093875 | 18.17036240625 | 0.965864444274338 | 0.367187029691361 | 0.592934810580657 | -5.92907599527097 |
| **Naxd** | 0.224288625 | 20.0281334525 | 0.965592124574391 | 0.367313919354902 | 0.592934810580657 | -5.92933340328928 |
| **Retreg3** | 0.125689717499998 | 26.76198363375 | 0.960686105129397 | 0.369605688122089 | 0.594815837657702 | -5.93396141957865 |
| **Serbp1** | -0.4950851975 | 18.89850095875 | -0.960575781413971 | 0.369657349768089 | 0.594815837657702 | -5.93406528752819 |
| **Capn2** | 0.46261968 | 22.935251715 | 0.959014396152521 | 0.370389097738256 | 0.594815837657702 | -5.9355343409998 |
| **Csnk2a2** | 0.0888887724999989 | 29.98284935625 | 0.958986533148108 | 0.370402165881517 | 0.594815837657702 | -5.9355605399616 |
| **Ntm** | 4.5684785025 | 17.49884205375 | 0.958822037362321 | 0.370479323922438 | 0.594815837657702 | -5.935715199973 |
| **Ssr4** | -0.264373727499997 | 24.63719441625 | -0.958145710089465 | 0.37079668977232 | 0.594815837657702 | -5.93635087681619 |
| **Rps13** | 0.303401694999991 | 22.62810482 | 0.958141816099729 | 0.370798517624793 | 0.594815837657702 | -5.93635453577836 |
| **Synpo** | -0.363593712500005 | 21.59369012375 | -0.95742483105316 | 0.371135190395618 | 0.594815837657702 | -5.93702805470348 |
| **Gpm6a** | 4.5568944625 | 17.46951111375 | 0.956920929994454 | 0.371371945712788 | 0.594815837657702 | -5.93750117961078 |
| **Rap2a** | -0.298254699999994 | 24.2893461175 | -0.956699645902173 | 0.371475951380649 | 0.594815837657702 | -5.93770888912816 |
| **Ranbp1** | 0.343958089999997 | 21.9333834975 | 0.95625775545505 | 0.371683710790768 | 0.594815837657702 | -5.93812356335138 |
| **Slc12a6** | 0.270487389999992 | 25.3420555325 | 0.955867308488256 | 0.371867357392319 | 0.594815837657702 | -5.93848984199548 |
| **Ehd1** | -0.170757347499997 | 25.13677243125 | -0.955463493582583 | 0.372057364519777 | 0.594815837657702 | -5.93886854212379 |
| **Ppa2** | 0.226524352500007 | 24.35173628625 | 0.955336352035471 | 0.372117203812071 | 0.594815837657702 | -5.9389877512057 |
| **Slc6a17** | -0.136065129999992 | 29.5737346225 | -0.954094847921791 | 0.372701905298546 | 0.595334725211704 | -5.94015116599179 |
| **Nckipsd** | -1.1203715525 | 21.69960936125 | -0.952950079645911 | 0.373241669112019 | 0.595781158017721 | -5.94122291456753 |

| | | | | | | |
|---|---|---|---|---|---|---|
| **Slc7a5** | 0.124470394999996 | 27.3012482075 | 0.94770846967348 | 0.37572072968965 | 0.599016214397003 | -5.94611772116301 |
| **Scamp4** | 0.19443429 | 25.9174123625 | 0.946785847955419 | 0.376158385289689 | 0.599016214397003 | -5.94697717697072 |
| **Rpl4** | 0.100079685000008 | 26.602124875 | 0.946200013313482 | 0.376436483494446 | 0.599016214397003 | -5.94752257297084 |
| **Snph** | 0.201460417500002 | 19.51558326125 | 0.945333859818705 | 0.376847936626478 | 0.599016214397003 | -5.9483284676872 |
| **Acadm** | 0.0992478099999907 | 30.633137205 | 0.945247430877653 | 0.376889012137671 | 0.599016214397003 | -5.94840885290767 |
| **Gng4** | 0.441064782499993 | 20.98397910375 | 0.944803166839109 | 0.377100203121923 | 0.599016214397003 | -5.94882196260613 |
| **Erlin2** | -0.212477405000005 | 24.718055495 | -0.943387968009888 | 0.377773549323906 | 0.5990968372805 | -5.9501369338479 |
| **Nebl** | -0.299690965000003 | 22.711295035 | -0.94251509602941 | 0.378189313418593 | 0.5990968372805 | -5.95094723835573 |
| **Slc1a1** | -0.104487772499994 | 24.79587900625 | -0.94242607626206 | 0.378231734588059 | 0.5990968372805 | -5.95102984506412 |
| **Rpl24** | 0.0940474174999935 | 25.51549734125 | 0.941995730034506 | 0.378436861199873 | 0.5990968372805 | -5.95142910491103 |
| **Pigs** | 0.0877287899999999 | 27.8594477125 | 0.941948033857788 | 0.378459601008127 | 0.5990968372805 | -5.95147334716098 |
| **Prkce** | 0.462385062500012 | 22.39257307625 | 0.940803282734373 | 0.379005687881641 | 0.599226736037522 | -5.95253468835414 |
| **Khdrbs1** | 0.328215355000001 | 22.7840619025 | 0.940435265296792 | 0.379181372107687 | 0.599226736037522 | -5.95287568181668 |
| **Cbln1** | -4.7197487025 | 18.38575617875 | -0.94013027251323 | 0.379327016390725 | 0.599226736037522 | -5.95315820145358 |
| **Atp9a** | 0.152074015 | 26.79181043 | 0.936977778011037 | 0.380834922574816 | 0.600742062781015 | -5.95607430582683 |
| **Arpc3** | -0.341985535000006 | 23.2432910975 | -0.936717356531258 | 0.380959690404472 | 0.600742062781015 | -5.95631486480961 |
| **Fxyd7** | 0.110656517499997 | 23.27528834375 | 0.936479650896199 | 0.381073602078652 | 0.600742062781015 | -5.95653439585471 |
| **Gfap** | 0.455019665000002 | 21.00272554 | 0.93380817237184 | 0.382355582017291 | 0.60221778390419 | -5.95899868461547 |
| **Ctnnb1** | -0.499501500000001 | 22.883863005 | -0.933432972296643 | 0.38253589244067 | 0.60221778390419 | -5.95934435382708 |
| **Coro1c** | -0.104474937499997 | 24.91116488375 | -0.932374151590164 | 0.383045077723666 | 0.602418096179149 | -5.96031926379905 |
| **Clasp2** | 0.059306620000001 | 26.9583663375 | 0.93207404331838 | 0.383189492370835 | 0.602418096179149 | -5.96059543442919 |

| | | | | | | |
|---|---|---|---|---|---|---|
| **Rph3a** | -0.453403447499998 | 20.40519484625 | -0.929916548442464 | 0.38422890690887 | 0.603637589508857 | -5.96257883236164 |
| **Gpr158** | 0.105065155000005 | 28.245029025 | 0.927386819735756 | 0.385450357978281 | 0.605141199871252 | -5.96489993059012 |
| **Megf9** | -0.119169592500015 | 27.98530104875 | -0.926382716615601 | 0.385935987748552 | 0.605214438348412 | -5.96581987614605 |
| **Pip4k2b** | -0.406012124999997 | 22.0333025775 | -0.926197100767863 | 0.386025810392609 | 0.605214438348412 | -5.96598985088936 |
| **Ap2a2** | 0.0844499750000018 | 25.7492492975 | 0.920646388393737 | 0.388719163807695 | 0.60783632489812 | -5.97106069868688 |
| **Copb1** | 0.108997785 | 23.6299098125 | 0.920480104653096 | 0.38880006620089 | 0.60783632489812 | -5.97121224434248 |
| **Stt3b** | 0.256418825000004 | 19.900257055 | 0.920376519204523 | 0.38885047024155 | 0.60783632489812 | -5.97130663810801 |
| **Dnajc8** | -0.138439380000001 | 24.1360273475 | -0.920305116137342 | 0.388885217387775 | 0.60783632489812 | -5.9713717004295 |
| **Uqcc1** | 4.80173211 | 19.03425389 | 0.92001613369469 | 0.389025869801549 | 0.60783632489812 | -5.97163498080492 |
| **Purb** | 0.12473750749999 | 26.22502568375 | 0.918370934450434 | 0.389827341685073 | 0.608673113995315 | -5.97313263928946 |
| **Add2** | 0.218658965000003 | 25.299401505 | 0.917312483816 | 0.390343628071156 | 0.609063779587509 | -5.9740950732573 |
| **Snap47** | -4.211718155 | 16.8045309575 | -0.914767259595501 | 0.391587222020483 | 0.610587977660003 | -5.97640589355591 |
| **Tubb5** | -0.209556567500002 | 24.24000932125 | -0.913251262030346 | 0.392329343812153 | 0.611328705232144 | -5.97777991133562 |
| **Sec23a** | 0.155509127499997 | 23.85719845375 | 0.911107805791655 | 0.393380416339082 | 0.612549505442285 | -5.97971960783965 |
| **Tusc2** | 0.442505957499996 | 22.29038336125 | 0.90982867476398 | 0.394008656121375 | 0.612694167025697 | -5.98087545758632 |
| **Nit2** | 0.127407622500005 | 30.56580109875 | 0.908439259160125 | 0.39469190932303 | 0.613339973143527 | -5.98212953353748 |
| **Golph3** | 0.229236405000005 | 22.7248997375 | 0.906588401264902 | 0.395603450626549 | 0.614339415525218 | -5.98379778911596 |
| **Ddx5** | -0.111071114999998 | 23.9657642225 | -0.905197300228991 | 0.396289594428495 | 0.61498771637073 | -5.98504990383777 |
| **Pkm** | 0.128069232499989 | 22.51757923375 | 0.903418387787633 | 0.397168311084301 | 0.615933123252227 | -5.98664890120855 |
| **Get3** | 0.104208274999994 | 23.5490086075 | 0.902310239332521 | 0.397716426919893 | 0.615933123252227 | -5.98764373468759 |
| **Rps28** | 0.721467249999996 | 22.038250215 | 0.901962543312369 | 0.397888521178182 | 0.615933123252227 | -5.98795568038671 |

| | | | | | | |
|---|---|---|---|---|---|---|
| **Rrbp1** | 0.217016212500006 | 24.16918457375 | 0.901787587965375 | 0.397975137304519 | 0.615933123252227 | -5.98811261124544 |
| **Synpr** | -0.678761612500004 | 21.76236250375 | -0.899893400306385 | 0.398913799887067 | 0.616968708068579 | -5.98981013198648 |
| **Usp14** | -0.149535374999999 | 19.669620715 | -0.899346872514147 | 0.399184936241173 | 0.616971180996654 | -5.99029939756142 |
| **Slc27a4** | -0.181259974999996 | 26.7046816675 | -0.89814745876293 | 0.39978045302424 | 0.617474667322865 | -5.99137232903605 |
| **Twf2** | -0.460827587500006 | 23.66708141125 | -0.89713673160056 | 0.400282796693531 | 0.617833662597096 | -5.99227560238376 |
| **Ccdc47** | 0.229184380000003 | 23.02167704 | 0.896073526456372 | 0.400811726997635 | 0.61783893750838 | -5.99322491588858 |
| **Ckmt1** | 0.109096159999996 | 26.52367218 | 0.896044755662053 | 0.400826047269526 | 0.61783893750838 | -5.99325059248375 |
| **Abi1** | 0.511185779999998 | 22.59750858 | 0.894214016852923 | 0.401738051479735 | 0.618791680940755 | -5.99488311415803 |
| **Dstn** | 0.109190737500001 | 27.18177520125 | 0.893344174247198 | 0.40217191132289 | 0.618791680940755 | -5.99565786075088 |
| **Snx12** | 0.187042025000007 | 23.69446781 | 0.893000072174885 | 0.402343638029659 | 0.618791680940755 | -5.9959641805345 |
| **Dagla** | 0.134308705000002 | 24.79520415 | 0.891583208097959 | 0.403051305503967 | 0.618791680940755 | -5.99722449828627 |
| **Rps19** | 0.0794051550000034 | 29.754953175 | 0.891574179477013 | 0.403055817888119 | 0.618791680940755 | -5.99723252432009 |
| **Mbp** | 0.189761157500001 | 22.88386353875 | 0.891553525127762 | 0.403066140796272 | 0.618791680940755 | -5.99725088485834 |
| **Pabpc1** | 0.141114192500005 | 27.09025600375 | 0.890274493241708 | 0.40370577290745 | 0.61894526265402 | -5.9983872207713 |
| **Hacd3** | -0.200349522500002 | 20.67515300125 | -0.889946892020363 | 0.403869723708713 | 0.61894526265402 | -5.99867806691969 |
| **Ivd** | 0.104649764999994 | 26.043334105 | 0.889731830402926 | 0.403977379818744 | 0.61894526265402 | -5.99886895408631 |
| **Hbb-b1** | 0.511164152499997 | 21.73856824375 | 0.887338341700762 | 0.405176947391791 | 0.620367914769104 | -6.00099095047238 |
| **Cldn11** | 0.112441079999996 | 28.775203725 | 0.886347528592989 | 0.40567428998456 | 0.620714204394824 | -6.00186806053555 |
| **Mtdh** | 0.214397339999994 | 22.913904055 | 0.885000204858529 | 0.406351305412782 | 0.620813130260891 | -6.00305953154834 |
| **Gpr37l1** | 0.270648149999992 | 26.63855884 | 0.884954580708434 | 0.406374245592341 | 0.620813130260891 | -6.00309985310907 |
| **Acsl1** | 0.113907767499992 | 27.79115248875 | 0.8845999450912 | 0.40655259163874 | 0.620813130260891 | -6.00341321598368 |

| | | | | | | |
|---|---|---|---|---|---|---|
| **Fdx2** | 4.581450375 | 18.513877815 | 0.88397510255301 | 0.406866964648356 | 0.620878988053391 | -6.00396509816685 |
| **Gmfb** | -0.347825814999993 | 21.6277784425 | -0.882607153832518 | 0.407555835747203 | 0.621515861442603 | -6.00517224404549 |
| **Sh3glb2** | -0.248078680000003 | 22.704608525 | -0.881985314335145 | 0.407869264610657 | 0.621579724829424 | -6.00572049875269 |
| **Rgs7** | -0.367198832500002 | 23.36991250625 | -0.877991070597051 | 0.409886722766378 | 0.62416646730493 | -6.00923481399721 |
| **P4hb** | 0.126310342500002 | 26.78254801875 | 0.877436782961322 | 0.410167265672015 | 0.62416646730493 | -6.00972150526292 |
| **Tubb3** | 0.231341307499999 | 24.59702298625 | 0.876059719173599 | 0.41086485086789 | 0.62416646730493 | -6.01092958143905 |
| **Bsn** | -0.0661212099999986 | 27.76736516 | -0.875994126094477 | 0.410898100295475 | 0.62416646730493 | -6.01098708774762 |
| **Bcs1l** | 0.299568817500003 | 22.76539169625 | 0.875931116417023 | 0.41093004203955 | 0.62416646730493 | -6.01104232594543 |
| **Mlycd** | 0.332980737499998 | 21.04328152125 | 0.874431752008706 | 0.41169065684839 | 0.624601691856942 | -6.01235583390583 |
| **Lin7a** | 0.5595557975 | 21.66499534875 | 0.874290588080646 | 0.411762321106215 | 0.624601691856942 | -6.01247940781737 |
| **Gnao1** | 0.338015710000004 | 21.5135127375 | 0.873226763967469 | 0.412302682786464 | 0.625007179402793 | -6.01341016356051 |
| **Coasy** | 0.124039535000005 | 23.74810345 | 0.872287453345633 | 0.412780229338738 | 0.625316972174964 | -6.01423123390499 |
| **Gria2** | 4.8300937575 | 19.37505920875 | 0.868622860898586 | 0.414647173012658 | 0.627324192334687 | -6.01742782301777 |
| **Pdk3** | -0.120380545000007 | 25.0947732725 | -0.868572385438145 | 0.414672930881445 | 0.627324192334687 | -6.01747177766812 |
| **Mpdu1** | -0.410224679999999 | 23.77409443 | -0.867934165267404 | 0.414998718263441 | 0.627324192334687 | -6.01802737253685 |
| **Cd200** | -4.4102001925 | 18.33818201625 | -0.867537168673783 | 0.415201464127152 | 0.627324192334687 | -6.01837280937504 |
| **Prodh** | 0.138289882500001 | 26.68642973125 | 0.866759789608008 | 0.415598679932061 | 0.627510144039899 | -6.01904886274781 |
| **Dnm3** | 0.0823395924999995 | 29.68827500125 | 0.86474141989968 | 0.416631293361855 | 0.628654601519634 | -6.02080190135526 |
| **Fmnl1** | -0.262878752500008 | 23.13689159375 | -0.860595799266068 | 0.418758071198423 | 0.631091285845362 | -6.02439233317017 |
| **Rab35** | 0.0824302374999881 | 27.46796502375 | 0.860518929303273 | 0.418797581126738 | 0.631091285845362 | -6.02445877863453 |

| | | | | | | |
|---|---|---|---|---|---|---|
| **Gsk3a** | 0.5924888925 | 23.35222488375 | 0.857952698769274 | 0.42011813406957 | 0.632664742687661 | -6.02667427718684 |
| **Pip4k2a** | 0.230997910000017 | 23.774545895 | 0.851113725736836 | 0.423652103088595 | 0.637567168947925 | -6.03255269711516 |
| **Adcyap1r1** | 0.233582457499999 | 20.79672978125 | 0.850101697502117 | 0.424176875175565 | 0.63793749492567 | -6.03341938068895 |
| **Nsf** | -0.108852262500005 | 25.73496645375 | -0.849361134427088 | 0.424561179808972 | 0.638096218373432 | -6.0340530619057 |
| **Fabp5** | -0.127215679999999 | 26.8741106175 | -0.848436580001099 | 0.425041316079068 | 0.638398669622695 | -6.03484355763033 |
| **Rpl13** | 0.336691062500002 | 21.03903035625 | 0.844278329979296 | 0.427205594717618 | 0.638994619799008 | -6.03839031337534 |
| **Pals2** | 0.299845175000002 | 22.6386914925 | 0.844137382419268 | 0.427279093348912 | 0.638994619799008 | -6.03851028818217 |
| **Septin2** | 0.169606825000002 | 24.0942578625 | 0.843819043295135 | 0.427445128144225 | 0.638994619799008 | -6.03878119966355 |
| **Pgk1** | -0.19753197 | 24.76772514 | -0.843404027811967 | 0.427661655570953 | 0.638994619799008 | -6.0391342606768 |
| **Scrn1** | -0.195904389999995 | 23.7290571175 | -0.842644621796708 | 0.428058066828792 | 0.638994619799008 | -6.03977993867535 |
| **Slc2a13** | -0.0957845099999943 | 25.23875394 | -0.842279741873913 | 0.428248628518412 | 0.638994619799008 | -6.0400900078486 |
| **Golga7** | -0.330455599999997 | 23.409580575 | -0.842207103380544 | 0.428286571860226 | 0.638994619799008 | -6.04015172197549 |
| **Cacnb4** | 0.0625407375000009 | 27.71781829625 | 0.842074672999877 | 0.42835575421376 | 0.638994619799008 | -6.04026422463475 |
| **Atl1** | 0.183467637500005 | 23.21477689875 | 0.842024506851233 | 0.428381963376527 | 0.638994619799008 | -6.04030683821102 |
| **Arhgef40** | 0.114581802500005 | 25.43601925875 | 0.841970635125549 | 0.42841010978814 | 0.638994619799008 | -6.04035259720746 |
| **Gpx4** | -0.2650474625 | 24.17480982625 | -0.8416160785172 | 0.428595388401993 | 0.638994619799008 | -6.04065370104965 |
| **Armc1** | 0.239405909999999 | 24.4181249625 | 0.841137895394476 | 0.428845360793469 | 0.638994619799008 | -6.04105963164394 |
| **Wdr7** | -4.8054416675 | 20.61784699875 | -0.8407137720781 5 | 0.429067160607722 | 0.638994619799008 | -6.0414195151256 |
| **Cat** | -0.0740923699999954 | 25.96067375 | -0.838736020175772 | 0.430102532256272 | 0.640120088644087 | -6.04309577354859 |
| **Nomo1** | 0.436332605000004 | 23.72954432 | 0.837544974909189 | 0.430726917706256 | 0.640632823021196 | -6.04410371546568 |

| | | | | | | |
|---|---|---|---|---|---|---|
| **Acox1** | 0.299054364999989 | 23.4126936675 | 0.835280355457397 | 0.431915893603066 | 0.641338687752704 | -6.04601699530074 |
| **Cyb5b** | 0.148304580000008 | 24.421619605 | 0.835153976682877 | 0.431982314289196 | 0.641338687752704 | -6.04612364394052 |
| **Ckap5** | 0.195047400000004 | 27.3712611 | 0.835040325897262 | 0.43204205177574 | 0.641338687752704 | -6.04621954052879 |
| **Crk** | -0.142839767500003 | 26.66230200625 | -0.832352055883292 | 0.433456788469296 | 0.643021768506947 | -6.04848477744139 |
| **Gnb2** | -0.115216432499992 | 26.24873357625 | -0.831457845484107 | 0.433928109480011 | 0.64330404313455 | -6.04923696175058 |
| **Marcks** | 0.0996954399999979 | 26.6731937875 | 0.830650024786552 | 0.434354209962217 | 0.64336572151162 | -6.04991591428329 |
| **Cldnd1** | 0.17340722749999 | 24.44806971875 | 0.830177785845151 | 0.434603439383747 | 0.64336572151162 | -6.05031257122096 |
| **Ppp2r2a** | 0.206868822499999 | 21.13257352875 | 0.82978101411314 | 0.43481291881978 | 0.64336572151162 | -6.05064569812603 |
| **Mlc1** | -0.322563247499993 | 21.53222904625 | -0.827678147923125 | 0.435924346555017 | 0.644593558956352 | -6.05240909374943 |
| **C3orf18** | 0.116321260000007 | 20.794878325 | 0.826998887384376 | 0.43628378723518 | 0.644708579071225 | -6.05297792364967 |
| **Ubl4a** | 0.463155499999999 | 23.479139155 | 0.826457807997434 | 0.436570257956571 | 0.644715690621027 | -6.05343076579806 |
| **Ngef** | -3.9015934 | 16.91412485 | -0.823258722984076 | 0.438266720199226 | 0.646747247447696 | -6.05610323300138 |
| **Phf24** | -0.102490532499999 | 25.04714482125 | -0.82279875835215 | 0.438511021423689 | 0.646747247447696 | -6.056486787333 |
| **Coro1a** | -0.173271274999998 | 21.8786678525 | -0.821048018087526 | 0.439441775245127 | 0.647702655206759 | -6.05794509478052 |
| **Ccz1** | 0.340550445000002 | 23.6110899925 | 0.819661995124622 | 0.440179623771285 | 0.647878837942065 | -6.05909781052131 |
| **Micos13** | 0.241029734999998 | 23.122577185 | 0.819482551284604 | 0.440275214616582 | 0.647878837942065 | -6.05924693279808 |
| **Rpl18** | 0.0781002200000032 | 29.6317352 | 0.818697724688715 | 0.440693469058889 | 0.647878837942065 | -6.05989883034094 |
| **Src** | -0.147852087499992 | 23.85821078625 | -0.816506035701297 | 0.441862963280788 | 0.648405061383468 | -6.06171660619073 |
| **Dpysl4** | -0.118229197499996 | 22.86554972875 | -0.81626014349659 | 0.441994308909496 | 0.648405061383468 | -6.06192029982406 |
| **Eef1d** | -0.396587310000005 | 24.363036385 | -0.815961196733465 | 0.442154031227358 | 0.648405061383468 | -6.06216787563338 |

| | | | | | | |
|---|---|---|---|---|---|---|
| **Atp6v1c1** | 0.078906994999997 | 27.36728298 | 0.815904194122601 | 0.442184491402181 | 0.648405061383468 | -6.06221507452033 |
| **Ppp3cb** | 0.132380667499991 | 26.79878227875 | 0.810396656035919 | 0.44513450266037 | 0.652312981171311 | -6.06676267081726 |
| **Mgll** | -0.119250767500006 | 22.78060439125 | -0.8094274060813 | 0.445655092404427 | 0.652658033598038 | -6.06756037571251 |
| **Slc20a2** | -0.215166240000002 | 21.9345465975 | -0.808123212138744 | 0.44635625708044 | 0.653266926123483 | -6.06863250929004 |
| **Nucb1** | 0.124854044999996 | 24.0163920475 | 0.80645837648711 | 0.447252435373967 | 0.654149277588392 | -6.06999905432031 |
| **Itga6** | 0.213978604999998 | 22.8285838925 | 0.805941921515412 | 0.447530698428755 | 0.654149277588392 | -6.07042250557754 |
| **Agap3** | 0.162979992499995 | 24.45170639125 | 0.804891037370414 | 0.448097283440303 | 0.654462452370448 | -6.0712834579882 |
| **Myl6** | 0.1633178875 | 21.4524014471429 | 0.704287435428755 | 0.508519424584654 | 0.700062728583937 | -6.07140934628417 |
| **Palm** | 0.104986897499998 | 21.65056618875 | 0.804484159247614 | 0.448316786944894 | 0.654462452370448 | -6.07161655123124 |
| **Esd** | -0.145769982499999 | 25.70662598625 | -0.802418964389731 | 0.449432083223514 | 0.655672427341379 | -6.07330510177326 |
| **Plscr3** | 0.249168014999995 | 26.7231968275 | 0.799427758492919 | 0.451050902893163 | 0.657614978804109 | -6.07574445051692 |
| **Acsl6** | 0.0855558450000053 | 30.40244494 | 0.797962101177242 | 0.451845589973938 | 0.658354268268837 | -6.07693696607669 |
| **Rdh14** | 0.129691702499997 | 27.03191876125 | 0.795545793188205 | 0.453157854268433 | 0.6598462648985 | -6.07889902984403 |
| **Synj2bp** | 4.06699323 | 18.056760635 | 0.79334209669705 | 0.454356962452101 | 0.660963311882812 | -6.08068417069219 |
| **Rab3a** | -0.18837562500001 | 23.10961067 | -0.793074881618058 | 0.454502513282458 | 0.660963311882812 | -6.08090035477492 |
| **Chst3** | 0.371083547500007 | 23.77589870125 | 0.792243713816012 | 0.454955453627754 | 0.661201925939002 | -6.08157240698662 |
| **Sv2a** | 0.0949361824999961 | 26.24763181875 | 0.787609597026463 | 0.45748653173556 | 0.664458547679376 | -6.0853087028977 |
| **Efr3b** | 0.198002732499994 | 25.65168905375 | 0.784897732721411 | 0.458972220614612 | 0.666193667081069 | -6.08748674717562 |
| **Tmpo** | -0.419904915 | 21.6094780325 | -0.781439369633159 | 0.460871707679668 | 0.668526830721647 | -6.09025530361927 |
| **Aldh4a1** | 0.218557500000003 | 24.9422188925 | 0.78085541889719 | 0.461192973495812 | 0.668569168037944 | -6.09072177756488 |
| **Cadm4** | 0.0729107975000076 | 33.01190271125 | 0.7756019717285 | 0.46409013816675 | 0.67234324447069 | -6.09490531613591 |

| | | | | | | |
|---|---|---|---|---|---|---|
| **Ubqln2** | -0.178299572499999 | 26.70830680125 | -0.772861572880944 | 0.465606358795912 | 0.673811332166873 | -6.09707827492169 |
| **Slc2a3** | 0.164855347499994 | 24.56502758375 | 0.772706543171914 | 0.465692235687196 | 0.673811332166873 | -6.09720101180161 |
| **H1-0** | 0.103131922500005 | 20.02226400125 | 0.770794991661221 | 0.466752008650999 | 0.674918097158646 | -6.09871269701143 |
| **Cacng8** | 0.1578616 | 25.855455645 | 0.766948284043705 | 0.468889636550716 | 0.677581046757948 | -6.10174525444542 |
| **Atad1** | 3.722727525 | 17.293742185 | 0.762085878035295 | 0.471601232168476 | 0.680940788721494 | -6.10556037139529 |
| **Rap2c** | 0.0878943950000064 | 28.0222994325 | 0.76117891992485 | 0.472108189127552 | 0.680940788721494 | -6.10626973459254 |
| **Emc3** | -0.178892052500004 | 23.18415754625 | -0.761141726004283 | 0.472128987091067 | 0.680940788721494 | -6.1062988101211 |
| **Cadm1** | 0.0992210174999997 | 32.20094729375 | 0.760649062808941 | 0.472404531450298 | 0.680940788721494 | -6.10668382630647 |
| **Gdap1l1** | 0.522786222499999 | 21.40506067375 | 0.751198977609714 | 0.477711034838605 | 0.687829025544517 | -6.11402850941629 |
| **Rab27b** | -0.445688810000007 | 21.0967319625 | -0.750603086917738 | 0.478046988741784 | 0.687829025544517 | -6.11448905047066 |
| **Rps16** | 0.310896774999996 | 22.0800664725 | 0.749989263655733 | 0.478393219216886 | 0.687829025544517 | -6.11496312887988 |
| **Wdr37** | 5.12264229749999 | 11.96180749125 | 0.749926574515425 | 0.478428588886557 | 0.687829025544517 | -6.11501152762345 |
| **Slc25a18** | 0.0951418849999968 | 19.6854570325 | 0.74914815046167 | 0.47886792814687 | 0.687829025544517 | -6.11561222072617 |
| **Vcam1** | -4.30106773 | 10.0331377525 | -0.74816901896473 | 0.479420931440435 | 0.687829025544517 | -6.11636704835279 |
| **Sec62** | -5.176380265 | 12.1162455375 | -0.747994534091117 | 0.479519523776545 | 0.687829025544517 | -6.11650147401854 |
| **Tomm34** | 0.0825738849999951 | 27.6314811 | 0.747874758788162 | 0.479587210471406 | 0.687829025544517 | -6.11659373532911 |
| **Cycs** | -0.116106207499993 | 25.79304172875 | -0.746195224500002 | 0.480537015473186 | 0.688759692184172 | -6.11788614460988 |
| **Pdzd7** | 0.726955552499998 | 21.05551616125 | 0.740723361431071 | 0.483640202093234 | 0.692036800163639 | -6.12207973273196 |
| **Pspc1** | 0.6027033075 | 20.22597216125 | 0.740647841144965 | 0.483683124590036 | 0.692036800163639 | -6.12213742814793 |
| **Hsp90b1** | 5.078972995 | 11.9912454975 | 0.740564671386627 | 0.4837303976679 | 0.692036800163639 | -6.1222009617877 |
| **Lin7c** | -5.52986153 | 13.0412190925 | -0.739506816785703 | 0.484331944052526 | 0.692464597086965 | -6.12300853313526 |

| | | | | | | |
|---|---|---|---|---|---|---|
| **Nrxn1** | -3.748211365 | 17.9178746325 | -0.738600093542216 | 0.484847947774278 | 0.69253969749693 | -6.12369995182269 |
| **Ina** | 0.161387542499998 | 26.22514986375 | 0.738351340133072 | 0.484989574088064 | 0.69253969749693 | -6.1238895122421 |
| **Bckdhb** | -0.196460530000007 | 27.5901796425 | -0.737660056263042 | 0.485383297607298 | 0.69257844612381 | -6.12441601507508 |
| **Anxa2** | 0.109265877499997 | 25.97122053625 | 0.734537279484087 | 0.487164544916518 | 0.69257844612381 | -6.12678920403523 |
| **Tst** | -0.406714402499993 | 21.74099990625 | -0.734416534197872 | 0.487233505955196 | 0.69257844612381 | -6.12688079420152 |
| **Cdk5** | -5.5306842875 | 13.17878041125 | -0.733689304046783 | 0.487648985047761 | 0.69257844612381 | -6.12743215721459 |
| **Slc25a46** | -3.60666819 | 17.4424462275 | -0.733609880541346 | 0.487694375339693 | 0.69257844612381 | -6.1274923455221 |
| **Mapre1** | 0.347277052499997 | 20.26738425875 | 0.733460614148182 | 0.487779688235835 | 0.69257844612381 | -6.12760544683103 |
| **Nefl** | -0.129106727500009 | 22.64818791375 | -0.733412506669647 | 0.487807186080652 | 0.69257844612381 | -6.12764189439965 |
| **Atp6ap1** | -0.0924001325000035 | 25.65890042625 | -0.732936920450982 | 0.488079082798133 | 0.69257844612381 | -6.12800210253105 |
| **Ppp1r9b** | 4.81196309 | 11.4830565925 | 0.732610524062326 | 0.488265744724362 | 0.69257844612381 | -6.12824919970079 |
| **Hnrnpa2b1** | -0.0717945674999996 | 29.17314403375 | -0.732532663674111 | 0.48831027909705 | 0.69257844612381 | -6.12830812980397 |
| **Cadm3** | 0.155008222500005 | 20.87106132125 | 0.732020709994264 | 0.488603172219405 | 0.69257844612381 | -6.12869547905589 |
| **Gdap1** | 0.0656137199999947 | 26.9221053925 | 0.73188121923706 | 0.488682996323134 | 0.69257844612381 | -6.12880097925745 |
| **Vti1a** | 5.2480584175 | 12.54229951625 | 0.731414579461499 | 0.488950095647041 | 0.69257844612381 | -6.12915378589352 |
| **Gnpda2** | 0.345314859999998 | 21.4565116425 | 0.730598859025302 | 0.489417237664212 | 0.692676982174218 | -6.12977005818738 |
| **Mrpl12** | -0.336518924999996 | 21.9896878875 | -0.730236436028596 | 0.489624882987281 | 0.692676982174218 | -6.1300436791359 |
| **Abhd16a** | 0.0852728975000048 | 25.70685604625 | 0.72896584544288 | 0.490353312243634 | 0.692889854736484 | -6.13100203328731 |
| **Slc39a10** | -5.6119212525 | 13.45487051375 | -0.72891798929173 | 0.490380762198823 | 0.692889854736484 | -6.13103810146054 |
| **Erp44** | 0.244948552500002 | 24.62998404875 | 0.728139622090187 | 0.490827371230487 | 0.693044630067344 | -6.13162445744889 |
| **Stxbp5l** | -0.0889720724999883 | 24.69283248125 | -0.727671941049251 | 0.491095845333872 | 0.693044630067344 | -6.13197651227819 |

| | | | | | | |
|---|---|---|---|---|---|---|
| **Tceal5** | 0.0854149849999928 | 22.9708523925 | 0.726307934387489 | 0.491879413960848 | 0.6930688830958 | -6.13300219112021 |
| **Coq6** | -5.24357843 | 12.6287675275 | -0.725386758478073 | 0.492409061326997 | 0.6930688830958 | -6.13369395122227 |
| **Comtd1** | -0.280326897500004 | 22.49572388125 | -0.725133990545426 | 0.492554460901384 | 0.6930688830958 | -6.13388363730977 |
| **Pdia6** | 5.53181093 | 13.33637632 | 0.724550581682096 | 0.492890163228118 | 0.6930688830958 | -6.13432123299985 |
| **Rpn1** | -4.860685975 | 11.7262627425 | -0.724066611130701 | 0.493168761933326 | 0.6930688830958 | -6.13468401539231 |
| **Scp2** | 0.123441210000003 | 19.1848996625 | 0.723658693129556 | 0.493403661582563 | 0.6930688830958 | -6.13498962853382 |
| **Mtnd4** | 0.226770145000003 | 24.0585561575 | 0.723461150020983 | 0.493517443360743 | 0.6930688830958 | -6.13513757550559 |
| **Syn2** | 5.4083378975 | 13.05060463625 | 0.722868942604469 | 0.493858649469262 | 0.6930688830958 | -6.13558089392889 |
| **Rab8b** | -0.237244737499999 | 24.08321862625 | -0.722178460742431 | 0.494256673863886 | 0.6930688830958 | -6.13609738807608 |
| **Hmgb1** | -0.209604370000008 | 22.9790245825 | -0.721946406823898 | 0.494390487529672 | 0.6930688830958 | -6.13627087442995 |
| **Adcy9** | 0.0917713424999924 | 25.24191762375 | 0.721854248025078 | 0.494443637437939 | 0.6930688830958 | -6.1363397602732 |
| **Sec22b** | 5.11162432 | 12.39689653 | 0.720025536967413 | 0.495499072225243 | 0.694123241324102 | -6.13770511197777 |
| **Omg** | 5.6040592125 | 13.61000664375 | 0.718919482102012 | 0.496138147685645 | 0.694593406759903 | -6.13852947823484 |
| **Crkl** | -0.106930922500002 | 23.69328311125 | -0.716325705858109 | 0.497638949642543 | 0.696268677097666 | -6.14045842421688 |
| **Por** | 0.308216512499996 | 22.17799291125 | 0.715325415662215 | 0.498218529747742 | 0.696416088366534 | -6.14120072849506 |
| **Nap1l4** | 4.81737552 | 11.75694824 | 0.71509379407002 | 0.498352797179722 | 0.696416088366534 | -6.1413724856955 |
| **Eef1a2** | -0.236162207499994 | 24.00950691625 | -0.713835128520371 | 0.499082840739596 | 0.6970107519542 | -6.14230500658451 |
| **Scai** | 0.0931972074999976 | 25.35951103875 | 0.712901532762217 | 0.499624791647908 | 0.697075488259865 | -6.14299578040434 |
| **Thsd7a** | 0.161507400000005 | 23.3896355125 | 0.712534830415338 | 0.499837767146005 | 0.697075488259865 | -6.14326689402616 |
| **L2hgdh** | -0.120603737499998 | 21.54668701125 | -0.71191055315133 | 0.500200475127768 | 0.697075488259865 | -6.1437281651404 |
| **Inpp1** | 0.105833527500007 | 27.03212947375 | 0.711657885516086 | 0.500347325124927 | 0.697075488259865 | -6.14391475973204 |

| | | | | | | |
|---|---|---|---|---|---|---|
| **Cse1l** | -0.133281857499998 | 25.87245267875 | -0.710674072330096 | 0.500919384223607 | 0.697447974749293 | -6.14464076274839 |
| **RTRAF** | -4.9747451075 | 12.22791259625 | -0.709672398442673 | 0.501502267727653 | 0.697835070412521 | -6.14537906073378 |
| **Syt1** | 5.3074766375 | 13.06108138375 | 0.70886957386259 | 0.501969758524772 | 0.698061225554802 | -6.14597014875511 |
| **Stx16** | 0.350147557499998 | 21.99939470875 | 0.707333424552989 | 0.502865062052982 | 0.698881680048133 | -6.1470995533343 |
| **Cap1** | -4.721057875 | 11.663809505 | -0.705711476506284 | 0.503811499794864 | 0.69940922657129 | -6.14828975369398 |
| **Hnrnpk** | -4.73539796000001 | 11.70743701 | -0.705184490054463 | 0.504119255879773 | 0.69940922657129 | -6.14867595561732 |
| **Radil** | 0.0653759300000019 | 25.016997995 | 0.704940736681786 | 0.5042616473568 | 0.69940922657129 | -6.1488545063169 |
| **Scamp3** | 5.30589107 | 13.1311341 | 0.70458952848058 | 0.504466855862472 | 0.69940922657129 | -6.1491116749212 |
| **Mapre3** | 4.6053178025 | 11.40576770375 | 0.703658780332906 | 0.505010948086131 | 0.699728609495818 | -6.14979267246711 |
| **H4c1** | 0.0784874500000008 | 21.24612182 | 0.70313243606277 | 0.505318804577076 | 0.699728609495818 | -6.15017743823149 |
| **Tma7** | 0.252302784999994 | 24.97240538 | 0.702211842085029 | 0.505857548946762 | 0.699728609495818 | -6.15084981077743 |
| **Them6** | -0.0804448224999987 | 28.45261720125 | -0.70207395265079 | 0.505938275845284 | 0.699728609495818 | -6.15095045552535 |
| **Cntn2** | -0.408226687500004 | 22.78450079375 | -0.701583157232358 | 0.506225678167355 | 0.699728609495818 | -6.15130854632353 |
| **Atp2b2** | -4.742770535 | 11.7944360875 | -0.7006076566399 | 0.5067972307691 | 0.700062728583937 | -6.15201964395595 |
| **Tagln2** | 0.211405352499998 | 24.88853245625 | 0.699101314559521 | 0.507680628210709 | 0.700062728583937 | -6.15311602708134 |
| **Pmpcb** | -0.082786902499997 | 19.84687128625 | -0.698492275261197 | 0.508038083397787 | 0.700062728583937 | -6.15355873553533 |
| **Gas7** | 5.2933755975 | 13.21700070875 | 0.697847178590961 | 0.508416878774242 | 0.700062728583937 | -6.15402729105236 |
| **Rmnd1** | -0.233668564999995 | 25.3840358675 | -0.697648259177868 | 0.508533719423033 | 0.700062728583937 | -6.15417169761241 |
| **Tubb4a** | -0.0816641924999963 | 26.23071359125 | -0.697521362621634 | 0.50860826458501 | 0.700062728583937 | -6.15426380025768 |
| **Cdipt** | -5.54222433 | 13.8799954 | -0.695491734908309 | 0.509801525525278 | 0.70118091957119 | -6.15573495673142 |
| **Acly** | 0.0835258000000003 | 19.5007495325 | 0.69509785807214 | 0.510033303226751 | 0.70118091957119 | -6.15602002579595 |

| | | | | | | |
|---|---|---|---|---|---|---|
| **Slc4a4** | 0.196359994999998 | 23.11154242 | 0.694478429485312 | 0.510397944848637 | 0.701261041871867 | -6.1564680561833 |
| **Sptan1** | -4.64305566 | 11.66298884 | -0.693314067669759 | 0.511083829390317 | 0.701482221806598 | -6.15730930064752 |
| **Adck1** | 4.8684788075 | 12.23744315375 | 0.692749527278082 | 0.5114165938904 | 0.701482221806598 | -6.15771673853879 |
| **Rdh11** | 0.224270300000001 | 19.3588788375 | 0.692471787128198 | 0.51158035712225 | 0.701482221806598 | -6.15791708277309 |
| **Scamp5** | -0.0967502474999975 | 19.71703328375 | -0.691727455975976 | 0.512019401657492 | 0.701482221806598 | -6.15845365377823 |
| **Cotl1** | 5.26945322 | 13.2799332175 | 0.690977986739374 | 0.512461721723786 | 0.701482221806598 | -6.15899342439896 |
| **Capn5** | 0.0440419999999904 | 26.35914574 | 0.690896792049394 | 0.512509655768163 | 0.701482221806598 | -6.15905187073532 |
| **Actr1a** | 4.954204995 | 12.498711155 | 0.690064831386669 | 0.513000977565827 | 0.701482221806598 | -6.15965039815388 |
| **Rap1gap2** | 0.271976572500002 | 22.06996707125 | 0.690048560543925 | 0.51301058947324 | 0.701482221806598 | -6.15966209746764 |
| **Fus** | -5.1661350725 | 13.05178889125 | -0.689207126208023 | 0.513507819664562 | 0.701742924902795 | -6.16026679315088 |
| **Bphl** | -4.9957236375 | 12.63245140875 | -0.687829194076492 | 0.514322751212735 | 0.702437218094243 | -6.16125566242603 |
| **Ssb** | 4.96211940999999 | 12.56857929 | 0.68728007480774 | 0.514647740278068 | 0.702461942454679 | -6.16164925865949 |
| **Syt12** | -0.243223830000002 | 24.5131292925 | -0.686499541373717 | 0.515109915214892 | 0.702504087090899 | -6.16220825851257 |
| **Gabrb3** | 0.106038730000009 | 22.6329797275 | 0.686043798890957 | 0.515379895534469 | 0.702504087090899 | -6.1625343960585 |
| **Krt2** | 0.244249722500005 | 25.20822042125 | 0.685673493376712 | 0.515599329974972 | 0.702504087090899 | -6.16279925509773 |
| **Slc5a3** | 4.734626345 | 12.02223405 | 0.68494959460405 | 0.51602846821808 | 0.702670531678277 | -6.1633166616651 |
| **Rap2b** | -4.9469318175 | 12.60488046125 | -0.682893507800317 | 0.51724859308156 | 0.703913216149637 | -6.1647836632723 |
| **Rpl26** | 4.618843505 | 11.7952530675 | 0.681245521070919 | 0.518227871601031 | 0.704826855671277 | -6.1659567242948 |
| **Ubxn6** | 0.160147002500004 | 22.36469422375 | 0.68056090885203 | 0.518635034133499 | 0.70496175364108 | -6.16644331742964 |
| **Fahd2** | 0.0925162449999952 | 22.803180815 | 0.678249643244514 | 0.520011128398149 | 0.706412743562826 | -6.16808292480844 |
| **Cd47** | -4.76531877 | 12.23553507 | -0.677340641668187 | 0.520552970532871 | 0.706594255026354 | -6.16872643964926 |

| | | | | | | |
|---|---|---|---|---|---|---|
| **Rab12** | -0.196838225000008 | 21.77936575 | -0.676829191637892 | 0.520857995841584 | 0.706594255026354 | -6.16908818336502 |
| **Srcin1** | -0.282126720000008 | 22.08183465 | -0.676344690589495 | 0.521147053640576 | 0.706594255026354 | -6.16943064697336 |
| **Podxl2** | 5.3728849775 | 13.82765727625 | 0.675597492780922 | 0.521593038336083 | 0.706594255026354 | -6.16995837605385 |
| **Pip5k1c** | 0.1043169525 | 23.39883731125 | 0.675438111595457 | 0.52168820052186 | 0.706594255026354 | -6.17007087769244 |
| **Psmc5** | 0.071589837500003 | 27.66711822625 | 0.673910002527965 | 0.522601151839182 | 0.707412203761022 | -6.1711483432513 |
| **Hnrnpc** | -0.0670595824999936 | 19.31307347375 | -0.672696848435407 | 0.523326657175662 | 0.707742424396972 | -6.17200222061027 |
| **Snap25** | 5.0602107775 | 13.08448078375 | 0.672232978204945 | 0.523604234941743 | 0.707742424396972 | -6.17232836077782 |
| **Mfsd6** | 0.067495802499991 | 25.44758604875 | 0.671951557665002 | 0.523772681052193 | 0.707742424396972 | -6.17252612771366 |
| **Arf4** | -0.326017629999996 | 22.002199135 | -0.670785156468437 | 0.52447120340109 | 0.707759690638744 | -6.17334504050461 |
| **Pin1** | 0.0667605550000019 | 26.262494785 | 0.670453993167928 | 0.524669634301839 | 0.707759690638744 | -6.17357731906561 |
| **Sfpq** | 0.0539115449999912 | 29.70169015 | 0.670381529648607 | 0.524713060294429 | 0.707759690638744 | -6.17362813176814 |
| **Psmc6** | 4.756463225 | 12.356302455 | 0.6685775745081 | 0.525794867243163 | 0.708540154353128 | -6.17489155179426 |
| **Tuba4a** | -0.213996555000001 | 21.133373215 | -0.668232508155088 | 0.526001958911489 | 0.708540154353128 | -6.17513288417608 |
| **Faah** | -0.1667476975 | 23.14206796375 | -0.666838946437971 | 0.526838828255022 | 0.708540154353128 | -6.17610640554123 |
| **Micu3** | -0.0565601999999963 | 20.520166935 | -0.6667982329558 | 0.526863290336942 | 0.708540154353128 | -6.17613482067341 |
| **Prrt3** | -0.144128115000004 | 22.9171480375 | -0.66671172670976 | 0.526915268684135 | 0.708540154353128 | -6.17619519088426 |
| **Epb41l3** | -0.123335965000003 | 20.42284893 | -0.666322929445846 | 0.527148922177098 | 0.708540154353128 | -6.176466436745 |
| **Synj1** | -4.626322165 | 12.04172732 | -0.66548884947 | 0.527650394831856 | 0.708797977564623 | -6.17704786911613 |
| **Lypla2** | 0.161424602499995 | 20.72033535875 | 0.663374121566331 | 0.528923173040988 | 0.710090993015145 | -6.17851917902892 |
| **Rab3b** | -0.362191054999993 | 20.32430378 | -0.662651765862515 | 0.529358374011113 | 0.710258685880093 | -6.17902081501751 |
| **Krt76** | 4.55533618 | 11.9546871375 | 0.661701291496417 | 0.529931352724406 | 0.710610935199863 | -6.17968013780569 |

| | | | | | | |
|---|---|---|---|---|---|---|
| **Slc7a10** | 0.0979479749999967 | 27.90217541 | 0.661114422252367 | 0.530285331733978 | 0.710669276545126 | -6.18008682203304 |
| **Prkcb** | -4.6151933575 | 12.13767144375 | -0.660127208578351 | 0.530881118470224 | 0.711051421988498 | -6.18077022080718 |
| **Ttf2** | 0.191776677500009 | 23.56101238125 | 0.659228322973732 | 0.531423963095613 | 0.711362252354303 | -6.18139169606283 |
| **Hsd11b1** | -0.129443139999999 | 24.181275645 | -0.658533675896186 | 0.53184370399214 | 0.711508029478672 | -6.18187145555061 |
| **Krt79** | -5.2831213475 | 13.91549319625 | -0.657195481237274 | 0.53265289255 | 0.712165038868266 | -6.18279443220829 |
| **Gnai1** | -0.102043170000002 | 25.1583330725 | -0.656692791940323 | 0.532957060542307 | 0.712165038868266 | -6.18314072041001 |
| **Slc25a25** | 5.12776052000001 | 13.5648423725 | 0.65608724414183 | 0.533323610146638 | 0.712239056957791 | -6.18355755612281 |
| **H2az1** | 0.0576322624999861 | 26.59878921125 | 0.655009714192366 | 0.533976248052563 | 0.712535211252926 | -6.18429845138009 |
| **Scamp1** | 5.0709203 | 13.439857635 | 0.65469337374669 | 0.534167943429454 | 0.712535211252926 | -6.18451575983073 |
| **Cdc37** | 0.244327994999999 | 23.80313634 | 0.651766593356821 | 0.535943538936714 | 0.714487338745567 | -6.18652192238346 |
| **Hspa5** | 0.116611625000004 | 26.506618165 | 0.65076399420606 | 0.536552629781217 | 0.714838415391947 | -6.18720733726268 |
| **Dync1i1** | 0.0649212500000047 | 25.4483364 | 0.650305241666171 | 0.53683147053681 | 0.714838415391947 | -6.18752064846519 |
| **Rps27** | 4.4514089075 | 11.87774998875 | 0.649655621734736 | 0.537226478740253 | 0.714948494090953 | -6.18796398260449 |
| **Rheb** | -0.273875725000007 | 21.6835062125 | -0.647617688494266 | 0.538466833648427 | 0.716182790366792 | -6.1893522424621 |
| **Ncstn** | 4.57968031 | 12.3323618325 | 0.643633904081482 | 0.540896608562065 | 0.718996711381282 | -6.19205493558614 |
| **Tecpr1** | 0.203376069999997 | 23.6387310625 | 0.642061915310648 | 0.54185724705309 | 0.719855623043833 | -6.19311736297025 |
| **Dync1li1** | 0.161444784999997 | 22.31793826 | 0.639608258540849 | 0.543358767083956 | 0.721179737322006 | -6.1947710810892 |
| **Arl6ip5** | 4.845497695 | 13.1307102625 | 0.639403679956347 | 0.543484074652888 | 0.721179737322006 | -6.1949087105996 |
| **Bpnt2** | 0.100498442499998 | 19.47698351125 | 0.638451201608645 | 0.544067715952632 | 0.721203154850035 | -6.19554897479141 |
| **Myo5a** | -0.239488902500003 | 20.44765545375 | -0.63834654453811 | 0.544131869124513 | 0.721203154850035 | -6.19561927474763 |
| **Rit1** | 0.068848437499998 | 26.80269641125 | 0.635526883950067 | 0.545862022397246 | 0.723077644251908 | -6.19750945328327 |

| | | | | | | |
|---|---|---|---|---|---|---|
| **Aldh1b1** | 0.083589857499998 | 21.69471431375 | 0.633713397554911 | 0.546976559704791 | 0.724098733449329 | -6.19872122376247 |
| **Rpl27a** | 0.0962163875000002 | 25.83522592625 | 0.63320527871525 | 0.547289089771025 | 0.724098733449329 | -6.19906019862528 |
| **Sptbn1** | 4.4678275325 | 12.22270984125 | 0.632729425138832 | 0.547581873132717 | 0.724098733449329 | -6.1993774304197 |
| **Rab1A** | 0.216280050000016 | 26.01383295 | 0.631017916843525 | 0.548635719864966 | 0.725073419613687 | -6.20051667511494 |
| **Sik3** | -0.179933342499996 | 24.73120664625 | -0.630285837872789 | 0.54908686817884 | 0.725250918211982 | -6.20100313946306 |
| **Septin9** | 0.0604812449999912 | 25.1856862075 | 0.629751749703224 | 0.549416145774892 | 0.725267334301458 | -6.20135772362735 |
| **Hba** | 0.0657975324999889 | 27.75511537375 | 0.627898263588767 | 0.550559793403014 | 0.726358136656772 | -6.20258619529607 |
| **Nmt1** | -0.330846552499999 | 21.33113520625 | -0.625515983078558 | 0.552031840823916 | 0.727880693344438 | -6.20416042613321 |
| **Dclk2** | -0.0910958349999973 | 26.8599021675 | -0.622970113190648 | 0.553607604514193 | 0.729342079331783 | -6.20583688167087 |
| **Sel1l** | 0.143683162500004 | 23.30383933875 | 0.622662950837866 | 0.553797905946622 | 0.729342079331783 | -6.20603873724157 |
| **Coro1b** | 0.0772484624999983 | 24.91103974375 | 0.62218183730761 | 0.554096057648742 | 0.729342079331783 | -6.20635472908011 |
| **Msn** | 0.150494442500005 | 24.44541484625 | 0.620132367273553 | 0.555367223130345 | 0.730371234245114 | -6.20769837161125 |
| **Nfs1** | 0.1410491625 | 25.16854878125 | 0.619623051536734 | 0.55568339405619 | 0.730371234245114 | -6.20803166943628 |
| **Cltc** | -4.36641989 | 12.169078555 | -0.619378629482686 | 0.555835163851021 | 0.730371234245114 | -6.20819153343248 |
| **Psmd3** | 0.0673288399999947 | 27.55360422 | 0.618646085505086 | 0.556290174226211 | 0.730549746875386 | -6.20867031670131 |
| **Ppp3ca** | -0.114037645000003 | 20.657112825 | -0.616611885873151 | 0.557554865593504 | 0.731790761091474 | -6.2099972020588 |
| **Septin11** | 0.0933419025000006 | 28.42343875125 | 0.61529561893368 | 0.558374126875097 | 0.73244606098401 | -6.21085371253117 |
| **Cfl2** | 0.156107375000005 | 21.85504109 | 0.613602834107346 | 0.55942879780282 | 0.733203686921553 | -6.211952828187 |
| **Tbc1d24** | -0.134514099999997 | 21.8813089475 | -0.613140133241826 | 0.559717285861095 | 0.733203686921553 | -6.21225278713842 |
| **Srgap3** | 0.181863560000004 | 21.6891773575 | 0.61213453379458 | 0.560344570991677 | 0.733203686921553 | -6.21290399909798 |
| **Entpd2** | -0.204606957499998 | 20.35665540375 | -0.612095918323789 | 0.56036866739689 | 0.733203686921553 | -6.21292898690686 |

| | | | | | | |
|---|---|---|---|---|---|---|
| **Slc12a9** | 0.0800193000000036 | 25.4612493975 | 0.611800106086759 | 0.560553277462961 | 0.733203686921553 | -6.21312035831378 |
| **Rtn4** | -0.0912110050000052 | 22.62702912 | -0.607491963577866 | 0.563246009352207 | 0.736305034498688 | -6.21589808067089 |
| **Atp6v1g2** | -0.1255763375 | 22.67762414875 | -0.606895267050619 | 0.563619570459954 | 0.736353193237795 | -6.21628142378822 |
| **Ctnna1** | 0.0809007650000027 | 24.177938245 | 0.605975089562118 | 0.564195934969138 | 0.736353193237795 | -6.21687192423995 |
| **Agpat1** | 1.16948562249999 | 23.27814190125 | 0.605573111068209 | 0.564447828967853 | 0.736353193237795 | -6.21712963209394 |
| **Chmp6** | 0.0892311300000017 | 20.1404653675 | 0.605378796419682 | 0.564569617357942 | 0.736353193237795 | -6.21725415207843 |
| **Dbi** | -0.0948796350000087 | 24.8480110075 | -0.604065994714433 | 0.565392835633028 | 0.73700694804328 | -6.21809447952946 |
| **H2ac11** | -0.0669968250000039 | 27.561850495 | -0.597581017402814 | 0.56946978456316 | 0.741372384779438 | -6.22222154050108 |
| **Camk2g** | -0.320288522499997 | 21.08527996625 | -0.597488822227501 | 0.569527870069343 | 0.741372384779438 | -6.22227992575641 |
| **Cacng3** | 0.124118557499997 | 21.40414636625 | 0.597194344037134 | 0.569713422816527 | 0.741372384779438 | -6.22246635841028 |
| **Dtna** | -0.0856516949999993 | 27.2356121675 | -0.596269419750054 | 0.570296455270594 | 0.741586797752468 | -6.22305138673698 |
| **Rgs7bp** | 0.136608504999991 | 23.6145474825 | 0.595905188574717 | 0.570526147156879 | 0.741586797752468 | -6.22328154490985 |
| **Flot1** | 0.323711839999994 | 21.843437805 | 0.594931400355868 | 0.571140505474566 | 0.741964027827061 | -6.22389626271648 |
| **Trip10** | 0.177034105000004 | 20.8303209375 | 0.594012211234218 | 0.571720773628686 | 0.742041226764258 | -6.22447568537138 |
| **Sirt4** | -0.16842384000001 | 24.0726537175 | -0.593810297067466 | 0.57184828484585 | 0.742041226764258 | -6.22460285670818 |
| **Phgdh** | -0.313148330000001 | 21.18287251 | -0.592256493054704 | 0.572830088315444 | 0.742220614265325 | -6.22558018672132 |
| **Cmpk1** | 0.188998997500008 | 21.08905391625 | 0.592062363186857 | 0.572952822639159 | 0.742220614265325 | -6.22570213102142 |
| **Rp1** | 0.05469140000001 | 31.1293100475 | 0.591943815353446 | 0.573027779458049 | 0.742220614265325 | -6.22577658015407 |
| **Rac1** | -0.0707984699999997 | 29.36260688 | -0.591381275498237 | 0.573383546981967 | 0.742220614265325 | -6.22612967759272 |
| **Plxnb1** | 0.204951592500002 | 24.56413970625 | 0.591026782769506 | 0.57360780543266 | 0.742220614265325 | -6.22635203200515 |
| **Tubb6** | 0.397744522500002 | 19.82886458375 | 0.590225948367999 | 0.574114616324372 | 0.742228044912948 | -6.2268539111838 |

| | | | | | | |
|---|---|---|---|---|---|---|
| **Krt10** | -0.0908018825000028 | 17.92043727125 | -0.589533422100916 | 0.574553094841073 | 0.742228044912948 | -6.22728742039374 |
| **Actn4** | 0.0929979225000039 | 23.21578842125 | 0.589480953088733 | 0.574586323978045 | 0.742228044912948 | -6.22732024640061 |
| **Napg** | -2.74443116750001 | 15.61084138125 | -0.587834769165928 | 0.575629437645871 | 0.743156109854145 | -6.22834880676727 |
| **Gja1** | -0.143502667500005 | 23.27815194625 | -0.58647562736408 | 0.576491496366711 | 0.743849512504736 | -6.22919606632601 |
| **Slc4a8** | 0.0857957200000001 | 24.10322154 | 0.585055955705717 | 0.577392748604748 | 0.744592676933109 | -6.23007917159222 |
| **Prkca** | -0.22503558 | 21.9947451575 | -0.582814376180016 | 0.578817438550923 | 0.746009637862085 | -6.23146961410526 |
| **Rock2** | -0.0509083024999946 | 26.72485838625 | -0.581914867781605 | 0.579389715142314 | 0.746106680970553 | -6.23202622149091 |
| **Caskin1** | 0.189943427499998 | 20.45805637625 | 0.581671445789243 | 0.579544639041347 | 0.746106680970553 | -6.23217671534397 |
| **Itm2c** | -0.0661653625000014 | 26.26117670375 | -0.579054858655932 | 0.581211456680164 | 0.747821301430128 | -6.23379081050796 |
| **Dhodh** | -0.108230279999997 | 19.9519548825 | -0.578555473175518 | 0.581529889272883 | 0.747821301430128 | -6.23409811959665 |
| **Hibch** | -0.206141695000014 | 25.1225802375 | -0.576338794491902 | 0.582944565571742 | 0.748861877573482 | -6.23545931408573 |
| **Tstd3** | 0.189169282500004 | 24.48777492625 | 0.575459149342946 | 0.583506500324781 | 0.748861877573482 | -6.23599816764202 |
| **Akap5** | 0.0649943324999924 | 25.46901722125 | 0.575154728032323 | 0.583701043190528 | 0.748861877573482 | -6.23618447681733 |
| **Syngr3** | 0.118155172499989 | 24.83042084125 | 0.574742310484353 | 0.583964661345535 | 0.748861877573482 | -6.23643673860244 |
| **Slmap** | -0.247347415 | 23.09619548 | -0.57472635944217 | 0.58397485865822 | 0.748861877573482 | -6.23644649202371 |
| **Nono** | -0.429117729999998 | 20.4517457225 | -0.573625648971807 | 0.584678776765359 | 0.74934474804922 | -6.23711894051999 |
| **Fam136a** | -0.1383506325 | 25.06286031375 | -0.572195014845697 | 0.585594412090534 | 0.750098270439413 | -6.2379912025269 |
| **Plxna4** | 0.364818570000001 | 22.1361367275 | 0.57116146118622 | 0.586256418375498 | 0.75052625372568 | -6.23862013596195 |
| **Stk24** | -0.201183465 | 23.585882695 | -0.569643752793007 | 0.587229308221867 | 0.751351529636587 | -6.23954181775283 |
| **Homer3** | 0.243087644999989 | 23.77029853 | 0.56684885303528 | 0.589023319759607 | 0.753124347447631 | -6.24123330089824 |

| | | | | | | |
|---|---|---|---|---|---|---|
| **Stt3a** | 0.1061981575 | 24.45065077125 | 0.566460389234303 | 0.589272916679208 | 0.753124347447631 | -6.24146780317251 |
| **Sccpdh** | -0.117250335000001 | 25.513324735 | -0.5656227502977 | 0.589811323661336 | 0.753391807958035 | -6.24197296059957 |
| **H2bc7** | 0.0512151874999986 | 24.75169874375 | 0.564908893191467 | 0.590270388344055 | 0.753557679263548 | -6.2424029329776 |
| **Calr** | 3.865550685 | 11.8036143 | 0.562940956887243 | 0.591536971879666 | 0.754753694889942 | -6.24358571222024 |
| **Ern1** | -0.0501012974999995 | 28.35900512875 | -0.562155027457872 | 0.592043234079332 | 0.75497880936356 | -6.24405702775333 |
| **Lrrc57** | -0.104488455000002 | 25.284817295 | -0.561151623372653 | 0.59268994054493 | 0.755214694847958 | -6.24465789130651 |
| **Myh10** | -0.112726457500003 | 22.21809023375 | -0.560844327758843 | 0.592888076296103 | 0.755214694847958 | -6.24484171244604 |
| **Rps18** | 0.0682981300000023 | 25.6144248225 | 0.558985298064416 | 0.594087525414599 | 0.756321660552845 | -6.24595181258535 |
| **Atp6v1a** | 0.219185827500006 | 22.33333332625 | 0.558191417606912 | 0.59460015531516 | 0.756423171559388 | -6.24642484831493 |
| **Tuba1a** | -0.0832614524999968 | 23.72364027375 | -0.557838409625971 | 0.594828182091262 | 0.756423171559388 | -6.24663499253608 |
| **Preb** | -0.184536272499997 | 22.90792413125 | -0.557118442712184 | 0.595293399967699 | 0.756594443379269 | -6.24706321086956 |
| **Septin8** | 0.0436034199999966 | 26.1596884125 | 0.552435840901087 | 0.598324125667604 | 0.760024374946252 | -6.24983601139245 |
| **Hnrnpa1** | -0.239329189999996 | 23.3950515575 | -0.549617521737379 | 0.600152386995121 | 0.761923912275004 | -6.25149459044352 |
| **Arf2** | -0.0987327799999989 | 25.0552718575 | -0.548559608559764 | 0.600839465948396 | 0.76237335784694 | -6.25211517333014 |
| **Gnai3** | 0.0835401299999958 | 25.627171485 | 0.54707465190634 | 0.601804634220549 | 0.763174962731766 | -6.25298442260976 |
| **Jup** | 0.272801024999996 | 24.0450355475 | 0.546223357704352 | 0.602358333496666 | 0.763454166873681 | -6.25348177422524 |
| **Ppp1cc** | 0.0362736724999984 | 29.15041059375 | 0.545625174247342 | 0.602747573632569 | 0.763524734944633 | -6.25383082760828 |
| **Plxna1** | 0.0896421299999979 | 25.3816878375 | 0.545032104771529 | 0.603133623930543 | 0.76359118649171 | -6.25417655180619 |
| **Krt5** | 0.0672211474999962 | 29.78845542875 | 0.544477138909479 | 0.603494995446677 | 0.763626337522081 | -6.25449975259791 |
| **Ubqln1** | -0.0703242550000027 | 24.73294983 | -0.542880875612743 | 0.60453508702438 | 0.764519786850168 | -6.25542770535461 |
| **Tsfm** | 0.06956315750001 | 25.00330156625 | 0.541411212677393 | 0.60549356505838 | 0.765309094654131 | -6.25627985811538 |

| | | | | | | |
|---|---|---|---|---|---|---|
| **Gnal** | -0.20770066 | 23.251605985 | -0.539447551460757 | 0.606775525806456 | 0.765802803853788 | -6.25741514538624 |
| **Gria4** | 0.167957137499997 | 22.82074150375 | 0.539402673157805 | 0.606804841734366 | 0.765802803853788 | -6.25744104755343 |
| **Krt31** | -0.0701002474999974 | 28.91092399875 | -0.539275608038754 | 0.606887848925632 | 0.765802803853788 | -6.25751437433697 |
| **Krt16** | 0.0759405374999922 | 27.96181106375 | 0.536043795576168 | 0.609001179737658 | 0.768046115933608 | -6.25937406886719 |
| **Atp4a** | 0.0880621650000002 | 19.6270937528571 | 0.382389806478777 | 0.715785734110501 | 0.852462822777802 | -6.25952722141178 |
| **Apex1** | -0.0654530150000028 | 27.7224045725 | -0.534671656466549 | 0.609899662649293 | 0.768755687116868 | -6.26016054536354 |
| **Gpd1l** | 0.0610523875000055 | 28.63668593875 | 0.534024169383085 | 0.610323892275168 | 0.768867027747859 | -6.26053102706903 |
| **Gphn** | 0.0546290574999979 | 21.87073429875 | 0.531562927808085 | 0.611937955721958 | 0.770369475325902 | -6.26193555078433 |
| **Syngap1** | 0.0628171749999993 | 25.544542765 | 0.531179467933143 | 0.61218963548179 | 0.770369475325902 | -6.26215383845535 |
| **Hnrnph1** | -0.169603984999998 | 22.3622050975 | -0.528705535782577 | 0.613814730383452 | 0.771990064751495 | -6.26355866664119 |
| **Tfrc** | 0.0689082900000031 | 32.1851535075 | 0.527830288109745 | 0.614390231136545 | 0.772289532713647 | -6.26405423501172 |
| **Ube2v1** | 0.0579276149999934 | 26.696266375 | 0.526124339629042 | 0.615512783707134 | 0.772696420736967 | -6.26501797968579 |
| **Tln2** | -0.0523438175000024 | 20.62671351875 | -0.525905078930893 | 0.615657142812342 | 0.772696420736967 | -6.26514163897878 |
| **Sfn** | -0.0774436199999968 | 19.982723815 | -0.525799537644636 | 0.615726636707832 | 0.772696420736967 | -6.26520114556159 |
| **Snd1** | -0.186486692500004 | 24.53144434375 | -0.520780830139674 | 0.619036113398021 | 0.776380122679685 | -6.26801811658631 |
| **Gabrb1** | 0.0465231450000019 | 27.10416135 | 0.520320164758737 | 0.61934036872569 | 0.776380122679685 | -6.26827543866812 |
| **Ephb2** | 0.111953857499998 | 22.55916803625 | 0.517362026171932 | 0.621296043610541 | 0.778405387971827 | -6.26992281736228 |
| **Kif5b** | 0.0925391324999971 | 20.08358688125 | 0.515147713549209 | 0.622762126835551 | 0.779471791594996 | -6.27115029228512 |
| **Aplp2** | 0.176387864999992 | 26.0778968475 | 0.514964740380459 | 0.622883354983706 | 0.779471791594996 | -6.27125150374208 |
| **Eci2** | 0.0798904699999987 | 24.3987923625 | 0.514533988081359 | 0.62316879799862 | 0.779471791594996 | -6.2714896429984 |
| **Kcna6** | 0.146930455 | 21.98472235 | 0.510001275624042 | 0.626176680463896 | 0.782806347122806 | -6.27398437136278 |

| | | | | | | |
|---|---|---|---|---|---|---|
| **Gcat** | 0.092118469999992 | 24.7071075 | 0.507230392374334 | 0.628019213898956 | 0.784466748483055 | -6.27549936896937 |
| **Bag6** | -0.155075230000001 | 21.104237855 | -0.506973354786308 | 0.628190279584727 | 0.784466748483055 | -6.27563951892748 |
| **Dnaja1** | 0.0690111149999986 | 26.9343886925 | 0.506424762386065 | 0.62855546553538 | 0.78449479858805 | -6.27593841954144 |
| **Stx4** | -0.226534062500001 | 21.25088915875 | -0.503594979834969 | 0.630440969544794 | 0.786419280265959 | -6.27747546866912 |
| **Arl8b** | 0.0950702949999922 | 21.2922005125 | 0.50246360207951 | 0.631195647190501 | 0.78658524432449 | -6.27808776554607 |
| **Hnrnpd** | 0.352529939999997 | 21.34604598 | 0.502365226396778 | 0.631261290443027 | 0.78658524432449 | -6.27814094576169 |
| **Ero1a** | 0.219376212499995 | 20.34324627125 | 0.501419110471096 | 0.63189278940385 | 0.786943740449083 | -6.27865190725041 |
| **Akap12** | -0.120333989999999 | 22.520617985 | -0.499306143408941 | 0.633304316424459 | 0.788272746218372 | -6.2797898171772 |
| **Esyt2** | 0.118799245000005 | 26.3545841375 | 0.497976188377548 | 0.634193613048832 | 0.788950641450422 | -6.28050376127809 |
| **Armc10** | 0.0685218499999962 | 28.3061746225 | 0.494018473256108 | 0.636843858061786 | 0.79155530772337 | -6.28261787906674 |
| **Enpep** | -0.120659584999999 | 24.035283065 | -0.493816925739059 | 0.636978976333092 | 0.79155530772337 | -6.28272512180544 |
| **Psd3** | 0.0642180700000026 | 25.7492400375 | 0.491606878702278 | 0.638461576890116 | 0.792967199946541 | -6.28389841262435 |
| **Gabra3** | 0.037931835000002 | 29.1666652875 | 0.489724868620972 | 0.639725522949735 | 0.794106139930555 | -6.2848936950458 |
| **Rab8a** | -0.0615212075000038 | 29.26077156875 | -0.48829052817679 | 0.640689684217492 | 0.794871917167392 | -6.28564984754979 |
| **Kcnd2** | -0.0599279775000063 | 23.78136219125 | -0.485688454336353 | 0.642440703808588 | 0.796401738426378 | -6.28701633418435 |
| **Adgrb1** | 0.208528855000004 | 21.66218471 | 0.485424275808296 | 0.642618615497387 | 0.796401738426378 | -6.28715468820293 |
| **Atl3** | 0.0521395524999981 | 28.71126848875 | 0.483602348004417 | 0.643846285598622 | 0.797210767124827 | -6.28810694826704 |
| **Stxbp3** | 0.176187832499998 | 23.20383669625 | 0.48332109427347 | 0.644035909878234 | 0.797210767124827 | -6.28825365284165 |
| **Vbp1** | -0.109537884999995 | 23.11886896 | -0.482905352166894 | 0.644316260017881 | 0.797210767124827 | -6.28847036214766 |
| **Grm4** | 0.093223452500002 | 24.18387786875 | 0.482306892690466 | 0.644719932776116 | 0.797247443902425 | -6.28878200935941 |
| **Dync1i2** | -0.0423754474999924 | 29.84945273625 | -0.481828823401087 | 0.645042492838484 | 0.797247443902425 | -6.28903070468211 |

| | | | | | | |
|---|---|---|---|---|---|---|
| **Arsa** | 0.0653597224999984 | 24.04005340875 | 0.47998188029994 | 0.646289425087269 | 0.798357525107803 | -6.28998933831602 |
| **Camlg** | -0.0750255274999887 | 22.82122694375 | -0.47447922667086 | 0.650011708971697 | 0.802363870872944 | -6.29282505534151 |
| **Txnrd2** | -0.106601417499995 | 25.18798515875 | -0.474098276774282 | 0.650269804410688 | 0.802363870872944 | -6.293020242506 |
| **Ubxn4** | 0.0734937499999972 | 25.8718009225 | 0.473537827208023 | 0.65064960584787 | 0.802363870872944 | -6.29330713357921 |
| **Sh3gl1** | -0.0475570675000014 | 28.12375714375 | -0.47311712390709 | 0.650934778598103 | 0.802363870872944 | -6.29352228094201 |
| **Tmem266** | -0.106549494999996 | 19.058339905 | -0.471936719228304 | 0.651735249601017 | 0.802504192091906 | -6.29412498473442 |
| **Hnrnpab** | -0.2587457825 | 21.45818266875 | -0.471892057532702 | 0.65176554588916 | 0.802504192091906 | -6.29414776095299 |
| **Timm22** | -0.378801425000002 | 19.70507268 | -0.4713985090634 | 0.652100392001287 | 0.802504192091906 | -6.2943993227099 |
| **Nlgn1** | 0.3711006375 | 24.43301263875 | 0.470794823655533 | 0.65251007776586 | 0.80257687695113 | -6.29470668661336 |
| **Caprin1** | 0.216508410000003 | 19.6068507075 | 0.470274611684154 | 0.652863218792201 | 0.802579971973871 | -6.29497125489225 |
| **Acsf2** | -0.0572357950000004 | 21.4689006525 | -0.46821746317387 | 0.654260634379965 | 0.803866125655254 | -6.29601479434233 |
| **Cryzl2** | 0.0529420000000087 | 23.5131606925 | 0.467552391558162 | 0.65471273611032 | 0.80399004986938 | -6.29635125257662 |
| **Hexb** | 0.0985192950000027 | 23.065034985 | 0.466143705541886 | 0.655670847113636 | 0.804734889567353 | -6.29706242514341 |
| **Rbmxl1** | 0.169968667499997 | 21.85476890875 | 0.46420480079086 | 0.656990727596123 | 0.805922709253765 | -6.29803799076474 |
| **Rpl5** | 0.0494479575000035 | 28.91102467625 | 0.461708025987926 | 0.658692315674537 | 0.807103732261585 | -6.29928863738559 |
| **Cox7a2l** | 0.0472183175000076 | 23.95235563875 | 0.461471962422027 | 0.658853309486257 | 0.807103732261585 | -6.29940655559945 |
| **Slc7a8** | 0.0560591825000003 | 27.65673829375 | 0.461240324024918 | 0.659011304323679 | 0.807103732261585 | -6.29952220839215 |
| **Sh3bgrl** | -0.0595620974999989 | 25.27678934375 | -0.460063620317941 | 0.659814194710901 | 0.807654915343985 | -6.30010887393723 |
| **Myo1d** | 0.105403232499995 | 24.68734032875 | 0.458650711677185 | 0.660778891779733 | 0.807747505465517 | -6.30081144595032 |
| **Padi2** | 0.0850027974999961 | 23.74523103625 | 0.458636110501419 | 0.660788864715693 | 0.807747505465517 | -6.30081869583267 |

| | | | | | | |
|---|---|---|---|---|---|---|
| **Pam16** | -0.211562309999994 | 22.7555281375 | -0.458402431274016 | 0.660948483065493 | 0.807747505465517 | -6.30093469447688 |
| **Pdap1** | 0.226341259999998 | 22.6968653025 | 0.454755545849598 | 0.66344200774755 | 0.810362196229531 | -6.30273782014154 |
| **Dnpep** | 0.292639157499998 | 21.49865663875 | 0.45323033682674 | 0.664486223492408 | 0.811204781639531 | -6.30348791540138 |
| **Sncg** | 0.0919678300000051 | 21.42274773 | 0.449656766775525 | 0.666935969940435 | 0.81352526710828 | -6.30523611281133 |
| **Cops4** | 0.317400452499996 | 21.44419617375 | 0.448750490894415 | 0.667557936958024 | 0.81352526710828 | -6.30567739563564 |
| **Macrod1** | 0.0538928899999931 | 30.3884592425 | 0.448553861431727 | 0.6676929187872 | 0.81352526710828 | -6.30577302764006 |
| **H3-5** | 0.102405185000002 | 24.14540965 | 0.448385304534621 | 0.667808639972241 | 0.81352526710828 | -6.30585497497614 |
| **Thnsl1** | 0.1460567775 | 22.22659537125 | 0.44600282272012 | 0.669445351928514 | 0.815085324768281 | -6.30701016549971 |
| **Rhot2** | 0.0551645450000002 | 26.6059977325 | 0.439299623740447 | 0.674060679952903 | 0.820268419145239 | -6.31022922290551 |
| **Capza1** | -0.244761005000004 | 23.86349892 | -0.434714665721987 | 0.677226298461578 | 0.823682782772875 | -6.312404555802 |
| **H3c1** | 0.098723415000002 | 20.713242845 | 0.432096603776183 | 0.679037067144422 | 0.825035124796934 | -6.31363702614511 |
| **Chchd2** | 0.118473965 | 24.5278676775 | 0.432064834565692 | 0.679059054223427 | 0.825035124796934 | -6.31365193851227 |
| **Gabra2** | 0.229899865000004 | 23.774824355 | 0.431266213416421 | 0.679611880721056 | 0.825268750647479 | -6.314026468266 |
| **Iscu** | -0.0626430975000076 | 24.73434242625 | -0.43015843973713 | 0.680379062660197 | 0.825762287608266 | -6.31454489686108 |
| **Agpat5** | 0.0474521625000008 | 23.09230011625 | 0.428484734719218 | 0.681538951173907 | 0.826357958038476 | -6.31532578453635 |
| **Plxnc1** | -0.0878085325000093 | 24.10140793125 | -0.427693425213594 | 0.682087657578505 | 0.826357958038476 | -6.31569397704785 |
| **Tmed4** | -0.0527950299999951 | 27.8043548975 | -0.427542852492047 | 0.682192090565029 | 0.826357958038476 | -6.31576396488553 |
| **Kbtbd11** | 0.119859657499997 | 22.49061295875 | 0.427367225019834 | 0.682313910306998 | 0.826357958038476 | -6.31584556898043 |
| **Reep2** | -0.0509940425000046 | 21.95759010125 | -0.425756780145041 | 0.68343143222299 | 0.827273690300595 | -6.31659237145523 |
| **Syncrip** | 0.0442925324999983 | 28.91228710375 | 0.422769444905421 | 0.685506673497875 | 0.829347133000336 | -6.31797060167774 |
| **Fam177a1** | 0.134116192500002 | 24.20249875125 | 0.421114762291051 | 0.686657410491546 | 0.83030048209992 | -6.31873004551177 |

| | | | | | | |
|---|---|---|---|---|---|---|
| **Sigmar1** | 0.153799974999995 | 20.22681591 | 0.413443782497308 | 0.692003810016862 | 0.836323506403694 | -6.32221385945445 |
| **Tdrkh** | -0.0502757749999958 | 23.792089895 | -0.412660736191486 | 0.692550638962463 | 0.836542697934078 | -6.32256606408808 |
| **Psmd2** | -0.246197192499999 | 19.17478210125 | -0.41180246461117 | 0.693150227539851 | 0.836713683406109 | -6.32295137556603 |
| **Tubal3** | -0.0421866450000046 | 29.6766040175 | -0.411411739566891 | 0.693423266676011 | 0.836713683406109 | -6.32312653476888 |
| **Tjp2** | -0.0475455599999997 | 31.99546702 | -0.406766216179038 | 0.696673318379355 | 0.840192426644017 | -6.3251969772075 |
| **Sar1b** | -0.184975512499989 | 22.06115238125 | -0.405502642253822 | 0.697558519990691 | 0.840816983811844 | -6.32575626539136 |
| **Rpl22** | -0.0621309725000039 | 24.63450011875 | -0.404760327364139 | 0.698078788571926 | 0.841001235284809 | -6.32608405972869 |
| **Grm7** | -0.353540337500004 | 22.57080543375 | -0.401635494141198 | 0.700270817527669 | 0.843198264766351 | -6.32745766615596 |
| **Aplp1** | 0.0715365199999987 | 28.654187195 | 0.400960694436085 | 0.700744586163606 | 0.843325109215823 | -6.32775296190374 |
| **Abhd3** | -0.0987065149999964 | 22.839124065 | -0.398879824122103 | 0.702206443524953 | 0.844640330650876 | -6.32866058235835 |
| **Fabp7** | -0.0338714033333289 | 23.0068916842857 | -0.144305315557519 | 0.890140688168173 | 0.96550795329598 | -6.32975756962939 |
| **Slc25a42** | 0.0574598575000032 | 20.27858392875 | 0.395159997455363 | 0.70482309035007 | 0.847342465237033 | -6.33027185402274 |
| **Rps21** | 0.0317295524999963 | 30.84827790875 | 0.392909800977066 | 0.706408048846645 | 0.848668752222709 | -6.33123955317842 |
| **Prep** | -0.0818174075000044 | 20.66985730875 | -0.392469708923998 | 0.70671821812234 | 0.848668752222709 | -6.33142819858631 |
| **Tpd52** | 0.095938827500003 | 20.80425891125 | 0.392015243772224 | 0.707038580379513 | 0.848668752222709 | -6.33162279321816 |
| **Plxna2** | 0.0841669500000002 | 25.0509935575 | 0.390811142673636 | 0.707887686520655 | 0.849070238793673 | -6.33213732914225 |
| **Psmd1** | -0.0999212449999973 | 23.52786003 | -0.389034263499023 | 0.709141522116687 | 0.849070238793673 | -6.3328938629683 |
| **Fam241b** | 0.0499423149999956 | 27.09578497 | 0.388935900480605 | 0.709210959369319 | 0.849070238793673 | -6.33293564635181 |
| **C1orf198** | -0.0932661200000098 | 23.001066275 | -0.38861304473782 | 0.70943889336697 | 0.849070238793673 | -6.33307272050502 |
| **Maip1** | -0.0685393999999917 | 24.9467257675 | -0.388397009412353 | 0.709591430786443 | 0.849070238793673 | -6.33316438141434 |

| | | | | | | |
|---|---|---|---|---|---|---|
| **Ppp5c** | -0.0421891524999971 | 22.79064307125 | -0.388386750263743 | 0.709598674885232 | 0.849070238793673 | -6.33316873302315 |
| **Rras** | 0.0496159324999965 | 25.24596711625 | 0.385883289691979 | 0.711367362866706 | 0.850721497155705 | -6.33422733821269 |
| **Hebp1** | -0.191952185000002 | 20.800488275 | -0.385381645838767 | 0.711722003955078 | 0.850721497155705 | -6.33443867469099 |
| **Plxnb2** | 0.123231574999995 | 21.63941659 | 0.381799539766034 | 0.714256634555368 | 0.852016252909094 | -6.33594012977478 |
| **Rbm3** | 0.0866888000000001 | 20.98776168 | 0.381725650938269 | 0.714308958001134 | 0.852016252909094 | -6.33597095938875 |
| **Rpl9** | -0.0406432300000041 | 26.9567485925 | -0.381693219033962 | 0.714331924770386 | 0.852016252909094 | -6.33598448957844 |
| **Nampt** | -0.181500020000005 | 23.57162239 | -0.381257166600205 | 0.714640747858927 | 0.852016252909094 | -6.33616629843146 |
| **Prpsap2** | -0.0611828224999975 | 28.43691410375 | -0.381221063416981 | 0.7146663196092 | 0.852016252909094 | -6.33618134247282 |
| **Emb** | 0.0510645000000025 | 23.25015453 | 0.379946024261845 | 0.71556967834731 | 0.852462822777802 | -6.33671177131156 |
| **Uqcc2** | -0.120898205000003 | 20.85770229 | -0.37890642046402 | 0.716306596265732 | 0.852639520984014 | -6.33714299764005 |
| **Pgp** | 0.047864917499993 | 25.65174028625 | 0.373492277243571 | 0.720149640368388 | 0.856414469567864 | -6.33937047839648 |
| **Comt** | -0.0471035300000011 | 21.2355277525 | -0.373384493785557 | 0.720226235875115 | 0.856414469567864 | -6.33941451062216 |
| **Sorcs2** | 0.0426525574999985 | 28.81490361375 | 0.370429710877395 | 0.72232737689399 | 0.858466960389587 | -6.34061686586694 |
| **Dip2b** | -0.288266055000001 | 27.41965566 | -0.367611693131376 | 0.72433367512597 | 0.86036624008142 | -6.3417550293176 |
| **Ppp2r5e** | -0.0487845325000009 | 23.89387529125 | -0.366667222072534 | 0.725006618836104 | 0.86036624008142 | -6.34213462303532 |
| **Chgb** | -0.059040600000003 | 22.433859745 | -0.366602041890622 | 0.725053069950659 | 0.86036624008142 | -6.34216078510983 |
| **Fam171a2** | 0.0939588900000068 | 24.0427625675 | 0.36538687301878 | 0.725919296996448 | 0.860947808717549 | -6.34264771271343 |
| **Grid1** | -0.050781527500007 | 23.53137474875 | -0.363381210979433 | 0.727349969211643 | 0.862079536383043 | -6.34344800002882 |
| **Mark2** | -0.254147637500004 | 22.37754972125 | -0.362993361021451 | 0.727626764653578 | 0.862079536383043 | -6.34360226924143 |
| **Vps37b** | 0.048123989999997 | 22.357394295 | 0.361703032765778 | 0.728547943850145 | 0.862390363513756 | -6.3441143639808 |
| **Rab31** | 0.236561789999996 | 21.7478040425 | 0.361001092955773 | 0.729049269553582 | 0.862390363513756 | -6.34439220783549 |

| | | | | | | |
|---|---|---|---|---|---|---|
| **Abcb9** | -0.038946637500004 | 26.06049425375 | -0.36059302463334 | 0.729340777860603 | 0.862390363513756 | -6.34455349211472 |
| **Nedd4** | 0.0553421225000008 | 24.30845052875 | 0.360104278368416 | 0.729689982925059 | 0.862390363513756 | -6.34474643257808 |
| **Tpp1** | 0.258137827499997 | 21.48224081875 | 0.359579822985602 | 0.730064778799769 | 0.862390363513756 | -6.34495318989734 |
| **Lxn** | -0.0723995424999977 | 24.15665765625 | -0.359461092271943 | 0.730149639357649 | 0.862390363513756 | -6.3449999571657 |
| **Sh3gl3** | -0.128860375000006 | 23.7650142825 | -0.358163533663485 | 0.731077310613071 | 0.863040724081134 | -6.34551008889428 |
| **Celf2** | 0.0478823549999987 | 20.57303653 | 0.355959861587416 | 0.732653906546284 | 0.864456078394044 | -6.34637238977222 |
| **Ube2o** | 0.0692753299999964 | 22.59422808 | 0.354774356498947 | 0.733502641529761 | 0.865011615899857 | -6.34683416112473 |
| **Naca** | -0.243954067500002 | 21.86507111375 | -0.353300145291863 | 0.734558629776448 | 0.86581086691982 | -6.34740631877231 |
| **Rpl21** | 0.204438972500007 | 22.06746683625 | 0.352301966432903 | 0.735273984541555 | 0.866208003404848 | -6.34779242032137 |
| **Ptpn5** | 0.0721357299999923 | 20.525504205 | 0.351508962588709 | 0.735842500662114 | 0.866431833341347 | -6.34809840891496 |
| **Rps23** | 0.0393493474999964 | 29.91921241625 | 0.350193652913324 | 0.736785859046066 | 0.867096571391488 | -6.3486044697464 |
| **Fsd1** | -0.116881232500006 | 21.92041240875 | -0.349478564716539 | 0.737298935827653 | 0.86725450365339 | -6.34887883030167 |
| **Ppp1r14a** | 0.152668687500004 | 24.74684790625 | 0.347771254554508 | 0.73852451714836 | 0.868249933103542 | -6.34953169427991 |
| **Pomgnt2** | -0.0433472275000035 | 22.68192959875 | -0.347011084109956 | 0.739070464626179 | 0.86844573589801 | -6.34982138658529 |
| **Ptpn11** | -0.195441622499992 | 21.43606915125 | -0.346418208863689 | 0.739496374912429 | 0.868500360274268 | -6.3500469003002 |
| **Blvrb** | 0.0659442074999923 | 24.46310538875 | 0.345208813408832 | 0.740365487154478 | 0.869075179536719 | -6.35050576845823 |
| **Ddrgk1** | 0.0530055150000024 | 19.50055312 | 0.338369580142163 | 0.745288053961669 | 0.874296356652579 | -6.35307155749938 |
| **Eif2s1** | 0.128351722499993 | 22.69122744375 | 0.337968241157346 | 0.745577321182104 | 0.874296356652579 | -6.35322058266523 |
| **Nucb2** | 0.0817619074999953 | 19.84680777375 | 0.337209591296783 | 0.746124243012168 | 0.874489704175552 | -6.35350181769137 |
| **Timmdc1** | -0.0840084875000073 | 22.81379129625 | -0.334989640260545 | 0.747725544089084 | 0.875917999191358 | -6.35432125526654 |
| **Rpl11** | 0.0247252000000096 | 30.56915014 | 0.333342993128592 | 0.748914176831813 | 0.876861662796941 | -6.35492569258132 |

| | | | | | | |
|---|---|---|---|---|---|---|
| **Gcdh** | 0.0802548025000007 | 23.99615690625 | 0.330966393847206 | 0.750631023654088 | 0.878422501607468 | -6.35579299549985 |
| **Dhrs7b** | 0.103846502500001 | 24.02792942875 | 0.329790696872505 | 0.751480907362047 | 0.878967704114321 | -6.35621982817984 |
| **Kcnj6** | -0.177264697499993 | 22.39095604875 | -0.328929232525223 | 0.75210387596932 | 0.879147331400729 | -6.35653164669864 |
| **Mogs** | 0.196184904999999 | 22.080176675 | 0.327970779568657 | 0.75279721621011 | 0.879147331400729 | -6.35687764381615 |
| **Sh3glb1** | -0.050318502500005 | 24.85651822875 | -0.327627596725249 | 0.753045532898579 | 0.879147331400729 | -6.3570012936159 |
| **Akr1b8** | -0.0533550624999997 | 23.79965411125 | -0.327454516253435 | 0.753170780636448 | 0.879147331400729 | -6.35706360748366 |
| **Tjp1** | 0.0577504599999941 | 19.9385801725 | 0.326869975747316 | 0.753593836033425 | 0.879192808705663 | -6.35727382295852 |
| **Mif** | -0.1112064975 | 21.25110372875 | -0.323500297886924 | 0.756034382701435 | 0.881308265918932 | -6.35847855101051 |
| **Snx27** | 0.0423581875000032 | 26.31104848125 | 0.322293015108009 | 0.756909510937168 | 0.881308265918932 | -6.35890723611488 |
| **Strap** | 0.187523212499986 | 23.62161378875 | 0.322287610111888 | 0.756913429745947 | 0.881308265918932 | -6.35890915184248 |
| **Susd2** | 0.0652082450000044 | 24.544648815 | 0.322241087340607 | 0.756947160680044 | 0.881308265918932 | -6.35892563992064 |
| **Hapln1** | -0.044931717499999 | 27.75537424625 | -0.321110109939015 | 0.757767341683558 | 0.881814664521436 | -6.35932575848159 |
| **Anxa4** | 0.141838812499998 | 21.24054850875 | 0.319056850214871 | 0.759257218323547 | 0.883099478019614 | -6.36004867762586 |
| **Mboat7** | 0.0286509674999991 | 25.41791567375 | 0.318385708862165 | 0.759744448323571 | 0.883217390661582 | -6.36028400036106 |
| **H3c2** | 0.0623384900000019 | 23.3745671675 | 0.31687565526997 | 0.760841135264618 | 0.883652929363435 | -6.36081171413197 |
| **Pepd** | 0.0273297274999997 | 22.00297676625 | 0.316407431119796 | 0.761181306935719 | 0.883652929363435 | -6.36097484865478 |
| **Anln** | -0.0994392225000027 | 21.16888429125 | -0.316275415108248 | 0.761277228791915 | 0.883652929363435 | -6.36102080221026 |
| **Mrps35** | 0.228025952499998 | 22.07805377875 | 0.31124113418366 | 0.764938463194753 | 0.887452682337957 | -6.36275929873048 |
| **Eprs1** | 0.0371341675000068 | 29.72847183875 | 0.306994116599336 | 0.76803220526479 | 0.89059053589215 | -6.36420485806575 |
| **Ddb1** | -0.103724267499995 | 21.86082880875 | -0.303421039988805 | 0.770638554193753 | 0.893160329392153 | -6.36540607403804 |

| | | | | | | |
|---|---|---|---|---|---|---|
| **Cryz** | 0.1799268125 | 21.43894124125 | 0.301487993752896 | 0.772049933450298 | 0.894319447306478 | -6.36605023454391 |
| **Ewsr1** | 0.127827414999995 | 21.50681153 | 0.300981256236366 | 0.772420073099566 | 0.894319447306478 | -6.36621843509322 |
| **Pnpla6** | 0.233612205000004 | 25.8801621225 | 0.299214446955647 | 0.77371111458421 | 0.895308032035667 | -6.36680273549961 |
| **Rpl13a** | 0.0517499224999973 | 25.99950672125 | 0.298471098409595 | 0.774254524879727 | 0.895308032035667 | -6.3670475671449 |
| **Rtcb** | 0.108456210000003 | 24.10073165 | 0.298207419811606 | 0.774447314735963 | 0.895308032035667 | -6.36713427072794 |
| **Ube2k** | -0.0228265900000011 | 25.5839293375 | -0.297217806014972 | 0.775171028539243 | 0.895692319195521 | -6.36745901304367 |
| **Trio** | -0.104400147500002 | 20.57696525875 | -0.295875711380326 | 0.776152899894212 | 0.896374363197706 | -6.3678977427502 |
| **Acp1** | 0.0501456700000027 | 22.90451375 | 0.295215546761232 | 0.77663603598488 | 0.896480023383455 | -6.3681128405077 |
| **Rpl36** | -0.0496851074999931 | 26.26831815875 | -0.294643149131814 | 0.777055026849471 | 0.896511570795584 | -6.36829896283041 |
| **Tmem256** | -0.0312705125000079 | 26.78215625875 | -0.29381576524304 | 0.777660807211985 | 0.896758482472662 | -6.36856737520603 |
| **Nrn1** | -0.0356991325000067 | 28.27905474375 | -0.29242780337321 | 0.778677396803325 | 0.897478631058817 | -6.36901599356483 |
| **Nars1** | 0.0440403774999965 | 24.11946836375 | 0.289132112882629 | 0.781093139885886 | 0.898768528036293 | -6.37007293437344 |
| **Psmd4** | 0.0918749074999994 | 17.52846672625 | 0.288988340675859 | 0.781198584595265 | 0.898768528036293 | -6.37011877681963 |
| **Hsdl2** | 0.0886918625000064 | 23.89295707125 | 0.288778284740834 | 0.781352651717105 | 0.898768528036293 | -6.37018571417316 |
| **Pgm2l1** | -0.106046940000009 | 23.7484563975 | -0.288279540469208 | 0.781718501903944 | 0.898768528036293 | -6.37034445609391 |
| **Hccs** | -0.0589549350000098 | 22.252388395 | -0.288063274289256 | 0.781877160901123 | 0.898768528036293 | -6.3704132068091 |
| **Tpp2** | -0.0713972400000031 | 26.1413383875 | -0.28726437750564 | 0.782463351272519 | 0.898768528036293 | -6.3706667386818 |
| **Slc20a1** | 0.0542387400000024 | 24.9764969275 | 0.287153016006391 | 0.78254507486952 | 0.898768528036293 | -6.37070202499191 |
| **Cnnm2** | -0.0391057325000013 | 31.73803606375 | -0.285837963750746 | 0.783510361026183 | 0.899425885852022 | -6.37111770664806 |
| **Ppp1r2** | -0.0311655925000025 | 24.52724641625 | -0.283874246582249 | 0.784952551079377 | 0.900519811801275 | -6.37173496241235 |
| **Lyrm4** | -0.0427519175000057 | 25.28459743125 | -0.283469213267826 | 0.785250128595607 | 0.900519811801275 | -6.37186176004906 |

| | | | | | | |
|---|---|---|---|---|---|---|
| **Tmx1** | 0.0380558874999934 | 27.84741483625 | 0.282152321248212 | 0.786217915111524 | 0.901178171101792 | -6.37227279810088 |
| **Clpb** | 0.0547286199999952 | 20.6238742 | 0.280545120532314 | 0.787399602410472 | 0.901564689321028 | -6.37277191744692 |
| **Mrps34** | 0.0384460199999985 | 27.9395993425 | 0.28030283836711 | 0.787577791804079 | 0.901564689321028 | -6.37284691716516 |
| **Myadm** | -0.0613911824999995 | 22.69850816625 | -0.28008674540545 | 0.787736731604218 | 0.901564689321028 | -6.37291375648822 |
| **Srr** | 0.0551945474999975 | 20.91299301625 | 0.279489048573444 | 0.788176403893057 | 0.901617085712747 | -6.37309836681802 |
| **Ntrk3** | 0.0238715575000015 | 28.24307439375 | 0.275046818553788 | 0.791446756058444 | 0.904905906402486 | -6.37445836228425 |
| **Actr3b** | 0.0641529675000001 | 23.34958923625 | 0.273604927243026 | 0.792509249000997 | 0.905381304647758 | -6.37489522152533 |
| **Pacs1** | -0.0936854325000027 | 23.34938686875 | -0.273409054642684 | 0.792653619272218 | 0.905381304647758 | -6.37495439314611 |
| **Kiaa0513** | -0.0305792400000051 | 27.1675597125 | -0.270604722581107 | 0.794721545269532 | 0.906923285989301 | -6.37579701672905 |
| **Atg9a** | 0.0378646675000134 | 27.20007187875 | 0.270210260164553 | 0.795012566782836 | 0.906923285989301 | -6.37591486005462 |
| **Camk4** | -0.0396273324999967 | 26.44006216375 | -0.269966748998889 | 0.795192238960475 | 0.906923285989301 | -6.3759875236362 |
| **Klc1** | -0.0269033074999996 | 22.75931926375 | -0.269386430605719 | 0.795620474861597 | 0.90695979430189 | -6.37616043192636 |
| **Tmub1** | 0.0956505749999899 | 25.6951689475 | 0.26205512751092 | 0.801036977269148 | 0.91267976155753 | -6.37831345866603 |
| **Slc22a23** | 0.0325434049999984 | 25.1795367125 | 0.261455063668575 | 0.801480843334415 | 0.91273116934949 | -6.37848710750676 |
| **Mesd** | 0.0243733299999889 | 30.8693920675 | 0.258567713740599 | 0.803617713418275 | 0.91470957037018 | -6.37931720668196 |
| **Daam2** | 0.0358625775000121 | 25.75641639375 | 0.256955815644125 | 0.804811436723677 | 0.915613011262673 | -6.37977669021433 |
| **Cul1** | 0.0570703499999965 | 22.85669236 | 0.256308014178963 | 0.80529133746355 | 0.915703860632919 | -6.3799605573958 |
| **Pbxip1** | 0.035304005000004 | 25.195738505 | 0.255723377361687 | 0.805724522639262 | 0.91574152548226 | -6.38012610575748 |
| **Tex2** | -0.0604610874999949 | 21.12872514875 | -0.254110154884081 | 0.806920216958405 | 0.916645348197414 | -6.38058099033128 |
| **Serinc1** | -0.0381836924999917 | 28.96448445625 | -0.252641702094636 | 0.808009094736064 | 0.917165097243472 | -6.38099259921205 |
| **Rgs20** | 0.0385355300000043 | 26.3550300675 | 0.25128209216933 | 0.80901767345449 | 0.917165097243472 | -6.38137161298179 |

| | | | | | | |
|---|---|---|---|---|---|---|
| **Fn3krp** | -0.0708810349999958 | 24.065919965 | -0.251069637039247 | 0.80917531123987 | 0.917165097243472 | -6.38143065709823 |
| **Tamm41** | -0.0698810625000021 | 21.87661715375 | -0.250916079412367 | 0.809289254115102 | 0.917165097243472 | -6.3814733023049 |
| **Rnh1** | 0.109970809999997 | 21.9518708 | 0.250792212844188 | 0.80938116925811 | 0.917165097243472 | -6.38150768321561 |
| **Lypla1** | 0.0580115874999976 | 24.65044875875 | 0.2500228680909 6 | 0.809952133875419 | 0.917357958654544 | -6.38172085264069 |
| **Banf1** | -0.0276588700000069 | 28.500079015 | -0.247270438756003 | 0.811995852670423 | 0.918922192110773 | -6.38247822990918 |
| **Bles03** | 0.0453417025000036 | 24.52705848625 | 0.246961140376832 | 0.81222561037526 | 0.918922192110773 | -6.38256282419356 |
| **Pnpt1** | -0.0650396174999983 | 23.95997223625 | -0.246541214036904 | 0.812537578345219 | 0.918922192110773 | -6.38267750927451 |
| **Dazap1** | 0.0483261350000035 | 27.6967442025 | 0.244918469628237 | 0.813743477909477 | 0.919831516511008 | -6.38311889189605 |
| **Pnpla8** | 0.0368309149999959 | 26.526236575 | 0.242823970048884 | 0.81530075513421 | 0.921136934107704 | -6.38368435870762 |
| **Pip4k2c** | 0.0633569525000013 | 23.47610931625 | 0.242130264110931 | 0.815816730015123 | 0.921265167737848 | -6.38387059168065 |
| **Serpinh1** | 0.0722490825000008 | 21.40568491375 | 0.239744946686515 | 0.817591667312283 | 0.922606094438694 | -6.38450696131298 |
| **Aldh9a1** | 0.0337934725000082 | 27.74534297875 | 0.239369239381517 | 0.817871340224339 | 0.922606094438694 | -6.38460663041447 |
| **Iqsec1** | 0.0198718399999969 | 30.7058450075 | 0.238400297081049 | 0.818592743509839 | 0.922606094438694 | -6.38486296613858 |
| **Ppm1l** | -0.0910509475000012 | 20.66843092125 | -0.238080950919195 | 0.818830546784189 | 0.922606094438694 | -6.38494722590796 |
| **Mgst3** | 0.050506572499998 | 22.59718768875 | 0.237827255492016 | 0.819019477457154 | 0.922606094438694 | -6.38501408453476 |
| **Gmps** | 0.0206070700000076 | 26.2589802925 | 0.236559507796007 | 0.819963781654394 | 0.923215492477574 | -6.38534713533328 |
| **Pip4p1** | 0.119295357500008 | 22.41834689375 | 0.234087301253301 | 0.8218061736692 | 0.924271571975325 | -6.38599157507797 |
| **Spcs2** | -0.128452065000001 | 20.7804365025 | -0.233945567490158 | 0.821911836509607 | 0.924271571975325 | -6.3860283195568 |
| **Rps20** | -0.0271799350000066 | 29.25911343 | -0.233675591651783 | 0.822113115134015 | 0.924271571975325 | -6.38609825023377 |
| **Lamtor1** | -0.189462784999993 | 23.227223605 | -0.2320653718779 | 0.823313902945422 | 0.924946483160671 | -6.38651368874615 |

| | | | | | | |
|---|---|---|---|---|---|---|
| **Rraga** | -0.0195653700000022 | 26.75470565 | -0.231373732505887 | 0.82382983557577 | 0.924946483160671 | -6.38669126477689 |
| **Clvs1** | 0.0383028900000006 | 28.4880339 | 0.231245262432391 | 0.823925678970995 | 0.924946483160671 | -6.38672419159358 |
| **Cdc42bpb** | 0.109795275 | 23.2859205375 | 0.227275478025357 | 0.826888872968531 | 0.92772145830084 | -6.38773277387387 |
| **Adpgk** | -0.044316500000015 | 25.523227095 | -0.226847963758692 | 0.827208167930107 | 0.92772145830084 | -6.38784036475353 |
| **Prpf19** | 0.016295842500007 | 27.95180820875 | 0.224377897107717 | 0.829053659814127 | 0.928789952890979 | -6.38845809112236 |
| **Cnr1** | 0.0297625075000063 | 26.60238969625 | 0.224236751711496 | 0.829159150843247 | 0.928789952890979 | -6.38849318828123 |
| **Dmac2l** | -0.0219226574999993 | 23.06045388125 | -0.223943704599663 | 0.829378184232923 | 0.928789952890979 | -6.38856598779604 |
| **Ablim2** | 0.0661825024999985 | 21.03166139125 | 0.219763922573858 | 0.832504067671371 | 0.931834626356855 | -6.38959413015819 |
| **Kctd8** | 0.0468294524999955 | 23.78359966375 | 0.219124111598409 | 0.832982845628108 | 0.93191482582734 | -6.38974982621678 |
| **Alad** | -0.0892621700000014 | 22.9488825 | -0.217356818031453 | 0.834305728498311 | 0.932651947985802 | -6.39017756674092 |
| **Cmtm5** | 0.0214944924999969 | 30.66787744875 | 0.217155293113544 | 0.834456614012635 | 0.932651947985802 | -6.39022612521317 |
| **Wdr48** | 0.0226963900000001 | 24.74028868 | 0.214385727089756 | 0.836530999648027 | 0.934514132842526 | -6.39088896706924 |
| **Tbc1d17** | 0.105408919999999 | 22.97137427 | 0.210370977842842 | 0.839540506609258 | 0.937418643721264 | -6.39183491925502 |
| **Scg3** | 0.0414630599999946 | 25.951221655 | 0.20890366784517 | 0.84064114666379 | 0.938189948665731 | -6.39217624302024 |
| **Cdv3** | 0.0556019725000034 | 24.74398676625 | 0.20319139548492 | 0.844929603823585 | 0.942516502510812 | -6.39348256141835 |
| **Dgkb** | -0.104782379999996 | 20.0779478 | -0.201277231544157 | 0.846367929621691 | 0.943661076913809 | -6.39391230417219 |
| **Pef1** | 0.181672995 | 20.878816895 | 0.198360305100518 | 0.84856095665864 | 0.945069403561647 | -6.39455944608382 |
| **H1-5** | 0.0587438924999901 | 22.73461768625 | 0.197903594523024 | 0.848904456045532 | 0.945069403561647 | -6.39465992552762 |
| **Grik3** | 0.0346892599999968 | 25.6121835775 | 0.197688883167643 | 0.849065956157959 | 0.945069403561647 | -6.3947070843926 |
| **Mpp1** | 0.0462613725000054 | 24.62217013125 | 0.197400938464369 | 0.849282552698256 | 0.945069403561647 | -6.39477024869652 |
| **Scfd1** | 0.0140571850000057 | 27.7567518875 | 0.194769994891623 | 0.851262237722691 | 0.946325174972381 | -6.39534316480748 |

| | | | | | | |
|---|---|---|---|---|---|---|
| **Tmcc1** | -0.0370521325000013 | 20.90327768625 | -0.194270151639983 | 0.851638482060459 | 0.946325174972381 | -6.39545115201345 |
| **Trappc2** | 0.114411172499995 | 20.67821946125 | 0.193714673171237 | 0.852056653287066 | 0.946325174972381 | -6.39557083691825 |
| **Arpc5** | 0.0393782025000036 | 22.17488961625 | 0.193703934306318 | 0.852064738146823 | 0.946325174972381 | -6.39557314740468 |
| **Cops5** | -0.145603012500001 | 21.04044987375 | -0.192546241576218 | 0.852936430772646 | 0.946506771983652 | -6.39582148431298 |
| **Ado** | 1.2866421025 | 15.05085299125 | 0.192388458957593 | 0.853055251464515 | 0.946506771983652 | -6.39585521630945 |
| **Ptgds** | -0.140761344999991 | 22.7831123775 | -0.191334289201426 | 0.853849216070042 | 0.946900105731106 | -6.39607988320441 |
| **Scn2b** | 1.46841508 | 17.139767555 | 0.190819374834871 | 0.854237098442435 | 0.946900105731106 | -6.39618917904044 |
| **Prune2** | 0.0183152050000004 | 26.0305936325 | 0.187339326319487 | 0.856859735746555 | 0.949347500059953 | -6.39692021590733 |
| **Scoc** | -0.203185415000007 | 24.3014528825 | -0.184944872163006 | 0.858665382035715 | 0.950887788814587 | -6.3974154755763 |
| **Eif3j2** | -0.118528105000001 | 23.02704529 | -0.18322834426166 | 0.859960369808939 | 0.951861357104768 | -6.39776663518246 |
| **G6pdx** | 0.0514417700000038 | 25.2484125075 | 0.182124518671912 | 0.860793366888313 | 0.951990277004962 | -6.39799073846567 |
| **Lhfpl4** | 0.143396799999998 | 22.9663287125 | 0.181971789387503 | 0.860908638444854 | 0.951990277004962 | -6.39802164062306 |
| **Lamtor5** | -0.0644162849999993 | 23.133544415 | -0.179913733426852 | 0.8624622999113 | 0.95249023077502 | -6.39843554945596 |
| **Kctd16** | 0.0577668325000005 | 23.26499987125 | 0.179341603064397 | 0.862894327848685 | 0.95249023077502 | -6.39854978612111 |
| **Sri** | -0.105821235000008 | 23.330384295 | -0.179198509611908 | 0.86300238870895 | 0.95249023077502 | -6.3985783011083 |
| **Stx5** | -0.0181375025000037 | 24.72367934375 | -0.17916827129616 | 0.863025224389424 | 0.95249023077502 | -6.39858432397203 |
| **Rbm39** | 0.0198012824999978 | 26.46539188625 | 0.178237101575948 | 0.86372850369252 | 0.952807009615507 | -6.39876930143665 |
| **Lman2l** | -0.062874067500001 | 22.82165918125 | -0.177654192187025 | 0.864168822248454 | 0.952833542450246 | -6.39888461089523 |
| **Ipo5** | 0.0778126274999948 | 24.31934548125 | 0.176701735879927 | 0.864888401408538 | 0.953167814551826 | -6.39907221790241 |
| **Arpp21** | 0.0728290249999972 | 24.28822131 | 0.174942922578549 | 0.86621754487304 | 0.953737758961415 | -6.39941602848121 |

| | | | | | | |
|---|---|---|---|---|---|---|
| **Chordc1** | 0.102381685000005 | 21.671903545 | 0.174906444043344 | 0.866245116832855 | 0.953737758961415 | -6.39942312318944 |
| **Abcb10** | -0.0397018424999978 | 22.60197283375 | -0.173840242119324 | 0.867051084162567 | 0.953737758961415 | -6.39962984112211 |
| **Copg1** | -1.19691426 | 15.523076475 | -0.173812305213864 | 0.867072204630277 | 0.953737758961415 | -6.39963524076155 |
| **Gart** | 0.0262552050000053 | 19.335352275 | 0.172442343094671 | 0.868108047519657 | 0.953904425798944 | -6.39989897165583 |
| **Cox15** | 0.0615732399999942 | 23.049949535 | 0.172428066686132 | 0.868118843538779 | 0.953904425798944 | -6.39990170911323 |
| **Pofut2** | 0.120928149999997 | 25.86970623 | 0.171958532215536 | 0.868473928949322 | 0.953904425798944 | -6.39999161572484 |
| **Trnt1** | 0.0721109450000021 | 18.828366665 | 0.169575437023889 | 0.870276647563536 | 0.955225149777898 | -6.4004441849601 |
| **Usp12** | 0.0335961100000048 | 21.56079854 | 0.169265726615142 | 0.870510992763956 | 0.955225149777898 | -6.40050254178765 |
| **Mia3** | -0.128762497500002 | 25.75490727625 | -0.16710530743459 | 0.872146082589947 | 0.956197519256008 | -6.40090667398929 |
| **Sec11c** | -0.0180509750000013 | 25.4338077925 | -0.166175855317679 | 0.872849736329421 | 0.956197519256008 | -6.40107895541169 |
| **Rpl23a** | 0.0144670474999984 | 20.73380228625 | 0.165892957855952 | 0.873063932187901 | 0.956197519256008 | -6.40113120356846 |
| **Pls1** | -0.0162681274999983 | 31.74320330625 | -0.1658874912836 | 0.873068071317194 | 0.956197519256008 | -6.40113221231684 |
| **Nmral1** | 0.014372242499995 | 27.11586460875 | 0.16422810449575 | 0.874324708212536 | 0.956849067501091 | -6.40143689555479 |
| **Ostc** | 0.0359752525000019 | 25.04982093625 | 0.163996215943282 | 0.87450034632614 | 0.956849067501091 | -6.40147923111576 |
| **Akap7** | 0.0272257324999998 | 24.38070890375 | 0.163446114483394 | 0.874917037256349 | 0.956849067501091 | -6.40157942500463 |
| **Islr2** | -0.0316051749999957 | 26.346586405 | -0.162397996756826 | 0.875711083983184 | 0.95726011042861 | -6.40176940180989 |
| **Tor1aip1** | 0.0274045200000081 | 25.7903906725 | 0.159357735235676 | 0.878015235338296 | 0.959320703431676 | -6.40231360482728 |
| **Dner** | 0.0323032375000025 | 24.27388791625 | 0.157615572984711 | 0.879336160705588 | 0.959835477320529 | -6.40262084991988 |
| **Stmn3** | 0.0336311675000047 | 22.14655172625 | 0.157389691205136 | 0.879507457053732 | 0.959835477320529 | -6.40266044060557 |
| **Lamtor2** | 0.0768169100000016 | 19.6246796925 | 0.15707731666244 | 0.879744356233495 | 0.959835477320529 | -6.40271509816837 |
| **Mrpl49** | -0.0174471949999999 | 19.6972315825 | -0.155002963220903 | 0.881317844020148 | 0.960542243889696 | -6.40307532243319 |

| | | | | | | |
|---|---|---|---|---|---|---|
| **Gng13** | -1.3176450525 | 10.98107987125 | -0.154847157235835 | 0.88143605311765 | 0.960542243889696 | -6.403102187158 |
| **Dctn3** | -0.0324170475000045 | 20.19064080375 | -0.154563790170541 | 0.88165105042038 | 0.960542243889696 | -6.4031509777488 |
| **Cystm1** | -0.0164738524999919 | 28.83598345375 | -0.153572301418375 | 0.882403401007 | 0.960821878245475 | -6.4033209954842 |
| **Gbf1** | -0.0851098575000009 | 22.88878193625 | -0.153119242002131 | 0.882747230209801 | 0.960821878245475 | -6.40339832322803 |
| **Vac14** | 0.0359941974999956 | 24.31097105875 | 0.152366683738693 | 0.88331841090402 | 0.96098661718598 | -6.40352626776733 |
| **Fkbp4** | 0.0344413175000042 | 24.41073554375 | 0.151028383365801 | 0.88433434601158 | 0.961634830413542 | -6.40375224948099 |
| **Tmem222** | -0.0294091174999949 | 25.80641284125 | -0.149677591846738 | 0.885360001501127 | 0.962292993084558 | -6.40397833257488 |
| **Bcap29** | -0.0351495800000023 | 26.1606511675 | -0.147353062981015 | 0.887125569310724 | 0.963754356028593 | -6.40436266739437 |
| **Galnt16** | 0.904925882499997 | 14.15631699125 | 0.144085628300712 | 0.889608476700874 | 0.96550795329598 | -6.40489279605712 |
| **Hdac11** | 0.0446769549999964 | 21.9065300325 | 0.142958102947029 | 0.890465589412862 | 0.96550795329598 | -6.40507299125747 |
| **Hmga1** | 0.0897443575000061 | 22.20837040875 | 0.142565633169068 | 0.890763970911479 | 0.96550795329598 | -6.40513538363068 |
| **Dcakd** | 0.0827987775000025 | 22.04638979875 | 0.142454120428713 | 0.890848753761952 | 0.96550795329598 | -6.40515308012393 |
| **Dpm1** | 0.0973539449999947 | 21.852170265 | 0.139189196450613 | 0.893331744909295 | 0.967740825412862 | -6.40566510535947 |
| **Sec11a** | -0.0119022274999949 | 27.33250650375 | -0.135341608133228 | 0.8962595100216 | 0.970453178069746 | -6.40625335689647 |
| **Pafah1b3** | -1.021580415 | 17.0312057375 | -0.133426540887625 | 0.897717405233525 | 0.971183274827103 | -6.40654003633634 |
| **Tra2b** | -0.0150037050000122 | 23.09064588 | -0.133341237631368 | 0.897782354536544 | 0.971183274827103 | -6.40655271143248 |
| **Cryab** | -0.0250524300000023 | 20.1715157875 | -0.132454228615955 | 0.898457766988419 | 0.971454808047468 | -6.40668403308235 |
| **Psmd11** | 0.0549108099999991 | 25.69683516 | 0.13048113140513 | 0.899960505341193 | 0.972620206197352 | -6.40697302311386 |
| **Hook3** | -0.133966467499995 | 23.55782191625 | -0.129211163911096 | 0.900927964884825 | 0.973206282029903 | -6.40715674664604 |
| **Abhd17c** | -0.0321856975000046 | 24.08674494125 | -0.127044432129497 | 0.902578996693725 | 0.974529869543366 | -6.40746607558556 |

| | | | | | | |
|---|---|---|---|---|---|---|
| **Huwe1** | -0.0103075099999934 | 21.4606281075 | -0.126149124594432 | 0.90326136603967 | 0.974806820775485 | -6.40759237242593 |
| **Pfas** | 0.00971287500000173 | 27.8404180475 | 0.123885293062325 | 0.904987164382997 | 0.976209057150179 | -6.40790775479123 |
| **Lrrc4b** | 0.0221842650000007 | 24.09646009 | 0.121922899821597 | 0.906483614341314 | 0.976642483754178 | -6.40817654297329 |
| **Dnaja4** | 0.864458817499999 | 15.96375895875 | 0.121864156688775 | 0.906528416075203 | 0.976642483754178 | -6.40818452313626 |
| **Lgmn** | 0.0200413824999997 | 23.22706431125 | 0.121679865095549 | 0.90666897246729 | 0.976642483754178 | -6.40820953401476 |
| **Dpp7** | 0.0355277525000002 | 25.95388845375 | 0.117144603593335 | 0.910129070846147 | 0.979587935273021 | -6.40881315451241 |
| **Vkorc1** | -0.0315838074999988 | 23.53827067375 | -0.116973962234835 | 0.910259300273357 | 0.979587935273021 | -6.40883542038385 |
| **Slc39a11** | 0.0652975550000008 | 23.7244970625 | 0.11271004389642 | 0.913514376454914 | 0.982355021020338 | -6.40938129450231 |
| **Mrpl23** | 0.0601420124999947 | 20.61863088375 | 0.112481527448998 | 0.913688877130756 | 0.982355021020338 | -6.4094099795893 |
| **Gpc4** | -0.0333679774999993 | 23.49090630625 | -0.110653352456896 | 0.915085099705988 | 0.982943096016245 | -6.40963737768619 |
| **Anp32b** | 0.0109259450000039 | 23.419380265 | 0.110618780228818 | 0.915111506506355 | 0.982943096016245 | -6.40964164219759 |
| **Abhd17b** | -0.0168671399999987 | 22.4544038125 | -0.109467200857496 | 0.915991166421866 | 0.982943096016245 | -6.40978293172856 |
| **Efnb2** | -0.0763032425000141 | 25.81425435125 | -0.109404293540872 | 0.91603922326625 | 0.982943096016245 | -6.40979060751404 |
| **Tars1** | -0.114888724999989 | 21.14987518 | -0.108954366423922 | 0.916382947531091 | 0.982943096016245 | -6.40984537826782 |
| **Pm20d1** | -0.00852247499999237 | 29.7107574825 | -0.107778052570268 | 0.917281689851825 | 0.983280283176221 | -6.40998751121768 |
| **Suclg2** | 0.0275903950000007 | 23.0361602975 | 0.107418485141662 | 0.917556437249632 | 0.983280283176221 | -6.41003065062478 |
| **Clcn4** | 0.739591194999999 | 15.957359875 | 0.104393812433796 | 0.919868082993138 | 0.985293166655718 | -6.41038785074076 |
| **Tmcc2** | 0.0259731974999973 | 21.61184484875 | 0.103713288094101 | 0.920388298695191 | 0.985293166655718 | -6.41046681638449 |
| **Cd44** | -0.0175093674999971 | 23.72435208875 | -0.103271396898831 | 0.920726117726772 | 0.985293166655718 | -6.41051781622335 |
| **C2orf88(AKAP)** | -0.684373387500001 | 15.41084799625 | -0.10021944223308 | 0.92305976825104 | 0.98732888295637 | -6.41086412339884 |
| **Sec61a1** | -0.0164149299999963 | 25.4582961825 | -0.099442155598132 | 0.923654245313399 | 0.987503301038006 | -6.4109506679408 |

| | | | | | | |
|---|---|---|---|---|---|---|
| **Tbrg4** | 0.0105332950000054 | 28.8391136775 | 0.0964856382996127 | 0.925915898457515 | 0.987839700677734 | -6.41127371399508 |
| **Lrtm2** | -0.00757268250000465 | 25.15996146125 | -0.0962209943697405 | 0.926118380042878 | 0.987839700677734 | -6.41130215645708 |
| **Spg7** | -0.0178364900000041 | 27.2547053875 | -0.0960959534292421 | 0.926214052110093 | 0.987839700677734 | -6.41131556806447 |
| **Rilp** | -0.6767934125 | 15.91856041125 | -0.0959097663555474 | 0.926356511109635 | 0.987839700677734 | -6.41133550583948 |
| **Slc22a17** | 0.117684372500001 | 23.73177981125 | 0.0958052423108476 | 0.926436487814749 | 0.987839700677734 | -6.41134668185881 |
| **Penk** | -0.0174374874999934 | 22.60915382625 | -0.0956461080507506 | 0.926558251356529 | 0.987839700677734 | -6.41136367362795 |
| **Slc9a7** | -0.0326222175000019 | 24.43212021375 | -0.0945383239867131 | 0.927405944631804 | 0.988283150494507 | -6.41148117811279 |
| **Gipc1** | -0.0117747250000022 | 22.5052204675 | -0.0935061090858871 | 0.928195902785211 | 0.988406549839808 | -6.41158943816051 |
| **Atp8a2** | 0.0212133724999966 | 22.33673104375 | 0.0930537381658611 | 0.928542131627104 | 0.988406549839808 | -6.41163650982739 |
| **Trappc4** | 0.0138125349999996 | 25.2495788625 | 0.0922695004246636 | 0.929142399214151 | 0.988406549839808 | -6.41171757454185 |
| **Arl6ip1** | 0.0615625249999958 | 23.895128025 | 0.0921302729402211 | 0.929248971277967 | 0.988406549839808 | -6.41173189461354 |
| **Aspa** | -0.0150426924999962 | 27.24341153875 | -0.090098240625764 | 0.930804573061485 | 0.988708442992103 | -6.41193844201317 |
| **Gfm1** | -0.621477667500001 | 15.60349099125 | -0.089923460280967 | 0.930938389730779 | 0.988708442992103 | -6.41195599307464 |
| **Cntn4** | -0.0155224400000016 | 19.6520908875 | -0.089713705251054 | 0.931098987167583 | 0.988708442992103 | -6.41197701133536 |
| **Trappc3** | -0.0303416500000004 | 23.2667740775 | -0.0885532004372197 | 0.931987581720331 | 0.988708442992103 | -6.41209241347796 |
| **Mobp** | -0.591964302499996 | 15.07689536125 | -0.0885398516141995 | 0.931997803479568 | 0.988708442992103 | -6.41209373218264 |
| **Bcl2l1** | 0.6534908575 | 16.66361611375 | 0.088374488886496 | 0.932124429871978 | 0.988708442992103 | -6.41211005160365 |
| **Exoc7** | 0.015650265000005 | 22.44269919 | 0.0872386991096294 | 0.932994217177854 | 0.988994211769476 | -6.41222131867259 |
| **Rpl36a** | -0.0267888025000005 | 22.60257097375 | -0.0861482325497053 | 0.933829388915994 | 0.988994211769476 | -6.41232679451265 |
| **Sumo3** | 0.0674760699999979 | 22.694700205 | 0.0860695821828561 | 0.933889629523952 | 0.988994211769476 | -6.41233435082214 |

| | | | | | | |
|---|---|---|---|---|---|---|
| **Bckdk** | -0.00895303749999954 | 21.93899127625 | -0.0856655938461336 | 0.934199063318396 | 0.988994211769476 | -6.41237305534434 |
| **Eif4g1** | 0.58434901 | 15.542857595 | 0.0848500278786248 | 0.934823781236728 | 0.988994211769476 | -6.41245063783416 |
| **Extl2** | 0.0112586174999905 | 26.18583028125 | 0.0841119507044988 | 0.935389185985305 | 0.988994211769476 | -6.41252021069123 |
| **Tmsb4x** | -0.636281649999999 | 9.9068530825 | -0.0836334340447397 | 0.935755775658435 | 0.988994211769476 | -6.4125649926853 |
| **Xxylt1** | -0.009938274999989 | 26.53339668 | -0.0821907905123217 | 0.936861080372139 | 0.988994211769476 | -6.41269845948583 |
| **Hexa** | -0.0630577149999958 | 20.96475125 | -0.0819231077143094 | 0.937066186481291 | 0.988994211769476 | -6.41272296937419 |
| **Anxa3** | -0.0455597149999996 | 22.550103115 | -0.0816601511166677 | 0.937267676260907 | 0.988994211769476 | -6.41274696882378 |
| **Srsf10** | -0.0284859524999952 | 23.13438748375 | -0.0814340292661266 | 0.937440945480843 | 0.988994211769476 | -6.41276754487178 |
| **Gatc** | -0.0202769874999973 | 22.12087198125 | -0.0809803869382562 | 0.937788566654003 | 0.988994211769476 | -6.4128086525296 |
| **Tmem94** | 0.0355860800000052 | 23.16455964 | 0.0805608089457082 | 0.938110097726776 | 0.988994211769476 | -6.41284646936744 |
| **Ptdss1** | -0.0152360250000001 | 23.4009261125 | -0.0796366564544485 | 0.938818338243434 | 0.988994211769476 | -6.41292907231784 |
| **Cfap418** | 0.0120256474999962 | 26.85917969375 | 0.0794716627978634 | 0.93894479033831 | 0.988994211769476 | -6.4129437197711 |
| **Cpne1** | 0.0354989549999942 | 20.88358875 | 0.0786370692834815 | 0.939584456679082 | 0.988994211769476 | -6.41301734692703 |
| **Scn3b** | -0.0291804949999914 | 24.24561156 | -0.0780918074120824 | 0.940002393521869 | 0.988994211769476 | -6.41306503046049 |
| **Plpp6** | 0.545848605000003 | 15.822397825 | 0.0778718503702465 | 0.940170993800952 | 0.988994211769476 | -6.41308417211537 |
| **Trim3** | 0.00921422000000405 | 25.3967502075 | 0.0746086105680097 | 0.942672698420962 | 0.991038540548411 | -6.41336182282936 |
| **Mrps5** | -0.0168315349999979 | 22.61076077 | -0.0737753069637726 | 0.94331164888141 | 0.991038540548411 | -6.4134308223441 |
| **Csnk1g1** | -0.00811800000001028 | 28.5946452575 | -0.0736427778593949 | 0.943413272107902 | 0.991038540548411 | -6.41344172475782 |
| **Crot** | -0.0050071349999925 | 25.8007595925 | -0.0721252524440129 | 0.944576990082368 | 0.991805839586486 | -6.41356516784364 |
| **Atxn10** | -0.018175179999993 | 23.3652454575 | -0.0712198830822067 | 0.945271344459621 | 0.99184254841994 | -6.41363759288529 |
| **Fbxl20** | -0.0309162150000013 | 22.53415982 | -0.0709496979586457 | 0.945478567344827 | 0.99184254841994 | -6.41365902937269 |

| | | | | | | |
|---|---|---|---|---|---|---|
| **Rpl34** | -0.00993731750000393 | 29.22829267875 | -0.0667297769137034 | 0.948715684590731 | 0.994323454464234 | -6.41398327886885 |
| **Nedd8** | -0.421070437499999 | 14.38139269625 | -0.0661074315636172 | 0.949193178045102 | 0.994323454464234 | -6.41402941898675 |
| **Pam** | -0.0162621524999977 | 25.25849309125 | -0.0659659956081161 | 0.949301697699145 | 0.994323454464234 | -6.4140398447079 |
| **Glrx3** | -0.452400890000003 | 15.56458752 | -0.0656018894745681 | 0.949581070973707 | 0.994323454464234 | -6.41406658159837 |
| **Myo18a** | 0.0273434200000011 | 22.940493995 | 0.0644475629611857 | 0.950466819074384 | 0.99479586139061 | -6.41415036860799 |
| **Plcxd3** | -0.00688524749999431 | 29.50838088125 | -0.0634064325894124 | 0.951265773331547 | 0.995177036177291 | -6.41422466507889 |
| **Mlf2** | -0.534578587500002 | 11.29790454375 | -0.0616710495454224 | 0.952597624856199 | 0.996115104292297 | -6.4143458182286 |
| **Mapk8ip3** | 0.0133088074999996 | 23.79829048125 | 0.0589560816895061 | 0.954681602783468 | 0.99783844236135 | -6.41452862309413 |
| **Mydgf** | -0.0106387500000054 | 23.79324506 | -0.0579718154662883 | 0.955437209530672 | 0.99792395841252 | -6.41459286571818 |
| **Tesc** | 0.428555482499998 | 16.94849080125 | 0.0569971353144234 | 0.956185505904667 | 0.99792395841252 | -6.4146554179311 |
| **Myorg** | 0.369272379999996 | 14.9336582075 | 0.0558380711697539 | 0.957075422353189 | 0.99792395841252 | -6.41472842418072 |
| **Tcerg1** | 0.0141281175000039 | 23.26237053375 | 0.0558254010961211 | 0.957085150660952 | 0.99792395841252 | -6.41472921395418 |
| **Has1** | 0.0166423599999916 | 24.065327425 | 0.0555406044943575 | 0.957303824632694 | 0.99792395841252 | -6.41474691915264 |
| **Astn2** | -0.400225379999998 | 9.606239375 | -0.0543161942334818 | 0.958244002452448 | 0.99792395841252 | -6.414822007526 |
| **Nf2** | -0.365919267499997 | 15.19436557625 | -0.0539449212479231 | 0.958529103020077 | 0.99792395841252 | -6.41484444584745 |
| **Alg5** | -0.0153560624999969 | 23.74902294125 | -0.0533446103848367 | 0.958990095630888 | 0.99792395841252 | -6.41488040102968 |
| **Adss2** | 0.0149572349999936 | 24.347013335 | 0.0532372935081475 | 0.959072508534074 | 0.99792395841252 | -6.41488678633877 |
| **Reep5** | 0.00819714000000005 | 24.348367435 | 0.0531714527282741 | 0.959123070558123 | 0.99792395841252 | -6.41489069747841 |
| **Mthfd1** | -0.0130556474999999 | 24.78461829125 | -0.0514319366976055 | 0.960458993994215 | 0.998421528814378 | -6.41499227822735 |
| **Smdt1** | -0.373687460000006 | 16.4133104775 | -0.0514128417157436 | 0.960473659436112 | 0.998421528814378 | -6.41499337456603 |
| **Ttn** | -0.0139961025000055 | 23.98763767625 | -0.0484515423215069 | 0.962748210257763 | 0.998963492980642 | -6.41515847413331 |

| | | | | | | |
|---|---|---|---|---|---|---|
| **Plgrkt** | -0.0462712224999997 | 23.17167191875 | -0.0480885695020525 | 0.963027033036451 | 0.998963492980642 | -6.41517803759589 |
| **Get4** | 0.348459267500001 | 9.65130008625 | 0.0470455232637845 | 0.9638282948815 | 0.998963492980642 | -6.41523343736578 |
| **Ccdc110** | -0.3099583 | 14.911614795 | -0.0469329662962872 | 0.963914763179606 | 0.998963492980642 | -6.41523934308315 |
| **Selenbp1** | -0.341138815000001 | 16.47127523 | -0.0467684604325068 | 0.964041140501177 | 0.998963492980642 | -6.41524794906157 |
| **Rapgef4** | 0.340387130000003 | 17.0003279775 | 0.0451574565648702 | 0.965278810345791 | 0.998963492980642 | -6.41533063136645 |
| **Pnkd** | -0.0103184025000083 | 26.14072581125 | -0.0428575923847541 | 0.967045882024486 | 0.998963492980642 | -6.4154436496284 |
| **Pacsin2** | 0.303304580000001 | 9.3739427925 | 0.0421840986281683 | 0.967563390757448 | 0.998963492980642 | -6.41547562852441 |
| **Ttc19** | -0.00417043000000206 | 22.2082235775 | -0.0420783613189929 | 0.967644640236447 | 0.998963492980642 | -6.41548060317475 |
| **Arfgef1** | 0.295219539999998 | 9.2257249 | 0.0417192872298989 | 0.967920559014772 | 0.998963492980642 | -6.41549740348685 |
| **Abcg5** | -0.294092532500008 | 16.11321262125 | -0.0412060975087463 | 0.968314911144375 | 0.998963492980642 | -6.41552116473631 |
| **Phactr4** | 0.311199192499998 | 17.09373697375 | 0.0410730483770356 | 0.968417152108731 | 0.998963492980642 | -6.41552727707486 |
| **U2af2** | -0.346993557499999 | 11.02154556625 | -0.0410438823379205 | 0.968439564696686 | 0.998963492980642 | -6.41552861433587 |
| **Gcn1** | -0.289609265000001 | 15.93862536 | -0.0410345540202206 | 0.968446733031389 | 0.998963492980642 | -6.41552904183824 |
| **Sco2** | 0.0127917049999908 | 23.795567805 | 0.0386267769677738 | 0.970297089636414 | 0.998963492980642 | -6.4156361385801 |
| **Dnajb14** | 0.00815767499999964 | 23.2449205875 | 0.0385584179253243 | 0.970349625930199 | 0.998963492980642 | -6.41563908468944 |
| **Amigo1** | 0.3214993925 | 11.00288637875 | 0.0380956251740228 | 0.970705302364736 | 0.998963492980642 | -6.41565889274445 |
| **Abhd10** | 0.00519609750000427 | 19.09958334375 | 0.037845579552507 | 0.970897476339515 | 0.998963492980642 | -6.41566949549977 |
| **Akap10** | 0.249066290000002 | 15.3662821825 | 0.0365938186786425 | 0.971859555306254 | 0.998963492980642 | -6.41572152502979 |
| **Sycp1** | 0.0038407149999955 | 24.463616565 | 0.0360634650793134 | 0.972267190070118 | 0.998963492980642 | -6.41574304168048 |
| **Hira** | 0.219898502500001 | 13.79878343125 | 0.0359654443740743 | 0.972342530688822 | 0.998963492980642 | -6.41574698403926 |
| **Itga3** | 0.00861618749999948 | 24.87407034375 | 0.0358847628757106 | 0.972404544284774 | 0.998963492980642 | -6.41575022097357 |

| | | | | | | |
|---|---|---|---|---|---|---|
| **Os9** | 0.219736964999997 | 13.858054005 | 0.0357990801945065 | 0.972470402126636 | 0.998963492980642 | -6.41575365059848 |
| **Psen1** | 0.00625383499999899 | 26.5686195775 | 0.0343544209284683 | 0.973580837644711 | 0.998963492980642 | -6.41581024201886 |
| **Kif1b** | -0.0236399749999947 | 21.46208368 | -0.0340591424087203 | 0.973807810971645 | 0.998963492980642 | -6.41582152214262 |
| **Ddi2** | -0.229282634999997 | 15.296445175 | -0.0338294421517328 | 0.973984377714295 | 0.998963492980642 | -6.41583022975735 |
| **Smc4** | -0.253560027499997 | 17.10242468375 | -0.0334553866911538 | 0.974271911248333 | 0.998963492980642 | -6.41584428361197 |
| **Vps29** | -0.232372597500003 | 16.13708794125 | -0.0323197867879829 | 0.975144863183616 | 0.998963492980642 | -6.4158859929143 |
| **Rabac1** | 0.230382262499999 | 9.29982735375 | 0.0323004461053661 | 0.975159730965669 | 0.998963492980642 | -6.41588669080656 |
| **Rpap1** | 0.252527095000001 | 10.58388128 | 0.0311085381090719 | 0.976076007884328 | 0.998963492980642 | -6.41592889388293 |
| **Dchs1** | 0.00787269999999651 | 23.680444205 | 0.0308933122142448 | 0.976241466562571 | 0.998963492980642 | -6.41593634553667 |
| **Psmd5** | -0.00482957250000737 | 24.14944457125 | -0.0307079042607394 | 0.976384003187835 | 0.998963492980642 | -6.41594272335219 |
| **Tgfbrap1** | 0.00429715250000484 | 25.75915065125 | 0.0302319177858077 | 0.976749932977739 | 0.998963492980642 | -6.41595892098026 |
| **Scn4b** | -0.0059953200000038 | 19.390612305 | -0.0300396726564244 | 0.97689772927239 | 0.998963492980642 | -6.41596539128827 |
| **Ddx42** | -0.00582601750000578 | 24.47281470125 | -0.0298459207585037 | 0.977046684947286 | 0.998963492980642 | -6.41597187055772 |
| **Srrm3** | 0.204736774999999 | 9.11483301 | 0.0292882931322352 | 0.97747539229988 | 0.998963492980642 | -6.41599028430961 |
| **Lamtor3** | -0.198862915000007 | 16.0553782025 | -0.0279478250895108 | 0.978505984464599 | 0.998963492980642 | -6.41603312825014 |
| **Nipal3** | -0.00288176750001057 | 27.46191345875 | -0.0265514676668454 | 0.979579593123969 | 0.998963492980642 | -6.41607562495349 |
| **Arhgef2** | -0.187880780000008 | 16.01053286 | -0.0264963789138947 | 0.979621949823124 | 0.998963492980642 | -6.41607725688176 |
| **Clstn3** | 0.177882405000002 | 15.13384937 | 0.0263900603590167 | 0.979703696347405 | 0.998963492980642 | -6.4160803968412 |
| **Psph** | -0.174709277500002 | 15.77533416625 | -0.0250035142269201 | 0.980769811726033 | 0.998963492980642 | -6.41612019072364 |
| **Cnpy3** | 0.169790412499999 | 15.47263294125 | 0.0247815518133489 | 0.980940482632349 | 0.998963492980642 | -6.41612636170715 |
| **Dhcr7** | 0.166944847499998 | 15.82402188875 | 0.0238257060380523 | 0.98167546188297 | 0.998963492980642 | -6.41615230748468 |

| | | | | | | |
|---|---|---|---|---|---|---|
| **Dnaaf3** | 0.00207986249999692 | 29.68341527375 | 0.0235095040925392 | 0.981918603559476 | 0.998963492980642 | -6.41616066598961 |
| **Nedd4l** | 0.145262859999995 | 14.963004055 | 0.0218514542367338 | 0.983193584797268 | 0.998963492980642 | -6.41620266726868 |
| **Parn** | 0.147912342500003 | 16.03115625125 | 0.0208377523212017 | 0.98397311179133 | 0.998963492980642 | -6.41622683383458 |
| **Irgm1** | -0.00178288249999881 | 22.04902119875 | -0.0200000629698472 | 0.984617301113374 | 0.998963492980642 | -6.41624593828833 |
| **Rgs18** | -0.124762705 | 14.0981319275 | -0.0199825220016943 | 0.984630790382062 | 0.998963492980642 | -6.41624632995327 |
| **Aspm** | -0.148360334999996 | 17.17366587 | -0.0194980382712509 | 0.985003367794113 | 0.998963492980642 | -6.41625701197426 |
| **Gnptab** | 0.1191698175 | 14.06505104625 | 0.0191326934364896 | 0.985284327753338 | 0.998963492980642 | -6.41626489381975 |
| **Ddr1** | -0.119689897500001 | 14.39664043625 | -0.0187678397774153 | 0.985564912233317 | 0.998963492980642 | -6.4162726163026 |
| **Wdr76** | 0.107905149999997 | 13.684653985 | 0.0177842965187651 | 0.986321300247945 | 0.998963492980642 | -6.41629269339361 |
| **Wipf2** | 0.00372621999999367 | 21.855976905 | 0.0171543710783762 | 0.986805748635599 | 0.998963492980642 | -6.41630498452732 |
| **Eml5** | 0.111954097499998 | 15.16098081125 | 0.0166746672839422 | 0.987174672097206 | 0.998963492980642 | -6.41631404728806 |
| **Tp53bp1** | -0.116450215 | 16.1204372025 | -0.0163141553222685 | 0.987451931509302 | 0.998963492980642 | -6.41632068907782 |
| **Fam83d** | -0.109474234999999 | 15.296442735 | -0.0161589996166041 | 0.98757125789157 | 0.998963492980642 | -6.41632350286612 |
| **Rars1** | -0.0993167074999999 | 14.63046183875 | -0.0153289357437407 | 0.988209645054336 | 0.998963492980642 | -6.41633809960715 |
| **Acacb** | -0.0127771125000038 | 23.59322324625 | -0.0147900056632991 | 0.988624131484804 | 0.998963492980642 | -6.41634716473057 |
| **Plvap** | 0.00130691999999399 | 22.93152065 | 0.0134200385319327 | 0.989677778012354 | 0.998963492980642 | -6.41636874800363 |
| **Emsy** | -0.1093311975 | 10.73087528375 | -0.0132844348111164 | 0.98978207255536 | 0.998963492980642 | -6.41637077037262 |
| **Alpk3** | 0.0166914800000058 | 22.590739055 | 0.0132732922637811 | 0.989790642439567 | 0.998963492980642 | -6.4163709356374 |
| **Abracl** | -0.10056696 | 10.020564875 | -0.0130824722180179 | 0.989937404948194 | 0.998963492980642 | -6.41637374433312 |
| **Dpp4** | -0.0982721099999999 | 9.7956691675 | -0.0130812332912467 | 0.989938357826407 | 0.998963492980642 | -6.4163737624361 |
| **Zdhhc17** | -0.0868073900000006 | 15.570273125 | -0.0125916915918254 | 0.990314873474231 | 0.998963492980642 | -6.41638078136266 |

| | | | | | | |
|---|---|---|---|---|---|---|
| **Tktl1** | -0.00890073249999901 | 21.14871009375 | -0.0124683811025295 | 0.990409714294286 | 0.998963492980642 | -6.41638250715492 |
| **Pkhd1l1** | -0.0974896224999977 | 10.21530487125 | -0.0124416881198424 | 0.990430244478738 | 0.998963492980642 | -6.41638287850056 |
| **Icmt** | -0.00161054750000034 | 25.68372090125 | -0.0113769622437611 | 0.991249155740098 | 0.998963492980642 | -6.41639704172391 |
| **Mat2b** | -0.00280296249999878 | 25.48579729625 | -0.0113175989768729 | 0.991294814078929 | 0.998963492980642 | -6.41639779412186 |
| **Fat3** | 0.0802382075000025 | 16.57086650375 | 0.010933904154762 | 0.99158992785908 | 0.998963492980642 | -6.41640256231278 |
| **Agpat4** | 0.0630857924999972 | 16.47189055125 | 0.00864959639885831 | 0.993346900543564 | 0.998963492980642 | -6.41642754591338 |
| **Cirbp** | 0.053485310000001 | 14.4064393225 | 0.0083690913721063 | 0.993562653619371 | 0.998963492980642 | -6.41643021202563 |
| **Atp6ap2** | 0.0501497274999991 | 14.73425912625 | 0.00768451938355478 | 0.994089200993552 | 0.998963492980642 | -6.41643634970336 |
| **Adgrl2** | -0.00129730749999979 | 23.91690035125 | -0.00545825821277817 | 0.995801578983774 | 0.998963492980642 | -6.41645269063513 |
| **Kcnc2** | 0.0399637899999981 | 10.3311662175 | 0.00504417230691032 | 0.996120085235318 | 0.998963492980642 | -6.41645511945279 |
| **Dync1li2** | -0.0419123100000043 | 10.908716795 | -0.00500986295083487 | 0.996146475312286 | 0.998963492980642 | -6.41645531210224 |
| **Smap1** | -0.0360173600000024 | 16.46715142 | -0.00493519300820695 | 0.996203909970755 | 0.998963492980642 | -6.41645572683456 |
| **Atp12a** | -0.0364607700000015 | 17.2296607825 | -0.00477946649747701 | 0.996323691832694 | 0.998963492980642 | -6.41645657173405 |
| **Ccdc9** | 0.027341242499995 | 13.45366924125 | 0.00458902313006678 | 0.996470177377309 | 0.998963492980642 | -6.41645756817244 |
| **Aldh1a7** | -0.0265789399999932 | 14.1635442225 | -0.00423502087148938 | 0.996742469806124 | 0.998963492980642 | -6.41645931274436 |
| **Cbr3** | -0.0325863925000007 | 10.07638123125 | -0.00421276844370604 | 0.996759586005649 | 0.998963492980642 | -6.41645941773142 |
| **Itpka** | -0.024786347500001 | 9.62858039625 | -0.00335665349704204 | 0.997418096772702 | 0.998963492980642 | -6.41646303691239 |
| **Farsa** | 0.0212989100000023 | 15.1745461975 | 0.00316952186476745 | 0.997562035923761 | 0.998963492980642 | -6.41646371897101 |
| **Mtch1** | 0.0204003950000011 | 10.2379365625 | 0.00259840236615391 | 0.998001333991724 | 0.998963492980642 | -6.41646555873639 |
| **Fau** | 0.0160836599999996 | 9.8771925275 | 0.00212339582377803 | 0.998366703736857 | 0.998963492980642 | -6.41646681136723 |

| | | | | | | |
|---|---|---|---|---|---|---|
| **Wars1** | 0.011738602500003 | 15.01686824875 | 0.00176503474076544 | 0.998642351377886 | 0.998963492980642 | -6.41646758960337 |
| **Stub1** | 0.000532995000003922 | 24.9284269925 | 0.00158394571117725 | 0.998781643266316 | 0.998963492980642 | -6.41646792830792 |
| **Arhgap26** | 0.00949164999999752 | 15.8936585575 | 0.0013475286712082 | 0.998963492980642 | 0.998963492980642 | -6.4164683153705 |

#### Advanced stage (10M) samples

| Proteins | logFC | AveExpr | t | P.Value | adj.P.Val | B |
|---|---|---|---|---|---|---|
| **Hprt1** | -1.3591323725 | 24.17226623875 | -8.92385089918692 | 5.33737222260029E-05 | 0.123133177175389 | -0.856109487469427 |
| **Glo1** | 0.913138952500002 | 21.87902612125 | 6.09250387307869 | 0.00055195638943727 | 0.433405417825426 | -1.47806998524073 |
| **Atp6v1g1** | 1.609594225 | 22.36168014 | 5.79499310080259 | 0.000738898442121101 | 0.433405417825426 | -1.57910923092069 |
| **S100a1** | -0.554221285000004 | 23.446748925 | -5.77812052555108 | 0.000751461496012876 | 0.433405417825426 | -1.5851229860015 |
| **Kcnj10** | -1.11930644249999 | 21.53325515375 | -5.2587230073605 | 0.00128496357841983 | 0.592882195082911 | -1.78658915080341 |
| **Ndufaf2** | 0.536130594999996 | 25.725255245 | 4.73051394886262 | 0.00230038498642447 | 0.795708781321242 | -2.02788382226174 |
| **Rida** | 0.563373502499999 | 22.30107751625 | 4.68818245610298 | 0.00241437428229246 | 0.795708781321242 | -2.04899363720814 |
| **Ppp1ca** | 0.785653814999996 | 23.90994288 | 4.53657282016592 | 0.0028770903477836 | 0.829680929042095 | -2.12690503773288 |
| **Fabp3** | 0.881626275000002 | 21.524043775 | 4.37895571230097 | 0.00346472801053013 | 0.843284351171898 | -2.21184566281906 |
| **Lrch2** | -0.349673440000004 | 26.11364856 | -4.22945030915694 | 0.00414677904201104 | 0.843284351171898 | -2.29626911718359 |
| **Tusc3** | 0.81559011249999 | 25.41481022125 | 4.12456277062395 | 0.00471336089474102 | 0.843284351171898 | -2.35780674852181 |
| **Gsta4** | 0.512738652499987 | 23.40882585625 | 4.11585820744777 | 0.0047640783124083 | 0.843284351171898 | -2.36300084748268 |
| **Psap** | 0.585611067500011 | 26.71220375375 | 4.06469839446909 | 0.00507455996002373 | 0.843284351171898 | -2.39380111103271 |
| **Ank3** | -0.372480110000001 | 25.0323151925 | -3.85339966993277 | 0.00661403727773626 | 0.843284351171898 | -2.52602656066083 |
| **Arf6** | 0.61333384000001 | 23.419748425 | 3.82702018831084 | 0.00683971830057956 | 0.843284351171898 | -2.54310964946601 |

| | | | | | | |
|---|---|---|---|---|---|---|
| **Sacm1l** | -0.678637300000005 | 27.34605794 | -3.76564083175835 | 0.0073980916903606 | 0.843284351171898 | -2.58335925685958 |
| **Aco1** | -0.441581762499993 | 24.56868023375 | -3.72602764089303 | 0.00778477794585615 | 0.843284351171898 | -2.60970944838824 |
| **App** | 2.00422629999999 | 22.002186305 | 3.58599039770136 | 0.00933866732341837 | 0.843284351171898 | -2.70522762066284 |
| **Dclk1** | 0.727239887500001 | 18.73106556375 | 3.57724352392855 | 0.00944634644706298 | 0.843284351171898 | -2.71131675529712 |
| **Samm50** | 0.6878859025 | 24.48884188625 | 3.52449649899784 | 0.0101249564337241 | 0.843284351171898 | -2.74834400627994 |
| **Chp1** | 0.365929612500004 | 22.68541638375 | 3.51651739201056 | 0.0102321377511076 | 0.843284351171898 | -2.75399111311216 |
| **Gsr** | 0.353042235000004 | 26.232087305 | 3.48046405289571 | 0.0107320062124218 | 0.843284351171898 | -2.77965799605811 |
| **Got1** | 1.23645398 | 19.442056375 | 3.45636373568028 | 0.0110808701496151 | 0.843284351171898 | -2.79695289415089 |
| **Cds1** | -0.998655187499988 | 26.04665433625 | -3.44696557661205 | 0.0112202238631155 | 0.843284351171898 | -2.80372707514486 |
| **Ppp2r5c** | 0.909040934999993 | 21.6671231125 | 3.4327210634667 | 0.0114350663011167 | 0.843284351171898 | -2.81402643841493 |
| **Rab5a** | 0.266474377500003 | 26.20875381375 | 3.3710983738577 | 0.0124170921141761 | 0.843284351171898 | -2.85902510494965 |
| **Dnajc6** | -0.320608282499997 | 22.71985841125 | -3.33887449545894 | 0.01296656318109 | 0.843284351171898 | -2.8828419622915 |
| **Tpt1** | 0.807498322499995 | 19.04512329125 | 3.32889843365951 | 0.0131419450578406 | 0.843284351171898 | -2.89025506557811 |
| **Timm10** | 0.327602190000007 | 26.662912235 | 3.30746152256388 | 0.0135275235447728 | 0.843284351171898 | -2.90624805899656 |
| **Cbarp** | 0.411546830000002 | 26.4189625525 | 3.30375263313442 | 0.0135954623487286 | 0.843284351171898 | -2.90902385311262 |
| **Ist1** | -16.975260255 | 13.9905649725 | -3.28373277492072 | 0.013968584297042 | 0.843284351171898 | -2.92405170127833 |
| **Uqcrh** | 0.523540147499997 | 21.81451638125 | 3.26249844383252 | 0.0143764196155465 | 0.843284351171898 | -2.94007340656224 |
| **Cct2** | 0.329011642500003 | 26.27517555625 | 3.23368320406298 | 0.0149504647761904 | 0.843284351171898 | -2.96195003745307 |
| **Sncb** | 0.553029259999995 | 21.152068515 | 3.22433411708995 | 0.0151419680642783 | 0.843284351171898 | -2.96908120746865 |
| **Txnrd1** | -1.41301874000001 | 21.73151016 | -3.14622628021952 | 0.0168486803601684 | 0.843284351171898 | -3.02929323275496 |

| | | | | | | |
|---|---|---|---|---|---|---|
| **Rab18** | 0.334424870000003 | 22.8357626825 | 3.14300085612572 | 0.0169234525963839 | 0.843284351171898 | -3.03180387815877 |
| **Calb2** | 0.614758627499999 | 23.4622562462 5 | 3.13922606187689 | 0.0170114119645357 | 0.843284351171898 | -3.03474457554496 |
| **Vamp2** | -0.800978377499998 | 29.6407304787 5 | -3.11572938959283 | 0.0175700480578894 | 0.843284351171898 | -3.0531080244788 |
| **Uqcrq** | -0.628277714999996 | 25.63665504 | -3.08332163140994 | 0.0183729673833798 | 0.843284351171898 | -3.07860106334684 |
| **Pdcd6** | 15.11918016 | 12.78877465 | 3.07661944001773 | 0.0185438601734608 | 0.843284351171898 | -3.08389702746355 |
| **Mdp1** | 0.3274238475 | 26.7453103237 5 | 3.06830417877422 | 0.0187582462891665 | 0.843284351171898 | -3.09047890359894 |
| **Pa2g4** | 0.325908934999994 | 27.480544255 | 3.06742761276338 | 0.0187809998498088 | 0.843284351171898 | -3.09117346961972 |
| **Ncald** | -0.4112331 | 22.20895414 | -3.05518658180108 | 0.0191018449285045 | 0.843284351171898 | -3.10088738475102 |
| **Sez6l2** | -0.447350167500005 | 22.0812627137 5 | -3.0314021632627 | 0.0197420909569884 | 0.843284351171898 | -3.11983854621327 |
| **Lgi3** | 0.865667232500009 | 24.9640087112 5 | 3.02756051440483 | 0.0198476319889045 | 0.843284351171898 | -3.12290902401247 |
| **Pvalb** | -0.548574459999998 | 21.8407185475 | -3.01647370593538 | 0.0201556040232911 | 0.843284351171898 | -3.13178502038703 |
| **Uso1** | 0.35869519249999 | 24.7364048462 5 | 3.00746799751115 | 0.0204095138636421 | 0.843284351171898 | -3.13901099249547 |
| **Akr1b1** | 0.333350414999998 | 20.9481004775 | 3.00393220701588 | 0.0205101324756162 | 0.843284351171898 | -3.141851961065 |
| **Plec** | -0.386204882500003 | 26.0062281612 5 | -2.99848217680737 | 0.0206662608009918 | 0.843284351171898 | -3.14623533413354 |
| **Gstm1** | 0.372881972499989 | 26.9827007712 5 | 2.97417016246156 | 0.0213783001162197 | 0.843284351171898 | -3.16585289990969 |
| **Kcna1** | 1.18960055 | 22.5914117675 | 2.95001137435578 | 0.0221117477720426 | 0.843284351171898 | -3.18544942341859 |
| **Hcn2** | -1.08254028999999 | 24.3701038925 | -2.93652000687469 | 0.0225329307957871 | 0.843284351171898 | -3.1964371810307 |
| **C1qbp** | 0.364046335000001 | 24.7090340725 | 2.92725828877739 | 0.0228269948985677 | 0.843284351171898 | -3.20399840272001 |
| **Pdp1** | 0.760536442500005 | 23.4722154662 5 | 2.92369715757571 | 0.0229411441824912 | 0.843284351171898 | -3.20690962542445 |
| **Lgi2** | 0.359566760000003 | 25.80550044 | 2.90986479220764 | 0.0233903064812466 | 0.843284351171898 | -3.21823821936866 |
| **Atp5if1** | -1.21104420500001 | 20.64253287 | -2.9021728544936 | 0.0236441040162393 | 0.843284351171898 | -3.22455200857099 |

| | | | | | | |
|---|---|---|---|---|---|---|
| **Arpc4** | -0.604612785 | 23.910263895 | -2.89019044776154 | 0.0240452980355455 | 0.843284351171898 | -3.23440761002469 |
| **Ndrg4** | -0.422864162500002 | 23.31985615625 | -2.87604359874374 | 0.024528263725841 | 0.843284351171898 | -3.24607476895839 |
| **Smpd3** | 0.492702280000003 | 27.632693515 | 2.87566883458363 | 0.0245411967719415 | 0.843284351171898 | -3.24638430245737 |
| **Uchl1** | 0.476437290000007 | 23.3487959425 | 2.79973483491188 | 0.0273164474061825 | 0.843284351171898 | -3.30958266186512 |
| **Ube2l3** | 0.896058842500004 | 24.75694673125 | 2.79416116223202 | 0.027532804707882 | 0.843284351171898 | -3.31425871057522 |
| **Cndp2** | 0.52157892000001 | 24.2390237625 | 2.77210898295874 | 0.0284066896480774 | 0.843284351171898 | -3.33280835294669 |
| **Rab2a** | 0.587777992499998 | 25.60488516875 | 2.75400788257259 | 0.0291458193061146 | 0.843284351171898 | -3.34809226366168 |
| **Krt73** | 1.07196737749999 | 22.44166053125 | 2.74216731934613 | 0.0296402350420921 | 0.843284351171898 | -3.3581178866595 |
| **Tkt** | 0.306840857499996 | 26.58885228625 | 2.74209891192506 | 0.0296431169560412 | 0.843284351171898 | -3.35817587212955 |
| **Acyp2** | 0.793148102499998 | 21.81625554375 | 2.72515735810758 | 0.0303659899288669 | 0.843284351171898 | -3.37255872693131 |
| **Hagh** | -0.417484879999996 | 27.8302634825 | -2.72236614689719 | 0.030486853030064 | 0.843284351171898 | -3.37493263409634 |
| **Psma6** | 0.212610704999996 | 26.06646001 | 2.71668995637545 | 0.0307342003199677 | 0.843284351171898 | -3.37976388854414 |
| **Vamp3** | 1.0455621375 | 24.59370940375 | 2.70883039967721 | 0.0310801739112277 | 0.843284351171898 | -3.38646163921771 |
| **DSCC1** | 0.233378572499991 | 26.30954901125 | 2.70850824818307 | 0.0310944417567898 | 0.843284351171898 | -3.38673637094925 |
| **Pfn1** | -0.339543612499998 | 25.32984351875 | -2.70819695778491 | 0.0311082351019135 | 0.843284351171898 | -3.38700185530704 |
| **Rpl28** | 0.903150872500007 | 22.04182686125 | 2.7079216639542 | 0.0311204387808735 | 0.843284351171898 | -3.38723665223457 |
| **Vat1** | 0.618086029999994 | 20.2796693975 | 2.6988786678532 | 0.0315241167017986 | 0.843284351171898 | -3.39495577833069 |
| **Ldhb** | 0.648043672500002 | 23.10908644375 | 2.69229947758492 | 0.0318212596425491 | 0.843284351171898 | -3.40057956019426 |
| **Pde1b** | 1.3239008225 | 21.37399285625 | 2.68885498275996 | 0.0319779967276016 | 0.843284351171898 | -3.40352645323887 |
| **Nrxn2** | 0.692200467500001 | 22.93547731875 | 2.68278261768607 | 0.0322562852383783 | 0.843284351171898 | -3.40872591443638 |

| | | | | | | |
|---|---|---|---|---|---|---|
| **Ca8** | 0.255409430000007 | 21.974899415 | 2.67517296732916 | 0.0326086121631426 | 0.843284351171898 | -3.41524943748105 |
| **Arpc2** | 0.3255493825 | 24.14485322375 | 2.67457883312619 | 0.0326362897006717 | 0.843284351171898 | -3.41575913273588 |
| **Ppp2r1a** | 0.359409750000005 | 26.23332034 | 2.66384820827694 | 0.033140428779045 | 0.843284351171898 | -3.42497368882475 |
| **Anxa5** | -0.476353194999998 | 24.720124635 | -2.6636037391207 | 0.0331520087945061 | 0.843284351171898 | -3.42518381593123 |
| **Atp5me** | 0.421752492500008 | 23.77902614375 | 2.65898552427125 | 0.0333715617947225 | 0.843284351171898 | -3.42915492915283 |
| **Slc39a12** | 0.494913430000004 | 24.5928421225 | 2.6553051442817 | 0.0335476186280943 | 0.843284351171898 | -3.43232185060634 |
| **Dpm3** | 0.335125875000006 | 24.0681521975 | 2.65204386800667 | 0.0337044389487378 | 0.843284351171898 | -3.43512978912441 |
| **Slc38a3** | 0.6723611675 | 20.72829833375 | 2.64931163019516 | 0.0338364103535421 | 0.843284351171898 | -3.4374834194979 |
| **Kcnab2** | 0.639822744999996 | 22.1007952225 | 2.64205921059876 | 0.0341893408973315 | 0.843284351171898 | -3.44373611290985 |
| **Marcksl1** | 0.878668524999995 | 24.8236681725 | 2.63919319897709 | 0.0343298709315131 | 0.843284351171898 | -3.44620914521719 |
| **Atp6v1e1** | 1.2570940125 | 25.44076545375 | 2.63403842068561 | 0.0345841480741145 | 0.843284351171898 | -3.45066008493311 |
| **Ddt** | 0.273439987499998 | 25.30625370375 | 2.62945564869245 | 0.0348118613997377 | 0.843284351171898 | -3.45462031397082 |
| **Hpcal4** | 0.421622914999993 | 23.723388655 | 2.6084494850991 | 0.0358758484861385 | 0.843284351171898 | -3.4728110147343 |
| **Pex5l** | 0.6856917725 | 19.73815471875 | 2.60789377019732 | 0.0359044525761387 | 0.843284351171898 | -3.47329308939011 |
| **Hspa4l** | 0.527184927500002 | 24.03521735125 | 2.59327629658997 | 0.03666545500689 | 0.843284351171898 | -3.4859889481677 |
| **Anxa6** | 13.0834674425 | 15.85681676875 | 2.58967950800062 | 0.0368552729841638 | 0.843284351171898 | -3.48911742637551 |
| **Ddah2** | -0.296859169999998 | 25.892201125 | -2.57728407420842 | 0.0375173137592074 | 0.843284351171898 | -3.49991249224938 |
| **Pfn2** | 1.383959255 | 22.211542025 | 2.56762366642885 | 0.0380418558650054 | 0.843284351171898 | -3.50834008099498 |
| **Actr1b** | 0.361914687499993 | 26.70931047125 | 2.56719309113875 | 0.0380654122112348 | 0.843284351171898 | -3.50871600008402 |
| **Mpp3** | 0.40202800749999 | 25.72916894875 | 2.53887668329031 | 0.0396483580343319 | 0.843284351171898 | -3.53349191721287 |

| | | | | | | |
|---|---|---|---|---|---|---|
| **Gda** | -0.219473174999994 | 28.440009375 | -2.52594449108418 | 0.040393895819457 | 0.843284351171898 | -3.54484193792561 |
| **Nectin1** | 0.345986987500005 | 24.81130851125 | 2.5238829615303 | 0.0405140804341945 | 0.843284351171898 | -3.54665323371082 |
| **Acyp1** | -1.7228525075 | 23.25144490125 | -2.51585334066455 | 0.0409857447489374 | 0.843284351171898 | -3.55371334360656 |
| **Praf2** | 0.649884610000001 | 26.3906922175 | 2.51278404736475 | 0.0411675378276193 | 0.843284351171898 | -3.55641419616247 |
| **Nme1** | 0.878776585000001 | 24.0273758675 | 2.50343339346995 | 0.0417265429047105 | 0.843284351171898 | -3.56464965997384 |
| **Cyrib** | 0.439532912499999 | 22.09918833375 | 2.49476504932487 | 0.0422517830882857 | 0.843284351171898 | -3.57229387887734 |
| **Napa** | 0.566751992500002 | 20.64007027375 | 2.49167497272167 | 0.0424406711180398 | 0.843284351171898 | -3.57502111040017 |
| **Cyc1** | -0.910877437500005 | 21.20111087875 | -2.48246217082751 | 0.0430090294090309 | 0.843284351171898 | -3.5831590218811 |
| **Zfyve28** | 0.677248510000009 | 22.99712352 | 2.47990186091492 | 0.0431683749090979 | 0.843284351171898 | -3.58542243426971 |
| **Ppa1** | 0.322764897500001 | 20.97382326875 | 2.47443938335348 | 0.0435103863915429 | 0.843284351171898 | -3.59025410366419 |
| **Nbea** | -1.5452809975 | 23.19386343625 | -2.47320307222857 | 0.0435881812308666 | 0.843284351171898 | -3.5913481399997 |
| **Gpi** | 0.316490810000001 | 25.201245755 | 2.47216341106556 | 0.0436537131843843 | 0.843284351171898 | -3.59226829769437 |
| **Otub1** | 0.379263545000001 | 19.13276686 | 2.46765728873073 | 0.043938922660679 | 0.843284351171898 | -3.59625794792047 |
| **Gnai2** | 0.321297519999995 | 19.73373165 | 2.4661257491258 | 0.0440362975177722 | 0.843284351171898 | -3.5976144958086 |
| **Ndufab1** | 0.505582705000002 | 22.2668400225 | 2.45546666957438 | 0.0447202034887777 | 0.843284351171898 | -3.60706330263791 |
| **Calb1** | -1.1225658575 | 21.03986194875 | -2.45340048152636 | 0.0448540381651448 | 0.843284351171898 | -3.60889641875673 |
| **Aldh6a1** | 1.003151 | 23.390614235 | 2.43554930413405 | 0.046027679699848 | 0.843284351171898 | -3.62475431653373 |
| **Uqcrb** | 0.686955727499999 | 23.03204138375 | 2.42284057061226 | 0.0468824885431477 | 0.843284351171898 | -3.63606586962825 |
| **Trpv2** | 0.456408875000001 | 24.0374965025 | 2.42130610682735 | 0.0469867999433604 | 0.843284351171898 | -3.63743284721808 |
| **Enoph1** | -0.965727865000005 | 28.8423717675 | -2.41116223872074 | 0.0476824096435146 | 0.843284351171898 | -3.64647598882604 |

| | | | | | | |
|---|---|---|---|---|---|---|
| **Cntnap1** | 0.937916592499992 | 23.25893855875 | 2.40852314310687 | 0.0478651158670276 | 0.843284351171898 | -3.64883054062257 |
| **Pgam1** | 0.38824121 | 23.04664471 | 2.39756946721387 | 0.0486311776922028 | 0.843284351171898 | -3.65861116484856 |
| **Lmbrd2** | -0.230635145000008 | 28.7972163875 | -2.39537621369788 | 0.048786075113112 | 0.843284351171898 | -3.66057106536862 |
| **Slc12a2** | 0.200443320000002 | 27.65135022 | 2.39088164783487 | 0.0491050880680085 | 0.843284351171898 | -3.66458900322414 |
| **Oxct1** | 0.316042149999991 | 23.560586385 | 2.38849789262184 | 0.0492751502156107 | 0.843284351171898 | -3.66672082652034 |
| **Apmap** | 0.443977327500004 | 20.75846578625 | 2.37625468473076 | 0.0501581886305558 | 0.843284351171898 | -3.67767930715652 |
| **Prkaca** | -0.945865897499999 | 25.71221670125 | -2.37593091287347 | 0.0501817598880581 | 0.843284351171898 | -3.67796931189498 |
| **Nme2** | 1.464215915 | 20.946102645 | 2.36807136981278 | 0.050757451565309 | 0.843284351171898 | -3.68501239726105 |
| **Ugp2** | 1.13185363499999 | 23.682175175 | 2.36687976733829 | 0.0508453231471504 | 0.843284351171898 | -3.68608075429841 |
| **Capza2** | -0.501828195000002 | 23.42300776 | -2.36534067635963 | 0.0509590503247077 | 0.843284351171898 | -3.68746086843885 |
| **Slc16a1** | 1.84224424249999 | 23.80508819125 | 2.35476444606284 | 0.0517476409983802 | 0.843284351171898 | -3.69695095881255 |
| **Pak1** | -0.251723152500006 | 25.27396840625 | -2.34892182269021 | 0.0521886339967721 | 0.843284351171898 | -3.70219823023686 |
| **Sgta** | 0.256770294999999 | 25.9034077025 | 2.34195393089812 | 0.0527195971926785 | 0.843284351171898 | -3.70846037626716 |
| **Aldoc** | 0.467479227499993 | 25.01434175375 | 2.33743121187081 | 0.0530671915854806 | 0.843284351171898 | -3.71252746425475 |
| **Ephb3** | 0.382343922499999 | 25.19756478875 | 2.3361122979565 | 0.0531689980193911 | 0.843284351171898 | -3.71371386765122 |
| **Ndufs4** | 0.310235922499995 | 25.24144844375 | 2.32710846945355 | 0.0538693587503424 | 0.843284351171898 | -3.72181738708097 |
| **Uqcrfs1** | 0.246585502500004 | 21.62451815625 | 2.3218445573895 | 0.0542831736771464 | 0.843284351171898 | -3.7265583849313 |
| **Idh1** | 0.684507052499992 | 23.00136728125 | 2.31932568896715 | 0.0544823388753655 | 0.843284351171898 | -3.7288279149998 |
| **Ca2** | 1.88835757 | 22.200731985 | 2.30846206923206 | 0.055349907437091 | 0.843284351171898 | -3.73862262834672 |
| **Lactb** | 0.299551419999993 | 24.2870186 | 2.29173347706489 | 0.0567135197174291 | 0.843284351171898 | -3.75372526235816 |

| | | | | | | |
|---|---|---|---|---|---|---|
| **Ociad2** | -0.880102044999994 | 21.97807939 | -2.29127631312055 | 0.0567512621844735 | 0.843284351171898 | -3.75413832438992 |
| **Rab5b** | 0.721139442499993 | 24.01488874625 | 2.28767468615657 | 0.0570495015418145 | 0.843284351171898 | -3.75739311577984 |
| **Nrcam** | 0.494954422499998 | 21.20871281375 | 2.28516941880223 | 0.0572578963650445 | 0.843284351171898 | -3.75965775916593 |
| **Hmox2** | 0.717201199999991 | 24.8476871475 | 2.27057046857257 | 0.0584877951350545 | 0.843284351171898 | -3.77286465044299 |
| **Adsl** | 0.944859212499999 | 22.06638539875 | 2.26742077749856 | 0.0587566509744543 | 0.843284351171898 | -3.7757162329285 |
| **Ckb** | -0.367768567500001 | 26.83053823625 | -2.26422140006146 | 0.059031035360552 | 0.843284351171898 | -3.77861359141968 |
| **Ndufb1** | -0.427243295 | 25.2555298475 | -2.25378480935331 | 0.0599351873090218 | 0.843284351171898 | -3.7880704344018 |
| **Osbpl1a** | 0.802354407500005 | 21.21132556625 | 2.25326338676675 | 0.059980727100305 | 0.843284351171898 | -3.78854312384925 |
| **Lrpap1** | 1.32439771249999 | 21.83522919625 | 2.2460319356664 | 0.0606159411582205 | 0.843284351171898 | -3.79510079001401 |
| **Ganab** | -0.692494335000006 | 29.0025607525 | -2.24507594695834 | 0.060700425439202 | 0.843284351171898 | -3.79596799293574 |
| **Mdh1** | -0.244898729999999 | 19.4669479975 | -2.23649974941674 | 0.061463702581734 | 0.843284351171898 | -3.80375064222006 |
| **Rad23b** | 0.405690515000003 | 19.9658942425 | 2.23509510755969 | 0.0615896400673029 | 0.843284351171898 | -3.80502581377878 |
| **Neo1** | 0.1977148 | 26.2149117775 | 2.23437005572703 | 0.0616547492240172 | 0.843284351171898 | -3.80568408969712 |
| **Svip** | 0.338563247499991 | 29.31865595125 | 2.23432318765434 | 0.0616589603465116 | 0.843284351171898 | -3.80572664257995 |
| **Adgrg1** | 0.846654302499996 | 23.61165685875 | 2.23066389692338 | 0.0619886518628652 | 0.843284351171898 | -3.80904949311604 |
| **Clta** | 0.201597192500003 | 24.97052938125 | 2.22890237819569 | 0.0621479964756425 | 0.843284351171898 | -3.8106493872773 |
| **Ehd3** | 0.450356642500012 | 23.79438233125 | 2.22772071858392 | 0.0622551204897902 | 0.843284351171898 | -3.81172274606604 |
| **Gdi1** | -0.288803097499994 | 25.02902827375 | -2.22372992643039 | 0.0626182935055383 | 0.843284351171898 | -3.81534848248069 |
| **Nnt** | 0.718211012499999 | 21.08285069125 | 2.22239330993242 | 0.0627404086747794 | 0.843284351171898 | -3.81656307436915 |
| **Cox6c** | -0.885604827499982 | 24.92324363625 | -2.21599567721063 | 0.0633282551250325 | 0.843284351171898 | -3.82237829992712 |

| | | | | | | |
|---|---|---|---|---|---|---|
| **Mcam** | -0.232607115 | 22.75911185 | -2.21261247032912 | 0.0636413728907154 | 0.843284351171898 | -3.82545460979513 |
| **Vdac1** | 0.710957239999999 | 23.529062585 | 2.21239241501766 | 0.0636617933487443 | 0.843284351171898 | -3.82565472933231 |
| **Tppp3** | 0.429297404999996 | 20.82356948 | 2.21183234088341 | 0.0637137964413858 | 0.843284351171898 | -3.82616407806999 |
| **Atp5pf** | 0.360621272500001 | 25.72861780875 | 2.20963318742332 | 0.0639184049093744 | 0.843284351171898 | -3.82816425230263 |
| **Ptprn2** | 0.290164507500002 | 25.22404374125 | 2.20642781264736 | 0.0642178235250042 | 0.843284351171898 | -3.83108016051943 |
| **Itfg1** | 0.418685992500002 | 24.04957903875 | 2.20595152722397 | 0.0642624349781936 | 0.843284351171898 | -3.83151349001762 |
| **Rev1** | -0.471978229999998 | 21.8084394875 | -2.20244765662752 | 0.0645915915308378 | 0.843284351171898 | -3.83470178804958 |
| **Endod1** | 0.4275415125 | 23.33621402125 | 2.2007512125274 | 0.0647515691079237 | 0.843284351171898 | -3.83624571964794 |
| **Aqp4** | 0.915153554999993 | 24.8769786175 | 2.19887383202437 | 0.064929076424718 | 0.843284351171898 | -3.83795452917552 |
| **Gde1** | -0.813901850000008 | 22.6362723975 | -2.19836116024263 | 0.064977635284083 | 0.843284351171898 | -3.83842120565623 |
| **Pcyox1l** | 0.505851499999999 | 23.8169472825 | 2.19671401984996 | 0.0651338965085657 | 0.843284351171898 | -3.83992067883521 |
| **Adam11** | 0.335165089999997 | 22.4786349075 | 2.19488195670334 | 0.0653081473138829 | 0.843284351171898 | -3.8415886906349 |
| **Vamp1** | 0.205855062499992 | 27.85762607375 | 2.1922985427503 | 0.0655546613761071 | 0.843284351171898 | -3.84394111866789 |
| **Mag** | 0.431761367499998 | 24.44746613125 | 2.17553859499974 | 0.0671769147080035 | 0.843284351171898 | -3.85921205006532 |
| **Dpysl3** | 0.3345040275 | 25.47439321625 | 2.16624477268342 | 0.0680938958073244 | 0.843284351171898 | -3.8676869801564 |
| **Prps1** | 0.324261897499998 | 21.26618521875 | 2.1618629143436 | 0.0685306018927588 | 0.843284351171898 | -3.87168435197342 |
| **Lamp5** | 0.622595195000002 | 21.819508035 | 2.16023634865845 | 0.0686934264530059 | 0.843284351171898 | -3.8731684498083 |
| **Sar1a** | 0.319053125000003 | 26.3545986 | 2.15467591383678 | 0.0692529929216444 | 0.843284351171898 | -3.87824288010209 |
| **Rap1gds1** | -0.214579825000001 | 25.819612585 | -2.15266219736546 | 0.0694567706037948 | 0.843284351171898 | -3.88008097405627 |
| **Psmd6** | 0.381283332500001 | 20.32460041375 | 2.13847463287064 | 0.0709096825384377 | 0.843284351171898 | -3.89303676813174 |

| | | | | | | |
|---|---|---|---|---|---|---|
| **Cadps** | 0.971924719999993 | 24.4712951475 | 2.13719379607636 | 0.071042344575955 | 0.843284351171898 | -3.89420686684748 |
| **Cab39** | 0.337011759999992 | 25.2381304775 | 2.13469761387814 | 0.0713016026369496 | 0.843284351171898 | -3.89648744862893 |
| **Tcp1** | -0.239430902500001 | 28.55891975125 | -2.13362224229915 | 0.0714135855279931 | 0.843284351171898 | -3.8974700242752 |
| **Kcnc3** | 0.206176747500002 | 24.98824564625 | 2.13051590568557 | 0.0717380546114868 | 0.843284351171898 | -3.90030859635162 |
| **Etfdh** | 0.481344602500002 | 24.20784045375 | 2.12069224070768 | 0.072773951477937 | 0.843284351171898 | -3.90928819550715 |
| **Tmem163** | 0.182560369999997 | 27.6490831175 | 2.12047187013068 | 0.0727973607488047 | 0.843284351171898 | -3.90948967768925 |
| **Cox5b** | 0.360714364999996 | 25.5686953825 | 2.11234947533003 | 0.0736654677361697 | 0.843284351171898 | -3.91691724414071 |
| **Npepps** | 0.288146779999998 | 24.1858897175 | 2.11102845778875 | 0.0738076339988801 | 0.843284351171898 | -3.91812549996184 |
| **Ndufb8** | 0.331567869999994 | 22.2623377225 | 2.09900302164445 | 0.0751145060888774 | 0.843284351171898 | -3.92912743141069 |
| **Pura** | 1.3084787625 | 22.24332050625 | 2.08658347259154 | 0.0764885331323506 | 0.843284351171898 | -3.94049520630911 |
| **Agpat3** | 0.253235385000007 | 27.8784526925 | 2.08598024776875 | 0.0765559070094988 | 0.843284351171898 | -3.94104747294673 |
| **Rufy3** | 0.6659818175 | 22.52636631125 | 2.08503811372389 | 0.0766612522127461 | 0.843284351171898 | -3.94191004160042 |
| **Psat1** | 0.348591772499994 | 25.34422506625 | 2.08208007137256 | 0.07699294994039 | 0.843284351171898 | -3.9446184468964 |
| **Lasp1** | 0.808520852499996 | 23.42258920125 | 2.07905554663916 | 0.0773335857688025 | 0.843284351171898 | -3.94738799462153 |
| **Gdi2** | 0.329207234999991 | 22.886387435 | 2.07488986929962 | 0.0778052097726659 | 0.843284351171898 | -3.95120292549407 |
| **Tnc** | 0.337837522499999 | 23.84756400125 | 2.07124074730336 | 0.0782207107851564 | 0.843284351171898 | -3.95454519205944 |
| **Atp5pd** | 11.97056221 | 16.6492285875 | 2.06903543615427 | 0.0784728878920503 | 0.843284351171898 | -3.95656523178667 |
| **Cntnap2** | 0.717844845000002 | 24.7127109725 | 2.06809294529767 | 0.0785809089756254 | 0.843284351171898 | -3.95742858152063 |
| **Hars1** | 0.374367382499997 | 22.58900565875 | 2.06568010580869 | 0.0788581268236145 | 0.843284351171898 | -3.95963891843701 |
| **Emc1** | 0.4073764125 | 23.33505568125 | 2.06431215230776 | 0.0790157280069127 | 0.843284351171898 | -3.96089212882063 |

| | | | | | | |
|---|---|---|---|---|---|---|
| **Rab10** | 0.31958032 | 24.7101464575 | 2.06132057150714 | 0.0793614821089293 | 0.843284351171898 | -3.9636329377405 |
| **Itgav** | 0.209226369999996 | 26.09410211 | 2.05320661690561 | 0.0803068711139043 | 0.843284351171898 | -3.97106778359167 |
| **Cacybp** | -0.480683472499997 | 23.76499543625 | -2.05194766050357 | 0.080454559562736 | 0.843284351171898 | -3.972221500201 |
| **Gpr37** | 0.272016757500001 | 24.87547188375 | 2.05147001551232 | 0.0805106629773648 | 0.843284351171898 | -3.97265922622407 |
| **Ppp1r7** | 0.349654942499999 | 25.45632430875 | 2.04368097588587 | 0.0814310686636009 | 0.843284351171898 | -3.97979794898012 |
| **Ndufb9** | 0.358878090000001 | 22.9835874 | 2.04304447967412 | 0.0815067427388269 | 0.843284351171898 | -3.98038135493168 |
| **Psma4** | 0.404335060000001 | 22.9918558275 | 2.03671713987219 | 0.0822628262486482 | 0.843284351171898 | -3.986181321493 |
| **St13** | 0.208484150000004 | 27.2218203975 | 2.03111999738606 | 0.0829374672485792 | 0.843284351171898 | -3.99131250152218 |
| **Pllp** | 0.340126990000009 | 26.701267965 | 2.03096633038832 | 0.0829560665795812 | 0.843284351171898 | -3.99145338247117 |
| **Cend1** | 0.273658829999995 | 26.324390885 | 2.02803950817208 | 0.0833111115953575 | 0.843284351171898 | -3.99413673963613 |
| **Nipsnap1** | 0.527261344999996 | 24.05587214 | 2.02738261649464 | 0.0833910046186604 | 0.843284351171898 | -3.99473900482633 |
| **Gosr2** | -1.540139455 | 21.7850021 | -2.02461391548399 | 0.0837285791684814 | 0.843284351171898 | -3.99727752647201 |
| **Mcee** | 11.1435693575 | 15.61814585625 | 2.02357054032911 | 0.0838561447694908 | 0.843284351171898 | -3.99823418492739 |
| **Mmut** | 0.287124549999994 | 26.7330487375 | 2.02096471142893 | 0.0841755833321715 | 0.843284351171898 | -4.00062349691283 |
| **Them4** | 0.620329410000004 | 22.9353860425 | 2.0054168960401 | 0.086106781444019 | 0.843284351171898 | -4.01488089238173 |
| **Prrt1** | 0.822216297500006 | 22.74211039875 | 1.99694759537264 | 0.0871771677449542 | 0.843284351171898 | -4.02264802136248 |
| **Btbd17** | 0.193752237499993 | 25.72019428875 | 1.99570284387461 | 0.0873355914485233 | 0.843284351171898 | -4.02378960183587 |
| **Uqcrc1** | 0.326286924999994 | 25.930309745 | 1.9944667054639 | 0.0874932009719059 | 0.843284351171898 | -4.02492328860165 |
| **Scarb2** | 0.294029057500001 | 26.40782335375 | 1.98830102639323 | 0.0882835474238358 | 0.843284351171898 | -4.03057801426792 |
| **Ndufb11** | 11.06092526 | 15.821645385 | 1.97992250594067 | 0.0893688738144205 | 0.843284351171898 | -4.03826226024942 |
| **Tollip** | -0.192711862500001 | 27.54865102875 | -1.97970946477684 | 0.0893966417127454 | 0.843284351171898 | -4.03845764747295 |

| | | | | | | |
|---|---|---|---|---|---|---|
| **Arhgdia** | -0.534942049999998 | 21.8922020375 | -1.97181306039271 | 0.090431897111987 | 0.843284351171898 | -4.04569962818862 |
| **Ndufa8** | 0.213252555 | 26.7026917575 | 1.96591534500665 | 0.0912128275657673 | 0.843284351171898 | -4.05110840279534 |
| **Psma3** | -1.0129367525 | 22.28926255125 | -1.96135795347725 | 0.0918208357337931 | 0.843284351171898 | -4.05528782883344 |
| **Eps15l1** | -0.289718026666666 | 18.9149197914286 | -1.99268382134536 | 0.0949709499662323 | 0.843284351171898 | -4.05810236620717 |
| **Pebp1** | 0.185760855000009 | 26.4777911025 | 1.94842063179435 | 0.0935686704492373 | 0.843284351171898 | -4.06715123063087 |
| **Ube2n** | 0.235853000000002 | 24.468273625 | 1.94617600391117 | 0.0938752377254254 | 0.843284351171898 | -4.06920934995847 |
| **Gstm5** | 0.542274817500005 | 23.88877071625 | 1.94601696795769 | 0.0938969960338208 | 0.843284351171898 | -4.06935516905884 |
| **Adam10** | 11.662959835 | 16.890253 | 1.9437797030144 | 0.0942036111118145 | 0.843284351171898 | -4.07140646887109 |
| **Eif5a** | 0.343891734999989 | 24.06410083 | 1.93605941271052 | 0.0952692511199631 | 0.843284351171898 | -4.07848449259679 |
| **Sars1** | 0.303092990000003 | 27.5941763325 | 1.93475748063816 | 0.0954501222254794 | 0.843284351171898 | -4.07967802378886 |
| **Mal2** | 0.278770587499995 | 25.43361079875 | 1.93392715728838 | 0.095565651205681 | 0.843284351171898 | -4.08043919849719 |
| **Grb2** | -0.897838727500002 | 24.67518608625 | -1.93133570601442 | 0.0959271030474378 | 0.843284351171898 | -4.08281476017427 |
| **Lrp1** | -0.227223817499997 | 30.79927742625 | -1.92845469183341 | 0.0963305190710899 | 0.843284351171898 | -4.08545562151123 |
| **Me1** | -10.5504591525 | 15.37174557875 | -1.92538939298879 | 0.0967615687568343 | 0.843284351171898 | -4.08826523505913 |
| **Vmp1** | 10.93004365 | 15.9328041525 | 1.92478569568598 | 0.096846685010285 | 0.843284351171898 | -4.08881855463719 |
| **Cyb5a** | -0.388265600000008 | 20.590063355 | -1.92448642605724 | 0.0968889067553467 | 0.843284351171898 | -4.08909284792077 |
| **Atp1b3** | -0.303611099999994 | 29.1372355225 | -1.91850312524624 | 0.0977368467343256 | 0.843284351171898 | -4.09457640607829 |
| **Tpi1** | -0.786431289999996 | 20.08262551 | -1.91804593334076 | 0.0978019374763753 | 0.843284351171898 | -4.09499538026814 |
| **Psmb1** | -11.00906546 | 16.0562661175 | -1.91471623950507 | 0.0982772720747436 | 0.843284351171898 | -4.09804659442733 |
| **Kpnb1** | 9.851674805 | 14.4481379425 | 1.90748572476512 | 0.0993172830315206 | 0.843284351171898 | -4.10467146885127 |

| | | | | | | |
|---|---|---|---|---|---|---|
| **Creld1** | -0.551292455000002 | 21.79791046 | -1.90630855334516 | 0.0994876204389513 | 0.843284351171898 | -4.10574991304613 |
| **Itgb1** | 0.979146517500002 | 22.53588873375 | 1.90325868388701 | 0.0999302699654242 | 0.843284351171898 | -4.10854382360837 |
| **Etfb** | 0.230265525 | 25.7869296675 | 1.89939851124976 | 0.100493289169615 | 0.843284351171898 | -4.11207966469626 |
| **Fkbp2** | 0.463816487500004 | 25.13120393625 | 1.89895235531777 | 0.100558562262035 | 0.843284351171898 | -4.11248830726217 |
| **Sh3bgrl3** | 0.917086009999998 | 23.2330004375 | 1.89818952501212 | 0.100670261240792 | 0.843284351171898 | -4.1131869844017 |
| **Alb** | 0.472760895 | 21.8234287375 | 1.89776149119309 | 0.100732990106153 | 0.843284351171898 | -4.11357901359932 |
| **Aars1** | -0.262782762499995 | 29.40705011625 | -1.89622434141998 | 0.100958576533301 | 0.843284351171898 | -4.11498681931562 |
| **Timm9** | 1.6924330075 | 21.53652556125 | 1.8944127167874 | 0.101225078108239 | 0.843284351171898 | -4.11664591296164 |
| **Psmb6** | -9.73399531 | 14.3409834525 | -1.89201834936599 | 0.101578360233525 | 0.843284351171898 | -4.11883853024846 |
| **Ppid** | 0.215343239999996 | 23.8129849175 | 1.89143265642516 | 0.101664960843769 | 0.843284351171898 | -4.11937484521076 |
| **Wdr1** | -0.417178635000003 | 26.2921345875 | -1.88476877377127 | 0.102655376540138 | 0.843284351171898 | -4.12547613263121 |
| **Trpc3** | -0.39786118 | 20.5772274925 | -1.88424274661361 | 0.102733957071376 | 0.843284351171898 | -4.12595768707853 |
| **Chl1** | 11.2737283475 | 16.69809267375 | 1.87965668251558 | 0.103421538003901 | 0.843284351171898 | -4.13015561688073 |
| **Pea15** | 0.565007207499995 | 21.73871649875 | 1.87843736741783 | 0.103605102301306 | 0.843284351171898 | -4.13127161136863 |
| **Mtnd5** | 10.366159335 | 15.3653463775 | 1.87700671991087 | 0.103820887345137 | 0.843284351171898 | -4.13258096164848 |
| **Ywhaq** | 0.398188830000002 | 24.530268215 | 1.87475108030431 | 0.104161996907405 | 0.843284351171898 | -4.13464520334298 |
| **Basp1** | 0.529099850000001 | 22.6318994275 | 1.87413391763136 | 0.104255517719252 | 0.843284351171898 | -4.13520996460956 |
| **Snca** | 0.230443282499998 | 25.11848140125 | 1.87372187975916 | 0.104318000824106 | 0.843284351171898 | -4.13558700955719 |
| **Slc8a1** | 11.00245126 | 16.337755035 | 1.87218851455967 | 0.104550847469984 | 0.843284351171898 | -4.13699009442488 |
| **Psma7** | 9.868889475 | 14.66727853 | 1.8697516421246 | 0.104921937368104 | 0.843284351171898 | -4.13921973370416 |

| | | | | | | |
|---|---|---|---|---|---|---|
| **Inpp5a** | 0.223200630000008 | 26.08884282 | 1.86854005120435 | 0.105106917314769 | 0.843284351171898 | -4.14032820309984 |
| **Lamp1** | 0.802434132500011 | 22.88243709625 | 1.86845480907204 | 0.105119943635971 | 0.843284351171898 | -4.14040618785906 |
| **Fundc2** | 10.2669096975 | 15.26280302375 | 1.8676180206395 | 0.10524790142943 | 0.843284351171898 | -4.14117171804552 |
| **Csnk2a1** | -0.145095135000002 | 28.76648592 | -1.86461274090767 | 0.105708706918221 | 0.843284351171898 | -4.1439208433662 |
| **Mtfp1** | 11.10216502 | 16.5323226425 | 1.86383808783672 | 0.105827804140261 | 0.843284351171898 | -4.14462940878883 |
| **Ndufa12** | 0.156512297500008 | 28.52405607375 | 1.86222801953888 | 0.106075758408948 | 0.843284351171898 | -4.14610203774513 |
| **Krt222** | 0.216482372500007 | 28.30728609875 | 1.8597940798201 | 0.106451664722075 | 0.843284351171898 | -4.14832800177955 |
| **Idi1** | 11.539435295 | 17.20825135 | 1.8597680330338 | 0.106455694483228 | 0.843284351171898 | -4.14835182154131 |
| **Nsfl1c** | 10.8947886525 | 16.24292579625 | 1.85848996584804 | 0.106653609782428 | 0.843284351171898 | -4.14952057665531 |
| **Zmpste24** | 9.9546035875 | 14.85763591875 | 1.85768794815497 | 0.106777989127355 | 0.843284351171898 | -4.15025396202823 |
| **Crym** | 12.1345708375 | 18.11494400875 | 1.8576533032744 | 0.106783365139217 | 0.843284351171898 | -4.15028564155208 |
| **Rexo2** | -10.7602256525 | 16.06415983625 | -1.85589563034178 | 0.107056457459449 | 0.843284351171898 | -4.15189280049541 |
| **Myo6** | -0.4117934575 | 21.81070455125 | -1.85522241993638 | 0.107161235002652 | 0.843284351171898 | -4.15250832577374 |
| **Bcap31** | -0.278737010000015 | 30.872314775 | -1.85515495564594 | 0.107171740548423 | 0.843284351171898 | -4.15257000815733 |
| **Nsdhl** | 11.427285245 | 17.0748393875 | 1.85319284413536 | 0.107477719247718 | 0.843284351171898 | -4.15436387051004 |
| **Ctsb** | 0.405296749999991 | 21.535360985 | 1.85078884194877 | 0.107853765860341 | 0.843284351171898 | -4.1565614920534 |
| **Gnas** | -0.516102954999997 | 23.7189630675 | -1.84852240658015 | 0.108209464468527 | 0.843284351171898 | -4.15863311055329 |
| **Mog** | -0.182082522500011 | 27.84008055375 | -1.84839811410206 | 0.10822900409753 | 0.843284351171898 | -4.15874671217563 |
| **Prrt2** | -0.77717954500001 | 21.621360865 | -1.84802251641762 | 0.108288071460477 | 0.843284351171898 | -4.15908999885954 |
| **Slc9a6** | 9.05389603 | 13.55208388 | 1.84780460612435 | 0.108322354893703 | 0.843284351171898 | -4.15928916022407 |

| | | | | | | |
|---|---|---|---|---|---|---|
| **Pafah1b2** | 0.327584272500012 | 24.72700043625 | 1.84696606774158 | 0.108454379009449 | 0.843284351171898 | -4.16005552990954 |
| **Uggt1** | 11.32622074 | 16.9684396275 | 1.84626659104809 | 0.108564628049772 | 0.843284351171898 | -4.16069478044786 |
| **Mtatp8** | -0.15475134499999 | 31.5391369925 | -1.84508615678859 | 0.108750930826744 | 0.843284351171898 | -4.1617735237112 |
| **Cds2** | 9.7228819575 | 14.58310792625 | 1.84408622858554 | 0.108908987987767 | 0.843284351171898 | -4.16268725797946 |
| **Ndufa2** | 10.37159565 | 15.56479051 | 1.84273276289877 | 0.109123283628164 | 0.843284351171898 | -4.16392397660405 |
| **Uqcrc2** | -0.157984185000011 | 28.4416504925 | -1.83432270739823 | 0.110464057156679 | 0.843284351171898 | -4.17160652874534 |
| **Mtcl3** | -10.534288925 | 15.8667694725 | -1.83276621064053 | 0.110713948013995 | 0.843284351171898 | -4.17302798025811 |
| **Gclm** | 9.8124595375 | 14.77844686625 | 1.83271381140381 | 0.110722370082342 | 0.843284351171898 | -4.17307583097823 |
| **Camkv** | -0.272481290000002 | 26.3484695625 | -1.83095146174233 | 0.111005992485994 | 0.843284351171898 | -4.17468511426291 |
| **Lonp1** | 0.787034775000002 | 22.6573129175 | 1.83030873364964 | 0.111109604582228 | 0.843284351171898 | -4.17527197751406 |
| **Hspa9** | 9.7836469775 | 14.74260359875 | 1.83028326730028 | 0.111113711859078 | 0.843284351171898 | -4.17529522991068 |
| **Dcps** | 11.0541666525 | 16.66402183375 | 1.83021625954708 | 0.111124519740939 | 0.843284351171898 | -4.17535641208302 |
| **Eno2** | 9.8383226725 | 14.84541802125 | 1.8272364808993 | 0.11160616842157 | 0.843284351171898 | -4.17807688409564 |
| **Hsph1** | 9.6308594625 | 14.54145696875 | 1.82567894904708 | 0.111858730662875 | 0.843284351171898 | -4.17949868118002 |
| **Hycc2** | -0.572556129999999 | 19.48380437 | -1.82529526443015 | 0.111921032134945 | 0.843284351171898 | -4.17984890791109 |
| **Ndufs6** | -9.3658863475 | 14.15384257875 | -1.82320037628218 | 0.112261786364105 | 0.843284351171898 | -4.18176097234494 |
| **Dlg2** | -0.435010669999997 | 19.918515245 | -1.82033777737446 | 0.112729040693189 | 0.843284351171898 | -4.18437334381674 |
| **Pgrmc2** | 9.6111197175 | 14.54526468125 | 1.81868679775285 | 0.112999380901733 | 0.843284351171898 | -4.18587979131494 |
| **Ndufs2** | 9.67053588 | 14.6457729775 | 1.81769673662229 | 0.113161799558449 | 0.843284351171898 | -4.18678310271422 |
| **Ptpa** | -9.26111051499999 | 14.0257696625 | -1.81738400981613 | 0.113213149034825 | 0.843284351171898 | -4.18706841620996 |

| | | | | | | |
|---|---|---|---|---|---|---|
| **Vapa** | 0.358268825000003 | 26.19950158 | 1.8158871978069 | 0.113459236573199 | 0.843284351171898 | -4.18843393891479 |
| **Exoc2** | -0.175204230000006 | 25.5676840975 | -1.81437367001516 | 0.113708598324032 | 0.843284351171898 | -4.18981457516607 |
| **Ran** | -0.398338047499998 | 21.34386373625 | -1.81419274888481 | 0.113738441479726 | 0.843284351171898 | -4.18997960175018 |
| **Cd9** | 0.352518387500005 | 26.24538407375 | 1.81377704161945 | 0.113807041582779 | 0.843284351171898 | -4.190358780206 |
| **Akr1a1** | 0.434251302499995 | 27.10431453625 | 1.81364121274058 | 0.113829464763639 | 0.843284351171898 | -4.19048267133534 |
| **Gstk1** | 0.230042435000009 | 24.8180214325 | 1.8115364791267 | 0.11417746904474 | 0.843284351171898 | -4.19240228035253 |
| **Taldo1** | 10.53995372 | 16.0219355725 | 1.8070044811989 | 0.114930300132195 | 0.843284351171898 | -4.19653473249074 |
| **Ermp1** | 0.243026925000002 | 20.14871002 | 1.80494065111671 | 0.115274719405774 | 0.843284351171898 | -4.19841618548056 |
| **Syt2** | -10.860107935 | 16.53234189 | -1.80242492963451 | 0.11569589992178 | 0.843284351171898 | -4.20070922831682 |
| **Tnr** | 10.3135549525 | 15.71931125625 | 1.80085571410952 | 0.115959368877725 | 0.843284351171898 | -4.20213933731411 |
| **Ptprs** | 0.374077725000006 | 23.29591108 | 1.79882313341322 | 0.116301496972109 | 0.843284351171898 | -4.20399149536179 |
| **Rps24** | -0.155457427500014 | 27.84123042625 | -1.79634665090516 | 0.116719659715358 | 0.843284351171898 | -4.20624778149525 |
| **Adam22** | 10.7799620875 | 16.41609130125 | 1.79576063638376 | 0.116818822286543 | 0.843284351171898 | -4.20678163038704 |
| **Ociad1** | -0.5143517825 | 24.38766775625 | -1.79567164568078 | 0.116833887970851 | 0.843284351171898 | -4.20686269732158 |
| **Timm50** | -0.345245792500002 | 26.78269304625 | -1.79202219418396 | 0.117453338214178 | 0.843284351171898 | -4.21018673613934 |
| **Apoe** | -10.5231753525 | 16.11428344625 | -1.78741259718533 | 0.118240286573993 | 0.843284351171898 | -4.21438399098157 |
| **Atcay** | 9.50561474 | 14.5980578025 | 1.78018513159474 | 0.119484387738012 | 0.843284351171898 | -4.22096189481784 |
| **Atp2a3** | -0.14640920250001 | 29.54310087125 | -1.7788501226618 | 0.119715563989232 | 0.843284351171898 | -4.22217650448934 |
| **Ppia** | 0.32912511 | 24.72378076 | 1.77712873213768 | 0.12001428293418 | 0.843284351171898 | -4.22374245418838 |
| **Psmc2** | 0.34007390750001 | 22.96616180375 | 1.77658978576439 | 0.120107955450944 | 0.843284351171898 | -4.22423268842261 |
| **Pgm1** | -1.10569543999999 | 20.6413420175 | -1.77629479896233 | 0.120159255914965 | 0.843284351171898 | -4.2245010038996 |

| | | | | | | |
|---|---|---|---|---|---|---|
| **Mecp2** | 0.593879427500013 | 24.53404373875 | 1.77473772898565 | 0.120430391590099 | 0.843284351171898 | -4.22591718221425 |
| **Psmd8** | 0.809334785000004 | 24.441829765 | 1.77148166040121 | 0.120999279256871 | 0.843284351171898 | -4.22887802906209 |
| **Cd81** | -9.3926083275 | 14.47691609125 | -1.76967469833856 | 0.12131609788133 | 0.843284351171898 | -4.23052080525826 |
| **Prdx2** | -10.328963815 | 15.70351939 | -1.76760402664055 | 0.121680133060205 | 0.843284351171898 | -4.23240301695395 |
| **Ktn1** | -0.238383790000004 | 19.9181547375 | -1.76587803453014 | 0.121984373108249 | 0.843284351171898 | -4.23397166204589 |
| **Stxbp5** | 1.3754903075 | 20.92886851875 | 1.76476460227111 | 0.122181024814592 | 0.843284351171898 | -4.23498346487176 |
| **Cisd1** | -1.068891485 | 22.0880071025 | -1.76231968523881 | 0.12261390833644 | 0.843284351171898 | -4.2372048720942 |
| **Hspa4** | 0.6490751575 | 24.06613048125 | 1.75802927790855 | 0.123377103247323 | 0.843284351171898 | -4.24110188706382 |
| **Slc6a11** | -0.350554304999992 | 25.7365778725 | -1.75191692757034 | 0.124472263508992 | 0.843284351171898 | -4.24665115677208 |
| **Hdhd2** | -0.217961334999998 | 26.8570957 | -1.75006182234725 | 0.124806484235435 | 0.843284351171898 | -4.24833474227163 |
| **Pdcd5** | 0.260701612500004 | 26.24540963125 | 1.74921623462588 | 0.124959112597723 | 0.843284351171898 | -4.24910205066398 |
| **Picalm** | -9.70311519 | 15.0853000425 | -1.74733099725531 | 0.125300041123501 | 0.843284351171898 | -4.25081254224111 |
| **Ube2d2** | 0.496578449999994 | 24.694856575 | 1.74373290456488 | 0.125953197216386 | 0.843284351171898 | -4.25407626218113 |
| **Dnajc5** | -0.440674032500002 | 25.67752694625 | -1.74319619209907 | 0.126050904579688 | 0.843284351171898 | -4.254563000066 |
| **Pi4ka** | -0.192614747499995 | 22.08649931125 | -1.7422699510114 | 0.12621969536779 | 0.843284351171898 | -4.25540293657213 |
| **Hspa12a** | 0.371922399999999 | 22.349087065 | 1.74020550690776 | 0.126596681426455 | 0.843284351171898 | -4.25727474634426 |
| **Rap1b** | 0.371922399999999 | 22.349087065 | 1.74020550690776 | 0.126596681426455 | 0.843284351171898 | -4.25727474634426 |
| **Ndufv3** | -0.706882707499997 | 22.88598820125 | -1.72560034713245 | 0.129294612362172 | 0.843284351171898 | -4.27050594734503 |
| **Atic** | 0.190972522500005 | 24.46758887875 | 1.72553049275672 | 0.129307647114365 | 0.843284351171898 | -4.27056918216873 |
| **Nipsnap2** | 0.215292252500006 | 27.15871959625 | 1.72333372283734 | 0.129718202170413 | 0.843284351171898 | -4.27255754193551 |

| | | | | | | |
|---|---|---|---|---|---|---|
| **Ndufs1** | 7.3568592925 | 17.05563837625 | 1.71803197827195 | 0.13071417267567 | 0.843284351171898 | -4.27735437080907 |
| **Cox7a2** | -0.211830307500001 | 24.31263295125 | -1.71774923485872 | 0.130767492227874 | 0.843284351171898 | -4.27761010931024 |
| **D11Wsu47e** | 0.483845347499994 | 25.03467226125 | 1.71753972518162 | 0.130807014780185 | 0.843284351171898 | -4.27779960354041 |
| **Hspd1** | 0.196234930000003 | 20.9604674425 | 1.71442402248541 | 0.131396115130235 | 0.843284351171898 | -4.28061713232379 |
| **Gstp1** | 0.372300287499996 | 21.31798458375 | 1.712792180663 | 0.131705662368477 | 0.843284351171898 | -4.28209241762164 |
| **Lingo1** | 0.306929705000002 | 26.35064096 | 1.71077817148182 | 0.132088660883342 | 0.843284351171898 | -4.28391283385756 |
| **Thop1** | -0.250784722499997 | 26.60848992875 | -1.70938731279935 | 0.132353775134449 | 0.843284351171898 | -4.28516975730327 |
| **Nceh1** | 0.743834602500005 | 23.41940645125 | 1.70894473633262 | 0.132438241623593 | 0.843284351171898 | -4.28556967347576 |
| **Epn2** | -0.226196267500001 | 23.92974389125 | -1.70818444175331 | 0.132583464907923 | 0.843284351171898 | -4.28625663561149 |
| **Grm5** | 10.4254915575 | 16.48720998375 | 1.70542161571462 | 0.133112466550434 | 0.843284351171898 | -4.28875247620373 |
| **Pitpna** | 0.431616605000002 | 22.8380947475 | 1.70384754505767 | 0.133414752641581 | 0.843284351171898 | -4.29017408111532 |
| **Spata20** | 0.2759819925 | 23.19263899125 | 1.70115839500354 | 0.133932689897038 | 0.843284351171898 | -4.29260215511456 |
| **Txndc12** | 0.295722899999998 | 24.80462911 | 1.70072899649169 | 0.134015569873497 | 0.843284351171898 | -4.29298979462698 |
| **Dlg4** | 0.472029692499998 | 24.59097186875 | 1.6976068415229 | 0.13461965659914 | 0.843284351171898 | -4.29580772842506 |
| **Asah1** | 0.3056819275 | 18.78388086375 | 1.69749383352948 | 0.134641570252999 | 0.843284351171898 | -4.29590970537573 |
| **Cnot11** | 0.450111847500004 | 24.58099562375 | 1.69582585309545 | 0.134965406593198 | 0.843284351171898 | -4.29741470902138 |
| **Rab4b** | 0.510927555000002 | 24.0525509125 | 1.69185441850756 | 0.135739434220422 | 0.843284351171898 | -4.30099688070513 |
| **Uba1** | -0.290878324999994 | 29.0593736025 | -1.69022374939794 | 0.136058467791399 | 0.843284351171898 | -4.30246721720583 |
| **Itm2b** | 0.210242342499996 | 25.21374879375 | 1.68779001430867 | 0.136535940917541 | 0.843284351171898 | -4.30466111101488 |
| **Nefh** | 0.282581839999999 | 25.4149633375 | 1.68131893263988 | 0.137813236735824 | 0.843284351171898 | -4.31049123674825 |

| | | | | | | |
|---|---|---|---|---|---|---|
| **Kcnj3** | 0.6837523875 | 24.77213522125 | 1.6811942573888 | 0.137837956584244 | 0.843284351171898 | -4.3106035162416 |
| **Psmb5** | 0.575123474999998 | 23.5807896725 | 1.67955035105054 | 0.138164293242173 | 0.843284351171898 | -4.31208381117287 |
| **Gng3** | 0.402421239999995 | 25.0993117775 | 1.67102062666119 | 0.139869326928149 | 0.843284351171898 | -4.31975955538418 |
| **Slc44a1** | 0.244425957499992 | 24.80336376875 | 1.67090572676591 | 0.139892429903598 | 0.843284351171898 | -4.31986289316319 |
| **Chid1** | 0.517205792499997 | 23.76437141375 | 1.66897247808478 | 0.140281690475501 | 0.843284351171898 | -4.32160136695752 |
| **Acot2** | -0.249937120000002 | 25.599199275 | -1.66788088612507 | 0.14050193501589 | 0.843284351171898 | -4.32258278305216 |
| **Blmh** | 0.412441015000006 | 24.895099365 | 1.66724586107884 | 0.14063021083104 | 0.843284351171898 | -4.32315364813729 |
| **Prdx6** | -0.239370307499993 | 21.26602194375 | -1.66270784180398 | 0.141550121214589 | 0.843284351171898 | -4.32723174299495 |
| **Cnpy2** | 0.161451602500005 | 26.94452669625 | 1.66197120883812 | 0.141699980418912 | 0.843284351171898 | -4.32789348143841 |
| **Gprc5b** | 1.30609012250001 | 25.14013207875 | 1.65400916640814 | 0.143329345260139 | 0.843284351171898 | -4.33504171303943 |
| **Adam23** | 0.154930745000009 | 23.51120463 | 1.65221151003051 | 0.143699657146701 | 0.843284351171898 | -4.3366545261624 |
| **Plppr4** | 0.160664167499998 | 24.79183130125 | 1.65195617635303 | 0.143752328197108 | 0.843284351171898 | -4.33688357193831 |
| **Arpc5l** | 0.190503774999996 | 20.2394824325 | 1.64792537563565 | 0.144586228282963 | 0.843284351171898 | -4.34049827514843 |
| **Mecr** | -1.00638461 | 22.4452320625 | -1.64647293822269 | 0.144887826355138 | 0.843284351171898 | -4.34180026494613 |
| **Cnnm1** | 0.498165004999997 | 25.4685263525 | 1.64395327676916 | 0.145412438284097 | 0.843284351171898 | -4.3440582817644 |
| **Diras2** | 1.1036953625 | 23.08873796875 | 1.64334301775927 | 0.145539767292978 | 0.843284351171898 | -4.34460504588278 |
| **Hspa8** | -0.210052294999993 | 25.3583767 | -1.64099781105938 | 0.146030065484699 | 0.843284351171898 | -4.34670578771193 |
| **Rps5** | 1.124993965 | 21.758322995 | 1.62837780386141 | 0.148695217737287 | 0.843284351171898 | -4.35799763996157 |
| **Hpca** | -1.0889717875 | 21.91604375625 | -1.62738979726778 | 0.148905785840429 | 0.843284351171898 | -4.35888075072794 |
| **Ndufs5** | 0.370419120000001 | 25.60156904 | 1.62732934576883 | 0.14891867859488 | 0.843284351171898 | -4.35893477975904 |

| | | | | | | |
|---|---|---|---|---|---|---|
| **F3** | -0.167288245000002 | 19.67883578 | -1.62577731407055 | 0.149250045930089 | 0.843284351171898 | -4.36032174746647 |
| **Slc6a9** | 0.453603735000005 | 22.6006871925 | 1.62479142213621 | 0.14946089833696 | 0.843284351171898 | -4.36120261233704 |
| **Glod4** | -0.885391955000003 | 22.731325605 | -1.62411524382824 | 0.149605673835895 | 0.843284351171898 | -4.3618066789406 |
| **Nptx1** | 0.598499667500004 | 24.60413859125 | 1.62342036404302 | 0.149754590460422 | 0.843284351171898 | -4.36242738592611 |
| **Tpm4** | 8.6352674 | 21.27676513 | 1.62171557742587 | 0.150120524600839 | 0.843284351171898 | -4.36394991318153 |
| **Rab5c** | -0.328295139999998 | 22.952072815 | -1.61392310674759 | 0.151803877248191 | 0.843284351171898 | -4.37090403593294 |
| **Igsf8** | -0.349100715000002 | 23.6014293825 | -1.61015225147696 | 0.152624797328273 | 0.843284351171898 | -4.37426606799469 |
| **Dld** | 0.557701892499995 | 22.53140805875 | 1.6077167860207 | 0.153157206071292 | 0.843284351171898 | -4.37643638471656 |
| **Rps6** | -0.475489007499991 | 24.27738358875 | -1.60202339110176 | 0.154408598616792 | 0.843284351171898 | -4.38150651866879 |
| **Sptb** | 0.243186534999992 | 32.0450961825 | 1.5992291449457 | 0.155026253504305 | 0.843284351171898 | -4.38399310285991 |
| **Sez6l** | 0.214428679999994 | 22.5449111225 | 1.59709193770406 | 0.155500228556274 | 0.843284351171898 | -4.38589419510531 |
| **Atp5f1c** | 0.335347672500003 | 22.87186957375 | 1.59686316245858 | 0.155551044779463 | 0.843284351171898 | -4.38609765452559 |
| **Ndufb4** | 0.263416004999993 | 25.4431011225 | 1.59641234736265 | 0.155651226482935 | 0.843284351171898 | -4.38649855995181 |
| **Ndrg1** | -11.2711552941667 | 16.0044244885714 | -1.5875360818239 | 0.165141277530699 | 0.843284351171898 | -4.38789369541213 |
| **Myl12b** | 0.422527605000006 | 25.01564193 | 1.59076281242989 | 0.156911797806719 | 0.843284351171898 | -4.39151999283305 |
| **Tmem33** | 0.541747429999997 | 27.4613715725 | 1.59034090047477 | 0.157006319139368 | 0.843284351171898 | -4.3918947998799 |
| **Slc24a2** | -0.523899210000003 | 21.4714130225 | -1.58439423993291 | 0.158344208820698 | 0.843284351171898 | -4.39717457941377 |
| **Got2** | -0.222363324999989 | 27.2680977225 | -1.58366581262048 | 0.158508819283677 | 0.843284351171898 | -4.39782093564091 |
| **Stx12** | 9.78878551583334 | 13.99504514 | 1.57479582978132 | 0.168020509779419 | 0.843284351171898 | -4.39817925891223 |
| **Lmnb1** | 0.169231619999998 | 29.2342060025 | 1.57911005203093 | 0.159541949617334 | 0.843284351171898 | -4.40186148913204 |

| | | | | | | |
|---|---|---|---|---|---|---|
| **Grid2** | -0.132985085000005 | 27.737082415 | -1.57801445533259 | 0.15979133525632 | 0.843284351171898 | -4.40283269021284 |
| **Ndufb3** | 0.309343722500003 | 27.01967249875 | 1.57695037592763 | 0.160033893569154 | 0.843284351171898 | -4.40377576740335 |
| **Plpbp** | -0.317793842499999 | 21.69585246375 | -1.57111672408155 | 0.161369771557752 | 0.843284351171898 | -4.4089427767706 |
| **Tmco1** | 1.640021615 | 22.565709255 | 1.56647753098315 | 0.16243950575879 | 0.843284351171898 | -4.41304783679433 |
| **Rhog** | -0.173980610000001 | 23.42605321 | -1.55987668907005 | 0.163972907067452 | 0.843284351171898 | -4.41888250249398 |
| **Slc25a22** | 0.238306787499994 | 19.53834050625 | 1.55477518286773 | 0.165167180483545 | 0.843284351171898 | -4.42338680751922 |
| **Rab22a** | -0.491600822499997 | 23.58995685625 | -1.5537490581474 | 0.165408368808028 | 0.843284351171898 | -4.42429227139624 |
| **Cplx1** | 0.245355562499991 | 26.75111828125 | 1.55357815564736 | 0.165448570697166 | 0.843284351171898 | -4.42444306002988 |
| **Nfasc** | 0.555762502500009 | 24.25289827375 | 1.55231521882844 | 0.165745935260513 | 0.843284351171898 | -4.42555720276567 |
| **B3gat3** | -0.277708682499998 | 26.81886229625 | -1.55105198592027 | 0.166043863914412 | 0.843284351171898 | -4.42667133031622 |
| **Sh3gl2** | 0.265647795 | 25.52804383 | 1.54931884952941 | 0.166453423183053 | 0.843284351171898 | -4.42819944487874 |
| **Rab3c** | 0.597275682499998 | 23.61586500875 | 1.54862674545508 | 0.166617235751021 | 0.843284351171898 | -4.42880952988053 |
| **Atp5f1b** | 0.282248290000005 | 26.0715635575 | 1.54814826227905 | 0.166730573954118 | 0.843284351171898 | -4.42923126052931 |
| **Fkbp1a** | -0.299752282500013 | 25.98421569375 | -1.54402245633639 | 0.167710811594132 | 0.843284351171898 | -4.4328660380831 |
| **Col7a1** | 0.197447874999991 | 28.90863757 | 1.5421577971868 | 0.168155573914153 | 0.843284351171898 | -4.43450778927148 |
| **Sema4b** | 0.594901902500002 | 20.11595015625 | 1.54138597662339 | 0.168339988495004 | 0.843284351171898 | -4.435187162671 |
| **Gnb5** | 0.430483302499997 | 21.54978609875 | 1.53936328615839 | 0.16882416477934 | 0.843284351171898 | -4.43696707433734 |
| **Romo1** | 0.45703228 | 20.2045815925 | 1.53704177403284 | 0.169381453677367 | 0.843284351171898 | -4.43900903587601 |
| **Mapk3** | 0.615536147499995 | 24.65929254125 | 1.53318906378081 | 0.170310057047815 | 0.843284351171898 | -4.44239566253165 |
| **Eno1** | -0.6877445175 | 22.26673921875 | -1.53251339780341 | 0.170473392762523 | 0.843284351171898 | -4.44298931124421 |
| **Bcat1** | 0.4798983875 | 22.47377499875 | 1.53129800923149 | 0.170767564433053 | 0.843284351171898 | -4.44405695702745 |

| | | | | | | |
|---|---|---|---|---|---|---|
| **Gap43** | 0.207930117499998 | 19.96804173375 | 1.53007118051315 | 0.171064979077562 | 0.843284351171898 | -4.44513437770575 |
| **Ndufb10** | 0.651121724999996 | 23.691384395 | 1.52889234180921 | 0.171351208902495 | 0.843284351171898 | -4.44616939206115 |
| **Plp1** | -0.575717344999994 | 25.037152495 | -1.52786145378383 | 0.171601876550171 | 0.843284351171898 | -4.44707429641799 |
| **Opa3** | -0.608757674999996 | 23.1537758025 | -1.52385460574881 | 0.172579377121144 | 0.843284351171898 | -4.45058960058444 |
| **Hsd17b11** | 0.441524950000002 | 20.0158372925 | 1.52317679907942 | 0.172745238619178 | 0.843284351171898 | -4.45118396082352 |
| **Ndufa7** | -0.217590169999994 | 29.242778105 | -1.52127816862666 | 0.17321062007822 | 0.843284351171898 | -4.45284838787237 |
| **Slc30a3** | 0.343114462499997 | 23.07177369625 | 1.51903772523088 | 0.173761266547 | 0.843284351171898 | -4.4548115913317 |
| **Ndufv1** | 0.266373072500002 | 22.76317526375 | 1.51406401748977 | 0.174989433285632 | 0.843284351171898 | -4.45916643314588 |
| **Dnajc3** | 0.26655718 | 21.99334042 | 1.51190268251451 | 0.175525614660176 | 0.843284351171898 | -4.46105736446605 |
| **Cand1** | 0.264885260000007 | 22.6555248125 | 1.51143096081805 | 0.17564283896641 | 0.843284351171898 | -4.46146994976812 |
| **Ndufb5** | -0.293418342500011 | 24.13467034125 | -1.50377300129183 | 0.177555931586921 | 0.843284351171898 | -4.4681618432606 |
| **Ndufa4** | -0.173006699999995 | 25.09667768 | -1.5035803052174 | 0.177604315695659 | 0.843284351171898 | -4.46833008251601 |
| **Edil3** | 0.223579415000003 | 27.840137065 | 1.50321217902614 | 0.177696782157218 | 0.843284351171898 | -4.46865146613416 |
| **Rpl10** | 0.205456084999994 | 24.152701555 | 1.5009696703437 | 0.178261011139167 | 0.843284351171898 | -4.47060865637339 |
| **Hint1** | 0.188517327499998 | 25.88666144625 | 1.49964387396743 | 0.178595360300626 | 0.843284351171898 | -4.47176530113783 |
| **Asrgl1** | 0.349633645000004 | 20.4629638675 | 1.49891059519427 | 0.178780530270128 | 0.843284351171898 | -4.47240487466371 |
| **Fahd1** | -0.684003490000002 | 25.555821015 | -1.49807275281715 | 0.178992319906875 | 0.843284351171898 | -4.47313551863905 |
| **Gabarapl2** | -0.389887392499993 | 21.52379610125 | -1.49621560368678 | 0.17946258777437 | 0.843284351171898 | -4.47475455373575 |
| **Faim2** | 0.285072722499997 | 24.87546454125 | 1.49331125399167 | 0.180200292465904 | 0.843284351171898 | -4.47728513591127 |
| **Grm1** | 0.148240160000004 | 26.04457401 | 1.49200389031491 | 0.180533266115077 | 0.843284351171898 | -4.47842369696796 |

| | | | | | | |
|---|---|---|---|---|---|---|
| **Cnp** | -0.197699802499997 | 25.11535589875 | -1.49116598947557 | 0.180746967212094 | 0.843284351171898 | -4.4791532288181 |
| **Tspan2** | -0.216928827499999 | 27.79604389875 | -1.4907841375867 | 0.180844432619694 | 0.843284351171898 | -4.47948564707749 |
| **Gcsh** | 0.888747074999991 | 23.915775285 | 1.48537444664352 | 0.182230386947791 | 0.843284351171898 | -4.48419181807511 |
| **Sorl1** | 0.478921749999998 | 24.364988245 | 1.48429282845225 | 0.182508655387844 | 0.843284351171898 | -4.48513205405925 |
| **Ndufa3** | 0.151652155000001 | 21.0959686425 | 1.48287088477046 | 0.182875068928797 | 0.843284351171898 | -4.48636776305618 |
| **Eef1b** | -0.211764777499997 | 24.31042700625 | -1.48158430717567 | 0.183207178292805 | 0.843284351171898 | -4.48748547447882 |
| **Pcmt1** | -0.617031307500003 | 21.64875191625 | -1.4810554098909 | 0.183343863966467 | 0.843284351171898 | -4.48794485318695 |
| **Tomm22** | 0.1560818375 | 25.41055180375 | 1.48016868863289 | 0.18357323240948 | 0.843284351171898 | -4.48871489274236 |
| **Alg2** | 0.463546050000005 | 21.6595077425 | 1.48005926430074 | 0.183601555343635 | 0.843284351171898 | -4.48880990681858 |
| **Hadh** | 0.287739165000012 | 24.9006152575 | 1.47847611141526 | 0.184011777554724 | 0.843284351171898 | -4.49018429235237 |
| **Hnrnpm** | 0.413211420000007 | 25.5725458175 | 1.47808154331036 | 0.18411414679704 | 0.843284351171898 | -4.49052674796847 |
| **Cdh10** | 0.394687832500001 | 18.02539941875 | 1.47746737874262 | 0.184273592700726 | 0.843284351171898 | -4.49105973202328 |
| **Fasn** | -0.393463022500001 | 23.29976159875 | -1.47430593795204 | 0.185096338232163 | 0.843284351171898 | -4.49380203828665 |
| **Coa3** | 0.405102767500011 | 23.48997015625 | 1.46903314076115 | 0.186475987548279 | 0.843284351171898 | -4.49837107773668 |
| **Rps14** | 0.195728642500008 | 24.78468286125 | 1.46771786958667 | 0.186821587013283 | 0.843284351171898 | -4.49950987635197 |
| **Cdh2** | -0.322545444999999 | 26.04865981 | -1.4668235443008 | 0.187056911310732 | 0.843284351171898 | -4.500283996287 |
| **Aldh18a1** | 6.834663915 | 18.3023407875 | 1.46662732050334 | 0.187108579784912 | 0.843284351171898 | -4.50045382284946 |
| **Txndc5** | 0.208087769999992 | 23.1119892 | 1.46382158005597 | 0.18784878920363 | 0.843284351171898 | -4.50288120999165 |
| **Gpx1** | -0.172313047499998 | 27.90317369625 | -1.46281938503388 | 0.18811383150742 | 0.843284351171898 | -4.50374784669878 |
| **Slc25a5** | 0.157737687500013 | 26.39519076625 | 1.46049907482188 | 0.188728767465702 | 0.843284351171898 | -4.50575347102469 |

| | | | | | | |
|---|---|---|---|---|---|---|
| **Ndufv2** | -0.340150335000004 | 23.2495000325 | -1.4594426112497 | 0.189009358339953 | 0.843284351171898 | -4.50666626570325 |
| **Sucla2** | 0.217542552500007 | 25.50876664125 | 1.45846392946746 | 0.189269628224496 | 0.843284351171898 | -4.50751163834442 |
| **Ndufs8** | 0.646265925000005 | 20.49795846 | 1.45734166732048 | 0.189568481581935 | 0.843284351171898 | -4.50848077536285 |
| **Atp1a3** | 0.405401332500006 | 23.52785350625 | 1.45413702350728 | 0.190424218937326 | 0.843284351171898 | -4.51124664075502 |
| **Ak1** | 1.1863351875 | 21.97152193875 | 1.45360548314083 | 0.190566494184109 | 0.843284351171898 | -4.51170518343639 |
| **Stx7** | 0.531409562500002 | 22.86257010875 | 1.45260775682428 | 0.190833811434841 | 0.843284351171898 | -4.5125657204755 |
| **Cct4** | -0.462614920000004 | 23.7568162025 | -1.45072553548868 | 0.191339032613375 | 0.843284351171898 | -4.51418853015866 |
| **Prkacb** | 0.671298077499998 | 22.75449592375 | 1.44716108915036 | 0.19229910943712 | 0.843284351171898 | -4.51725954825006 |
| **Tmem35a** | 0.399730627499999 | 24.40982808875 | 1.44699847306571 | 0.19234301349508 | 0.843284351171898 | -4.51739958530674 |
| **Cul3** | 0.847880437499999 | 21.50476352125 | 1.44269146365505 | 0.193509148100035 | 0.843284351171898 | -4.52110639937938 |
| **Tm9sf4** | 0.147267347499994 | 27.72083089875 | 1.44213800415963 | 0.193659460971104 | 0.843284351171898 | -4.52158242778204 |
| **Dgke** | -0.197864900000003 | 23.5666183525 | -1.4404861798604 | 0.194108703472041 | 0.843284351171898 | -4.52300274085595 |
| **Letm1** | 0.201530820000006 | 29.471346355 | 1.43707634910791 | 0.195039042775707 | 0.843284351171898 | -4.52593269407388 |
| **Psma5** | 0.302236512499992 | 26.05051057375 | 1.43657158468099 | 0.195177104160633 | 0.843284351171898 | -4.52636619438839 |
| **Micos10** | -0.388621995000001 | 22.4209601625 | -1.43570772449297 | 0.195413588656259 | 0.843284351171898 | -4.52710795607455 |
| **Uqcr10** | 0.316610472499995 | 23.30940678375 | 1.42686359974406 | 0.19784959238588 | 0.843284351171898 | -4.53469208123573 |
| **Dynll2** | 0.229526399999994 | 22.1704794725 | 1.42655846161372 | 0.197934124676605 | 0.843284351171898 | -4.53495342068874 |
| **Vps35** | 0.444264437499996 | 22.34648583125 | 1.42549576489407 | 0.198228776814874 | 0.843284351171898 | -4.53586340964714 |
| **Nfu1** | 0.459245472499997 | 25.90256927625 | 1.42433669647991 | 0.198550599563878 | 0.843284351171898 | -4.53685561793262 |
| **Lrrc59** | -0.179981095000002 | 27.730520605 | -1.42258084114736 | 0.199039019021665 | 0.843284351171898 | -4.53835809358637 |

| | | | | | | |
|---|---|---|---|---|---|---|
| **Slc7a14** | -0.248816127500003 | 25.23487423625 | -1.42155009009411 | 0.199326241892912 | 0.843284351171898 | -4.53923976060675 |
| **Sparcl1** | -1.019121435 | 21.598411565 | -1.42011975815508 | 0.199725426600269 | 0.843284351171898 | -4.540462795504 |
| **Folh1** | 0.163017005 | 27.211958865 | 1.41837333599803 | 0.200213801134657 | 0.843284351171898 | -4.54195544734719 |
| **Pfkm** | 0.312007299999998 | 28.339477305 | 1.4181230304632 | 0.200283885098916 | 0.843284351171898 | -4.5421693214812 |
| **Rpl27** | 0.24412852999999 | 19.27315007 | 1.41773162468015 | 0.2003935203942 | 0.843284351171898 | -4.54250372892361 |
| **Ndufb6** | -0.517586099999999 | 23.7677931425 | -1.4157338322287 | 0.200953954753654 | 0.843284351171898 | -4.54421002002571 |
| **Slc44a2** | 0.894755692500002 | 26.67569518125 | 1.41399882502199 | 0.201441811590793 | 0.843284351171898 | -4.54569108891907 |
| **Prdx1** | 0.267058347499997 | 23.40646958625 | 1.41099058354717 | 0.202290200232048 | 0.843284351171898 | -4.54825731187577 |
| **Etfa** | -0.621045405 | 26.045948385 | -1.40856433496872 | 0.202976786279563 | 0.843284351171898 | -4.55032545289927 |
| **Nol3** | 0.290967337500003 | 22.39414667875 | 1.40785628730243 | 0.203177544598234 | 0.843284351171898 | -4.55092872343972 |
| **Atp5pb** | 0.699609700000003 | 22.718605225 | 1.40603876245612 | 0.203693696102784 | 0.843284351171898 | -4.5524767273294 |
| **Mcts1** | 0.29351923250001 | 23.78036597625 | 1.40508589670051 | 0.203964765399801 | 0.843284351171898 | -4.55328796793192 |
| **Dpysl5** | -0.231599049999996 | 29.898924165 | -1.40504423563114 | 0.203976624419665 | 0.843284351171898 | -4.55332343178063 |
| **Rps12** | 0.444596492499997 | 22.50374109375 | 1.40461599101985 | 0.204098562049489 | 0.843284351171898 | -4.5536879487847 |
| **Crmp1** | -0.492942039999999 | 23.57517347 | -1.40223637167097 | 0.204777319782441 | 0.843284351171898 | -4.55571262839005 |
| **Hras** | -0.183790267499997 | 24.24893255125 | -1.40087302596923 | 0.205167106888486 | 0.843284351171898 | -4.55687198747337 |
| **Glg1** | -0.209138272499999 | 27.58295010125 | -1.39092747435066 | 0.208030705118509 | 0.843284351171898 | -4.56531539218546 |
| **Ddah1** | 0.601647152500007 | 22.27688665875 | 1.38869544296596 | 0.208678248630822 | 0.843284351171898 | -4.56720687120115 |
| **Twf1** | -0.263817490000001 | 22.4904528025 | -1.3870852146 | 0.209146513538989 | 0.843284351171898 | -4.568570630008 |
| **Pde10a** | 0.170574637499996 | 24.95102304375 | 1.38199276641276 | 0.210633598264228 | 0.843284351171898 | -4.5728792241331 |
| **Atp5f1d** | -0.396735637500001 | 23.89309822125 | -1.3806374594097 | 0.211030954381799 | 0.843284351171898 | -4.57402478861064 |

| | | | | | | |
|---|---|---|---|---|---|---|
| **Gjc3** | -0.658287699999995 | 21.223491885 | -1.38038223481492 | 0.211105857064599 | 0.843284351171898 | -4.5742404622879 |
| **Adcy5** | 0.824033235000002 | 21.695770095 | 1.38037561362439 | 0.21110780054992 | 0.843284351171898 | -4.57424605719976 |
| **Tmx2** | 0.229900182499996 | 22.50303879375 | 1.3760526750905 | 0.2123800929575 | 0.843284351171898 | -4.57789650877134 |
| **Cs** | 0.598459375000001 | 23.2541461475 | 1.37345124776081 | 0.213149004158065 | 0.843284351171898 | -4.58009089688278 |
| **Epb41l1** | 0.282716255000004 | 24.5429797225 | 1.37306282020485 | 0.213264024682032 | 0.843284351171898 | -4.58041839552997 |
| **Pld3** | 0.284543169999992 | 26.303935575 | 1.37092697679729 | 0.213897472191835 | 0.843284351171898 | -4.58221849895802 |
| **Atp6v0d1** | 0.600827067499999 | 21.81298875625 | 1.36675196991802 | 0.215140516885203 | 0.843284351171898 | -4.58573373464245 |
| **Rab14** | 0.148308644999993 | 26.5162244575 | 1.36344491003888 | 0.2161296847602 | 0.843284351171898 | -4.58851488669302 |
| **Gsn** | 6.784081925 | 19.15537756 | 1.36058142660529 | 0.216989427309752 | 0.843284351171898 | -4.59092063173211 |
| **Cox5a** | 0.226884637500003 | 28.23294062625 | 1.35947320370054 | 0.217322975799587 | 0.843284351171898 | -4.5918511077346 |
| **Dnm1l** | 0.177794349999999 | 25.64030874 | 1.35813477424078 | 0.217726415663692 | 0.843284351171898 | -4.59297442465956 |
| **Dpp10** | 0.274615862500003 | 23.85529097625 | 1.3579570100196 | 0.217780048551728 | 0.843284351171898 | -4.59312358207038 |
| **Cox6b1** | 0.131528830000001 | 24.0817131625 | 1.3560336728141 | 0.218361082258169 | 0.843284351171898 | -4.5947368562584 |
| **Gnaq** | 0.239654225000002 | 23.02785739 | 1.35389430492022 | 0.219008987606796 | 0.843284351171898 | -4.59653015086641 |
| **Ank1** | -0.3287494275 | 25.15061055625 | -1.35204742152674 | 0.219569679783253 | 0.843284351171898 | -4.59807726796904 |
| **Crocc** | -0.610523314999991 | 26.42776704 | -1.35041056502036 | 0.220067669457376 | 0.843284351171898 | -4.59944766502239 |
| **Decr1** | -0.703316790000006 | 23.6991298225 | -1.34900057872837 | 0.220497436190816 | 0.843284351171898 | -4.60062753209166 |
| **Try5** | -0.302798850000006 | 24.5962780375 | -1.34872977401449 | 0.220580062756719 | 0.843284351171898 | -4.60085407686997 |
| **Ppp1r1b** | 0.17476882750001 | 26.03289406625 | 1.34700257902276 | 0.221107698740264 | 0.843284351171898 | -4.60229850448295 |
| **Ddx6** | 0.236299227500002 | 26.26430085625 | 1.34607602047449 | 0.221391209014097 | 0.843284351171898 | -4.60307303110525 |

| | | | | | | |
|---|---|---|---|---|---|---|
| **Dnm1** | -0.126178237500003 | 26.03244460875 | -1.3431979137681 | 0.222273902756912 | 0.843284351171898 | -4.60547736910143 |
| **Hint2** | 0.573521015000011 | 22.81807528 | 1.34274654832649 | 0.222412614092421 | 0.843284351171898 | -4.60585422525163 |
| **L1cam** | 0.296534812499996 | 23.14917483625 | 1.34149890135339 | 0.222796431375546 | 0.843284351171898 | -4.60689562018444 |
| **Sema4d** | 0.364681135000001 | 23.5044207025 | 1.34021034234991 | 0.223193446695424 | 0.843284351171898 | -4.60797070614664 |
| **Hcn1** | 0.511350262500002 | 23.61173981875 | 1.33713044210706 | 0.224144912440544 | 0.843284351171898 | -4.61053847421031 |
| **Fbxo2** | 0.245644379999998 | 20.876681695 | 1.33656177643451 | 0.224320978589227 | 0.843284351171898 | -4.61101228842857 |
| **Gaa** | -0.145594479999993 | 27.824945845 | -1.33519750909254 | 0.224743869122391 | 0.843284351171898 | -4.61214862772122 |
| **Sv2c** | 0.160225957500007 | 28.40177745125 | 1.33420850693958 | 0.225050874393254 | 0.843284351171898 | -4.61297206753824 |
| **Cyfip2** | 6.542679775 | 18.8956358025 | 1.33321513210611 | 0.225359607990219 | 0.843284351171898 | -4.61379886838173 |
| **Ola1** | 0.498504722500009 | 23.52699279875 | 1.33201378946429 | 0.225733473540169 | 0.843284351171898 | -4.61479838878951 |
| **Oxr1** | 0.188819245000005 | 21.216987715 | 1.33172751663502 | 0.225822643824796 | 0.843284351171898 | -4.61503650788391 |
| **Vat1l** | -0.783309712500003 | 19.38336641625 | -1.33100031198845 | 0.226049297786619 | 0.843284351171898 | -4.61564128480757 |
| **Cct3** | -0.609841265000004 | 21.122065655 | -1.33071016108292 | 0.226139787287364 | 0.843284351171898 | -4.61588254552041 |
| **Ndufa13** | 0.438299174999994 | 20.7028649075 | 1.3305850620655 | 0.226178811789132 | 0.843284351171898 | -4.61598655802956 |
| **Clec2l** | -0.240088794999998 | 27.508792185 | -1.32562303569499 | 0.22773148484211 | 0.843284351171898 | -4.62010857224708 |
| **Phb1** | 6.3394763775 | 17.88730008125 | 1.32376792585876 | 0.22831436548025 | 0.843284351171898 | -4.62164781239491 |
| **Emc4** | 0.117707725000013 | 25.457395465 | 1.32163817592316 | 0.228985150492306 | 0.843284351171898 | -4.62341370038928 |
| **Rad18** | 0.48266392 | 23.346479565 | 1.31333012638517 | 0.231618369340329 | 0.843284351171898 | -4.6302896858344 |
| **Atp5f1a** | -0.179722674999997 | 23.6103476125 | -1.31171687252459 | 0.232132745040673 | 0.843284351171898 | -4.63162250861124 |
| **Pdhb** | 0.288874569999997 | 21.671289385 | 1.31128239394186 | 0.232271446204259 | 0.843284351171898 | -4.63198133035589 |

| | | | | | | |
|---|---|---|---|---|---|---|
| **Canx** | -0.685608874999989 | 24.7687058575 | -1.30972485168853 | 0.232769264526106 | 0.843284351171898 | -4.63326719377605 |
| **Ppt1** | -0.142918007500008 | 27.63449325375 | -1.30882258365357 | 0.233058071365096 | 0.843284351171898 | -4.63401175137105 |
| **Lrpprc** | 0.1549037375 | 26.74470226875 | 1.30641898096384 | 0.233828966704321 | 0.843284351171898 | -4.63599403637016 |
| **Atp1b2** | 0.485019795 | 21.1691227175 | 1.30614717301583 | 0.23391628211992 | 0.843284351171898 | -4.63621809162568 |
| **Pfdn5** | 0.360731947499993 | 21.77970461875 | 1.30556526525154 | 0.234103309405989 | 0.843284351171898 | -4.63669769235374 |
| **Itgb8** | 0.176393392500003 | 25.55464338625 | 1.3035207354771 | 0.234761462846318 | 0.843284351171898 | -4.63838196207106 |
| **Echdc1** | 0.245817532499998 | 20.69528540125 | 1.30349195350483 | 0.234770739534084 | 0.843284351171898 | -4.63840566350558 |
| **Pfkl** | -0.355237774999999 | 28.4445696 | -1.30270499497436 | 0.235024507116075 | 0.843284351171898 | -4.63905361310893 |
| **Tufm** | 0.266010930000007 | 27.158382155 | 1.30266838070564 | 0.235036319798774 | 0.843284351171898 | -4.63908375526958 |
| **Ndufa5** | 0.201370780000001 | 22.8692207725 | 1.30036974314135 | 0.235778954131347 | 0.843284351171898 | -4.64097526582005 |
| **Bsg** | -0.162896217500009 | 25.71228586625 | -1.29959404137098 | 0.236030025319023 | 0.843284351171898 | -4.64161321772259 |
| **Nlgn3** | 0.168316679999997 | 27.80074987 | 1.29639831479063 | 0.23706684066625 | 0.843284351171898 | -4.64423952012722 |
| **Mtch2** | 0.138311404999993 | 25.5853852325 | 1.29397653224047 | 0.23785519515572 | 0.843284351171898 | -4.64622771192962 |
| **Svop** | 0.323086424999993 | 23.110361275 | 1.29383080210479 | 0.237902706783104 | 0.843284351171898 | -4.64634729372021 |
| **Ywhae** | 0.150494192499991 | 29.16280714625 | 1.29329943007522 | 0.238076017067441 | 0.843284351171898 | -4.64678326667294 |
| **Maob** | -0.486834439999999 | 23.41698436 | -1.29252797862983 | 0.238327825974402 | 0.843284351171898 | -4.64741606285677 |
| **Aco2** | -0.360406002499992 | 21.97068277875 | -1.29194602034991 | 0.238517935618967 | 0.843284351171898 | -4.64789330331387 |
| **Dctn2** | 0.239223757499996 | 24.16350138625 | 1.29062883897456 | 0.238948708721988 | 0.843284351171898 | -4.64897308645541 |
| **Kcna4** | -0.828225485000004 | 21.1263214325 | -1.290210593461 | 0.239085633628233 | 0.843284351171898 | -4.64931583915454 |
| **Chchd3** | 0.3697334875 | 23.05077991875 | 1.28944362175715 | 0.239336901230811 | 0.843284351171898 | -4.64994423342392 |

| | | | | | | |
|---|---|---|---|---|---|---|
| **Cox7c** | 0.375133907499993 | 20.95765625375 | 1.28865483883663 | 0.239595553427934 | 0.843284351171898 | -4.65059030886957 |
| **Slc25a27** | 0.171473432500008 | 26.25758607125 | 1.28846880199559 | 0.23965659267017 | 0.843284351171898 | -4.65074265973671 |
| **Rplp2** | 0.227487062500007 | 23.64537441375 | 1.28758827939539 | 0.239945677839323 | 0.843284351171898 | -4.65146359969787 |
| **Fscn1** | -0.163458759999997 | 28.198311475 | -1.28758353620374 | 0.239947235899632 | 0.843284351171898 | -4.65146748260206 |
| **Acot13** | 0.1592511325 | 25.89743896625 | 1.28451142732715 | 0.240958217962629 | 0.843284351171898 | -4.65398092902017 |
| **Eif4b** | -5.845546375 | 17.325398985 | -1.28377515618628 | 0.241201060836524 | 0.843284351171898 | -4.65458287397368 |
| **Trp53i11** | -1.055354565 | 21.6848948775 | -1.28337369022989 | 0.241333564889946 | 0.843284351171898 | -4.65491102489843 |
| **Ndufb2** | 0.790300932499996 | 20.64599949125 | 1.28312597281808 | 0.241415355611695 | 0.843284351171898 | -4.65511347952323 |
| **Pcp4** | 6.03502249 | 18.0591729525 | 1.280962743748 | 0.242130626124519 | 0.843284351171898 | -4.6568806307204 |
| **Rab6b** | 6.05319558749999 | 18.02198814875 | 1.28057868704859 | 0.242257805886867 | 0.843284351171898 | -4.65719421538788 |
| **Opa1** | -0.570820140000002 | 20.8082288975 | -1.28040664945502 | 0.242314794612201 | 0.843284351171898 | -4.65733467016837 |
| **Immt** | 0.800617772499997 | 23.56509710875 | 1.27910850907528 | 0.242745187442494 | 0.843284351171898 | -4.65839419768646 |
| **Aldh2** | 0.164846537500001 | 31.54238729875 | 1.27671044475696 | 0.24354199492458 | 0.843284351171898 | -4.6603500784082 |
| **Cltb** | 0.652091652499998 | 23.82674960875 | 1.27472686591671 | 0.244202788917329 | 0.843284351171898 | -4.66196653167708 |
| **Tom1l2** | -6.482939695 | 19.4799859675 | -1.27428700846757 | 0.244349529323406 | 0.843284351171898 | -4.66232481078073 |
| **Acsl3** | -0.355222692500003 | 21.33410087375 | -1.27411393487208 | 0.244407289117257 | 0.843284351171898 | -4.66246576841916 |
| **Aldh5a1** | 0.196051054999998 | 26.2473624125 | 1.27347079670993 | 0.244622026775301 | 0.843284351171898 | -4.66298948115474 |
| **Rab24** | 0.174223612499993 | 23.98688038625 | 1.27262285717414 | 0.244905394719223 | 0.843284351171898 | -4.66367976489094 |
| **S100b** | 0.2531736575 | 21.26018042125 | 1.27045431870041 | 0.245631375609828 | 0.843284351171898 | -4.66544407180052 |
| **Suclg1** | -0.298014870000006 | 25.90527483 | -1.2701538858188 | 0.245732100556392 | 0.843284351171898 | -4.66568838370031 |

| | | | | | | |
|---|---|---|---|---|---|---|
| **Slc6a1** | 6.40623879 | 19.14085128 | 1.26882579692353 | 0.246177790814132 | 0.843284351171898 | -4.6667680400191 |
| **Gabra6** | 0.336690012500007 | 24.42306851125 | 1.26573322161435 | 0.247218323819919 | 0.843284351171898 | -4.66927993061954 |
| **Paics** | -0.307195870000001 | 21.823716945 | -1.26534982209625 | 0.247347586824957 | 0.843284351171898 | -4.66959112644702 |
| **Gad2** | 0.608153509999998 | 21.41963848 | 1.26504945844428 | 0.247448895002593 | 0.843284351171898 | -4.66983489111713 |
| **Mapk10** | 0.171073262499995 | 26.31379534625 | 1.2620574018372 | 0.248460024127061 | 0.843284351171898 | -4.67226155301243 |
| **Cox4i1** | -0.1509238575 | 23.09393841125 | -1.26120052541764 | 0.248750249578102 | 0.843284351171898 | -4.67295597701264 |
| **Ywhab** | 0.276443025000006 | 20.12680056 | 1.25795553774774 | 0.249851977919528 | 0.843284351171898 | -4.67558359998694 |
| **Ckap4** | 7.13326329916666 | 19.18133032 | 1.21611583497246 | 0.271067551560651 | 0.843284351171898 | -4.6780070798505 |
| **Gsk3b** | -0.181100432499992 | 26.62111042375 | -1.25487302284453 | 0.250902427166591 | 0.843284351171898 | -4.67807648575416 |
| **Sod2** | 5.36146959 | 16.30170952 | 1.25472042306566 | 0.250954528044544 | 0.843284351171898 | -4.67819981530464 |
| **Necap1** | 0.556597042500002 | 22.46520161375 | 1.25425488402424 | 0.251113530604329 | 0.843284351171898 | -4.67857601198867 |
| **Hsdl1** | -0.375362812500001 | 24.37389920375 | -1.2531313714234 | 0.251497617173395 | 0.843284351171898 | -4.67948361685511 |
| **Rab6a** | 6.5941476575 | 20.07864825375 | 1.25301911996158 | 0.251536019414505 | 0.843284351171898 | -4.67957427396584 |
| **Ndufa9** | 0.245830877499991 | 24.76066865125 | 1.25290953797447 | 0.25157351326008 | 0.843284351171898 | -4.67966277115065 |
| **Acat2** | 0.241582910000002 | 24.96326838 | 1.25042257444284 | 0.252425726922128 | 0.843284351171898 | -4.68167015327175 |
| **Nherf1** | -0.510767464999997 | 23.66778641 | -1.24986018164558 | 0.252618786409838 | 0.843284351171898 | -4.68212381281673 |
| **Bpnt1** | 0.329885405000006 | 24.68713512 | 1.24985708606608 | 0.252619849417836 | 0.843284351171898 | -4.68212630960681 |
| **Dlst** | 0.1666554725 | 29.83973387375 | 1.24889928166113 | 0.252948939393204 | 0.843284351171898 | -4.68289869036769 |
| **Rps27a** | 6.2555404875 | 19.11595728875 | 1.24432164552574 | 0.254526839339341 | 0.843284351171898 | -4.68658593356354 |
| **Dbnl** | -0.525486399999998 | 25.9863046775 | -1.24370885066597 | 0.254738706708859 | 0.843284351171898 | -4.68707900542127 |

| | | | | | | |
|---|---|---|---|---|---|---|
| **Rpl30** | -0.170411107500009 | 24.58033355125 | -1.24062371868358 | 0.255807652907009 | 0.843284351171898 | -4.68955948371844 |
| **Mrps26** | -5.65028663 | 17.38069554 | -1.23772587073411 | 0.256815200037472 | 0.843284351171898 | -4.69188647796831 |
| **Slc25a12** | 0.331217022499992 | 25.11076743375 | 1.23743311873091 | 0.256917174780907 | 0.843284351171898 | -4.69212140299754 |
| **Eif5** | 6.0925810025 | 18.71014933375 | 1.23545288221084 | 0.25760786225788 | 0.843284351171898 | -4.69370972648941 |
| **Actb** | 0.171528502499999 | 25.11164645625 | 1.23454767219408 | 0.25792411850588 | 0.843284351171898 | -4.6944353430797 |
| **Sod1** | 0.129007412499995 | 23.73420613125 | 1.23391762060615 | 0.258144437304248 | 0.843284351171898 | -4.69494022885798 |
| **Ykt6** | 0.209039367500001 | 24.21983660375 | 1.23368697600757 | 0.258225130122538 | 0.843284351171898 | -4.69512502005112 |
| **Ppp1cb** | -5.88294869 | 18.155319905 | -1.23337742698852 | 0.258333462099508 | 0.843284351171898 | -4.69537300068338 |
| **Capzb** | -0.140346297500002 | 22.85441233875 | -1.23226559028429 | 0.258722888215654 | 0.843284351171898 | -4.69626342815385 |
| **Vps26b** | 6.3030865625 | 19.47460107625 | 1.23175354613726 | 0.258902402473136 | 0.843284351171898 | -4.69667336345834 |
| **Gria3** | -0.163614520000007 | 28.48178905 | -1.23151769389188 | 0.258985124119392 | 0.843284351171898 | -4.69686215343441 |
| **Ermn** | -0.176829944999998 | 25.3129083475 | -1.23094653061469 | 0.259185544480116 | 0.843284351171898 | -4.69731926759265 |
| **H1-3** | -0.197966904999991 | 32.062035345 | -1.23022258588144 | 0.259439765649875 | 0.843284351171898 | -4.69789849631824 |
| **Idh3a** | 0.272714507500002 | 25.38457243875 | 1.22514195958782 | 0.261229868499353 | 0.843284351171898 | -4.70195846879356 |
| **Mtx2** | -5.5267725125 | 17.21448063375 | -1.2191443402787 | 0.263356581450412 | 0.843284351171898 | -4.70673978916476 |
| **Hepacam** | 0.264829622499995 | 24.90215224625 | 1.21910232827221 | 0.263371530350093 | 0.843284351171898 | -4.70677323735489 |
| **Ass1** | 0.447765947499995 | 23.13721469375 | 1.21860419793861 | 0.263548832250798 | 0.843284351171898 | -4.70716978102124 |
| **Pcyox1** | 0.125327022500002 | 20.84159801375 | 1.21786949478109 | 0.263810523647707 | 0.843284351171898 | -4.70775449441184 |
| **Ndufa10** | -5.6601580875 | 17.64430223125 | -1.21661131964875 | 0.264259181135096 | 0.843284351171898 | -4.70875537657471 |
| **Ctnnd2** | 0.309004754999997 | 24.67704712 | 1.21485323109712 | 0.264887189179494 | 0.843284351171898 | -4.71015301677375 |

| | | | | | | |
|---|---|---|---|---|---|---|
| **Cd99l2** | -0.560628800000007 | 22.6019968075 | -1.21263570627277 | 0.265681117729746 | 0.843284351171898 | -4.71191435681657 |
| **Cpt1c** | -0.115925082500002 | 26.8965464937 5 | -1.20783902841829 | 0.267405347294494 | 0.843284351171898 | -4.71571836126107 |
| **Slc30a9** | 0.186433089999998 | 27.87783205 | 1.20699001785753 | 0.267711520096072 | 0.843284351171898 | -4.71639082336435 |
| **Hpcal1** | 0.656871519999999 | 22.59573735 | 1.20675934351302 | 0.267794757812533 | 0.843284351171898 | -4.71657348580059 |
| **Gstz1** | -5.998492655 | 18.85175756 | -1.20649546854415 | 0.267890002660589 | 0.843284351171898 | -4.71678241546777 |
| **Map2k1** | 0.368692929999998 | 22.24075702 | 1.20648343909184 | 0.267894345336813 | 0.843284351171898 | -4.71679193950239 |
| **Micu1** | -0.73167566 | 22.458616905 | -1.20625617135722 | 0.267976401009942 | 0.843284351171898 | -4.71697186372546 |
| **Napb** | 6.09457115 | 19.0189861 | 1.20594104755417 | 0.268090212545302 | 0.843284351171898 | -4.71722131189793 |
| **Tpbg** | 0.263449529999999 | 25.5934076825 | 1.20506618812215 | 0.268406395034698 | 0.843284351171898 | -4.71791365534667 |
| **Vdac2** | -0.321528527499993 | 26.01073817125 | -1.20416306687624 | 0.268733122295487 | 0.843284351171898 | -4.71862807943038 |
| **Vdac3** | 0.212416667499998 | 28.52576524125 | 1.20402080245609 | 0.268784620735576 | 0.843284351171898 | -4.71874059284547 |
| **Acaa1a** | 0.21972087499999 | 21.1962621425 | 1.20037180599002 | 0.270108377423808 | 0.843284351171898 | -4.72162402905177 |
| **Ndufc2** | -5.7600817875 | 18.17038852125 | -1.19922579904873 | 0.270525251848402 | 0.843284351171898 | -4.72252862184329 |
| **Slc6a5** | -0.434995230000006 | 23.58080305 | -1.19764965419667 | 0.271099481083949 | 0.843284351171898 | -4.72377197205566 |
| **Hk1** | -6.27255259499999 | 19.7387146125 | -1.19739217050067 | 0.27119338650917 | 0.843284351171898 | -4.72397500467774 |
| **Atp1b1** | 5.5915339225 | 17.74204597125 | 1.19288645101942 | 0.272841085753875 | 0.843284351171898 | -4.72752401673335 |
| **Ppm1h** | 6.2563671775 | 19.83839818875 | 1.19284514681649 | 0.272856229236147 | 0.843284351171898 | -4.72755651682983 |
| **Lancl1** | -0.566930105000004 | 22.160600505 | -1.19212225273314 | 0.273121380644324 | 0.843284351171898 | -4.72812522399395 |
| **Flot2** | 5.330235775 | 16.948211465 | 1.19137089692604 | 0.273397201441175 | 0.843284351171898 | -4.72871612153524 |
| **Ccny** | 0.19059979 | 22.725212295 | 1.19095518441854 | 0.273549909160114 | 0.843284351171898 | -4.72904296719012 |

| | | | | | | |
|---|---|---|---|---|---|---|
| **Tagln3** | 0.1889508525 | 23.50719052375 | 1.18821372514792 | 0.274558754239974 | 0.843284351171898 | -4.73119681173163 |
| **Kcnc1** | 5.7401928325 | 18.27745500875 | 1.18742116582132 | 0.274850994973165 | 0.843284351171898 | -4.73181898080771 |
| **Pfkp** | 0.270864824999993 | 24.4887230675 | 1.18471498806082 | 0.275850815966183 | 0.843284351171898 | -4.73394163464044 |
| **Pptc7** | -0.237606102499999 | 28.20306827875 | -1.1846210126407 | 0.275885590820197 | 0.843284351171898 | -4.73401529829313 |
| **Cox6a1** | 0.334714952500008 | 24.29594553625 | 1.1842992145102 | 0.276004697536426 | 0.843284351171898 | -4.7342675187199 |
| **Ndufa6** | 0.158752645000014 | 26.342433525 | 1.18299566346215 | 0.276487620911531 | 0.843284351171898 | -4.73528883320867 |
| **Rpl15** | 0.207370697499996 | 26.69693906875 | 1.18273673376426 | 0.276583630347593 | 0.843284351171898 | -4.73549162679017 |
| **Ndufb7** | -0.297076814999997 | 22.8155371025 | -1.18202007262878 | 0.27684950949315 | 0.843284351171898 | -4.73605278678521 |
| **Abcb1a** | -0.710766225 | 24.8578074125 | -1.18127246493306 | 0.277127098217817 | 0.843284351171898 | -4.73663797707495 |
| **Bri3bp** | 0.153617494999999 | 27.5629502375 | 1.18025539112942 | 0.277505115232836 | 0.843284351171898 | -4.73743376133587 |
| **Rmdn3** | 6.0461979325 | 19.37027731875 | 1.17995030659345 | 0.277618590628719 | 0.843284351171898 | -4.73767239277725 |
| **Ccdc136** | -5.48401855999999 | 17.606601045 | -1.17837000308841 | 0.278207003172221 | 0.843284351171898 | -4.73890792627321 |
| **Epha4** | 0.189176407500003 | 23.53901421375 | 1.17783413695322 | 0.278406765498952 | 0.843284351171898 | -4.73932667435738 |
| **Rps10** | -5.84949248 | 18.7896876425 | -1.17646129998759 | 0.278919085488749 | 0.843284351171898 | -4.7403989806808 |
| **Gabbr2** | 0.129842414999999 | 26.0506158225 | 1.17451332308338 | 0.279647392786862 | 0.843284351171898 | -4.74191932074046 |
| **Gng2** | 1.11205407500001 | 22.23672469 | 1.1743799755171 | 0.279697306772836 | 0.843284351171898 | -4.742023343066 |
| **Atp5po** | 0.285426722500006 | 24.85528104375 | 1.17362105450165 | 0.279981524213032 | 0.843284351171898 | -4.74261523920274 |
| **Guk1** | -5.33518562 | 17.1783952475 | -1.17315786211698 | 0.28015510939635 | 0.843284351171898 | -4.74297638575264 |
| **Hsp90ab1** | 0.269116035 | 21.0080655325 | 1.17194637775312 | 0.280609548844826 | 0.843284351171898 | -4.74392058956982 |
| **Tprg1l** | 0.233693879999997 | 24.1572132875 | 1.1707612672563 | 0.281054691389432 | 0.843284351171898 | -4.74484370735852 |
| **Rap1a** | -0.117135944999998 | 25.0435891 | -1.16934995760901 | 0.281585566921676 | 0.843284351171898 | -4.74594233221961 |

| | | | | | | |
|---|---|---|---|---|---|---|
| **Mras** | -0.147121512499993 | 27.48040411875 | -1.16913215116723 | 0.281667571148049 | 0.843284351171898 | -4.74611181569619 |
| **Nckap1** | 0.38232953 | 25.8746102375 | 1.16889709553579 | 0.28175609208137 | 0.843284351171898 | -4.7462947014434 |
| **Mcu** | 0.320264687499993 | 21.55480535125 | 1.16765559523283 | 0.282224021231337 | 0.843284351171898 | -4.74726031015177 |
| **Abhd6** | 5.6862085475 | 18.35047483125 | 1.16732486045821 | 0.282348786605742 | 0.843284351171898 | -4.74751744983387 |
| **Usp5** | 6.24635216500001 | 20.22176332 | 1.16541371188931 | 0.283070644355557 | 0.843284351171898 | -4.74900252264946 |
| **Mettl26** | -0.127229377499997 | 21.13308634875 | -1.16531274324415 | 0.283108823891465 | 0.843284351171898 | -4.74908094280644 |
| **Actn1** | 0.108515780000005 | 27.5702327225 | 1.16531088630851 | 0.283109526099584 | 0.843284351171898 | -4.74908238501204 |
| **Slc25a4** | -0.220373535 | 18.74561895 | -1.16527865931857 | 0.28312171310446 | 0.843284351171898 | -4.74910741418331 |
| **Slc2a1** | 0.466698212499992 | 23.46084425875 | 1.16475711830774 | 0.283319000654613 | 0.843284351171898 | -4.74951241575895 |
| **Psmb2** | -5.2986207025 | 17.15060510125 | -1.16402380163263 | 0.283596592231377 | 0.843284351171898 | -4.75008169740297 |
| **Sirt2** | -0.293220564999999 | 25.7599020525 | -1.16299130171869 | 0.283987821785926 | 0.843284351171898 | -4.75088289332272 |
| **Acat1** | 0.234330140000004 | 26.3596439525 | 1.16267455326805 | 0.284107932615622 | 0.843284351171898 | -4.75112860182256 |
| **Adss1** | 0.136996025000006 | 28.5631575625 | 1.16155055531531 | 0.28453449368722 | 0.843284351171898 | -4.7520002038624 |
| **Ndufs3** | 0.394410780000008 | 25.57783478 | 1.16146374128128 | 0.284567462096068 | 0.843284351171898 | -4.7520675036919 |
| **Bin1** | -0.134465469999995 | 28.0727758825 | -1.1612301515806 | 0.284656185682548 | 0.843284351171898 | -4.75224857249544 |
| **Ppm1e** | 0.291374400000002 | 22.5330559375 | 1.15900332555134 | 0.28550315069148 | 0.843284351171898 | -4.75397367166749 |
| **Cd82** | -0.408739197499997 | 26.29052453625 | -1.15856930007832 | 0.285668474715785 | 0.843284351171898 | -4.75430968684034 |
| **Slc25a51** | 0.694035517499991 | 22.90924436125 | 1.15843642669994 | 0.285719103250632 | 0.843284351171898 | -4.75441254079838 |
| **Eif4a2** | 5.96074873 | 19.3147101675 | 1.15813419318825 | 0.285834290618651 | 0.843284351171898 | -4.75464646711485 |
| **Mpc1** | 0.179827472500005 | 25.37459456375 | 1.1580402099803 | 0.285870117414823 | 0.843284351171898 | -4.75471920225999 |

| | | | | | | |
|---|---|---|---|---|---|---|
| **Ntrk2** | -0.427558955000006 | 23.6422985625 | -1.1575946666371 | 0.286040011263486 | 0.843284351171898 | -4.75506396967322 |
| **Pc** | -0.275633540000001 | 23.7616707225 | -1.15639144300365 | 0.286499242229516 | 0.843284351171898 | -4.75599466126197 |
| **Emd** | -5.74876144 | 18.6074856175 | -1.15614861331756 | 0.286591996667469 | 0.843284351171898 | -4.75618242249666 |
| **Slc25a3** | 5.91941786 | 18.95799605 | 1.15289310085639 | 0.287837927632466 | 0.843284351171898 | -4.75869747322245 |
| **Hsd17b4** | 5.76696145 | 18.8473133075 | 1.15236267256853 | 0.288041355756447 | 0.843284351171898 | -4.7591068706794 |
| **Rgs17** | -0.175463379999997 | 26.6882076725 | -1.15136482158798 | 0.288424371823441 | 0.843284351171898 | -4.75987674278087 |
| **Syt7** | 0.244760782500002 | 23.75592640875 | 1.15129555299494 | 0.288450975631515 | 0.843284351171898 | -4.75993017134743 |
| **Gpd1** | 6.40706457333333 | 18.5824012628571 | 1.10301661571028 | 0.313634697748701 | 0.843284351171898 | -4.76005966940806 |
| **Hsp90aa1** | 5.7627039425 | 18.88221639625 | 1.15051415678293 | 0.288751225536981 | 0.843284351171898 | -4.76053275325826 |
| **Gprin1** | 0.569940642500001 | 22.92105400125 | 1.14811365427704 | 0.289675236014805 | 0.843284351171898 | -4.76238245083094 |
| **Rpl3** | -5.98433442750001 | 19.60683914625 | -1.1476698976669 | 0.289846316720751 | 0.843284351171898 | -4.76272414150546 |
| **Anxa7** | 0.184109262500002 | 20.89195492875 | 1.14716092103919 | 0.290042644701306 | 0.843284351171898 | -4.76311595733638 |
| **Ncam1** | 0.153505489999997 | 27.126608365 | 1.14604172723512 | 0.290474739892334 | 0.843284351171898 | -4.76397717158933 |
| **Rps4x** | 0.165375885000003 | 24.3539119025 | 1.14457385103086 | 0.2910422616369 | 0.843284351171898 | -4.76510595746922 |
| **Maoa** | 0.285472697499994 | 23.45219242375 | 1.14435598471563 | 0.291126573014546 | 0.843284351171898 | -4.76527342359186 |
| **Ncam2** | -0.1661416 | 31.4894502875 | -1.14359528863005 | 0.291421110894099 | 0.843284351171898 | -4.76585799874338 |
| **Lmna** | 6.277778715 | 20.647433455 | 1.14357353425184 | 0.291429537711753 | 0.843284351171898 | -4.76587471309988 |
| **Retreg2** | -5.1436042875 | 16.94029782125 | -1.14307446925156 | 0.291622911878172 | 0.843284351171898 | -4.76625810478406 |
| **Baiap2** | -5.18376243 | 17.0183121725 | -1.14302122148233 | 0.291643550214035 | 0.843284351171898 | -4.76629900504561 |
| **Col9a1** | 5.89025943 | 19.40584084 | 1.14146693400223 | 0.292246510162857 | 0.843284351171898 | -4.76749238437052 |

| | | | | | | |
|---|---|---|---|---|---|---|
| **Lpin3** | -5.610234945 | 18.448339755 | -1.13985969895356 | 0.29287109417314 | 0.843284351171898 | -4.76872542304099 |
| **Ap3b2** | 5.436044105 | 17.928090425 | 1.13983028145515 | 0.29288253631722 | 0.843284351171898 | -4.76874798212885 |
| **Cap2** | -0.328250747500004 | 26.75260721375 | -1.13930895491156 | 0.293085371265963 | 0.843284351171898 | -4.76914771004177 |
| **Cpd** | -0.219266244999993 | 24.5257737975 | -1.13909492265325 | 0.29316867940314 | 0.843284351171898 | -4.76931178872745 |
| **Sntb1** | -5.028942095 | 16.5942161625 | -1.13825707581454 | 0.293494984184419 | 0.843284351171898 | -4.76995391521234 |
| **Sfxn5** | 0.389118515000007 | 23.0561040575 | 1.13767977835753 | 0.293719990895284 | 0.843284351171898 | -4.77039619595473 |
| **Kcnab1** | -0.287551220000005 | 25.8072578125 | -1.13717783673664 | 0.293915742787435 | 0.843284351171898 | -4.77078063835652 |
| **Plcb1** | 0.213173507500002 | 20.47112563875 | 1.13662818923375 | 0.294130223029533 | 0.843284351171898 | -4.77120150545698 |
| **Cntn1** | -0.168737310000001 | 24.79230914 | -1.1362017656003 | 0.294296708571922 | 0.843284351171898 | -4.77152793762694 |
| **Fkbp8** | -5.98267948749999 | 19.78026624625 | -1.13576138181595 | 0.294468726120165 | 0.843284351171898 | -4.77186498122942 |
| **Jam3** | -0.186477342500005 | 22.64555982125 | -1.1345437279649 | 0.294944783736806 | 0.843284351171898 | -4.7727965029491 |
| **Ddx3x** | -5.97965454499999 | 19.82920279 | -1.13377394886418 | 0.295246066345783 | 0.843284351171898 | -4.77338509227424 |
| **Ap1b1** | 5.40787189749999 | 17.70979272625 | 1.13351255126056 | 0.295348432049756 | 0.843284351171898 | -4.77358490910169 |
| **Iars2** | -0.124121767500004 | 27.91440837875 | -1.13341850359048 | 0.295385269140715 | 0.843284351171898 | -4.77365679414698 |
| **Tmem65** | 5.5537904225 | 18.40346178375 | 1.13327961011024 | 0.295439678606192 | 0.843284351171898 | -4.77376295054695 |
| **Kcna2** | 0.270109057500001 | 21.93342255625 | 1.13327578452766 | 0.295441177338494 | 0.843284351171898 | -4.77376587433478 |
| **Stmn1** | 5.8609340225 | 19.41076830625 | 1.13212564725219 | 0.295892046079562 | 0.843284351171898 | -4.7746446299198 |
| **Map7d2** | -5.1391281775 | 17.08429822625 | -1.13100054189754 | 0.29633365058572 | 0.843284351171898 | -4.77550375259953 |
| **Ahcy** | -0.162354037499998 | 25.95431110625 | -1.13090799635442 | 0.29636999892446 | 0.843284351171898 | -4.77557439737508 |
| **Scg2** | 0.282252722500001 | 22.66626015375 | 1.12966236418611 | 0.296859592955422 | 0.843284351171898 | -4.77652492134071 |

| | | | | | | |
|---|---|---|---|---|---|---|
| **Mapre2** | 5.07276126 | 16.883356445 | 1.12884792719513 | 0.297180066280515 | 0.843284351171898 | -4.77714607279446 |
| **Mccc2** | -0.258291877499996 | 24.57046744125 | -1.12812442743333 | 0.297464995495418 | 0.843284351171898 | -4.77769764702666 |
| **Septin6** | 0.480912905000004 | 23.066421125 | 1.12764146151339 | 0.297655322605493 | 0.843284351171898 | -4.77806572941622 |
| **Arf5** | -0.149823575000003 | 30.7850543625 | -1.12746790120255 | 0.297723743688781 | 0.843284351171898 | -4.77819798205347 |
| **Asph** | 0.183749197499992 | 23.47310320625 | 1.1273533665535 | 0.29776890271635 | 0.843284351171898 | -4.77828525066083 |
| **Eef1g** | 0.381078277500002 | 23.32146219375 | 1.12642201555544 | 0.298136327543875 | 0.843284351171898 | -4.77899469037951 |
| **Apoo** | -5.29072298 | 17.579489795 | -1.12569227468594 | 0.29842447617925 | 0.843284351171898 | -4.77955031501644 |
| **Slc9a1** | 0.601307479999996 | 21.9765330175 | 1.12515496763315 | 0.298636785916089 | 0.843284351171898 | -4.7799592843769 |
| **Ahsa1** | 0.752821849999997 | 22.155484365 | 1.12451521656063 | 0.298889736972163 | 0.843284351171898 | -4.78044607787015 |
| **Shisa4** | -0.121854837499995 | 26.29179414375 | -1.12446475081871 | 0.298909698104011 | 0.843284351171898 | -4.78048447081946 |
| **Ncan** | 0.396437957499995 | 25.64613689875 | 1.12398688798729 | 0.299098765448292 | 0.843284351171898 | -4.78084796512008 |
| **Clptm1** | 0.940236970000001 | 21.36736117 | 1.12389599152982 | 0.299134739928037 | 0.843284351171898 | -4.78091709665453 |
| **Tmed2** | 5.3989139375 | 18.04017197625 | 1.12332250372615 | 0.299361793635243 | 0.843284351171898 | -4.78135318784057 |
| **Septin4** | -5.87554899249999 | 19.63219406625 | -1.12302886199633 | 0.299478106255944 | 0.843284351171898 | -4.78157642757007 |
| **Syt3** | 5.92526668 | 19.8077312975 | 1.12288045362286 | 0.299536905505482 | 0.843284351171898 | -4.78168924115717 |
| **Acad8** | 5.3548814275 | 17.90651810625 | 1.12279164822482 | 0.299572094650313 | 0.843284351171898 | -4.78175674292498 |
| **Gng12** | 0.262977415000002 | 24.2225549925 | 1.12254690311581 | 0.299669092490233 | 0.843284351171898 | -4.78194275944472 |
| **Aldoa** | 5.7133886325 | 19.09093820875 | 1.12205897978319 | 0.299862544148592 | 0.843284351171898 | -4.782313529739 |
| **Git1** | 5.8665511825 | 19.63402054625 | 1.12159922371668 | 0.300044921887418 | 0.843284351171898 | -4.7826628082647 |
| **Mrrf** | 5.6383516025 | 18.66467528875 | 1.12104276814573 | 0.300265780546932 | 0.843284351171898 | -4.78308543599784 |
| **Hdhd3** | 5.72111057 | 19.1815345125 | 1.11956323055516 | 0.300853661962119 | 0.843284351171898 | -4.78420853717085 |

| | | | | | | |
|---|---|---|---|---|---|---|
| **Chmp5** | 5.09632671 | 17.0793080475 | 1.11914102583881 | 0.301021594424114 | 0.843284351171898 | -4.78452886630366 |
| **Pdxp** | -0.201045745000002 | 29.146840995 | -1.11839066782768 | 0.301320240019049 | 0.843284351171898 | -4.78509798952561 |
| **Rpl6** | 5.7267874775 | 19.20313063875 | 1.11807250194057 | 0.301446944692533 | 0.843284351171898 | -4.78533923975086 |
| **Timm8a1** | 4.9704717725 | 16.68701149125 | 1.1170898672719 | 0.301838539649905 | 0.843284351171898 | -4.78608406681749 |
| **Ttyh1** | -0.834295675 | 21.389340695 | -1.11661930943442 | 0.302026211741314 | 0.843284351171898 | -4.78644060641542 |
| **Sdha** | 5.6340742925 | 18.94983712875 | 1.11535456356133 | 0.30253110291361 | 0.843284351171898 | -4.78739845386126 |
| **Thy1** | -0.264193755000004 | 30.250493055 | -1.11351833118145 | 0.303265363768062 | 0.843284351171898 | -4.78878795679286 |
| **Rpl8** | -5.925419955 | 19.9523374275 | -1.11334345907956 | 0.303335366523213 | 0.843284351171898 | -4.78892021345291 |
| **Mtco2** | 6.10492173 | 20.5762688875 | 1.11224059422586 | 0.303777157228147 | 0.843284351171898 | -4.78975402873859 |
| **Rab39b** | -0.219780372500001 | 27.05210958625 | -1.11166535756164 | 0.304007797004365 | 0.843284351171898 | -4.79018873667366 |
| **Hspe1** | 5.4345229525 | 18.30231720375 | 1.1114274829076 | 0.304103214134713 | 0.843284351171898 | -4.79036845974241 |
| **Prxl2a** | 5.8277692625 | 19.65905768875 | 1.11134822732866 | 0.304135010854765 | 0.843284351171898 | -4.79042833512312 |
| **Lmnb2** | -4.8853088775 | 16.44897874375 | -1.1111770539028 | 0.30420369357549 | 0.843284351171898 | -4.79055764312312 |
| **Nudt3** | 6.0121862425 | 20.30735329375 | 1.10961694534907 | 0.304830266238486 | 0.843284351171898 | -4.79173562996663 |
| **Cttn** | 0.106562347499992 | 25.94504240875 | 1.10927157687538 | 0.304969116049759 | 0.843284351171898 | -4.79199627177953 |
| **Acadl** | -0.454785347499996 | 24.08886068625 | -1.10828132747174 | 0.3053675163504 | 0.843284351171898 | -4.79274331982986 |
| **Park7** | 0.288384372500001 | 23.52308721625 | 1.1075063976628 | 0.305679585028906 | 0.843284351171898 | -4.79332764949319 |
| **Krt42** | 0.117297992500003 | 24.64408954125 | 1.10724576481817 | 0.305784601880181 | 0.843284351171898 | -4.79352412223238 |
| **Emc8** | 5.1250996175 | 17.35252828875 | 1.10673949223749 | 0.305988678594159 | 0.843284351171898 | -4.79390568573091 |
| **Psmc3** | 5.79189792 | 19.610376115 | 1.10672994653339 | 0.30599252750201 | 0.843284351171898 | -4.79391287904988 |

| | | | | | | |
|---|---|---|---|---|---|---|
| **Rpl32** | 0.172231892500005 | 24.33882666125 | 1.10654678479642 | 0.306066387503705 | 0.843284351171898 | -4.79405089627159 |
| **Wfs1** | 5.848807945 | 19.7723290775 | 1.10631742885445 | 0.306158895847525 | 0.843284351171898 | -4.79422370260427 |
| **Lta4h** | -0.154626239999995 | 25.0382424075 | -1.10568390388011 | 0.306414540169672 | 0.843284351171898 | -4.79470091433109 |
| **Sdhc** | 5.4969850825 | 18.63259843625 | 1.10523471055168 | 0.306595907251183 | 0.843284351171898 | -4.79503917554369 |
| **Srsf2** | 5.58782448 | 18.94333718 | 1.10488095001718 | 0.306738803865679 | 0.843284351171898 | -4.79530551341716 |
| **Coro2b** | 4.88836013 | 16.5701701925 | 1.10486368970489 | 0.306745777317074 | 0.843284351171898 | -4.79531850697577 |
| **Tmed10** | 0.168381622499997 | 23.87671223125 | 1.10396244720749 | 0.307110073879889 | 0.843284351171898 | -4.79599679148773 |
| **Ache** | 4.9925913375 | 16.86507618125 | 1.10383817817459 | 0.307160333070296 | 0.843284351171898 | -4.79609029138156 |
| **Slc30a1** | -0.441841182500003 | 20.80141956625 | -1.10370654084174 | 0.307213579600672 | 0.843284351171898 | -4.79618932822694 |
| **Ca14** | 1.17686006750001 | 20.89970608875 | 1.10278708885066 | 0.3075857024476 | 0.843284351171898 | -4.79688087493666 |
| **C2orf72** | -0.674373475000003 | 25.095352405 | -1.10143943706128 | 0.30813179117273 | 0.843284351171898 | -4.79789385186 |
| **Dixdc1** | -0.302327640000001 | 22.9130422075 | -1.1009711923455 | 0.308321715771402 | 0.843284351171898 | -4.79824563709112 |
| **Gtpbp1** | 0.181621857500005 | 26.53319310375 | 1.09745119366498 | 0.309752515133234 | 0.843284351171898 | -4.80088724741606 |
| **Vti1b** | 5.8163335 | 19.84424552 | 1.09680999301784 | 0.310013729388006 | 0.843284351171898 | -4.80136788680827 |
| **Cask** | 5.545343035 | 18.9025143025 | 1.0966456718932 | 0.310080699853596 | 0.843284351171898 | -4.801491033173 |
| **Atp5mf** | 5.8705910975 | 19.89525110375 | 1.09594519922848 | 0.310366315387242 | 0.843284351171898 | -4.80201585867216 |
| **Clu** | 5.41846608 | 18.5008905175 | 1.09591825760531 | 0.310377305019651 | 0.843284351171898 | -4.80203604045693 |
| **Dnajc11** | 0.133452967499998 | 25.80846290125 | 1.09553303415907 | 0.310534474326799 | 0.843284351171898 | -4.80232457557089 |
| **Cnrip1** | 0.12510849500001 | 26.1646403525 | 1.09547224834371 | 0.310559280554503 | 0.843284351171898 | -4.8023700989241 |
| **Dync1h1** | -0.2278007275 | 26.51691889625 | -1.09541295183253 | 0.310583480560413 | 0.843284351171898 | -4.80241450543036 |
| **Cyb5r3** | 5.2935387875 | 18.10150361125 | 1.09377363533007 | 0.311253123069112 | 0.843284351171898 | -4.80364158994507 |

| | | | | | | |
|---|---|---|---|---|---|---|
| **Pcbp2** | 0.203451502499995 | 22.88466723375 | 1.0928824041617 | 0.311617672904647 | 0.843284351171898 | -4.80430823535309 |
| **Rpn2** | 0.665714767500013 | 23.04322282375 | 1.09194967934049 | 0.311999566529127 | 0.843284351171898 | -4.80500556189534 |
| **Trap1** | -0.203448962499998 | 18.98374930375 | -1.09172309717545 | 0.312092395363258 | 0.843284351171898 | -4.80517490486696 |
| **Psma2** | -5.6351367225 | 19.28170423625 | -1.09107022263486 | 0.31235999799624 | 0.843284351171898 | -4.80566272974811 |
| **Odf2l** | 5.9554283375 | 20.41854790875 | 1.09056792107937 | 0.312566009914508 | 0.843284351171898 | -4.80603792535724 |
| **Gpd2** | -5.6049800875 | 19.20925836375 | -1.09048644646978 | 0.312599435966951 | 0.843284351171898 | -4.80609877306251 |
| **Fam131b** | -0.2460093175 | 28.35464591375 | -1.09021879790661 | 0.312709262792627 | 0.843284351171898 | -4.80629864147613 |
| **Pdha1** | 0.293189062499994 | 25.40908882125 | 1.08938070347976 | 0.31305336863418 | 0.843284351171898 | -4.80692429942342 |
| **Csnk2b** | 0.159472692499989 | 33.74240004375 | 1.089350160917 | 0.313065914633732 | 0.843284351171898 | -4.80694709461077 |
| **Atp8a1** | -0.739470797499994 | 22.36143391875 | -1.08872473789806 | 0.313322909936987 | 0.843284351171898 | -4.80741378750656 |
| **Psmb7** | 0.313128660000011 | 24.66886746 | 1.08857008384343 | 0.313386485876908 | 0.843284351171898 | -4.80752916550932 |
| **Echs1** | -5.430239205 | 18.55737889 | -1.08839955813985 | 0.313456598532266 | 0.843284351171898 | -4.80765637270317 |
| **Vim** | 0.520105852500006 | 23.56849742875 | 1.08812629329779 | 0.313568979504918 | 0.843284351171898 | -4.80786019473618 |
| **Coq9** | 0.400290919999996 | 22.103447565 | 1.08538018180347 | 0.31470013932402 | 0.843284351171898 | -4.80990670696167 |
| **Fmnl2** | 0.133882502500001 | 26.33119237625 | 1.08521207942029 | 0.31476949011513 | 0.843284351171898 | -4.81003188004921 |
| **Slc4a10** | 1.45491533 | 23.3633259875 | 1.08454972089674 | 0.31504286713769 | 0.843284351171898 | -4.81052497173069 |
| **Atp5f1e** | 5.7191960475 | 19.67389678875 | 1.0836162075728 | 0.315428484808656 | 0.843284351171898 | -4.81121960851826 |
| **Srsf1** | -5.43543504 | 18.73317034 | -1.08353422118862 | 0.315462370152676 | 0.843284351171898 | -4.81128059777067 |
| **Ftl1** | 0.137222104999996 | 23.0782781925 | 1.08325914122699 | 0.315576083462135 | 0.843284351171898 | -4.81148520751165 |
| **Tmx4** | 0.239134592500001 | 20.83423955125 | 1.08155538951186 | 0.316281123728094 | 0.843284351171898 | -4.81275177474678 |

| | | | | | | |
|---|---|---|---|---|---|---|
| **Acot7** | 0.229107030000005 | 23.8376079475 | 1.08146183884114 | 0.316319873373221 | 0.843284351171898 | -4.81282128448057 |
| **Cdh13** | 5.2207566575 | 18.03289915125 | 1.0812870142115 | 0.31639229780965 | 0.843284351171898 | -4.81295117216218 |
| **Mpp2** | 0.237824197499997 | 20.64511224875 | 1.08013559159854 | 0.316869631422322 | 0.843284351171898 | -4.8138063078785 |
| **Acot9** | 5.600877445 | 19.360932495 | 1.08006411952992 | 0.316899280042392 | 0.843284351171898 | -4.81385936996195 |
| **Cspg5** | 0.348900569999998 | 26.3688963425 | 1.08003260816618 | 0.316912352552312 | 0.843284351171898 | -4.81388276384454 |
| **Gatd3** | 5.1533138425 | 17.82073170625 | 1.07968956223194 | 0.317054693568924 | 0.843284351171898 | -4.8141374120819 |
| **Rnaseh2a** | 0.382804732499999 | 24.48986490875 | 1.07948848898364 | 0.317138149460314 | 0.843284351171898 | -4.814286648498 |
| **Dmtn** | 4.62954291 | 16.0045902825 | 1.07931218510677 | 0.317211339367187 | 0.843284351171898 | -4.81441748691433 |
| **Gng7** | 0.522854427499993 | 24.01277960625 | 1.07877016139365 | 0.317436437805452 | 0.843284351171898 | -4.81481964980826 |
| **Chchd6** | 5.6409968275 | 19.51992604375 | 1.07857043561328 | 0.317519414955625 | 0.843284351171898 | -4.8149678078225 |
| **Actr2** | 5.21800521 | 18.0471039 | 1.07852940557234 | 0.317536463274943 | 0.843284351171898 | -4.81499824209133 |
| **Ly6h** | 5.4335221225 | 18.64676085875 | 1.07760771608436 | 0.31791962761137 | 0.843284351171898 | -4.81568172099319 |
| **Phb2** | 5.36980970000001 | 18.6071021675 | 1.07700165014122 | 0.318171784361419 | 0.843284351171898 | -4.81613095127329 |
| **Tmem9b** | 5.809183975 | 20.1338981275 | 1.07684455711674 | 0.318237170023287 | 0.843284351171898 | -4.81624736665344 |
| **Calm2** | 5.967200325 | 20.67913182 | 1.07678300255988 | 0.318262793374862 | 0.843284351171898 | -4.81629297940183 |
| **Npm1** | 5.21176649 | 18.068103445 | 1.07668734741711 | 0.318302615100273 | 0.843284351171898 | -4.81636385792047 |
| **Slc4a3** | -0.144663512499999 | 23.95652508375 | -1.07629107893354 | 0.318467626515641 | 0.843284351171898 | -4.81665744308143 |
| **Rpl19** | -0.221434287499999 | 29.27980858375 | -1.07617234873634 | 0.31851708076541 | 0.843284351171898 | -4.81674539414456 |
| **Ptges3** | 4.977161255 | 17.13910824 | 1.07580014253292 | 0.318672154583407 | 0.843284351171898 | -4.81702107197099 |
| **Ncdn** | 5.33790882 | 18.506653955 | 1.07514808421192 | 0.318943971106027 | 0.843284351171898 | -4.81750388157413 |
| **Rps8** | 5.50587693499999 | 19.109305895 | 1.07497398165772 | 0.319016578971086 | 0.843284351171898 | -4.81763276309504 |

| | | | | | | |
|---|---|---|---|---|---|---|
| **Fxyd1** | -5.1326543075 | 17.81668071875 | -1.0749078243479 | 0.319044172760754 | 0.843284351171898 | -4.81768173343292 |
| **Map1b** | -5.6145289025 | 19.49359771875 | -1.07453694344887 | 0.319198900433847 | 0.843284351171898 | -4.81795622859287 |
| **Asphd2** | 0.271378712499995 | 27.35221215375 | 1.074205099555 | 0.319337393535663 | 0.843284351171898 | -4.81820178174341 |
| **Tomm40** | -5.02182074750001 | 17.45983016125 | -1.07302618484996 | 0.319829798296165 | 0.843284351171898 | -4.81907375564463 |
| **Ndrg3** | -0.153224280000011 | 21.7416918425 | -1.07291650578613 | 0.319875639727606 | 0.843284351171898 | -4.81915484840801 |
| **Rtn1** | -0.379064650000004 | 23.0051447825 | -1.07258503852672 | 0.320014211858751 | 0.843284351171898 | -4.81939989193579 |
| **Tm9sf2** | 5.27634391 | 18.3573392325 | 1.07214656610072 | 0.320197592496233 | 0.843284351171898 | -4.81972396840642 |
| **Fis1** | -5.3350759725 | 18.56474303625 | -1.07156527414681 | 0.32044083443205 | 0.843284351171898 | -4.82015347558738 |
| **Glul** | 0.142005002499999 | 26.71846165875 | 1.07090217448308 | 0.320718490456393 | 0.843284351171898 | -4.82064325127551 |
| **Rogdi** | 0.340116957500005 | 20.55551160125 | 1.06946962039654 | 0.321318996805633 | 0.843284351171898 | -4.82170070997857 |
| **Scn1b** | 4.9665325225 | 17.29158978375 | 1.06859636887765 | 0.321685494935233 | 0.843284351171898 | -4.82234487695744 |
| **Cpt1a** | 5.0536016325 | 17.64127032125 | 1.06778207881008 | 0.322027550092765 | 0.843284351171898 | -4.82294525276286 |
| **Prkar1b** | -6.27948384 | 21.9315441325 | -1.06742746617659 | 0.32217660197804 | 0.843284351171898 | -4.82320661875845 |
| **Spock2** | 0.244347367499991 | 23.85298172875 | 1.06678691412813 | 0.322445981331328 | 0.843284351171898 | -4.82367859720759 |
| **Stim1** | -5.46685224750001 | 19.09826549125 | -1.06625170163132 | 0.322671199676541 | 0.843284351171898 | -4.82407282172688 |
| **Pdhx** | 0.31420662 | 21.12054957 | 1.06599992025342 | 0.322777193413817 | 0.843284351171898 | -4.82425823476024 |
| **Ssr1** | 5.1201055875 | 17.86088997875 | 1.0658516304271 | 0.322839632850345 | 0.843284351171898 | -4.82436742322788 |
| **Vps45** | 5.15516164 | 18.009032005 | 1.06578766095365 | 0.322866571057 | 0.843284351171898 | -4.8244145221532 |
| **Lsamp** | 4.9795511 | 17.3971875 | 1.06548644440767 | 0.322993440722692 | 0.843284351171898 | -4.82463627553259 |
| **Clpp** | 0.172609274999999 | 25.1734907 | 1.06533958986459 | 0.323055309034099 | 0.843284351171898 | -4.82474437445593 |

| | | | | | | |
|---|---|---|---|---|---|---|
| **Ptprd** | 5.1683800975 | 18.06253013875 | 1.06520483052238 | 0.323112090133754 | 0.843284351171898 | -4.82484356191153 |
| **Sfxn1** | -5.2476279825 | 18.37361177375 | -1.06431480420992 | 0.323487305622727 | 0.843284351171898 | -4.8254984531789 |
| **Sult4a1** | 4.910277325 | 17.1653727225 | 1.0640469504114 | 0.323600295326212 | 0.843284351171898 | -4.82569547550211 |
| **Myh9** | -5.011001475 | 17.5524469225 | -1.06312033819724 | 0.323991415705977 | 0.843284351171898 | -4.82637681258386 |
| **Rps3** | 0.476530677500005 | 23.52710132125 | 1.0611000076277 | 0.324845506238632 | 0.843284351171898 | -4.82786106123998 |
| **Ddx39b** | 5.720308125 | 20.0667593275 | 1.06105872972106 | 0.324862975186433 | 0.843284351171898 | -4.82789136773167 |
| **Ogdh** | 5.20592913 | 18.297627745 | 1.05923867024514 | 0.325633979161455 | 0.843284351171898 | -4.82922692545833 |
| **Scn2a** | 5.900472785 | 20.748391695 | 1.05882473579577 | 0.325809532317893 | 0.843284351171898 | -4.82953046763281 |
| **Hspa1b** | 0.212019224999999 | 28.01716925 | 1.05861629631886 | 0.325897961978792 | 0.843284351171898 | -4.8296832899001 |
| **Tubb2a** | -0.123263499999997 | 26.0943569875 | -1.05780903059384 | 0.326240622767702 | 0.843284351171898 | -4.83027497575904 |
| **Kcna10** | -0.158778640000005 | 22.9607971125 | -1.0575363323189 | 0.326356440391293 | 0.843284351171898 | -4.8304747854767 |
| **Adcy2** | -4.88420116749999 | 17.08788611125 | -1.05710224894935 | 0.326540867723483 | 0.843284351171898 | -4.83079277686092 |
| **Bcan** | 5.3379728175 | 18.78860325125 | 1.05637090040512 | 0.326851781582682 | 0.843284351171898 | -4.83132834502266 |
| **Rps17** | 5.10121743 | 17.890201075 | 1.05633084768345 | 0.326868815796345 | 0.843284351171898 | -4.83135766892654 |
| **Cbln3** | 0.886144182499997 | 22.45647363625 | 1.05576200880117 | 0.327110816677063 | 0.843284351171898 | -4.83177405822854 |
| **Auh** | 5.3572731525 | 18.79500116625 | 1.05478066901149 | 0.327528644449371 | 0.843284351171898 | -4.83249206274831 |
| **Pycr2** | -4.865306195 | 17.1662560375 | -1.05470710398046 | 0.327559983622333 | 0.843284351171898 | -4.8325458700515 |
| **Gapdh** | -0.143525410000002 | 20.25869178 | -1.05463099757745 | 0.327592407958792 | 0.843284351171898 | -4.83260153366704 |
| **Ndufa11** | -0.198962337499999 | 25.80177975625 | -1.05421989705488 | 0.327767597396219 | 0.843284351171898 | -4.83290216515047 |
| **Spryd7** | 0.261983772499995 | 21.08244259875 | 1.05348793403766 | 0.328079706909036 | 0.843284351171898 | -4.83343725388252 |

| | | | | | | |
|---|---|---|---|---|---|---|
| **Hadhb** | -5.0872988525 | 18.05858335375 | -1.04738279413077 | 0.330692197151148 | 0.843284351171898 | -4.83789107309852 |
| **Prepl** | 0.300052367500001 | 25.84102959125 | 1.04492400609306 | 0.33174902679234 | 0.843284351171898 | -4.83968012907082 |
| **Stxbp1** | -0.229109432500003 | 24.76244363875 | -1.04455387458387 | 0.33190834835767 | 0.843284351171898 | -4.83994920952993 |
| **Hibadh** | 5.48895487 | 19.5295882525 | 1.04450877999447 | 0.331927763297272 | 0.843284351171898 | -4.83998198848812 |
| **Fth1** | 5.25199978 | 18.6820547475 | 1.0439182047116 | 0.332182111913006 | 0.843284351171898 | -4.84041118988346 |
| **Septin7** | 5.1878902525 | 18.47157992625 | 1.04275680382945 | 0.332682755733488 | 0.843284351171898 | -4.84125478466707 |
| **Tmem109** | -5.953413375 | 21.215743285 | -1.04259861468811 | 0.332750992574095 | 0.843284351171898 | -4.8413696401764 |
| **Itpr1** | 0.15959865999999 | 30.927915335 | 1.04204352005349 | 0.332990527567213 | 0.843284351171898 | -4.84177258598613 |
| **Aldh7a1** | -0.136556742499998 | 25.78126310875 | -1.04167382617961 | 0.333150134289508 | 0.843284351171898 | -4.84204087191667 |
| **Sytl4** | -0.123170637499985 | 25.51568814375 | -1.04121031967916 | 0.333350328349083 | 0.843284351171898 | -4.8423771508676 |
| **Tf** | -0.0922942899999981 | 23.7134059475 | -1.04053926017026 | 0.33364033647686 | 0.843284351171898 | -4.84286384097853 |
| **Omp** | -4.84447014000001 | 17.2895881475 | -1.04019138590563 | 0.333790754286695 | 0.843284351171898 | -4.84311605940838 |
| **Me2** | -5.180245615 | 18.4999806625 | -1.03956444519549 | 0.33406197404145 | 0.843284351171898 | -4.84357047143314 |
| **Kif2a** | 4.57217394 | 16.32516361 | 1.03953942758826 | 0.334072800496976 | 0.843284351171898 | -4.84358860074523 |
| **Ppp2ca** | 4.89345684 | 17.473447705 | 1.03919119027007 | 0.334223530332917 | 0.843284351171898 | -4.84384092591607 |
| **Map2** | 5.171332365 | 18.50348372 | 1.03764734054372 | 0.334892415404923 | 0.843284351171898 | -4.84495890857959 |
| **Cpox** | -5.232926695 | 18.72502774 | -1.0373566624117 | 0.335018472777709 | 0.843284351171898 | -4.84516928379447 |
| **Cplx2** | -0.248891935 | 20.3373991575 | -1.03727210352578 | 0.335055150204186 | 0.843284351171898 | -4.84523047524013 |
| **Ppib** | 5.8203321975 | 20.83151657875 | 1.03679564746978 | 0.335261872630107 | 0.843284351171898 | -4.84557520469838 |
| **Grm2** | 5.3369763275 | 19.06342704125 | 1.03676872380901 | 0.33527355715641 | 0.843284351171898 | -4.84559468167354 |

| | | | | | | |
|---|---|---|---|---|---|---|
| **Epn1** | -0.179613385000003 | 29.8226560225 | -1.03658001503403 | 0.335355463432149 | 0.843284351171898 | -4.84573118721841 |
| **Surf1** | 0.127756589999997 | 25.24845161 | 1.03569390513223 | 0.335740278698138 | 0.843284351171898 | -4.84637195420346 |
| **Hnrnpdl** | -0.417207762500006 | 27.50388584125 | -1.03538419903351 | 0.335874858826892 | 0.843284351171898 | -4.84659582636092 |
| **Prkar2b** | 5.1805362575 | 18.57278064625 | 1.03484627970121 | 0.336108708710492 | 0.843284351171898 | -4.84698456022869 |
| **Faf2** | -0.179816542499996 | 24.90636908375 | -1.03450682588031 | 0.336256346010987 | 0.843284351171898 | -4.8472298032717 |
| **Mff** | -5.23478183500001 | 18.78233518 | -1.03403641053305 | 0.336461026846068 | 0.843284351171898 | -4.84756957506831 |
| **Nlgn2** | 0.333939522499993 | 22.73646733375 | 1.03380032638017 | 0.336563785825185 | 0.843284351171898 | -4.84774005628027 |
| **Sdhb** | -0.156676139999998 | 26.7390942225 | -1.03377894168223 | 0.336573095046617 | 0.843284351171898 | -4.84775549736301 |
| **Kif5c** | 0.417957125000008 | 20.9293747275 | 1.03258321796166 | 0.337093944092658 | 0.843284351171898 | -4.84861855461566 |
| **Exog** | 0.626020294999996 | 22.67768612 | 1.03170712449535 | 0.337475969397833 | 0.843284351171898 | -4.84925049550781 |
| **Ak3** | -5.0154931775 | 18.03379676625 | -1.03123776321879 | 0.33768077787216 | 0.843284351171898 | -4.84958891031837 |
| **Cacna2d1** | 5.1716632575 | 18.52368747125 | 1.03081786974889 | 0.3378640841393 | 0.843284351171898 | -4.84989157346202 |
| **Rpl10a** | 0.126208049999992 | 25.9763023225 | 1.03078166646404 | 0.337879892522553 | 0.843284351171898 | -4.84991766537639 |
| **Atp2b4** | 4.9607248675 | 17.85542223875 | 1.03061402027134 | 0.337953103876453 | 0.843284351171898 | -4.85003848117809 |
| **Serpinb1a** | -4.816158 | 17.3453735025 | -1.02973101662438 | 0.338338920101771 | 0.843284351171898 | -4.85067461507202 |
| **Eci1** | -5.69861050666667 | 18.4878337057143 | -0.968779951524447 | 0.371267665336423 | 0.850791505843268 | -4.85166456904528 |
| **Dlat** | -0.292998332500009 | 29.37627993125 | -1.02822416799663 | 0.338998121084304 | 0.843284351171898 | -4.85175935985669 |
| **Slc12a5** | -0.190641100000001 | 27.44867513 | -1.0280600221689 | 0.339069991229023 | 0.843284351171898 | -4.85187746205805 |
| **Ggt7** | 0.186672870000002 | 26.13023118 | 1.02787428325542 | 0.339151330282544 | 0.843284351171898 | -4.85201108556791 |
| **Hoxa2** | 5.535373735 | 19.895626915 | 1.02747928270116 | 0.339324360759077 | 0.843284351171898 | -4.85229520287507 |

| | | | | | | |
|---|---|---|---|---|---|---|
| **Isoc2a** | -0.116092317500001 | 27.2220499262 5 | -1.02713444672025 | 0.339475473600895 | 0.843284351171898 | -4.8525431794147 |
| **Tacc1** | -5.11616972 | 18.4539595275 | -1.02706717072904 | 0.339504961252122 | 0.843284351171898 | -4.85259155222274 |
| **Coq7** | -0.1068232125 | 26.7656100937 5 | -1.02530644847606 | 0.340277420661201 | 0.843284351171898 | -4.85385681188139 |
| **Cyp2d26** | -5.20032368 | 18.76934088 | -1.02498351444845 | 0.34041924788726 | 0.843284351171898 | -4.85408871922164 |
| **Yars1** | 5.19821451 | 18.76825396 | 1.02492335075913 | 0.340445675928804 | 0.843284351171898 | -4.8541319190582 |
| **Map4** | 0.354175732499993 | 22.4271123312 5 | 1.02444388085644 | 0.340656350050934 | 0.843284351171898 | -4.8544761375847 |
| **Fgf14** | 0.138654349999996 | 25.228178445 | 1.02426891771956 | 0.340733252659263 | 0.843284351171898 | -4.85460171997538 |
| **Spr** | -4.9547181475 | 17.9298614887 5 | -1.02403031333705 | 0.340838149966101 | 0.843284351171898 | -4.85477295920166 |
| **Stip1** | -5.35366607 | 19.361358055 | -1.02386092619918 | 0.340912632855198 | 0.843284351171898 | -4.85489450744127 |
| **Apool** | 5.711053585 | 20.644592555 | 1.02384417907275 | 0.340919997596782 | 0.843284351171898 | -4.85490652407198 |
| **Sympk** | 0.177035062500003 | 25.1319622062 5 | 1.02325179582104 | 0.34118058568864 | 0.843284351171898 | -4.85533149637481 |
| **Fxyd6** | 4.75051682 | 17.2095472575 | 1.02293847130788 | 0.341318479957283 | 0.843284351171898 | -4.85555620845356 |
| **Gk** | 4.83251844 | 17.502276435 | 1.02283352641885 | 0.341364676077005 | 0.843284351171898 | -4.85563146342549 |
| **Dctn1** | -0.124846667500005 | 26.1454615012 5 | -1.02116452307779 | 0.342100024224506 | 0.843284351171898 | -4.85682760954569 |
| **Ccdc51** | 4.6390969125 | 16.7864265987 5 | 1.02102950786732 | 0.342159565267355 | 0.843284351171898 | -4.85692431663745 |
| **Impa1** | -0.176728825000001 | 28.58934622 | -1.02083807716683 | 0.342243999257345 | 0.843284351171898 | -4.85706141796739 |
| **Cacna2d2** | 0.400984825000005 | 24.2286873325 | 1.02067070077428 | 0.342317837106663 | 0.843284351171898 | -4.85718127794859 |
| **Rack1** | 4.6675698075 | 16.9234881362 5 | 1.02001745048223 | 0.342606137573856 | 0.843284351171898 | -4.85764895387281 |
| **Acsbg1** | 5.5565835825 | 20.1736951412 5 | 1.01897692047147 | 0.343065751892621 | 0.843284351171898 | -4.85839348557376 |
| **Cpne5** | 5.1728258225 | 18.7859502087 5 | 1.01886808207224 | 0.343113855118098 | 0.843284351171898 | -4.85847133399961 |

| | | | | | | |
|---|---|---|---|---|---|---|
| **Rhoa** | 0.13671483249999 | 28.6956817412 5 | 1.01846383954926 | 0.34329256435816 4 | 0.843284351171 898 | - 4.858760427146 27 |
| **Homer1** | - 0.13754116500000 2 | 28.5458088225 | - 1.01782115404141 | 0.34357683626676 6 | 0.843284351171 898 | - 4.859219886938 98 |
| **Cep164** | 5.51123248 | 20.0219682775 | 1.01746572148308 | 0.34373413029881 8 | 0.843284351171 898 | - 4.859473905899 16 |
| **Rps2** | -5.2436982975 | 19.0663071862 5 | - 1.01700154119655 | 0.34393963511206 | 0.843284351171 898 | - 4.859805556270 64 |
| **Lman1** | -4.69274 | 17.0536492075 | - 1.01600064866485 | 0.34438308531448 3 | 0.843284351171 898 | - 4.860520340916 63 |
| **Tmx3** | 5.0505039225 | 18.1077397337 5 | 1.01576087972933 | 0.34448938278911 3 | 0.843284351171 898 | - 4.860691502387 78 |
| **Sema4a** | 4.9909779875 | 18.1801177312 5 | 1.01544206763268 | 0.34463076264288 6 | 0.843284351171 898 | - 4.860919048369 89 |
| **Rgs6** | -4.8592115625 | 17.7098372237 5 | - 1.01528824119669 | 0.34469899455161 5 | 0.843284351171 898 | - 4.861028822176 06 |
| **Rab11b** | 4.8829408825 | 17.8011354262 5 | 1.01498084040055 | 0.34483537836001 1 | 0.843284351171 898 | - 4.861248157042 94 |
| **Ttyh3** | 5.268635225 | 19.2210587925 | 1.01388535524047 | 0.34532175453314 5 | 0.843284351171 898 | - 4.862029444978 98 |
| **Dnajb11** | 4.9435647425 | 17.9860502912 5 | 1.01368079522931 | 0.34541263525557 8 | 0.843284351171 898 | - 4.862175273209 95 |
| **Scn1a** | 0.24954838250000 9 | 24.2800886512 5 | 1.0130976924399 | 0.34567179581547 7 | 0.843284351171 898 | - 4.862590853109 23 |
| **Rps9** | - 0.36259152749999 4 | 27.1924396037 5 | - 1.01256490114494 | 0.34590872882644 7 | 0.843284351171 898 | - 4.862970437636 32 |
| **Abr** | 5.3151713025 | 19.4175216687 5 | 1.01192042105189 | 0.34619550031662 | 0.843284351171 898 | - 4.863429417865 18 |
| **Sdcbp** | 0.27796802249999 8 | 28.2852379912 5 | 1.01101467401693 | 0.34659884187272 7 | 0.843284351171 898 | - 4.864074138127 06 |
| **Astn1** | 5.566939935 | 20.3741375175 | 1.01044187072715 | 0.34685410921114 | 0.843284351171 898 | - 4.864481668251 22 |
| **Lpgat1** | 0.23069865250000 3 | 23.8583155762 5 | 1.00959286854919 | 0.34723273454152 4 | 0.843284351171 898 | - 4.865085423069 5 |
| **Acot11** | 0.38388807250000 3 | 21.5529487262 5 | 1.00949505160233 | 0.34727637829048 3 | 0.843284351171 898 | - 4.865154962476 51 |
| **Dnaja2** | -5.08887521 | 18.615944095 | - 1.00790618094168 | 0.34798589926571 3 | 0.843284351171 898 | - 4.866283886490 85 |
| **Cyfip1** | 0.1379085425 | 20.4928366087 5 | 1.0078579134619 | 0.34800747119502 5 | 0.843284351171 898 | - 4.866318163021 08 |

| | | | | | | |
|---|---|---|---|---|---|---|
| **Naxe** | 0.244980442500008 | 24.87659991125 | 1.00763902812301 | 0.34810530960499 | 0.843284351171898 | -4.86647358795334 |
| **Lrrc8a** | 0.418648445000002 | 22.65945195 | 1.00702577299139 | 0.348379540014802 | 0.843284351171898 | -4.86690892547923 |
| **Cpe** | -5.30222059500001 | 19.4480866075 | -1.00481386225305 | 0.349370049257351 | 0.843284351171898 | -4.86847765262397 |
| **Isyna1** | 5.04713392 | 18.5875352925 | 1.00303332702462 | 0.35016898456177 | 0.843284351171898 | -4.86973877196196 |
| **Dpysl2** | 0.283331530000002 | 21.2129652075 | 1.00288708434652 | 0.350234667810108 | 0.843284351171898 | -4.8698422866299 |
| **Ctsd** | 4.856044485 | 17.6898482625 | 1.00262751944717 | 0.350351272162491 | 0.843284351171898 | -4.87002598915673 |
| **Copa** | 5.2635413275 | 19.40136853375 | 1.00184847370196 | 0.350701425048931 | 0.843284351171898 | -4.87057715469257 |
| **Pitpnm1** | -0.144814600000007 | 27.4049596675 | -1.00118958784877 | 0.350997783701609 | 0.843284351171898 | -4.87104308533584 |
| **Ctbp1** | -0.871286412500005 | 24.42395685875 | -1.00016892052102 | 0.351457253164551 | 0.843284351171898 | -4.87176444520976 |
| **Vma21** | 0.830793177499999 | 19.95431445875 | 0.999997409110719 | 0.351534507774632 | 0.843284351171898 | -4.87188561318697 |
| **Sv2b** | -0.152220087500002 | 24.07274015875 | -0.998194505463745 | 0.352347399170555 | 0.843284351171898 | -4.8731584726745 |
| **Prkar1a** | -0.343943117500004 | 23.76681836875 | -0.997122621267416 | 0.35283138357321 | 0.843284351171898 | -4.8739144995553 |
| **Ap2a1** | -0.100359365000003 | 30.994443695 | -0.996762132329589 | 0.352994270356296 | 0.843284351171898 | -4.87416863914833 |
| **Cep112** | 5.5540478025 | 20.51197291625 | 0.996107617239615 | 0.353290162497709 | 0.843284351171898 | -4.87462990550303 |
| **Snx3** | 0.409096242500002 | 23.63326020875 | 0.995747777054406 | 0.353452920767818 | 0.843284351171898 | -4.8748834145507 |
| **Gnb4** | -0.726269275 | 21.51467885 | -0.994789117737275 | 0.353886814197766 | 0.843284351171898 | -4.87555849431272 |
| **Hmgcl** | -5.075211725 | 18.8358241175 | -0.994596864727657 | 0.353973878656066 | 0.843284351171898 | -4.87569382468827 |
| **Lgi1** | 0.636766799999982 | 22.95551288 | 0.993676899195173 | 0.354390728674022 | 0.843284351171898 | -4.87634116181617 |
| **Pcbp1** | -0.280699052500001 | 25.09828656875 | -0.993559024967418 | 0.354444166829617 | 0.843284351171898 | -4.87642407534952 |
| **Alcam** | -4.85823809 | 17.92987019 | -0.993303980093388 | 0.354559812608778 | 0.843284351171898 | -4.87660345301202 |

| | | | | | | |
|---|---|---|---|---|---|---|
| **Tomm40l** | -0.310760264999995 | 21.6496487425 | -0.993210736393435 | 0.354602099712362 | 0.843284351171898 | -4.8766690252583 |
| **Septin3** | -5.28365229 | 19.58673996 | -0.993175272936543 | 0.354618183832776 | 0.843284351171898 | -4.87669396332064 |
| **Timm29** | 0.260627302499994 | 24.37231559125 | 0.99296229623899 | 0.354714789367534 | 0.843284351171898 | -4.87684371689301 |
| **Ank2** | 0.198959654999999 | 26.35266682 | 0.992380294264946 | 0.354978887961171 | 0.843284351171898 | -4.87725283866273 |
| **Prkcsh** | 5.15221890999999 | 19.150499455 | 0.991722653248055 | 0.355277493716579 | 0.843284351171898 | -4.87771493705382 |
| **Grik2** | 4.339833405 | 16.1297498425 | 0.991160297164779 | 0.355532989634614 | 0.843284351171898 | -4.87810991885197 |
| **Pccb** | -0.101810284999999 | 30.56443905 | -0.991124668238582 | 0.355549181776747 | 0.843284351171898 | -4.87813493844141 |
| **Rpl23** | -5.10044705 | 18.7103623375 | -0.9910495192455 | 0.355583336332057 | 0.843284351171898 | -4.8781877080968 |
| **Abcd3** | -0.0814803224999849 | 25.37532407125 | -0.990962643671595 | 0.355622823692494 | 0.843284351171898 | -4.8782487088023 |
| **Hnrnpa0** | 0.249546714999994 | 26.1809365725 | 0.990873179992778 | 0.355663490979739 | 0.843284351171898 | -4.87831152300988 |
| **Ugt8** | 5.0855953425 | 18.95895765375 | 0.989098784610294 | 0.356470820268727 | 0.843392410586336 | -4.87955657073791 |
| **Clip2** | -0.852324169999992 | 21.7158943075 | -0.988220200264243 | 0.356871092306496 | 0.843392410586336 | -4.88017249334366 |
| **Vsnl1** | -0.275387862500004 | 25.72177254375 | -0.987646312949092 | 0.357132736472631 | 0.843392410586336 | -4.8805746115734 |
| **Gltp** | -5.11749081 | 19.1440154175 | -0.985266131105607 | 0.358219487139368 | 0.843392410586336 | -4.88224069939209 |
| **Epb41l2** | -0.871483857499996 | 23.34041079875 | -0.985112005510805 | 0.358289946548931 | 0.843392410586336 | -4.88234849094877 |
| **Clic4** | 0.260775004999992 | 23.7368350125 | 0.984225839915161 | 0.358695270688983 | 0.843392410586336 | -4.88296803114243 |
| **Pclo** | 0.247836417500004 | 25.02848063875 | 0.983013146915593 | 0.359250520997251 | 0.843392410586336 | -4.88381524205347 |
| **Wdr33** | 0.165015289999992 | 32.521077535 | 0.982831614226494 | 0.359333695790768 | 0.843392410586336 | -4.88394200334569 |
| **Ywhah** | 0.305717300000008 | 21.866531685 | 0.980974596529453 | 0.360185401930514 | 0.843392410586336 | -4.88523781509079 |
| **H1-4** | -4.83644562 | 18.1554325275 | -0.979841023313993 | 0.360706073094252 | 0.843392410586336 | -4.88602799441078 |

| | | | | | | |
|---|---|---|---|---|---|---|
| **Afg3l2** | -4.94277054 | 18.209998945 | -0.979692060283893 | 0.360774537805766 | 0.843392410586336 | -4.88613178579525 |
| **Lamp2** | 0.120105464999995 | 25.0350158025 | 0.978730047977454 | 0.3612169289743 | 0.843392410586336 | -4.88680181818023 |
| **Mtnd1** | 0.358929549999999 | 21.614341185 | 0.978172931387319 | 0.36147331629487 | 0.843392410586336 | -4.88718963958281 |
| **Gad1** | 0.312867307499985 | 21.81326685625 | 0.977223183250096 | 0.361910718260765 | 0.843392410586336 | -4.88785043368417 |
| **Amph** | 0.410277760000007 | 25.2822629325 | 0.977222716204579 | 0.361910933456832 | 0.843392410586336 | -4.88785075852679 |
| **Psmd12** | -0.221433350000005 | 24.36895136 | -0.976867054278525 | 0.362074837060362 | 0.843392410586336 | -4.88809810014832 |
| **Mlec** | -0.182851602500005 | 28.17034064875 | -0.976437580787867 | 0.362272832420675 | 0.843392410586336 | -4.88839669143888 |
| **Cmtr1** | -4.63233789 | 17.444832325 | -0.975242086214934 | 0.362824417900222 | 0.843392410586336 | -4.88922738660877 |
| **Cct5** | 4.67534542 | 17.65490713 | 0.974759985977663 | 0.363047035815883 | 0.843392410586336 | -4.88956217961423 |
| **Ksr2** | 0.359092642500002 | 21.82004613625 | 0.974646450404177 | 0.363099478102482 | 0.843392410586336 | -4.88964100758271 |
| **Ldha** | 0.177213720000012 | 20.50488013 | 0.973569492425392 | 0.3635972174354 | 0.843392410586336 | -4.89038842939505 |
| **Rps3a** | 0.413065137500006 | 19.30875143375 | 0.973235210538481 | 0.363751819910448 | 0.843392410586336 | -4.89062031011045 |
| **Rab21** | 0.427887277499998 | 21.79023714375 | 0.970795310606777 | 0.36488178667146 | 0.8451629335854 | -4.89231113943073 |
| **Tmem43** | 5.50255433 | 20.88166059 | 0.968954571385699 | 0.365736056567921 | 0.846291958377326 | -4.89358482940926 |
| **Ech1** | 5.37400576583333 | 18.5610230657143 | 0.902578001707494 | 0.40264635060299 | 0.86351411562879 | -4.89410577498788 |
| **Me3** | -0.452198699999993 | 20.4459253475 | -0.967532722378324 | 0.366396975593091 | 0.846971766225713 | -4.89456753393293 |
| **Txn** | 0.140657457499994 | 23.77342987625 | 0.966478164526517 | 0.366887758917347 | 0.847257317139459 | -4.895295745931 |
| **Cct8** | -0.538475837500005 | 21.73622356875 | -0.964869005306343 | 0.367637622289652 | 0.848139994622228 | -4.89640587750652 |
| **Rtn4ip1** | 4.74113651 | 18.0713058275 | 0.96320720197985 | 0.368413251546924 | 0.849080291027725 | -4.89755098899726 |
| **Actr3** | -5.31631225 | 20.2991763425 | -0.961870962624412 | 0.36903783687187 | 0.849093688987669 | -4.89847077350978 |
| **Gh1** | 0.403432474999995 | 23.774459855 | 0.961620139875233 | 0.369155166907079 | 0.849093688987669 | -4.89864332606428 |

| | | | | | | |
|---|---|---|---|---|---|---|
| **Dnaja3** | 0.0687276099999963 | 26.409834715 | 0.958932311524186 | 0.370414275352908 | 0.850390655724049 | -4.90049045343039 |
| **Cct7** | 0.0979177824999979 | 27.4765442587 5 | 0.95884279126948 | 0.370456267448058 | 0.850390655724049 | -4.90055191189596 |
| **Rab1b** | -0.337165604999999 | 27.226234205 | -0.956900050871648 | 0.371368463972332 | 0.850791505843268 | -4.90188468458796 |
| **Tubb2b** | 4.86554769500001 | 18.6375862525 | 0.954010499279481 | 0.372728401563663 | 0.851029347266327 | -4.90386352448919 |
| **Actc1** | -0.073323685000009 | 27.2103060275 | -0.953435551145712 | 0.372999447921082 | 0.851029347266327 | -4.90425676770902 |
| **Tomm70** | 0.102878867499999 | 26.98275791875 | 0.952498696615467 | 0.373441429057767 | 0.851029347266327 | -4.9048971880934 |
| **Emilin3** | 0.134616064999996 | 24.931244055 | 0.95215777524138 | 0.37360236509268 | 0.851029347266327 | -4.90513012830665 |
| **Kcnk1** | 5.21863775249999 | 20.10723640375 | 0.952075384913736 | 0.373641266372045 | 0.851029347266327 | -4.90518641415873 |
| **Capns1** | -0.569313640000011 | 25.15870371 | -0.951191450580206 | 0.374058817661492 | 0.851029347266327 | -4.90579007005447 |
| **Ndufs7** | 0.200570904999999 | 25.5316067175 | 0.950785721065322 | 0.374250594484959 | 0.851029347266327 | -4.90606701975249 |
| **Fah** | 0.241450385000011 | 25.23276975 | 0.949871190432908 | 0.374683141904817 | 0.851029347266327 | -4.9066909732993 |
| **Kgd4** | -0.0920696075000009 | 31.15152268125 | -0.948864991379259 | 0.375159485756721 | 0.851029347266327 | -4.90737698498752 |
| **Gabrb2** | 0.195892215000004 | 26.07316267 | 0.948861410723774 | 0.375161181694779 | 0.851029347266327 | -4.90737942531925 |
| **Mapt** | 0.165690725000001 | 26.33931875 | 0.947245937172654 | 0.375926928347168 | 0.85192870697143 | -4.90847976608102 |
| **Idh2** | -0.273338895000013 | 27.1122343375 | -0.945079750828285 | 0.376955582104191 | 0.853421519052373 | -4.90995315292641 |
| **Sfxn3** | -0.635159782500004 | 22.26050137875 | -0.94228274909611 | 0.378286951899067 | 0.85453171840017 | -4.91185210763245 |
| **Pdpr** | 5.169244215 | 20.0987235 | 0.942136334652476 | 0.378356743191027 | 0.85453171840017 | -4.9119514032937 |
| **Mtco1** | 0.105176577499993 | 23.65595424375 | 0.941106584929583 | 0.37884786938338 | 0.85453171840017 | -4.9126494549956 |
| **Osbpl8** | 0.144087455000001 | 24.353340405 | 0.940939531355381 | 0.378927589043508 | 0.85453171840017 | -4.91276264751037 |
| **Glrx5** | -0.275450679999999 | 24.0046512325 | -0.939959276361774 | 0.379395632963023 | 0.854751684810249 | -4.91342656608344 |

| | | | | | | |
|---|---|---|---|---|---|---|
| **Aldh3b1** | -0.201622820000001 | 22.4529983675 | -0.937564289370979 | 0.380541013679122 | 0.85602692265566 | -4.91504662196886 |
| **Plpp3** | 4.94812383750001 | 19.33446983625 | 0.937224386698953 | 0.380703780946991 | 0.85602692265566 | -4.91527630848973 |
| **Cbr1** | 0.185405075000002 | 25.8874328075 | 0.934739292916867 | 0.38189540445986 | 0.857870202618204 | -4.91695380541364 |
| **Rhot1** | 0.224101787499993 | 26.01086545875 | 0.93338941326632 | 0.382543863747475 | 0.857982692973403 | -4.9178636874411 |
| **Htra2** | -0.365176520000006 | 23.9371634375 | -0.933086921673393 | 0.382689289583716 | 0.857982692973403 | -4.91806745341575 |
| **Pls3** | 0.156042719999995 | 28.54779834 | 0.931924578580097 | 0.38324848596392 | 0.85840219137744 | -4.91885000226487 |
| **Ap2m1** | 0.164052130000002 | 28.2481062375 | 0.930010835591403 | 0.384170519444097 | 0.859525162751165 | -4.92013692675163 |
| **Set** | 0.295924879999998 | 24.672889555 | 0.929338175497603 | 0.384495001282706 | 0.859525162751165 | -4.92058882141247 |
| **Ndrg2** | 5.43535937 | 21.4187177825 | 0.927916217856066 | 0.38518161352257 | 0.859625580223351 | -4.92154333219582 |
| **Atm** | 0.1813566975 | 25.47522908375 | 0.927701953988554 | 0.385285153858234 | 0.859625580223351 | -4.92168707011263 |
| **Cetn2** | 0.192790870000003 | 26.3295829925 | 0.92421716649595 | 0.386972076413824 | 0.862555150035451 | -4.92402150899657 |
| **Stx1a** | -0.132644792499995 | 29.48517860625 | -0.922166967983084 | 0.387967131349658 | 0.86351411562879 | -4.92539200096472 |
| **Slc25a1** | -0.0811743900000081 | 27.5485062425 | -0.921493787937074 | 0.388294275031343 | 0.86351411562879 | -4.92584152675204 |
| **Calu** | 0.153277957499995 | 30.11281367125 | 0.920517257088003 | 0.388769205313747 | 0.86351411562879 | -4.92649320282685 |
| **Rpl12** | -0.149773334999995 | 27.0516609325 | -0.918079078207125 | 0.389956901676148 | 0.86351411562879 | -4.92811813567612 |
| **Hspa2** | -0.0964782274999969 | 31.45911430125 | -0.916922280277129 | 0.390521356214497 | 0.86351411562879 | -4.92888800878803 |
| **Arrb1** | -0.340302575000003 | 21.6811652925 | -0.915380188800526 | 0.391274764029264 | 0.86351411562879 | -4.92991322098783 |
| **Cst3** | -0.148037672499996 | 24.08226585875 | -0.91419965170914 | 0.391852264471915 | 0.86351411562879 | -4.93069722822698 |
| **Ndel1** | 0.155760417500002 | 25.22585712875 | 0.913905370340779 | 0.391996321540973 | 0.86351411562879 | -4.9308925505201 |
| **Mtarc2** | 0.126585949999999 | 23.2600192525 | 0.913506767767801 | 0.392191509207558 | 0.86351411562879 | -4.93115704152701 |

| | | | | | | |
|---|---|---|---|---|---|---|
| **Slc23a2** | 0.278971814999995 | 25.21163925 | 0.913160152662409 | 0.392361298691899 | 0.86351411562879 | -4.93138696909613 |
| **Dpp6** | 5.0938855925 | 20.34900752625 | 0.912701683959335 | 0.392585964047247 | 0.86351411562879 | -4.93169099856648 |
| **Ptprz1** | -0.164377897500003 | 22.57227876625 | -0.911835205834092 | 0.3930108303874 | 0.86351411562879 | -4.93226529609934 |
| **Gnaz** | -0.258248762500003 | 29.55216434125 | -0.911681302615003 | 0.393086330735587 | 0.86351411562879 | -4.93236726138348 |
| **Adgrl1** | -0.196034945000001 | 24.344731285 | -0.911631683316688 | 0.393110674797961 | 0.86351411562879 | -4.93240013294824 |
| **Cryl1** | -0.0829052399999952 | 26.00058538 | -0.911295606091168 | 0.393275589567848 | 0.86351411562879 | -4.9326227419585 |
| **Frrs1l** | 0.252106299999998 | 25.7609292625 | 0.910986118168212 | 0.393427502505259 | 0.86351411562879 | -4.93282768664783 |
| **Abat** | -0.373073424999998 | 25.80087713 | -0.909336351358381 | 0.394238033669022 | 0.86351411562879 | -4.93391932585758 |
| **Mt-Cyb** | 0.464626069999998 | 22.623646035 | 0.907550645348896 | 0.395116754355273 | 0.86351411562879 | -4.93509930745558 |
| **Slc1a4** | -0.127905169999998 | 26.6472883225 | -0.90712052486207 | 0.395328628478644 | 0.86351411562879 | -4.93538327785369 |
| **Tspan9** | -0.119755995000002 | 25.7911401 | -0.90613574015371 | 0.395814044639569 | 0.86351411562879 | -4.93603307775866 |
| **Anp32a** | -0.103789169999999 | 29.4669530275 | -0.902284408706284 | 0.397716686915341 | 0.86351411562879 | -4.93856943601558 |
| **Psmc1** | -0.416460715000007 | 26.68268946 | -0.901612849301336 | 0.398049146670041 | 0.86351411562879 | -4.93901090153278 |
| **Ncln** | -0.106735397500003 | 30.12676689375 | -0.900838607533301 | 0.398432696018658 | 0.86351411562879 | -4.93951957195064 |
| **Negr1** | 0.322407782500001 | 25.00551536625 | 0.900336983887729 | 0.398681340269792 | 0.86351411562879 | -4.93984896529268 |
| **Rasal1** | -0.318245392499993 | 21.31758450375 | -0.900068381607032 | 0.398814528072587 | 0.86351411562879 | -4.9400252894101 |
| **Stx1b** | 0.284345752499995 | 23.17485928875 | 0.89966680919559 | 0.399013711349526 | 0.86351411562879 | -4.94028883061306 |
| **Emc2** | 0.2295088025 | 27.21509845875 | 0.899639572418865 | 0.39902722369036 | 0.86351411562879 | -4.94030670228706 |
| **Sepsecs** | 0.32370057 | 25.61460038 | 0.899068315334283 | 0.399310706227781 | 0.86351411562879 | -4.94068144759154 |
| **Gna11** | 0.303239260000009 | 28.44477974 | 0.898764615885873 | 0.39946147584911 | 0.86351411562879 | -4.94088060438256 |

| | | | | | | |
|---|---|---|---|---|---|---|
| **Tomm20** | 4.91371101 | 19.9163969675 | 0.89837201204174 | 0.399656444017026 | 0.86351411562879 | -4.94113798949662 |
| **Fyn** | -0.267731104999999 | 23.7609446675 | -0.897864362726568 | 0.399908648635253 | 0.86351411562879 | -4.9414706754934 |
| **Pdia3** | 0.165376914999996 | 29.139496175 | 0.8960581369481 | 0.400806953516156 | 0.86351411562879 | -4.94265326925085 |
| **Glud1** | 0.0928565399999925 | 26.792395165 | 0.894754940481093 | 0.4014560098287 | 0.86351411562879 | -4.94350543676035 |
| **Gls** | 0.198657742499996 | 22.67719386125 | 0.894159080940111 | 0.401753036254163 | 0.86351411562879 | -4.94389477155804 |
| **Atl2** | 0.268026397500005 | 22.45369985375 | 0.893298613119673 | 0.402182252378686 | 0.86351411562879 | -4.94445666769978 |
| **Pdxk** | 0.3010911125 | 26.33980722375 | 0.892769726910126 | 0.402446238030616 | 0.86351411562879 | -4.94480184108173 |
| **Ppm1a** | 0.214070947500005 | 25.13247747625 | 0.891491470134015 | 0.40308478936244 | 0.86351411562879 | -4.94563546890592 |
| **Tom1** | 0.124098840000009 | 26.2402847775 | 0.891200632819543 | 0.403230181144944 | 0.86351411562879 | -4.94582501949877 |
| **Alyref** | -0.162860467499989 | 25.72122870875 | -0.890027000080655 | 0.403817282354465 | 0.86351411562879 | -4.94658946469284 |
| **Ywhag** | 0.173671455000004 | 24.106650325 | 0.889544007144144 | 0.404059079137956 | 0.86351411562879 | -4.94690384795326 |
| **Srsf3** | 0.168622684999995 | 25.1136228725 | 0.887742293296184 | 0.404961998297091 | 0.86351411562879 | -4.94807549306275 |
| **Cfl1** | -0.525867470000005 | 22.31929683 | -0.886557629432089 | 0.405556495789466 | 0.86351411562879 | -4.94884492533039 |
| **Stk39** | -4.478313885 | 17.9741106575 | -0.886449904357122 | 0.405610587108114 | 0.86351411562879 | -4.94891485477352 |
| **Timm44** | 0.3808404175 | 23.65320334125 | 0.886246937871678 | 0.405712515831965 | 0.86351411562879 | -4.94904659298965 |
| **Acad9** | -0.360166037500001 | 23.47416523125 | -0.884739352329716 | 0.406470207681248 | 0.86351411562879 | -4.9500244194219 |
| **Slc12a7** | -0.688332992500001 | 21.04945094625 | -0.884183852048316 | 0.406749656788479 | 0.86351411562879 | -4.95038441110535 |
| **Kcnj16** | 0.401786487500001 | 21.96570074125 | 0.883772740590133 | 0.406956560821599 | 0.86351411562879 | -4.95065072473248 |
| **Pcp4l1** | -0.121931002499998 | 29.38448138375 | -0.883297188496738 | 0.407195993012729 | 0.86351411562879 | -4.95095866877772 |
| **Slc3a2** | 0.318019947500005 | 23.22060377125 | 0.883237231933598 | 0.407226187445245 | 0.86351411562879 | -4.95099748503796 |

| | | | | | | |
|---|---|---|---|---|---|---|
| **Cacna1a** | 0.145674185000004 | 28.698297775 | 0.882808790936207 | 0.407442000415456 | 0.86351411562879 | -4.95127480408174 |
| **Fh** | 0.231637612499998 | 23.59301604625 | 0.881892604983421 | 0.407903780696128 | 0.86351411562879 | -4.95186749601813 |
| **Pacsin1** | 0.215866402500001 | 25.26063435875 | 0.879857913820929 | 0.408930689236219 | 0.86351411562879 | -4.9531821421905 |
| **Arpc1a** | -0.162990800000006 | 22.8142645175 | -0.878915362718969 | 0.409407036961039 | 0.86351411562879 | -4.95379038096158 |
| **Nln** | -0.151858610000005 | 20.5337330625 | -0.878796875320993 | 0.409466947051379 | 0.86351411562879 | -4.953866808191 |
| **Snap91** | -0.319414940000001 | 29.178538495 | -0.878476494618432 | 0.409628971451772 | 0.86351411562879 | -4.95407342341162 |
| **Psma1** | -0.11312048500001 | 28.4016191225 | -0.877046011263761 | 0.410352975260755 | 0.86351411562879 | -4.95499527135161 |
| **Rassf4** | 0.0836909974999998 | 29.23130123875 | 0.876756762332414 | 0.410499485302656 | 0.86351411562879 | -4.95518153752511 |
| **Cnih2** | -0.420877814999997 | 22.5846747525 | -0.876140621717957 | 0.410811699818315 | 0.86351411562879 | -4.95557815915768 |
| **Fdps** | 0.123296292500001 | 24.18950581125 | 0.875994031374214 | 0.410886006556258 | 0.86351411562879 | -4.95567249185626 |
| **Atp1a2** | -0.114547957500005 | 25.16015258625 | -0.87516990199463 | 0.411303941344847 | 0.86351411562879 | -4.95620261208795 |
| **Uqcr11** | -4.5359950125 | 18.84029322375 | -0.874046315573688 | 0.411874238455882 | 0.86351411562879 | -4.95692476307668 |
| **Atp6v1b2** | 0.265815787499996 | 21.99228484625 | 0.873969649881062 | 0.411913172607682 | 0.86351411562879 | -4.9569740125982 |
| **Crip2** | -0.406788675000001 | 23.0274779025 | -0.873921842064235 | 0.411937452844923 | 0.86351411562879 | -4.95700472239708 |
| **Rps7** | -1.40568013 | 22.7500783775 | -0.873118063727307 | 0.412345825761778 | 0.86351411562879 | -4.95752085068645 |
| **Mapk1** | 0.141879360000004 | 24.7174695875 | 0.87259272620369 | 0.412612891995856 | 0.86351411562879 | -4.95785799434845 |
| **Sbf1** | -0.0831647649999994 | 29.48463859 | -0.871170785776627 | 0.413336398583534 | 0.86351411562879 | -4.95876979259139 |
| **Cyp46a1** | 0.116658052499993 | 26.21534207875 | 0.871065271764682 | 0.413390122724018 | 0.86351411562879 | -4.95883740794109 |
| **Mccc1** | -0.280556944999994 | 22.4992519625 | -0.869690449405878 | 0.414090601170684 | 0.86351411562879 | -4.95971786426801 |
| **Snx6** | 0.0917297324999957 | 28.50553043375 | 0.869576185520039 | 0.41414885809347 | 0.86351411562879 | -4.95979099406155 |

| | | | | | | |
|---|---|---|---|---|---|---|
| **Slc25a20** | 0.0668905075000019 | 29.54861077625 | 0.868662793323437 | 0.414614761748453 | 0.86351411562879 | -4.96037531575252 |
| **Idh3g** | 0.524726989999998 | 22.404867005 | 0.867843837215566 | 0.415032820030746 | 0.86351411562879 | -4.96089883610082 |
| **Slc1a3** | 0.105135282500004 | 27.04096299125 | 0.866724802565344 | 0.415604557764004 | 0.86351411562879 | -4.96161358891383 |
| **Slc17a7** | 0.127516935000003 | 24.956588985 | 0.861818599377016 | 0.418118001342921 | 0.86351411562879 | -4.96473918038592 |
| **Ak4** | -0.0887139425000036 | 29.05651640625 | -0.861689313383058 | 0.418184383392341 | 0.86351411562879 | -4.96482136550612 |
| **Fbxl8** | 0.0931150900000048 | 24.0211059675 | 0.860724193987771 | 0.41868016674887 | 0.86351411562879 | -4.96543458604271 |
| **Kras** | -0.275440730000007 | 24.349319655 | -0.86053101798449 | 0.418779452755298 | 0.86351411562879 | -4.96555726512644 |
| **Trim56** | 0.0909718949999956 | 29.3304075275 | 0.859712550831224 | 0.419200306921206 | 0.86351411562879 | -4.96607681557444 |
| **Agfg1** | 0.0767545125000026 | 27.99447026375 | 0.859432204948469 | 0.41934453067996 | 0.86351411562879 | -4.966254689848 |
| **Dhx57** | 0.228735719999996 | 25.7530424725 | 0.859129123249278 | 0.419500491314766 | 0.86351411562879 | -4.966446940755 |
| **Serpina3n** | -0.323490615000001 | 21.9788549275 | -0.859082114518118 | 0.419524684964769 | 0.86351411562879 | -4.96647675480773 |
| **Phyhipl** | 0.0731743525000148 | 27.27120547875 | 0.858752833470165 | 0.41969418204788 | 0.86351411562879 | -4.96668555843656 |
| **Snx1** | -0.511331922499998 | 22.55669027375 | -0.857064097758147 | 0.420564236563146 | 0.86351411562879 | -4.967755477188 |
| **Mblac2** | 0.122698632499997 | 25.11016028375 | 0.85569130353989 | 0.421272475993571 | 0.86351411562879 | -4.96862406437485 |
| **Kctd12** | -0.0890066299999965 | 22.730186075 | -0.854912836503155 | 0.421674478607761 | 0.86351411562879 | -4.96911614794168 |
| **Anpep** | 0.480828647500001 | 21.94176017625 | 0.854050928238451 | 0.422119893861832 | 0.86351411562879 | -4.96966058411483 |
| **Atp6v0a1** | -0.389607174999998 | 24.1536444375 | -0.85356981552761 | 0.42236867010819 | 0.86351411562879 | -4.96996430620969 |
| **Sco1** | -0.0982948424999996 | 26.71559109125 | -0.853564236104285 | 0.422371555766219 | 0.86351411562879 | -4.96996782769503 |
| **Abcb7** | 0.246936872500001 | 21.83377771625 | 0.853529538988558 | 0.422389501311448 | 0.86351411562879 | -4.96998972659239 |
| **Surf4** | 0.119276315000004 | 24.5903189075 | 0.85330385691299 | 0.422506238800156 | 0.86351411562879 | -4.97013214829451 |
| **Hnrnpu** | 0.208397974999997 | 21.1955937625 | 0.852063044618357 | 0.423148483953331 | 0.86351411562879 | -4.97091468485852 |

| | | | | | | |
|---|---|---|---|---|---|---|
| **Ap3d1** | 0.0899662775000039 | 23.83294930375 | 0.851456504689364 | 0.423462685585032 | 0.86351411562879 | -4.97129689660376 |
| **Ctnna2** | -0.311578665000003 | 23.35036064 | -0.848569805773149 | 0.42496036761114 | 0.86351411562879 | -4.97311314411381 |
| **Lnpk** | -0.101025017500003 | 26.57141777375 | -0.84756008409033 | 0.425485132576327 | 0.86351411562879 | -4.97374734132458 |
| **Brk1** | 0.221033617499998 | 25.07502322875 | 0.847502520444507 | 0.425515063165952 | 0.86351411562879 | -4.97378347937823 |
| **Galnt2** | -0.250445222500009 | 27.21877815875 | -0.845469145721181 | 0.426573301328175 | 0.86351411562879 | -4.9750588289964 |
| **Uba6** | 4.495089755 | 19.2038873625 | 0.844831182665264 | 0.426905708768773 | 0.86351411562879 | -4.97545848781585 |
| **Gjb6** | -0.172575347499997 | 27.81905139125 | -0.844702254116128 | 0.42697290893779 | 0.86351411562879 | -4.9755392287959 |
| **Hnrnpa3** | 4.5523968975 | 19.48303965875 | 0.844455409364215 | 0.427101590616697 | 0.86351411562879 | -4.97569378832696 |
| **Slc7a11** | -4.48861012999999 | 18.97583137 | -0.84380748095163 | 0.427439492151956 | 0.86351411562879 | -4.97609932013007 |
| **Slc32a1** | 0.141614122500002 | 24.52288259875 | 0.842958222677742 | 0.427882679907325 | 0.86351411562879 | -4.97663050562933 |
| **Prkcg** | 0.159152772500008 | 27.87283760625 | 0.84239235950003 | 0.428178160136308 | 0.86351411562879 | -4.97698421127913 |
| **Slc25a13** | -0.112730750000004 | 27.26597787 | -0.84203010586573 | 0.428367397150602 | 0.86351411562879 | -4.97721055173549 |
| **Arfgap1** | -0.152662905 | 22.6073350675 | -0.841493824147632 | 0.42864765447751 | 0.86351411562879 | -4.97754549165949 |
| **Vwa8** | -0.244701537499999 | 21.27894549625 | -0.84004778952404 | 0.429403997643905 | 0.86351411562879 | -4.97844782054746 |
| **Arl3** | -6.29111692 | 13.2208868225 | -0.839668451887226 | 0.429602567007417 | 0.86351411562879 | -4.9786843332461 |
| **Cadm2** | 0.243031429999995 | 22.7487380675 | 0.839140917901146 | 0.42987882101865 | 0.86351411562879 | -4.9790131099559 |
| **Adh5** | -0.154243644999998 | 25.7131574375 | -0.838936712017052 | 0.429985791758721 | 0.86351411562879 | -4.97914033575282 |
| **Rtn3** | 0.197121862499998 | 25.43049082125 | 0.838780991032095 | 0.430067377083483 | 0.86351411562879 | -4.97923733835001 |
| **Ptpra** | -0.183098569999999 | 24.2907587725 | -0.838592173584797 | 0.430166317163553 | 0.86351411562879 | -4.97935493924712 |
| **M6pr** | 0.0920618824999941 | 20.13031804875 | 0.838496941773903 | 0.430216224678883 | 0.86351411562879 | -4.97941424471264 |

| | | | | | | |
|---|---|---|---|---|---|---|
| **Stum** | -0.0940541450000012 | 23.48423517 | -0.837994802863913 | 0.430479445863389 | 0.86351411562879 | -4.97972686637726 |
| **Aak1** | 0.243496567500003 | 24.34546993375 | 0.837170664006228 | 0.430911709046949 | 0.86351411562879 | -4.98023965044381 |
| **Gpm6b** | 0.131866792499999 | 19.03419979375 | 0.837163720084285 | 0.430915352471849 | 0.86351411562879 | -4.9802439693644 |
| **Acadsb** | -0.119520712499998 | 30.34864165125 | -0.836640803920675 | 0.431189786023615 | 0.86351411562879 | -4.98056913001356 |
| **Pcca** | 0.276104782499996 | 24.20530436125 | 0.835994932373623 | 0.431528920633589 | 0.86351411562879 | -4.98097053377679 |
| **Syn1** | 0.25101071000001 | 24.208274355 | 0.835144800807513 | 0.431975598632073 | 0.86351411562879 | -4.98149852397425 |
| **Dbt** | -0.419324439999997 | 22.567094425 | -0.834526265005207 | 0.432300798499722 | 0.86351411562879 | -4.98188242033464 |
| **Nt5dc3** | 0.125509452499998 | 29.33215665125 | 0.834274968060556 | 0.43243296967344 | 0.86351411562879 | -4.98203832678429 |
| **Ap3m2** | -0.583391409999994 | 23.17378673 | -0.834206340211142 | 0.432469069925754 | 0.86351411562879 | -4.98208089778531 |
| **Adgrl3** | -0.247939600000002 | 28.839237815 | -0.833716325490128 | 0.432726894471965 | 0.86351411562879 | -4.98238478458827 |
| **Atad3** | 0.112007210000005 | 21.00671651 | 0.833070031805051 | 0.433067113906593 | 0.86351411562879 | -4.98278538109541 |
| **Slc25a10** | 0.0931675750000025 | 20.906375795 | 0.832218860203249 | 0.433515475351733 | 0.863596664365105 | -4.98331260715937 |
| **Tardbp** | -0.180008364999992 | 23.504168225 | -0.831570571389411 | 0.433857188556201 | 0.863596664365105 | -4.98371388942411 |
| **Eef2** | 0.235355179999999 | 23.68386367 | 0.830398978971013 | 0.434475221869661 | 0.863693707905613 | -4.98443848444605 |
| **Dhrs4** | 0.362223629999992 | 24.116588925 | 0.830058966882534 | 0.434654700857571 | 0.863693707905613 | -4.98464862578475 |
| **Cdc42** | 0.124097419999995 | 22.202930285 | 0.827719355363046 | 0.435891120171583 | 0.864896436805497 | -4.98609281785481 |
| **Vcp** | 0.253757667500004 | 23.98306715375 | 0.827495075136888 | 0.436009777201904 | 0.864896436805497 | -4.98623109751455 |
| **Abcb8** | -0.213978967499994 | 28.41891419875 | -0.826623154530945 | 0.436471290970401 | 0.86494729861782 | -4.98676840651248 |
| **Ptges2** | -0.0804824374999917 | 27.13950843625 | -0.826030352239941 | 0.436785263497945 | 0.86494729861782 | -4.98713346509005 |
| **Ak2** | -0.194120907500015 | 23.84184619375 | -0.823097123091679 | 0.438341181365814 | 0.867283966904745 | -4.988936847901 |

| | | | | | | |
|---|---|---|---|---|---|---|
| **Dhrs1** | -0.284718207500003 | 23.24932449375 | -0.822183143423007 | 0.43882679944812 | 0.86750079376762 | -4.98949776740015 |
| **Pcsk1n** | -0.0960026925000008 | 26.68059642375 | -0.819274517940081 | 0.440374752467083 | 0.869815542758185 | -4.99127963645757 |
| **Arcn1** | -0.218510564999995 | 24.3932324875 | -0.816473813189976 | 0.441868914528097 | 0.870480174853973 | -4.99299079879134 |
| **Gabrg2** | 0.108890485 | 23.715188895 | 0.816320776935482 | 0.441950661587016 | 0.870480174853973 | -4.99308417013417 |
| **Ptn** | -0.152268217500001 | 26.20273499875 | -0.816297588339655 | 0.441963049121988 | 0.870480174853973 | -4.99309831691401 |
| **Map1a** | 0.258566110000004 | 26.23652787 | 0.814549209455489 | 0.442897752670935 | 0.870480174853973 | -4.99416406624718 |
| **Fam234b** | -0.390991149999994 | 23.0763595175 | -0.813485707963017 | 0.443466993842665 | 0.870480174853973 | -4.99481147611532 |
| **Slc25a11** | 0.349427954999996 | 22.7788600875 | 0.813359771567655 | 0.443534435640818 | 0.870480174853973 | -4.99488809703904 |
| **Mdh2** | -0.283503765000003 | 20.3423952425 | -0.813202960267648 | 0.443618421745631 | 0.870480174853973 | -4.99498348974627 |
| **Lancl2** | -0.150491762500003 | 22.88483873375 | -0.812995007085683 | 0.4437298160504 | 0.870480174853973 | -4.9951099715781 |
| **Fkbp3** | -0.119551310000002 | 20.496180415 | -0.81076907820986 | 0.444923412855426 | 0.87176256029369 | -4.99646226539639 |
| **Rab7a** | -0.158585372499996 | 25.19400838375 | -0.809886687399857 | 0.445397196335454 | 0.87176256029369 | -4.9969975405796 |
| **Cacna1e** | -0.121615970000004 | 24.585612395 | -0.809534436571504 | 0.445586429885185 | 0.87176256029369 | -4.99721109666468 |
| **Qdpr** | -0.340385317500004 | 24.95927991625 | -0.808628014113778 | 0.446073630937234 | 0.87176256029369 | -4.99776029480884 |
| **Fech** | -0.196406592500001 | 24.81644877375 | -0.808257477716343 | 0.446272901476743 | 0.87176256029369 | -4.99798466422512 |
| **Prdx5** | 0.141984477500003 | 27.74416947625 | 0.805554371270537 | 0.447728493109821 | 0.872480017134706 | -4.99961905203047 |
| **Selenof** | 0.0950271899999962 | 29.3553922625 | 0.804664322023861 | 0.448208501574102 | 0.872480017134706 | -5.00015627520113 |
| **Gdpd1** | -0.341360587499999 | 21.88227250875 | -0.804431523564229 | 0.448334110491957 | 0.872480017134706 | -5.00029671347548 |
| **D2hgdh** | 0.178311650000001 | 27.12482014 | 0.804147192463421 | 0.448487557825721 | 0.872480017134706 | -5.00046819655824 |
| **Lpcat4** | 0.518525880000006 | 22.5934910125 | 0.803392758743479 | 0.448894887486713 | 0.872480017134706 | -5.000922975096 |
| **Syn3** | 0.0864652449999994 | 25.34969028 | 0.802618932388595 | 0.449312956288332 | 0.872480017134706 | -5.00138909882246 |

| | | | | | | |
|---|---|---|---|---|---|---|
| **Diras1** | -0.129434995 | 24.236198685 | -0.801515499747018 | 0.449909569473739 | 0.872480017134706 | -5.00205316039304 |
| **Pgrmc1** | 5.3785453475 | 11.81578814375 | 0.800657446156555 | 0.450373891449661 | 0.872480017134706 | -5.00256905774027 |
| **Kars1** | 0.446348077499994 | 24.11454701625 | 0.800298848735515 | 0.450568039817063 | 0.872480017134706 | -5.002784533833 |
| **Opalin** | -0.196630949999999 | 24.020889485 | -0.79987584548683 | 0.45079713326511 | 0.872480017134706 | -5.00303861378412 |
| **Skp1** | -0.297426542500002 | 23.14674090875 | -0.799870088772216 | 0.450800251592791 | 0.872480017134706 | -5.00304207087425 |
| **Abhd12** | 0.113510399999996 | 24.321684285 | 0.797744157562586 | 0.451952866406405 | 0.872857972209275 | -5.0043174334666 |
| **Manf** | -0.18216701250001 | 26.38164557875 | -0.797572380156775 | 0.452046088473873 | 0.872857972209275 | -5.00442036843349 |
| **Txndc15** | -0.111304672499998 | 27.44924918875 | -0.797416688628104 | 0.452130592453439 | 0.872857972209275 | -5.00451364923848 |
| **Sugt1** | -0.0966340524999936 | 31.71117934125 | -0.792743993681937 | 0.454671889847546 | 0.874685461889378 | -5.00730661629678 |
| **Dag1** | 0.381087205 | 20.6682892825 | 0.792686899404045 | 0.454703002471675 | 0.874685461889378 | -5.00734066332142 |
| **Syngr1** | 0.126705155000007 | 25.5647983225 | 0.792030691271226 | 0.455060698867367 | 0.874685461889378 | -5.00773184196802 |
| **Tspan7** | 0.0892814149999985 | 26.25007956 | 0.790851643443321 | 0.455703883704648 | 0.874685461889378 | -5.00843405792753 |
| **Map2k4** | -0.186877054999997 | 22.409131085 | -0.79018545306306 | 0.456067577288043 | 0.874685461889378 | -5.00883046423974 |
| **Bcl2l13** | 0.192632247499994 | 22.67435210875 | 0.790051470937871 | 0.456140746504742 | 0.874685461889378 | -5.00891015659513 |
| **Atp6v1h** | -0.247650627500001 | 22.37985953875 | -0.790016047714711 | 0.456160092904282 | 0.874685461889378 | -5.00893122450046 |
| **Tppp** | 0.298318927499995 | 24.08243547125 | 0.789666908706735 | 0.456350805665586 | 0.874685461889378 | -5.00913883472302 |
| **Rpl7a** | 0.172938457499999 | 24.36094657125 | 0.78941304809551 | 0.456489508502302 | 0.874685461889378 | -5.00928974388257 |
| **Akr7a2** | -0.419303682500004 | 21.87969140625 | -0.788390167737385 | 0.457048679671459 | 0.875030127802536 | -5.00989741616701 |
| **Clybl** | 0.221064587500003 | 23.94453757125 | 0.78384832388347 | 0.45953726387714 | 0.878611355526132 | -5.01258815156823 |
| **Ttc7b** | -0.28665582499999 | 22.37650218 | -0.78358664186716 | 0.459680930264431 | 0.878611355526132 | -5.01274280774079 |
| **Erp29** | -0.168666649999999 | 24.7859077675 | -0.782103665197277 | 0.460495686715822 | 0.878910964637001 | -5.01361849019824 |

| | | | | | | |
|---|---|---|---|---|---|---|
| **Nefm** | 6.1826767925 | 13.88782549125 | 0.781846213497118 | 0.460637233634044 | 0.878910964637001 | -5.01377037957649 |
| **Gabra1** | 0.220293284999997 | 20.7818850475 | 0.781221892157156 | 0.460980609974327 | 0.878910964637001 | -5.01413854812091 |
| **Efhd2** | -0.138829617500001 | 19.89174457875 | -0.77981574343084 | 0.461754636626196 | 0.879659741285411 | -5.01496691815134 |
| **Vgf** | -0.385155239999996 | 22.5286690775 | -0.777377374416555 | 0.463098977208577 | 0.879758643122545 | -5.01640058045604 |
| **Phyhip** | 0.217247557499999 | 26.24799891625 | 0.777213171591333 | 0.463189603372808 | 0.879758643122545 | -5.01649699752794 |
| **Kcnip4** | 0.108702597500002 | 23.80817401625 | 0.776121867349479 | 0.463792221195191 | 0.879758643122545 | -5.01713738321024 |
| **Rps26** | 0.148709937499998 | 25.44275778625 | 0.776047652033966 | 0.463833222407748 | 0.879758643122545 | -5.01718090747024 |
| **Map6** | -0.194492267499996 | 24.15460123625 | -0.775153834566858 | 0.464327218880647 | 0.879758643122545 | -5.01770483622537 |
| **Scrn3** | -0.496387185000003 | 23.5234249525 | -0.772228045681283 | 0.465946771898555 | 0.879758643122545 | -5.01941649878385 |
| **Lrp6** | 0.522503202500005 | 20.67847575125 | 0.772126027387953 | 0.466003313238656 | 0.879758643122545 | -5.01947608953401 |
| **Sirpa** | 0.231288017500002 | 25.77647895625 | 0.770444003426483 | 0.46693621415002 | 0.879758643122545 | -5.02045768959234 |
| **Stam** | -0.162207040000006 | 20.005775435 | -0.770212929546728 | 0.467064474370557 | 0.879758643122545 | -5.02059240751217 |
| **Map1lc3a** | -0.2405092475 | 24.87392684625 | -0.76925808647016 | 0.46759472639384 | 0.879758643122545 | -5.02114874821302 |
| **Stoml2** | 0.227328607499995 | 22.32955304125 | 0.768876381909297 | 0.467806813029178 | 0.879758643122545 | -5.02137099547332 |
| **Rps11** | -0.304312900000003 | 24.8390546875 | -0.767886347297755 | 0.468357212454107 | 0.879758643122545 | -5.02194703377343 |
| **Ddost** | 0.151647334999993 | 20.56473735 | 0.766652541127253 | 0.469043752530046 | 0.879758643122545 | -5.0226640807461 |
| **Rpl14** | 0.109919717499995 | 30.69485304375 | 0.765928921795793 | 0.469446722984666 | 0.879758643122545 | -5.02308419710171 |
| **Hnrnpl** | -0.204594262499995 | 23.54155649125 | -0.765572025580736 | 0.469645558870365 | 0.879758643122545 | -5.02329128629845 |
| **Srgap2** | -0.460139689999998 | 23.02698161 | -0.76539176976602 | 0.469746005741174 | 0.879758643122545 | -5.02339585059631 |
| **Psmb4** | 0.351879482499992 | 22.90884762875 | 0.765012086165573 | 0.46995763087254 | 0.879758643122545 | -5.02361603644514 |

| | | | | | | |
|---|---|---|---|---|---|---|
| **Pde2a** | -0.0961358099999963 | 24.522209995 | -0.763985207861886 | 0.470530309668973 | 0.879758643122545 | -5.02421110673566 |
| **Aifm1** | -0.192906399999998 | 25.2358040375 | -0.762386601816929 | 0.47142277961577 | 0.879758643122545 | -5.02513622125624 |
| **Prkar2a** | -0.236700989999999 | 25.8074511725 | -0.761735238299079 | 0.471786752720614 | 0.879758643122545 | -5.02551272233549 |
| **Pdzd8** | -0.132986889999998 | 24.7111267125 | -0.761248697031766 | 0.472058749944886 | 0.879758643122545 | -5.02579378520578 |
| **Wbp2** | -0.0967829225000045 | 27.0422325112 5 | -0.761141255906719 | 0.472118828451325 | 0.879758643122545 | -5.02585583195523 |
| **Stx6** | 0.154213779999999 | 31.806959645 | 0.760397376627555 | 0.472534930385732 | 0.879758643122545 | -5.026285226843 |
| **Selenom** | -0.0753402400000027 | 21.6261005075 | -0.758823600083801 | 0.473416070127192 | 0.879758643122545 | -5.02719256315837 |
| **Dguok** | 0.159479895 | 24.1776221475 | 0.758696829722105 | 0.473487095888112 | 0.879758643122545 | -5.0272655852023 |
| **Lman2** | 0.343254722500003 | 21.4649375587 5 | 0.758590699651243 | 0.473546563047882 | 0.879758643122545 | -5.02732671055563 |
| **Ahcyl1** | 0.227061277499999 | 23.6857403387 5 | 0.75811028316206 | 0.473815814970434 | 0.879758643122545 | -5.02760331966273 |
| **Cpne6** | 0.0956695274999966 | 27.0515342137 5 | 0.75808616536801 | 0.473829334648364 | 0.879758643122545 | -5.02761720225362 |
| **Cmtm4** | -0.0996866900000057 | 27.9464734075 | -0.757145637870354 | 0.474356768696722 | 0.879758643122545 | -5.02815830929993 |
| **Nudc** | 0.103846202500005 | 26.3150097737 5 | 0.757124063219565 | 0.474368872108709 | 0.879758643122545 | -5.02817071538123 |
| **Eif2s2** | 0.195303947500008 | 25.3693737612 5 | 0.756072682957251 | 0.474958951446403 | 0.879758643122545 | -5.02877494790827 |
| **Cyb5r1** | -0.0855416824999935 | 28.0725326212 5 | -0.755886090455731 | 0.475063726980547 | 0.879758643122545 | -5.0288821130889 |
| **Mpst** | -0.115394892499999 | 24.5600093062 5 | -0.755204843431 | 0.475446393984086 | 0.879758643122545 | -5.02927319189589 |
| **Gucy1b1** | -4.78151436 | 11.1381484775 | -0.751997428138447 | 0.47725084843777 | 0.879758643122545 | -5.03111064879797 |
| **Trim9** | 0.115972949999993 | 23.7287863175 | 0.751434490218495 | 0.477568026934536 | 0.879758643122545 | -5.03143249558474 |
| **Ryr2** | -0.278921982500002 | 23.1626681337 5 | -0.748888122114521 | 0.479004512976022 | 0.879758643122545 | -5.03288590185927 |
| **Icam5** | 0.244405945000004 | 27.9459921825 | 0.748881481239472 | 0.479008263101522 | 0.879758643122545 | -5.0328896871236 |
| **Mthfd1l** | -5.612615315 | 13.1394135 | -0.747864036689857 | 0.479583051045945 | 0.879758643122545 | -5.03346930601007 |

| | | | | | | |
|---|---|---|---|---|---|---|
| **Pafah1b1** | -5.52486368 | 12.9077003475 | -0.746483257958603 | 0.480363840093904 | 0.879758643122545 | -5.03425489425461 |
| **Csrp1** | 0.21473661000001 | 22.64940083 | 0.746333118653942 | 0.480448790841778 | 0.879758643122545 | -5.03434024485826 |
| **Aldh3a2** | 0.19946191 | 23.554847335 | 0.746195072696532 | 0.480526907924195 | 0.879758643122545 | -5.03441870847145 |
| **Erc2** | -5.34685515 | 12.549121795 | -0.745286016762587 | 0.481041535157316 | 0.879758643122545 | -5.03493511232088 |
| **Ap2s1** | 0.272306925000002 | 25.78365158 | 0.744182525270478 | 0.481666731605844 | 0.879758643122545 | -5.03556128605188 |
| **Hadha** | 0.638826394999999 | 21.102302015 | 0.743249438338431 | 0.482195808382899 | 0.879758643122545 | -5.03609017972753 |
| **Tmem126a** | 0.251093245000003 | 24.8877796425 | 0.743218370220064 | 0.482213431254053 | 0.879758643122545 | -5.03610778058604 |
| **Dtd1** | 0.218004867500014 | 24.14391674875 | 0.743055265782225 | 0.482305956603684 | 0.879758643122545 | -5.03620017354154 |
| **Mtpn** | -0.1751524375 | 24.64939246375 | -0.739939798132345 | 0.484075572001839 | 0.879758643122545 | -5.03796182999985 |
| **Ephx1** | -0.260422845000001 | 27.883650585 | -0.738867917497531 | 0.484685413175565 | 0.879758643122545 | -5.0385665456255 |
| **Slc1a6** | -0.146164507499996 | 23.04980065375 | -0.738767585207047 | 0.48474252299638 | 0.879758643122545 | -5.03862311308888 |
| **Pdia4** | -0.257243527500005 | 23.77891955875 | -0.737669319359884 | 0.485367957226381 | 0.879758643122545 | -5.03924190983692 |
| **Ppme1** | 0.131578662500004 | 26.67526044375 | 0.737083604968719 | 0.485701726678588 | 0.879758643122545 | -5.03957161435322 |
| **Adcy1** | -0.0895133274999935 | 27.33209795125 | -0.736941035807464 | 0.485782992908765 | 0.879758643122545 | -5.03965183584474 |
| **Brinp1** | 0.165319997499999 | 25.75513985625 | 0.735994692406814 | 0.486322650533966 | 0.879758643122545 | -5.04018400988461 |
| **Mpc2** | 0.215393850000002 | 24.0865354675 | 0.735981799267016 | 0.486330005677775 | 0.879758643122545 | -5.04019125648304 |
| **Arl8a** | 0.0632827850000055 | 21.8604284675 | 0.735893554947426 | 0.48638034836653 | 0.879758643122545 | -5.04024085150183 |
| **Tpd52l2** | 0.275653529999989 | 24.759416935 | 0.734345791517872 | 0.487263899814861 | 0.879758643122545 | -5.04110993997796 |
| **Rps15a** | -5.0205918775 | 11.95910895375 | -0.734185527888278 | 0.487355448437844 | 0.879758643122545 | -5.04119984514599 |
| **Slc1a2** | 0.4896788 | 23.3748889675 | 0.733654102287281 | 0.487659100682296 | 0.879758643122545 | -5.04149785183691 |

| | | | | | | |
|---|---|---|---|---|---|---|
| **Hdgf** | -0.262444642500004 | 19.02759058375 | -0.733441642223471 | 0.487780533844614 | 0.879758643122545 | -5.04161694369088 |
| **C2cd2l** | -0.155089922500004 | 24.40518778375 | -0.730888594323639 | 0.48924132042357 | 0.879758643122545 | -5.04304583025156 |
| **Mfn2** | 0.20585234 | 20.8720334975 | 0.729369788489042 | 0.49011171776521 | 0.879758643122545 | -5.04389395054421 |
| **Hsd17b12** | 0.1528746125 | 25.29653883625 | 0.729147961355215 | 0.490238928382208 | 0.879758643122545 | -5.04401770140008 |
| **Hyou1** | -0.262946812499997 | 24.98734521125 | -0.72908897188488 | 0.490272760601808 | 0.879758643122545 | -5.04405060473679 |
| **Smap2** | 0.0998966650000064 | 22.71677763 | 0.728138996780323 | 0.490817812814685 | 0.879758643122545 | -5.04458018625652 |
| **Fundc1** | -0.0887706250000022 | 22.607822345 | -0.727269261239971 | 0.491317179145481 | 0.879758643122545 | -5.04506454332191 |
| **Slc25a23** | -5.8117869275 | 13.97314587625 | -0.726616483568233 | 0.491692198114595 | 0.879758643122545 | -5.04542776616333 |
| **Vars1** | -5.3567335275 | 12.87820882375 | -0.726516906980876 | 0.491749421237555 | 0.879758643122545 | -5.0454831498402 |
| **Matr3** | 5.4314638625 | 13.04495122625 | 0.726458769875618 | 0.491782832601482 | 0.879758643122545 | -5.04551548235643 |
| **Atp6v0c** | 0.155231872499993 | 24.42006099375 | 0.726313238368167 | 0.491866476068438 | 0.879758643122545 | -5.0455964093619 |
| **Chmp4b** | 0.299769272499997 | 22.94669184375 | 0.725151704922651 | 0.492534398567972 | 0.879758643122545 | -5.04624183947083 |
| **Khsrp** | 0.19005774999999 | 27.9340516725 | 0.724986875421684 | 0.492629229874608 | 0.879758643122545 | -5.0463333620863 |
| **Rpsa** | -5.6514827725 | 13.62487582875 | -0.724717822341048 | 0.492784050024407 | 0.879758643122545 | -5.04648271900768 |
| **Prnp** | 0.0963665299999974 | 31.161793905 | 0.723839976345149 | 0.493289409003657 | 0.879758643122545 | -5.04696971461596 |
| **S1pr1** | 0.134077570000002 | 22.8347195325 | 0.723236008734254 | 0.493637300141726 | 0.879758643122545 | -5.04730449290284 |
| **Cct6a** | 5.27068082 | 12.73152598 | 0.72323075095019 | 0.493640329386643 | 0.879758643122545 | -5.04730740628164 |
| **Actbl2** | 4.687012805 | 11.319388925 | 0.722833300293461 | 0.49386935402853 | 0.879758643122545 | -5.0475275866531 |
| **Ilvbl** | -0.144349814999995 | 29.9651765825 | -0.720961960638305 | 0.494948625626369 | 0.879758643122545 | -5.04856294442807 |
| **Tmod2** | 5.624654035 | 13.6576407375 | 0.719256934162753 | 0.495933330057234 | 0.879758643122545 | -5.04950437357349 |

| | | | | | | |
|---|---|---|---|---|---|---|
| **Tmem11** | -0.0806211250000004 | 22.8103856575 | -0.718333012075519 | 0.49646746119439 | 0.879758643122545 | -5.05001375377804 |
| **Ergic1** | 5.4183402225 | 13.18985017125 | 0.717296024952558 | 0.497067406955874 | 0.879758643122545 | -5.05058483025662 |
| **Sdc4** | 0.425333700000003 | 22.053141745 | 0.716530449660073 | 0.497510633691966 | 0.879758643122545 | -5.05100600413395 |
| **Rcn2** | 6.614917965 | 16.11847166 | 0.716523519452255 | 0.49751464709264 | 0.879758643122545 | -5.05100981503617 |
| **Tmed9** | -0.0932891599999905 | 28.6384753775 | -0.71600549095608 | 0.497814706333672 | 0.879758643122545 | -5.05129459186422 |
| **Rala** | 3.7740192075 | 18.63709505125 | 0.715983226564651 | 0.497827605267003 | 0.879758643122545 | -5.05130682752428 |
| **Tmem30a** | -5.483069265 | 13.37776321 | -0.715566622754269 | 0.498069006216763 | 0.879758643122545 | -5.05153571949741 |
| **Snta1** | 0.170690740000012 | 24.425687545 | 0.715039282075457 | 0.498374683707333 | 0.879758643122545 | -5.05182529609702 |
| **Syp** | 0.2007178675 | 20.03807069625 | 0.712656255266802 | 0.499757558583385 | 0.879758643122545 | -5.05313169162299 |
| **Acaa2** | -0.100956609999997 | 22.961070525 | -0.712306863056928 | 0.499960522478161 | 0.879758643122545 | -5.05332293008281 |
| **Atp2b1** | 0.117987205000006 | 26.7767393425 | 0.711129436196066 | 0.500644893339283 | 0.879758643122545 | -5.05396682207216 |
| **Ywhaz** | 4.997641935 | 12.2705683875 | 0.710732568291477 | 0.500875707875878 | 0.879758643122545 | -5.05418365720735 |
| **Atp2a2** | -0.0708701899999973 | 26.00111962 | -0.708607492962991 | 0.502112813644474 | 0.879758643122545 | -5.05534303023678 |
| **Prdx3** | 0.100698167499999 | 23.51795308375 | 0.708293057581487 | 0.5022960303666 | 0.879758643122545 | -5.05551433318957 |
| **Wasf3** | 5.3550707425 | 13.21895743625 | 0.706580107933891 | 0.503294905550513 | 0.879758643122545 | -5.05644643926559 |
| **Acadvl** | 0.242639354999998 | 23.7727224725 | 0.705717200956149 | 0.503798583622553 | 0.879758643122545 | -5.05691528712409 |
| **Sec61b** | 5.49940716750001 | 13.59481646375 | 0.705584924533949 | 0.503875822247326 | 0.879758643122545 | -5.05698711578049 |
| **Vcan** | 5.0319681625 | 12.45421530125 | 0.704677210311438 | 0.504406061026975 | 0.879758643122545 | -5.05747972226205 |
| **Tm9sf3** | 0.348872582499997 | 21.64999559125 | 0.703732512678313 | 0.504958288702067 | 0.879758643122545 | -5.05799184298793 |
| **Krt1** | 0.291057350000003 | 23.5843423525 | 0.703673207636451 | 0.504992968857966 | 0.879758643122545 | -5.05802397332395 |

| | | | | | | |
|---|---|---|---|---|---|---|
| **Prxl2b** | 4.64641583 | 11.515273075 | 0.703505104280612 | 0.50509128004896 | 0.879758643122545 | -5.05811503634176 |
| **Mtfr1l** | 0.131438987500001 | 20.30200602125 | 0.703112513676581 | 0.505320925606584 | 0.879758643122545 | -5.05832763597324 |
| **Ezr** | -0.581958449999995 | 22.98293799 | -0.70231738691389 | 0.505786242042068 | 0.879758643122545 | -5.0587579204612 |
| **Pck2** | 4.98088031250001 | 12.36035271125 | 0.702313689085457 | 0.505788406699113 | 0.879758643122545 | -5.05875992060817 |
| **Emc10** | 0.37426709 | 24.54323809 | 0.701705777045773 | 0.506144351632804 | 0.879758643122545 | -5.05908862041844 |
| **Psmb3** | 5.43478238 | 13.5062134375 | 0.701609542864423 | 0.506200713607988 | 0.879758643122545 | -5.05914063293281 |
| **Tecr** | 0.3316525525 | 23.94763912125 | 0.701413484083256 | 0.506315552971052 | 0.879758643122545 | -5.05924658024483 |
| **Asap2** | -0.0925163300000023 | 24.4116976725 | -0.701062670023901 | 0.506521080762309 | 0.879758643122545 | -5.05943609391227 |
| **Nptn** | 5.36435206 | 13.34716443 | 0.700409142640334 | 0.506904100264875 | 0.879758643122545 | -5.05978892743616 |
| **Ahcyl2** | 0.1444142625 | 27.34733318625 | 0.699183634028374 | 0.507622852397492 | 0.879758643122545 | -5.06044983385844 |
| **Txn2** | -5.14626776 | 12.8278413 | -0.69913771853057 | 0.507649794328766 | 0.879758643122545 | -5.06047457706535 |
| **Arhgap1** | 0.180913867500006 | 25.45939275875 | 0.699028929056196 | 0.507713632647011 | 0.879758643122545 | -5.06053319678783 |
| **Atp6v1d** | 4.762589265 | 11.914008265 | 0.696391407465853 | 0.50926293536192 | 0.879758643122545 | -5.06195207480582 |
| **Nrxn3** | 5.2616402125 | 13.18303586625 | 0.695444637679184 | 0.509819820083423 | 0.879758643122545 | -5.0624603125877 |
| **Bdh1** | 0.273894487499994 | 22.29056035375 | 0.695434486350428 | 0.509825793167153 | 0.879758643122545 | -5.06246575883714 |
| **Dynlrb1** | 5.372195235 | 13.4842115 | 0.694074557207859 | 0.51062638921288 | 0.879758643122545 | -5.06319477219734 |
| **Pygm** | 0.101859637500002 | 25.72121157875 | 0.694052012237856 | 0.510639668357746 | 0.879758643122545 | -5.0632068478273 |
| **Inpp4a** | -4.7406239625 | 11.86702418875 | -0.694012650765033 | 0.510662853073779 | 0.879758643122545 | -5.06322793000079 |
| **Rdx** | 0.225915082500006 | 24.11819417125 | 0.693919142830492 | 0.51071793388813 | 0.879758643122545 | -5.06327800926792 |
| **Tpm1** | -0.14824011 | 19.9555236725 | -0.692977552486803 | 0.511272790676409 | 0.879758643122545 | -5.06378197642278 |

| | | | | | | |
|---|---|---|---|---|---|---|
| **Cept1** | 0.126321700000002 | 24.437369845 | 0.69294747197366 | 0.511290522803951 | 0.879758643122545 | -5.06379806703209 |
| **Rpl18a** | 0.161737069999997 | 27.5281823025 | 0.692859348924945 | 0.511342472639183 | 0.879758643122545 | -5.06384520229875 |
| **Slc17a6** | -0.224635752499992 | 26.72669448625 | -0.692205570888957 | 0.511727990541609 | 0.879758643122545 | -5.06419473950434 |
| **Psip1** | 0.1351020725 | 29.53482444875 | 0.691359855878462 | 0.512226966651246 | 0.879758643122545 | -5.06464648711387 |
| **Vamp4** | 0.153595052499998 | 20.21335615875 | 0.690211492833485 | 0.512905007266905 | 0.879758643122545 | -5.06525916162008 |
| **Rpl35** | 0.289010295000004 | 33.20814492 | 0.689120561346623 | 0.513549671743982 | 0.879758643122545 | -5.06584040970291 |
| **Dmxl2** | 4.5470358275 | 11.49412515125 | 0.688766742209101 | 0.513758865844096 | 0.879758643122545 | -5.06602875998206 |
| **Dlg3** | 0.399565862499998 | 20.53124854875 | 0.688573599336276 | 0.513873083835042 | 0.879758643122545 | -5.06613154266789 |
| **Gars1** | -5.1510892475 | 13.02672368625 | -0.688471895630096 | 0.513933234423682 | 0.879758643122545 | -5.06618565553843 |
| **Arxes2** | -0.1518134875 | 24.32532664625 | -0.68778626172183 | 0.514338856492989 | 0.879758643122545 | -5.06655028273426 |
| **Lap3** | 5.32130209 | 13.494620085 | 0.686442417000992 | 0.51513447220517 | 0.879758643122545 | -5.06726407439596 |
| **Rftn2** | 0.142598414999995 | 21.17001757 | 0.686357330202152 | 0.515184873886592 | 0.879758643122545 | -5.06730922958909 |
| **Rplp0** | -0.296833989999996 | 21.54129467 | -0.685873502257015 | 0.515471532325212 | 0.879758643122545 | -5.06756590617731 |
| **Enpp6** | -0.126959494999994 | 20.0773641025 | -0.685308823595572 | 0.515806222282073 | 0.879758643122545 | -5.06786528400334 |
| **Ensa** | 0.0964386800000021 | 25.442114215 | 0.684693662712931 | 0.516170991672573 | 0.879758643122545 | -5.0681911918817 |
| **Gabbr1** | 4.6976635375 | 11.94968859625 | 0.684184128170212 | 0.516473253111206 | 0.879758643122545 | -5.06846095459387 |
| **Atp5mg** | 0.142416569999995 | 24.23207868 | 0.684112104501495 | 0.51651598745977 | 0.879758643122545 | -5.06849907253031 |
| **Camk2d** | -0.068681049999995 | 24.056473605 | -0.684064371193415 | 0.516544310669854 | 0.879758643122545 | -5.06852433314539 |
| **Abce1** | 0.148356317499999 | 28.03477761375 | 0.683047783961075 | 0.517147752197487 | 0.879758643122545 | -5.06906196456365 |
| **Grm3** | -5.3748671375 | 13.71913529625 | -0.681823250634808 | 0.517875226938022 | 0.879758643122545 | -5.06970868263929 |

| | | | | | | |
|---|---|---|---|---|---|---|
| **Eef1a1** | 0.186643480000001 | 26.0484833775 | 0.681269149446903 | 0.518204623590117 | 0.879758643122545 | -5.07000100353105 |
| **Esyt1** | 0.158849005 | 24.0164282575 | 0.680388007838261 | 0.518728711037591 | 0.879758643122545 | -5.07046544785839 |
| **Grhpr** | 0.0992193174999976 | 28.00928115125 | 0.680246505688456 | 0.518812905472257 | 0.879758643122545 | -5.07053998588738 |
| **Rpl7** | 0.107642092500004 | 26.11462294875 | 0.67950023535072 | 0.519257083760532 | 0.879758643122545 | -5.070932878377 |
| **Ddx1** | -0.232934765000003 | 22.939813345 | -0.679173120415296 | 0.519451858062987 | 0.879758643122545 | -5.07110498228 |
| **Slc24a4** | -0.20131301 | 25.64635964 | -0.678973892670841 | 0.519570507173471 | 0.879758643122545 | -5.07120976730384 |
| **Ptrh2** | -0.316709240000002 | 22.0701594025 | -0.67858095381445 | 0.519804570526759 | 0.879758643122545 | -5.0714163603544 |
| **Atp6v1f** | -0.273117749999997 | 25.26032533 | -0.677852569354295 | 0.520238627356978 | 0.879758643122545 | -5.0717990535063 |
| **Dlg1** | 0.107767602500008 | 28.31611662875 | 0.67686933849133 | 0.520824916814044 | 0.879758643122545 | -5.07231509646158 |
| **Vapb** | 0.301513790000001 | 20.318345985 | 0.676373269454126 | 0.521120876388213 | 0.879758643122545 | -5.07257521695828 |
| **Dnajb2** | 0.107930695 | 27.590014 | 0.676353605541807 | 0.521132610267655 | 0.879758643122545 | -5.07258552469988 |
| **Hsd17b10** | 5.3585923 | 13.7819283475 | 0.676285865075663 | 0.52117303374493 | 0.879758643122545 | -5.07262103204761 |
| **Numbl** | 5.3023505425 | 13.64230492375 | 0.675941533079387 | 0.521378541350009 | 0.879758643122545 | -5.07280147354246 |
| **Ethe1** | 5.121570535 | 13.1867983 | 0.675383827080268 | 0.521711505997433 | 0.879758643122545 | -5.07309356655874 |
| **Pgs1** | 0.241558665000003 | 27.0999581325 | 0.673994289525833 | 0.522541681608662 | 0.879758643122545 | -5.0738204427047 |
| **Agk** | -0.125970247499993 | 27.67364614125 | -0.673899808845025 | 0.522598159221451 | 0.879758643122545 | -5.07386982051628 |
| **Nr3c2** | -0.308422742499989 | 23.53289636625 | -0.673872063038774 | 0.52261474553604 | 0.879758643122545 | -5.07388432001846 |
| **Cyp4a10** | -0.0839964649999985 | 32.175448865 | -0.672999352848405 | 0.523136617809236 | 0.879758643122545 | -5.07434012800854 |
| **Oat** | -4.72645281 | 12.20258848 | -0.672663374428164 | 0.523337617574488 | 0.879758643122545 | -5.07451547391095 |
| **Atp5mk** | -0.279082895000009 | 22.9482319075 | -0.672509563174823 | 0.523429651770381 | 0.879758643122545 | -5.07459572287977 |
| **Camk2a** | 0.0823493650000006 | 29.6096687875 | 0.672149672278945 | 0.523645035377048 | 0.879758643122545 | -5.07478343086998 |

| | | | | | | |
|---|---|---|---|---|---|---|
| **Tpm3** | 0.0815607050000082 | 27.8011376025 | 0.67118697377982 | 0.524221456191677 | 0.879758643122545 | -5.07528512938964 |
| **Gria1** | 0.188025017499996 | 23.18803494375 | 0.671109653611496 | 0.524267769442291 | 0.879758643122545 | -5.0753253976076 |
| **Nap1l1** | -0.0875512450000038 | 23.4233274475 | -0.671061250597714 | 0.524296763205425 | 0.879758643122545 | -5.07535060383224 |
| **Sort1** | 0.189977429999999 | 28.0385480225 | 0.670548068362563 | 0.524604225465514 | 0.879758643122545 | -5.07561775316535 |
| **Olfm1** | 0.135332565000006 | 24.9413346 | 0.66932356997641 | 0.525338317582394 | 0.879758643122545 | -5.07625450026117 |
| **Sdhd** | 0.135797727499998 | 23.98506820125 | 0.668688793671424 | 0.525719123695602 | 0.879758643122545 | -5.07658420235702 |
| **Ncs1** | 0.3023460625 | 22.59575119625 | 0.66811865772833 | 0.526061299995683 | 0.879758643122545 | -5.07688010568311 |
| **Atp1a1** | 0.118200294999994 | 21.266850145 | 0.668042566571502 | 0.526106977947609 | 0.879758643122545 | -5.07691958127874 |
| **Timm13** | -0.0964577475000041 | 25.12457529625 | -0.66804126494216 | 0.52610775934481 | 0.879758643122545 | -5.07692025652259 |
| **Plp2** | -0.0894610324999938 | 26.52937141875 | -0.667514384112831 | 0.52642411774609 | 0.879758643122545 | -5.07719349445785 |
| **Gna13** | 0.121404847499996 | 25.40133643375 | 0.667139960598577 | 0.526649008050007 | 0.879758643122545 | -5.07738755829199 |
| **Hp1bp3** | -0.0546208525000011 | 29.56794564375 | -0.666528781575717 | 0.527016230947272 | 0.879758643122545 | -5.07770413535794 |
| **Grpel1** | 0.171865047499999 | 23.71768365125 | 0.665186200984434 | 0.527823477220047 | 0.880079062982755 | -5.07839870273351 |
| **Wasf1** | -4.6150170275 | 12.06491868875 | -0.664458157347249 | 0.528261548988197 | 0.880079062982755 | -5.07877485284671 |
| **Fam162a** | -0.088803277500002 | 29.52317799125 | -0.664306845849117 | 0.528352623420509 | 0.880079062982755 | -5.07885298561073 |
| **Pygb** | -4.4642817875 | 11.68775939875 | -0.662919997928198 | 0.529187826831791 | 0.880470079103157 | -5.07956841256191 |
| **Impact** | 5.0373484675 | 13.21712429875 | 0.661870636299812 | 0.529820335494977 | 0.880470079103157 | -5.0801089023892 |
| **Slc8a2** | -0.0703835774999995 | 27.66836882375 | -0.661238917759142 | 0.53020133579608 | 0.880470079103157 | -5.08043392968304 |
| **Sdr39u1** | -4.41899038250001 | 11.61954164625 | -0.660310592830648 | 0.530761535774386 | 0.880470079103157 | -5.0809110884556 |
| **Mtx1** | 0.322287425000003 | 21.8481642675 | 0.660255972960693 | 0.530794507805759 | 0.880470079103157 | -5.08093914538964 |

| | | | | | | |
|---|---|---|---|---|---|---|
| **Slc27a1** | 0.146339452499994 | 21.79132651125 | 0.660006612269286 | 0.530945054079371 | 0.880470079103157 | -5.08106721112472 |
| **Pigk** | -0.223798372500003 | 21.98826901875 | -0.659486860069346 | 0.531258929393842 | 0.880470079103157 | -5.08133401192713 |
| **Reep1** | -4.8647103375 | 12.83562915875 | -0.657888414556 | 0.532224948106654 | 0.881332095566797 | -5.08215341563567 |
| **Add3** | 0.208877939999997 | 24.05477486 | 0.657362366983051 | 0.532543104126621 | 0.881332095566797 | -5.08242271259143 |
| **Cpt2** | -0.0823491299999901 | 28.0856714575 | -0.654914737008244 | 0.534025002185613 | 0.882043269327246 | -5.08367331397233 |
| **Nptxr** | 0.337249849999996 | 22.145518065 | 0.6548632490434 | 0.534056202725898 | 0.882043269327246 | -5.08369957893541 |
| **Ndufaf3** | 0.2058077125 | 24.55257676625 | 0.654315941514026 | 0.534387928844011 | 0.882043269327246 | -5.08397866231682 |
| **Add1** | 0.101157737499999 | 27.11634297375 | 0.653260169531496 | 0.535028200248652 | 0.882043269327246 | -5.08451646260743 |
| **Cep135** | 0.0716021124999955 | 24.84744842875 | 0.653061555404334 | 0.535148702793322 | 0.882043269327246 | -5.0846175523607 |
| **Farp1** | 0.142644607500003 | 19.43583595125 | 0.65286688315252 | 0.535266830107561 | 0.882043269327246 | -5.08471661046119 |
| **Ncl** | 0.136075587499999 | 24.07285603375 | 0.65166157084166 | 0.535998575516524 | 0.882618639340914 | -5.08532936962715 |
| **Camk2b** | -0.177895975000006 | 20.3124536125 | -0.650727365675881 | 0.536566158755653 | 0.882888125380001 | -5.08580364075122 |
| **Pgd** | 4.4170468975 | 11.78708030375 | 0.649965859236766 | 0.537029093329958 | 0.882888125380001 | -5.08618980930891 |
| **Api5** | -0.134538454999994 | 26.9162052175 | -0.649029401916542 | 0.537598723267763 | 0.882888125380001 | -5.08666417005271 |
| **Pakap** | 0.610917942499995 | 22.84948195875 | 0.648874426218787 | 0.537693028243997 | 0.882888125380001 | -5.08674261658152 |
| **Bcas1** | -0.0954907524999982 | 25.73052803125 | -0.647927978846423 | 0.538269177563316 | 0.883205542417191 | -5.08722134890558 |
| **Pnp** | 0.125621655 | 20.648737385 | 0.643614727740485 | 0.540899698130895 | 0.886449011530399 | -5.08939553710737 |
| **Aldh1l1** | 0.200094597500005 | 23.10839330875 | 0.643200398953358 | 0.541152801260252 | 0.886449011530399 | -5.08960373647246 |
| **Macf1** | -0.115898282500012 | 25.44502620375 | -0.642798079917264 | 0.541398637731397 | 0.886449011530399 | -5.08980579144706 |
| **Ppp3r1** | -0.211808245 | 24.041870895 | -0.641852237814592 | 0.541976863984288 | 0.886718270333417 | -5.09028039247114 |

| | | | | | | |
|---|---|---|---|---|---|---|
| **Rpl17** | -0.0497896650000023 | 30.8006290175 | -0.641271941563053 | 0.542331807299719 | 0.886718270333417 | -5.0905712758349 |
| **Prdm11** | 0.118248599999994 | 23.389437165 | 0.640533824073784 | 0.542783489824436 | 0.886744039870382 | -5.09094094493037 |
| **Septin5** | -0.0899822299999968 | 20.803731245 | -0.639481730499989 | 0.543427705782003 | 0.886744039870382 | -5.09146723260648 |
| **S100a6** | -0.0998424149999977 | 25.50993814 | -0.638241203658092 | 0.544187905477275 | 0.886744039870382 | -5.09208682942798 |
| **Rhob** | 0.315896142500002 | 20.67407702375 | 0.638067768473199 | 0.544294239168924 | 0.886744039870382 | -5.09217337180653 |
| **Ap2b1** | -4.477682585 | 12.1575454975 | -0.63773103807218 | 0.544500726068415 | 0.886744039870382 | -5.09234133934362 |
| **Tubb4b** | -0.477868627499998 | 23.39039821625 | -0.637481470415199 | 0.544653794753503 | 0.886744039870382 | -5.09246577947658 |
| **Gnb1** | -0.102006929999998 | 23.4969075175 | -0.636699705577349 | 0.54513344941086 | 0.886899060501308 | -5.09285531536708 |
| **Sgip1** | 0.226019205 | 21.12566228 | 0.634036343129684 | 0.546769504806458 | 0.888813249674788 | -5.09417933469714 |
| **Dbn1** | -0.227884547499997 | 28.65599754875 | -0.631790308055425 | 0.548151531392273 | 0.888813249674788 | -5.09529219111666 |
| **Naxd** | 0.171942460000004 | 24.826041535 | 0.631312965045986 | 0.548445523390922 | 0.888813249674788 | -5.0955282663589 |
| **Retreg3** | -0.134381530000002 | 26.2861393625 | -0.631253868555542 | 0.548481927158026 | 0.888813249674788 | -5.09555748252125 |
| **Serbp1** | 0.105015170000001 | 29.8045064125 | 0.631086254926705 | 0.548585186091596 | 0.888813249674788 | -5.0956403346815 |
| **Capn2** | -0.220451197499997 | 22.65368962875 | -0.63102712557316 | 0.548621615750714 | 0.888813249674788 | -5.09566955807481 |
| **Csnk2a2** | -0.108108434999998 | 28.54640645 | -0.627180627695812 | 0.550994608709 | 0.890560676422597 | -5.09756554901922 |
| **Ntm** | -0.255698864999992 | 25.9628972725 | -0.626326321410146 | 0.551522494019761 | 0.890560676422597 | -5.09798529486666 |
| **Ssr4** | -0.0928034900000005 | 27.5167974 | -0.626143776119337 | 0.551635330511872 | 0.890560676422597 | -5.09807492089002 |
| **Rps13** | 0.107697229999999 | 22.5564018075 | 0.625768094240361 | 0.551867594294218 | 0.890560676422597 | -5.09825930227116 |
| **Synpo** | 0.261029430000004 | 23.1891699625 | 0.625752028947853 | 0.551877527916111 | 0.890560676422597 | -5.09826718485195 |
| **Gpm6a** | -0.0886689000000018 | 23.9084626875 | -0.625253478718503 | 0.552185849359922 | 0.890560676422597 | -5.09851171640499 |

| | | | | | | |
|---|---|---|---|---|---|---|
| **Rap2a** | -0.346598422500001 | 22.26038440375 | -0.624331052051049 | 0.552756586167687 | 0.890560676422597 | -5.09896371044494 |
| **Ranbp1** | 0.475621369999999 | 22.1686903 | 0.624279616490533 | 0.552788421602583 | 0.890560676422597 | -5.098988897224 |
| **Slc12a6** | -0.126482782499998 | 26.05243692625 | -0.623198665663154 | 0.553457719875464 | 0.891016719994903 | -5.09951779957347 |
| **Ehd1** | 0.0796260675000084 | 25.97923486375 | 0.620119444448772 | 0.555366979501174 | 0.893466960745612 | -5.10102010866556 |
| **Ppa2** | -0.0623649325000102 | 27.58882396375 | -0.618463091933059 | 0.556395634057876 | 0.894498068133464 | -5.10182556241182 |
| **Slc6a17** | 0.214499982500001 | 24.23469878875 | 0.616950349421063 | 0.557336101984963 | 0.894843886839778 | -5.1025595546007 |
| **Nckipsd** | 0.0836050549999925 | 26.9074968925 | 0.616869320635862 | 0.557386504286416 | 0.894843886839778 | -5.10259882642502 |
| **Slc7a5** | 0.1317813625 | 26.04369347875 | 0.615661370571787 | 0.558138209351072 | 0.894979423271633 | -5.10318374860156 |
| **Scamp4** | -0.133560262499998 | 24.47672316875 | -0.614855037296743 | 0.558640327237263 | 0.894979423271633 | -5.1035736448838 |
| **Rpl4** | -0.130909259999999 | 27.76455354 | -0.614593562957969 | 0.558803210001507 | 0.894979423271633 | -5.10369998383026 |
| **Snph** | 0.17183541 | 25.0358141075 | 0.614082429763136 | 0.55912169734265 | 0.894979423271633 | -5.103946818356 |
| **Acadm** | 0.370792235000007 | 20.6856680125 | 0.613393523215829 | 0.559551127249849 | 0.894979423271633 | -5.10427922112441 |
| **Gng4** | -0.0867862600000109 | 27.100207425 | -0.612399930315756 | 0.56017083080975 | 0.894979423271633 | -5.10475806811578 |
| **Erlin2** | 0.330467280000001 | 22.0374215625 | 0.611591064897303 | 0.560675622685527 | 0.894979423271633 | -5.10514739170686 |
| **Nebl** | 0.118349052500005 | 26.32624733875 | 0.611175057793916 | 0.560935347663344 | 0.894979423271633 | -5.10534745077467 |
| **Slc1a1** | -0.0832294675000043 | 26.23987307125 | -0.611131741557583 | 0.560962395340608 | 0.894979423271633 | -5.10536827489789 |
| **Rpl24** | 4.72474318 | 13.373633525 | 0.60945691350981 | 0.562008794105353 | 0.896029224603351 | -5.10617246056589 |
| **Pigs** | 0.148160037499995 | 25.28716802375 | 0.606040100685993 | 0.564147152431438 | 0.898817320897325 | -5.10780713493686 |
| **Prkce** | 0.121803170000007 | 20.9381392025 | 0.60516138135675 | 0.564697865417603 | 0.899073827134859 | -5.10822624136675 |
| **Khdrbs1** | 0.0525583975000039 | 26.61938835875 | 0.603888409001661 | 0.56549623098455 | 0.899569453579963 | -5.10883244917495 |
| **Cbln1** | -0.436589720000001 | 21.9313211425 | -0.602804177755811 | 0.566176751573783 | 0.899569453579963 | -5.10934789971055 |

| | | | | | | |
|---|---|---|---|---|---|---|
| **Atp9a** | 0.128168040000006 | 20.2399797325 | 0.602388979507016 | 0.56643748014761 | 0.899569453579963 | -5.10954507410928 |
| **Arpc3** | 0.375424077500004 | 21.39859048375 | 0.601635185550488 | 0.56691101532474 | 0.899569453579963 | -5.10990274257106 |
| **Fxyd7** | 0.250664077499998 | 22.07150115375 | 0.601559117113682 | 0.566958814696691 | 0.899569453579963 | -5.10993881469226 |
| **Gfap** | -0.0467258800000074 | 25.996665915 | -0.600595064327723 | 0.567564806039214 | 0.899670050196364 | -5.11039563012065 |
| **Ctnnb1** | 0.153842449999999 | 24.775375445 | 0.600217624114127 | 0.567802164319855 | 0.899670050196364 | -5.11057430584849 |
| **Coro1c** | -0.235613627500001 | 21.07790837125 | -0.598286466420426 | 0.569017514661142 | 0.900171067569701 | -5.11148696041967 |
| **Clasp2** | 0.113286274999997 | 25.3452804825 | 0.598103183158077 | 0.569132941316626 | 0.900171067569701 | -5.11157344578733 |
| **Rph3a** | 0.292631517499998 | 19.19529744125 | 0.597855509245819 | 0.569288941302208 | 0.900171067569701 | -5.1116902782601 |
| **Gpr158** | -0.214567522499994 | 24.06422857375 | -0.596958622779433 | 0.569854065154552 | 0.900447485144897 | -5.11211300319491 |
| **Megf9** | -4.2425882975 | 12.29611990625 | -0.595140198741808 | 0.571000857504726 | 0.900819772697047 | -5.11296837082728 |
| **Pip4k2b** | 0.154712097499999 | 25.52137666125 | 0.594634620667252 | 0.571319941836529 | 0.900819772697047 | -5.11320578441029 |
| **Ap2a2** | 0.0832139574999964 | 26.15104670875 | 0.594369788352402 | 0.571487126600628 | 0.900819772697047 | -5.11333007620766 |
| **Copb1** | 0.314511392500002 | 21.56610190625 | 0.594109359881916 | 0.571651559266787 | 0.900819772697047 | -5.11345225398544 |
| **Stt3b** | -0.115772447499992 | 27.70041155625 | -0.592375989858108 | 0.572746703202681 | 0.901929450026338 | -5.1142642566479 |
| **Dnajc8** | -0.115939972500005 | 29.70539084125 | -0.588992208964372 | 0.57488811187123 | 0.904684088735967 | -5.11584341515377 |
| **Uqcc1** | 0.288239702499997 | 21.79344741125 | 0.588219897962744 | 0.57537751904635 | 0.904837039154689 | -5.11620273001956 |
| **Purb** | 0.103631922500004 | 22.72001273875 | 0.587332533327205 | 0.575940134418182 | 0.905104829770263 | -5.11661506292794 |
| **Add2** | 0.0932475450000041 | 20.00931972 | 0.585007638132211 | 0.577415704288271 | 0.906615865585923 | -5.11769278538605 |
| **Snap47** | 0.0760866899999826 | 26.96642783 | 0.584579598530049 | 0.577687612662032 | 0.906615865585923 | -5.11789079738153 |
| **Tubb5** | 0.134228359999994 | 20.63078757 | 0.583808983105608 | 0.578177326610072 | 0.906767567973784 | -5.11824696483644 |

| | | | | | | |
|---|---|---|---|---|---|---|
| **Sec23a** | 0.1221386 | 28.545051855 | 0.583007635088123 | 0.578686825925183 | 0.90695007296834 | -5.11861689875296 |
| **Tusc2** | -0.388964979999997 | 21.541925375 | -0.580894769809101 | 0.580031438036455 | 0.908440276680314 | -5.11959013963796 |
| **Nit2** | 0.125205494999996 | 25.8222205125 | 0.58011596627839 | 0.580527518315336 | 0.908600396711994 | -5.11994809307501 |
| **Golph3** | 0.2812756975 | 21.45494865125 | 0.57906441058235 | 0.581197724283508 | 0.90882937218721 | -5.12043073832142 |
| **Ddx5** | 0.178891974999999 | 27.1613721925 | 0.578447861594372 | 0.581590887539886 | 0.90882937218721 | -5.12071336482552 |
| **Pkm** | -0.363716824999997 | 18.961580415 | -0.578032804248615 | 0.581855649206983 | 0.90882937218721 | -5.12090347811578 |
| **Get3** | 0.128961477500013 | 25.50856524375 | 0.574529083365739 | 0.584093409065795 | 0.911707371254932 | -5.12250353369437 |
| **Rps28** | -0.0555240775000101 | 26.64500184375 | -0.572038780282873 | 0.585686918305068 | 0.912841701006479 | -5.12363557121738 |
| **Rrbp1** | -0.0928956500000027 | 26.6266024475 | -0.571460161502824 | 0.586057524313014 | 0.912841701006479 | -5.1238979771653 |
| **Synpr** | 0.0632177150000004 | 21.252230285 | 0.571259040375439 | 0.586186374050977 | 0.912841701006479 | -5.12398913147709 |
| **Usp14** | -0.245565032500004 | 23.82361033125 | -0.570336986137741 | 0.586777302296298 | 0.912841701006479 | -5.12440667207816 |
| **Slc27a4** | 0.0439992099999849 | 28.4085724875 | 0.570303850482566 | 0.58679854468687 | 0.912841701006479 | -5.12442166604744 |
| **Twf2** | -0.3870877325 | 20.91755115125 | -0.568056866411242 | 0.588240051118828 | 0.914467518821521 | -5.12543663521335 |
| **Ccdc47** | 0.100313367499993 | 28.90396989125 | 0.565100351815684 | 0.590139813275833 | 0.916455932906665 | -5.12676669961251 |
| **Ckmt1** | 0.214227672499998 | 24.56814020125 | 0.564830137991859 | 0.590313617815043 | 0.916455932906665 | -5.12688795576952 |
| **Abi1** | -0.100398959999996 | 28.15843472 | -0.563776710883205 | 0.590991471308254 | 0.916891273912672 | -5.12736018161623 |
| **Dstn** | 0.197094494999995 | 20.1506811425 | 0.562220066121504 | 0.591993940212995 | 0.917604282196193 | -5.12805655728298 |
| **Snx12** | -3.6734518125 | 12.0199840485714 | -0.427271723454951 | 0.684574117483231 | 0.956590964044264 | -5.12822705865094 |
| **Dagla** | 0.104018767500015 | 27.90684670125 | 0.561828056293811 | 0.592246543645484 | 0.917604282196193 | -5.12823165643855 |
| **Rps19** | -0.129183654999999 | 25.409280955 | -0.560410265768892 | 0.593160649005514 | 0.918271001014205 | -5.12886403676908 |

| | | | | | | |
|---|---|---|---|---|---|---|
| **Mbp** | 0.063688302500001 | 28.55730957625 | 0.559926187766056 | 0.593472935635968 | 0.918271001014205 | -5.12907962644532 |
| **Pabpc1** | 0.101285067500008 | 22.90396801875 | 0.559203696963161 | 0.593939198897818 | 0.918271612449547 | -5.12940108833707 |
| **Hacd3** | 0.3277148775 | 21.95376772625 | 0.55869222489138 | 0.594269405022616 | 0.918271612449547 | -5.12962843765744 |
| **Ivd** | 0.2169844125 | 21.34930472625 | 0.55752961950683 | 0.595020367407235 | 0.918313138827896 | -5.13014452823286 |
| **Hbb-b1** | 0.0933181000000012 | 26.8329814525 | 0.556593653625874 | 0.595625324501972 | 0.918313138827896 | -5.13055931738668 |
| **Cldn11** | -0.148913297499988 | 23.56624576625 | -0.556350937784158 | 0.595782259213055 | 0.918313138827896 | -5.13066677986986 |
| **Mtdh** | 0.183595989999997 | 22.480322655 | 0.555533261760494 | 0.596311121564059 | 0.918313138827896 | -5.13102849936149 |
| **Gpr37l1** | 0.193479762499994 | 23.68136768375 | 0.555454474917301 | 0.596362093811692 | 0.918313138827896 | -5.13106332770629 |
| **Acsl1** | 0.147022045 | 21.346746215 | 0.552662483322532 | 0.598169990447452 | 0.918313138827896 | -5.13229471138111 |
| **Fdx2** | 3.35029986499999 | 20.717285655 | 0.552353119258005 | 0.598370501543145 | 0.918313138827896 | -5.13243081374777 |
| **Gmfb** | 0.167485259999999 | 24.49455336 | 0.552024921079175 | 0.598583260893499 | 0.918313138827896 | -5.13257512784209 |
| **Sh3glb2** | -0.103059634999994 | 29.6949188925 | -0.551905860197759 | 0.598660454342286 | 0.918313138827896 | -5.13262746196883 |
| **Rgs7** | -0.0465459225000089 | 27.62967177625 | -0.551763139167411 | 0.598752995252091 | 0.918313138827896 | -5.13269018284413 |
| **P4hb** | 0.607287794999991 | 25.5023475 | 0.550942363830828 | 0.599285344667762 | 0.918313138827896 | -5.13305060418907 |
| **Tubb3** | 0.268601132500002 | 22.52433423125 | 0.550793148246314 | 0.599382153309412 | 0.918313138827896 | -5.13311607682616 |
| **Bsn** | -0.101982479999997 | 24.303710585 | -0.550656223260097 | 0.599470995697794 | 0.918313138827896 | -5.13317614270272 |
| **Bcs1l** | -0.174813947499999 | 19.94254190625 | -0.549774044285537 | 0.600043564887809 | 0.918378561204771 | -5.13356281537197 |
| **Mlycd** | -0.0507514300000018 | 28.7143259675 | -0.549363897879217 | 0.600309870089638 | 0.918378561204771 | -5.13374240064233 |
| **Lin7a** | 0.0577831325000133 | 24.92762419625 | 0.546880051762792 | 0.601924021157153 | 0.919504767373831 | -5.13482741405317 |
| **Gnao1** | -0.129168860000007 | 21.546471665 | -0.54676166864393 | 0.602001013793439 | 0.919504767373831 | -5.13487901755296 |

| | | | | | | |
|---|---|---|---|---|---|---|
| **Coasy** | -0.127806919999998 | 33.73048903 | -0.545586335288164 | 0.602765710363002 | 0.919504767373831 | -5.1353908071482 |
| **Gria2** | 0.0893480374999953 | 24.00796819625 | 0.54534373082388 | 0.602923621061399 | 0.919504767373831 | -5.1354963248689 |
| **Pdk3** | 0.0746948074999914 | 28.56529590625 | 0.545166664978237 | 0.603038887315391 | 0.919504767373831 | -5.13557331093806 |
| **Mpdu1** | 0.120920452500002 | 24.56222564375 | 0.54441495608294 | 0.603528370599503 | 0.919643296547591 | -5.13589989608895 |
| **Cd200** | 0.0608680499999927 | 27.1119990825 | 0.542984096952833 | 0.604460698198223 | 0.920455993889967 | -5.13652042976619 |
| **Prodh** | -0.392203835000004 | 20.9282541325 | -0.540882459249711 | 0.605831539658303 | 0.921581632505938 | -5.13742921784685 |
| **Dnm3** | 0.2297699075 | 20.96273290875 | 0.539274084643551 | 0.606881797389515 | 0.921581632505938 | -5.13812257882138 |
| **Fmnl1** | 0.2619074425 | 23.44271614875 | 0.538259936621135 | 0.60754454462827 | 0.921581632505938 | -5.13855882152429 |
| **Rab35** | -0.0738696049999916 | 28.17282714 | -0.537227603831307 | 0.608219584524361 | 0.921581632505938 | -5.13900213083977 |
| **Gsk3a** | 0.364203267500002 | 22.68564242875 | 0.536369345645877 | 0.608781111164533 | 0.921581632505938 | -5.1393701074527 |
| **Pip4k2a** | 0.0812855600000049 | 25.6330683275 | 0.536068842926683 | 0.608977786263172 | 0.921581632505938 | -5.13949882275584 |
| **Adcyap1r1** | 0.0992865825000031 | 25.52439361625 | 0.534790412906297 | 0.609814891564195 | 0.921581632505938 | -5.1400456936914 |
| **Nsf** | 0.232999005000007 | 23.3715865575 | 0.534530642906665 | 0.609985063855032 | 0.921581632505938 | -5.1401566716795 |
| **Fabp5** | 0.210278589999994 | 22.9079681075 | 0.534431045708574 | 0.610050315714256 | 0.921581632505938 | -5.1401992083985 |
| **Rpl13** | 0.362850439999999 | 23.3452555975 | 0.53338708600355 | 0.610734503810704 | 0.921581632505938 | -5.14064464244193 |
| **Pals2** | -0.114147290000002 | 22.6754933125 | -0.532958451793106 | 0.611015542715135 | 0.921581632505938 | -5.14082730456476 |
| **Septin2** | 0.154160479999998 | 23.8630295525 | 0.532942767652171 | 0.611025827541792 | 0.921581632505938 | -5.14083398584824 |
| **Pgk1** | -3.26601712416667 | 11.7562291771429 | -0.387357962381178 | 0.712293536659626 | 0.957813491602516 | -5.1411397629906 |
| **Scrn1** | 0.0532312225000027 | 29.80873514375 | 0.531820431641684 | 0.611762041512273 | 0.921581632505938 | -5.14131163105619 |
| **Slc2a13** | 0.15913811250001 | 29.66572941875 | 0.531028885914338 | 0.612281559505073 | 0.921581632505938 | -5.14164795414202 |

| | | | | | | |
|---|---|---|---|---|---|---|
| **Golga7** | 0.224568662499998 | 19.07284547375 | 0.53095176062805 | 0.612332192280206 | 0.921581632505938 | -5.14168070015267 |
| **Cacnb4** | 0.0609043149999984 | 28.189597965 | 0.530620856045254 | 0.612549457158964 | 0.921581632505938 | -5.1418211477798 |
| **Atl1** | -0.116124527499995 | 23.85590901125 | -0.530420362057674 | 0.612681117671254 | 0.921581632505938 | -5.14190620624511 |
| **Arhgef40** | -0.0998053125000062 | 21.39572144375 | -0.530418143682706 | 0.612682574521333 | 0.921581632505938 | -5.14190714721681 |
| **Gpx4** | -0.104812945000006 | 30.99120316 | -0.529234963777736 | 0.613459860754885 | 0.921581632505938 | -5.14240851436111 |
| **Armc1** | -0.143190314999995 | 25.600644215 | -0.529039515018464 | 0.613588311827022 | 0.921581632505938 | -5.14249123800062 |
| **Wdr7** | -0.0850476999999997 | 27.92175202 | -0.529038755165457 | 0.613588811239324 | 0.921581632505938 | -5.14249155955459 |
| **Cat** | 0.203076375000002 | 24.8332733825 | 0.527147955210981 | 0.614832221308759 | 0.922408263298891 | -5.14329041977298 |
| **Nomo1** | 0.175238647500002 | 22.97647249875 | 0.526985918161516 | 0.614938842199261 | 0.922408263298891 | -5.14335876044774 |
| **Acox1** | 0.109481677499996 | 24.85179527625 | 0.524146597719645 | 0.616808750576652 | 0.924254561224128 | -5.14455320032679 |
| **Cyb5b** | -0.246200299999998 | 24.96109749 | -0.523900501365684 | 0.616970968480779 | 0.924254561224128 | -5.14465645389095 |
| **Ckap5** | -0.0543416700000066 | 27.6001097925 | -0.522138587209345 | 0.618133032341863 | 0.924637073857137 | -5.14539441558836 |
| **Crk** | -0.069865489999998 | 26.9082695 | -0.521836065254158 | 0.618332678278618 | 0.924637073857137 | -5.14552089865081 |
| **Gnb2** | 0.200689395000001 | 23.54053623 | 0.521122300186814 | 0.618803857184885 | 0.924637073857137 | -5.14581905853832 |
| **Marcks** | -0.129906950000002 | 23.2205440875 | -0.52074475798446 | 0.619053162928897 | 0.924637073857137 | -5.14597661972149 |
| **Cldnd1** | 0.1290164125 | 27.02319011375 | 0.520159978921907 | 0.619439421924328 | 0.924637073857137 | -5.14622046445873 |
| **Ppp2r2a** | 0.552005985000005 | 24.1526059875 | 0.519869880296569 | 0.619631086338593 | 0.924637073857137 | -5.1463413397828 |
| **Mlc1** | 0.0489607050000096 | 24.79523557 | 0.518469941020767 | 0.620556456196658 | 0.924693013692072 | -5.14692379628656 |
| **C3orf18** | 0.102171265000003 | 25.5236536425 | 0.51831067494425 | 0.620661779203948 | 0.924693013692072 | -5.14698997041392 |
| **Ubl4a** | 0.375322315000002 | 19.6728507 | 0.517630119497106 | 0.621111940081452 | 0.924693013692072 | -5.14727253032368 |

| | | | | | | |
|---|---|---|---|---|---|---|
| **Ngef** | -0.0820523849999972 | 26.939431815 | -0.517388421877798 | 0.621271855753234 | 0.924693013692072 | -5.14737280007602 |
| **Phf24** | -0.270862302499992 | 21.5223739712 5 | -0.515075599055395 | 0.622803216017404 | 0.925527660346887 | -5.14833014820753 |
| **Coro1a** | -0.179235385000009 | 23.215007 | -0.514682539155239 | 0.623063668317356 | 0.925527660346887 | -5.14849246213689 |
| **Ccz1** | 0.0544773975000012 | 27.1668796362 5 | 0.513327535959162 | 0.623961976473535 | 0.925527660346887 | -5.14905115076257 |
| **Micos13** | -0.0973133174999958 | 27.9736434462 5 | -0.512805770640163 | 0.624308068293686 | 0.925527660346887 | -5.14926592716688 |
| **Rpl18** | 0.132241472499999 | 23.5745593212 5 | 0.512315005680048 | 0.624633690625793 | 0.925527660346887 | -5.14946776230962 |
| **Src** | -0.0689149824999973 | 27.0768527137 5 | -0.510284736405252 | 0.625981733607913 | 0.925527660346887 | -5.15030088401189 |
| **Dpysl4** | 0.578620164999993 | 23.39737341 | 0.50998271747946 | 0.626182397863405 | 0.925527660346887 | -5.15042456148946 |
| **Eef1d** | 0.104848812499995 | 20.0842386162 5 | 0.509410815760112 | 0.626562468361806 | 0.925527660346887 | -5.15065857481075 |
| **Atp6v1c1** | 0.314126162500003 | 23.3889921087 5 | 0.508448064375289 | 0.62720256334337 | 0.925527660346887 | -5.15105198010471 |
| **Ppp3cb** | 0.289197337499999 | 22.5807573687 5 | 0.508405036447532 | 0.627231178983628 | 0.925527660346887 | -5.15106954668381 |
| **Mgll** | -0.194711155 | 20.70086201 | -0.507191478395648 | 0.628038538151277 | 0.925527660346887 | -5.15156443849317 |
| **Slc20a2** | -0.0769824949999958 | 22.5958585925 | -0.506729372630273 | 0.628346113593738 | 0.925527660346887 | -5.15175260422838 |
| **Nucb1** | 0.0622333325 | 26.1559020487 5 | 0.506623257376868 | 0.628416754653256 | 0.925527660346887 | -5.15179579152991 |
| **Itga6** | 0.182416732499995 | 21.1751899662 5 | 0.506156601812892 | 0.628727457668193 | 0.925527660346887 | -5.1519856158591 |
| **Agap3** | -0.0697979225000012 | 28.9202902887 5 | -0.505915632914982 | 0.62888792845777 | 0.925527660346887 | -5.15208357408821 |
| **Myl6** | -0.221105692499997 | 24.1056440162 5 | -0.50571200375654 | 0.629023550086367 | 0.925527660346887 | -5.15216632002754 |
| **Palm** | -0.204601227499992 | 22.9720856337 5 | -0.504440745078109 | 0.62987058604405 | 0.925527660346887 | -5.15268221984375 |
| **Esd** | 0.0960844699999939 | 23.7326152275 | 0.504129913527287 | 0.630077783642128 | 0.925527660346887 | -5.15280818150598 |
| **Plscr3** | 0.0764564275000019 | 29.2628739762 5 | 0.503204338407566 | 0.630694976500016 | 0.925527660346887 | -5.15318284469609 |

| | | | | | | |
|---|---|---|---|---|---|---|
| **Acsl6** | -0.101166150000001 | 27.002755385 | -0.503100417062986 | 0.630764293287965 | 0.925527660346887 | -5.15322487191475 |
| **Rdh14** | 0.112912527500008 | 22.36130689375 | 0.502571740982926 | 0.631116988641203 | 0.925527660346887 | -5.15343855363808 |
| **Synj2bp** | 0.124643204999998 | 25.2311556725 | 0.502453822519693 | 0.63119566965421 | 0.925527660346887 | -5.15348618639044 |
| **Rab3a** | 0.1858286175 | 20.51906182375 | 0.501672773573059 | 0.631716954157592 | 0.925527660346887 | -5.15380143185324 |
| **Chst3** | 0.158234077500008 | 25.44603376625 | 0.501333843820375 | 0.631943231587586 | 0.925527660346887 | -5.15393809129989 |
| **Sv2a** | -0.10679447750001 | 30.45429171625 | -0.501175656685707 | 0.632048855577312 | 0.925527660346887 | -5.15400184499372 |
| **Efr3b** | 0.519310767499999 | 25.24322466625 | 0.500744557725453 | 0.632336754016278 | 0.925527660346887 | -5.15417549665638 |
| **Tmpo** | -0.134335262500002 | 21.05784347375 | -0.500253839216556 | 0.632664551524508 | 0.925527660346887 | -5.15437299834881 |
| **Aldh4a1** | -0.0689303250000002 | 28.9861979775 | -0.497188802012937 | 0.634713993527006 | 0.927937378369329 | -5.15560260934248 |
| **Cadm4** | -0.386980755000003 | 21.93974771 | -0.494799859432193 | 0.636313763468763 | 0.928070410018091 | -5.15655621581595 |
| **Ubqln2** | 0.0517442900000056 | 26.4832849925 | 0.494475196686014 | 0.636531337701043 | 0.928070410018091 | -5.15668548999379 |
| **Slc2a3** | -0.079160009999999 | 23.5648654625 | -0.494183658604165 | 0.636726746245627 | 0.928070410018091 | -5.15680150867253 |
| **H1-0** | -0.0937644649999996 | 24.3003398775 | -0.494067972582383 | 0.636804295483578 | 0.928070410018091 | -5.15684752907091 |
| **Cacng8** | -0.11716162750001 | 24.56311821875 | -0.494049900341135 | 0.636816410515231 | 0.928070410018091 | -5.15685471739916 |
| **Atad1** | -0.113540177500003 | 24.06387145625 | -0.491497976389907 | 0.638528336936314 | 0.929977824060655 | -5.15786735003003 |
| **Rap2c** | -0.491001157500008 | 23.56163630375 | -0.488476443178395 | 0.640558371446773 | 0.932345844118426 | -5.15906013833031 |
| **Emc3** | -0.0600922775000008 | 30.26996676875 | -0.48781775089462 | 0.641001359273582 | 0.932402355513338 | -5.15931927324937 |
| **Cadm1** | 0.147476645000001 | 26.42209859 | 0.486791965423253 | 0.641691541000553 | 0.932818138051844 | -5.15972218914675 |
| **Gdap1l1** | -0.114592304999988 | 25.8550981975 | -0.485160043156944 | 0.642790338454219 | 0.9338270219231 | -5.16036158954039 |
| **Rab27b** | 0.172730010000006 | 23.01616057 | 0.483730659985577 | 0.643753556083367 | 0.934637793508073 | -5.16092001946631 |

| | | | | | | |
|---|---|---|---|---|---|---|
| **Rps16** | 0.100470200000004 | 27.37986727 | 0.482400315663461 | 0.644650698245609 | 0.935351673492214 | -5.16143840172125 |
| **Wdr37** | 0.0788689824999942 | 22.18207775375 | 0.481148254136908 | 0.645495632459606 | 0.935988952912829 | -5.16192508584407 |
| **Slc25a18** | -0.179666510000011 | 23.634847175 | -0.480169356196318 | 0.646156619935819 | 0.936358870723577 | -5.1623047822976 |
| **Vcam1** | -0.0501108700000046 | 25.782584915 | -0.478185510420732 | 0.64749724155049 | 0.937140625583078 | -5.16307210513449 |
| **Sec62** | 0.0706006374999859 | 26.65773572625 | 0.478168828502051 | 0.647508520667285 | 0.937140625583078 | -5.16307854510686 |
| **Tomm34** | -0.0475154775000064 | 21.03299464875 | -0.47412133519331 | 0.650248090404099 | 0.939462689260693 | -5.16463496468536 |
| **Cycs** | -0.212581740000008 | 24.1541441575 | -0.473886702513393 | 0.650407082430869 | 0.939462689260693 | -5.16472481755777 |
| **Pdzd7** | 0.126626215000009 | 22.1737804075 | 0.473090988586879 | 0.650946421007634 | 0.939462689260693 | -5.16502923288227 |
| **Pspc1** | 0.0550588699999999 | 30.2868877475 | 0.472932516949545 | 0.65105386075094 | 0.939462689260693 | -5.16508980305151 |
| **Hsp90b1** | -0.0864276924999992 | 25.26392638375 | -0.472792137232023 | 0.651149042101365 | 0.939462689260693 | -5.16514344264788 |
| **Lin7c** | -0.0470206250000018 | 26.6912123775 | -0.470826868635337 | 0.652482284762829 | 0.940628346012298 | -5.16589284170534 |
| **Nrxn1** | 0.0567388125000043 | 28.13027461375 | 0.46995537422198 | 0.653073947564361 | 0.940628346012298 | -5.16622424239615 |
| **Ina** | -0.168238210000002 | 24.6236825925 | -0.46911607220386 | 0.653644009281062 | 0.940628346012298 | -5.16654286737779 |
| **Bckdhb** | -0.114755385000002 | 24.61079124 | -0.46906049056838 | 0.653681769652702 | 0.940628346012298 | -5.16656394938052 |
| **Anxa2** | 0.179805640000005 | 22.044249835 | 0.468598596767408 | 0.653995607717263 | 0.940628346012298 | -5.16673905583214 |
| **Tst** | -0.0758210825000027 | 21.85637860875 | -0.467745070678462 | 0.654575742584386 | 0.940804238115597 | -5.16706221433569 |
| **Cdk5** | -0.04356297 | 30.1335555225 | -0.465651333512951 | 0.655999928608097 | 0.940804238115597 | -5.16785263830161 |
| **Slc25a46** | 0.173633555000002 | 23.95216488 | 0.464581123644414 | 0.656728495348582 | 0.940804238115597 | -5.1682554001215 |
| **Mapre1** | -0.0522386275000031 | 26.81276141875 | -0.464125215932895 | 0.657038986014709 | 0.940804238115597 | -5.16842671655693 |
| **Nefl** | -0.201796202500002 | 20.50259119375 | -0.463977039865193 | 0.657139915341167 | 0.940804238115597 | -5.16848236328645 |

| | | | | | | |
|---|---|---|---|---|---|---|
| **Atp6ap1** | -0.0664371749999937 | 24.784711725 | -0.463721856731422 | 0.657313750034107 | 0.940804238115597 | -5.16857815754584 |
| **Ppp1r9b** | -0.0824472375000002 | 25.74324029375 | -0.463522735567376 | 0.657449410334517 | 0.940804238115597 | -5.16865287271591 |
| **Hnrnpa2b1** | 0.0523681475000082 | 26.19505472625 | 0.463055233188844 | 0.657767972185559 | 0.940804238115597 | -5.16882817483387 |
| **Cadm3** | -0.187582584999987 | 21.4593750225 | -0.463025641739487 | 0.657788138743155 | 0.940804238115597 | -5.16883926542374 |
| **Gdap1** | 0.0877852025000045 | 22.86585909625 | 0.462180194855565 | 0.658364440095417 | 0.940881887463225 | -5.16915585458733 |
| **Vti1a** | -0.104994382500003 | 28.80896752375 | -0.461749523201225 | 0.658658105007849 | 0.940881887463225 | -5.1693169202716 |
| **Gnpda2** | 0.103137972500001 | 26.78128506625 | 0.460728996466166 | 0.659354237519832 | 0.941293456657335 | -5.16969803098683 |
| **Mrpl12** | -0.215995247500015 | 23.70846090125 | -0.459550478377606 | 0.660158593808468 | 0.941858921407628 | -5.17013717427162 |
| **Abhd16a** | -0.346452430000006 | 21.0677406325 | -0.457211396400384 | 0.661756487374638 | 0.943555139909327 | -5.17100569335432 |
| **Slc39a10** | 0.179066584999994 | 24.4973750375 | 0.456085507686534 | 0.662526292809322 | 0.944069275794382 | -5.17142228422767 |
| **Erp44** | 0.1538953375 | 24.65570795625 | 0.4527134679297 | 0.664834491794474 | 0.94621617371926 | -5.17266429114498 |
| **Stxbp5l** | -0.0752039599999996 | 22.89959337 | -0.452686106729236 | 0.664853236930612 | 0.94621617371926 | -5.17267433407923 |
| **Tceal5** | 0.108483232500014 | 24.12404270875 | 0.450276996602517 | 0.666504728492766 | 0.94798175624711 | -5.17355639489641 |
| **Coq6** | 0.0482198175000086 | 29.12899358125 | 0.44944277721603 | 0.667077067766102 | 0.948211210928155 | -5.17386081596605 |
| **Comtd1** | 0.0737806124999985 | 25.64210133625 | 0.447077408877385 | 0.66870119304488 | 0.949934514996637 | -5.17472113430029 |
| **Pdia6** | 0.108937127500006 | 30.34111142625 | 0.444116468720008 | 0.670736951744882 | 0.951960989316029 | -5.17579213504515 |
| **Rpn1** | 0.0699283649999956 | 28.15571829 | 0.443650904167554 | 0.671057317081531 | 0.951960989316029 | -5.17595993327411 |
| **Scp2** | -0.0841694824999983 | 19.69412855875 | -0.44302877035257 | 0.671485536359224 | 0.951960989316029 | -5.17618390690993 |
| **Mtnd4** | -0.110319560000001 | 25.85425228 | -0.442603571724311 | 0.671778279413305 | 0.951960989316029 | -5.17633681422699 |
| **Syn2** | -0.395481975000003 | 22.48300073 | -0.441038885450008 | 0.672856071694607 | 0.952555870517044 | -5.1768983225245 |

| | | | | | | |
|---|---|---|---|---|---|---|
| **Rab8b** | 0.386228864999996 | 23.1181838275 | 0.439918708483147 | 0.67362818536286 | 0.952555870517044 | -5.17729917764546 |
| **Hmgb1** | -0.113680844999998 | 26.7648543525 | -0.439432471658664 | 0.673963470005419 | 0.952555870517044 | -5.17747288243314 |
| **Adcy9** | 0.0523696375000036 | 26.8424689812 5 | 0.439001758694697 | 0.674260534910468 | 0.952555870517044 | -5.1776266025061 |
| **Sec22b** | 0.0919752524999957 | 22.2419029562 5 | 0.438998816156298 | 0.674262564609594 | 0.952555870517044 | -5.17762765220648 |
| **Omg** | -1.8861365475 | 14.6843206028 571 | -0.24325353318576 | 0.816175862020677 | 0.991389288010703 | -5.17772082254275 |
| **Crkl** | 0.26718383 | 24.4941849125 | 0.434716371621181 | 0.677219593682675 | 0.956147859624193 | -5.17914840213866 |
| **Por** | 0.0721911325000093 | 27.5751970187 5 | 0.43367615606141 | 0.677938792801482 | 0.956383769286733 | -5.17951570043659 |
| **Nap1l4** | 0.0904073574999877 | 29.6906555787 5 | 0.433275648232552 | 0.678215798245815 | 0.956383769286733 | -5.17965690047743 |
| **Eef1a2** | 0.0750759699999968 | 25.029804835 | 0.430688387382661 | 0.680006531076069 | 0.956590964044264 | -5.18056611571048 |
| **Scai** | -0.0702904750000073 | 31.72226373 | -0.430521589281301 | 0.68012205441746 | 0.956590964044264 | -5.18062455768875 |
| **Thsd7a** | 0.0905890149999955 | 25.3166121725 | 0.430073483572899 | 0.680432455543134 | 0.956590964044264 | -5.18078145842689 |
| **L2hgdh** | -0.211360647500005 | 24.6699389037 5 | -0.429900325364108 | 0.680552419475335 | 0.956590964044264 | -5.1808420476068 |
| **Inpp1** | 0.144697822500007 | 23.3880079837 5 | 0.426595182875281 | 0.682844129054825 | 0.956590964044264 | -5.18199417296708 |
| **Cse1l** | -0.0669252349999923 | 29.87336545 | -0.426210290979709 | 0.683111239445647 | 0.956590964044264 | -5.18212780112743 |
| **RTRAF** | 0.188386467499992 | 23.7840153387 5 | 0.425717538303969 | 0.683453275384356 | 0.956590964044264 | -5.18229871235479 |
| **Syt1** | -0.0650091150000023 | 29.040576035 | -0.425251723058463 | 0.683776686876745 | 0.956590964044264 | -5.18246011052784 |
| **Stx16** | 0.236725337499994 | 23.5012335862 5 | 0.425183764685731 | 0.68382387577158 | 0.956590964044264 | -5.18248364331322 |
| **Cap1** | 0.0885606125000074 | 25.0530077837 5 | 0.42493728689448 | 0.683995037656367 | 0.956590964044264 | -5.18256896474121 |
| **Hnrnpk** | 0.150280142499998 | 22.9218212987 5 | 0.424664452018433 | 0.684184526083315 | 0.956590964044264 | -5.18266335610629 |
| **Radil** | -0.282884390000007 | 22.0375998625 | -0.423479108942669 | 0.685008051523593 | 0.956590964044264 | -5.18307278577685 |

| | | | | | | |
|---|---|---|---|---|---|---|
| **Scamp3** | 0.0938214950000003 | 21.1681132275 | 0.423211334962252 | 0.685194153399084 | 0.956590964044264 | -5.18316512954561 |
| **Mapre3** | -0.0600830049999921 | 27.74689606 | -0.421559924729612 | 0.686342397585911 | 0.956590964044264 | -5.18373342313921 |
| **H4c1** | 0.131433472499992 | 26.74790930375 | 0.420961661555012 | 0.686758597431986 | 0.956590964044264 | -5.18393878842206 |
| **Tma7** | -0.0636586125000029 | 29.30635249125 | -0.420112713744673 | 0.687349394715835 | 0.956590964044264 | -5.18422973768636 |
| **Them6** | -0.055155120000002 | 27.97575794 | -0.419454453497906 | 0.68780765145693 | 0.956590964044264 | -5.18445495668219 |
| **Cntn2** | 0.127937992499998 | 20.56340418875 | 0.419433972738851 | 0.687821911683761 | 0.956590964044264 | -5.18446195872435 |
| **Atp2b2** | -0.133465817499992 | 26.21021562375 | -0.419311116869767 | 0.687907455946715 | 0.956590964044264 | -5.1845039544532 |
| **Tagln2** | -0.168305870000001 | 22.5638874625 | -0.418810246558352 | 0.68825626178854 | 0.956590964044264 | -5.18467504739567 |
| **Pmpcb** | -0.0496404900000087 | 27.38251688 | -0.418445716486656 | 0.688510171802385 | 0.956590964044264 | -5.1847994473351 |
| **Gas7** | -0.0380891200000022 | 21.9896543525 | -0.418042518392917 | 0.688791066110748 | 0.956590964044264 | -5.18493692503848 |
| **Rmnd1** | 0.179426290000002 | 25.0490057725 | 0.417681429753318 | 0.689042669147816 | 0.956590964044264 | -5.18505993942396 |
| **Tubb4a** | 0.145077375 | 25.55580167 | 0.416274926829738 | 0.690023110993424 | 0.956590964044264 | -5.1855381525669 |
| **Cdipt** | -0.0626924374999938 | 25.18605905625 | -0.416183865984051 | 0.690086609634202 | 0.956590964044264 | -5.1855690613105 |
| **Acly** | 0.0737815924999978 | 23.79412636125 | 0.416011036240879 | 0.690207134847033 | 0.956590964044264 | -5.18562770741222 |
| **Slc4a4** | -0.0652100649999952 | 27.45321943 | -0.41555366137933 | 0.690526138294763 | 0.956590964044264 | -5.18578279772748 |
| **Sptan1** | -0.124970647499993 | 24.67898967875 | -0.415471751466733 | 0.690583274842472 | 0.956590964044264 | -5.18581055551489 |
| **Adck1** | -0.2583243875 | 21.06606442625 | -0.412259115252393 | 0.692825972884334 | 0.956590964044264 | -5.18689521310507 |
| **Rdh11** | 0.0415096350000042 | 26.3784491375 | 0.412090886235234 | 0.692943503115244 | 0.956590964044264 | -5.18695179339737 |
| **Scamp5** | -0.120032977499999 | 24.89710305125 | -0.411406194215003 | 0.693421945144187 | 0.956590964044264 | -5.18718185162063 |
| **Cotl1** | 0.2537977275 | 22.45074848875 | 0.411071255622972 | 0.693656045036108 | 0.956590964044264 | -5.1872942610724 |

| | | | | | | |
|---|---|---|---|---|---|---|
| **Capn5** | 0.233942004999992 | 23.657082025 | 0.410510707258819 | 0.694047911913604 | 0.956590964044264 | -5.18748219567431 |
| **Actr1a** | -0.0575762650000016 | 29.74792936 | -0.410238141134049 | 0.694238493300833 | 0.956590964044264 | -5.18757349172679 |
| **Rap1gap2** | -0.0985755500000032 | 26.5978063825 | -0.409470736339526 | 0.694775199692095 | 0.956590964044264 | -5.18783022829409 |
| **Fus** | -0.182678689999999 | 19.9520624575 | -0.408917367311944 | 0.695162331256985 | 0.956590964044264 | -5.18801507876214 |
| **Bphl** | 0.0544421350000093 | 26.21107087 | 0.408643645719634 | 0.695353860469911 | 0.956590964044264 | -5.18810642747013 |
| **Ssb** | -0.0529158725000052 | 27.33630211875 | -0.408293664637255 | 0.695598785116503 | 0.956590964044264 | -5.18822314246413 |
| **Syt12** | 0.0850845824999986 | 24.10283725875 | 0.408060905110464 | 0.695761697175184 | 0.956590964044264 | -5.18830071330178 |
| **Gabrb3** | 0.194289952500007 | 24.25848204625 | 0.407290346166205 | 0.696301147084956 | 0.956590964044264 | -5.18855721791713 |
| **Krt2** | -0.113266352499995 | 27.90204275625 | -0.406685800483847 | 0.696724507812452 | 0.956590964044264 | -5.18875814094982 |
| **Slc5a3** | -0.1062521725 | 20.43045725125 | -0.406358681918178 | 0.696953636139723 | 0.956590964044264 | -5.18886674310152 |
| **Rap2b** | -0.105282340000002 | 23.7048388125 | -0.406097279561676 | 0.697136758443618 | 0.956590964044264 | -5.18895346867763 |
| **Rpl26** | 0.0916474725000036 | 24.15723506375 | 0.405646748984944 | 0.697452423354552 | 0.956590964044264 | -5.18910281829851 |
| **Ubxn6** | -0.123144945000004 | 20.9091900575 | -0.405073646884965 | 0.697854061546567 | 0.956590964044264 | -5.18929257482518 |
| **Fahd2** | -0.118072282500002 | 20.31604747625 | -0.404478754956584 | 0.698271080830862 | 0.956590964044264 | -5.1894892792493 |
| **Cd47** | 0.0553067600000077 | 27.93463466 | 0.404150352914603 | 0.698501338846019 | 0.956590964044264 | -5.1895977508096 |
| **Rab12** | 0.136702547500008 | 23.03423450125 | 0.403363018660784 | 0.69905351505433 | 0.956590964044264 | -5.18985747070905 |
| **Srcin1** | 0.0466035999999832 | 32.877774535 | 0.402612820059764 | 0.699579829939127 | 0.956590964044264 | -5.19010449732466 |
| **Podxl2** | -0.0551605374999973 | 27.29337477125 | -0.402446498252029 | 0.699696540028947 | 0.956590964044264 | -5.19015920544576 |
| **Pip5k1c** | 0.0613032275000052 | 27.19155959875 | 0.402121841204484 | 0.699924381147255 | 0.956590964044264 | -5.19026593338949 |
| **Psmc5** | 0.116662657499997 | 29.74646121625 | 0.400404298741056 | 0.701130289557678 | 0.957166117407006 | -5.19082921033352 |

| | | | | | | |
|---|---|---|---|---|---|---|
| **Hnrnpc** | 0.221436245000003 | 22.316429105 | 0.400340636069434 | 0.701175005816142 | 0.957166117407006 | -5.19085004521224 |
| **Snap25** | 0.0530207024999925 | 23.38931905875 | 0.398513465047051 | 0.702458942902256 | 0.957740463709318 | -5.19144669353557 |
| **Mfsd6** | -0.0475039275000029 | 26.43795255125 | -0.397804331245504 | 0.702957527587919 | 0.957740463709318 | -5.19167756303356 |
| **Arf4** | -0.0460758549999909 | 31.1726394725 | -0.397706379549866 | 0.703026408817071 | 0.957740463709318 | -5.19170942227267 |
| **Pin1** | 0.133817507500005 | 21.43210699875 | 0.397379458745414 | 0.703256326624874 | 0.957740463709318 | -5.19181570125136 |
| **Sfpq** | -0.09265173 | 21.3267517 | -0.396298468078905 | 0.704016807152083 | 0.957813491602516 | -5.19216653497675 |
| **Psmc6** | 0.0608627974999969 | 30.02677863875 | 0.39595894595067 | 0.704255737524183 | 0.957813491602516 | -5.1922765404109 |
| **Tuba4a** | 0.0509557799999989 | 26.7854887475 | 0.394850436082795 | 0.705036074776279 | 0.957813491602516 | -5.19263507950542 |
| **Faah** | -0.0815840850000029 | 27.96505313 | -0.394241225170318 | 0.705465092912433 | 0.957813491602516 | -5.19283172049905 |
| **Micu3** | -0.105627404999993 | 24.6159977875 | -0.3940955289461 | 0.705567712146486 | 0.957813491602516 | -5.19287870586439 |
| **Prrt3** | 0.150950147499994 | 24.04345250125 | 0.393141132910217 | 0.706240091857258 | 0.957813491602516 | -5.19318608246671 |
| **Epb41l3** | -0.116364415 | 25.018385245 | -0.39219985225558 | 0.706903508863966 | 0.957813491602516 | -5.19348854622579 |
| **Synj1** | -0.0527613474999988 | 30.50818065375 | -0.392056186100241 | 0.70700478931728 | 0.957813491602516 | -5.1935346505923 |
| **Lypla2** | 0.0790677224999996 | 20.25610653125 | 0.390346091981855 | 0.708210846712481 | 0.957813491602516 | -5.19408221769375 |
| **Rab3b** | -0.068284432500004 | 20.34938514625 | -0.389568847113556 | 0.708759303628911 | 0.957813491602516 | -5.19433034205658 |
| **Krt76** | -0.0905505900000065 | 28.070461365 | -0.389535613496875 | 0.70878275883356 | 0.957813491602516 | -5.19434094100222 |
| **Slc7a10** | -0.0509313724999956 | 24.73584053375 | -0.388851966848395 | 0.709265329930749 | 0.957813491602516 | -5.19455878173127 |
| **Prkcb** | 0.0898619375000074 | 24.42149975375 | 0.388674502655911 | 0.709390621535072 | 0.957813491602516 | -5.19461527074431 |
| **Ttf2** | -0.0816683100000013 | 27.011042465 | -0.387800491869962 | 0.710007823848785 | 0.957813491602516 | -5.19489312350559 |
| **Hsd11b1** | -0.0379610849999921 | 29.72778792 | -0.387240769691191 | 0.710403207570445 | 0.957813491602516 | -5.19507075170235 |

| | | | | | | |
|---|---|---|---|---|---|---|
| **Krt79** | 0.235857057499999 | 21.77899257875 | 0.387224533433923 | 0.710414678181849 | 0.957813491602516 | -5.19507590066996 |
| **Gnai1** | -0.114199022500006 | 23.11742058625 | -0.386289979378265 | 0.711075059415154 | 0.957813491602516 | -5.1953719297203 |
| **Slc25a25** | 0.092952179999994 | 25.0378523025 | 0.386157956052865 | 0.711168372262022 | 0.957813491602516 | -5.19541369486967 |
| **H2az1** | -0.0337646850000013 | 24.7987380375 | -0.385101208497231 | 0.711915463806386 | 0.957813491602516 | -5.19574750700225 |
| **Scamp1** | -0.0523003074999977 | 29.59343130375 | -0.384135916643718 | 0.712598197012017 | 0.957813491602516 | -5.19605167336818 |
| **Cdc37** | 0.0350774200000075 | 25.413268245 | 0.383998796407514 | 0.712695202708195 | 0.957813491602516 | -5.19609482178753 |
| **Hspa5** | 0.0789199000000025 | 24.7047056475 | 0.38376725089018 | 0.712859022575431 | 0.957813491602516 | -5.19616765047068 |
| **Dync1i1** | 0.0622514849999902 | 30.2878198325 | 0.381354054886084 | 0.714567344793369 | 0.958447319350789 | -5.19692420525296 |
| **Rps27** | -0.160802459999996 | 25.65792476 | -0.381346373284645 | 0.714572785490081 | 0.958447319350789 | -5.19692660627827 |
| **Rheb** | 0.0867295150000054 | 19.3532174075 | 0.380998912250366 | 0.714818902636406 | 0.958447319350789 | -5.19703516370746 |
| **Ncstn** | -0.0529416250000025 | 25.2377360625 | -0.380476845288701 | 0.715188767220936 | 0.958447319350789 | -5.19719809717562 |
| **Tecpr1** | -0.0680098800000053 | 29.1318662925 | -0.38016743427894 | 0.715408012103189 | 0.958447319350789 | -5.19729456233637 |
| **Dync1li1** | -0.15809131 | 21.560719215 | -0.379440626579154 | 0.715923133110237 | 0.958533707149497 | -5.19752086697742 |
| **Arl6ip5** | 0.0523825475000024 | 27.56961967125 | 0.378422942851078 | 0.716644679033463 | 0.958533707149497 | -5.19783705161124 |
| **Bpnt2** | -0.0432717774999993 | 21.87161453 | -0.0991242884678596 | 0.924373602235794 | 0.999885157113822 | -5.19789531381286 |
| **Myo5a** | 0.0449879774999964 | 27.25684202625 | 0.378188013414049 | 0.716811290267931 | 0.958533707149497 | -5.19790992762837 |
| **Rit1** | 0.137396810000002 | 23.7923878 | 0.376602012061806 | 0.717936512222847 | 0.958533707149497 | -5.19840078898754 |
| **Aldh1b1** | 0.0475849574999962 | 24.07317488875 | 0.376057587940585 | 0.718322939623215 | 0.958533707149497 | -5.19856883526305 |
| **Rpl27a** | -0.0400816275000047 | 26.88897527125 | -0.375581711296566 | 0.718660785455443 | 0.958533707149497 | -5.19871553430064 |
| **Sptbn1** | -0.0467877124999987 | 26.76652002625 | -0.37546224314019 | 0.718745611831114 | 0.958533707149497 | -5.19875233522056 |

| | | | | | | |
|---|---|---|---|---|---|---|
| **Rab1A** | 0.330997502499997 | 22.60075244125 | 0.374846693901892 | 0.7191827400652 | 0.958533707149497 | -5.19894177273019 |
| **Sik3** | -0.0903647925000115 | 25.36681790125 | -0.374805641344507 | 0.719211897302029 | 0.958533707149497 | -5.19895439631588 |
| **Septin9** | -0.0361273675000007 | 24.40487126625 | -0.372945085677364 | 0.720533869976119 | 0.959622570576207 | -5.1995251366892 |
| **Hba** | 0.0407273224999969 | 30.43007363125 | 0.372485155541419 | 0.720860821330112 | 0.959622570576207 | -5.19966580841519 |
| **Nmt1** | -0.0372021550000028 | 21.3043925675 | -0.371208002415112 | 0.721769043434078 | 0.959725301582682 | -5.20005556731086 |
| **Dclk2** | -0.0640194449999925 | 27.3754852425 | -0.370930144731553 | 0.72196670058656 | 0.959725301582682 | -5.20014019497593 |
| **Sel1l** | -0.0732258699999946 | 23.1559575375 | -0.37062188785733 | 0.722186009340068 | 0.959725301582682 | -5.20023401096507 |
| **Coro1b** | 0.0326204625000024 | 27.63396152625 | 0.368257960883117 | 0.723868755540971 | 0.961407725407611 | -5.20095099387556 |
| **Msn** | -0.0623173925000025 | 27.85534561375 | -0.366251185537258 | 0.725298559425855 | 0.961932576216894 | -5.20155622999857 |
| **Nfs1** | -0.0351786325000063 | 26.26642324125 | -0.366108156538702 | 0.725400511114608 | 0.961932576216894 | -5.2015992469795 |
| **Cltc** | -0.0435222825000068 | 28.32057247375 | -0.365574032635094 | 0.725781290011988 | 0.961932576216894 | -5.20175974725896 |
| **Psmd3** | 0.0476391275000054 | 25.94340701875 | 0.365030459976961 | 0.726168890837569 | 0.961932576216894 | -5.20192285799537 |
| **Ppp3ca** | 0.110185324999996 | 23.19456573 | 0.364347038037769 | 0.726656335592392 | 0.961932576216894 | -5.20212760590451 |
| **Septin11** | 0.0608162425000032 | 25.11195073625 | 0.364193723622287 | 0.726765704527979 | 0.961932576216894 | -5.20217348758269 |
| **Cfl2** | -0.0558956725000073 | 24.36624172625 | -0.362384296556548 | 0.72805700246844 | 0.963089165535947 | -5.20271359790585 |
| **Tbc1d24** | 0.0539487674999997 | 23.83441637875 | 0.360852205375453 | 0.729151124934268 | 0.963983750844331 | -5.20316892174749 |
| **Srgap3** | 0.053008782500001 | 28.21850163375 | 0.35976163223523 | 0.729930358206608 | 0.964230481199303 | -5.20349191104227 |
| **Entpd2** | -0.16242038 | 22.34534246 | -0.359421208452827 | 0.730173667384127 | 0.964230481199303 | -5.20359254184138 |
| **Slc12a9** | -0.142086057499998 | 22.27269246375 | -0.358452427454282 | 0.730866262077783 | 0.964592944286869 | -5.2038784211443 |
| **Rtn4** | 0.135956247500005 | 21.23776306875 | 0.355917870790141 | 0.732679530521302 | 0.966433205781957 | -5.20462286876826 |

| | | | | | | |
|---|---|---|---|---|---|---|
| **Atp6v1g2** | -0.123865784999996 | 24.2272930575 | -0.352822042082401 | 0.734896842281865 | 0.968342660587289 | -5.20552533684281 |
| **Ctnna1** | 0.0531368699999994 | 26.2250479025 | 0.352179687345114 | 0.735357255779365 | 0.968342660587289 | -5.20571164781882 |
| **Agpat1** | 0.0533074474999999 | 26.43951696125 | 0.351989824686992 | 0.735493364010682 | 0.968342660587289 | -5.20576665424602 |
| **Chmp6** | -0.0527195900000095 | 25.84077978 | -0.350824839439131 | 0.736328739655407 | 0.968342660587289 | -5.20610355020266 |
| **Dbi** | 0.101948742500007 | 19.90303876125 | 0.35042559083 | 0.736615117313764 | 0.968342660587289 | -5.20621876143653 |
| **H2ac11** | 0.0658182300000014 | 24.897126815 | 0.350383115659425 | 0.736645587052749 | 0.968342660587289 | -5.20623101112997 |
| **Camk2g** | -0.0708548325000073 | 27.25714558875 | -0.349586435127905 | 0.737217183723166 | 0.968542165631745 | -5.20646050829604 |
| **Cacng3** | -0.0525712099999964 | 24.0410184075 | -0.347463901803096 | 0.738740918546512 | 0.969530629770398 | -5.2070695032631 |
| **Dtna** | -0.049798305000003 | 27.314608815 | -0.346933346463374 | 0.739121994440924 | 0.969530629770398 | -5.20722117591132 |
| **Rgs7bp** | -0.104504332499989 | 26.37447271125 | -0.346782531793273 | 0.739230332798496 | 0.969530629770398 | -5.20726424965705 |
| **Flot1** | -0.124427832499997 | 24.27596674125 | -0.344955479894206 | 0.740543309894961 | 0.970648056303966 | -5.20778464621371 |
| **Trip10** | 0.166613694999985 | 23.4941214825 | 0.343231056946047 | 0.741783390639693 | 0.970648056303966 | -5.20827339918348 |
| **Sirt4** | -0.076515947499999 | 24.15494156375 | -0.342780545418674 | 0.74210750259265 | 0.970648056303966 | -5.20840070151671 |
| **Phgdh** | 0.128918595000002 | 23.9241950775 | 0.34211574396117 | 0.74258588443264 | 0.970648056303966 | -5.20858826410017 |
| **Cmpk1** | 0.0518573199999999 | 23.713982495 | 0.341720588385874 | 0.74287029101633 | 0.970648056303966 | -5.2086995854861 |
| **Rp1** | 0.0377746024999972 | 27.67124632625 | 0.34154833910423 | 0.742994278097665 | 0.970648056303966 | -5.20874807221172 |
| **Rac1** | -0.0487973425000092 | 21.71662408125 | -0.341502172442834 | 0.743027510807458 | 0.970648056303966 | -5.2087610637626 |
| **Plxnb1** | 0.175799562500007 | 22.50137816625 | 0.340917040611745 | 0.743448764551507 | 0.970648726553666 | -5.20892557737356 |
| **Tubb6** | -0.0323973325000004 | 24.39646134875 | -0.339588584123671 | 0.744405511367552 | 0.970819903234637 | -5.20929807869846 |
| **Krt10** | 0.279096095 | 21.86094677 | 0.339422774752803 | 0.744524960546759 | 0.970819903234637 | -5.20934447414574 |

| | | | | | | |
|---|---|---|---|---|---|---|
| **Actn4** | -0.0756777699999951 | 27.5061790475 | -0.338474433866087 | 0.745208291548507 | 0.970819903234637 | -5.20960941422681 |
| **Napg** | 0.117210787499996 | 23.3077388037 5 | 0.338398338181022 | 0.745263133345705 | 0.970819903234637 | -5.2096306424711 |
| **Gja1** | -0.0435197574999933 | 29.56243603875 | -0.335822580755126 | 0.747120407019018 | 0.971903454683127 | -5.2103464980511 |
| **Slc4a8** | 0.130895785000003 | 23.69262529 | 0.335724109850835 | 0.747191446437523 | 0.971903454683127 | -5.21037376108504 |
| **Prkca** | 0.147710457500004 | 21.51120807125 | 0.334870986475027 | 0.747807022225087 | 0.971903454683127 | -5.21060963922126 |
| **Rock2** | -0.0655417874999955 | 25.07466075125 | -0.334684652515246 | 0.747941498924466 | 0.971903454683127 | -5.21066108170412 |
| **Caskin1** | 0.0262488399999903 | 28.1664528025 | 0.33432462146695 | 0.748201359131874 | 0.971903454683127 | -5.21076040014588 |
| **Itm2c** | 0.246444937500005 | 22.64959395875 | 0.332937887982781 | 0.749202593587143 | 0.972656377830917 | -5.21114198752131 |
| **Dhodh** | -0.0924963475000098 | 23.34193103625 | -0.330882657011008 | 0.750687448701992 | 0.974035964091955 | -5.21170472629202 |
| **Hibch** | -0.0327319324999955 | 26.82731522875 | -0.329566495487969 | 0.751638943147628 | 0.974722339427532 | -5.2120633451363 |
| **Tstd3** | 0.0532173500000077 | 24.988720695 | 0.32694999161815 | 0.753531878498903 | 0.975489827828202 | -5.21277219566983 |
| **Akap5** | 0.0434996424999952 | 26.82537969875 | 0.326812474315748 | 0.75363141744962 | 0.975489827828202 | -5.21280930106334 |
| **Syngr3** | -0.171344954999995 | 26.067725545 | -0.326586567489085 | 0.753794946227192 | 0.975489827828202 | -5.21287022346726 |
| **Slmap** | -0.0628685024999953 | 27.47146864875 | -0.326410877026538 | 0.753922133947847 | 0.975489827828202 | -5.21291757557448 |
| **Nono** | 0.0650809499999951 | 21.8715894125 | 0.325461931124332 | 0.754609247325485 | 0.975831577118775 | -5.21317291225062 |
| **Fam136a** | -0.100408892500006 | 19.62826349375 | -0.324870235352828 | 0.755037804159952 | 0.975838775460509 | -5.21333176055702 |
| **Plxna4** | -0.1544076325 | 20.64947506875 | -0.323728748078501 | 0.755864829950632 | 0.976360673402077 | -5.21363742230479 |
| **Stk24** | -0.0908737474999946 | 26.30453236375 | -0.322862912381131 | 0.75649237209662 | 0.976372344345168 | -5.21386858150859 |
| **Homer3** | 0.0507997475000082 | 24.65842801625 | 0.321861750278368 | 0.75721824344863 | 0.976372344345168 | -5.21413512762455 |
| **Stt3a** | 0.0483042574999963 | 25.02088693375 | 0.321770493530023 | 0.757284420372899 | 0.976372344345168 | -5.21415938392378 |

| | | | | | | |
|---|---|---|---|---|---|---|
| **Sccpdh** | -0.246977380000004 | 20.954554145 | -0.321347803982497 | 0.757590972019976 | 0.976372344345168 | -5.21427164968316 |
| **H2bc7** | 0.0971556900000081 | 23.6589772125 | 0.319478552513251 | 0.758947192543285 | 0.976372344345168 | -5.21476641775175 |
| **Calr** | -0.0297554700000049 | 26.945684845 | -0.318907624940185 | 0.759361607227267 | 0.976372344345168 | -5.21491698148725 |
| **Ern1** | 0.0725265324999995 | 23.1056832187 5 | 0.318783452559393 | 0.75945175054655 | 0.976372344345168 | -5.21494969362568 |
| **Lrrc57** | -0.0494740975000063 | 30.0858071587 5 | -0.31769948906562 | 0.760238828256646 | 0.976372344345168 | -5.21523473355077 |
| **Myh10** | -0.0657117574999937 | 27.3306688437 5 | -0.317509884202033 | 0.760376533924932 | 0.976372344345168 | -5.2152844961403 |
| **Rps18** | -0.0252956499999968 | 31.40841245 | -0.317280706995293 | 0.7605429925862 | 0.976372344345168 | -5.21534460647041 |
| **Atp6v1a** | 0.185879734999997 | 24.7967293725 | 0.317005284572421 | 0.760743058711434 | 0.976372344345168 | -5.21541679104462 |
| **Tuba1a** | -0.0456887100000003 | 25.87080565 | -0.315345404268335 | 0.761949208944307 | 0.976372344345168 | -5.21585054515596 |
| **Preb** | -0.0342461250000028 | 30.32230827 | -0.315288352858958 | 0.76199067799821 | 0.976372344345168 | -5.21586541463477 |
| **Septin8** | 0.0565105275000057 | 24.3683006387 5 | 0.315009658997734 | 0.762193264792082 | 0.976372344345168 | -5.21593801418661 |
| **Hnrnpa1** | 0.0538937075000021 | 26.7207500312 5 | 0.314640604521159 | 0.762461567027329 | 0.976372344345168 | -5.21603405741498 |
| **Arf2** | -0.0772177000000056 | 27.1408648575 | -0.314387751058046 | 0.762645411577804 | 0.976372344345168 | -5.21609979770755 |
| **Plxna1** | -0.129611907500003 | 22.8056057512 5 | -0.312005519905965 | 0.764378292401324 | 0.977080923824173 | -5.21671666165533 |
| **Krt5** | 0.161425012499997 | 22.0844566437 5 | 0.31174800656661 | 0.764565699999369 | 0.977080923824173 | -5.21678307221157 |
| **Tsfm** | 0.0569290850000002 | 26.6512335875 | 0.311170717722365 | 0.764985888838052 | 0.977080923824173 | -5.21693175807841 |
| **Gnal** | 0.24693317749999 | 20.3928897637 5 | 0.310346744035191 | 0.765585778996176 | 0.977080923824173 | -5.21714351938758 |
| **Gria4** | 0.103763099999995 | 26.21083595 | 0.310079899972896 | 0.76578009077225 | 0.977080923824173 | -5.21721198223705 |
| **Krt31** | 0.0443056624999869 | 26.3603311287 5 | 0.309959688908474 | 0.765867632577831 | 0.977080923824173 | -5.21724280562937 |
| **Atp4a** | 0.1232382775 | 20.6081725262 5 | 0.309021579553272 | 0.766550922506759 | 0.977080923824173 | -5.21748295071439 |

| | | | | | | |
|---|---|---|---|---|---|---|
| **Apex1** | 0.211236650000011 | 23.982730195 | 0.308971899671561 | 0.766587114053643 | 0.977080923824173 | -5.21749564860502 |
| **Gpd1l** | 0.0569926125000038 | 23.36010775875 | 0.306924363908475 | 0.768079279284369 | 0.977873393715142 | -5.21801727286051 |
| **Gphn** | -0.0981018075000044 | 26.25802900375 | -0.30666779360142 | 0.768266332792914 | 0.977873393715142 | -5.21808239993103 |
| **Syngap1** | 0.0610223224999942 | 28.11177683375 | 0.305808399605807 | 0.768892998314062 | 0.977873393715142 | -5.21830016297656 |
| **Hnrnph1** | -0.0996027224999985 | 22.08655852375 | -0.305792834090145 | 0.76890435032478 | 0.977873393715142 | -5.21830410170391 |
| **Tfrc** | 0.176748804999999 | 23.4587290525 | 0.304206640951398 | 0.770061489209326 | 0.978805430085904 | -5.21870445978783 |
| **Ube2v1** | 0.0885578974999994 | 19.10377752875 | 0.303409465679694 | 0.770643272894298 | 0.979005523440059 | -5.21890490890904 |
| **Tln2** | 0.0461992749999958 | 28.67316652 | 0.302795714118278 | 0.771091301295255 | 0.979035570769484 | -5.21905889004518 |
| **Hnrnph2** | -0.101299264999994 | 21.32378625 | -0.300891067841471 | 0.772482260558953 | 0.98026214252448 | -5.21953481892096 |
| **Snd1** | 0.120204740000005 | 25.62593035 | 0.300261569868006 | 0.772942180330087 | 0.980306547565426 | -5.21969147822644 |
| **Gabrb1** | 0.0398889600000025 | 24.27018391 | 0.29914482274758 | 0.773758332987719 | 0.980640420313128 | -5.21996861565025 |
| **Ephb2** | -0.041435829999994 | 25.8655224575 | -0.298172331152026 | 0.77446931084411 | 0.980640420313128 | -5.22020914039348 |
| **Aplp2** | 0.0439377850000042 | 23.966711835 | 0.298156831931765 | 0.774480644044439 | 0.980640420313128 | -5.22021296765725 |
| **Eci2** | 0.116686577500005 | 21.38199414125 | 0.295918330423393 | 0.776118082314619 | 0.981804543292429 | -5.22076370533515 |
| **Kcna6** | -0.030887879999991 | 25.3044765275 | -0.295039693142742 | 0.7767611316857 | 0.981804543292429 | -5.22097877846371 |
| **Gcat** | 0.0797153424999983 | 22.20210850875 | 0.294838292865249 | 0.776908557377234 | 0.981804543292429 | -5.22102799008177 |
| **Bag6** | 0.0365243325000009 | 27.30872596125 | 0.294573585719016 | 0.777102338990887 | 0.981804543292429 | -5.2210926210891 |
| **Stx4** | 0.0471773999999954 | 20.7385341875 | 0.293197047573458 | 0.778110323610861 | 0.982410934355161 | -5.22142781111151 |
| **Hnrnpd** | 0.136085654999995 | 20.737206525 | 0.292755187231103 | 0.778433978327366 | 0.982410934355161 | -5.22153508274024 |
| **Ero1a** | -0.0952018899999949 | 24.145628225 | -0.291650501397459 | 0.779243348099875 | 0.982894698778792 | -5.22180258472536 |

| | | | | | | |
|---|---|---|---|---|---|---|
| **Akap12** | 0.395611997500001 | 24.30249354375 | 0.287953310084715 | 0.781954311408338 | 0.985253396035326 | -5.22269073903002 |
| **Esyt2** | 0.0678861074999944 | 25.70938400875 | 0.287325086690683 | 0.782415282114755 | 0.985253396035326 | -5.22284056171094 |
| **Armc10** | -0.0422308724999922 | 31.98828792375 | -0.286215331899789 | 0.783229815831128 | 0.985253396035326 | -5.22310444771375 |
| **Enpep** | 0.0661408725000001 | 23.41415613125 | 0.285741584365989 | 0.783577624669564 | 0.985253396035326 | -5.22321679739996 |
| **Psd3** | -0.038992287499994 | 29.56531474625 | -0.284418186701401 | 0.784549499367357 | 0.985253396035326 | -5.22352968585134 |
| **Gabra3** | 0.0925723574999928 | 29.42930163375 | 0.284383803347869 | 0.784574755295418 | 0.985253396035326 | -5.22353779626299 |
| **Rab8a** | 0.159155725000002 | 22.7355146525 | 0.283754138058981 | 0.785037318733004 | 0.985253396035326 | -5.22368615468116 |
| **Trim2** | 0.0449807299999918 | 26.5984127975 | 0.282942483125052 | 0.785633713233263 | 0.985253396035326 | -5.22387692175139 |
| **Kcnd2** | 0.0549157850000022 | 20.2435481875 | 0.282616599719395 | 0.785873212239319 | 0.985253396035326 | -5.22395336644623 |
| **Adgrb1** | -0.0435014625000107 | 28.41714433625 | -0.282317270936778 | 0.786093217692961 | 0.985253396035326 | -5.22402350670324 |
| **Atl3** | -0.0373247624999955 | 25.13644205875 | -0.282184486944684 | 0.786190820144368 | 0.985253396035326 | -5.22405459822955 |
| **Vbp1** | 0.0418035725000045 | 28.40503982875 | 0.281750354724345 | 0.786509956581631 | 0.985253396035326 | -5.22415615169805 |
| **Grm4** | 0.0479965050000004 | 25.7344228125 | 0.281539111549298 | 0.786665260293486 | 0.985253396035326 | -5.22420551141763 |
| **Dync1i2** | -0.124807337499995 | 23.08454494375 | -0.279786555717422 | 0.787954123688315 | 0.986332155913697 | -5.2246136327038 |
| **Camlg** | 0.0924168850000022 | 23.61254506 | 0.27851292288005 | 0.788891227843715 | 0.986969665203607 | -5.22490867328741 |
| **Txnrd2** | -0.0350827299999992 | 26.9919427975 | -0.277451181712514 | 0.789672716144141 | 0.987069306004667 | -5.22515362910988 |
| **Ubxn4** | -0.0350555649999968 | 27.8252901525 | -0.276457149730076 | 0.790404604200481 | 0.987069306004667 | -5.22538213966471 |
| **Sh3gl1** | -0.0603776975000052 | 28.67462617375 | -0.276104579722855 | 0.79066425007301 | 0.987069306004667 | -5.22546299785281 |
| **Tmem266** | -0.0558964350000011 | 26.7133364625 | -0.276080064684215 | 0.79068230494002 | 0.987069306004667 | -5.22546861638772 |
| **Hnrnpab** | -0.0568167474999939 | 27.85879473625 | -0.275480753656492 | 0.791123729376856 | 0.987086232380967 | -5.2256058200244 |

| | | | | | | |
|---|---|---|---|---|---|---|
| **Timm22** | -0.0649213025000037 | 25.40788778125 | -0.273770709350246 | 0.792383721883783 | 0.988123916965344 | -5.22599571653691 |
| **Nlgn1** | -0.0725338450000024 | 23.132005415 | -0.272556089333516 | 0.793279083333943 | 0.988706021205514 | -5.22627122050258 |
| **Golga5** | 0.071604709999999 | 29.3283309925 | 0.270651868408941 | 0.79468346305285 | 0.98992157087631 | -5.22670074341929 |
| **Caprin1** | 0.0752637549999982 | 25.91950373 | 0.269578237138362 | 0.795475638094787 | 0.990373608788275 | -5.2269416238091 |
| **Acsf2** | 0.0640687950000043 | 26.80899759 | 0.268436917462874 | 0.79631804212717 | 0.990887660834618 | -5.2271966689938 |
| **Cryzl2** | 0.143742325000009 | 22.7102629475 | 0.267681956644441 | 0.796875437107084 | 0.991046702644767 | -5.22736479760118 |
| **Hexb** | 0.0471672174999931 | 27.40825519875 | 0.266236696910364 | 0.797942842959808 | 0.991389288010703 | -5.22768536858343 |
| **Rbmxl1** | -0.0791899049999998 | 25.15785487 | -0.265972493965392 | 0.798138022222394 | 0.991389288010703 | -5.22774378834294 |
| **Rpl5** | -0.0957501399999963 | 21.646126755 | -0.265306462050085 | 0.798630120694472 | 0.991389288010703 | -5.22789080860605 |
| **Cox7a2l** | 0.0728096649999941 | 28.2608279725 | 0.26489082983941 | 0.7989372610874 | 0.991389288010703 | -5.22798237355107 |
| **Slc7a8** | 0.0544071200000005 | 21.18423801 | 0.263683048240607 | 0.799829995043541 | 0.991389288010703 | -5.22824765753727 |
| **Sh3bgrl** | 0.0975835575000019 | 24.12234530375 | 0.262730993600938 | 0.800533935571132 | 0.991389288010703 | -5.22845593961892 |
| **Myo1d** | 0.138789875000008 | 25.495315645 | 0.262376056236616 | 0.800796424228465 | 0.991389288010703 | -5.22853340179228 |
| **Padi2** | 0.286851332500007 | 25.28603357125 | 0.257935550605623 | 0.804082669142407 | 0.991389288010703 | -5.22949387680072 |
| **Pam16** | -0.0401479175000006 | 24.46388108625 | -0.256773938389776 | 0.804943041648268 | 0.991389288010703 | -5.22974249332409 |
| **Pdap1** | -0.0841299599999985 | 23.286132345 | -0.256101901169349 | 0.805440933352436 | 0.991389288010703 | -5.2298858275741 |
| **Dnpep** | 0.221161777500008 | 24.77031494125 | 0.256093166847989 | 0.805447404983838 | 0.991389288010703 | -5.22988768804533 |
| **Sncg** | -0.0433847274999906 | 29.55812442375 | -0.255442267230329 | 0.80592973052565 | 0.991389288010703 | -5.23002615994847 |
| **Cops4** | -0.0463719199999986 | 25.4499613925 | -0.254818744861559 | 0.806391854600204 | 0.991389288010703 | -5.23015848511162 |
| **Macrod1** | 0.0524628325000052 | 23.76409120625 | 0.254722220162394 | 0.806463401415608 | 0.991389288010703 | -5.23017894156292 |

| | | | | | | |
|---|---|---|---|---|---|---|
| **H3-5** | -0.0589532149999954 | 21.642835735 | -0.254594807179348 | 0.806557846553874 | 0.991389288010703 | -5.23020593257596 |
| **Thnsl1** | 0.0695716249999982 | 23.8438652275 | 0.254496026436153 | 0.806631070380353 | 0.991389288010703 | -5.23022684909876 |
| **Rhot2** | 0.203826605000003 | 19.838475435 | 0.254146881850429 | 0.806889899751612 | 0.991389288010703 | -5.23030071591134 |
| **Capza1** | -0.0330056474999942 | 30.45131907375 | -0.252868229704606 | 0.807838018214406 | 0.991389288010703 | -5.23057038891212 |
| **H3c1** | 0.0735837450000005 | 21.90699899 | 0.252358669472616 | 0.808215953462639 | 0.991389288010703 | -5.23067748717929 |
| **Chchd2** | 0.0313009175000261 | 34.13742942375 | 0.251710271815568 | 0.80869694274661 | 0.991389288010703 | -5.23081346098279 |
| **Ilf2** | 0.0889600225000002 | 22.65390288625 | 0.251653529535101 | 0.80873903911009 | 0.991389288010703 | -5.23082534400425 |
| **Iscu** | -0.02903410750001 | 24.37343227875 | -0.250285279811252 | 0.809754332374697 | 0.991389288010703 | -5.23111109182919 |
| **Clstn1** | -0.0397140699999952 | 20.87174571 | -0.250047875781764 | 0.809930535473547 | 0.991389288010703 | -5.23116051680291 |
| **Agpat5** | -0.0614314799999995 | 21.4481376725 | -0.2498344661865 | 0.81008893986536 | 0.991389288010703 | -5.23120490728418 |
| **Plxnc1** | -0.136437397499996 | 23.34545413375 | -0.249512058187098 | 0.810328267097153 | 0.991389288010703 | -5.23127189986142 |
| **Kbtbd11** | -0.088035289999997 | 24.5836710925 | -0.247599274993695 | 0.811748597605784 | 0.991389288010703 | -5.23166761515035 |
| **Reep2** | -0.037706914999994 | 28.5714913375 | -0.247488432120409 | 0.811830927092651 | 0.991389288010703 | -5.23169045502187 |
| **Syncrip** | 0.0994057999999924 | 26.5253875675 | 0.247456119600378 | 0.811854927975066 | 0.991389288010703 | -5.23169711133827 |
| **Fam177a1** | -0.133378927500004 | 22.79790963625 | -0.246790399947864 | 0.81234945537461 | 0.991389288010703 | -5.23183405924869 |
| **Sigmar1** | -0.0276800175000034 | 26.57350492625 | -0.246373831485595 | 0.812658948724896 | 0.991389288010703 | -5.2319195698181 |
| **Tdrkh** | 0.0456537049999959 | 25.54624112 | 0.245762229601778 | 0.813113409354351 | 0.991389288010703 | -5.2320448597432 |
| **Psmd2** | -0.0326567099999977 | 26.3904011 | -0.245601710049501 | 0.813232698866787 | 0.991389288010703 | -5.23207769259437 |
| **Mapk9** | 0.0510655149999977 | 21.820973135 | 0.244185075478375 | 0.814285697063736 | 0.991389288010703 | -5.2323665431859 |
| **Tubal3** | -0.0312504399999938 | 27.347030665 | -0.242836209441152 | 0.815288707269914 | 0.991389288010703 | -5.23264005701493 |

| | | | | | | |
|---|---|---|---|---|---|---|
| **Tjp2** | -0.0342268924999942 | 31.29523321875 | -0.242746166971182 | 0.815355675736356 | 0.991389288010703 | -5.23265826244491 |
| **Sar1b** | 0.0335095224999975 | 27.45634778875 | 0.242489215829269 | 0.815546790527407 | 0.991389288010703 | -5.23271017834548 |
| **Rpl22** | -0.0428203350000018 | 26.5521486675 | -0.2424221537001 | 0.815596672140715 | 0.991389288010703 | -5.2327237191163 |
| **Grm7** | -0.0542729875000063 | 23.16342149125 | -0.242312037501418 | 0.815678579885707 | 0.991389288010703 | -5.23274594515122 |
| **Aplp1** | -0.0521288424999966 | 30.41041798375 | -0.242223602148506 | 0.815744362564138 | 0.991389288010703 | -5.23276378793558 |
| **Cacna2d3** | 0.0451531174999964 | 22.45726117125 | 0.242223184637903 | 0.815744673133395 | 0.991389288010703 | -5.23276387215772 |
| **Abhd3** | -0.0939529325000024 | 20.62047676375 | -0.24203040455736 | 0.815888078255666 | 0.991389288010703 | -5.23280274546319 |
| **Fabp7** | -0.102169182499988 | 24.15974646125 | -0.240220483837938 | 0.817234811224957 | 0.991389288010703 | -5.23316623156027 |
| **Dusp3** | 0.0840077174999969 | 22.61730900125 | 0.239822698496686 | 0.817530886360596 | 0.991389288010703 | -5.23324576082431 |
| **Slc25a42** | 0.0982068599999906 | 23.04940891 | 0.239718903832427 | 0.817608146931012 | 0.991389288010703 | -5.23326649128038 |
| **Rps21** | -0.0384517149999937 | 26.8140879175 | -0.239204580989759 | 0.817991020396839 | 0.991389288010703 | -5.23336908513843 |
| **Prep** | 0.0356056625000178 | 28.15376502625 | 0.239123274848957 | 0.818051551419859 | 0.991389288010703 | -5.23338528382775 |
| **Tpd52** | 0.0762065324999952 | 27.57688958125 | 0.239119579670244 | 0.818054302448641 | 0.991389288010703 | -5.23338601989318 |
| **Psmd1** | -0.174153805000003 | 22.4139151875 | -0.237836773742153 | 0.81900950724044 | 0.991389288010703 | -5.23364087696932 |
| **Fam241b** | -0.0900504825000006 | 21.81072446875 | -0.2359271986047 | 0.820432031455696 | 0.991389288010703 | -5.23401776879609 |
| **C1orf198** | -0.0291079150000009 | 19.3945218075 | -0.235309093862004 | 0.820892640617858 | 0.991389288010703 | -5.23413912632439 |
| **Maip1** | 0.150311350000006 | 22.374910265 | 0.23520980118389 | 0.820966640231943 | 0.991389288010703 | -5.2341585921922 |
| **Rplp1** | 0.0470826124999988 | 23.11026945375 | 0.234746885292182 | 0.821311662378745 | 0.991389288010703 | -5.23424923847569 |
| **Ppp5c** | -0.0192566800000016 | 27.36984832 | -0.234602481949327 | 0.821419298333467 | 0.991389288010703 | -5.23427747914667 |
| **Rras** | 0.0350098799999934 | 23.84703761 | 0.234469936999666 | 0.82151809888233 | 0.991389288010703 | -5.2343033857111 |

| | | | | | | |
|---|---|---|---|---|---|---|
| **Rab15** | -0.0840225374999939 | 23.09195682625 | -0.233321188967079 | 0.822374535194553 | 0.991389288010703 | -5.23452731327575 |
| **Plxnb2** | 0.0829024474999969 | 29.06773374625 | 0.231737904907695 | 0.82355536290464 | 0.991389288010703 | -5.23483417935607 |
| **Rbm3** | 0.0265926524999927 | 23.93547221875 | 0.230995951606177 | 0.824108888333645 | 0.991389288010703 | -5.234977277196 |
| **Rpl9** | -0.0331209775000154 | 27.44784897375 | -0.229993761861015 | 0.824856731424759 | 0.991389288010703 | -5.23516985167779 |
| **Prpsap2** | 0.100820022500002 | 25.25583986875 | 0.228426455757081 | 0.826026662535457 | 0.991389288010703 | -5.23546936939281 |
| **Emb** | 0.096292637500003 | 20.25179208375 | 0.228331490684455 | 0.826097565503311 | 0.991389288010703 | -5.23548745301895 |
| **Pgp** | -0.159798059999996 | 21.5512653075 | -0.227769503869669 | 0.826517192756905 | 0.991389288010703 | -5.23559431781915 |
| **Comt** | -0.0833577500000047 | 22.1138915525 | -0.227110687593017 | 0.827009198985068 | 0.991389288010703 | -5.23571926636179 |
| **Sorcs2** | -0.065566900000011 | 24.5184824225 | -0.225313313475936 | 0.828351909610501 | 0.991389288010703 | -5.23605834360237 |
| **Myh14** | -0.0613173000000025 | 22.481095725 | -0.224976046338436 | 0.82860393083571 | 0.991389288010703 | -5.23612167498523 |
| **Dip2b** | 0.113866649999999 | 20.857384135 | 0.223925706166705 | 0.829388931754492 | 0.991389288010703 | -5.23631830941437 |
| **Ppp2r5e** | 0.0315321125000061 | 26.91995628375 | 0.223620385537524 | 0.829617161092816 | 0.991389288010703 | -5.23637529914667 |
| **Chgb** | 0.0389172349999996 | 25.5589105075 | 0.223055360504887 | 0.830039568099188 | 0.991389288010703 | -5.23648056278539 |
| **Fam171a2** | -0.0477905274999983 | 24.82047262875 | -0.222617980745361 | 0.830366590421842 | 0.991389288010703 | -5.23656186672971 |
| **Grid1** | -0.0456155174999964 | 24.07433275625 | -0.222168965824477 | 0.830702349899002 | 0.991389288010703 | -5.23664517055183 |
| **Mark2** | 0.0718108549999954 | 22.70133832 | 0.221924314263213 | 0.830885308830814 | 0.991389288010703 | -5.23669049021735 |
| **Vps37b** | -0.0320148999999965 | 22.29252358 | -0.221886936923018 | 0.830913261899955 | 0.991389288010703 | -5.23669740974155 |
| **Rab31** | -0.0290269049999949 | 26.4127499575 | -0.22181072339949 | 0.830970259868559 | 0.991389288010703 | -5.23671151531459 |
| **Nedd4** | -0.0418909100000064 | 25.2673340275 | -0.221612921091218 | 0.831118195816477 | 0.991389288010703 | -5.2367481023008 |
| **Tpp1** | -0.0409129700000044 | 27.5155070025 | -0.221370131524024 | 0.831299787739098 | 0.991389288010703 | -5.23679296665359 |
| **Lxn** | 0.0659609900000042 | 21.5025295575 | 0.220948624996152 | 0.831615075538003 | 0.991389288010703 | -5.23687074088716 |

| | | | | | | |
|---|---|---|---|---|---|---|
| **Sh3gl3** | -0.0890254274999975 | 24.39295485875 | -0.220542057623445 | 0.831919220500261 | 0.991389288010703 | -5.2369456207152 |
| **Celf2** | 0.0287572325000021 | 31.09770209125 | 0.220245452998814 | 0.832141124115584 | 0.991389288010703 | -5.23700016263603 |
| **Ube2o** | -0.0390284775000076 | 27.88504856625 | -0.220179884501057 | 0.832190181165273 | 0.991389288010703 | -5.23701221014252 |
| **Naca** | 0.053189502500004 | 22.93215420625 | 0.220109219293918 | 0.832243052374884 | 0.991389288010703 | -5.23702519017031 |
| **Rpl21** | -0.067955642500003 | 25.58530951625 | -0.220049475909406 | 0.832287752684454 | 0.991389288010703 | -5.23703616084813 |
| **Ptpn5** | 0.177990229999999 | 16.9760418075 | 0.219914374537169 | 0.832388838698193 | 0.991389288010703 | -5.23706095872793 |
| **Rps23** | 0.0575157750000059 | 22.84722667 | 0.216888229859421 | 0.834653963780021 | 0.993084733406119 | -5.23761248749862 |
| **Fsd1** | 0.0394365475000029 | 28.62390047625 | 0.216862409892311 | 0.834673297821615 | 0.993084733406119 | -5.23761716100255 |
| **Chrm1** | -0.0600354899999935 | 28.9632917975 | -0.214934259017301 | 0.836117448442533 | 0.994089572580149 | -5.23796461794193 |
| **Acbd3** | 0.0283358000000007 | 25.9590299975 | 0.213866346642735 | 0.836917589588608 | 0.994089572580149 | -5.2381557455966 |
| **Ppp1r14a** | 0.0480825275000001 | 26.15187718625 | 0.213041842796801 | 0.837535497611638 | 0.994089572580149 | -5.23830266942154 |
| **Pomgnt2** | 0.142349710000001 | 22.1790795675 | 0.212477164159772 | 0.837958756177764 | 0.994089572580149 | -5.23840297132327 |
| **Ptpn11** | 0.234290962499987 | 27.34890629125 | 0.211265371536918 | 0.838867258412595 | 0.994089572580149 | -5.23861733415808 |
| **Blvrb** | 0.0337473849999981 | 26.047104865 | 0.211112298871299 | 0.838982038566913 | 0.994089572580149 | -5.23864432652369 |
| **Pcdhga4** | -0.0948615824999983 | 22.09560010625 | -0.210679674247361 | 0.839306461073803 | 0.994089572580149 | -5.23872051018244 |
| **Eif2s1** | 0.0737954550000097 | 21.2696126825 | 0.210673456362661 | 0.839311124072542 | 0.994089572580149 | -5.23872160400935 |
| **Nucb2** | 0.0403730799999948 | 30.5133806725 | 0.208371302205097 | 0.841038062860144 | 0.994089572580149 | -5.23912440765822 |
| **Timmdc1** | 0.0417711699999899 | 23.07862655 | 0.208371250015028 | 0.841038102020764 | 0.994089572580149 | -5.23912441674046 |
| **Rpl11** | 0.0601653125000041 | 23.54777257625 | 0.208004389275368 | 0.841313386568652 | 0.994089572580149 | -5.2391882034252 |
| **Gcdh** | -0.0357666700000046 | 24.8923754425 | -0.20735601378305 | 0.841799972364387 | 0.994089572580149 | -5.23930066706181 |

| | | | | | | |
|---|---|---|---|---|---|---|
| **Dhrs7b** | 0.292036029999991 | 25.8181840575 | 0.207160569453608 | 0.841946661937818 | 0.994089572580149 | -5.23933449998327 |
| **Kcnj6** | -0.0531655200000003 | 25.0880929875 | -0.207131174688456 | 0.841968724588903 | 0.994089572580149 | -5.23933958572773 |
| **Mogs** | -0.0857746475000063 | 22.61181922875 | -0.206856136841312 | 0.8421751654978 | 0.994089572580149 | -5.23938713707389 |
| **Sh3glb1** | 0.0694738424999954 | 24.14443350625 | 0.206376273233441 | 0.842535378805907 | 0.994089572580149 | -5.23946995189354 |
| **Akr1b8** | -0.0450692024999988 | 21.24644537875 | -0.205966281231941 | 0.842843174671336 | 0.994089572580149 | -5.23954055831038 |
| **Tjp1** | 0.0352206949999854 | 25.1824978175 | 0.205051868103041 | 0.843529764419295 | 0.994390989532607 | -5.23969753529426 |
| **Mif** | 0.0276213275000075 | 28.12472176625 | 0.204236560570867 | 0.844142064227712 | 0.99460456699353 | -5.23983691898601 |
| **Snx27** | -0.0430956100000159 | 26.382832535 | -0.203096129605482 | 0.844998728183681 | 0.995105699805897 | -5.24003096797682 |
| **Strap** | 0.0294517100000036 | 28.74124149 | 0.201814079846211 | 0.845962042787732 | 0.995302152959441 | -5.24024783637148 |
| **Zdhhc5** | -0.0423401224999935 | 24.21943577875 | -0.20137306529474 | 0.846293480857191 | 0.995302152959441 | -5.24032212464473 |
| **Purg** | -0.0327395424999963 | 19.91316371125 | -0.20115173856286 | 0.846459828394635 | 0.995302152959441 | -5.2403593464912 |
| **Susd2** | -0.0487577375000008 | 19.54563714375 | -0.200502001451014 | 0.846948214758122 | 0.995369093961787 | -5.24046838375751 |
| **Hapln1** | 0.0761096150000071 | 24.4697361225 | 0.198156094605215 | 0.848712159279957 | 0.996656670771414 | -5.24085917506996 |
| **Anxa4** | 0.019747182499998 | 29.66478017375 | 0.197895948191998 | 0.848907827510112 | 0.996656670771414 | -5.24090223223046 |
| **Mboat7** | -0.0641170524999986 | 22.26815424625 | -0.197154871754258 | 0.849465288531383 | 0.996803876216633 | -5.24102458313101 |
| **Vcl** | -0.0356108124999963 | 27.32944500125 | -0.196056767924842 | 0.850291487502871 | 0.997266121845004 | -5.24120504693091 |
| **H3c2** | -0.0294048599999925 | 25.3015360125 | -0.192258276958148 | 0.853150980443671 | 0.999885157113822 | -5.24182163101311 |
| **Pepd** | 0.0391721975000046 | 23.64308895375 | 0.19149300805192 | 0.853727362516088 | 0.999885157113822 | -5.24194441225507 |
| **Anln** | 0.0181859150000108 | 30.1102107925 | 0.187435301438859 | 0.856785130599397 | 0.999885157113822 | -5.24258736558972 |
| **Sarnp** | 0.0298038449999893 | 28.781139265 | 0.187195525040113 | 0.856965902633969 | 0.999885157113822 | -5.24262493350991 |

| | | | | | | |
|---|---|---|---|---|---|---|
| **Srprb** | -0.0859639549999933 | 24.5795735325 | -0.186127563400371 | 0.85777117173709 | 0.999885157113822 | -5.24279168409334 |
| **Mrps35** | -0.0324017575000077 | 23.37225857875 | -0.185937285601626 | 0.857914665036867 | 0.999885157113822 | -5.2428212950544 |
| **Eprs1** | -0.097011162500003 | 25.29318666875 | -0.185511351982522 | 0.858235893340893 | 0.999885157113822 | -5.24288747031265 |
| **Ddb1** | -0.0374680425000022 | 26.17570147125 | -0.18182290497888 | 0.861018831046697 | 0.999885157113822 | -5.2434542585161 |
| **Cryz** | 0.0426182174999994 | 20.51903304875 | 0.181613825041157 | 0.861176646323526 | 0.999885157113822 | -5.24348605027556 |
| **Ewsr1** | 0.0320407775000007 | 26.41837412875 | 0.180520911052475 | 0.862001698052802 | 0.999885157113822 | -5.24365164577727 |
| **Pnpla6** | 0.045091780000007 | 23.396847095 | 0.179662237842322 | 0.862650049478236 | 0.999885157113822 | -5.24378105702884 |
| **Rpl13a** | -0.0218935250000065 | 29.392377075 | -0.179346774820146 | 0.862888272296336 | 0.999885157113822 | -5.24382844755657 |
| **Rtcb** | 0.0267413799999972 | 23.0429376575 | 0.179257749221861 | 0.862955503020507 | 0.999885157113822 | -5.24384180656551 |
| **Ube2k** | 0.0573556074999999 | 24.01300833875 | 0.178820773044134 | 0.863285518269829 | 0.999885157113822 | -5.24390728332686 |
| **Trio** | 0.0470163599999971 | 29.4630450275 | 0.178721770007939 | 0.863360291911772 | 0.999885157113822 | -5.24392209605712 |
| **Acp1** | -0.0798891725000033 | 22.32112525375 | -0.17816788040936 | 0.863778653816634 | 0.999885157113822 | -5.24400481889367 |
| **Rpl36** | 0.0772731150000077 | 26.315556995 | 0.177468432362786 | 0.864307025733016 | 0.999885157113822 | -5.24410891818625 |
| **Tmem256** | -0.0602575274999992 | 20.56650357875 | -0.177275122503424 | 0.86445306766989 | 0.999885157113822 | -5.24413761723522 |
| **Nrn1** | -0.0385121900000023 | 23.63182496 | -0.177088114847741 | 0.864594353848833 | 0.999885157113822 | -5.2441653512319 |
| **Nars1** | 0.0264613900000015 | 25.473808395 | 0.176950442241046 | 0.864698370312592 | 0.999885157113822 | -5.24418575015018 |
| **Hsdl2** | 0.0276680575000015 | 28.52319494375 | 0.176935020918974 | 0.864710021840362 | 0.999885157113822 | -5.2441880341473 |
| **Pgm2l1** | 0.061414172500001 | 20.08899489625 | 0.176819073320907 | 0.864797626818378 | 0.999885157113822 | -5.24420520043248 |
| **Hccs** | -0.0260632600000044 | 25.5230333225 | -0.176159636228472 | 0.865295907852 | 0.999885157113822 | -5.24430261995399 |
| **Tpp2** | 0.0270307649999957 | 31.127854455 | 0.174416018874024 | 0.866613730323307 | 0.999885157113822 | -5.24455847324789 |

| | | | | | | |
|---|---|---|---|---|---|---|
| **Slc20a1** | 0.034498627499989 | 22.98360231125 | 0.173951212274066 | 0.866965107660475 | 0.999885157113822 | -5.24462625281243 |
| **Cnnm2** | -0.0787973949999952 | 22.1489620325 | -0.172821677598073 | 0.867819130942307 | 0.999885157113822 | -5.24479021970947 |
| **Kcnma1** | -0.0416346875000002 | 23.58947649375 | -0.172285455650952 | 0.86822462658602 | 0.999885157113822 | -5.24486768964705 |
| **Ppp1r2** | -0.0637978975000024 | 23.76521919875 | -0.171887178851873 | 0.868525834619253 | 0.999885157113822 | -5.24492507604875 |
| **Hdgfl3** | -0.0520913275000048 | 23.02848167625 | -0.170935031834344 | 0.869246017932136 | 0.999885157113822 | -5.2450617353726 |
| **Lyrm4** | -0.0432836124999945 | 24.77877316875 | -0.170819283432856 | 0.869333576648532 | 0.999885157113822 | -5.24507829726828 |
| **Tmx1** | 0.063544097500003 | 24.42235228875 | 0.1707694679426 | 0.86937126054548 | 0.999885157113822 | -5.24508542171729 |
| **Clpb** | 0.0231460800000036 | 22.799049065 | 0.16864393476448 | 0.870979501235097 | 0.999885157113822 | -5.24538749332943 |
| **Mrps34** | -0.0240446875000089 | 26.71831475625 | -0.167413851399891 | 0.871910519718289 | 0.999885157113822 | -5.24556059737799 |
| **Rab9b** | -0.16254613000001 | 21.024680245 | -0.166587085125652 | 0.872536401034942 | 0.999885157113822 | -5.24567623972904 |
| **Myadm** | -0.0123236875000003 | 27.61672428625 | -0.166264869511372 | 0.87278035245005 | 0.999885157113822 | -5.24572115557706 |
| **Msra** | -0.0637534800000026 | 22.7745359925 | -0.166189829396627 | 0.872837167924884 | 0.999885157113822 | -5.24573160357786 |
| **Srr** | -0.0198349299999983 | 24.2213164625 | -0.166172541107239 | 0.872850257603428 | 0.999885157113822 | -5.2457340100029 |
| **Mrpl10** | -0.0347166225000066 | 22.50148103625 | -0.166030281238884 | 0.87295797006103 | 0.999885157113822 | -5.24575380230549 |
| **Ntrk3** | 0.0144665500000016 | 28.822272125 | 0.164975498130076 | 0.873756692543905 | 0.999885157113822 | -5.24590002854232 |
| **Actr3b** | 0.0548026849999985 | 28.0871304525 | 0.16447318714881 | 0.874137117683885 | 0.999885157113822 | -5.24596934043061 |
| **Pacs1** | -0.0708457800000026 | 23.61037883 | -0.163740566775833 | 0.874692031951625 | 0.999885157113822 | -5.24607005664442 |
| **Cbx1** | 0.0686288674999993 | 24.04204400875 | 0.163607123584116 | 0.874793115049498 | 0.999885157113822 | -5.24608835368137 |
| **Kiaa0513** | -0.0566184599999957 | 24.6138951675 | -0.163003473876573 | 0.875250410631514 | 0.999885157113822 | -5.24617093846342 |
| **Atg9a** | -0.0142040574999989 | 31.32447283625 | -0.159694374609154 | 0.877758131379765 | 0.999885157113822 | -5.24661828240688 |

| | | | | | | |
|---|---|---|---|---|---|---|
| **Camk4** | 0.0250961000000025 | 23.01015507 | 0.159502164674345 | 0.877903839862547 | 0.999885157113822 | -5.24664398726263 |
| **Klc1** | -0.0290233174999983 | 22.48770128125 | -0.158774149700115 | 0.878455772017873 | 0.999885157113822 | -5.24674106901339 |
| **Cacng2** | 0.0370670849999968 | 25.4491829175 | 0.158403638481522 | 0.878736696803684 | 0.999885157113822 | -5.2467903082328 |
| **Tmub1** | -0.069688962499999 | 22.63144944375 | -0.157643590151148 | 0.879313030892643 | 0.999885157113822 | -5.24689095843063 |
| **Slc22a23** | 0.0368543499999952 | 24.3539646875 | 0.156475153731031 | 0.880199193786794 | 0.999885157113822 | -5.24704475435609 |
| **Psmc4** | -0.0230085950000003 | 25.370459045 | -0.155842077598525 | 0.880679407381216 | 0.999885157113822 | -5.24712760969985 |
| **Mesd** | -0.0294370200000031 | 22.68341582 | -0.155731428266112 | 0.880763344895426 | 0.999885157113822 | -5.24714205702036 |
| **Cul1** | -0.0419090650000022 | 20.8583012225 | -0.155204471676266 | 0.881163111734446 | 0.999885157113822 | -5.24721072142894 |
| **Tex2** | 0.0462128100000001 | 26.062207605 | 0.15343369939241 | 0.882506751702232 | 0.999885157113822 | -5.24743976965931 |
| **Rgs20** | 0.116774679999999 | 22.02210138 | 0.153382604927661 | 0.882545527763066 | 0.999885157113822 | -5.24744634002468 |
| **Gba1** | -0.231550009999996 | 22.33097756 | -0.152852090933606 | 0.8829481602348 | 0.999885157113822 | -5.24751443195502 |
| **Fn3krp** | 0.0276843475000064 | 24.93489523375 | 0.150769541686771 | 0.884529066199655 | 0.999885157113822 | -5.24777946767724 |
| **Gabra4** | 0.101210307499997 | 22.31072277375 | 0.149346752415142 | 0.885609461575303 | 0.999885157113822 | -5.24795846611906 |
| **Tamm41** | -0.0403373925000139 | 22.77574115125 | -0.148682415215577 | 0.886114016227038 | 0.999885157113822 | -5.248041468703 |
| **Rnh1** | -0.0354663449999961 | 28.362440225 | -0.1485401414482 | 0.886222078545636 | 0.999885157113822 | -5.24805919673588 |
| **Lypla1** | -0.0480442799999921 | 28.1183481475 | -0.148509446672198 | 0.886245392733851 | 0.999885157113822 | -5.24806301925305 |
| **Banf1** | 0.0226112700000023 | 19.027107155 | 0.148196757272656 | 0.886482902591961 | 0.999885157113822 | -5.24810191480267 |
| **Bles03** | 0.0544906825000027 | 23.69742383125 | 0.147876315021602 | 0.886726314347506 | 0.999885157113822 | -5.24814169036893 |
| **Pnpt1** | -0.0326039399999978 | 25.877360785 | -0.147715623953008 | 0.886848382165931 | 0.999885157113822 | -5.24816160433431 |
| **Dazap1** | 0.0139954400000022 | 20.0898896725 | 0.147700831405482 | 0.886859619385191 | 0.999885157113822 | -5.24816343645116 |

| | | | | | | |
|---|---|---|---|---|---|---|
| **Pnpla8** | -0.0344068325000144 | 28.22637518125 | -0.147443790460031 | 0.887054886070427 | 0.999885157113822 | -5.24819524295459 |
| **Mpi** | 0.0217320000000001 | 27.255152565 | 0.146914308003052 | 0.887457145462203 | 0.999885157113822 | -5.24826058844847 |
| **Pip4k2c** | -0.0182089225000013 | 23.05905602125 | -0.145342633154019 | 0.888651391419914 | 0.999885157113822 | -5.24845318140673 |
| **Aldh9a1** | 0.0246599349999919 | 27.518376165 | 0.144465613965599 | 0.889317935193952 | 0.999885157113822 | -5.24855975790557 |
| **Iqsec1** | 0.0328071749999985 | 24.248374465 | 0.142900863688468 | 0.890507401289093 | 0.999885157113822 | -5.24874831857632 |
| **Ppm1l** | -0.0543395475000068 | 19.00075394875 | -0.142515169510982 | 0.890800638790649 | 0.999885157113822 | -5.248794483697 |
| **Isca1** | -0.0465619675000006 | 26.32312033625 | -0.141649043888642 | 0.891459208217701 | 0.999885157113822 | -5.24889770210348 |
| **Mgst3** | -0.0155478725000044 | 30.23083755875 | -0.141524714141377 | 0.891553751489872 | 0.999885157113822 | -5.24891246753935 |
| **Gmps** | -0.0329603375000076 | 24.30065002125 | -0.140777729527344 | 0.892121816202531 | 0.999885157113822 | -5.24900090869867 |
| **Pip4p1** | -0.0434664025000053 | 25.06669646625 | -0.140174450984665 | 0.892580645620945 | 0.999885157113822 | -5.24907199628135 |
| **Spcs2** | 0.020010599999992 | 28.3317257175 | 0.139854851982834 | 0.892823737773184 | 0.999885157113822 | -5.24910953356059 |
| **Rps20** | 0.0757100000000044 | 22.6834175775 | 0.139372779463026 | 0.893190433402028 | 0.999885157113822 | -5.24916599262754 |
| **Lamtor1** | 0.0182484000000045 | 25.06076171 | 0.139185327436615 | 0.89333302918675 | 0.999885157113822 | -5.2491878942556 |
| **Rraga** | -0.075311312500002 | 21.68318356625 | -0.138787018296143 | 0.893636039254422 | 0.999885157113822 | -5.24923433496965 |
| **Clvs1** | 0.0614990225000014 | 21.80435016625 | 0.138418385813355 | 0.893916490145342 | 0.999885157113822 | -5.24927719780519 |
| **Ptgr3** | 0.0125416875000006 | 26.65738183125 | 0.137052884395037 | 0.89495548848034 | 0.999885157113822 | -5.24943498557909 |
| **Atp2c1** | 0.0541158724999917 | 23.16668582375 | 0.136127085458033 | 0.895660047618862 | 0.999885157113822 | -5.24954108078632 |
| **Vps28** | 0.038069765000003 | 23.3799818525 | 0.135986152440192 | 0.89576731055046 | 0.999885157113822 | -5.24955716887293 |
| **Cdc42bpb** | 0.046360237499993 | 30.29190596375 | 0.135787805043845 | 0.895918275064341 | 0.999885157113822 | -5.24957978301074 |
| **Adpgk** | -0.0259297250000046 | 23.32341108 | -0.135311564098408 | 0.8962807665728 | 0.999885157113822 | -5.24963394670861 |

| | | | | | | |
|---|---|---|---|---|---|---|
| **Map6d1** | 0.0567592049999988 | 25.9579278775 | 0.13528754761746 | 0.896299047461728 | 0.999885157113822 | -5.24963667313812 |
| **Mrs2** | -0.0295371700000011 | 20.6178348875 | -0.134845702169544 | 0.896635383931775 | 0.999885157113822 | -5.24968674710929 |
| **Prpf19** | 0.0284195424999965 | 25.5227828262 5 | 0.133988675546802 | 0.897287825091675 | 0.999885157113822 | -5.24978340934173 |
| **Cnr1** | 0.049298069999999 | 24.247240565 | 0.133078822296543 | 0.897980576134771 | 0.999885157113822 | -5.24988535979784 |
| **Nipsnap3b** | 0.0221625900000006 | 26.15156958 | 0.132676918784163 | 0.898286611106797 | 0.999885157113822 | -5.2499301739662 |
| **Pdk1** | 0.037519189999994 | 21.8366986275 | 0.132643042968996 | 0.898312407167309 | 0.999885157113822 | -5.24993394512858 |
| **Dmac2l** | 0.0212975374999971 | 25.73524495875 | 0.131972444634265 | 0.898823087672211 | 0.999885157113822 | -5.2500084013165 |
| **Ablim2** | 0.031735087499996 | 23.97769852625 | 0.131288492597846 | 0.899343990676956 | 0.999885157113822 | -5.25008395393979 |
| **Kctd8** | 0.0415308549999978 | 23.0798056025 | 0.130232016719675 | 0.90014871571365 | 0.999885157113822 | -5.25019989071406 |
| **Ssr3** | -0.0217784275000064 | 25.68542204125 | -0.128530105942389 | 0.901445338999404 | 0.999885157113822 | -5.25038469934863 |
| **Serpinb6** | -0.0474648300000027 | 24.694254995 | -0.126996743755652 | 0.902613829367513 | 0.999885157113822 | -5.2505491363711 |
| **Alad** | -0.0343008625000039 | 25.70923178625 | -0.125739675291213 | 0.903571965975066 | 0.999885157113822 | -5.25068248043246 |
| **Cmtm5** | 0.11127243 | 20.8872911525 | 0.125447333705522 | 0.903794813406349 | 0.999885157113822 | -5.25071330169762 |
| **Wdr48** | -0.0296546474999992 | 22.82682958125 | -0.124815811386596 | 0.904276244914882 | 0.999885157113822 | -5.25077963897199 |
| **Scg3** | 0.0233174999999974 | 24.130969985 | 0.12416878964974 | 0.9047695372082 | 0.999885157113822 | -5.25084725914047 |
| **Cdv3** | -0.0277382324999991 | 21.18128788375 | -0.123626366155987 | 0.90518311828719 | 0.999885157113822 | -5.25090367849427 |
| **Tbc1d10b** | 0.0187919374999979 | 21.74180393875 | 0.123408272598406 | 0.905349416809885 | 0.999885157113822 | -5.25092629393658 |
| **Dgkb** | 0.0272696124999996 | 24.22010543375 | 0.122913231802766 | 0.905726909371102 | 0.999885157113822 | -5.25097748036214 |
| **Pef1** | -0.0236309699999993 | 28.2591725575 | -0.122074873760463 | 0.906366257845687 | 0.999885157113822 | -5.25106369861484 |
| **H1-5** | -0.0243340224999997 | 21.47995070625 | -0.121798457541902 | 0.906577074778511 | 0.999885157113822 | -5.25109199712129 |

| | | | | | | |
|---|---|---|---|---|---|---|
| **Grik3** | 0.0159329649999975 | 25.0795820075 | 0.121785726000444 | 0.906586785057682 | 0.999885157113822 | -5.25109329899423 |
| **Mpp1** | 0.0447085824999931 | 24.91473422625 | 0.118891075708897 | 0.90879496335062 | 0.999885157113822 | -5.2513857796034 |
| **Scfd1** | 0.0122245850000056 | 26.0254512425 | 0.118723459592401 | 0.908922855822453 | 0.999885157113822 | -5.25140250148795 |
| **Tmcc1** | -0.045633610000003 | 25.2849451725 | -0.11855013936047 | 0.909055103644546 | 0.999885157113822 | -5.25141976775233 |
| **Trappc2** | 0.0166491825000072 | 25.87478929125 | 0.118178961855582 | 0.909338332142978 | 0.999885157113822 | -5.25145666026256 |
| **Arpc5** | 0.0185378225000044 | 25.26718974875 | 0.117728653411396 | 0.909681960912508 | 0.999885157113822 | -5.25150126333626 |
| **Hapln4** | -0.0244734724999951 | 29.04081903625 | -0.117509923207893 | 0.909848880680707 | 0.999885157113822 | -5.25152286744703 |
| **Cops5** | 0.0155052849999961 | 33.939020855 | 0.117481993953784 | 0.909870194710156 | 0.999885157113822 | -5.25152562315821 |
| **Ado** | 0.0178076049999909 | 27.7157208375 | 0.117002909512972 | 0.910235817347384 | 0.999885157113822 | -5.2515727917797 |
| **Ptgds** | -0.0266661125000063 | 23.46129947625 | -0.116893686931876 | 0.910319175978211 | 0.999885157113822 | -5.25158351852895 |
| **Sf1** | -0.0379581699999996 | 22.648856925 | -0.116294715270819 | 0.91077633254293 | 0.999885157113822 | -5.25164216632225 |
| **Scn2b** | -0.0285291300000026 | 21.1498487425 | -0.116052608757443 | 0.910961127236467 | 0.999885157113822 | -5.25166578689934 |
| **Cpne2** | 0.0455535825000055 | 23.17075752625 | 0.114936256245464 | 0.911813291791186 | 0.999885157113822 | -5.25177406769107 |
| **Scoc** | 0.0146830824999959 | 30.46587854625 | 0.113802385037488 | 0.912678957323475 | 0.999885157113822 | -5.25188298155063 |
| **Tbl2** | -0.0270502450000052 | 23.9903857725 | -0.113493382196965 | 0.91291489091889 | 0.999885157113822 | -5.25191247645529 |
| **Eif3j2** | -0.0374580125000001 | 22.11004856125 | -0.112422903349389 | 0.913732308994956 | 0.999885157113822 | -5.25201403848991 |
| **G6pdx** | -0.0471005350000056 | 24.97951474 | -0.110988105798928 | 0.914828097437803 | 0.999885157113822 | -5.25214866284738 |
| **Rpl31** | -0.117164295000002 | 22.6739571275 | -0.110162120619082 | 0.915459012416163 | 0.999885157113822 | -5.25222538281084 |
| **Fbxo41** | 0.0309224999999955 | 23.3231315375 | 0.108991388398459 | 0.916353369153738 | 0.999885157113822 | -5.25233314650256 |
| **Lhfpl4** | -0.771866652499998 | 16.09737974875 | -0.10825329548119 | 0.916917287264742 | 0.999885157113822 | -5.25240049741346 |

| | | | | | | |
|---|---|---|---|---|---|---|
| **Cox20** | -0.0520715149999944 | 24.88080072 | -0.108143702475059 | 0.917001022974137 | 0.999885157113822 | -5.25241045891426 |
| **Lamtor5** | 0.0143417600000042 | 25.42299037 | 0.104502456910967 | 0.919783794775018 | 0.999885157113822 | -5.25273571872608 |
| **Kctd16** | 0.687808000000002 | 14.8680417025 | 0.104260515116849 | 0.919968738889495 | 0.999885157113822 | -5.25275693751221 |
| **Sri** | -0.0391705125000001 | 24.18622510625 | -0.102882326419454 | 0.921022349542832 | 0.999885157113822 | -5.25287687317397 |
| **Stx5** | -0.076087175000005 | 20.923411775 | -0.102807756029255 | 0.921079362715335 | 0.999885157113822 | -5.2528833172644 |
| **Map1lc3b** | -0.0203868949999979 | 22.1912598675 | -0.102162419335001 | 0.92157277905055 | 0.999885157113822 | -5.25293889039293 |
| **Rbm39** | 0.0581488950000093 | 22.903526985 | 0.101926844694064 | 0.921752905758025 | 0.999885157113822 | -5.25295909004027 |
| **Lman2l** | 0.0317840300000043 | 19.55900523 | 0.101870107825682 | 0.921796289034382 | 0.999885157113822 | -5.25296394807419 |
| **Ipo5** | 0.734569494999999 | 16.679129655 | 0.0993457859427614 | 0.923726774691142 | 0.999885157113822 | -5.25317736333416 |
| **Arpp21** | -0.0600596375000038 | 24.32290008125 | -0.0984798405292537 | 0.924389138492531 | 0.999885157113822 | -5.25324934450833 |
| **Abcb10** | -0.0143079950000029 | 22.890438825 | -0.0981682899647983 | 0.924627460149307 | 0.999885157113822 | -5.253275088365 |
| **Copg1** | -0.0299569749999975 | 26.40005525 | -0.0977004656715797 | 0.924985339594528 | 0.999885157113822 | -5.25331359267525 |
| **Gart** | 0.0116995125000017 | 21.30994782875 | 0.0976876560786246 | 0.924995139028969 | 0.999885157113822 | -5.25331464439178 |
| **Cox15** | -0.692708957500002 | 16.13829636875 | -0.0969025166405773 | 0.925595803315414 | 0.999885157113822 | -5.25337884500463 |
| **Pofut2** | 0.0256051850000034 | 24.16205283 | 0.0966899230171816 | 0.925758455292139 | 0.999885157113822 | -5.25339613989962 |
| **Trnt1** | -0.0127658575000069 | 23.57427341875 | -0.0962526092528192 | 0.926093049081678 | 0.999885157113822 | -5.2534315971855 |
| **Usp12** | -0.0329928849999988 | 25.3434584 | -0.0943774372955627 | 0.927527946699838 | 0.999885157113822 | -5.25358181956961 |
| **Mia3** | -0.5988999775 | 14.40844793375 | -0.0938087459048272 | 0.927963171967362 | 0.999885157113822 | -5.2536267960355 |
| **Dad1** | -0.635828590000001 | 15.3417600025 | -0.0935337900613133 | 0.928173608000765 | 0.999885157113822 | -5.25364844447943 |
| **Sec11c** | -0.0128642899999996 | 26.19622567 | -0.0931194292382301 | 0.92849074861966 | 0.999885157113822 | -5.25368094922028 |

| | | | | | | |
|---|---|---|---|---|---|---|
| **Mtres1** | 0.0142967000000027 | 26.4260234 | 0.091894977999484 | 0.929427992313022 | 0.999885157113822 | -5.25377616125413 |
| **Tmem205** | -0.59398264 | 14.8867724325 | -0.0900359132074521 | 0.930851224908948 | 0.999885157113822 | -5.25391831835653 |
| **Rpl23a** | 0.00808945500001101 | 27.714641325 | 0.089746849542129 | 0.931072546228946 | 0.999885157113822 | -5.25394016199053 |
| **Pls1** | 0.00963353749999385 | 24.70637121625 | 0.0893915728735711 | 0.931344572479895 | 0.999885157113822 | -5.25396691320078 |
| **Ostc** | 0.0546132774999997 | 23.05316660125 | 0.0885077391853639 | 0.932021344164703 | 0.999885157113822 | -5.25403300411134 |
| **Akap7** | 0.0135133199999942 | 27.68453704 | 0.0873508247637943 | 0.932907311583935 | 0.999885157113822 | -5.25411852579245 |
| **Islr2** | -0.0153225524999918 | 24.45767265625 | -0.0870330359536202 | 0.933150692917499 | 0.999885157113822 | -5.25414182101205 |
| **Tor1aip1** | -0.0108177949999987 | 26.2181107325 | -0.086795632865931 | 0.933332515178215 | 0.999885157113822 | -5.25415916837995 |
| **Dner** | 0.00997160500000049 | 28.0269475425 | 0.086696527475218 | 0.933408419278772 | 0.999885157113822 | -5.25416639616643 |
| **Lamtor2** | -0.0340719225000043 | 24.38885156875 | -0.0866210923180768 | 0.933466195019829 | 0.999885157113822 | -5.25417189215623 |
| **Mrpl49** | -0.0145244800000022 | 24.9778812925 | -0.0857456143427267 | 0.934136754612027 | 0.999885157113822 | -5.25423532798665 |
| **Mmab** | -0.0143220774999975 | 29.85748579125 | -0.0844205161137497 | 0.935151803804774 | 0.999885157113822 | -5.25433012014427 |
| **Cystm1** | -0.0335049400000003 | 22.3363425175 | -0.0843068091681449 | 0.935238911470177 | 0.999885157113822 | -5.25433818568526 |
| **Gbf1** | 0.00935468250000326 | 24.14241107375 | 0.0842430250073612 | 0.93528777512529 | 0.999885157113822 | -5.25434270532138 |
| **Vac14** | 0.645913737500003 | 17.42989256875 | 0.0833397985453185 | 0.935979749414066 | 0.999885157113822 | -5.25440634016697 |
| **Rap1gap** | 0.0215290650000064 | 24.7140961975 | 0.0819972876230909 | 0.93700837569763 | 0.999885157113822 | -5.25449965958338 |
| **Tmem222** | -0.0126057699999933 | 26.45345229 | -0.0816419025674506 | 0.937280692055278 | 0.999885157113822 | -5.25452410974287 |
| **Bckdha** | 0.0239351199999973 | 28.36381208 | 0.0811015529122406 | 0.937694756203693 | 0.999885157113822 | -5.25456108230283 |
| **Bcap29** | 0.0396853375000035 | 26.34298622625 | 0.0800586454938809 | 0.938493983651424 | 0.999885157113822 | -5.25463174903261 |
| **Hdac11** | -0.04733169 | 21.81527676 | -0.079985411523944 | 0.938550109062457 | 0.999885157113822 | -5.2546366770341 |

| | | | | | | |
|---|---|---|---|---|---|---|
| **Hmga1** | 0.0249368399999987 | 21.663794775 | 0.078458317273748 | 0.939720536399839 | 0.999885157113822 | -5.25473841190484 |
| **Dcakd** | 0.00992260499999986 | 19.7091950475 | 0.0773433572984076 | 0.940575188891495 | 0.999885157113822 | -5.25481145465534 |
| **Dpm1** | 0.0104498174999996 | 28.12096836875 | 0.0768117381582054 | 0.940982721781097 | 0.999885157113822 | -5.25484591459666 |
| **Sec11a** | 0.488246297500005 | 15.06231254875 | 0.0729589904436156 | 0.9439367539839 | 0.999885157113822 | -5.25508856451883 |
| **Pafah1b3** | 0.0166099024999937 | 23.76472103375 | 0.0729019087240495 | 0.94398052775055 | 0.999885157113822 | -5.25509206592269 |
| **Tra2b** | 0.00670508500001077 | 27.734245 | 0.0728266692601869 | 0.944038226309092 | 0.999885157113822 | -5.25509667694762 |
| **Cryab** | 0.570311867500003 | 10.36085999125 | 0.0716722834467579 | 0.944923530312405 | 0.999885157113822 | -5.2551668273083 |
| **Psmd11** | 0.0122227674999991 | 25.99363579125 | 0.0710600430352411 | 0.945393094568052 | 0.999885157113822 | -5.25520357828004 |
| **Hook3** | 0.00994728500000264 | 30.4574568875 | 0.0695682661995975 | 0.946537326209431 | 0.999885157113822 | -5.25529180763451 |
| **Abhd17c** | -0.0411864650000027 | 21.1395568925 | -0.068953515894224 | 0.94700889545376 | 0.999885157113822 | -5.25532762265616 |
| **Lrrc4b** | -0.0177259700000043 | 22.5108844975 | -0.0678058459696176 | 0.947889323539805 | 0.999885157113822 | -5.25539363607661 |
| **Dnaja4** | 0.0129050325000009 | 29.03995721625 | 0.0671659013467354 | 0.948380287399913 | 0.999885157113822 | -5.25542996506109 |
| **Lgmn** | 0.0101904874999974 | 25.25730941125 | 0.0668597099993649 | 0.94861520517944 | 0.999885157113822 | -5.25544722558002 |
| **Dpp7** | 0.0076246300000129 | 26.9202525525 | 0.0668485600179724 | 0.948623759832569 | 0.999885157113822 | -5.25544785263738 |
| **Vkorc1** | 0.0296060850000082 | 23.9149131075 | 0.0663065349222858 | 0.949039629108515 | 0.999885157113822 | -5.25547820936908 |
| **Slc39a11** | 0.0109339550000023 | 26.47139143 | 0.0662710884689503 | 0.949066826032253 | 0.999885157113822 | -5.25548018599256 |
| **Mrpl23** | 0.0415987849999979 | 19.51980129 | 0.0651603432789119 | 0.949919101997572 | 0.999885157113822 | -5.25554159056289 |
| **Mrpl43** | -0.015946894999999 | 23.48193721 | -0.0647297018268775 | 0.950249552728604 | 0.999885157113822 | -5.25556511866348 |
| **Parva** | 0.441284382500003 | 15.46037085375 | 0.0644494786517576 | 0.950464586364057 | 0.999885157113822 | -5.2555803450014 |
| **Gpc4** | -0.0212759274999961 | 23.89410589375 | -0.0634913223477925 | 0.951199876103021 | 0.999885157113822 | -5.25563190949146 |

| | | | | | | |
|---|---|---|---|---|---|---|
| **Anp32b** | 0.0156767950000045 | 28.3749669 | 0.0621123763916565 | 0.95225816988136 | 0.999885157113822 | -5.25570476575887 |
| **Efnb2** | -0.0134253925000003 | 26.90593092625 | -0.0608989700562879 | 0.953189503694468 | 0.999885157113822 | -5.25576755448817 |
| **Pm20d1** | -0.413262739999992 | 15.4571823075 | -0.0603373753626082 | 0.953620574996679 | 0.999885157113822 | -5.25579619592967 |
| **Suclg2** | -0.0364392575000068 | 23.53020330625 | -0.0589664496432503 | 0.954672946151696 | 0.999885157113822 | -5.25586500040463 |
| **Farsb** | -0.474689860000002 | 10.648500295 | -0.0580579502581288 | 0.955370396339003 | 0.999885157113822 | -5.25590972643465 |
| **Clcn4** | 0.385743527500003 | 15.07133997375 | 0.0577249216750899 | 0.955626071224166 | 0.999885157113822 | -5.25592594793305 |
| **Cd44** | -0.0307751000000067 | 21.3409752725 | -0.05771862467701 | 0.955630905651158 | 0.999885157113822 | -5.25592625375591 |
| **Ptbp2** | -0.0070254474999949 | 25.79381345375 | -0.0573876848242671 | 0.95588498265004 | 0.999885157113822 | -5.25594227943691 |
| **C2orf88(AKAP)** | -0.0172463024999985 | 24.89222832875 | -0.0573718975131228 | 0.955897103397771 | 0.999885157113822 | -5.25594304163356 |
| **Sec61a1** | 0.0179461049999929 | 25.13093676 | 0.0572779498971888 | 0.955969232175728 | 0.999885157113822 | -5.25594757300495 |
| **Tbrg4** | 0.00847851750000572 | 29.16237099625 | 0.0566393258592094 | 0.956459550906802 | 0.999885157113822 | -5.25597817915046 |
| **Lrtm2** | -0.396996507499997 | 9.19489026125 | -0.0562860483421589 | 0.956730796837398 | 0.999885157113822 | -5.25599496277954 |
| **Spg7** | -0.0258391150000108 | 25.1141103225 | -0.056099686125357 | 0.956873887952666 | 0.999885157113822 | -5.25600377427959 |
| **Rilp** | -0.0064609875000059 | 30.14360238875 | -0.0550153228548685 | 0.957706508744098 | 0.999885157113822 | -5.25605446564946 |
| **Slc22a17** | -0.00887390749999994 | 30.74203203875 | -0.0543939980829558 | 0.958183614381467 | 0.999885157113822 | -5.25608306572007 |
| **Atp5mpl** | -0.0211313424999986 | 24.59033500125 | -0.0539851544354457 | 0.958497569162628 | 0.999885157113822 | -5.25610170811282 |
| **Uap1l1** | 0.401363360000005 | 17.0207505725 | 0.0532318068748521 | 0.959076092415726 | 0.999885157113822 | -5.25613569118048 |
| **Penk** | 0.0106196974999975 | 25.27927795375 | 0.0517661657597709 | 0.960201687916667 | 0.999885157113822 | -5.2562004384903 |
| **Gipc1** | -0.0157148925000001 | 21.73675646875 | -0.0501863818165431 | 0.961415053365195 | 0.999885157113822 | -5.25626820602271 |
| **Atp8a2** | 0.332749977500001 | 15.29599208625 | 0.049111704971125 | 0.962240530594994 | 0.999885157113822 | -5.25631310696767 |

| | | | | | | |
|---|---|---|---|---|---|---|
| **Trappc4** | -0.0204309024999958 | 25.36521157125 | -0.0488076393850463 | 0.96247409756261 | 0.999885157113822 | -5.25632563485927 |
| **Arl6ip1** | 0.342795527500002 | 15.88335153125 | 0.0485313020253353 | 0.962686368633319 | 0.999885157113822 | -5.25633695288643 |
| **Gfm1** | -0.00766296249999954 | 25.16740509625 | -0.0484425329406322 | 0.962754558111318 | 0.999885157113822 | -5.2563405750016 |
| **Cntn4** | 0.319158975000002 | 14.8966804225 | 0.0482907824766445 | 0.962871128582312 | 0.999885157113822 | -5.25634675165148 |
| **Trappc3** | -0.0142215225000015 | 21.30669921375 | -0.0477418202445736 | 0.963292834205164 | 0.999885157113822 | -5.25636893416583 |
| **Mobp** | -0.013790692500006 | 23.52052335375 | -0.0477211529563982 | 0.963308710794672 | 0.999885157113822 | -5.25636976434201 |
| **Bcl2l1** | 0.0166874700000008 | 23.916170185 | 0.0469677198239586 | 0.963887509546466 | 0.999885157113822 | -5.25639978349512 |
| **Exoc7** | 0.00632022750000516 | 29.56093126625 | 0.0463111989486864 | 0.964391877994373 | 0.999885157113822 | -5.25642555217512 |
| **Rpl36a** | -0.0127421375000019 | 24.72677865875 | -0.0462988835364529 | 0.964401339408931 | 0.999885157113822 | -5.25642603209625 |
| **Sumo3** | -0.0269751674999981 | 23.66099350375 | -0.0462846388672006 | 0.964412282998434 | 0.999885157113822 | -5.25642658703981 |
| **Gfer** | -0.311253622500004 | 15.36348094625 | -0.0457209013439709 | 0.964845385808726 | 0.999885157113822 | -5.25644841211852 |
| **Bckdk** | 0.292367399999998 | 14.4713884275 | 0.0456326195323757 | 0.964913211268009 | 0.999885157113822 | -5.25645180574346 |
| **Eif4g1** | -0.0337495899999993 | 22.7800315325 | -0.0454670078513297 | 0.965040448834772 | 0.999885157113822 | -5.25645815431207 |
| **Extl2** | -0.0243354525000008 | 23.71971179375 | -0.045426936142386 | 0.965071235637194 | 0.999885157113822 | -5.25645968695794 |
| **Tmsb4x** | 0.305722275000001 | 15.3339243825 | 0.0450230337371833 | 0.965381554494831 | 0.999885157113822 | -5.25647505984711 |
| **Hexa** | -0.0098939775000062 | 24.89847878625 | -0.044603976751587 | 0.965703523479067 | 0.999885157113822 | -5.25649086452386 |
| **Anxa3** | -0.349243757499998 | 10.67437725625 | -0.0426639864177761 | 0.967194141681518 | 0.999885157113822 | -5.25656210656972 |
| **Srsf10** | 0.253695439999998 | 13.745197345 | 0.041662916723199 | 0.967963383064527 | 0.999885157113822 | -5.25659763063745 |
| **Tmem94** | -0.00733653249999833 | 25.33473237125 | -0.0414900869392038 | 0.968096192573627 | 0.999885157113822 | -5.2566036783723 |
| **Ptdss1** | 0.00774090999999899 | 23.1255048 | 0.0413267484174739 | 0.968221709610507 | 0.999885157113822 | -5.25660937089266 |

| | | | | | | |
|---|---|---|---|---|---|---|
| **Macroh2a1** | -0.00406521999999399 | 20.3078689875 | -0.0410881301319439 | 0.968405076919729 | 0.999885157113822 | -5.25661764665528 |
| **Cfap418** | 0.00879068500000457 | 28.6471155625 | 0.0372574324100926 | 0.971349068032226 | 0.999885157113822 | -5.25674394681026 |
| **Elavl1** | 0.00814898999999158 | 28.1161880075 | 0.0370938084725299 | 0.971474828377834 | 0.999885157113822 | -5.25674906668423 |
| **Cpne1** | 0.0132563949999991 | 24.49059655 | 0.0370729180043116 | 0.971490884725488 | 0.999885157113822 | -5.25674971873595 |
| **Scn3b** | 0.00302607499999752 | 30.27159407 | 0.0369177208233811 | 0.971610169232746 | 0.999885157113822 | -5.2567545513921 |
| **Plpp6** | -0.0111736425000011 | 29.51442131625 | -0.0366863237392994 | 0.971788022417363 | 0.999885157113822 | -5.25676171919415 |
| **Trim3** | 0.00883557999998885 | 30.347900765 | 0.0366043176667104 | 0.971851053191953 | 0.999885157113822 | -5.25676424861974 |
| **Mrps5** | -0.00367550500000391 | 28.8164456025 | -0.0361553851360724 | 0.972196111533509 | 0.999885157113822 | -5.256777995398 |
| **Crot** | 0.00510887500000479 | 26.6144759775 | 0.0361116736949007 | 0.972229709348284 | 0.999885157113822 | -5.25677932482926 |
| **Atxn10** | -0.249456942499998 | 15.77388370875 | -0.0357128124229585 | 0.972536287881825 | 0.999885157113822 | -5.25679138146288 |
| **Fbxl20** | -0.00652646000001056 | 28.452582165 | -0.0356030567598251 | 0.972620650751845 | 0.999885157113822 | -5.25679467563477 |
| **Nedd8** | -0.243878085000002 | 15.476000945 | -0.0355916984081693 | 0.972629381285544 | 0.999885157113822 | -5.25679501596208 |
| **Pam** | -0.0125331675000062 | 23.17394120375 | -0.035575540483775 | 0.972641800991054 | 0.999885157113822 | -5.2567954999107 |
| **Myo18a** | -0.255661940000001 | 9.52800889 | -0.0349920982794568 | 0.97309026629747 | 0.999885157113822 | -5.25681282752227 |
| **Plcxd3** | -0.264791340000002 | 10.630123245 | -0.0324841260741768 | 0.975018147072404 | 0.999885157113822 | -5.25688405003108 |
| **Mlf2** | 0.237491087499999 | 9.69004579875 | 0.0319568339685842 | 0.975423500429486 | 0.999885157113822 | -5.25689835099312 |
| **Mapk8ip3** | -0.209601579999999 | 14.8565596475 | -0.0318577123104326 | 0.975499700622345 | 0.999885157113822 | -5.25690101320033 |
| **Mydgf** | 0.0110609474999999 | 23.82158697375 | 0.0318204627257518 | 0.97552833646909 | 0.999885157113822 | -5.25690201151167 |
| **Tesc** | 0.219090609999999 | 15.5535493725 | 0.031767072579388 | 0.975569380539516 | 0.999885157113822 | -5.25690344036384 |
| **Hapln2** | -0.2654756375 | 10.99260025375 | -0.0314886918562489 | 0.975783389072358 | 0.999885157113822 | -5.25691085166242 |

| | | | | | | |
|---|---|---|---|---|---|---|
| **Myorg** | -0.195717664999997 | 14.3372685625 | -0.0308347932869343 | 0.976286089899083 | 0.999885157113822 | -5.25692800385006 |
| **Tcerg1** | -0.00584050750000387 | 23.8295845362 5 | -0.0302153841020867 | 0.976762286876066 | 0.999885157113822 | -5.25694391952811 |
| **Has1** | 0.258545529999996 | 11.21186332 | 0.0300726793969942 | 0.976871998602715 | 0.999885157113822 | -5.25694754055772 |
| **Nf2** | -0.0124546774999956 | 19.6867376437 5 | -0.0298935995925738 | 0.977009676358411 | 0.999885157113822 | -5.25695206033404 |
| **Gprin3** | 0.00837740999999426 | 26.813755815 | 0.0294338320617994 | 0.977363152580824 | 0.999885157113822 | -5.25696354077611 |
| **Alg5** | 0.21894715 | 9.84751976 | 0.0289967959144209 | 0.977699157659284 | 0.999885157113822 | -5.25697428870569 |
| **Adss2** | 0.00483289750000893 | 25.5352504587 5 | 0.0281756331290511 | 0.97833050271962 | 0.999885157113822 | -5.25699404865064 |
| **Slc22a4** | 0.0183349525000089 | 22.1593280412 5 | 0.0281312589603433 | 0.978364619959321 | 0.999885157113822 | -5.2569951002822 |
| **Reep5** | -0.182723240000003 | 14.9877572275 | -0.0275360636287518 | 0.978822242727657 | 0.999885157113822 | -5.25700904575084 |
| **Mthfd1** | 0.176387947500004 | 14.7278221362 5 | 0.0270129852416513 | 0.979224424662192 | 0.999885157113822 | -5.2570210553832 |
| **Fam210a** | -0.0130347975000014 | 21.7103552962 5 | -0.0269481387271574 | 0.979274283987559 | 0.999885157113822 | -5.25702252818625 |
| **Plgrkt** | -0.00790258249999098 | 21.8691490912 5 | -0.0264496814337903 | 0.979657542204838 | 0.999885157113822 | -5.2570337310679 |
| **Get4** | 0.1987591825 | 9.92198485625 | 0.0261254837078001 | 0.979906817313969 | 0.999885157113822 | -5.25704090522096 |
| **Ccdc110** | -0.210140614999998 | 10.745279265 | -0.025471894413037 | 0.980409368399995 | 0.999885157113822 | -5.25705509954264 |
| **Selenbp1** | 0.171166634999999 | 16.529751195 | 0.0233825565892395 | 0.982015943474283 | 0.999885157113822 | -5.25709806311334 |
| **Rapgef4** | -0.0042514649999994 | 23.505678005 | -0.0232855368273503 | 0.982090548082759 | 0.999885157113822 | -5.25709996889057 |
| **Pnkd** | -0.184487369999999 | 10.3738394525 | -0.0231895102475077 | 0.982164389161317 | 0.999885157113822 | -5.25710184735741 |
| **Pacsin2** | -0.170700275000002 | 9.594082885 | -0.0231882928398765 | 0.982165325306383 | 0.999885157113822 | -5.25710187112245 |
| **Ttc19** | -0.1569125775 | 15.3999651112 5 | -0.0230151431176282 | 0.98229847186216 | 0.999885157113822 | -5.25710523847676 |
| **Arfgef1** | -0.00305166500000098 | 29.607704315 | -0.0222725296757835 | 0.982869524509158 | 0.999885157113822 | -5.25711939436852 |

| | | | | | | |
|---|---|---|---|---|---|---|
| **Abcg5** | 0.00460677750000826 | 25.37496154625 | 0.0221040774069391 | 0.982999061947536 | 0.999885157113822 | -5.2571225408635 |
| **Phactr4** | -0.136017522500001 | 14.30205858625 | -0.0214765105390005 | 0.983481656925904 | 0.999885157113822 | -5.25713405285825 |
| **U2af2** | -0.0101004825000004 | 20.69176641375 | -0.0214454375752444 | 0.983505552035418 | 0.999885157113822 | -5.25713461424318 |
| **Gcn1** | -0.134898839999993 | 14.241118605 | -0.0213867306846932 | 0.983550697684795 | 0.999885157113822 | -5.25713567266295 |
| **Sco2** | -0.00469604750000485 | 27.09235714875 | -0.0211106923484386 | 0.983762972271528 | 0.999885157113822 | -5.25714061043887 |
| **Dnajb14** | -0.00372874000001033 | 23.0219952375 | -0.0200048229813194 | 0.984613404286664 | 0.999885157113822 | -5.25715974908695 |
| **Amigo1** | -0.128243179999998 | 14.8616597075 | -0.0194896746949114 | 0.985009569363041 | 0.999885157113822 | -5.25716831301881 |
| **Abhd10** | -0.0139054524999978 | 21.20352445625 | -0.0194074802309629 | 0.985072779890856 | 0.999885157113822 | -5.25716965877289 |
| **Akap10** | -0.00477119750000554 | 22.24153099875 | -0.0193968531596199 | 0.985080952503113 | 0.999885157113822 | -5.25716983235272 |
| **GOLGA7B** | 0.150304632500001 | 10.15819899375 | 0.0192693454731769 | 0.985179010796439 | 0.999885157113822 | -5.25717190761745 |
| **Gfra1** | 0.00428202999999527 | 23.6972143275 | 0.0190559231488655 | 0.98534314135348 | 0.999885157113822 | -5.25717535057165 |
| **Itga3** | 0.00528161749999612 | 29.94752230125 | 0.0189902535975943 | 0.985393644097381 | 0.999885157113822 | -5.25717640224699 |
| **Os9** | 0.00260999750000224 | 22.02860799875 | 0.0188665129545126 | 0.98548880623514 | 0.999885157113822 | -5.2571783740467 |
| **Psen1** | -0.152330580000005 | 10.6843459025 | -0.018593138506637 | 0.985699044421192 | 0.999885157113822 | -5.25718268456176 |
| **Kif1b** | -0.00339178000000118 | 22.5632033725 | -0.0182370467740513 | 0.985972898039239 | 0.999885157113822 | -5.25718820500888 |
| **Ddi2** | 0.133109532500001 | 9.74036410125 | 0.0178232492019824 | 0.986291133098683 | 0.999885157113822 | -5.25719448598029 |
| **Smc4** | 0.00428855749999713 | 24.80152859875 | 0.0178045863259163 | 0.986305486029489 | 0.999885157113822 | -5.25719476586439 |
| **Vps29** | 0.00333685500000414 | 28.4591411 | 0.0175923857857677 | 0.986468682034872 | 0.999885157113822 | -5.25719792758253 |
| **Abhd11** | 0.010003679999997 | 22.677419665 | 0.0174935699200371 | 0.986544678096201 | 0.999885157113822 | -5.25719938697125 |
| **Rabac1** | -0.1343612975 | 17.79455204625 | -0.0170533681886639 | 0.986883224717671 | 0.999885157113822 | -5.25720578834313 |

| | | | | | | |
|---|---|---|---|---|---|---|
| **Rpap1** | -0.00649177749999907 | 23.47641767875 | -0.0166483810890312 | 0.987194691344373 | 0.999885157113822 | -5.2572115335616 |
| **Dchs1** | 0.00382119249999846 | 26.89922331375 | 0.0165035066939653 | 0.98730611162734 | 0.999885157113822 | -5.25721355524693 |
| **Tgfbrap1** | 0.135563117499998 | 10.96366745875 | 0.0161263455094775 | 0.987596180914778 | 0.999885157113822 | -5.25721873556528 |
| **Scn4b** | 0.110968877499998 | 15.72626792875 | 0.0159344686850916 | 0.987743751423153 | 0.999885157113822 | -5.25722132504293 |
| **Impdh2** | 0.1050801425 | 15.19077280875 | 0.0156182117785788 | 0.987986982510363 | 0.999885157113822 | -5.25722552545625 |
| **Lamtor3** | -0.0959963099999968 | 13.9914258325 | -0.0154970146288412 | 0.988080194800338 | 0.999885157113822 | -5.25722711283907 |
| **Nipal3** | 0.0101602925000037 | 21.86874676375 | 0.015443568289719 | 0.988121300250544 | 0.999885157113822 | -5.2572278089252 |
| **Arhgef2** | 0.00134764500000273 | 26.50177721 | 0.0152865945272911 | 0.988242028616201 | 0.999885157113822 | -5.25722983945194 |
| **Psph** | 0.0118884175000034 | 21.64357436875 | 0.0152731032321614 | 0.988252404773515 | 0.999885157113822 | -5.25723001299997 |
| **Cnpy3** | 0.0940546324999971 | 14.07977473125 | 0.0150800845049246 | 0.98840085577599 | 0.999885157113822 | -5.25723247915859 |
| **Dnaaf3** | 0.107308175 | 9.5960097375 | 0.0145834966687589 | 0.988782784513319 | 0.999885157113822 | -5.25723867981604 |
| **Nedd4l** | 0.119467567500001 | 10.78126357875 | 0.0144518284887816 | 0.98888405185111 | 0.999885157113822 | -5.25724028907656 |
| **Pank3** | -0.0987513500000023 | 9.222670305 | -0.0139635080646608 | 0.989259625974985 | 0.999885157113822 | -5.25724612994252 |
| **Parn** | -0.00246570750000075 | 25.79889544875 | -0.0134863988786287 | 0.989626580215113 | 0.999885157113822 | -5.25725164283205 |
| **Rps29** | 0.0887793474999992 | 15.09640331625 | 0.0132820492630365 | 0.989783750435217 | 0.999885157113822 | -5.25725394543155 |
| **Irgm1** | 0.0841743600000004 | 14.38744913 | 0.0132085533885566 | 0.989840278005682 | 0.999885157113822 | -5.25725476498349 |
| **Rgs18** | 0.0924909525000004 | 9.23356036875 | 0.0130643871994901 | 0.989951160118191 | 0.999885157113822 | -5.2572563593715 |
| **Aspm** | -0.0952100775000009 | 9.71220939875 | -0.0127852959401746 | 0.990165817427297 | 0.999885157113822 | -5.25725939622485 |
| **Mrpl38** | -0.0910714624999969 | 16.12170671625 | -0.0127606882958499 | 0.99018474393387 | 0.999885157113822 | -5.2572596608404 |
| **Gnptab** | -0.00202139500000342 | 24.25328618 | -0.0127301805194096 | 0.990208208425496 | 0.999885157113822 | -5.25725998819461 |

| | | | | | | |
|---|---|---|---|---|---|---|
| **Ddr1** | 0.0776778350000029 | 14.1739635 | 0.0123671191417827 | 0.990487451160053 | 0.999885157113822 | -5.25726382376699 |
| **Wdr76** | -0.0867618625000013 | 9.55315490625 | -0.0118450515834368 | 0.990888993413852 | 0.999885157113822 | -5.25726914465903 |
| **Wipf2** | 0.00246576500000373 | 22.6805106925 | 0.0111814560029495 | 0.991399394411287 | 0.999885157113822 | -5.25727557681697 |
| **Eml5** | 0.0914100424999997 | 10.7756127462 5 | 0.0110640583313603 | 0.991489690663093 | 0.999885157113822 | -5.25727667614626 |
| **Tp53bp1** | -0.0854087299999957 | 10.27478343 | -0.0108410555569439 | 0.99166121329852 | 0.999885157113822 | -5.25727873242483 |
| **Fam83d** | -0.0791389175000017 | 10.0575244062 5 | -0.0102625757745734 | 0.992106153349494 | 0.999885157113822 | -5.25728387134929 |
| **Acacb** | 0.001361904999996 | 28.8723939625 | 0.00972386473156099 | 0.992520507842527 | 0.999885157113822 | -5.25728840365419 |
| **Plvap** | -0.0678691675000032 | 9.35606277625 | -0.00945632109212741 | 0.992726292397277 | 0.999885157113822 | -5.2572905637642 |
| **Emsy** | -0.0027154524999915 | 26.4805625962 5 | -0.00921724824108368 | 0.992910178792808 | 0.999885157113822 | -5.25729244302222 |
| **Tiprl** | -0.0596457899999905 | 15.774905645 | -0.00853699918025885 | 0.993433404853255 | 0.999885157113822 | -5.25729752695386 |
| **Alpk3** | 0.00363816000000128 | 23.523512425 | 0.00818890673590574 | 0.993701147916266 | 0.999885157113822 | -5.257299977791 |
| **Dpp4** | -0.00416933499999672 | 22.1398942575 | -0.0081464887464024 | 0.993733774719022 | 0.999885157113822 | -5.25730026947339 |
| **Tktl1** | 0.0559694625000056 | 15.8397820362 5 | 0.00798070569472058 | 0.993861290808907 | 0.999885157113822 | -5.25730139493207 |
| **Vps4b** | 0.00176646500000288 | 26.455860285 | 0.00782827040610499 | 0.993978540311148 | 0.999885157113822 | -5.25730240935737 |
| **Pkhd1l1** | 0.0016368249999914 | 28.671351245 | 0.00711483372379686 | 0.994527300470519 | 0.999885157113822 | -5.25730689709644 |
| **Icmt** | -0.00193537249999309 | 23.1950965512 5 | -0.00708745712839572 | 0.994548358024886 | 0.999885157113822 | -5.25730706076664 |
| **Fat3** | 0.00169226500000264 | 23.2445378675 | 0.00558844949279105 | 0.995701372729448 | 0.999885157113822 | -5.25731505939365 |
| **Agpat4** | -0.0384960374999981 | 15.8778559187 5 | -0.00546648559981402 | 0.995795186095862 | 0.999885157113822 | -5.25731562696552 |
| **Cirbp** | 0.0361763775000075 | 15.6420573012 5 | 0.00520599829943316 | 0.995995550474299 | 0.999885157113822 | -5.2573167972343 |
| **Atp6ap2** | 0.00162696250000494 | 21.4142791187 5 | 0.00459008584226769 | 0.996469305747651 | 0.999885157113822 | -5.25731933706875 |
| **Adgrl2** | 0.0308114850000045 | 15.22963939 | 0.00456873115853628 | 0.996485731640024 | 0.999885157113822 | -5.25731941940037 |

| | | | | | | |
|---|---|---|---|---|---|---|
| **Kcnc2** | 0.000625454999997999 | 27.8826947975 | 0.00357601140366298 | 0.997249327500712 | 0.999885157113822 | -5.25732282300046 |
| **Dync1li2** | 0.020915914999998 | 15.16984749 | 0.00311333779257422 | 0.997605215211324 | 0.999885157113822 | -5.25732412584785 |
| **Smap1** | -0.000424855000005664 | 27.2785133575 | -0.0029223094205729 | 0.997752154068434 | 0.999885157113822 | -5.25732461119988 |
| **Atp12a** | -0.000511244999991334 | 26.09161117 | -0.00275468915626984 | 0.997881087507223 | 0.999885157113822 | -5.25732501177181 |
| **Ccdc9** | 0.000489984999987314 | 26.57272565 | 0.00242877129849587 | 0.998131783581923 | 0.999885157113822 | -5.25732572292516 |
| **Aldh1a7** | -0.0160151750000015 | 15.582811675 | -0.00231950525150078 | 0.998215831097344 | 0.999885157113822 | -5.25732594132718 |
| **Cbr3** | -0.0147701975000025 | 17.17329512375 | -0.00194058166469004 | 0.998507299552196 | 0.999885157113822 | -5.25732662085241 |
| **Itpka** | -0.0125031824999962 | 15.41054270375 | -0.00182723340066556 | 0.998594487216016 | 0.999885157113822 | -5.25732680063185 |
| **Farsa** | -0.000545105000004042 | 21.2021437275 | -0.00169555409711358 | 0.99869577517033 | 0.999885157113822 | -5.25732699590367 |
| **Mtch1** | 0.00556344250000151 | 10.20538539375 | 0.000711011359557414 | 0.999453087877941 | 0.999885157113822 | -5.25732799330048 |
| **Fau** | -0.000165405000004171 | 25.00161566 | -0.000688891818485069 | 0.99947010229375 | 0.999885157113822 | -5.25732800633545 |
| **Wars1** | 0.00352715500000045 | 9.9481697075 | 0.000462371118580808 | 0.999644342695309 | 0.999885157113822 | -5.25732811611419 |
| **Stub1** | 0.00143468499999777 | 9.6381734825 | 0.000194128435353536 | 0.999850675797792 | 0.999885157113822 | -5.2573281902449 |
| **Arhgap26** | 5.96374999943805E-05 | 23.89565884375 | 0.000149301114019006 | 0.999885157113822 | 0.999885157113822 | -5.25732819672552 |
